## Supplementary text, figures, and tables for "Performance of *qpAdm*-based screens for genetic admixture on admixture-graph-shaped histories and stepping-stone landscapes"

### Supplementary text 1

#### ***An overview of published rotating and model competition qpAdm protocols***

In this section we outline two published rotating *qpAdm* protocols (Narasimhan *et al.* 2019; Lazaridis *et al.* 2022) that are typical for this class of protocols (see further examples in Skoglund *et al.* 2017; Harney *et al.* 2019; Olalde *et al.* 2019; Carlhoff *et al.* 2021; Fernandes *et al.* 2021; Librado *et al.* 2021; Bergström *et al.* 2022; Oliveira *et al.* 2022; Taylor *et al.* 2023; Allentoft *et al.* 2024). Lazaridis *et al.* relied on the following set of 15 reference populations: 1) Mbuti (present-day Africans); 2) a Paleolithic group from the Caucasus (CHG, Caucasian hunter-gatherers); 3) East European Mesolithic (EHG, East European hunter-gatherers); 4) Ganj Dareh (a Neolithic group from Iran); 5) Natufians (an Epipaleolithic group from Israel); 6) a Pre-pottery Neolithic (PPN) group from the Levant; 7) Taforalt (an Epipaleolithic group from Morocco); 8) Neolithic Mesopotamia; 9) Afontova Gora 3 (an individual from Late Upper Paleolithic Siberia); 10) Mal'ta 1 (an individual from Late Upper Paleolithic Siberia); 11) a Mesolithic group from the Iron Gates region (Serbia); 12) Boncuklu (a Pre-pottery Neolithic group from Central Turkey); 13) Barcın (a Neolithic group from Western Turkey); 14) Pınarbaşı (an Epipaleolithic individual from Turkey); and 15) Mesolithic and Paleolithic individuals from Western Europe (WHG, West European hunter-gatherers).

This reference set was divided into all possible "right" and proxy source subsets, except for the African group (Mbuti) which stayed in the "right" set in all models. Three Chalcolithic groups from Iran and an Early Bronze Age group from Russia (Yamnaya) were considered as proxy sources only and not rotated to the "right" set, and various clusters of Chalcolithic and Bronze Age individuals (most of them dated between ca. 5000 and 1000 years BCE) from the Balkans, Anatolia, Levant, Caucasus, Mesopotamia, and Iran were target groups for the *qpAdm* analyses. Thus, this protocol can be classified as a distal rotating protocol (**Box 1**) since most (but not all) targets do not pre-date the proxy sources (**Fig. S1a**). For each target, progressively more complex admixture models were tested, including from one to five proxy sources, and in most cases only the simplest feasible models were interpreted. Model feasibility criteria were as follows: estimated admixture proportions between 0 and 1, and *p*-

value > 0.01. Among alternative models for the same target, those having a higher *p*-value were considered fitting the data better (Lazaridis *et al.* 2022).

As shown in **Fig. S1a**, in this analytical setup there is a large temporal overlap between "left" groups (targets and, on average, earlier proxy sources) and "right" groups. For instance, such a divergent and ancient group as the Mal'ta 1 individual from the vicinity of Lake Baikal (dated to ca. 24,000 years before present, yBP; Raghavan *et al.* 2014) appeared "on the left" in some *qpAdm* models. Thus, "left-to-right" gene flows (that may lead to erroneous conclusions from a *qpAdm* analysis, see **Fig. 1**) are expected to be common in the analytical setup used by Lazaridis *et al.*

Narasimhan *et al.* (2019) used both proximal and distal *qpAdm* protocols (**Box 1**). The distal rotating protocol relied on the following set of 16 reference populations: 1) Mota (a 4500-years-old individual from Ethiopia); 2) Ust'-Ishim (an Upper Paleolithic individual from West Siberia); 3) Tianyuan (an Upper Paleolithic individual from Northeast China); 4) Late Upper Paleolithic individuals from Siberia (Afontova Gora 3 and Mal'ta 1, collectively labelled "ANE" or "Ancient North Eurasians"); 5) a Late Upper Paleolithic individual from Italy (Villabruna); 6) Natufians (an Epipaleolithic group from Raqefet, Israel); 7) a Mesolithic individual from Iran (Belt Cave); 8) present-day Andamanese; 9) East European Mesolithic individuals (EEHG, East European hunter-gatherers); 10) West Siberian Mesolithic (WSHG, West Siberian hunter-gatherers); 11) a Pre-pottery Neolithic (PPN) group from the Levant; 12) a Mesolithic group from the Iron Gates region (WEHG, West European hunter-gatherers); 13) Anatolian Neolithic individuals; 14) Ganj Dareh (a Neolithic group from Iran); 15) an Early Neolithic group from the Baikal region (ESHG, East Siberian hunter-gatherers); and 16) present-day Han Chinese. This reference set was split into all possible "right" and proxy source subsets, except for the Upper Paleolithic individuals/groups (Ust'-Ishim, Tianyuan, ANE, Villabruna) who stayed in the "right" set in all models. Diverse groups from Iran, Pakistan, Central Asia, and the Russian steppe zone (dated from the Chalcolithic to the historical period) were used as targets for the *qpAdm* protocol. For each target, generally post-dating the proxy sources (**Fig. S1b**), progressively more complex admixture models were tested, from one- to five-way mixture models, and in most cases only the simplest feasible models were interpreted. Model feasibility criteria were as follows: estimated admixture proportions ± 2 standard errors are between 0 and 1, and *p*-value > 0.01 (Narasimhan *et al.* 2019).

In summary, this *qpAdm* protocol rotated a diverse set of groups between the "right" and "left" sets: from present-day to Mesolithic groups older than 10,000 years, and from Africans to South and East Asians (see date distributions in **Fig. S1b**). Another *qpAdm* protocol used by Narasimhan *et al.*, termed "proximal" protocol (**Box 1**), relied on a smaller fixed set of groups that were kept always "on the right": 1) Mota (a 4500-years-old individual from Ethiopia); 2) East European Mesolithic individuals (EEHG, East European hunter-gatherers); 3) West Siberian Mesolithic (WSHG, West Siberian hunter-gatherers); 4) a Pre-pottery Neolithic (PPN) group from the Levant; 5) a Mesolithic group from the Iron Gates region (WEHG, West European hunter-gatherers); 6) Anatolian Neolithic individuals; 7) Ganj Dareh (a Neolithic group from Iran); 8) an Early Neolithic group from the Baikal region (ESHG, East Siberian hunter-gatherers). Thirty-one diverse Neolithic, Chalcolithic and Bronze Age groups from Eurasia were originally used as proxy sources in one- to three-way models, but if several feasible models were found for the target, the proxy sources from those models were moved one by one from the "left" to the "right" sets (i.e., model competition was performed). The set of targets and the model feasibility criteria matched those for the "distal" protocol. At the model competition step, groups very close in space and time appeared on both sides of the "left" - "right" and proxy source - target divides (**Fig. S1c**), making “left-to-right” gene flows (**Fig. 1**) highly likely. Of 45 target groups, 23 groups were also used as rotated proxy sources. For this reason, we interpreted the Narasimhan *et al.* proximal model competition protocol as follows (**Box 1**): its first step is a non-rotating *qpAdm* protocol with temporal stratification of “right” and “left” sets, but with no (or very limited) temporal stratification of targets and proxy sources; and the second step is a model competition protocol with no (or very limited, see **Fig. S1c**) temporal stratification of targets and proxy sources. We note that any such interpretation is an approximation that captures most important features of a published protocol and omits some details.

Since the sets of groups that are split into the "left" and "right" subsets in the protocols summarized above are very diverse chronologically and genetically, and since there are substantial overlaps in dates between the "left" and "right" subsets (**Fig. S1**), we argue that this approach is essentially similar to taking an admixture graph connecting populations sampled at widely different times in history, with divergence dates ranging from the Paleolithic (up to ca. 87,000 yBP in our simulations) to the "present" in the context of the dataset, and *randomly* splitting this graph into "left" and "right" population sets.

#### Supplementary text 2

***Case studies illustrating false and true feasible qpAdm models***

*False positive models*

This section is based on high-quality and noisy data generated by simulation setups no. 1 and 2 (**Fig. S2**).

In **Fig. S6b**, the target group, *G*, was simulated as a two-way mixture, and an incorrect proximal model "*G = C + J*" emerged as fitting in 18 of 40 simulation/subsampling replicates. Here, *C* is a group genetically similar to *G* ($F_{ST} < 0.015$ in **Fig. S6b**) and is largely its descendant (87% of ancestry in *C* is derived from the *G/I* lineage, see also **Fig. S3c**) rather than a proxy source (only 1.7% of its ancestry is derived from one of the true ancestry sources for *G*). Despite the small $F_{ST}$, the cladality of groups *G* and *C* was rejected in 31 of 40 simulation/subsampling replicates due to simulated gene flows from outgroup (*L* and *M*) lineages into *C* (**Fig. S6b**; this situation is schematically depicted in **Fig. 1a**). The model "*G = C + J*" was rejected according to *p*-values only when a lot of data was available (1,000-Mbp-sized genomes and high-quality data, **Fig. S6b**). This rejection of the incorrect two-way model does not mean that the outcome of *qpAdm* analysis for the target is acceptable since even more complex three-way false models may end up being fitting, although we did not fit three-way models in this study. A correct model we picked as an example, "*G = A + J*", was universally rejected according to *p*-values (**Fig. S6b**) due to multiple violations of the topological assumptions: for instance, outgroups *C* and *I* are derived from the target lineage after the admixture event (**Fig. 1d**). However, removal of various violating outgroups did not make the model "*G = A + J*" and similar models "*G = L/M + J*" fitting. On all datasets the correct model "*G = A + J*" was not supported by PCA and *ADMIXTURE*, whereas the FP model "*G = C*

*+ J*" was supported by these methods in most cases (**Fig. S6b**). Moreover, no true two-way model for group *G* emerged as feasible across the 40 simulation/subsampling replicates we explored in this analysis: other FP models were "*G = B + C*" (also supported by PCA, see **Fig. S6b**) and "*G = I + X*". This case study is another illustration of incorrect inference of gene flow direction by rotating *qpAdm* and of the risk associated with application of rotating protocols in the situation when proxy sources post-date putative target groups (group *C* is much younger than *G* and is a descendant of a population closely related to it, see **Fig. S3c**). This case study also shows that certain combinations of simulated admixture graph topologies and population sampling patterns create very unfavorable conditions for rotating *qpAdm* protocols and for certain target groups lead to false findings exclusively; and these false findings are supported by other methods such as PCA and *ADMIXTURE*.

In **Fig. S6c**, the target group, *A*, was simulated as a two-way mixture, and we expect that proximal models "*A = B + C/K*" would be fitting. The topological assumptions are violated when testing these models with the rotating protocol (**Fig. S6c**): groups *C* and *K* are cladal, with one of them appearing in the "left" set and the other one in the "right" set (this situation is schematically illustrated in **Fig. 1e**); a gene flow from an outgroup branch, *D*, enters the *C/K* branch after it splits from one of the true ancestry sources (**Fig. 1a**). However, *p*-values of the models "*A = B + C/K*" were high, but they were universally considered unfeasible due to estimated admixture proportions being negative ("*A = B + K*" is shown as an example in **Fig. S6c**). Removal of the outgroups *C, D,* and *K* does not make these models fitting. In contrast, incorrect proximal models "*A = B + G/J*" often emerged as fitting. The proxy sources *G* and *J* are symmetrically related to both true ancestry sources in the target. In this case proxy sources *B* and *G/J* do not have equal standing: while group *B* is indeed related to one of the two real sources for group *A*, *G* and *J* are groups completely "external" with respect to the admixture history of *A*. The model "*A = B + G*" shown as an example in **Fig. S6c** was rejected in some replicates according to *p*-values, but only if a large amount of data was available (1000-Mbp-sized genomes, high-quality data). Overall, it was fitting in 18 of 40 simulation/subsampling replicates. Both the false ("*A = B + G*") and true positive ("*A = B + K*") models discussed as examples here were universally supported by three-dimensional PCA and unsupervised *ADMIXTURE* (at some *K* values) on all datasets analyzed (**Fig. S6c**). Of 72 feasible two-way models found for group *A* as a target across the 40 simulation/subsampling replicates, all were classified as FP, and all followed the pattern "*A = B + X*", with *G*, *J*, *D*, *E*, and *I* being the most common second sources. Another example of an inappropriate proxy source symmetrically related to both true ancestry sources for a target is shown in **Fig. S6d**: that is distal model "*E = K + L*" which was usually rejected by the rotating protocol if enough data is available (feasible in 6 of 40 simulation/subsampling replicates) but was feasible in the case of the non-rotating protocol (in 34 of 40 simulation/subsampling replicates). This model was also supported by PCA on all four datasets used for this analysis.

In **Fig. S6e**, the target group, *A*, was simulated as a two-way mixture, and both correct and incorrect two-way models were fitting approximately equally often for this target: across the 40 simulation/subsampling replicates, 38 true positive (TP) models and 47 FP models were found. We selected two models as examples: both correct ("*A = B + F*") and incorrect ("*A = F + M*") proximal models fitted approximately equally often (in 28 and 23 of 40 replicates, respectively), suggesting that violations of the topological assumptions of *qpAdm* play no role in the emergence of this FP model. FDR for target *A* varied between 56% and 61% in the case of low-quality data and/or 300-Mbp-sized genomes; in contrast, for 1,000-Mbp-sized genomes and high-quality data FDR dropped to 14%. Failure to reject the model "*A = F + M*", where both proxy sources represent only one true ancestry source and are symmetrically related to the other, may be attributed to low $F_{ST}$ between all the populations involved in both models (**Fig. S6e**) and to the lack of data: increasing the simulated genome size to 1,000 Mbp and using high-quality data resulted in rejection (according to *p*-values) of the correct model "*A = B + F*" in 4 of 10 replicates, and in rejection of the incorrect model "*A = F + M*" in 9 of 10 replicates (**Fig. S6e**). Both models shown here as examples had universal support by PCA and support by unsupervised *ADMIXTURE* (at some *K* values; **Fig. S6e**). In this case study, the two-way model "*A = F + M*" would lead to misleading historical interpretations: although the target is indeed a two-way mixture, one of the true sources is completely missing in the model.

In **Fig. S6d**, the target group, *E*, was simulated as a two-way mixture, but no appropriate source proxy was sampled for one of the true ancestry sources: in groups *H* and *A*, 30% and 6% of their ancestry, respectively, is derived from that true source. Thus, multiple gene flows from the "right" set enter the *A* lineage after its split from the true ancestry source, and the same is true for the *H* lineage, and these are violations of the topological assumptions of *qpAdm* (**Fig. 1a**). The other true source for group *E* has a closely related proxy sampled (group *L*, see **Fig. S3g**), and $F_{ST}$ between groups *E* and *L* is low, < 0.015 (**Fig. S6d**). Despite this low $F_{ST}$, the (inaccurate) one-way model "*E = L*" was rejected by both the rotating and non-rotating *qpAdm* protocols in 39 of 40 simulation/subsampling replicates. The two-way model "*E = H + L*" was rejected according to *p*-values in 37 of 40 replicates; in contrast, the proximal model "*E = A + L*" was rejected according to *p*-values only in the case of 1,000-Mbp-sized genomes and high-quality data (**Fig. S6d**). A similar proximal model, "*E = A + M*", was usually rejected by the rotating protocol (feasible in 6 of 40 replicates; group *L* is closer to the respective real source than group M), but both models "*E = A + L/M*" were feasible if the non-rotating protocol was used (in 36 or 38 of 40 replicates). The ancestry proportion contributed by the proxy sources *L* or *M* was consistently overestimated by all the models and protocols we investigated (**Fig. S6d**); this appears to be related to the lack of appropriate proxies for the other true ancestry source. Accepting models like "*E = A + L/M*" at face value may lead to erroneous historical interpretations as the "present-day" group *A* is genetically far from the real source, having diverged from it 390 generations ago and having received 94% of its ancestry from other sources (**Fig. S3g**). But all thresholds for classifying models of this type into false and true are arbitrary. We chose 40% as a threshold percentage of proxy source's ancestry derived from the corresponding true source. All 212 two-way feasible models for target *E* (across 40 replicates and both rotating and non-rotating protocols) were classified as false. Since no true model was feasible for target *E*, rejection of an incorrect two-way model given ample data is not an acceptable result, and even more complex false models (which we did not test) may emerge as feasible. Models "*E = A + L/M*" were supported by PCA on all four datasets we explored and by unsupervised *ADMIXTURE* on two of these datasets (**Fig. S6d**). This case study illustrates again that applying *qpAdm* protocols to poorly sampled regions of the world (where no close proxies are available for some key ancestry sources) and without temporal stratification of "right" and "left" sets may be risky.

In **Fig. S6f**, the target group, *H*, was simulated as a two-way mixture, with one source being the deepest lineage in the graph, and the other source close to a sampled group, *M*. This situation is similar to the basal Eurasian hypothesis (Lazaridis *et al.* 2016). A group of FP models was found for this target. One way-models "*H = M*" were almost always rejected (in

39 of 40 simulation/subsampling replicates), despite low $F_{ST}$ for this population pair (**Fig. S6f**), and two-way distal models "*H = F/I/L + M*" were often fitting, especially the model "*H = F + M*" which was fitting in 35 of 40 simulation/subsampling replicates including those with the highest amount of data. Group *F* is the second-deepest lineage in this simulated graph after the true ancestry source for *H*, but it is distinct from the latter. The distal model "*H = F + M*" was universally supported by three-dimensional PCA and negative $f_3$-statistics $f_3$(target; proxy source$_1$, proxy source$_2$) (**Fig. S6f, Suppl. text 2**), as well as strongly supported by the unsupervised *ADMIXTURE* analysis (**Fig. S6f**) in the case of simulation setup no. 3 (at 7 of 8 *K* values tested). Given the simulated history and population sampling pattern, a researcher relying on the standard archaeogenetic toolkit would conclude that group *H*, sampled 44 generations ago, is derived from a lineage closely related to *M*, sampled 74 generation ago, and an extinct lineage related to a much more ancient group *F* sampled 604 generation ago. While the conclusion about deeply-divergent ancestry in *H* is correct, and the proportion of that ancestry estimated on low- or high-quality data is precise (**Fig. S6f**), its true source is a separate unsampled lineage not cladal with *F*. Interpretation of such a *qpAdm* model in terms of population history would be misleading if the true divergence point of the deep ancestry is not revealed by an admixture graph analysis similar to that supporting the basal Eurasian hypothesis (Lazaridis *et al.* 2016). Importantly, all 113 two-way feasible models for target *H* (across 40 replicates and both rotating and non-rotating protocols) were classified as false.

Another situation where deeply divergent ancestry in a target is misrepresented by an inappropriate proxy source is model "*F = A + E*" in **Fig. S6g**. In contrast to the situation in **Fig. S6f**, the deeply divergent ancestry in *F* is not unique to this group, but also contributed to several other sampled groups. The distal model "*F = A + E*" was feasible in 24 of 40 simulation/subsampling replicates (for both rotating and non-rotating *qpAdm* protocols). It was weakly rejected when a lot of data was available (**Fig. S6g**), and on the lowest-quality data we tested (300-Mbp-sized genomes, noisy data) it was often rejected (e.g., in 5 of 10 replicates analyzed with the rotating protocol) due to negative admixture proportion estimates and/or high *p*-values for one-way models. In contrast to most models discussed above, this model was not supported by PCA and unsupervised *ADMIXTURE* (**Fig. S6g**). Of 70 two-way feasible models for target *F* (across 40 replicates and both rotating and non-rotating protocols), 69 were classified as false. Thus, this situation would likely be ambiguous for a researcher encountering it in practice.

*True positive models*

Above we have discussed several examples of FP models that in practice would result in misleading conclusions about population history, and now we turn to feasible models that were interpreted as true in our analysis. We rely on a relaxed definition of true models: two-way admixture models for targets with population history best approximated with three-way and more complex models were considered as true positives if they included at least two source proxies *not* satisfying the topological criteria of false models listed above. We also note that TP models were not restricted to those including most optimal source proxies, if the models do *not* satisfy the false positivity criteria. Stricter definitions of TP models are possible (such as those in the companion paper; Williams *et al.* 2024), but we applied this lenient approach since in the archaeogenetic literature sampling of ancient individuals is often poor, and thus the task is not to find all groups which participated in the history of a given target group, but to identify at least predominant ancestry sources with some degree of precision.

On a random sample of 400 two-way admixture models from our 40 simulated histories, the fraction of models that were classified as appropriate (true) according to the rules described above was 17.7%. Since groups that are truly admixed are common in our simulations, we do not expect to encounter a "needle in a haystack" situation where finding true admixture models is exceedingly hard. Examples of simulations yielding TP *qpAdm* models are presented below and are based on high-quality and noisy data generated by simulation setups no. 1 and 2 (**Fig. S2**).

In **Figs. S3a** and **S6h**, a set of ideal distal TP models is presented. The target, group *K*, was simulated as a two way-mixture, and close (*H, I*) and more distant proxies (*G, M*) for both true sources were sampled. All four alternative proxy sources display no violations of the topological assumptions of *qpAdm* (**Fig. 1**), and outgroups such as *L* and *C* related differentially to the true and proxy sources are available (**Fig. S6h**). An ideal model including the closest proxy sources sampled, "*K = H + I*", and a nearly ideal model (with one proxy being more distant from the true source), "$K = G + I$", were nearly universally fitting when the rotating protocol was applied (in 38 and 35 of 40 simulation/subsampling iterations, respectively) and yielded precise estimates of admixture proportions (**Fig. S6h**). However, the non-rotating protocol demonstrated low power for this target on noisy pseudo-haploid data, resulting in models being unfeasible due to high *p*-values for one-way models and/or negative admixture proportion estimates: the "$K = H + I$" and "$K = G + I$" models were feasible in just 2 or 4 of 20 subsampling iterations, respectively. The two-way models were not rejected according to *p*-values in this case, and had usually non-negative, but imprecise, admixture proportion estimates (**Fig. S6h**). This observation illustrates the power of rotating protocols (Harney *et al.* 2021) and the motivation behind their adoption. In the case of high-quality data though, both models were unfeasible in just 1 of 20 simulation iterations analyzed by the non-rotating protocol. Non-optimal models "$K = H + M$" and "$K = G + M$" were rejected according to *p*-values by the rotating protocol (but not the non-rotating one) if enough data was available (**Fig. S6h**). All 112 two-way feasible models for target *K* (across 40 replicates analyzed by the rotating protocol) were classified as true, but of 142 feasible models that were outcomes of the non-rotating protocol, just 86 were classified as true. These observations demonstrate again the advantage of the rotating protocol (in rejecting models with non-optimal or inappropriate proxy sources) that exist when there are no violations of the topological assumptions of *qpAdm*. All four models for *K* had universal support by three-dimensional PCA and unsupervised *ADMIXTURE* analyses (at least at some *K* values, see **Fig. S6h**).

Another series of examples (**Figs. S3c,e, S6i-j**) illustrates models that very often emerge as feasible despite imperfect cladality of proxy and true ancestry sources (see schematic representations in **Fig. 1a,b**). The targets in these cases were simulated as two-way mixtures. Despite non-cladality of one proxy source with the corresponding true source, distal model "$C = I + K$" (**Fig. S6i**) was usually feasible when both the rotating and non-rotating protocols were applied (in 37 and 30 of 40 simulation/subsampling replicates, respectively) and yielded precise estimates of admixture proportions (**Fig. S6i**). The proximal models illustrated in **Fig. S6j**, "$K =C + D$" and "$K = D + L$", demonstrated violations of the topological assumptions at one or two proxy sources, were almost universally rejected by the rotating (but not the non-rotating) protocol, and produced estimates of admixture proportions that were always biased

(**Fig. S6j**). Given that the true model in this case is two-way, rejection of a model due to non-cladality of true and proxy sources may lead to erroneous interpretations if more complex (and false) three-way models emerge as feasible. However, as stated above, three-way models were not explored in this study. All true (or false) two-way models for target *K* were rejected by the rotating protocol on high-quality data (300- or 1000-Mbp-sized genomes). Thus, this case study (**Fig. S6j**) demonstrates a disadvantage of the rotating protocol when all proxies sampled for a certain true ancestry source demonstrate violations of the topological assumptions of *qpAdm*.

Finally, we consider examples of targets that were simulated as three-way mixtures, but for whom two-way *qpAdm* models were often feasible (**Fig. S6i,k-m**). In this case rejection of such a two-way model is considered an ideal outcome and was indeed observed very often when the rotating protocol was applied to high-quality data. In **Fig. S6i** (target *E*), all three true sources were represented by sampled proxies, however two of them demonstrated non-cladality with the true sources. A two-way model, "*E = C +K*", was rejected by the rotating (but not the non-rotating) protocol only when a large amount of data was available and, surprisingly, yielded precise estimates of admixture proportions (**Fig. S6i**). This incomplete (two-way) model for target *E* was also not supported by PCA and *ADMIXTURE* (**Fig. S6i**). In **Fig. S6k**, two-way models for a three-way admixed target were almost always rejected by the rotating protocol (but not the non-rotating protocol) according to *p*-values. In one case ("*I = D + J*") rejection was driven also by negative estimates of admixture proportions (**Fig. S6k**). The two-way models for target *I* were overwhelmingly supported by PCA and *ADMIXTURE* (**Fig. S6k**). That is, the third ancestry component in group *I* is largely invisible to these methods. A similar situation is illustrated in **Fig. S6l**, where both the rotating and non-rotating protocols tend to reject all three two-way models shown (according to *p*-values), but only if a lot of data is available (1000-Mbp-sized genomes and high-quality data). The two-way models for target *H* were also overwhelmingly supported by PCA and *ADMIXTURE* (**Fig. S6l**). In **Fig. S6m**, an opposite situation is shown. The model "*D = F + I*" was feasible in nearly all cases (35 of 40 replicates when the rotating protocol was applied and 38 of 40 replicates when the non-rotating protocol was applied) since proxy source *I* represents two true sources at the same time (**Fig. S6m**), however it lacked support by PCA and *ADMIXTURE* (**Fig. S6m**). Ancestry proportions were estimated with a small bias since proxy source *I* does not have the correct ratio of ancestries. We stress that whenever two-way models were feasible for targets modelled as more complex mixtures, they were nevertheless considered as TP in our subsequent analyses.

#### Supplementary text 3

***There is no practical way of reducing FDR of the proximal rotating qpAdm protocol***

This section is based on high-quality data generated by simulation setups no. 1, 2, 4, and 5 (**Fig. S2**).

We observe that among two-way admixture models supported by both proximal rotating *qpAdm* and PCA, FDR remains very high: from 60% to 77%, and this comes at a cost of rejecting 41% or 61% of TP *qpAdm* models (false omission rate, FOR) as those were not supported by PCA (**Table S2**). Adding an unsupervised *ADMIXTURE* analysis applied to all 13 simulated groups as another level of control helped to drive FDR below 50% (to 42%) only in the case of the fourth simulation setup where all groups were sampled at “present” (**Table S2**). Considering only admixture models supported by all three methods increases FOR to 69 – 76% (**Table S2**).

There exists a test for admixture that helped to reduce the FDR of the proximal rotating *qpAdm* protocol more substantially than PCA or *ADMIXTURE*, to 14 – 33% on high-quality data (**Table S2**), and these are "admixture" $f_3$-statistics of the form $f_3$(target; proxy source$_1$, proxy source$_2$) (Patterson *et al.* 2012). FDR of models supported by $f_3$-statistics not combined with *qpAdm* was even lower, 6% (7 of 115 statistics with Z-score < -2 in the case of at least one of four sets of simulated high-quality data; simulation replicate no. 1 was considered for each simulation setup). However, "admixture" $f_3$-statistics tend to become positive when genetic drift on the target lineage moves allele frequencies away from their original values intermediate between those in the real ancestry sources (Patterson *et al.* 2012). Indeed, FOR for models supported by both *qpAdm* and $f_3$-statistics varied from 93% to 100% (**Table S2**), and just 115 of 34,320 population triplets tested with $f_3$-statistics demonstrated Z-scores < -

2 under at least one simulation setup (one replicate per setup considered). Thus, using $f_3$-statistics is impractical in large-scale screens for admixture.

Moreover, in the case of one simulated history, we found a class of false two-way admixture models consistently supported by negative $f_3$-statistic Z-scores down to -18.4 (on 1000-Mbp-sized genomes and high-quality data), and having strong support by *qpAdm*, PCA, and *ADMIXTURE* (**Fig. S6f**). In this case the target receives a gene flow from the most divergent lineage that remains unsampled, and other deep-branching lineages unrelated to the true source emerge as proxy sources in fitting *qpAdm* models (**Fig. S6f**). $f_4$-statistics are defined on unrooted trees, and $f_3$-statistics, being just a special case of them (Patterson *et al.* 2012), have the same property, which explains our observation.

Another way of reducing FDR of the proximal rotating *qpAdm* protocol is adjustment of *p*-value thresholds. All published rotating *qpAdm* protocols, to our knowledge, rely on one *p*-value threshold (usually 0.01 or 0.05), both for rejection of simple models and for acceptance of more complex models with an additional ancestry source. Here we compare distributions of *p*-values for FP and TP two-way models and the corresponding one-way models (we note that the distributions are conditional on the admixture graphs simulated). The means of the distributions of $log_{10}$(*p*-values) for one-way models were significantly different for TP and FP two-way models, although the distributions largely overlap (**Fig. S13b**; a one-way model with the highest *p*-value was considered for each population triplet). On high-quality diploid data, average $log_{10}$(p-values) for two-way models were significantly different for true and false *qpAdm* models only in the case of the fourth simulation setup (with groups sampled at "present") (**Fig. S13b**).

These *p*-value distributions (**Fig. S13**) suggest that introducing different thresholds for rejecting one-way *qpAdm* models and for accepting two-way models may reduce FDR. Indeed, setting the former threshold to $10^{-50}$ and the latter to 0.1 reduces FDR from 55% – 66% to 33% – 59% on high-quality data, with FOR from 63% to 71% (**Table S3**). FDR can be reduced even further, down to 27%, if support by proximal rotating *qpAdm*, PCA and *ADMIXTURE* is required to declare a positive result. This approximately two-fold reduction in FDR is associated with a very high FOR of up to 89% (**Table S3**).

In summary, we conclude that without the introduction of temporal stratification, on our simulated data it was impossible to reduce FDR of the proximal rotating *qpAdm* protocol even below 20%.

#### Supplementary text 4

***AGS histories: model competition qpAdm protocols and admixture inference pipelines***

This section is based on high-quality data generated by simulation setups no. 1, 2, 4, and 5 (**Fig. S2**).

An implicit assumption of many archaeogenetic studies relying on *qpAdm* protocols is that admixture models supported by clines observed in (usually two-dimensional) spaces of principal components, and/or by an *ADMIXTURE* analysis, and/or by individual *D*-, $f_4$- or $f_3$-statistics are especially robust. And, *vice versa*, *qpAdm* results are often interpreted as a formal test of hypotheses about admixture formulated based on PCA and/or *ADMIXTURE* results. We constructed "admixture inference pipelines" composed of a *qpAdm* protocol and one or two further methods to test these assumptions on AGS simulated data. We note that all patterns in our PCA or *ADMIXTURE* analyses consistent with admixture were not explored with *qpAdm*. *Vice versa*, all feasible *qpAdm* models were checked by the PCA and/or *ADMIXTURE* methods.

We considered a two-way admixture model to be supported by PCA if the target group was located on a straight line between the two proxy source groups in the space of first three PCs when all 13 simulated groups were co-analyzed on high-quality data. Deviation from the straight line due to post-admixture genetic drift was acceptable to a certain extent (see Methods) considering theoretical results by Peter (2022). Non-linear PCA clines are often observed on real data (de Barros Damgaard *et al.* 2018; Jeong *et al.* 2019), and deviations from straight lines were also common among TP two-way *qpAdm* models in this study (see Methods for details and **Fig. 2** and **Fig. S6** for examples). This situation is expected since many target groups in our simulations represent three-way and more complex mixtures (see examples in **Fig. S6k-m**), and since arrangement of populations in PC spaces is influenced not only by admixture of previously isolated groups or continuous gene flows decaying with distance (Duforet-Frebourg and Slatkin 2016), but also by genetic drift (McVean 2009; Peter 2022). Our requirements for a model to be declared supported by PCA were more stringent than those usually applied in the literature since we considered three-dimensional PC spaces instead of two-dimensional ones (however, 2D PCAs are shown in **Fig. S6** for clarity). Also see Methods for the rules we used to judge if an admixture model is supported by an unsupervised *ADMIXTURE* analysis and see **Fig. 2** and **Fig. S6** for examples.

We explored FDR of model competition protocols as well. A typical protocol of this type (Narasimhan *et al.* 2019; Maróti *et al.* 2022; Brielle *et al.* 2023) consists of two stages. First, the oldest, e.g., Paleolithic, populations (and/or those most different genetically from the target group) are used as a fixed "right" set, and populations sampled at later dates are used as proxy sources and targets. As usual, progressively more complex models are tested for targets of interest, and a composite feasibility criterion is applied.

In many publications (e.g., Haak *et al.* 2015; Mathieson *et al.* 2015, 2018; Antonio *et al.* 2019; Prendergast *et al.* 2019; Marcus *et al.* 2020; Papac *et al.* 2021; Wang *et al.* 2021; Yaka *et al.* 2021; Changmai *et al.* 2022a, 2022b; Patterson *et al.* 2022) this first non-rotating step remains the only *qpAdm* protocol used (in its distal or proximal forms). In a model competition protocol, subsequent analysis is focused on targets for whom two or more alternative *qpAdm* models emerge as feasible at the first step. For each target, alternative proxy sources are pooled and rotated between the "left" and "right" sets, testing only the models that emerged as feasible at the first step and applying a composite feasibility criterion. This rotation can be performed in various ways: "whatever is not on the left is on the right" (Brielle *et al.* 2023) or placing alternative sources in the "right" set one by one (Carlhoff *et al.* 2021; Maróti *et al.* 2022; Brielle *et al.* 2023).

In the latter case several "right" sets are tested for each model, and the model is considered supported by the model competition protocol only if it is not rejected under any of these "right" sets (Maróti *et al.* 2022). The reasoning behind this protocol is as follows: model rejection due to violations of the topological assumptions of *qpAdm* is not expected for a model composed of sources very close to the true ones since in this case branches private to the proxy sources are short, and it is unlikely that gene flows to or from the "right" population set happened on these short branches. Models composed of sources closely related to the true ones are also not expected to be rejected when more distant proxy sources are placed in the "right" set (Harney *et al.* 2021; see also a case study in **Fig. S6h**).

For reasons detailed in the Discussion section, we explored the *qpAdm* model competition protocol and multi-method admixture inference pipelines on one replicate per simulation setup, and three (**Table S5**) or five (**Table S4**) simulation setups were involved in this analysis. The two alternative model competition protocols described above were applied to targets for whom more than one model was feasible given a fixed "right" set composed of six groups. If only one model was feasible for a target, such a model was evaluated as passing model competition (such models accounted only for 7%–11% of models feasible at the first step). The model competition protocols failed to improve *qpAdm* performance: FDR ranged from 29% to 46% (as compared to 36%–46% prior to the model competition step), and both model competition protocol versions demonstrated very similar results (**Table S5**). FOR of the model competition protocols varied from 59% to 72%. FDR also remained high for models supported by proximal non-rotating *qpAdm* & PCA or by proximal non-rotating *qpAdm* & model competition & PCA (**Table S5**). Notably, FP models supported by the proximal non-rotating *qpAdm* protocol largely lacked support by an unsupervised *ADMIXTURE* analysis (**Table S5**), in contrast to outcomes of the proximal rotating protocol (**Table S2**). FDR of a pipeline composed of these two methods ranged from 5% to 21% across three simulation setups tested (**Table S5**). Adding a model competition step to this pipeline increased both FDR and FOR in 4 of 6 cases; and, in general, the proximal non-rotating *qpAdm* protocol combined with *ADMIXTURE* is the best-performing protocol in this analysis (**Table S5**) according to FDR and FOR.

The fact that our *ADMIXTURE* analysis supports a large fraction of FP two-way mixture models emerging as outcomes of the proximal rotating *qpAdm* protocol reflects known problems in modelling with *ADMIXTURE* very ancient individuals in the context of modern populations. These individuals are often modelled (Raghavan *et al.* 2014; Haak *et al.* 2015; Moreno-Mayar *et al.* 2018) as complex mixtures of ancestry components typical for modern populations, which is obviously an artefact. This artifact comes from the following facts: 1) *ADMIXTURE* uses an internal topology where each "component" is modeled as a population which descends from a root in a multifurcating tree such that all populations/components diverge simultaneously; 2) ancient populations have less drift, and thus are closer to the root; 3) due to the martingale/mean-zero nature of genetic drift, allele frequencies in the root population can be well approximated as a mixture of allele frequencies in all of its descendants (who each can be assumed to have a mean allele frequency of the root with some variance).

Above we have discussed multi-method pipelines based on the proximal non-rotating *qpAdm* protocol. Combining distal *qpAdm* protocols with PCA allows to reduce FDR of both rotating and non-rotating protocols further, to 10%–24%, and distal *qpAdm* protocols combined with an unsupervised *ADMIXTURE* analysis demonstrated even better FDR values ca. 0%–8% (**Table S4**).

A limitation of our study is that we had to use idealized versions of *qpAdm*, PCA, and *ADMIXTURE* protocols, while in the archaeogenetic literature manual adjustment of analytical protocols is common: protocols often vary from one target group to another (see, e.g., Lazaridis *et al.* 2016; Zhang *et al.* 2021; Brielle *et al.* 2023; Lee *et al.* 2023) and from study to study. These extensive details are very hard to formalize and reproduce. In the case of *qpAdm* protocols, certain groups of populations may be placed exclusively in “right" or in “left" sets, with the rest rotated between these sets, and relative sizes and compositions of these three groups vary from study to study: in the case of model competition protocols, this rotated subset is small, and rotation may be restricted to a particular model complexity class, but in other cases it may encompass all or nearly all populations analyzed (see, e.g., Narasimhan *et al.* 2019; Librado *et al.* 2021; Bergström *et al.* 2022; Lazaridis *et al.* 2022; Oliveira *et al.* 2022; Taylor *et al.* 2023). Reproducing all aspects of PCA and *ADMIXTURE* protocols used in the literature is also hardly possible on simulated data. For instance, PCs in archaeogenetic studies are usually calculated based on present-day populations, and ancient individuals are projected on the resulting PCs (e.g., Haak *et al.* 2015; Mathieson *et al.* 2018; Narasimhan *et al.* 2019; Furtwängler *et al.* 2020; Marcus *et al.* 2020; Lazaridis *et al.* 2022). In contrast, in our study all simulated individuals were co-analyzed for calculating PCs since this analysis was based on high-quality data only. Unsupervised *ADMIXTURE* analyses in the literature are usually performed on worldwide or continent-wide panels of populations that often overlap just partially with population sets used for *qpAdm* analyses (see, for instance, Rasmussen *et* *al.* 2010; Haak *et al.* 2015; Harney *et al.* 2018; Moreno-Mayar *et al.* 2018; Zhang *et al.* 2021; Changmai *et al.* 2022a; Brielle *et al.* 2023), while in our study identical population sets were used for *qpAdm*, PCA, and *ADMIXTURE* analyses.

Another limitation is that some comparisons of method performance in this study, such as *qpAdm* vs. "*qpAdm* combined with *ADMIXTURE*", are qualitative rather than quantitative: we applied the PCA and *ADMIXTURE* methods to one simulation replicate only per simulation setup since automated classifiers of admixture models into positive and negative ones based on PCA and *ADMIXTURE* results were not available.

#### Supplementary text 5

##### ***SSL histories: randomized sampling of landscapes***

*qpAdm performance at the level of models: model optimality metrics in spaces of admixture fractions SE and p-values*

Maximal standard errors of EAF (abbreviated as max. SE) correlate strongly with max|*EAF–EF*| (**Figs. S19, S20**): Spearman's $\rho$ range from 0.81 to 0.94 (when models are stratified by landscape type and model complexity) or from 0.76 to 0.95 (when models are stratified by landscape type, model complexity, *qpAdm* protocol, and sampling density; all the correlation coefficients are statistically significant). Therefore, if we consider max. SE instead of max|*EAF–EF*|, the same patterns are visible with respect to *p*-values and optimality metrics, but they are less clear-cut in the former case (**Figs. S19, S20, Tables S7, S8**). For instance, Spearman's $\rho$ for min. STS angle vs. max|*EAF–EF*| are in all cases higher, albeit slightly, than those vs. max. SE (for two- to four-way models on the "$10^{-4}$ to $10^{-3}$", "$10^{-5}$ to $10^{-2}$", and "$10^{-3}$ to $10^{-2}$" landscape types, and for *p*-values $\geq 10^{-55}$ or $\geq 10^{-5}$; **Table S7**). The same is true for correlations between *p*-values and optimality metrics in the most practically relevant parts of the two-dimensional spaces we inspect here (**Table S7**).

Inspecting the distributions of max. (or average) ST distances in the SE dimension, we note "striped" patterns, namely two or more separate SE intervals populated by sources relatively close to targets, probably arising due to the "quantized" nature of our landscape (**Figs. S19,**

**S20**). Interestingly, models having the lowest SE do not demonstrate the smallest median max. ST distances. In contrast, that part of the space is populated by the most symmetric models (**Figs. S19, S20**) suggesting that symmetric arrangement of sources is more important for achieving low SE than spatial proximity to the target.

*qpAdm performance at the level of models: one-way models*

High *p*-values of one-way *qpAdm* models (equivalent to "pairwise *qpWave* tests") are often used as evidence for "cladality" and "genetic continuity" in the archaeogenetic literature (see, e.g., Papac *et al.* 2021; Changmai *et al.* 2022a; Skourtanioti *et al.* 2023), and such *qpWave* tests have also become popular as a method for individual clustering that precedes other analyses (Fernandes *et al.* 2020, 2021; Lazaridis *et al.* 2022; Antonio *et al.* 2024; Speidel *et al.* 2025). We show in the context of our randomized *qpAdm* experiments that interpretation of such *qpWave* tests on SSL depends on properties of those landscapes (**Fig. S21**). For instance, of one-way models having *p*-value ≥0.05 on the sparsest landscapes ("$10^{-5}$ to $10^{-4}$"), just 8.7% include groups sampled from the same deme at different times (spatial distance = 0; **Table S10**), and there is a relatively flat distribution of models over spatial distances in this subset of models: 54.7% of "fitting" one-way models include demes at a distance of 3 or greater (**Fig. S21**, **Table S10**). Frequency of "fitting" one-way models in our equal-sized sets of random experiments is also highly variable across landscape types: e.g., it is ca. 10 times higher for the "$10^{-5}$ to $10^{-4}$" landscapes than for the "$10^{-4}$ to $10^{-3}$" landscapes (**Fig. S21**, **Table S10**). In the latter case ("$10^{-4}$ to $10^{-3}$"), most "fitting" one-way models (~85%) correspond to samples from the same deme (**Table S10**). With growing gene-flow intensity the rate of one-way model rejection grows, as expected, and the mode of this distribution moves from samples from the same deme to the nearest demes (**Table S10**).

The statements above describe various aspects of the shape of model distributions in the space formed by spatial distances between demes and *p*-values. There is a non-linear trend line that links *p*-value and spatial distance on all the landscapes, except for the sparsest landscape type where the shape of the distribution is strikingly different and more complex (**Fig. S21**). We also note that ideal one-way models (defined here as pairs of samples from the same deme) show very similar distributions over *p*-values for all the four sparsity levels, and with growing gene-flow intensity on the landscape these distributions become more different from the uniform one, as expected. Therefore, the differences in model distributions between the landscapes are mostly due to “non-ideal” models, which are much more common in our random model samples (**Fig. S21**).

To sum up, on the sparsest landscapes the high frequency of pairwise *qpWave* tests reporting high *p*-values reflects the initial multifurcation (**Fig. 4**) and a very low rate of subsequent gene flows, that is a situation that approaches all vs. all cladality (**Fig. S21**, **Table S10**). However, on landscapes of higher density the method works mainly as a test for genetic continuity since it identifies samples from the same deme, and, with increasing gene-flow intensity, clusters of nearby demes become progressively less structured, and the method identifies members of these clusters as “genetically continuous”. On the contrary, samples from the same deme at different time points become more dissimilar with increasing gene-flow intensities, and *qpWave* is sensitive to that (**Fig. S21**). Thus, interpretation of pairwise *qpWave* results in practice is confounded by these two very different scenarios producing equal outcomes, i.e., high *p*-values. In other words, interpretation of a high *p*-value encountered in practice could be hindered by a lack of knowledge about the sparsity of the underlying landscape.

#### Supplementary text 6

***Do estimated admixture fractions reflect gene flow intensities on SSL?***

Previously we did not assess the accuracy of admixture fraction estimates. On SSL with non-uniform gene flows (**Fig. S14b,c**) in all directions it is not trivial to derive the most accurate predictor of admixture fractions, and the concept of admixture fractions itself becomes questionable. It is more intuitive and useful to conceptualize ancestry on SSL in the form of clouds of ancestor density (Grundler *et al.* 2024). With these caveats in mind, we assessed correlation between simulated source-to-target gene-flow intensities and EAF. For simplicity, we considered the systematic sampling scenario and models with the nearest neighbors only as proxy sources; direct ancestors were not considered as sources (**Fig. S39**). More precisely, we used the following simple admixture fraction predictor:

predicted $AF_i = \frac{S_i \xrightarrow[gf]{} T}{\sum_{i=1}^{n} S_i \xrightarrow[gf]{} T}$ , where the numerator is a source-to-target gene-flow intensity in the last gene-flow epoch (lasting 153 generations, while proxy sources were sampled 100 generations in the past and targets at "present"; **Figs. 4, S16**), and the denominator is a sum of such intensities over all sources. Another analogous predictor can be constructed based on sums of source-to-target and target-to-source gene-flow intensities in the same epoch: $\frac{S_i \xrightarrow[gf]{} T + T \xrightarrow[gf]{} S_i}{\sum_{i=1}^{n} (S_i \xrightarrow[gf]{} T + T \xrightarrow[gf]{} S_i)}$.

The most notable pattern we observe when comparing EAF and AF calculated using the former simple predictor (**Fig. S39a**) comes from the linear algebra nature of *qpAdm* (Haak *et al.* 2015): symmetric and non-symmetric models produce EAF distributions of very different widths. In other words, only in the case of symmetric models nearly all EAF ∈ (0, 1). Thus, we conclude that EAF depend more strongly on STS angles than on gene-flow rates (**Fig. S39a**).

For highly non-symmetric two- or three-way models (on our SSL, all four-way models with nearest neighbors as sources are non-symmetric, as discussed in the main text), namely those with min. STS angles = 60°, Pearson's correlation coefficients for predicted AF and EAF were non-significantly different from 0 over the whole range of EAF, and non-significant or low (up to 0.27) over EAF in the range (0, 1) (**Fig. S39a**). The linear correlation becomes much stronger in the case of ideally symmetric two- and three-way models: Pearson's correlation coefficients vary from 0.5 to 0.58, and all are significantly different from 0 with Bonferroni correction (**Fig. S39a**).

However, even in the case of ideal symmetric models, the correlation is far from perfect (**Fig. S39a**). This may be a consequence of poor AF prediction since only direct source-to-target gene flows were considered. In other words, proxy sources in ideal symmetric models stand not only for themselves, but for equally sized sections of the landscape. Considering sums of reverse and forward ST gene-flow intensities (the second predictor above) allows linear correlation between EAF and predicted AF to be improved, but for two-way models only (**Fig. S39b**). Very similar results were obtained using both predictors on randomized *qpAdm* models (**Fig. S40a,b**).

**Figure S1.**

**a**, Lazaridis *et al.* 2022; distal rotating protocol

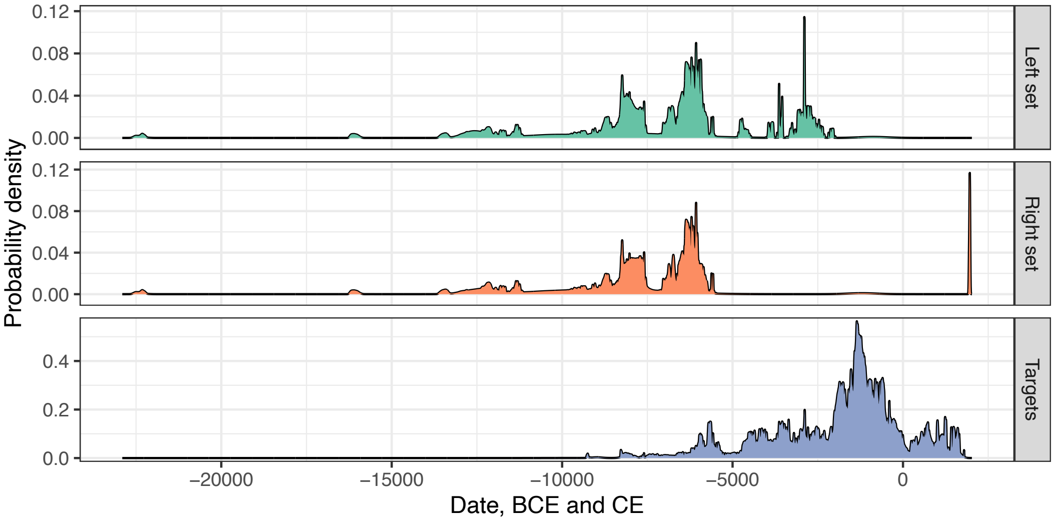


**b**, Narasimhan *et al.* 2019; distal rotating protocol

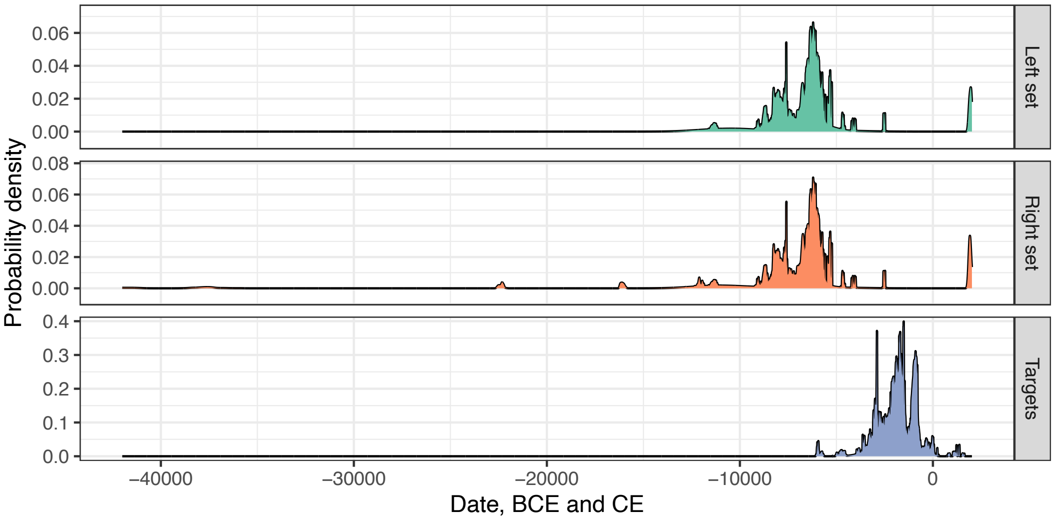


**c**, Narasimhan *et al.* 2019; proximal model competition protocol

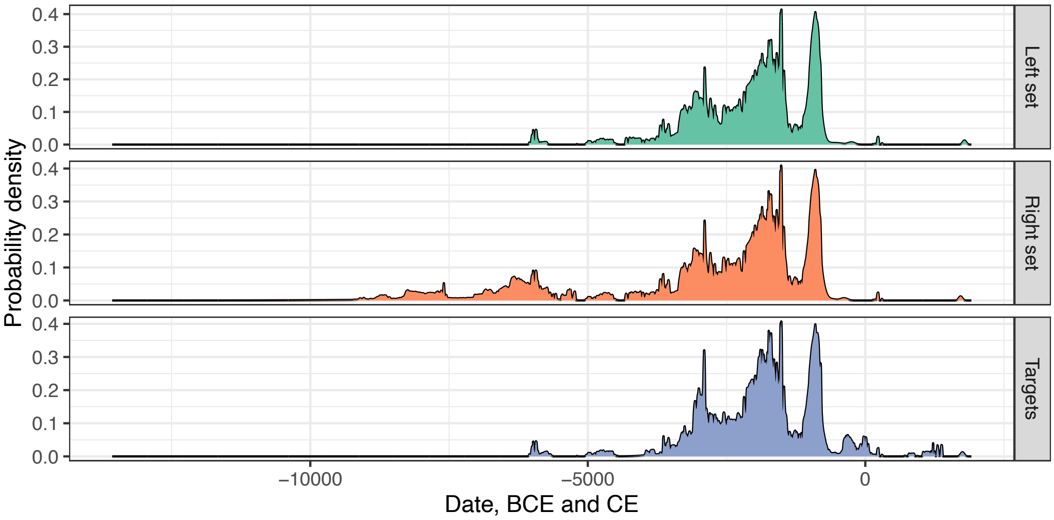


Distributions of radiocarbon and calendar dates for populations sets analyzed with the distal rotating *qpAdm* protocols by Lazaridis *et al.* 2022 (**a**) and Narasimhan *et al.* 2019 (**b**), and with

the proximal model competition protocol from Narasimhan *et al.* 2019 (**c**). Probability density curves are shown for three sets of groups: 1) those appearing in the "right" set in at least one *qpAdm* model; 2) those appearing in the "left" set in at least one *qpAdm* model; 3) target groups. Targets in the former study were composed of large clusters of West Eurasian individuals, some of them dating back to the Paleolithic (Lazaridis *et al.* 2022). For that reason, the date distribution for targets in panel **a** is very wide. See **Suppl. text 1** for a detailed description of the published protocols used for making this figure.

**Figure S2.**

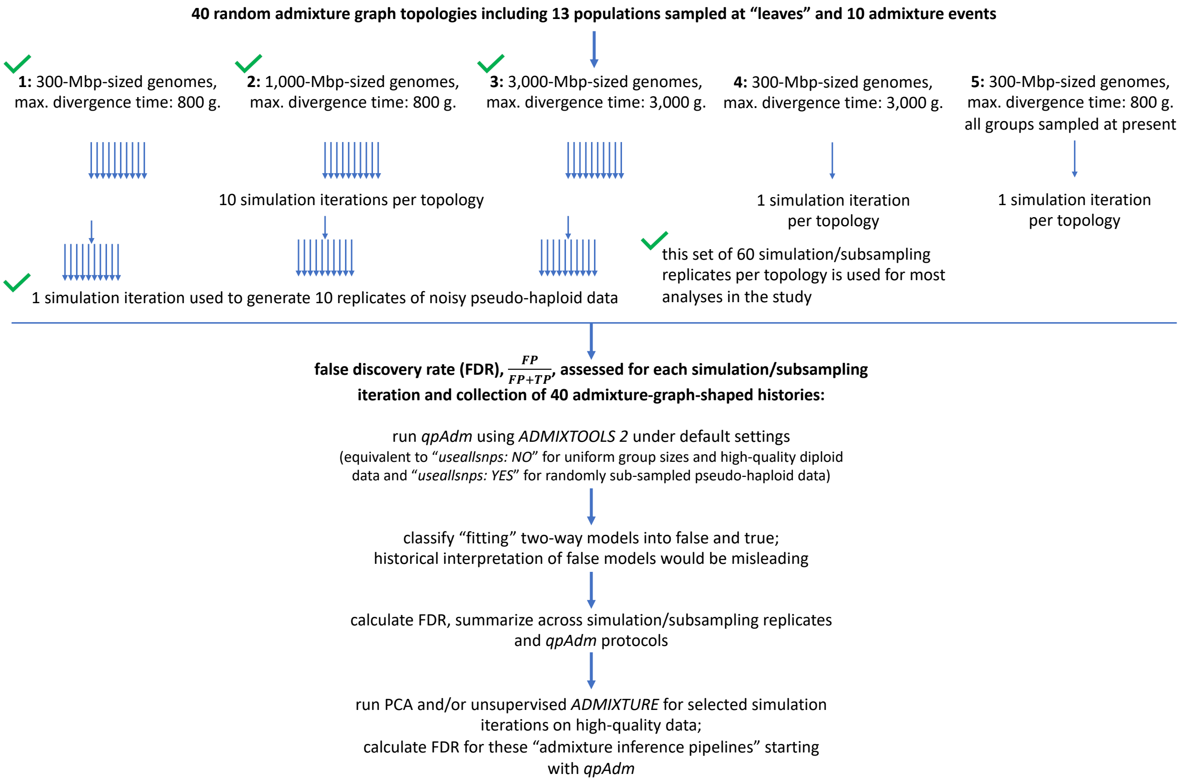


A diagram illustrating our analytical workflow in the case of AGS simulations. The simulation setups are numbered in bold, and these numbers are referred to in the text.

**Figure S3.**

**a**

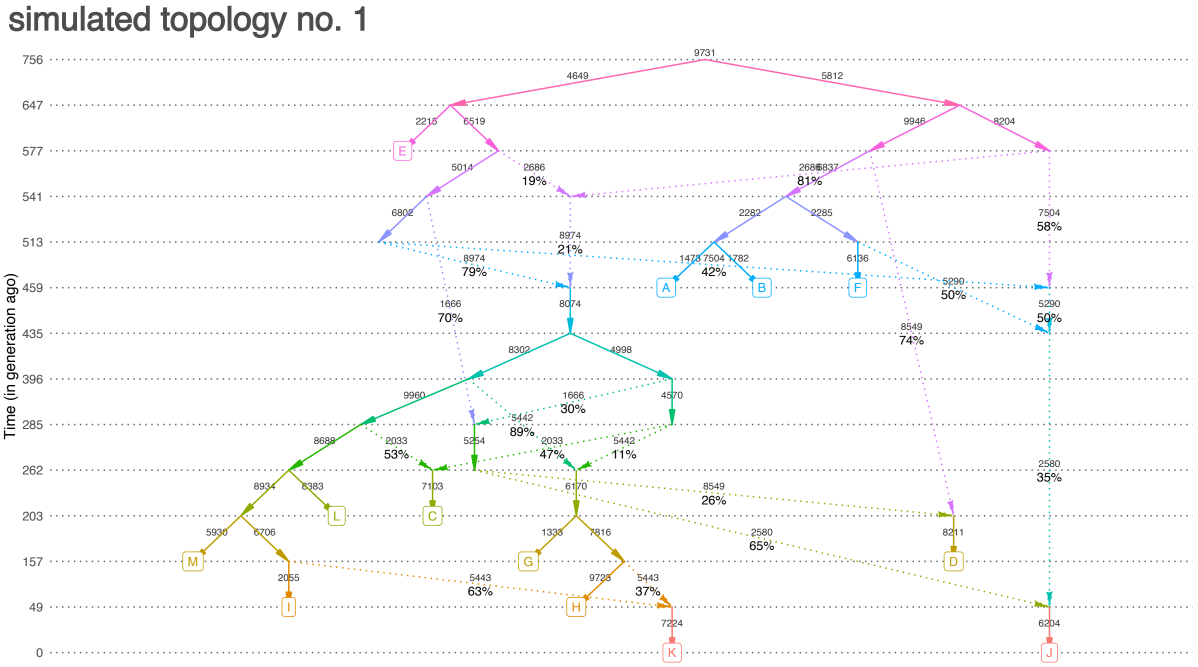

simulated topology no. 1
Time (in generation ago)
756
647
577
541
513
459
435
396
285
262
203
157
49
0
A
B
C
D
E
F
G
H
I
J
K
L
M


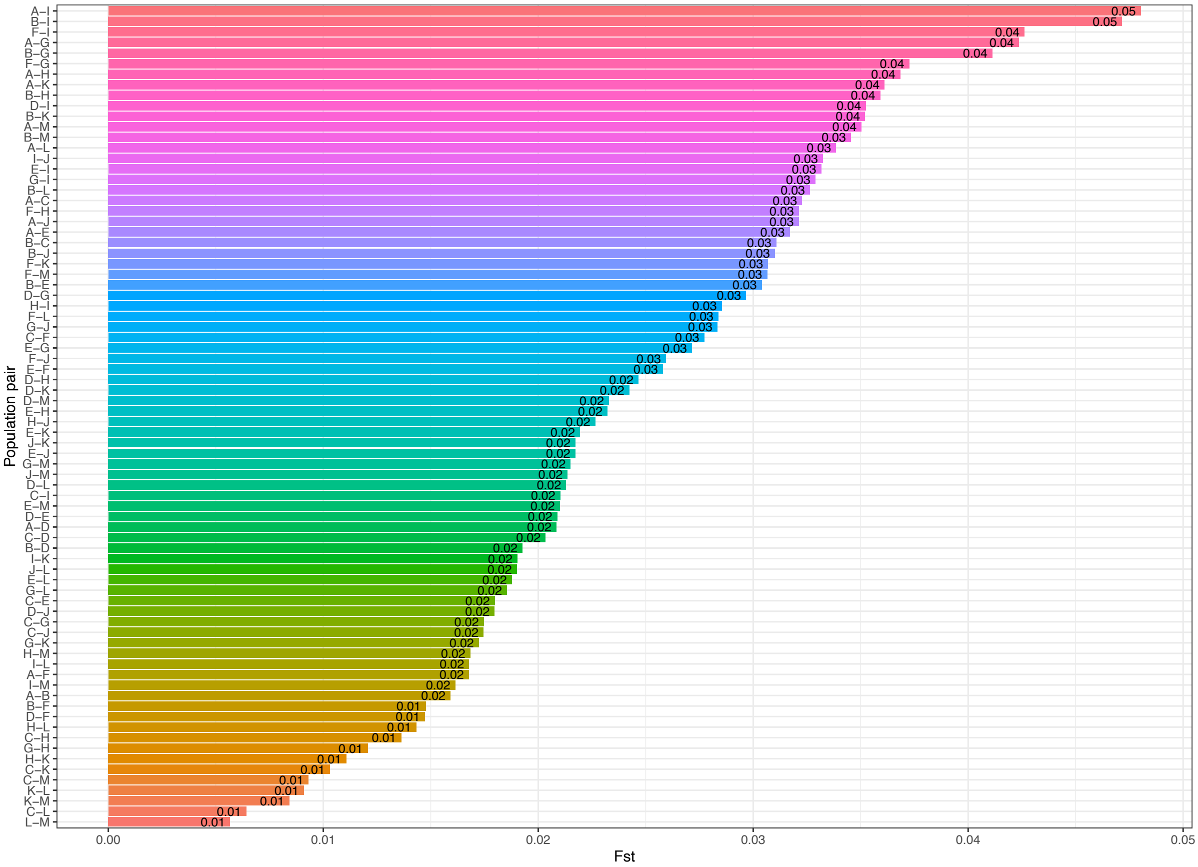

Population pair
Fst
0.00
0.01
0.02
0.03
0.04
0.05

**b**

simulated topology no. 1

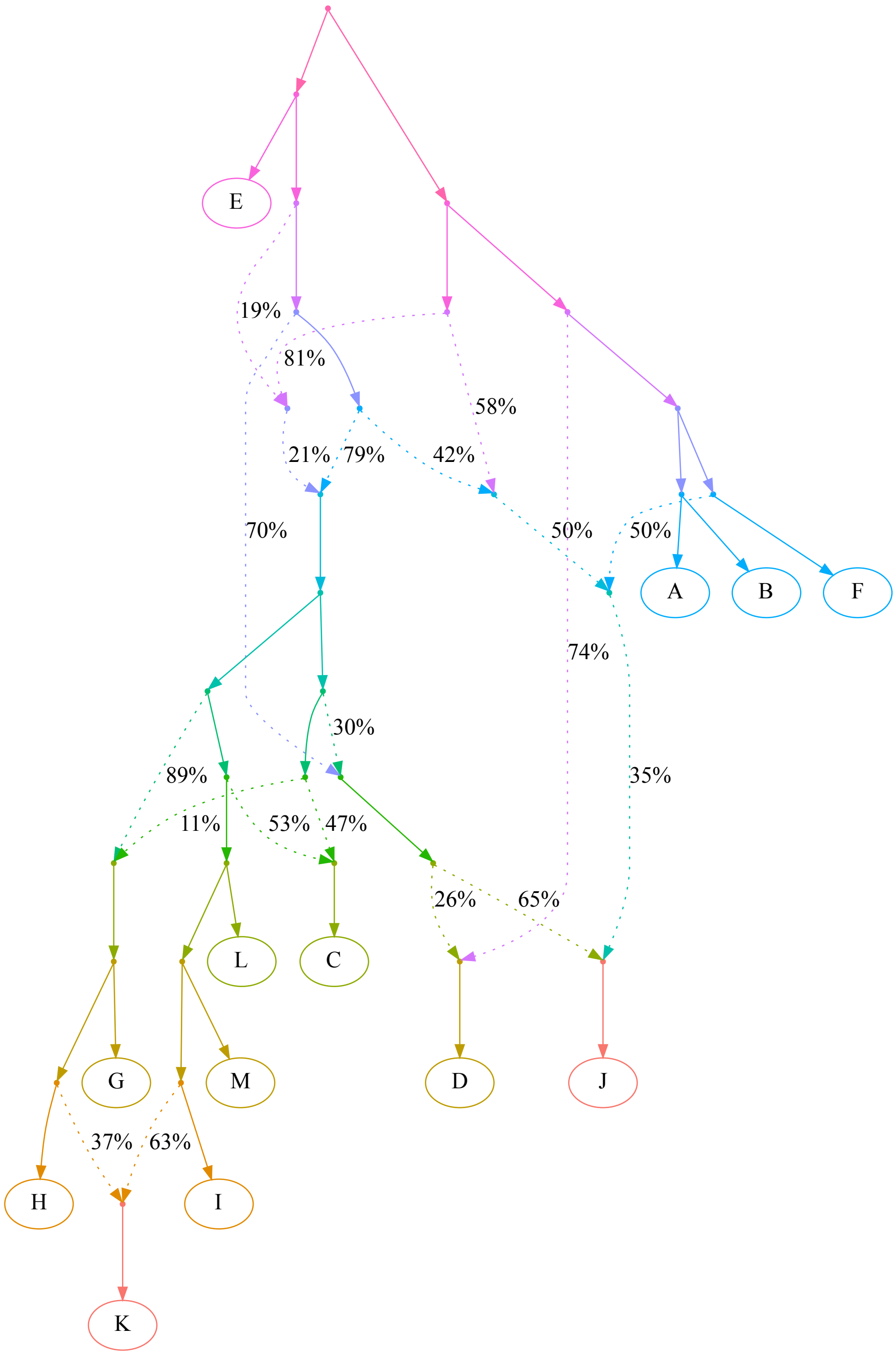

**c**

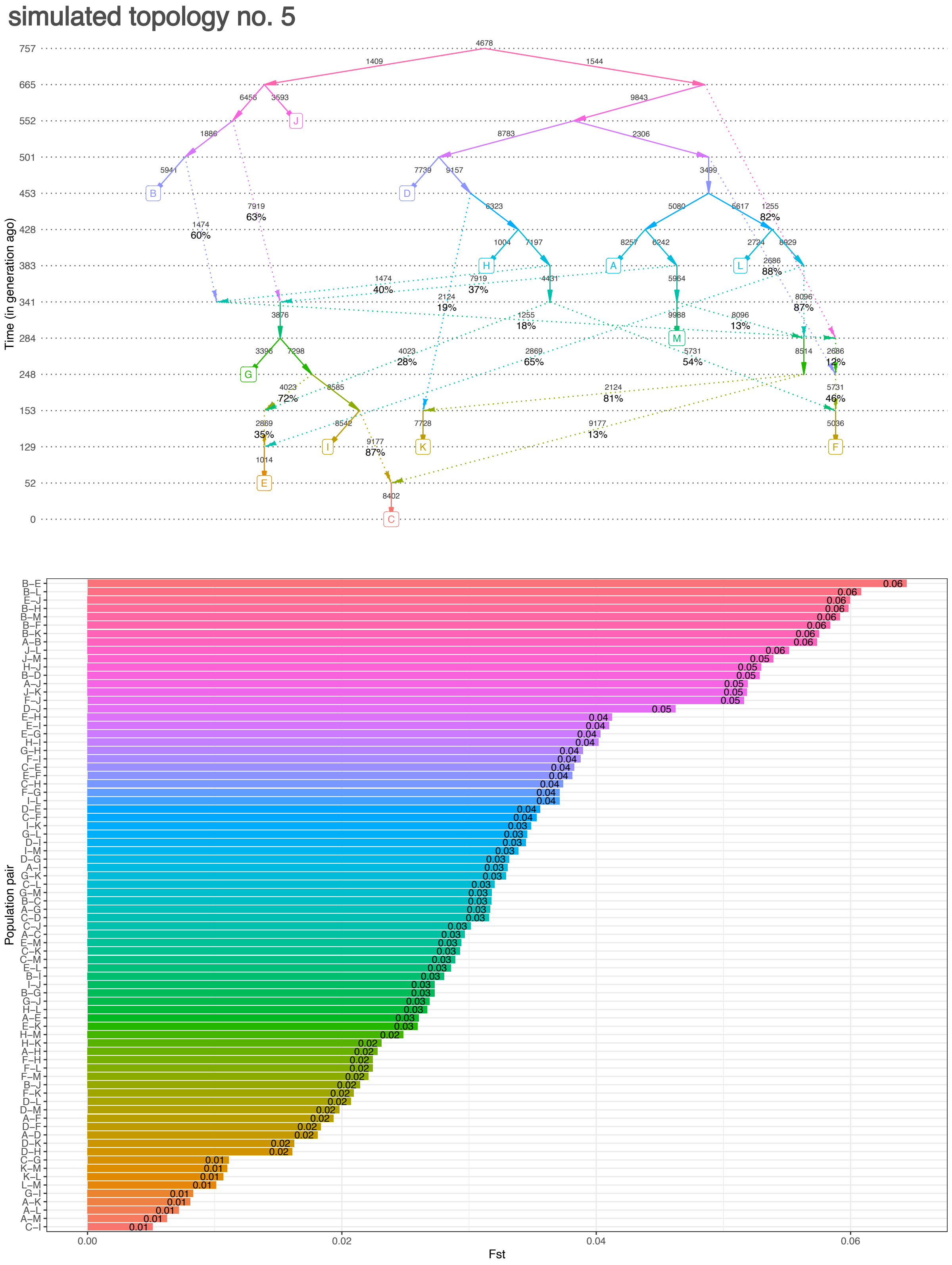
simulated topology no. 5
Time (in generation ago)
Population pair
Fst

**d**

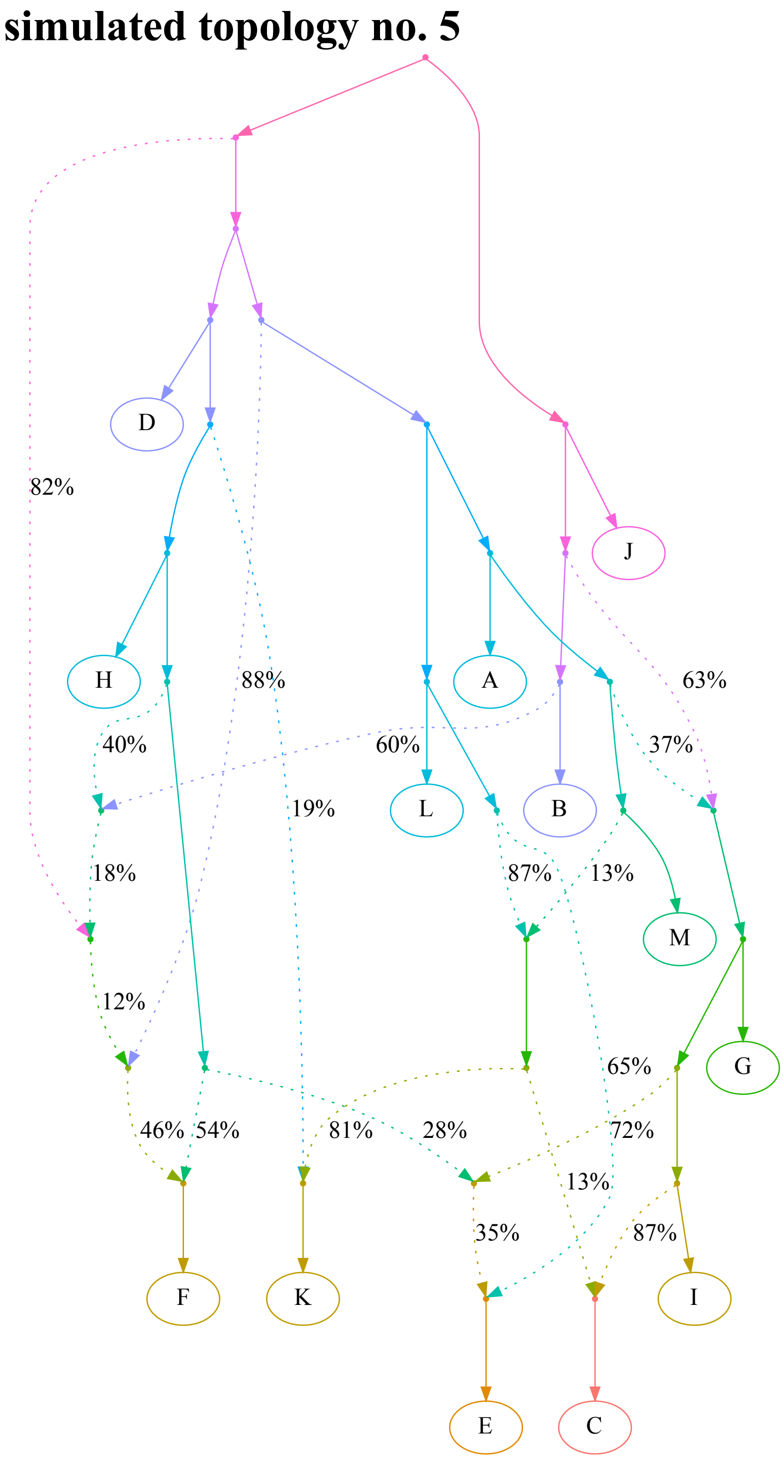
simulated topology no. 5
82%
88%
19%
63%
40%
60%
37%
18%
87%
13%
12%
65%
46%
54%
81%
28%
72%
13%
35%
87%
D
J
H
A
L
B
M
G
F
K
I
E
C

**e**

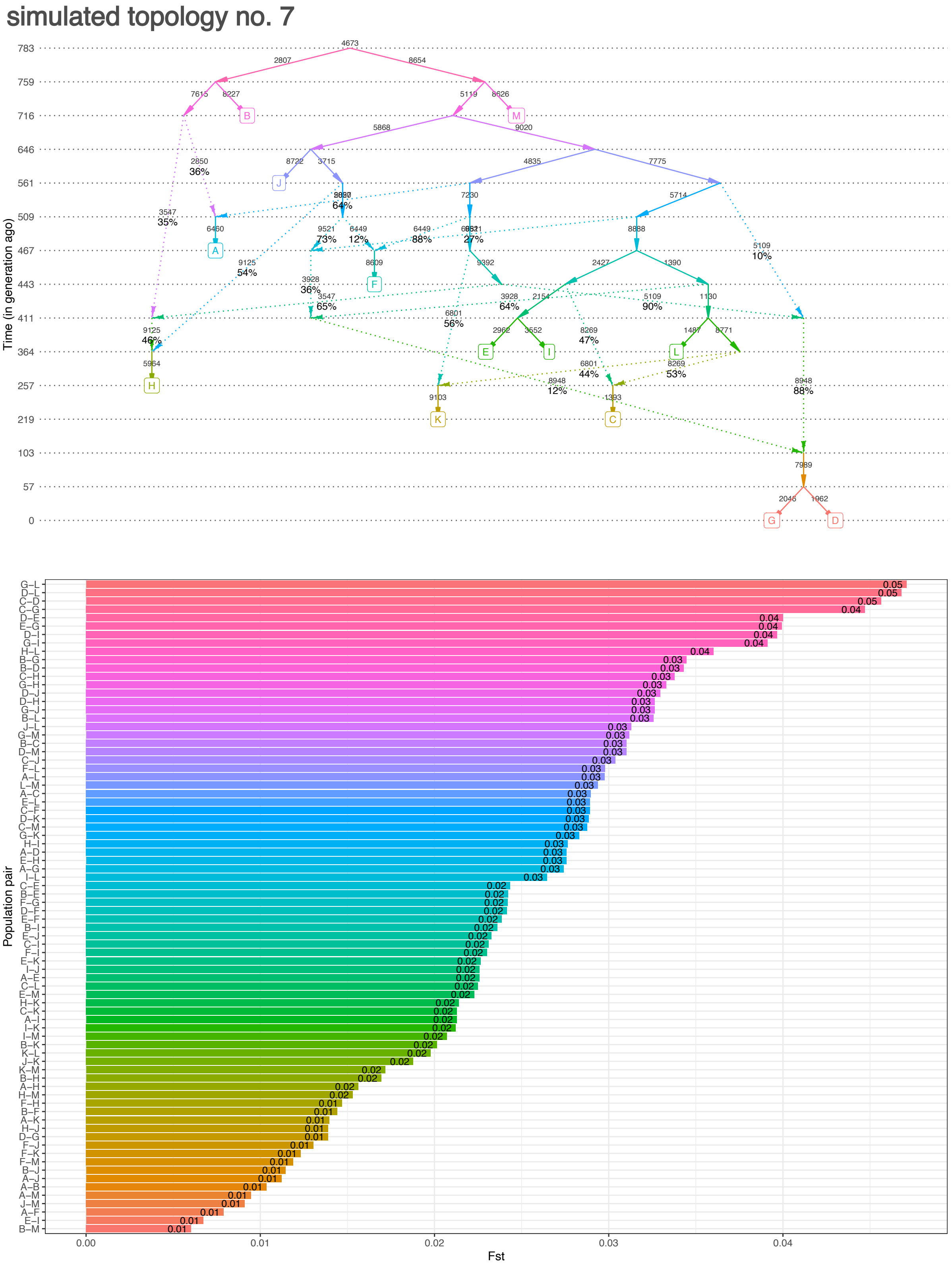
simulated topology no. 7
Time (in generation ago)
783
759
716
646
561
509
467
443
411
364
257
219
103
57
0
A
B
C
D
E
F
G
H
I
J
K
L
M
Population pair
Fst
0.00
0.01
0.02
0.03
0.04

**f**

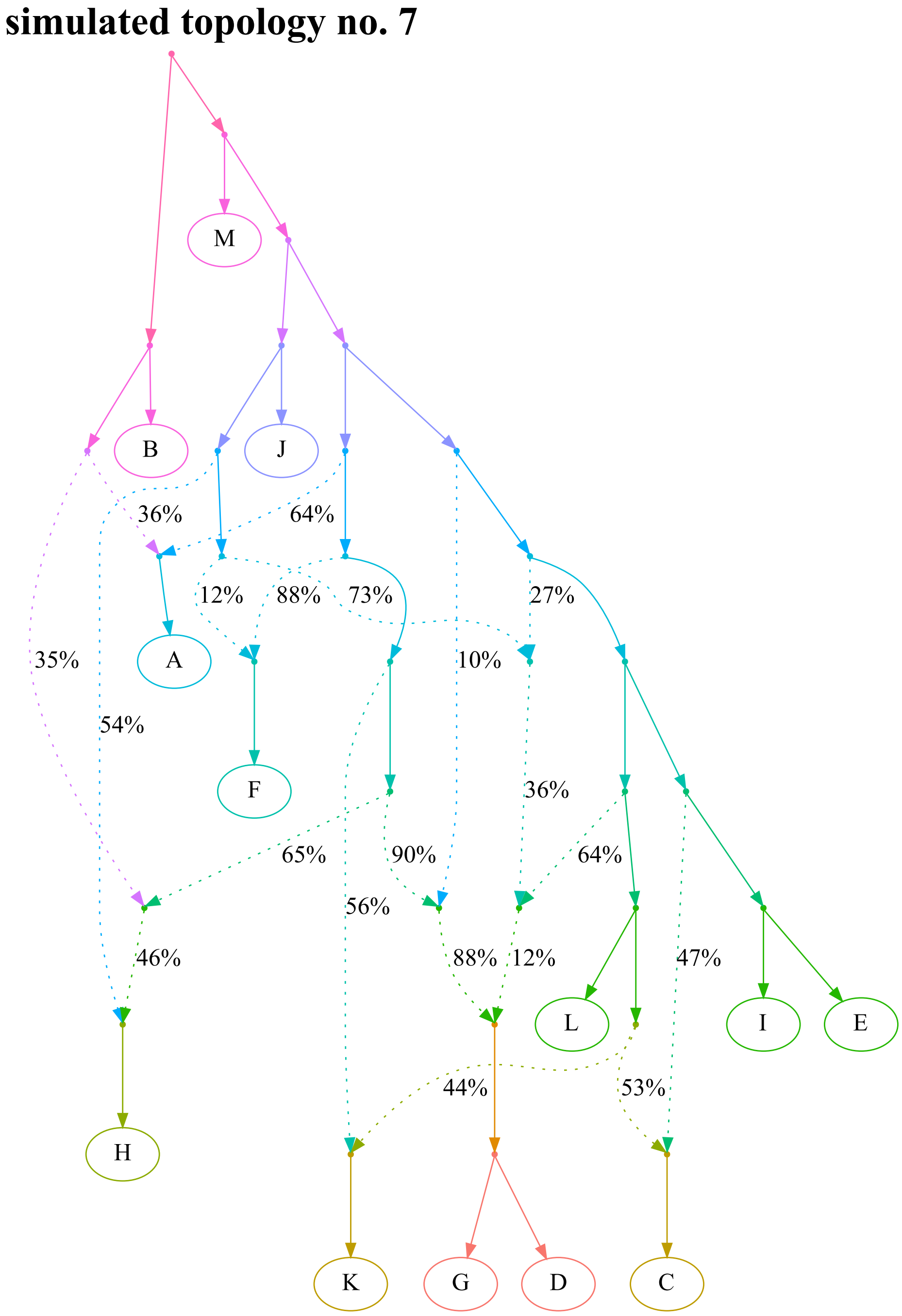
simulated topology no. 7
M
B
J
36%
64%
12%
88%
73%
27%
A
35%
10%
54%
F
36%
65%
90%
64%
56%
46%
88%
12%
47%
L
I
E
44%
53%
H
K
G
D
C

**g**

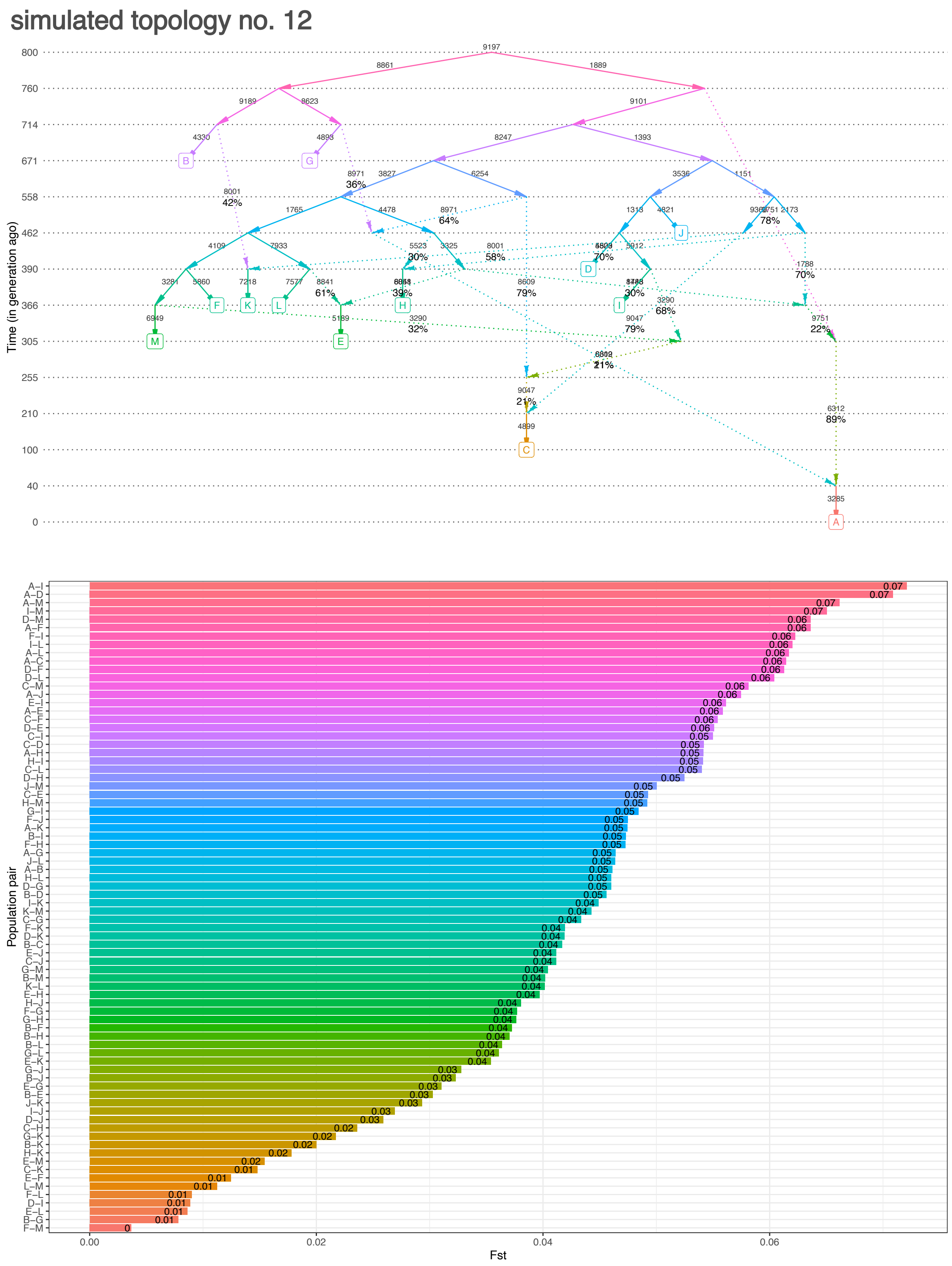
simulated topology no. 12
Time (in generation ago)
800
760
714
671
558
462
390
366
305
255
210
100
40
0
B
G
J
D
F
K
L
H
I
M
E
C
A
Population pair
Fst
0.00
0.02
0.04
0.06

**h**

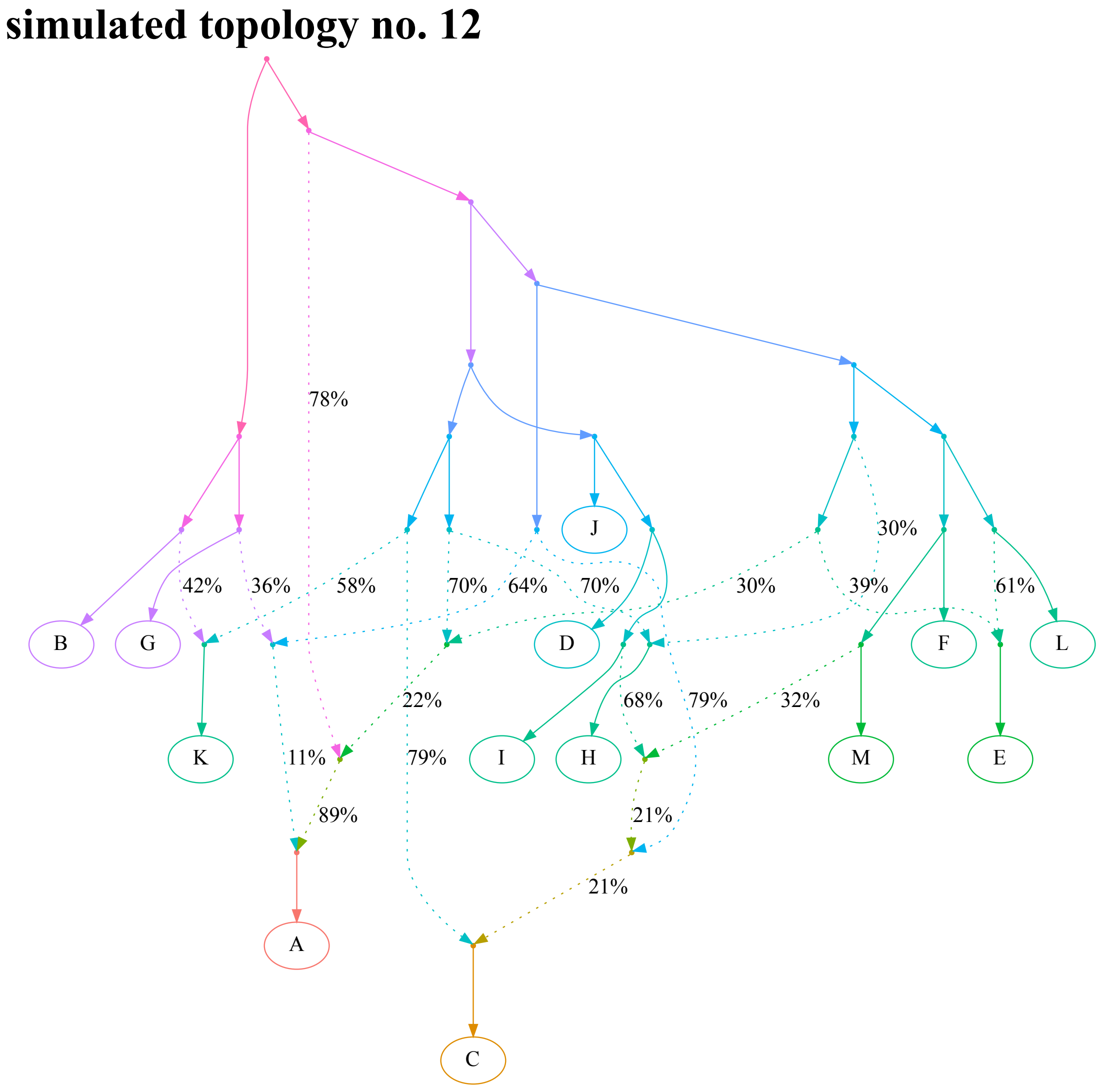
simulated topology no. 12
78%
J
30%
42%
36%
58%
70%
64%
70%
30%
39%
61%
B
G
D
F
L
22%
68%
79%
32%
K
11%
79%
I
H
M
E
89%
21%
21%
A
C

**i**

simulated topology no. 17

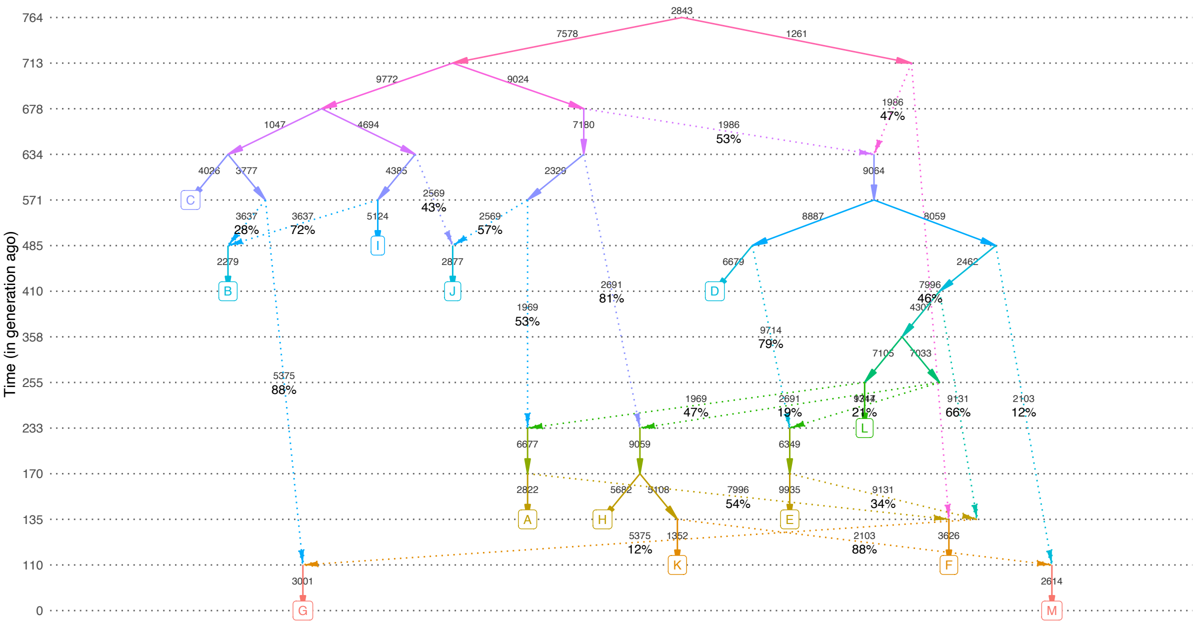


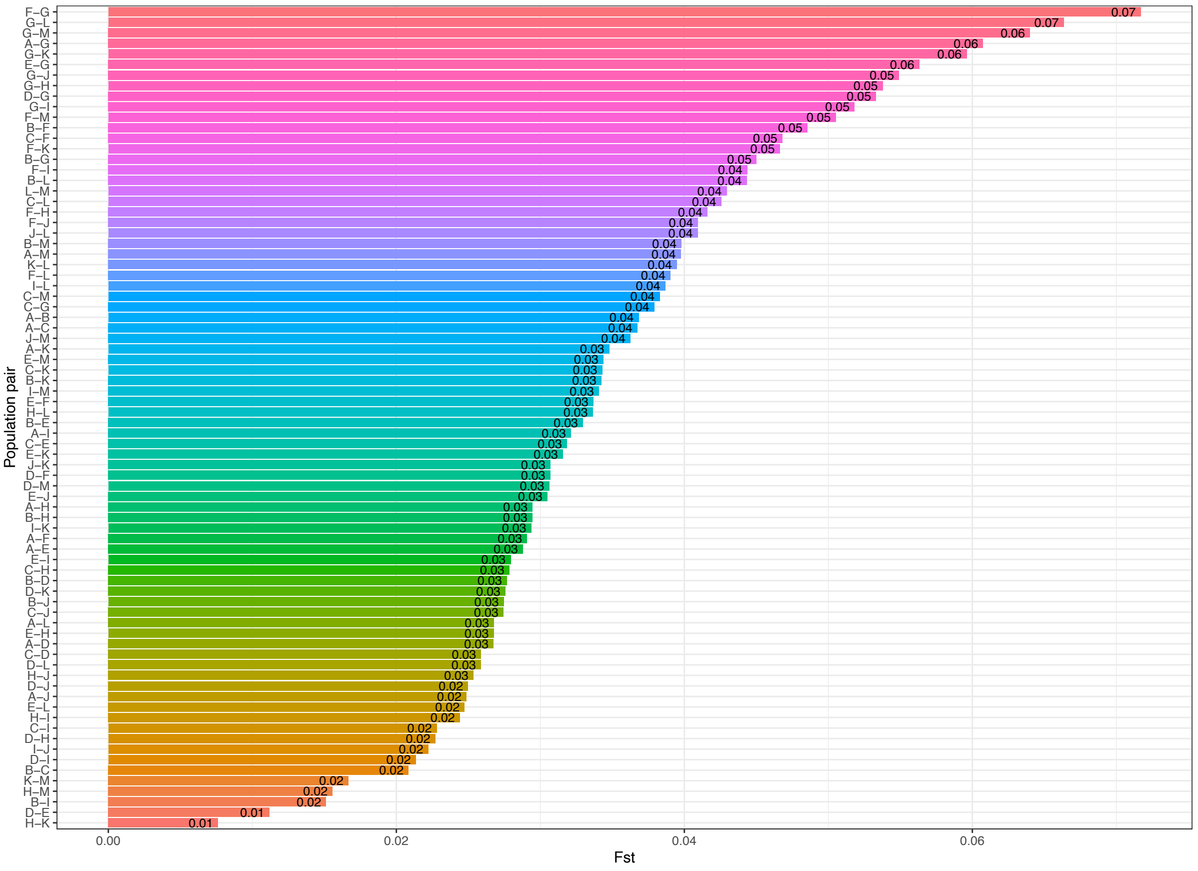

**j**

#### simulated topology no. 17

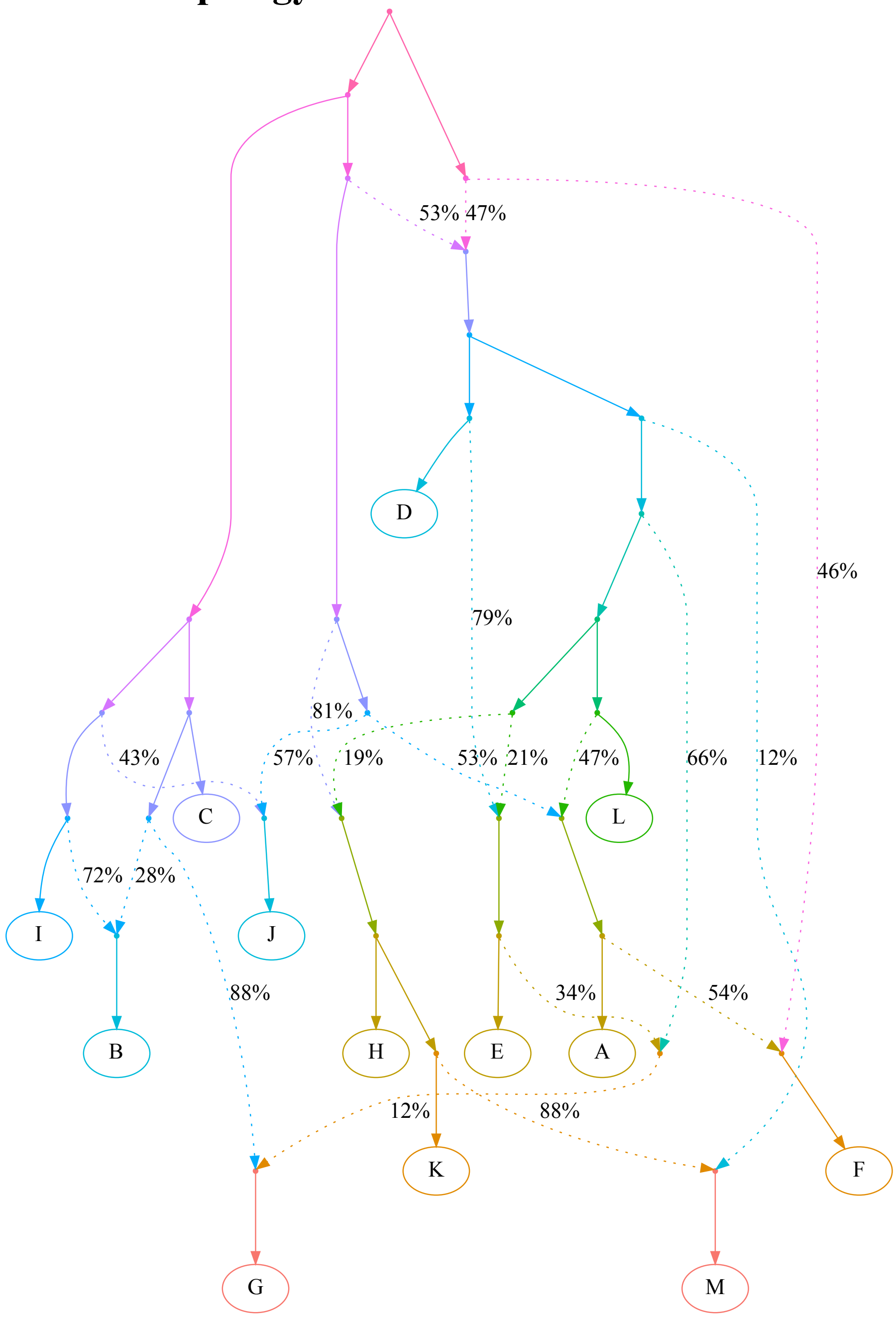

**k**

simulated topology no. 19

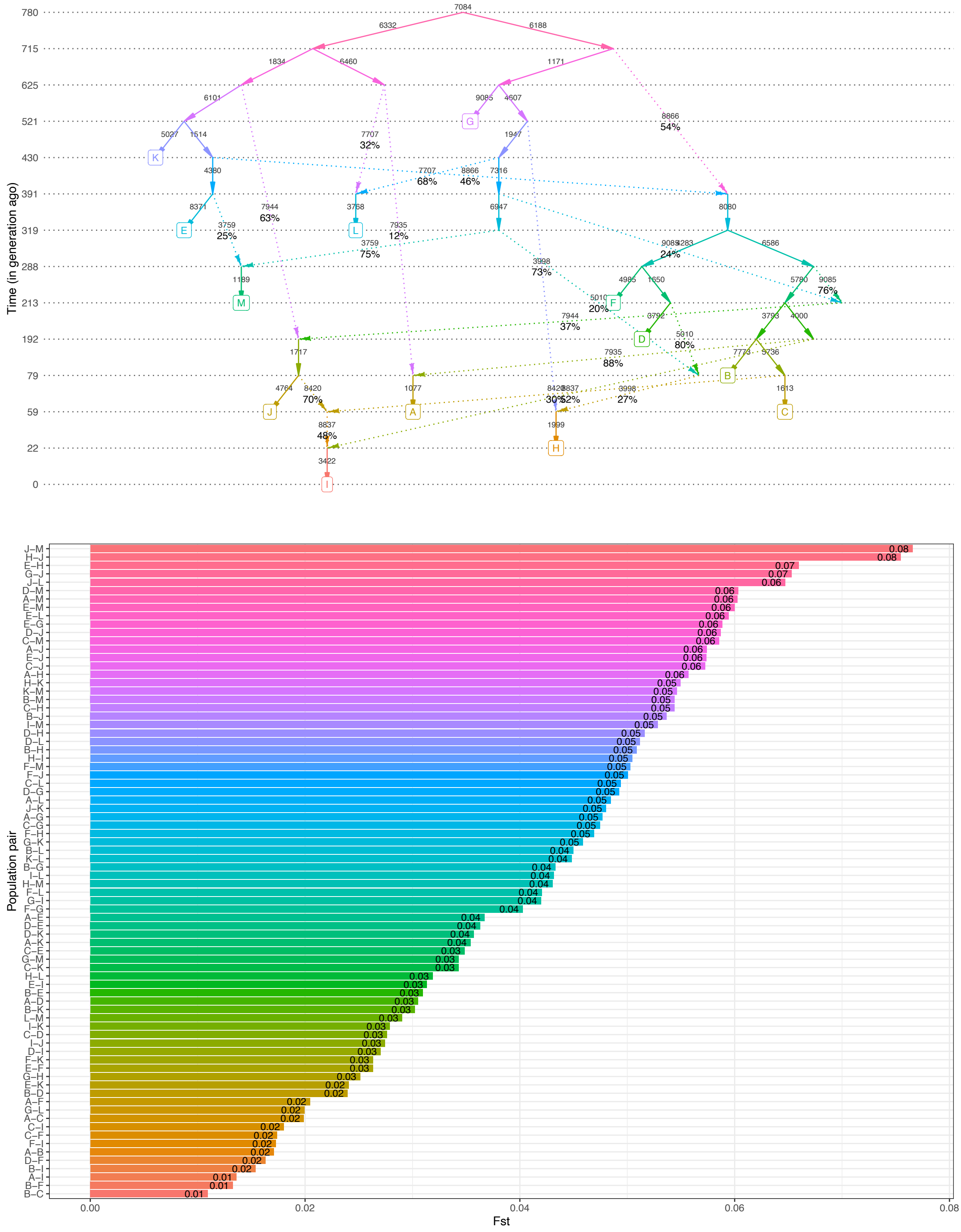

#### simulated topology no. 19

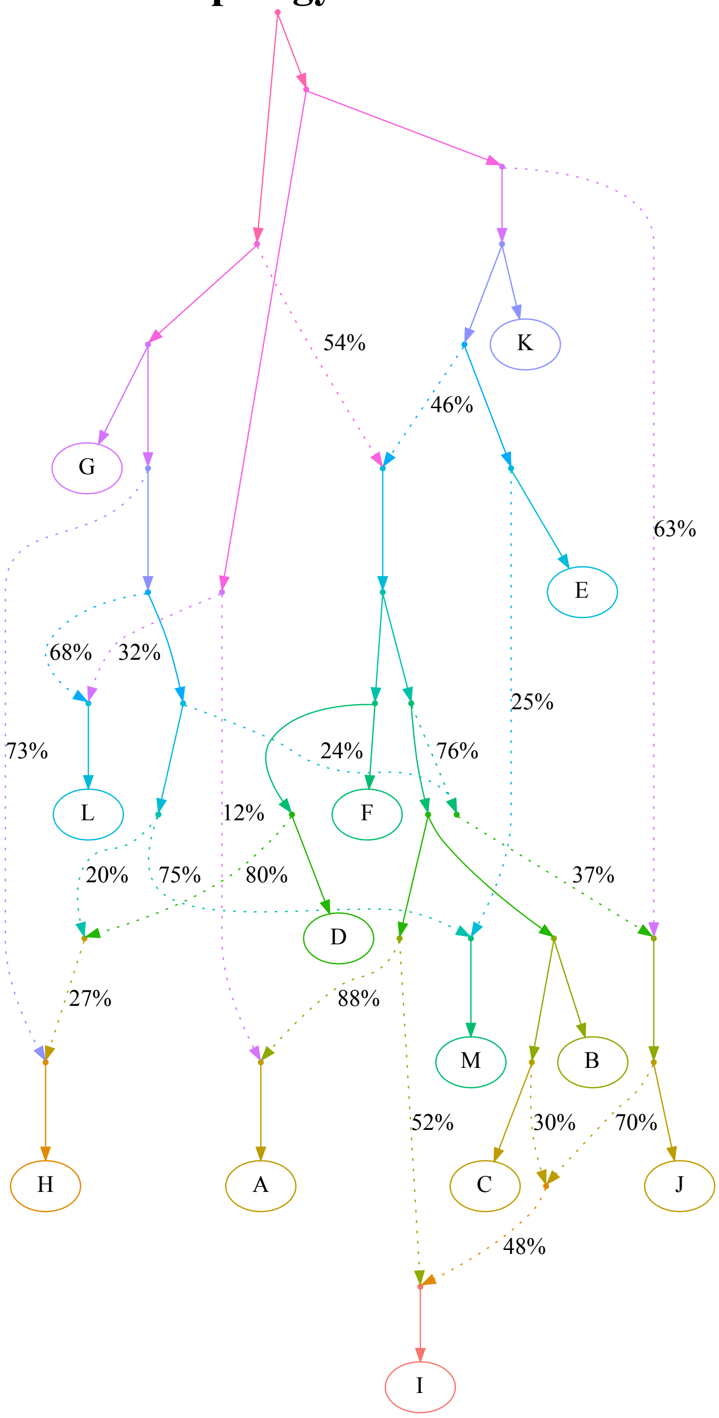

**m**

#### simulated topology no. 20

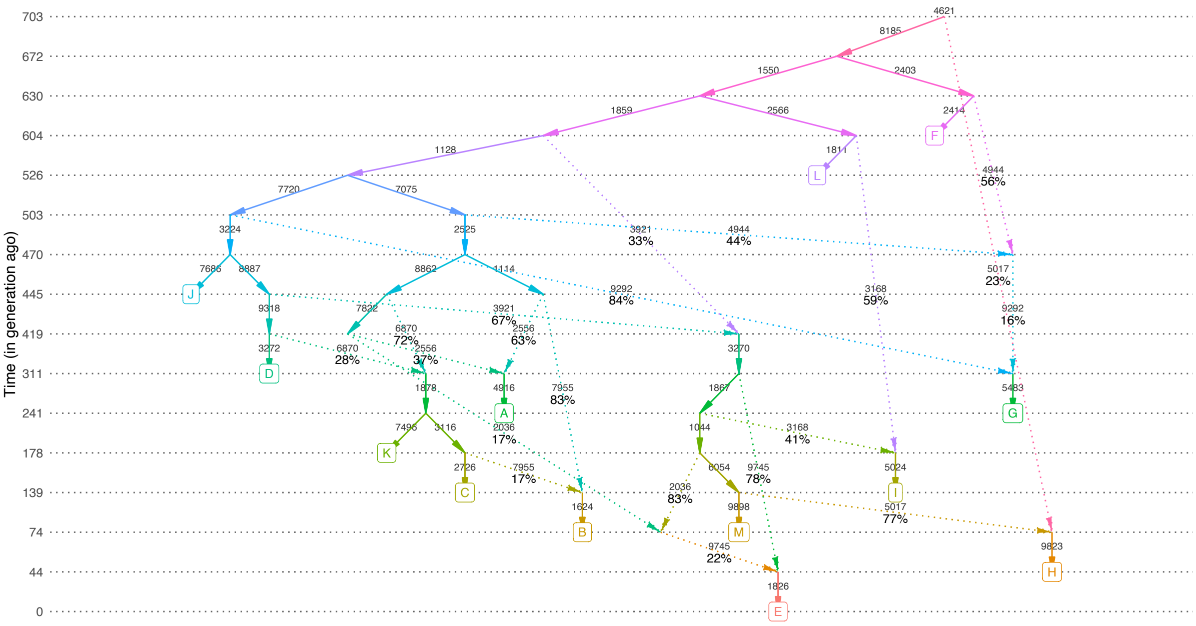


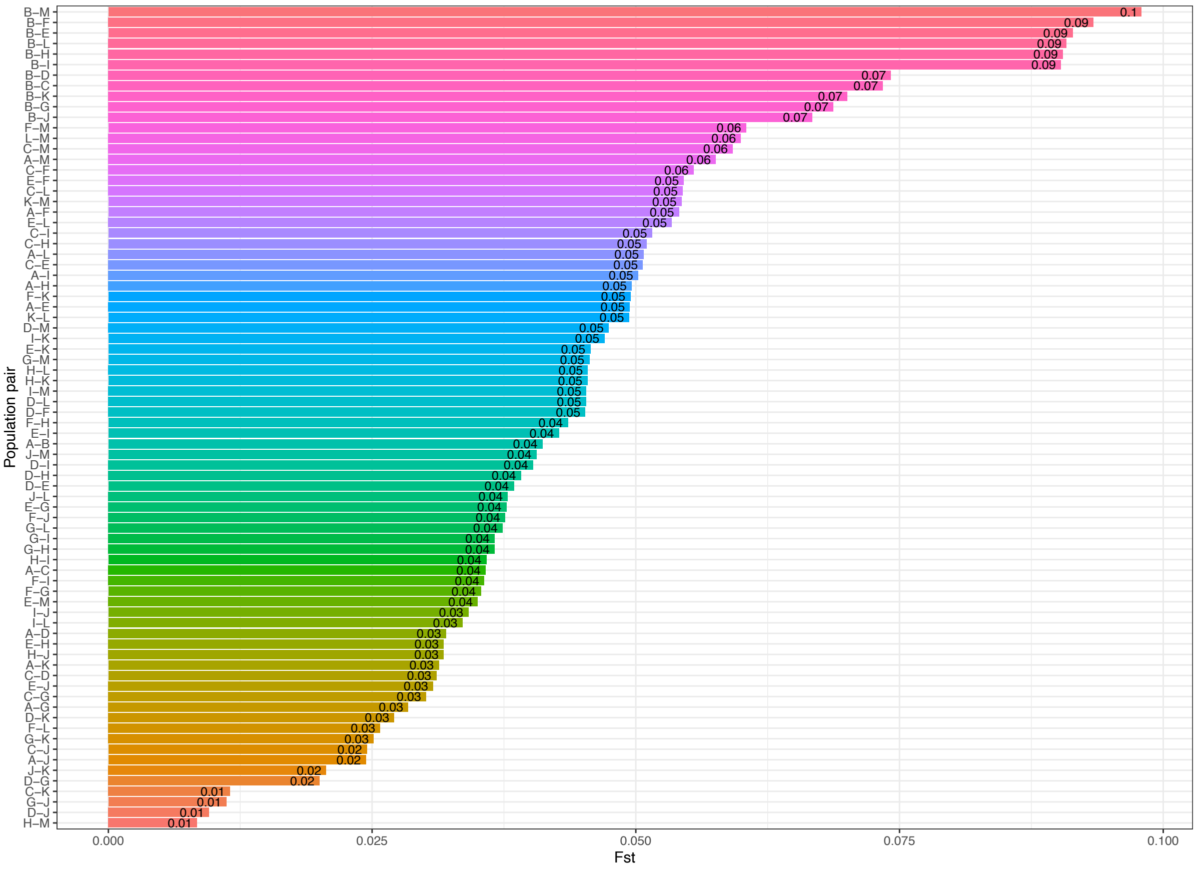

n

### simulated topology no. 20

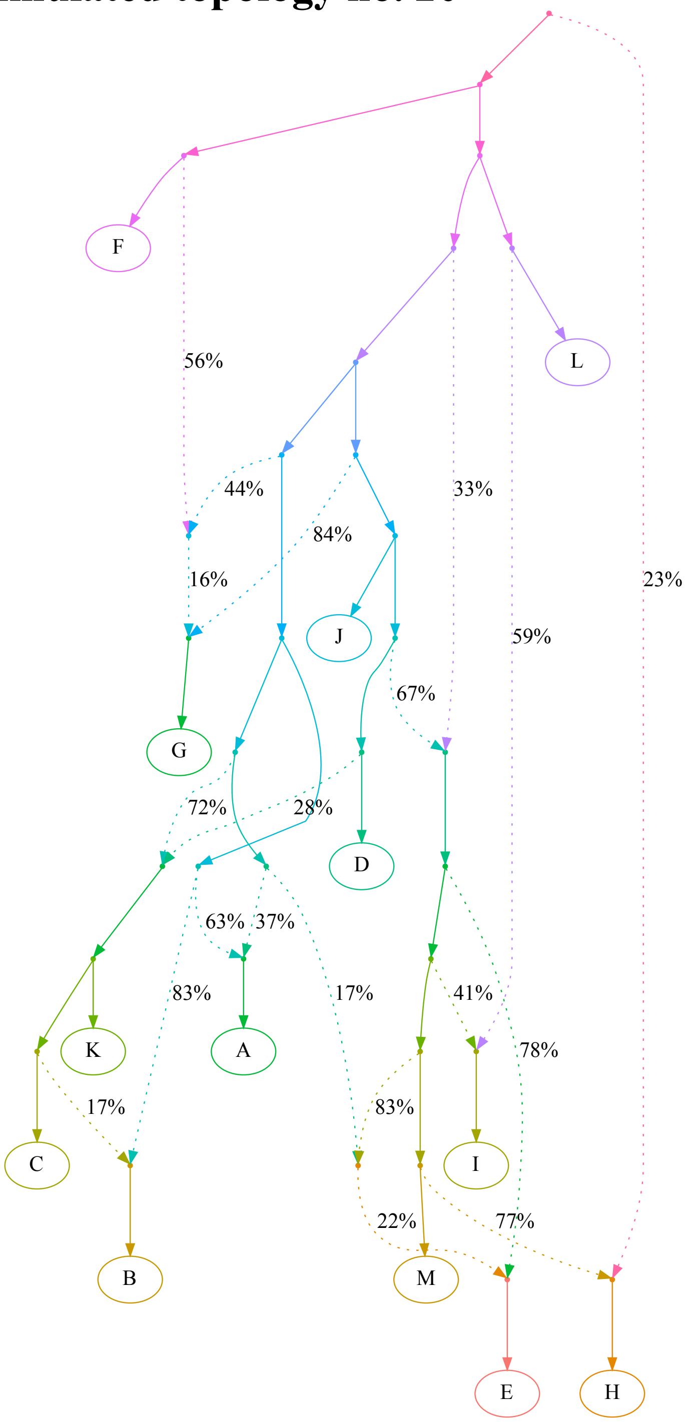

**o**

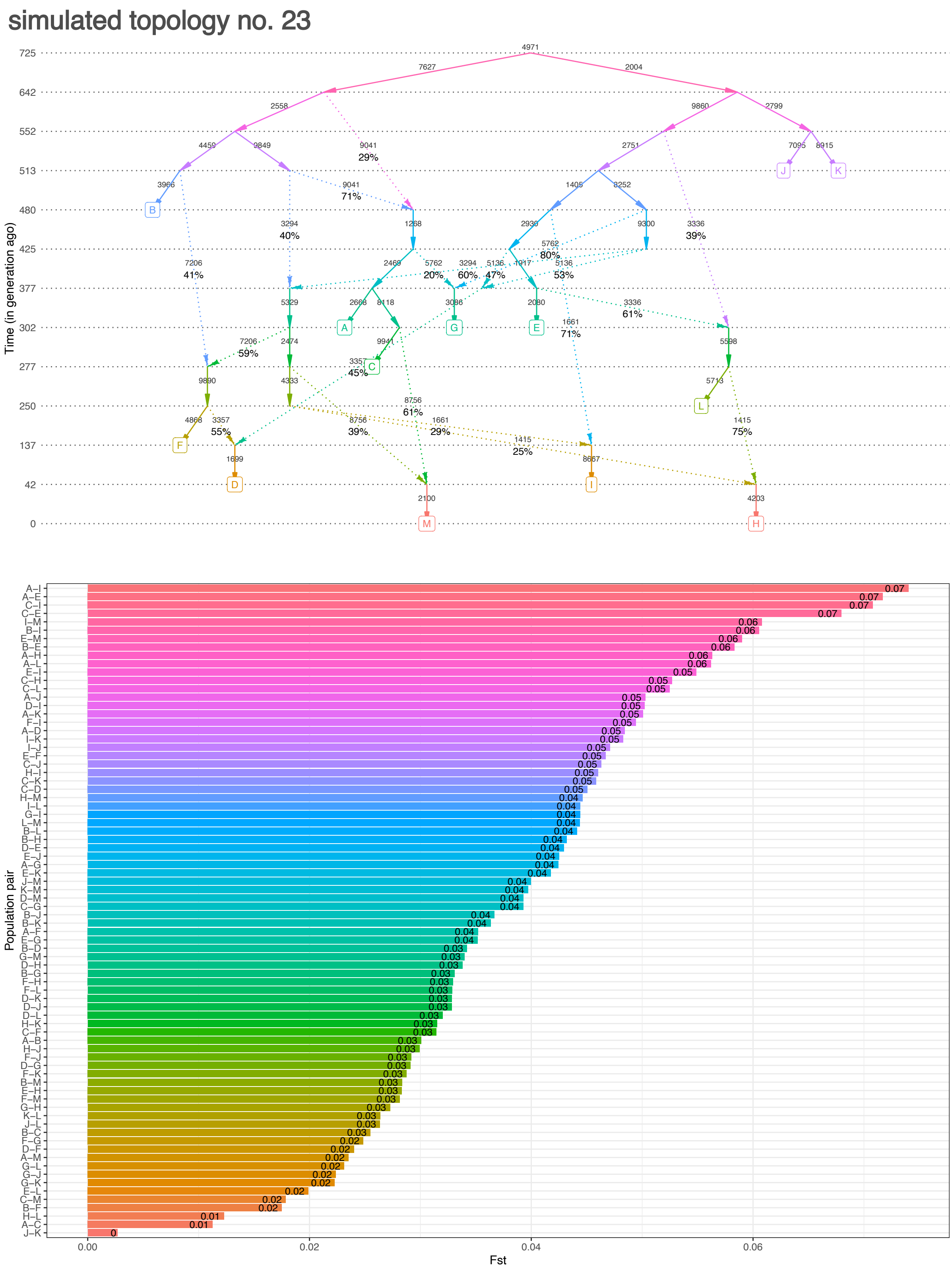

simulated topology no. 23
Time (in generation ago)
Population pair
Fst

**p**

### simulated topology no. 23

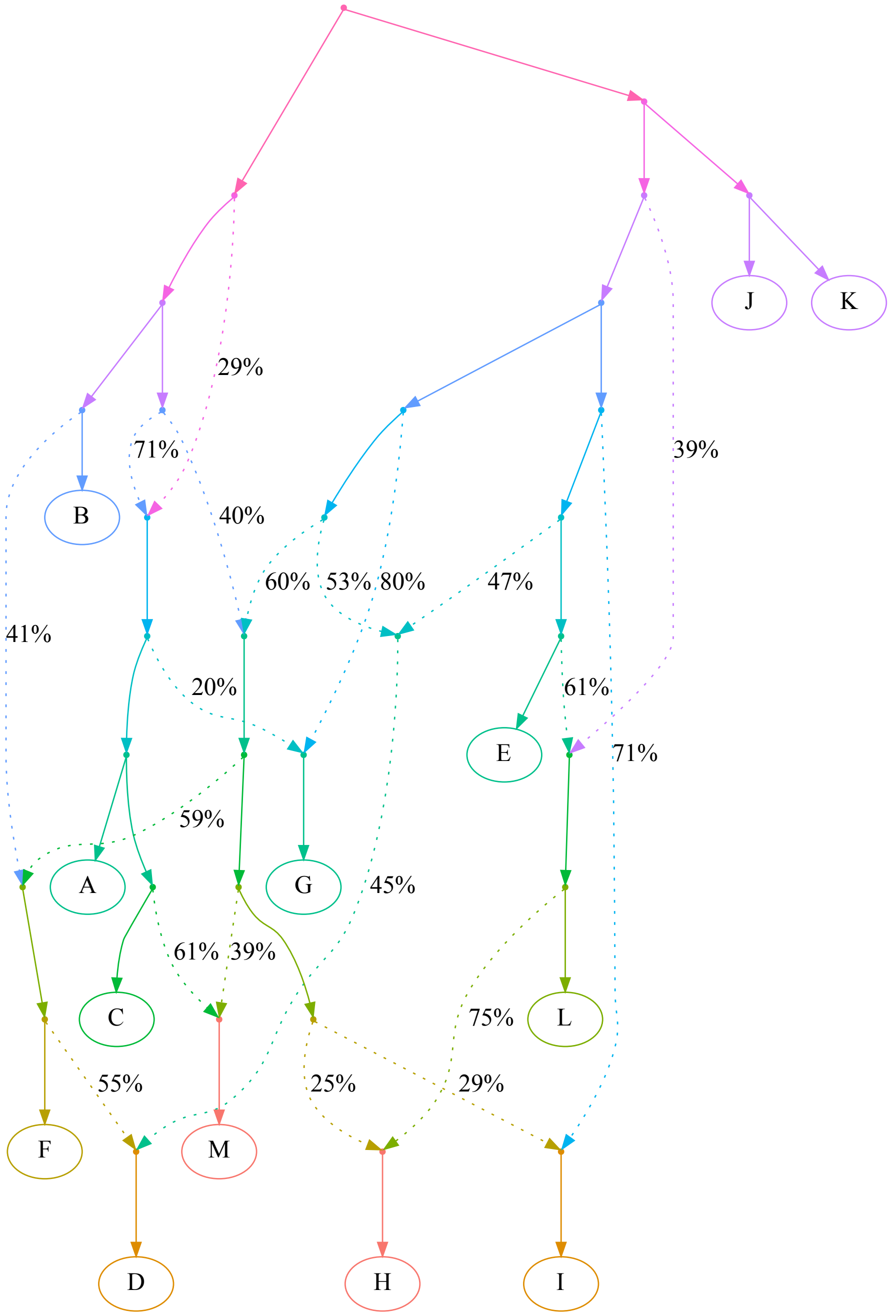

**q**

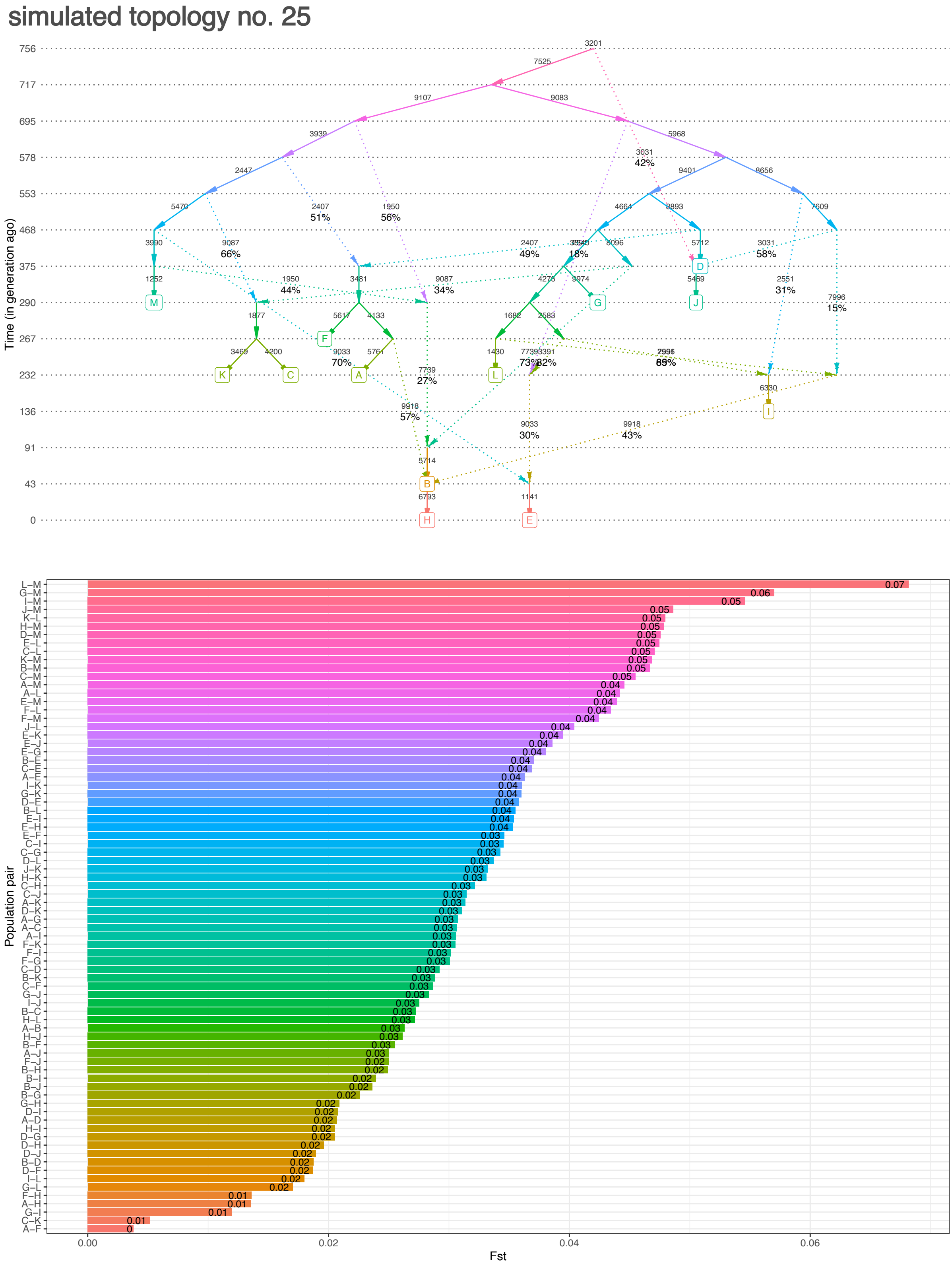

simulated topology no. 25
Time (in generation ago)
756
717
695
578
553
468
375
290
267
232
136
91
43
0
M
F
K
C
A
L
G
J
D
I
B
H
E
Population pair
Fst
0.00
0.02
0.04
0.06

r

### simulated topology no. 25

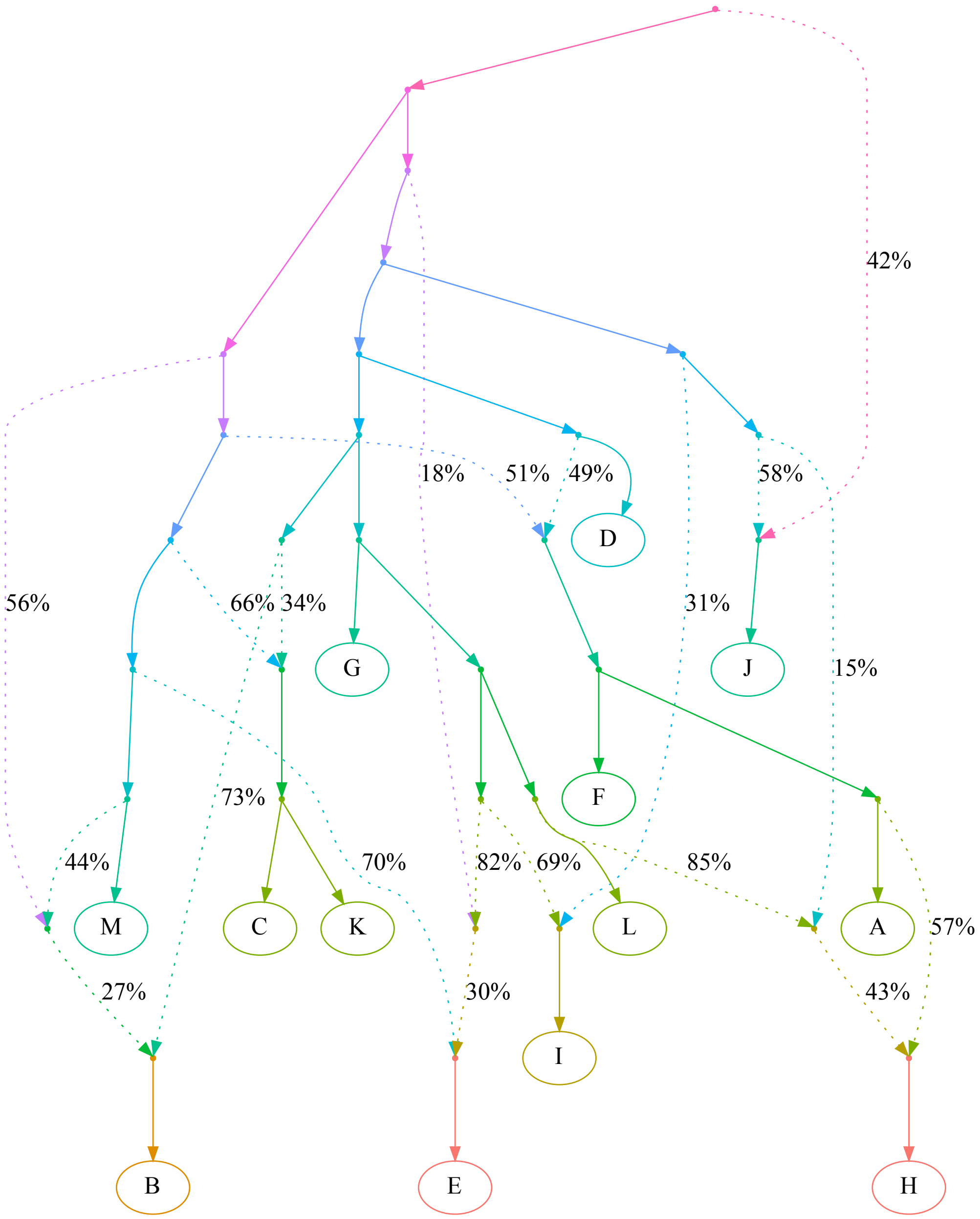

**s**

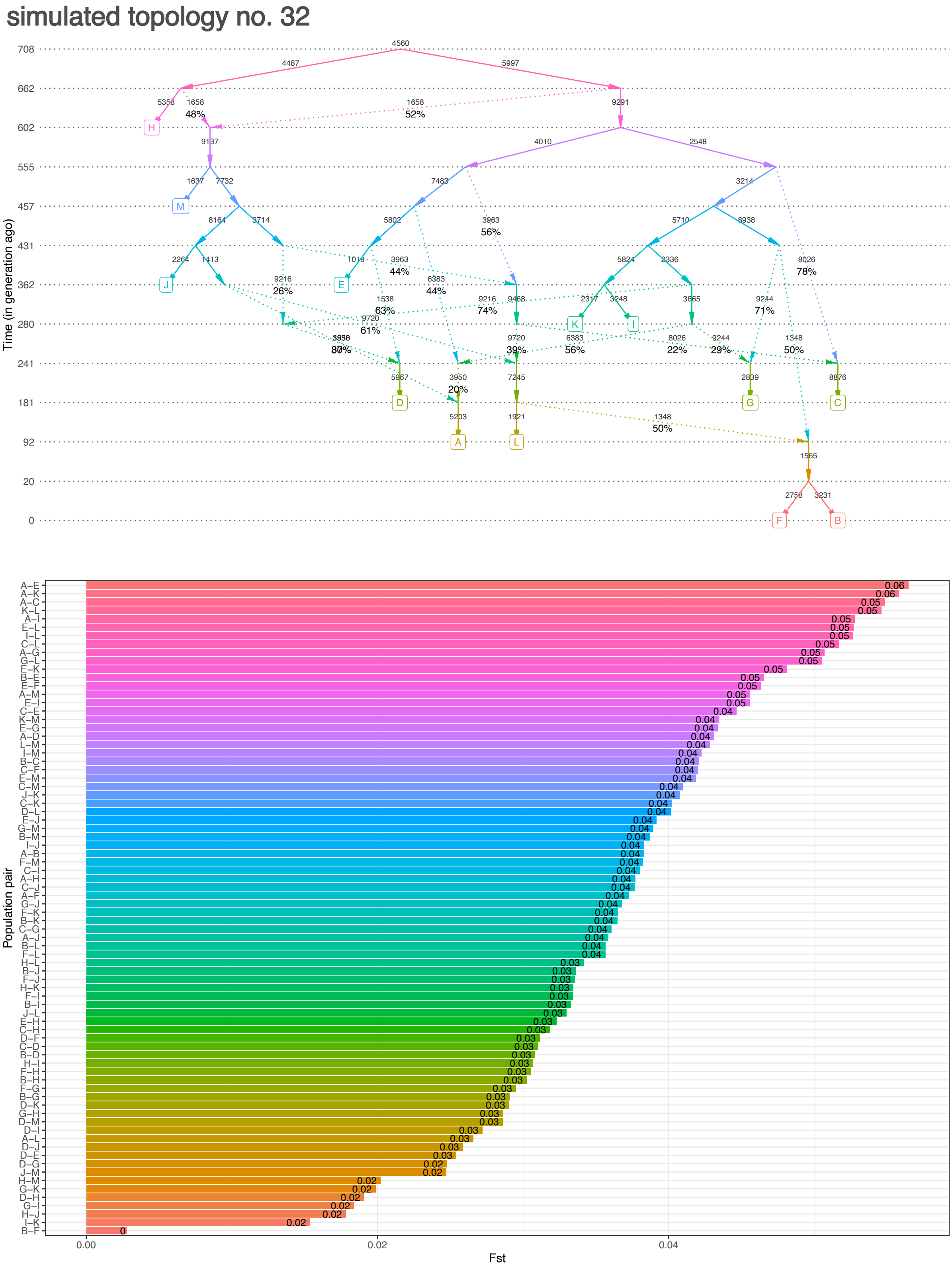

simulated topology no. 32
Time (in generation ago)
708
662
602
555
457
431
362
280
241
181
92
20
0
H
M
J
E
K
I
D
G
C
A
L
F
B
Population pair
Fst
0.00
0.02
0.04

**t**

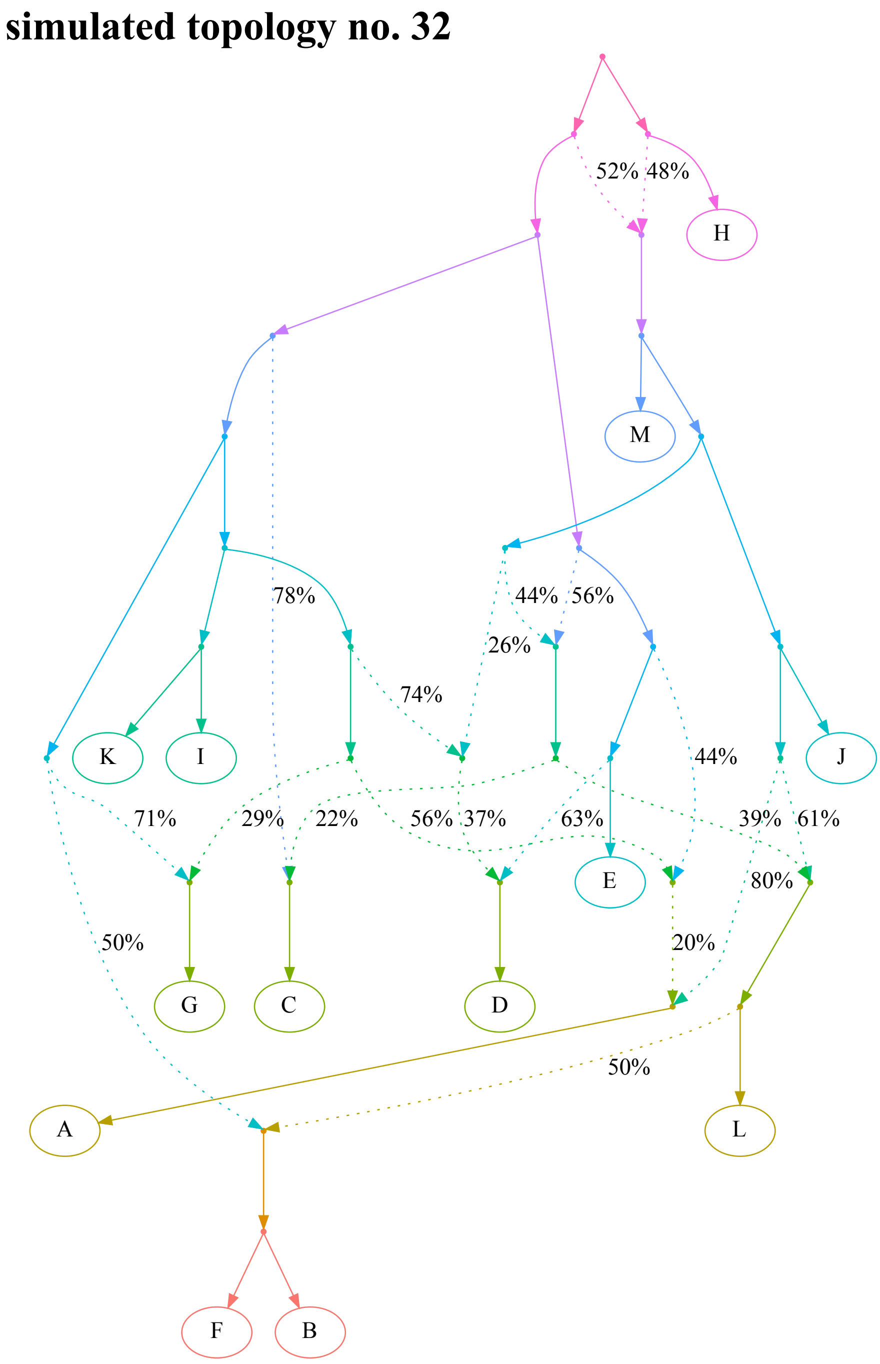


Simulated genetic histories in the shape of random admixture graphs including 13 populations and 10 pulse-like admixture events. Ten topologies are shown as examples, along with $F_{ST}$ values for all population pairs: topologies no. 1 (**a, b**), 5, (**c, d**), 7 (**e, f**), 12 (**g, h**), 17 (**i, j**), 19 (**k, l**), 20 (**m, n**), 23 (**o, p**), 25 (**q, r**), and 32 (**s, t**). These topologies correspond to the case studies presented in **Figs. 2** and **S6**. In all cases, $F_{ST}$ values are shown for simulation replicate no. 1, 300-Mbp-sized genomes, divergence times up to 800 generations, and diploid data. Time in generations is shown on the y-axis (for visual clarity, points on this axis are not spaced proportionately). Effective population sizes (in diploid individuals) are shown beside each edge. Sampled populations are labelled by letters in squares. Gene flows going from the past to the future were simulated as unsampled ghost populations. The same simulated histories are shown in a form more accessible to topological analysis, but without any additional information, in panels **b**, **d**, **f**, **h**, **j, l, n, p, r, t**. The color gradients have no special meaning and are used for visual clarity only.

**Figure S4.**

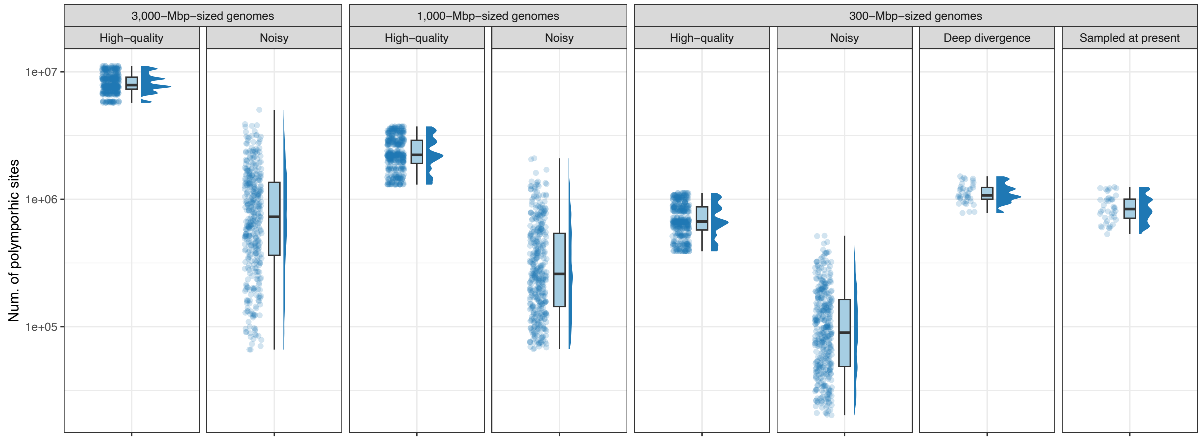


Number of SNPs with no missing data at the group level that are polymorphic in 13 simulated populations. SNP counts are summarized across simulated graph topologies and simulation replicates (for high-quality data) or subsampling replicates (for noisy pseudohaploid data). In the case of the “deep divergence” and “sampled at present” simulation setups, only one simulation replicate was generated.

**Figure S5.**

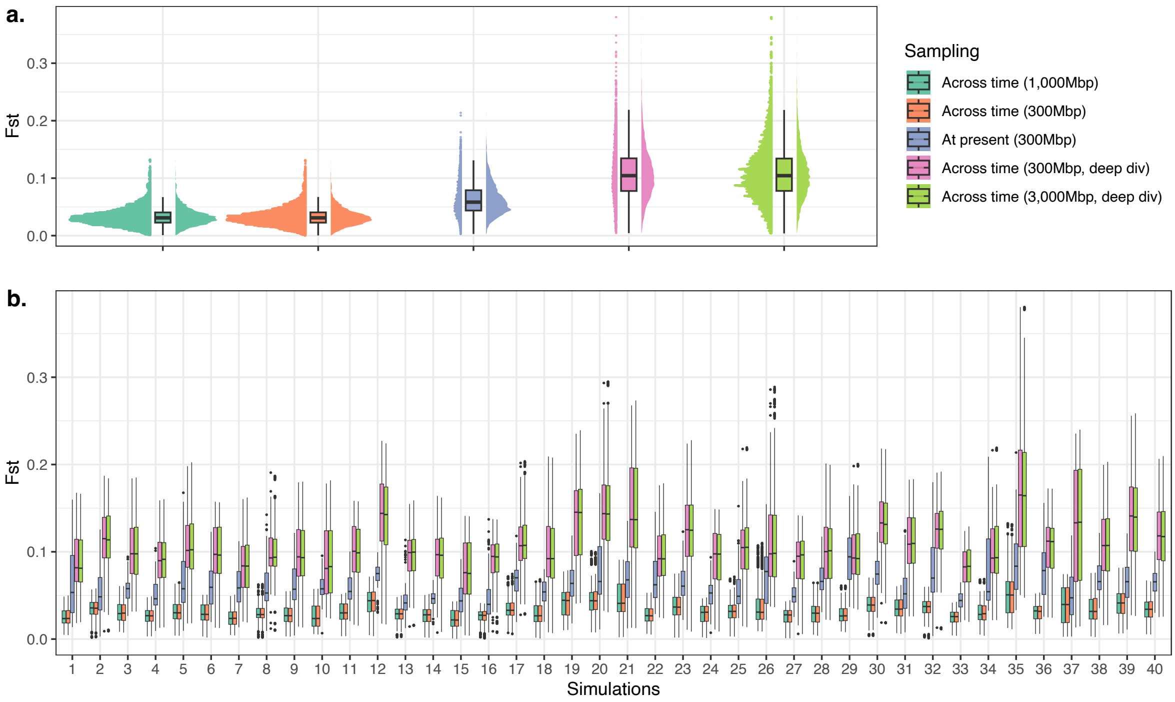


$F_{ST}$ distributions for five sets of simulations: 1 & 2) populations sampled at different points in time and with maximal graph depth at 800 generations, 1,000- or 300-Mbp-sized genomes, 10 simulation replicates per topology (in blue-green and orange, respectively); 3) all populations sampled at "present" and with maximal graph depth at 800 generations, 300-Mbp-sized genomes, one simulation replicate per topology (in blue); 4 & 5) populations sampled at different points in time and with maximal graph depth at 3,000 generations, 300- or 3,000-Mbp-sized genomes (in magenta and green), one or 10 simulation replicates per topology, respectively. In all cases, diploid data were used. The distributions are shown either for all 40 topologies pooled (**a**), or for each topology separately (**b**).

876 **Figure S6.**

877 **a**

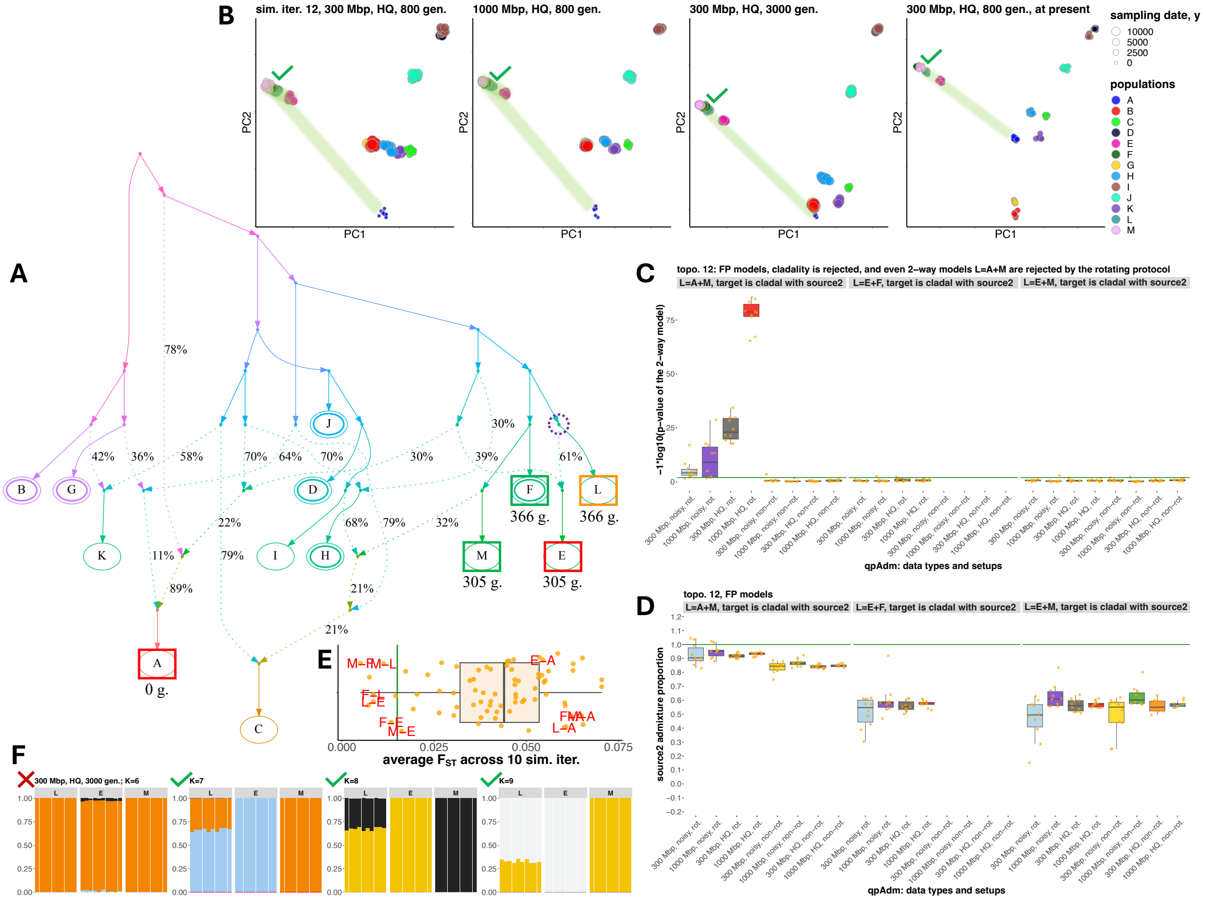


878

879

**b**

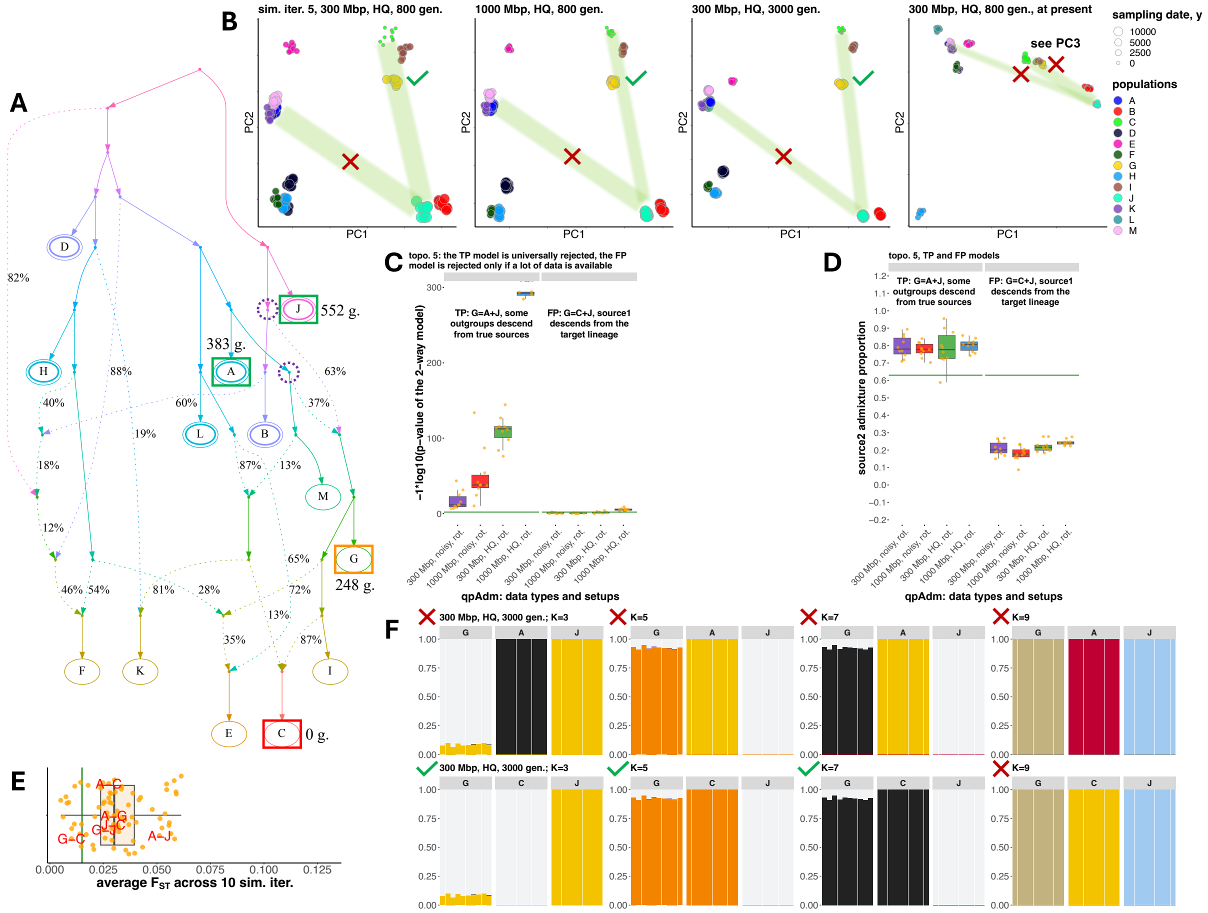

A
B
C
D
E
F
sim. iter. 5, 300 Mbp, HQ, 800 gen.
1000 Mbp, HQ, 800 gen.
300 Mbp, HQ, 3000 gen.
300 Mbp, HQ, 800 gen., at present
see PC3
PC1
PC2
sampling date, y
10000
5000
2500
0
populations
552 g.
383 g.
248 g.
0 g.
topo. 5: the TP model is universally rejected, the FP model is rejected only if a lot of data is available
TP: G=A+J, some outgroups descend from true sources
FP: G=C+J, source1 descends from the target lineage
−1*log10(p−value of the 2−way model)
qpAdm: data types and setups
topo. 5, TP and FP models
source2 admixture proportion
average F_ST across 10 sim. iter.
300 Mbp, HQ, 3000 gen.; K=3
K=5
K=7
K=9

c

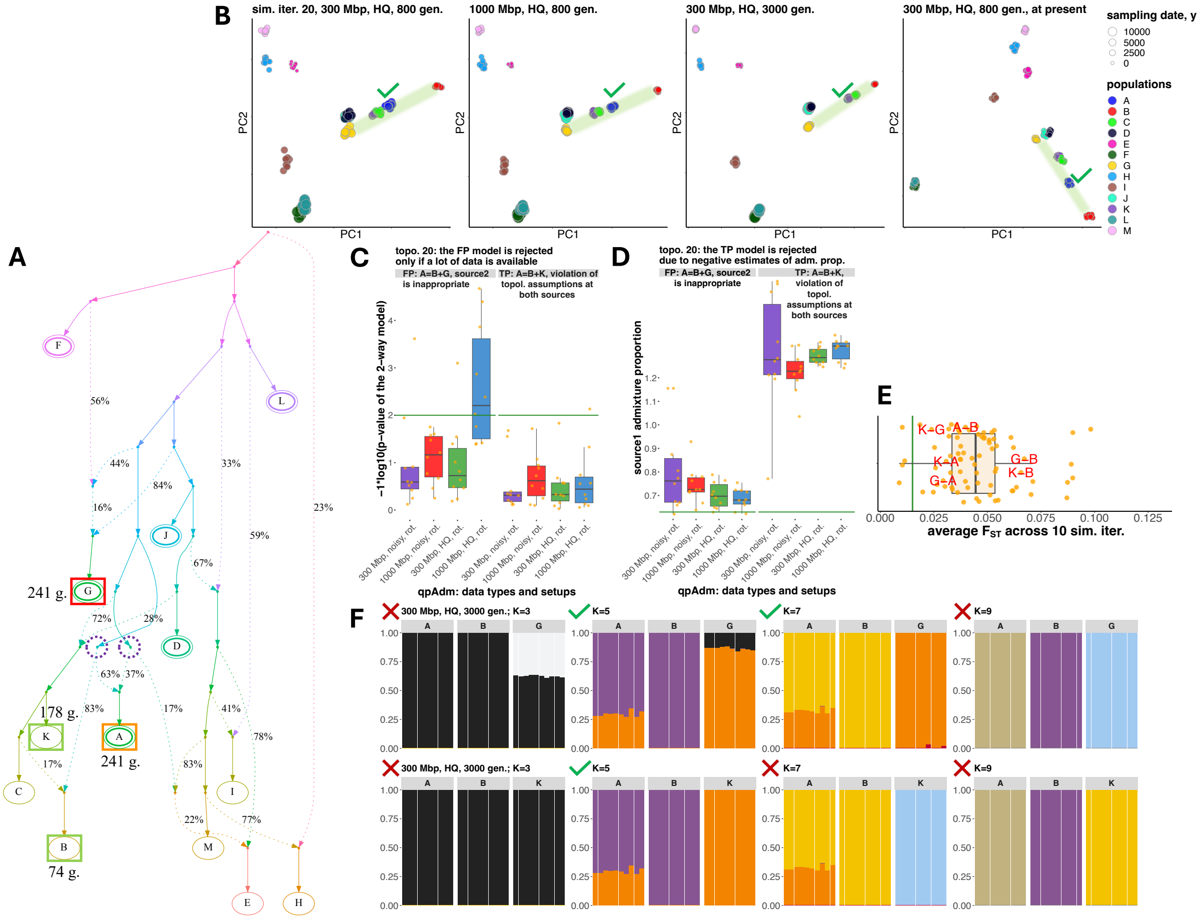

A
B
sim. iter. 20, 300 Mbp, HQ, 800 gen.
1000 Mbp, HQ, 800 gen.
300 Mbp, HQ, 3000 gen.
300 Mbp, HQ, 800 gen., at present
sampling date, y
10000
5000
2500
0
populations
A
B
C
D
E
F
G
H
I
J
K
L
M
PC1
PC2
C
topo. 20: the FP model is rejected only if a lot of data is available
FP: A=B+G, source2 is inappropriate
TP: A=B+K, violation of topol. assumptions at both sources
−1*log10(p-value of the 2-way model)
qpAdm: data types and setups
300 Mbp, noisy, rot.
1000 Mbp, noisy, rot.
300 Mbp, HQ, rot.
1000 Mbp, HQ, rot.
D
topo. 20: the TP model is rejected due to negative estimates of adm. prop.
FP: A=B+G, source2 is inappropriate
TP: A=B+K, violation of topol. assumptions at both sources
source1 admixture proportion
E
K–G
A–B
K–A
G–A
G–B
K–B
0.000
0.025
0.050
0.075
0.100
0.125
average $F_{ST}$ across 10 sim. iter.
F
300 Mbp, HQ, 3000 gen.; K=3
K=5
K=7
K=9
241 g.
178 g.
241 g.
74 g.
56%
44%
16%
84%
33%
23%
59%
67%
72%
28%
63%
37%
83%
17%
41%
78%
17%
83%
22%
77%

d

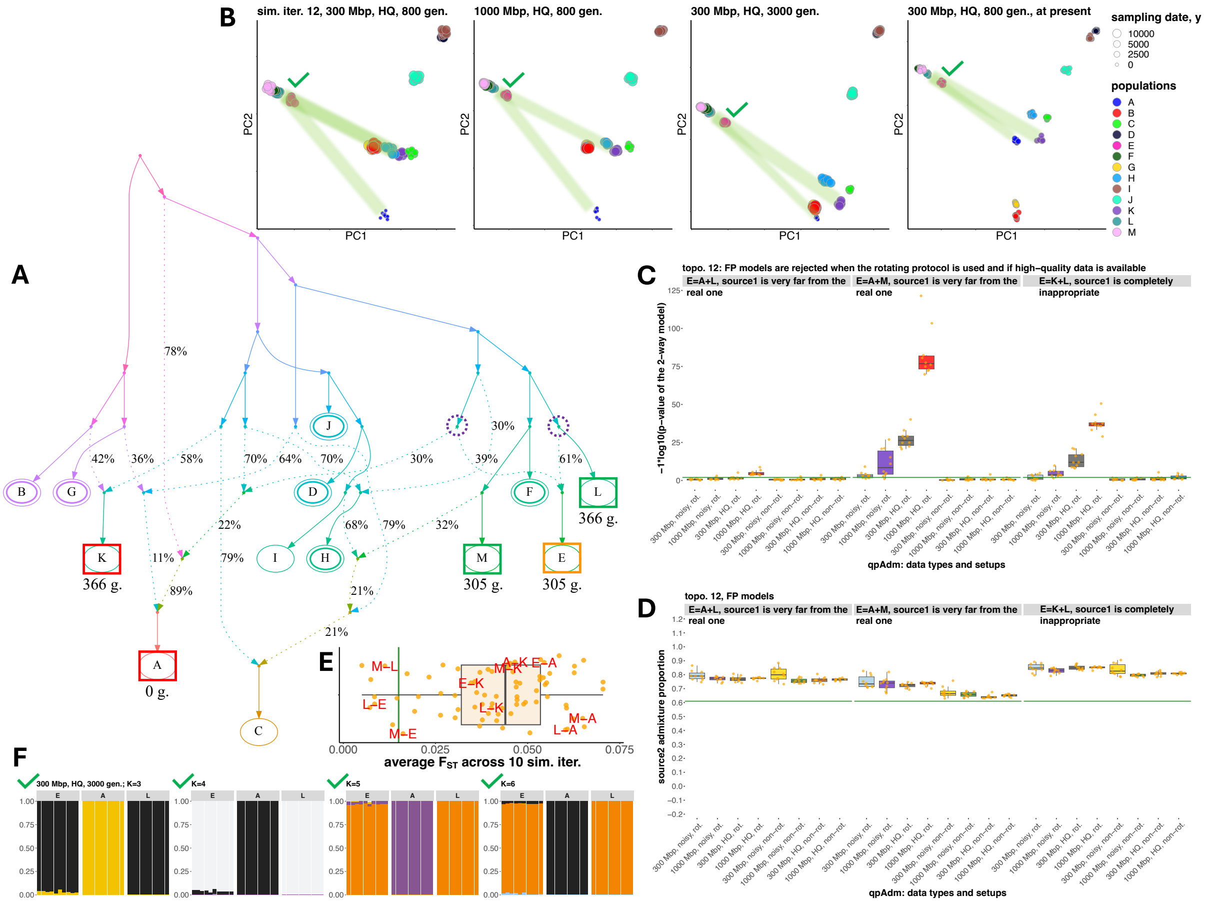
A
B
sim. iter. 12, 300 Mbp, HQ, 800 gen.
1000 Mbp, HQ, 800 gen.
300 Mbp, HQ, 3000 gen.
300 Mbp, HQ, 800 gen., at present
PC1
PC2
sampling date, y
10000
5000
2500
0
populations
C
topo. 12: FP models are rejected when the rotating protocol is used and if high-quality data is available
E=A+L, source1 is very far from the real one
E=A+M, source1 is very far from the real one
E=K+L, source1 is completely inappropriate
−1*log10(p-value of the 2-way model)
qpAdm: data types and setups
D
topo. 12, FP models
source2 admixture proportion
E
average F_ST across 10 sim. iter.
F
300 Mbp, HQ, 3000 gen.; K=3
K=4
K=5
K=6
366 g.
305 g.
0 g.

**e**

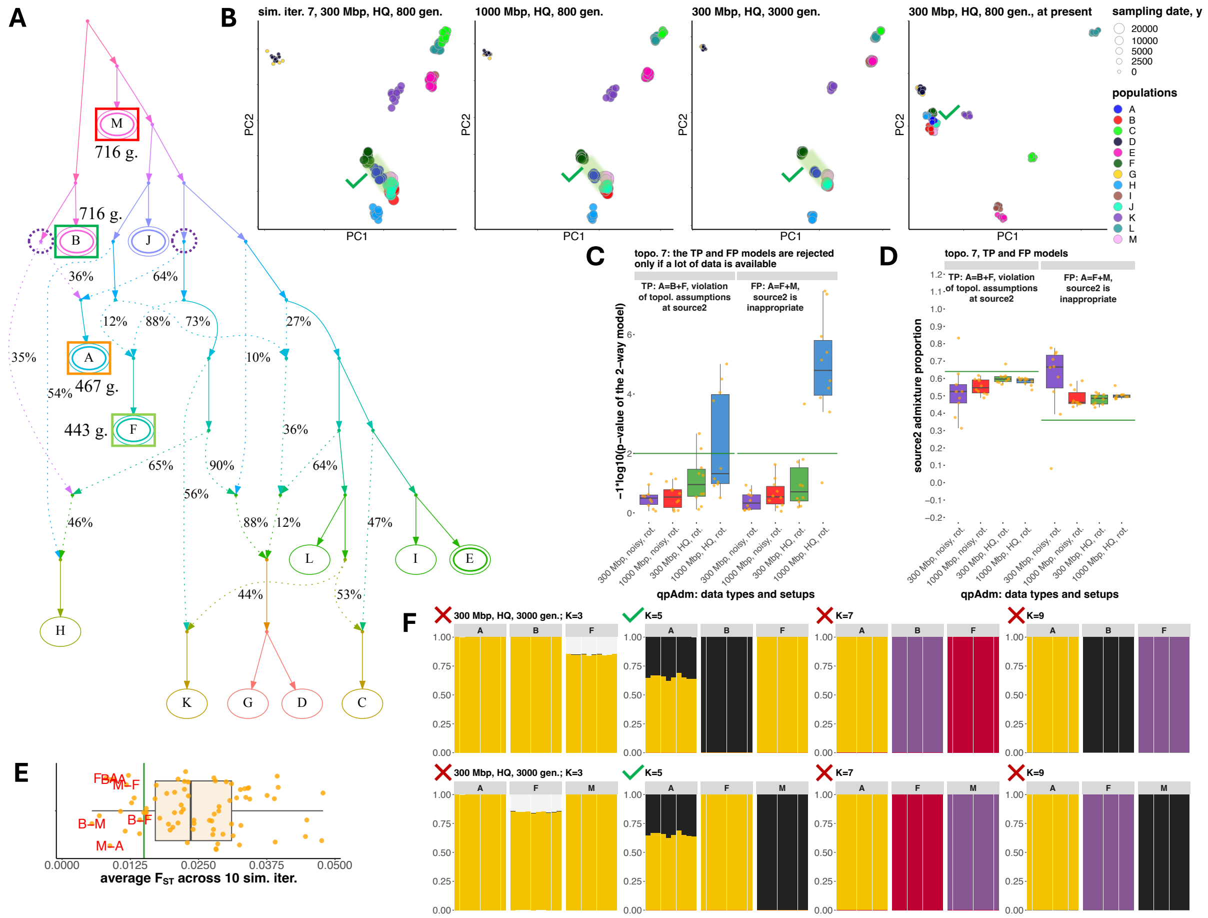

A
M
716 g.
716 g.
B
J
467 g.
A
443 g.
F
36%
64%
12%
88%
73%
27%
35%
10%
54%
36%
65%
90%
64%
56%
46%
88%
12%
47%
44%
53%
H
L
I
E
K
G
D
C
B
sim. iter. 7, 300 Mbp, HQ, 800 gen.
1000 Mbp, HQ, 800 gen.
300 Mbp, HQ, 3000 gen.
300 Mbp, HQ, 800 gen., at present
PC1
PC2
sampling date, y
20000
10000
5000
2500
0
populations
C
topo. 7: the TP and FP models are rejected only if a lot of data is available
TP: A=B+F, violation of topol. assumptions at source2
FP: A=F+M, source2 is inappropriate
−1*log10(p−value of the 2−way model)
300 Mbp, noisy, rot.
1000 Mbp, noisy, rot.
300 Mbp, HQ, rot.
1000 Mbp, HQ, rot.
qpAdm: data types and setups
D
topo. 7, TP and FP models
source2 admixture proportion
E
average $F_{ST}$ across 10 sim. iter.
F
300 Mbp, HQ, 3000 gen.; K=3
K=5
K=7
K=9

892 **f**

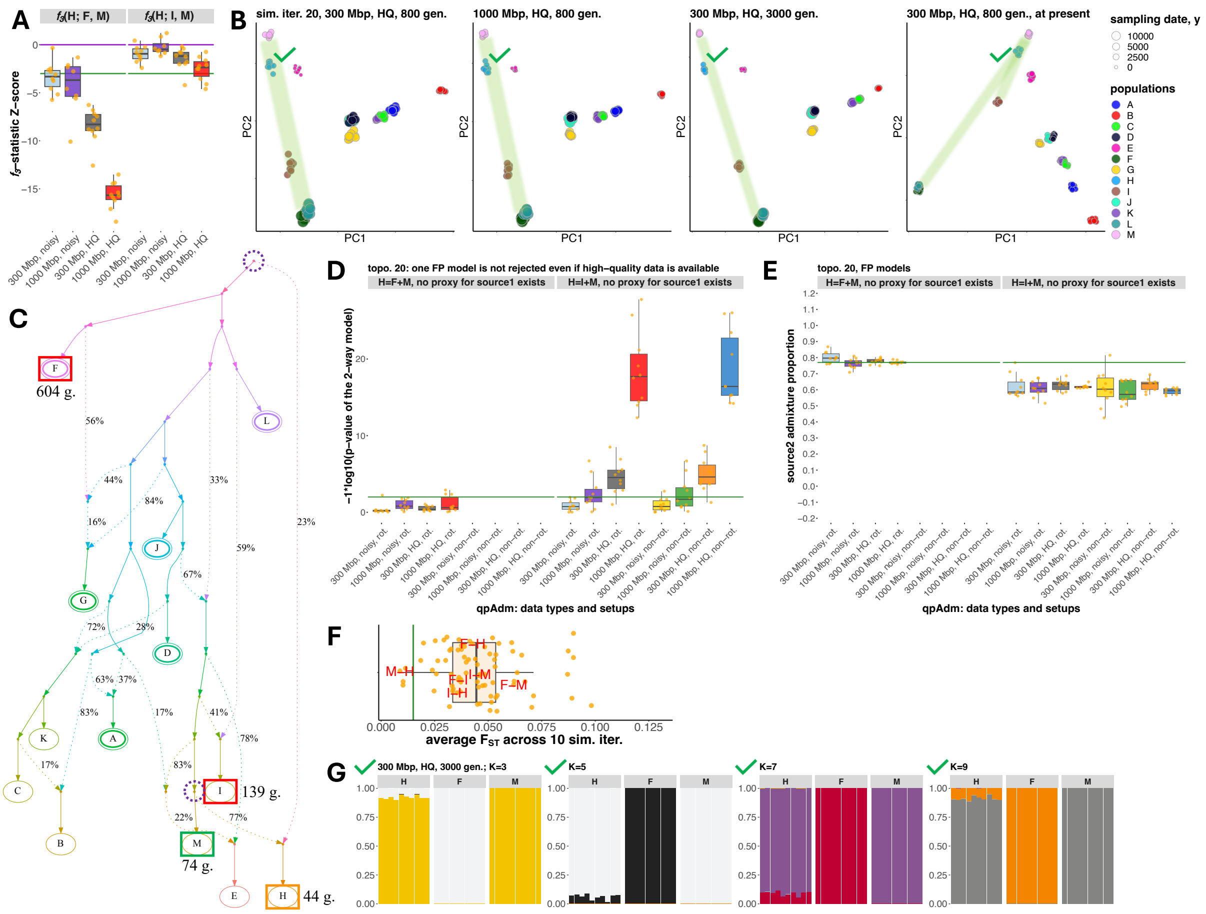

893
894

895 g

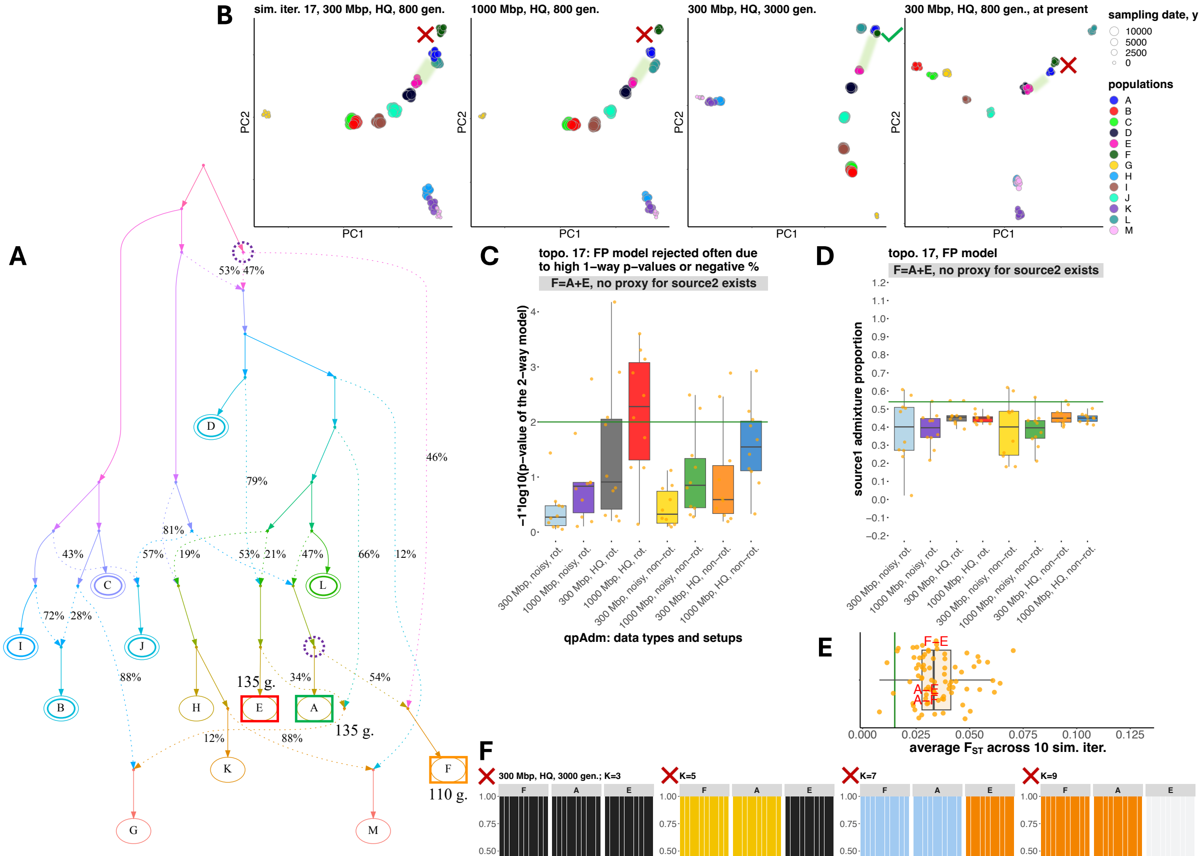

A
B
sim. iter. 17, 300 Mbp, HQ, 800 gen.
1000 Mbp, HQ, 800 gen.
300 Mbp, HQ, 3000 gen.
300 Mbp, HQ, 800 gen., at present
sampling date, y
populations
C
topo. 17: FP model rejected often due to high 1-way p-values or negative %
F=A+E, no proxy for source2 exists
qpAdm: data types and setups
D
topo. 17, FP model
F=A+E, no proxy for source2 exists
source1 admixture proportion
E
average $F_{ST}$ across 10 sim. iter.
F
300 Mbp, HQ, 3000 gen.; K=3
K=5
K=7
K=9


896
897

h

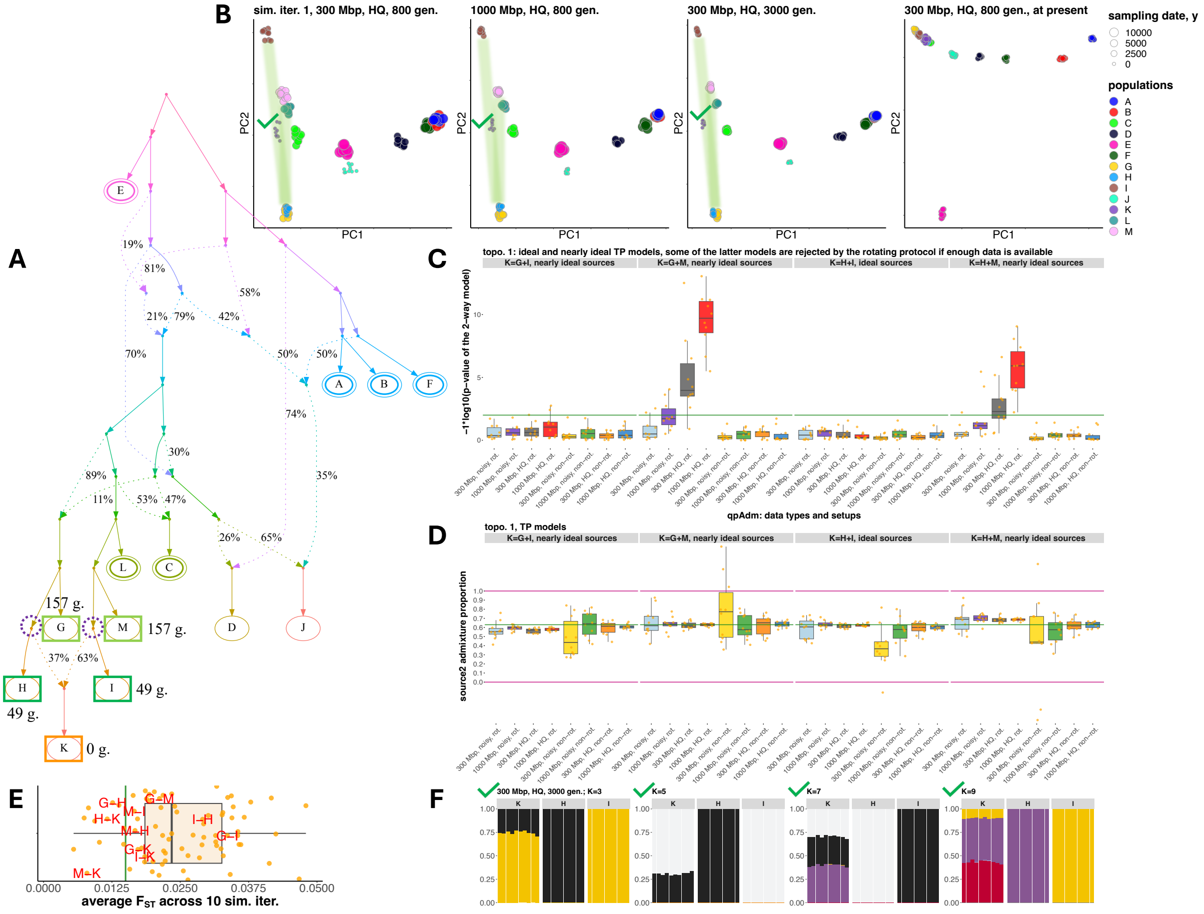

A
B
sim. iter. 1, 300 Mbp, HQ, 800 gen.
1000 Mbp, HQ, 800 gen.
300 Mbp, HQ, 3000 gen.
300 Mbp, HQ, 800 gen., at present
PC1
PC2
sampling date, y
10000
5000
2500
0
populations
A
B
C
D
E
F
G
H
I
J
K
L
M
C
topo. 1: ideal and nearly ideal TP models, some of the latter models are rejected by the rotating protocol if enough data is available
K=G+I, nearly ideal sources
K=G+M, nearly ideal sources
K=H+I, ideal sources
K=H+M, nearly ideal sources
−1*log10(p−value of the 2−way model)
qpAdm: data types and setups
D
topo. 1, TP models
source2 admixture proportion
E
average $F_{ST}$ across 10 sim. iter.
F
300 Mbp, HQ, 3000 gen.; K=3
K=5
K=7
K=9
157 g.
49 g.
0 g.

901 **i**

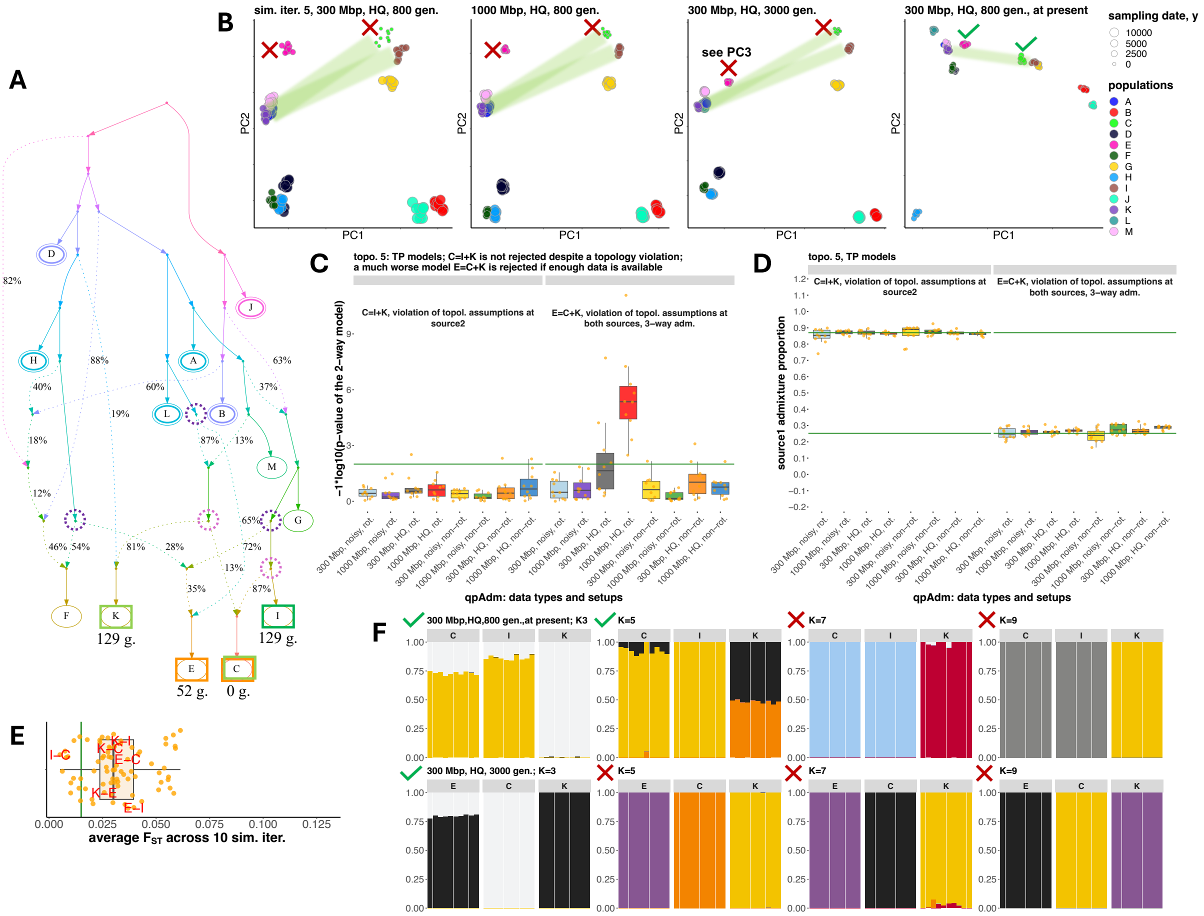

A
B
sim. iter. 5, 300 Mbp, HQ, 800 gen.
1000 Mbp, HQ, 800 gen.
300 Mbp, HQ, 3000 gen.
see PC3
300 Mbp, HQ, 800 gen., at present
sampling date, y
populations
C
topo. 5: TP models; C=I+K is not rejected despite a topology violation;
a much worse model E=C+K is rejected if enough data is available
C=I+K, violation of topol. assumptions at source2
E=C+K, violation of topol. assumptions at both sources, 3-way adm.
−1*log10(p−value of the 2−way model)
qpAdm: data types and setups
D
topo. 5, TP models
source1 admixture proportion
E
average F_ST across 10 sim. iter.
F
300 Mbp,HQ,800 gen.,at present; K3
300 Mbp, HQ, 3000 gen.; K=3


902
903

j

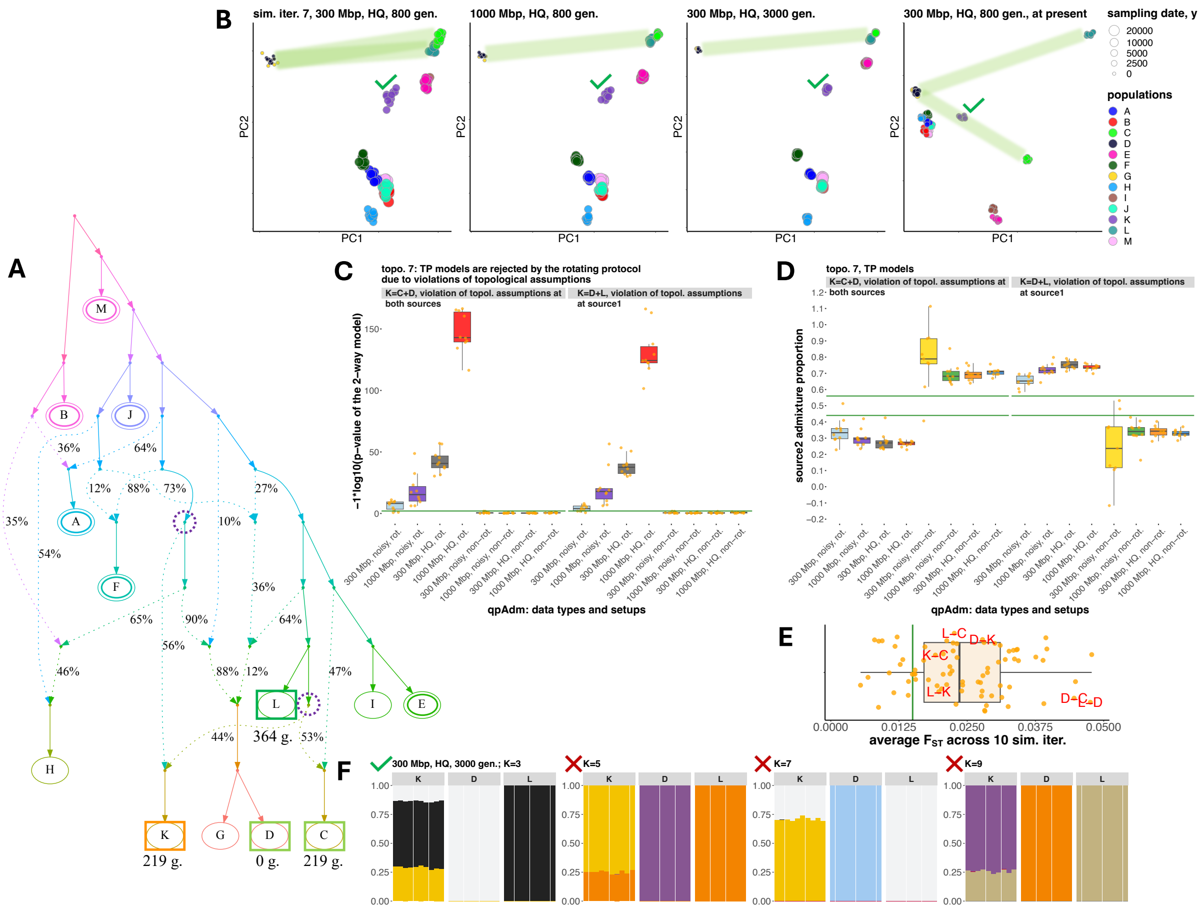
A
B
sim. iter. 7, 300 Mbp, HQ, 800 gen.
1000 Mbp, HQ, 800 gen.
300 Mbp, HQ, 3000 gen.
300 Mbp, HQ, 800 gen., at present
PC1
PC2
sampling date, y
20000
10000
5000
2500
0
populations
A
B
C
D
E
F
G
H
I
J
K
L
M
C
topo. 7: TP models are rejected by the rotating protocol due to violations of topological assumptions
K=C+D, violation of topol. assumptions at both sources
K=D+L, violation of topol. assumptions at source1
−1*log10(p−value of the 2−way model)
qpAdm: data types and setups
D
topo. 7, TP models
source2 admixture proportion
E
average F_ST across 10 sim. iter.
F
300 Mbp, HQ, 3000 gen.; K=3
K=5
K=7
K=9
36%
64%
12%
88%
73%
27%
35%
54%
10%
65%
90%
36%
64%
56%
46%
88%
12%
47%
44%
53%
364 g.
219 g.
0 g.
219 g.

**k**

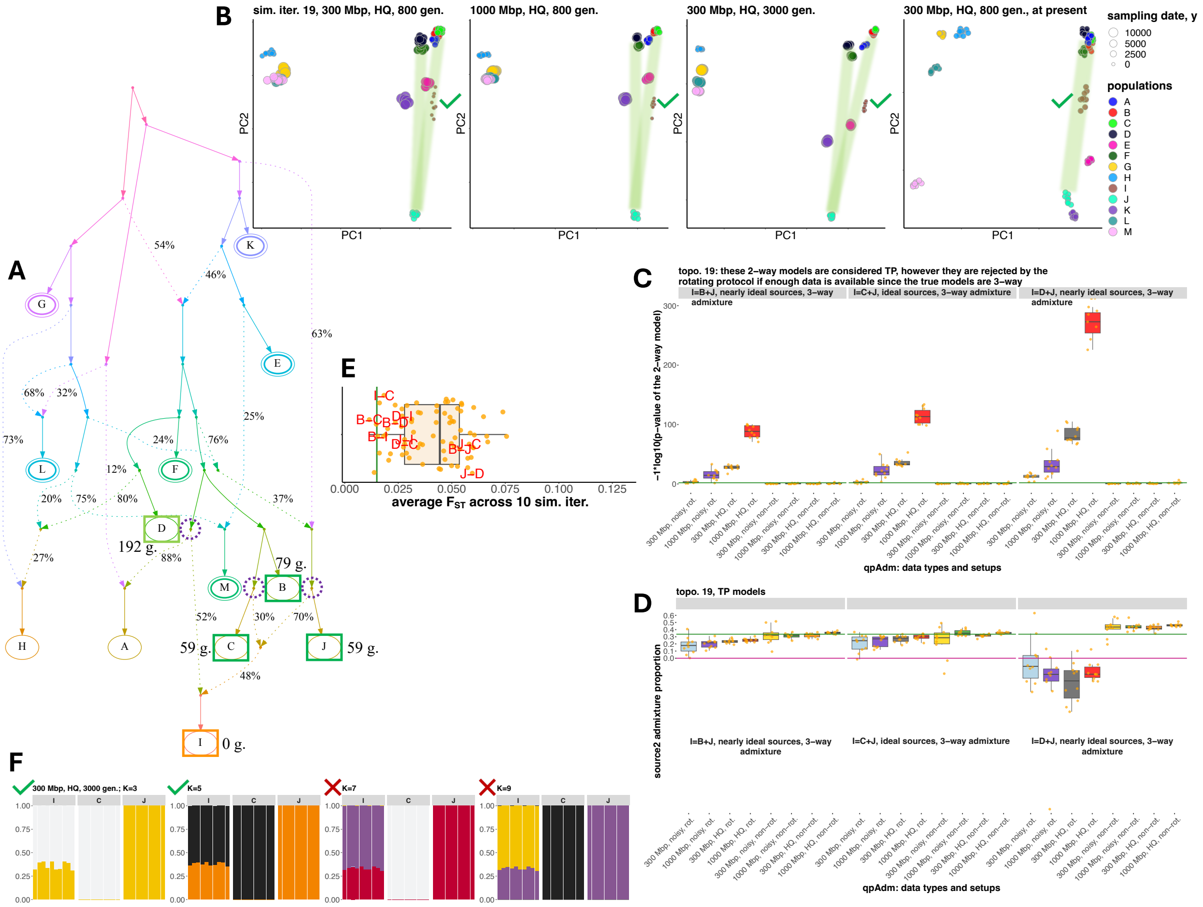
A
B
sim. iter. 19, 300 Mbp, HQ, 800 gen.
1000 Mbp, HQ, 800 gen.
300 Mbp, HQ, 3000 gen.
300 Mbp, HQ, 800 gen., at present
PC1
PC2
sampling date, y
10000
5000
2500
0
populations
C
topo. 19: these 2-way models are considered TP, however they are rejected by the rotating protocol if enough data is available since the true models are 3-way
I=B+J, nearly ideal sources, 3-way admixture
I=C+J, ideal sources, 3-way admixture
I=D+J, nearly ideal sources, 3-way admixture
−1*log10(p−value of the 2−way model)
qpAdm: data types and setups
D
topo. 19, TP models
source2 admixture proportion
E
average $F_{ST}$ across 10 sim. iter.
F
300 Mbp, HQ, 3000 gen.; K=3
K=5
K=7
K=9
192 g.
79 g.
59 g.
59 g.
0 g.

I

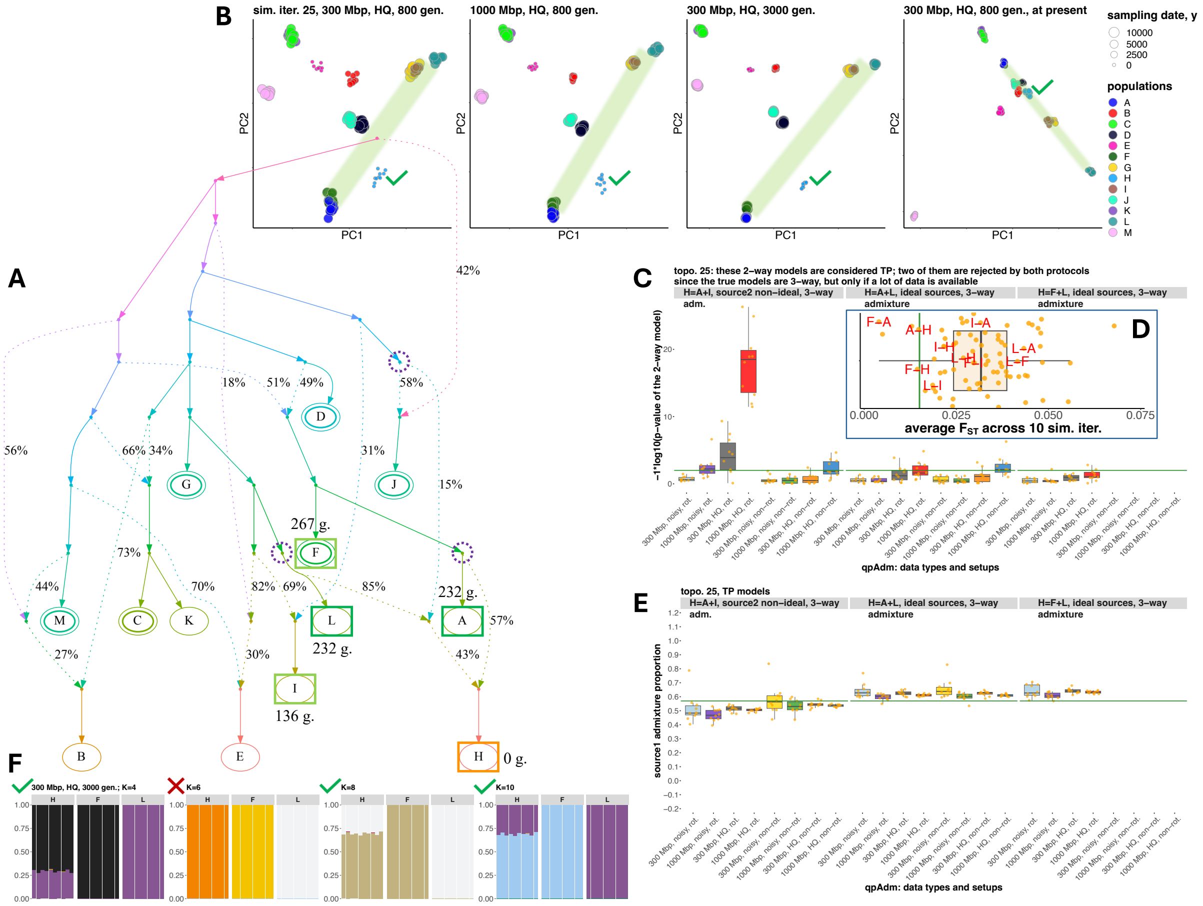

A
B
sim. iter. 25, 300 Mbp, HQ, 800 gen.
1000 Mbp, HQ, 800 gen.
300 Mbp, HQ, 3000 gen.
300 Mbp, HQ, 800 gen., at present
PC1
PC2
sampling date, y
10000
5000
2500
0
populations
A
B
C
D
E
F
G
H
I
J
K
L
M
42%
18%
51%
49%
58%
56%
66%
34%
31%
15%
267 g.
73%
44%
70%
82%
69%
85%
232 g.
57%
27%
30%
43%
136 g.
0 g.
C
topo. 25: these 2-way models are considered TP; two of them are rejected by both protocols
since the true models are 3-way, but only if a lot of data is available
H=A+I, source2 non-ideal, 3-way adm.
H=A+L, ideal sources, 3-way admixture
H=F+L, ideal sources, 3-way admixture
−1*log10(p-value of the 2-way model)
D
average F_ST across 10 sim. iter.
qpAdm: data types and setups
E
topo. 25, TP models
source1 admixture proportion
F
300 Mbp, HQ, 3000 gen.; K=4
K=6
K=8
K=10

**m**

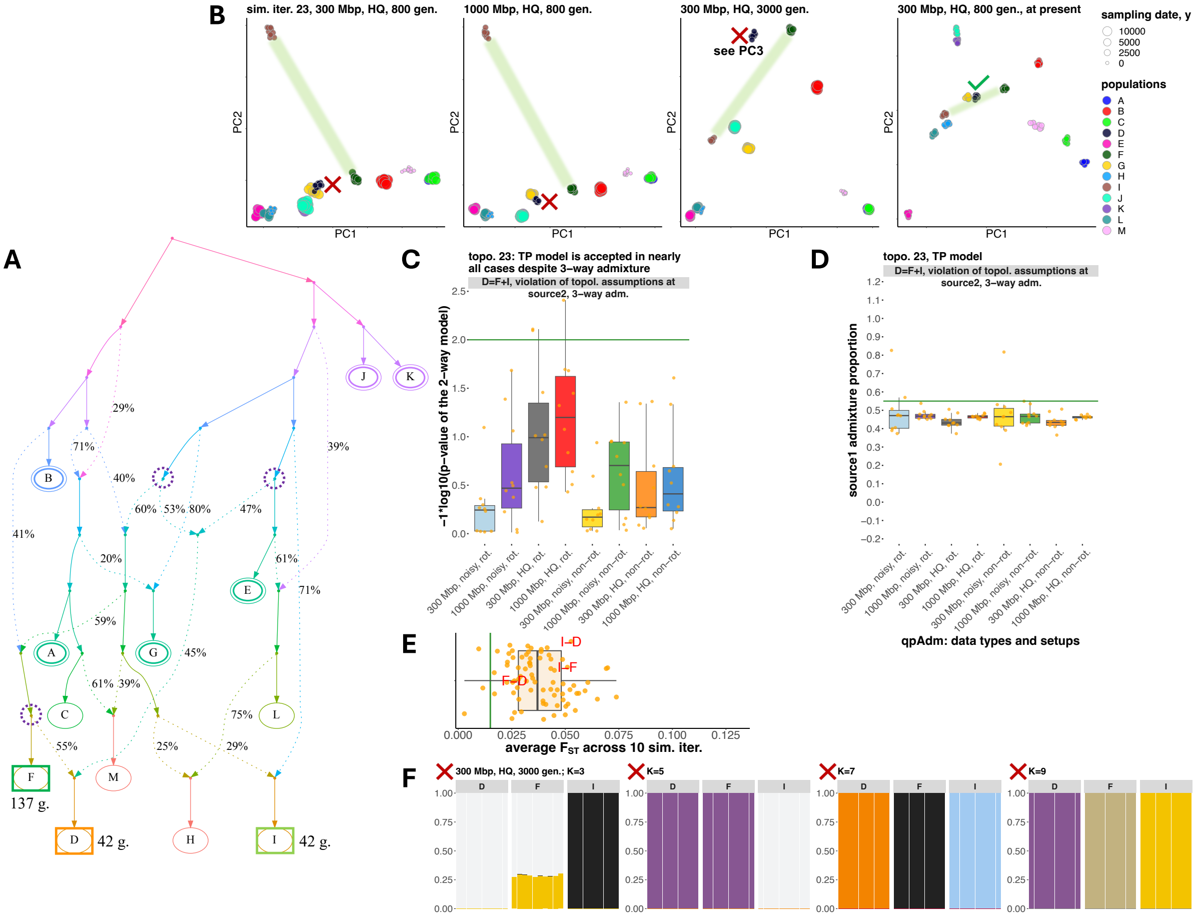

A
B
sim. iter. 23, 300 Mbp, HQ, 800 gen.
1000 Mbp, HQ, 800 gen.
300 Mbp, HQ, 3000 gen.
see PC3
300 Mbp, HQ, 800 gen., at present
PC1
PC2
sampling date, y
10000
5000
2500
0
populations
A
B
C
D
E
F
G
H
I
J
K
L
M
C
topo. 23: TP model is accepted in nearly all cases despite 3–way admixture
D=F+I, violation of topol. assumptions at source2, 3–way adm.
−1*log10(p−value of the 2−way model)
300 Mbp, noisy, rot.
1000 Mbp, noisy, rot.
300 Mbp, HQ, rot.
1000 Mbp, HQ, rot.
300 Mbp, noisy, non−rot.
1000 Mbp, noisy, non−rot.
300 Mbp, HQ, non−rot.
1000 Mbp, HQ, non−rot.
D
topo. 23, TP model
D=F+I, violation of topol. assumptions at source2, 3–way adm.
source1 admixture proportion
qpAdm: data types and setups
E
I–D
I–F
F–D
average F_ST across 10 sim. iter.
F
300 Mbp, HQ, 3000 gen.; K=3
K=5
K=7
K=9
137 g.
42 g.
42 g.

Case studies illustrating false (**a-g**) and true (**b, c, e, h-m**) positive *qpAdm* models of various topological types, with corresponding simulated histories, PCA, $F_{ST}$, and unsupervised *ADMIXTURE* results also shown. Each panel of this figure illustrates one case study and is composed of several standardized sub-panels described below. **A**) Simulated history in the form of an admixture graph (only topology is shown here, for divergence/admixture dates and effective population sizes see the respective simulated history presented in **Fig. S3**). Sampled populations are marked by letters. The target population from the *qpAdm* models illustrated here is enclosed in an orange rectangle; correct proxy sources – in green rectangles, correct proxy sources with violations of the topological assumptions of *qpAdm* – in green-yellow rectangles, and inappropriate proxy sources, if any – in red rectangles. Sampling dates for these groups (in generations before present) are shown beside the rectangles (the dates are from the "shallow" simulations up to 800 generations deep). True simulated ancestry source(s) for the target are enclosed in dashed circles. Six populations used as outgroups for the non-rotating *qpAdm* protocol are enclosed in double ovals. **B**) Two-dimensional PCA plots for four simulated high-quality datasets indicated in plot titles. Simulated populations are colored according to the legend on the right, and larger points correspond to older sampling dates. The space between the proxy sources is shaded in green. If admixture model(s) are supported by three-dimensional (sic!) PCA (PC1 vs. PC2 vs. PC3) according to the criteria listed in Methods, a green tick mark is placed beside the target group in the plot, and a red cross mark is placed otherwise. **C**) Boxplots summarizing *p*-values of two-way *qpAdm* models (indicated in plot titles) across simulation or subsampling replicates, grouped by simulation setups and *qpAdm* protocols as indicated on the x-axis. Green lines show the *p*-value threshold of 0.01 used in this study for model rejection. *P*-values for one-way models are not shown for brevity. Some *qpAdm* models were not tested using the non-rotating protocol since at least one "left" population belongs to the fixed set of six "right" populations. **D**) Boxplots summarizing EAF across simulation or subsampling replicates, grouped by simulation setups and *qpAdm* protocols. The admixture proportions are shown either for the first or second proxy source, as indicated on the y-axis; green lines show the simulated admixture proportion. **E**) All $F_{ST}$ values for population pairs from the simulated history. Average $F_{ST}$ values are shown across 10 simulation replicates for 1000-Mbp-sized genomes and high-quality data. Population pairs formed by components of the illustrated two-way *qpAdm* model(s) are labelled in red. The green line shows median $F_{ST}$ (0.015) for the Bronze Age West Asian populations analyzed by Lazaridis *et al.* (2016) as an example of human genetic diversity at the sub-continental scale. **F**) Ancestry proportions estimated with unsupervised *ADMIXTURE* for the groups constituting the *qpAdm* model(s). For brevity, results are shown for one or two *qpAdm* models, for one simulated dataset indicated in the plot title (usually simulation setup no. 3: 300-Mbp-sized genomes, max. divergence date of 3,000 generations, high-quality data), and for four selected *K* values. If admixture model(s) are supported by this analysis at a given *K* value, a green tick mark is placed beside the target group in the plot, and a red cross mark is placed otherwise. In panel **f**, $f_3$-statistics $f_3$(target; proxy source$_1$, proxy source$_2$) are shown in sub-panel **A**, and the other information is shown in sub-panels **B** to **G**.

**Figure S7.**

**a**

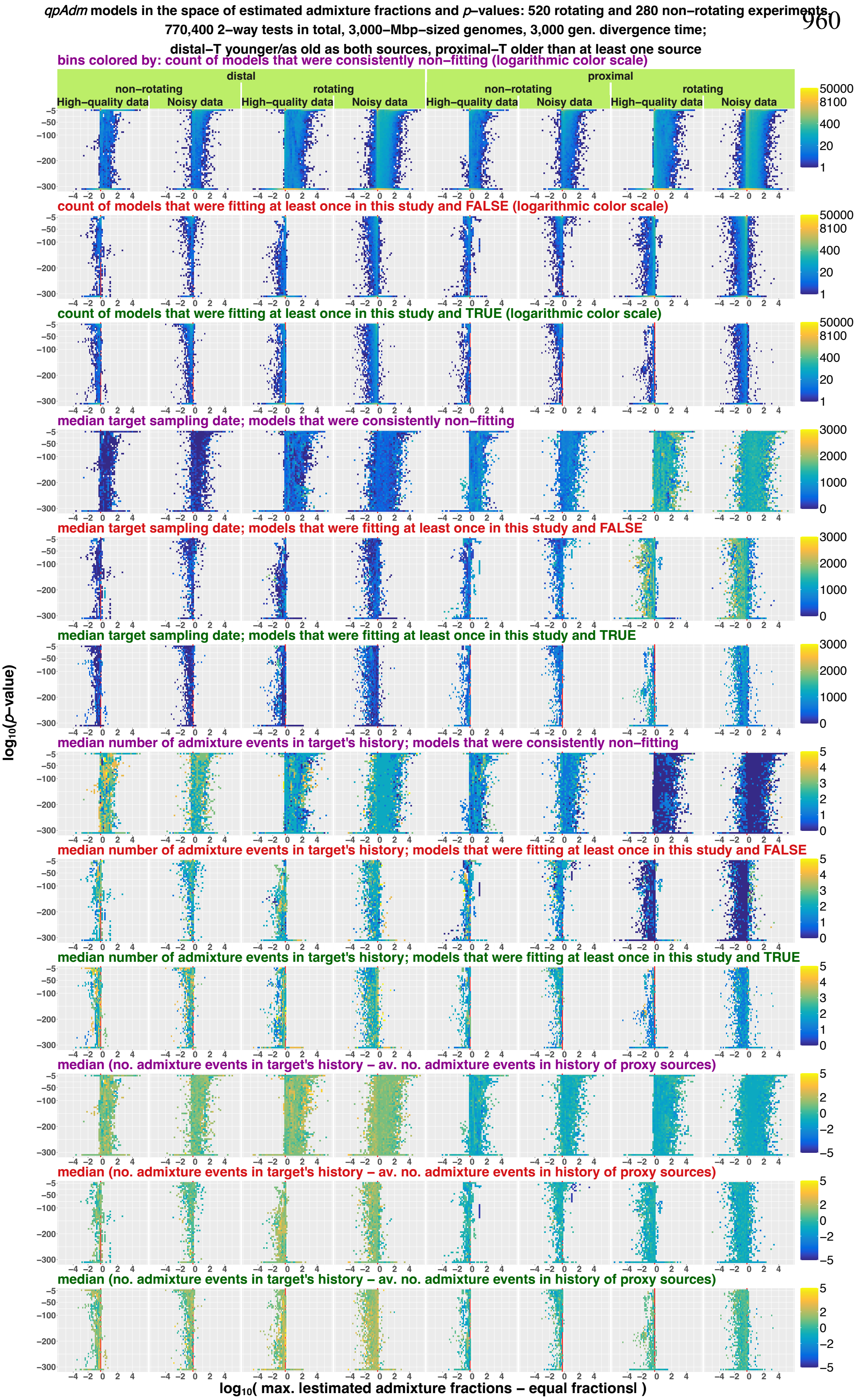

*qpAdm* models in the space of estimated admixture fractions and *p*–values: 520 rotating and 280 non–rotating experiments,
770,400 2–way tests in total, 3,000–Mbp–sized genomes, 3,000 gen. divergence time;
distal–T younger/as old as both sources, proximal–T older than at least one source
bins colored by: count of models that were consistently non–fitting (logarithmic color scale)
distal
proximal
non–rotating
rotating
High–quality data
Noisy data
count of models that were fitting at least once in this study and FALSE (logarithmic color scale)
count of models that were fitting at least once in this study and TRUE (logarithmic color scale)
median target sampling date; models that were consistently non–fitting
median target sampling date; models that were fitting at least once in this study and FALSE
median target sampling date; models that were fitting at least once in this study and TRUE
median number of admixture events in target's history; models that were consistently non–fitting
median number of admixture events in target's history; models that were fitting at least once in this study and FALSE
median number of admixture events in target's history; models that were fitting at least once in this study and TRUE
median (no. admixture events in target's history – av. no. admixture events in history of proxy sources)
median (no. admixture events in target's history – av. no. admixture events in history of proxy sources)
median (no. admixture events in target's history – av. no. admixture events in history of proxy sources)
log10(p–value)
log10( max. |estimated admixture fractions – equal fractions| )

961 **b**

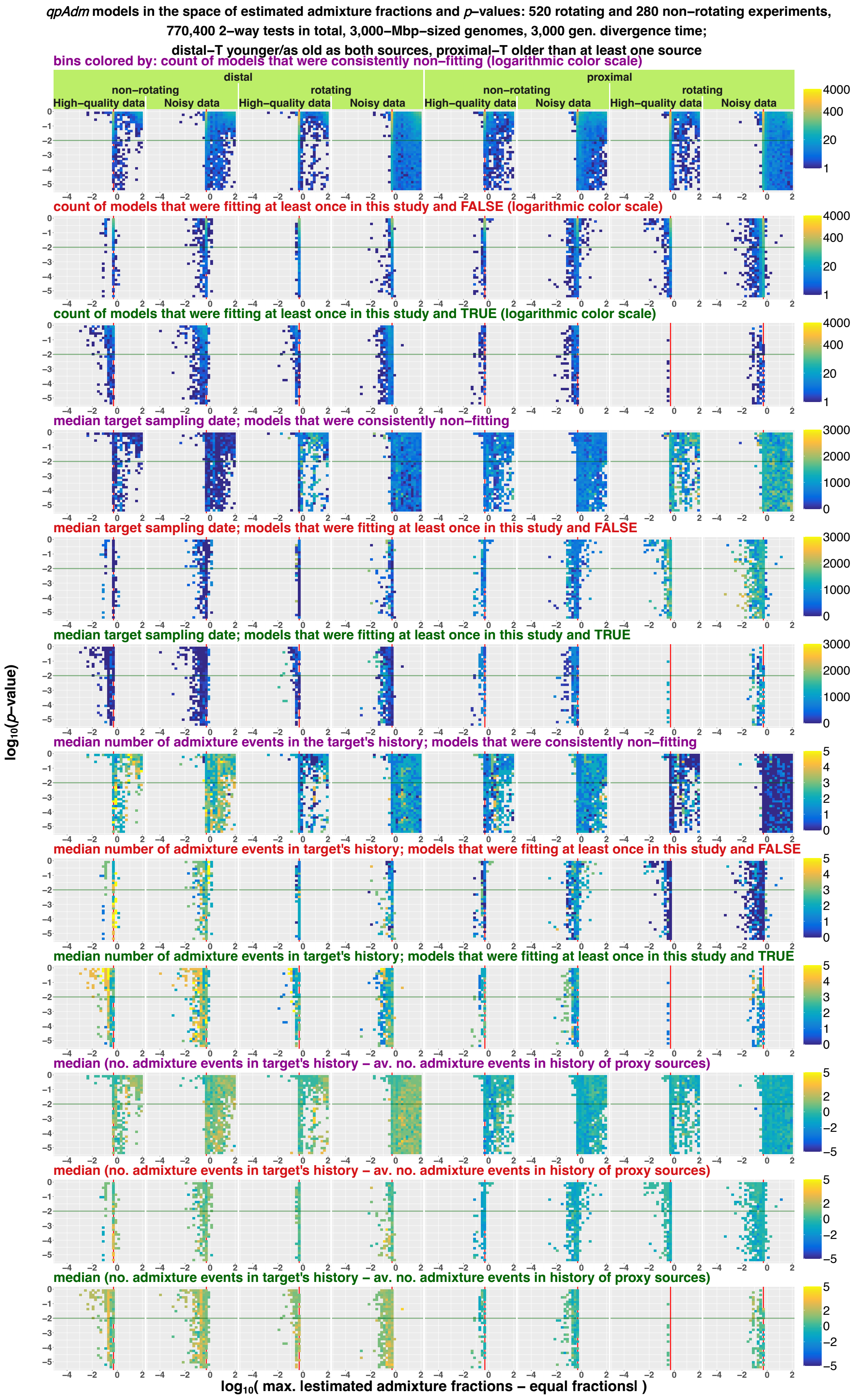

qpAdm models in the space of estimated admixture fractions and p–values: 520 rotating and 280 non–rotating experiments, 770,400 2–way tests in total, 3,000–Mbp–sized genomes, 3,000 gen. divergence time; distal–T younger/as old as both sources, proximal–T older than at least one source
bins colored by: count of models that were consistently non–fitting (logarithmic color scale)
distal
proximal
non–rotating
rotating
High–quality data
Noisy data
count of models that were fitting at least once in this study and FALSE (logarithmic color scale)
count of models that were fitting at least once in this study and TRUE (logarithmic color scale)
median target sampling date; models that were consistently non–fitting
median target sampling date; models that were fitting at least once in this study and FALSE
median target sampling date; models that were fitting at least once in this study and TRUE
median number of admixture events in the target's history; models that were consistently non–fitting
median number of admixture events in target's history; models that were fitting at least once in this study and FALSE
median number of admixture events in target's history; models that were fitting at least once in this study and TRUE
median (no. admixture events in target's history – av. no. admixture events in history of proxy sources)
median (no. admixture events in target's history – av. no. admixture events in history of proxy sources)
median (no. admixture events in target's history – av. no. admixture events in history of proxy sources)
log10(p–value)
log10( max. lestimated admixture fractions – equal fractionsl )


962

**c**

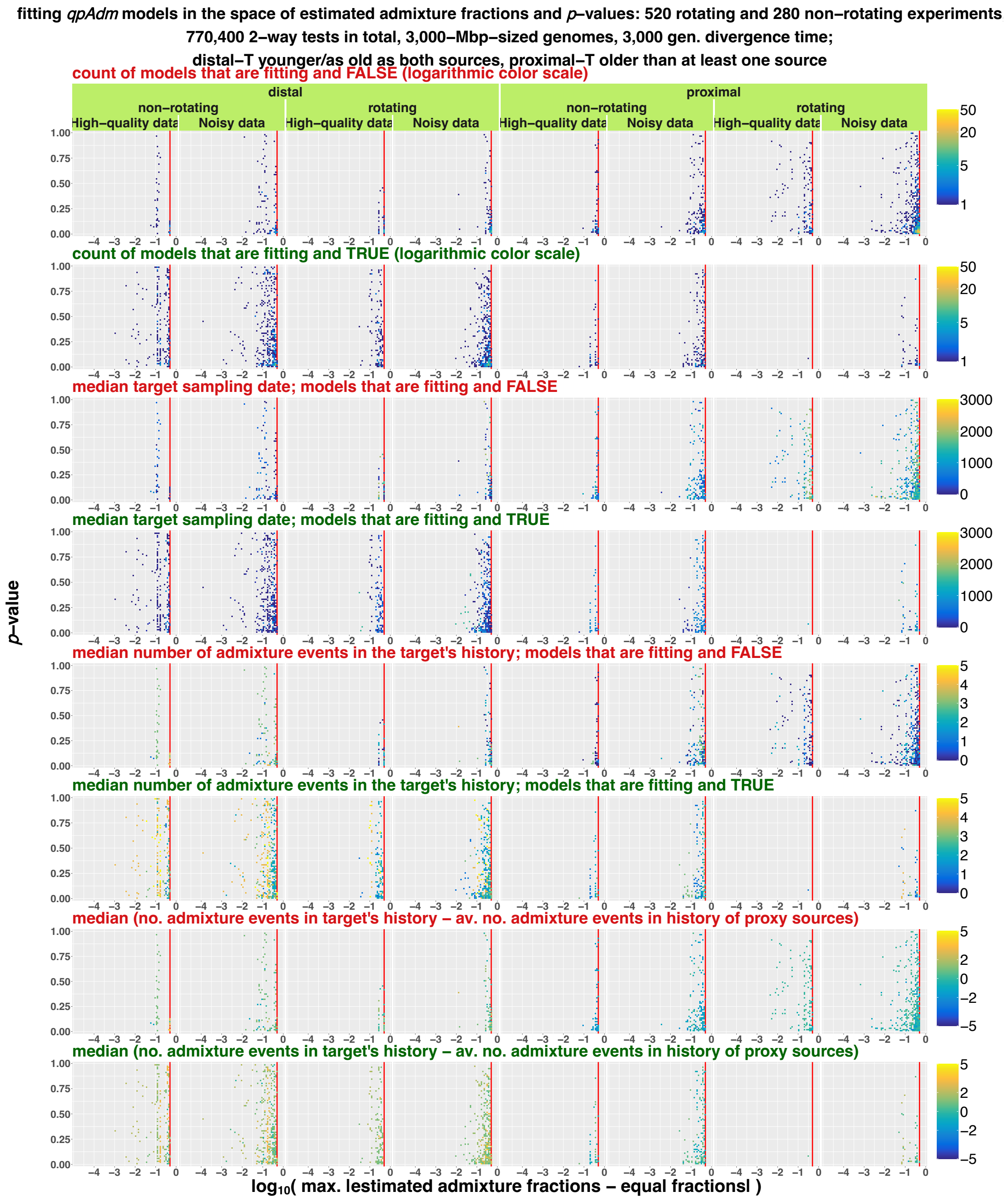


Two-way *qpAdm* models tested on AGS simulations visualized in the space of max|*EAF–EF*| and *p*-values (both axes are logarithmic). Results are shown for the 3,000-Mbp-sized genomes only and stratified in columns by: model subsets (distal or proximal), *qpAdm* protocols, and data quality. Results are also stratified by three model classes: consistently non-fitting (plot rows labelled in purple), fitting at least once in this study and evaluated as false (in red), and fitting at least once in this study and evaluated as true (in green). The whole space is shown in panel **a** (*p*-values from $10^{-306}$ to 1); *p*-values from $3.16\times10^{-6}$ to 1 are shown in **b**, and plots based on fitting models only are shown in panel **c**. The space is divided into rectangular bins (50×50 in **a**, 35×35 in **b**, and 75×75 in **c**), and they are colored by density of individual *qpAdm* models populating this space (logarithmic color scale) or by median values of metrics informative for understanding method performance: target sampling date, number of admixture events on all paths from the target to the root of an AGS simulation (number of admixture events in a target's history), and difference between number of admixture events in a target's history and average number of admixture events in the history of both proxy sources. The red vertical lines (or the red tick marks) mark max|*EAF*–*EF*| = 0.5, and the green horizontal lines mark the *p*-value threshold used in the AGS section of this study (0.01).

982 **Figure S8.**

983 **a**

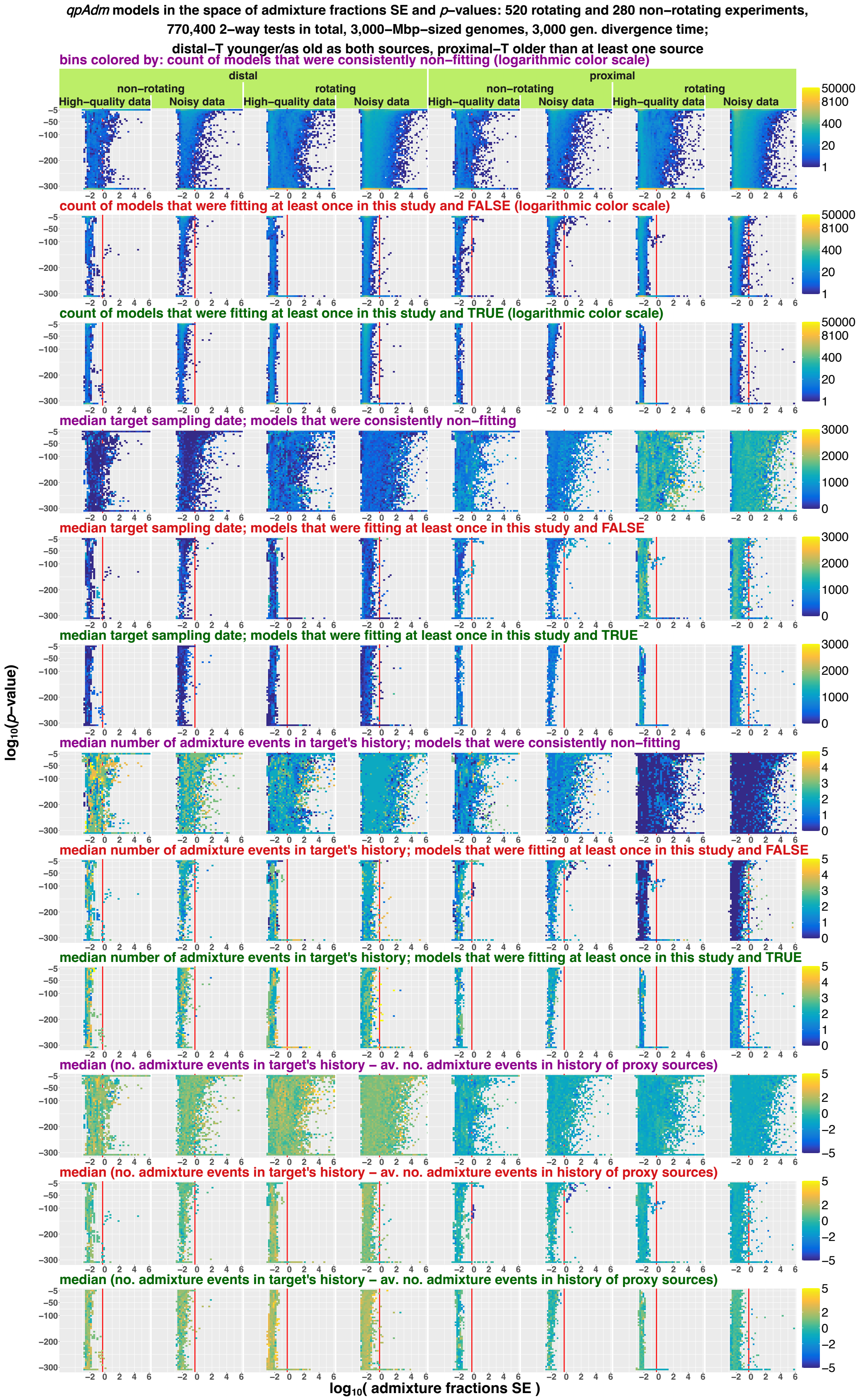


984

985 **b**

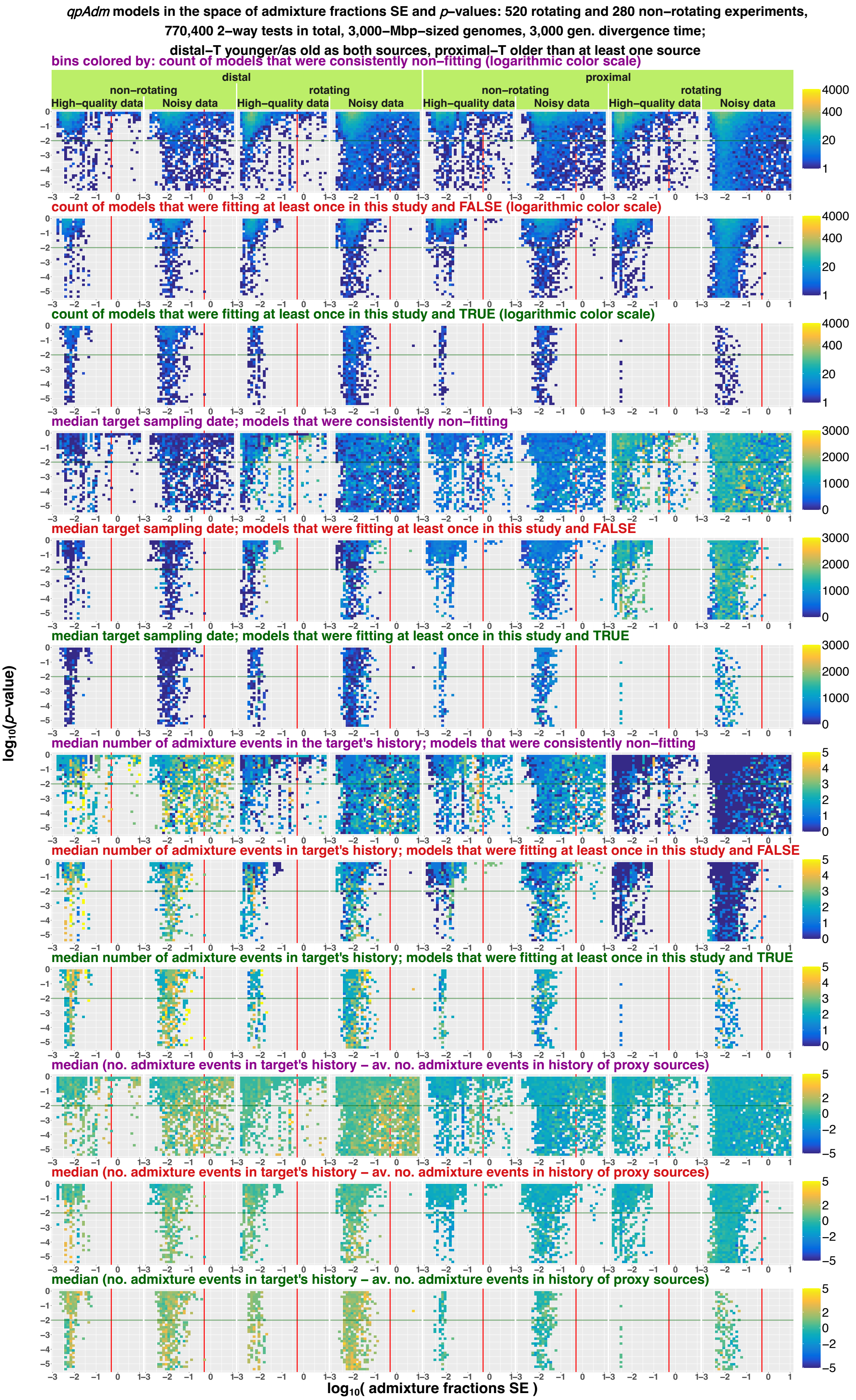

qpAdm models in the space of admixture fractions SE and p–values: 520 rotating and 280 non–rotating experiments,
770,400 2–way tests in total, 3,000–Mbp–sized genomes, 3,000 gen. divergence time;
distal–T younger/as old as both sources, proximal–T older than at least one source
bins colored by: count of models that were consistently non–fitting (logarithmic color scale)
distal
proximal
non–rotating
rotating
High–quality data
Noisy data
count of models that were fitting at least once in this study and FALSE (logarithmic color scale)
count of models that were fitting at least once in this study and TRUE (logarithmic color scale)
median target sampling date; models that were consistently non–fitting
median target sampling date; models that were fitting at least once in this study and FALSE
median target sampling date; models that were fitting at least once in this study and TRUE
median number of admixture events in the target's history; models that were consistently non–fitting
median number of admixture events in target's history; models that were fitting at least once in this study and FALSE
median number of admixture events in target's history; models that were fitting at least once in this study and TRUE
median (no. admixture events in target's history – av. no. admixture events in history of proxy sources)
median (no. admixture events in target's history – av. no. admixture events in history of proxy sources)
median (no. admixture events in target's history – av. no. admixture events in history of proxy sources)
log10(p–value)
log10( admixture fractions SE )


986

**c**

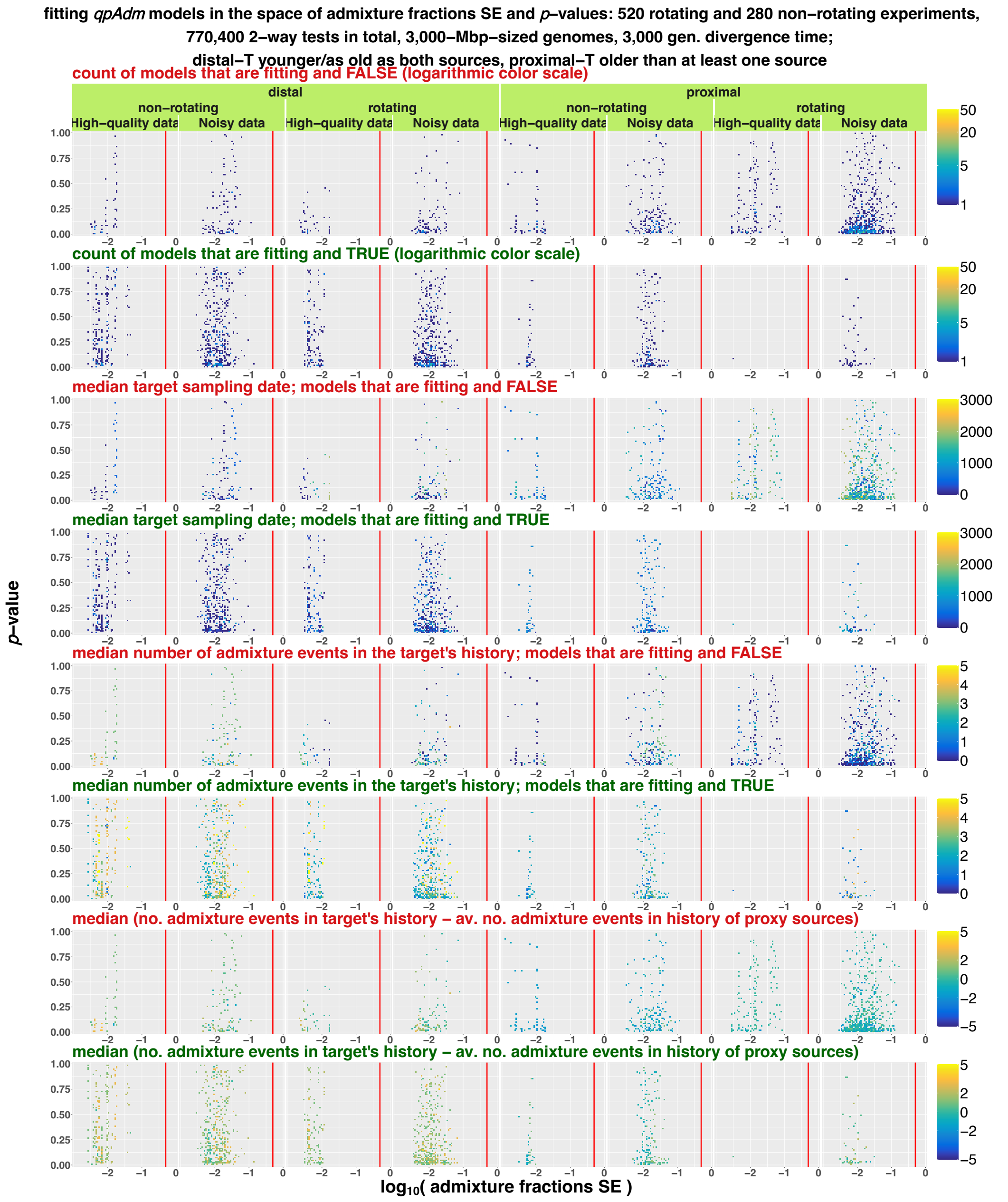


Two-way *qpAdm* models tested on AGS simulations visualized in the space of admixture fractions SE and *p*-values (both axes are logarithmic). Results are shown for the 3,000-Mbp-sized genomes only and stratified in columns by: model subsets (distal or proximal), *qpAdm* protocols, and data quality. Results are also stratified by three model classes: consistently non-fitting (plot rows labelled in purple), fitting at least once in this study and evaluated as false (in red), and fitting at least once in this study and evaluated as true (in green). The whole space is shown in panel **a** (*p*-values from $10^{-306}$ to 1); *p*-values from $3.16\times10^{-6}$ to 1 are shown in **b**, and plots based on fitting models only are shown in panel **c**. The space is divided into rectangular bins (50×50 in **a**, 35×35 in **b**, and 75×75 in **c**), and they are colored by density of individual *qpAdm* models populating this space (logarithmic color scale) or by median values of metrics informative for understanding method performance: target sampling date, number of admixture events on all paths from the target to the root of an AGS simulation (number of admixture events in a target's history), and difference between number of admixture events in a target's history and average number of admixture events in the history of both proxy sources. The red vertical lines (or the red tick marks) mark SE = 0.5, and the green horizontal lines mark the *p*-value threshold used in the AGS section of this study (0.01).

**Figure S9.**

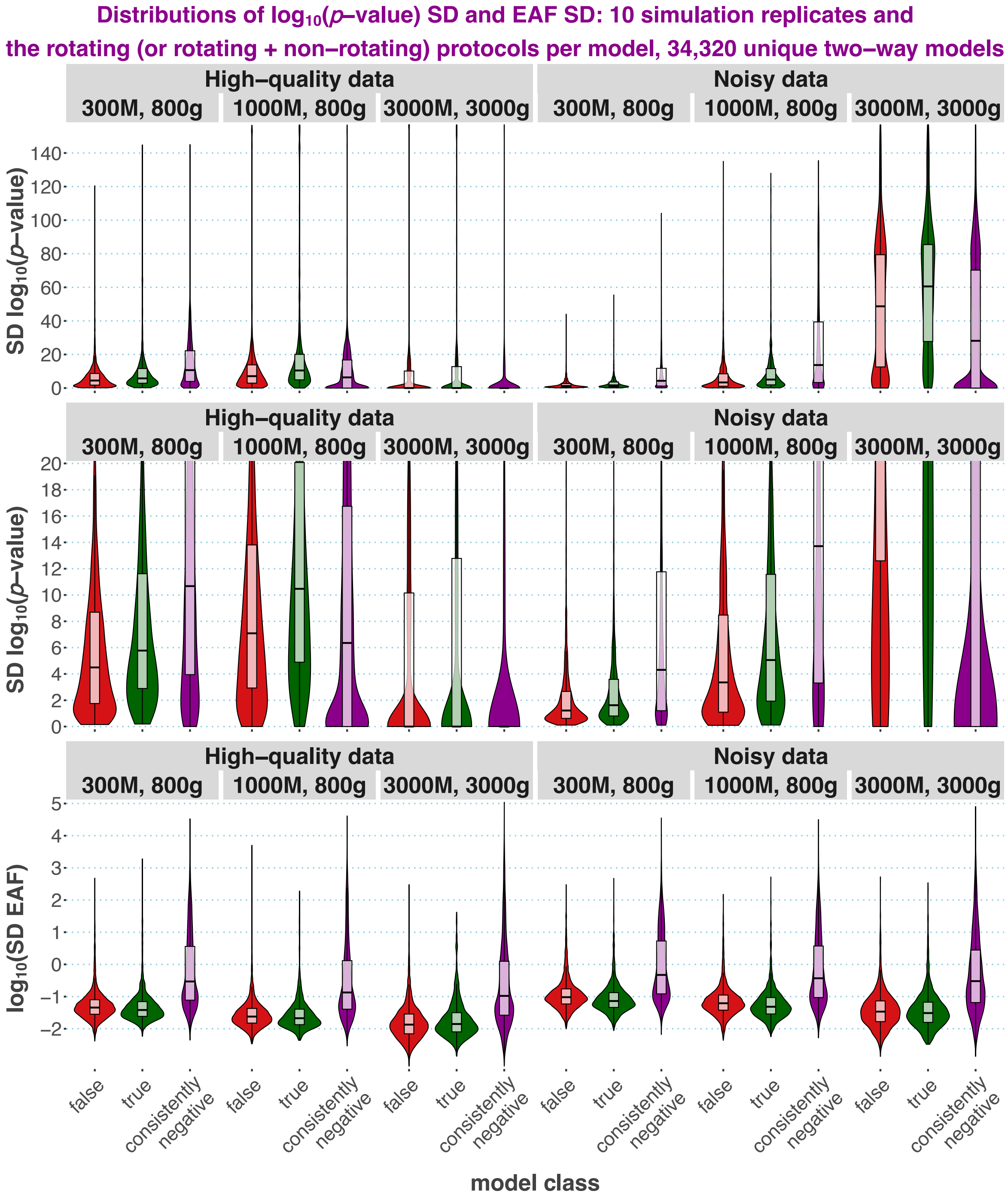


Exploring stability of $\log_{10}$(*p*-values) and EAF reported by *qpAdm*: SD or $\log_{10}$SD of these metrics were calculated for a two-way model tested on 10 simulation or data subsampling replicates. Some models were tested with both the rotating and non-rotating setups, and others with the rotating setup only. The violin plots and boxplots summarize distributions of SDs and are stratified by data quality and by three combinations of simulated genome sizes (300, 1,000, or 3,000 Mbp) and maximal simulation depths (800 or 3,000 generations). Results are also stratified by three model classes: fitting at least once in this study and evaluated as false (labelled “false” and shown in red), fitting at least once in this study and evaluated as true (labelled “true”, in green), and consistently non-fitting (labelled “consistently negative”, in purple).

**Figure S10.**

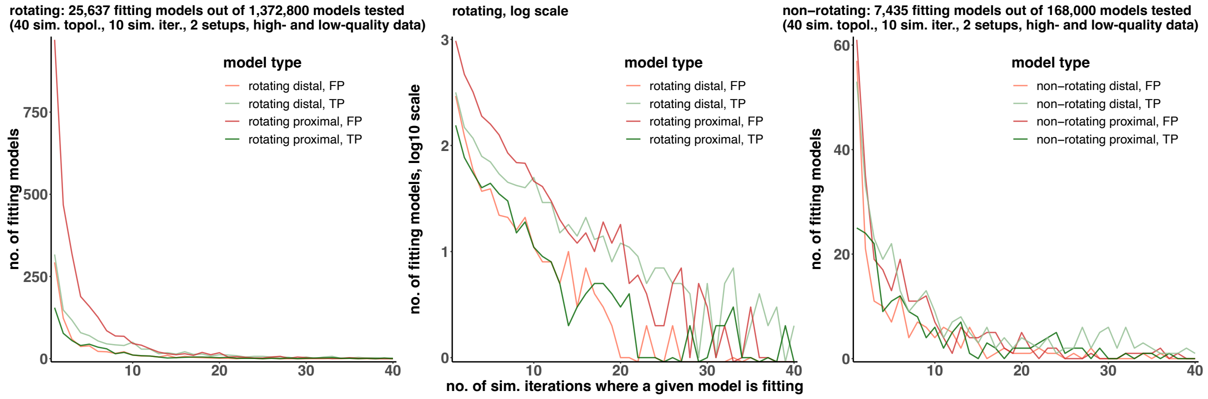


Plots illustrating that for any *qpAdm* protocol or model type (false or true), the vast majority of models fit the data only sporadically. Model feasibility was assessed across the 40 simulation or subsampling replicates (simulation setups no. 1 and 2) analyzed by the rotating and non-rotating *qpAdm* protocols. Models are binned by the number of simulation/subsampling replicates where a model is feasible (fitting) (on the x-axis). Separate decay curves are shown for TP and FP, proximal and distal two-way admixture models. In the case of the rotating protocol, two plots illustrate the same data: a linear-scale plot on the left and a log-scale plot in the center.

**Figure S11.**

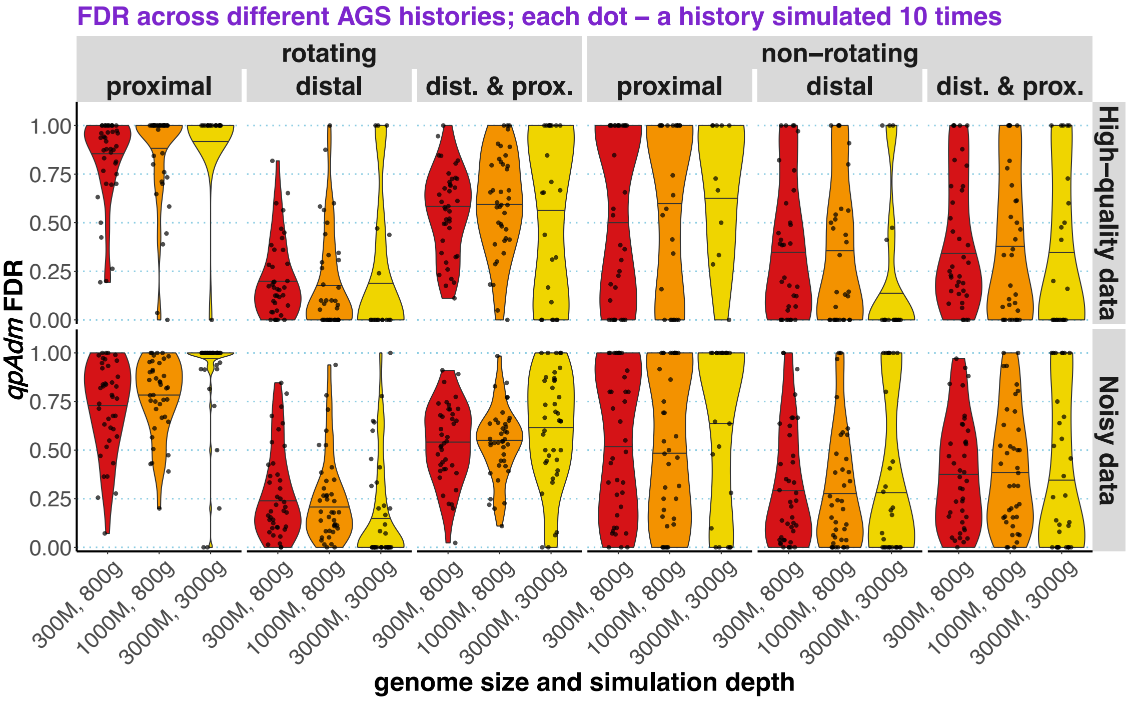


Variability of FDR across simulated AGS histories. Here FDR was calculated not per each simulation/subsampling replicate and all simulated histories combined, but for each AGS history and all simulation/subsampling replicates combined (shown as dots). FDR distributions are summarized as violin plots with medians and stratified by *qpAdm* protocols (rotating and non-rotating), by model subsets (proximal, distal, or both models included), by data quality, and by three combinations of simulated genome sizes (300, 1,000, or 3,000 Mbp) and maximal simulation depths (800 or 3,000 generations).

**Figure S12.**

**a**

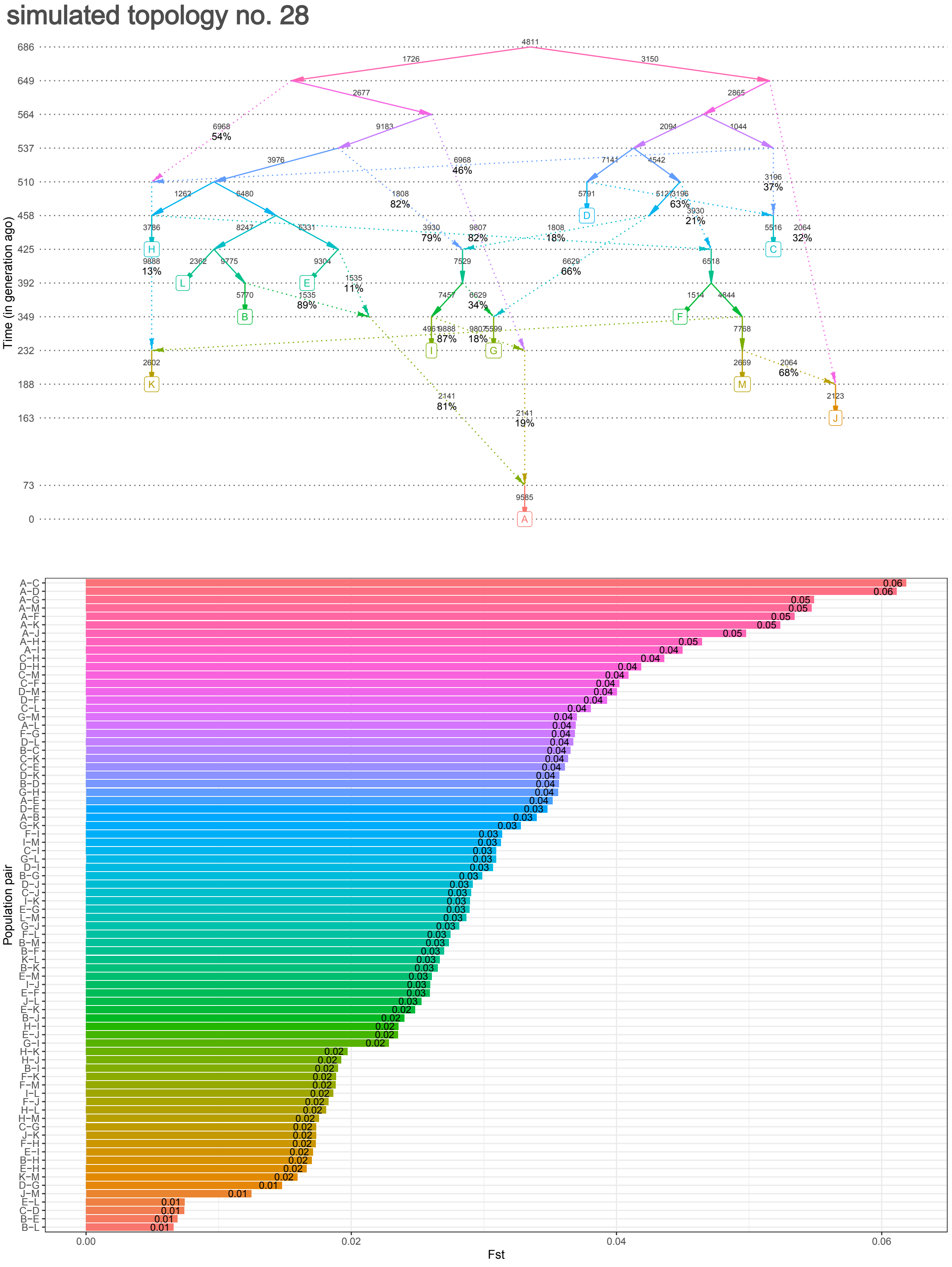
simulated topology no. 28
Time (in generation ago)
686
649
564
537
510
458
425
392
349
232
188
163
73
0
A
B
C
D
E
F
G
H
I
J
K
L
M
Population pair
Fst
0.00
0.02
0.04
0.06

**b**

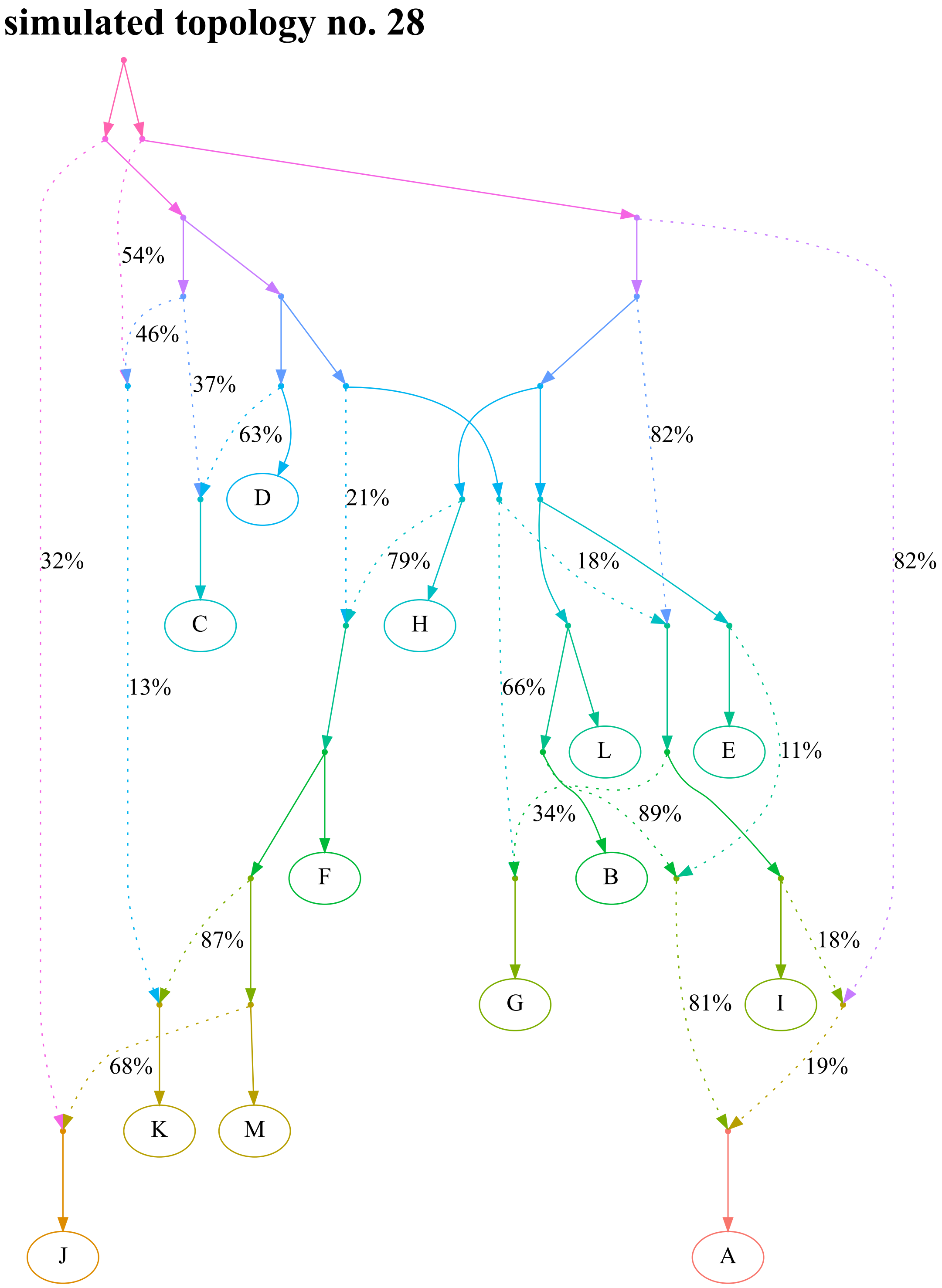

simulated topology no. 28
54%
46%
37%
63%
D
21%
82%
79%
18%
32%
82%
C
H
13%
66%
L
E
11%
34%
89%
F
B
87%
18%
G
81%
I
68%
19%
K
M
J
A

**c**

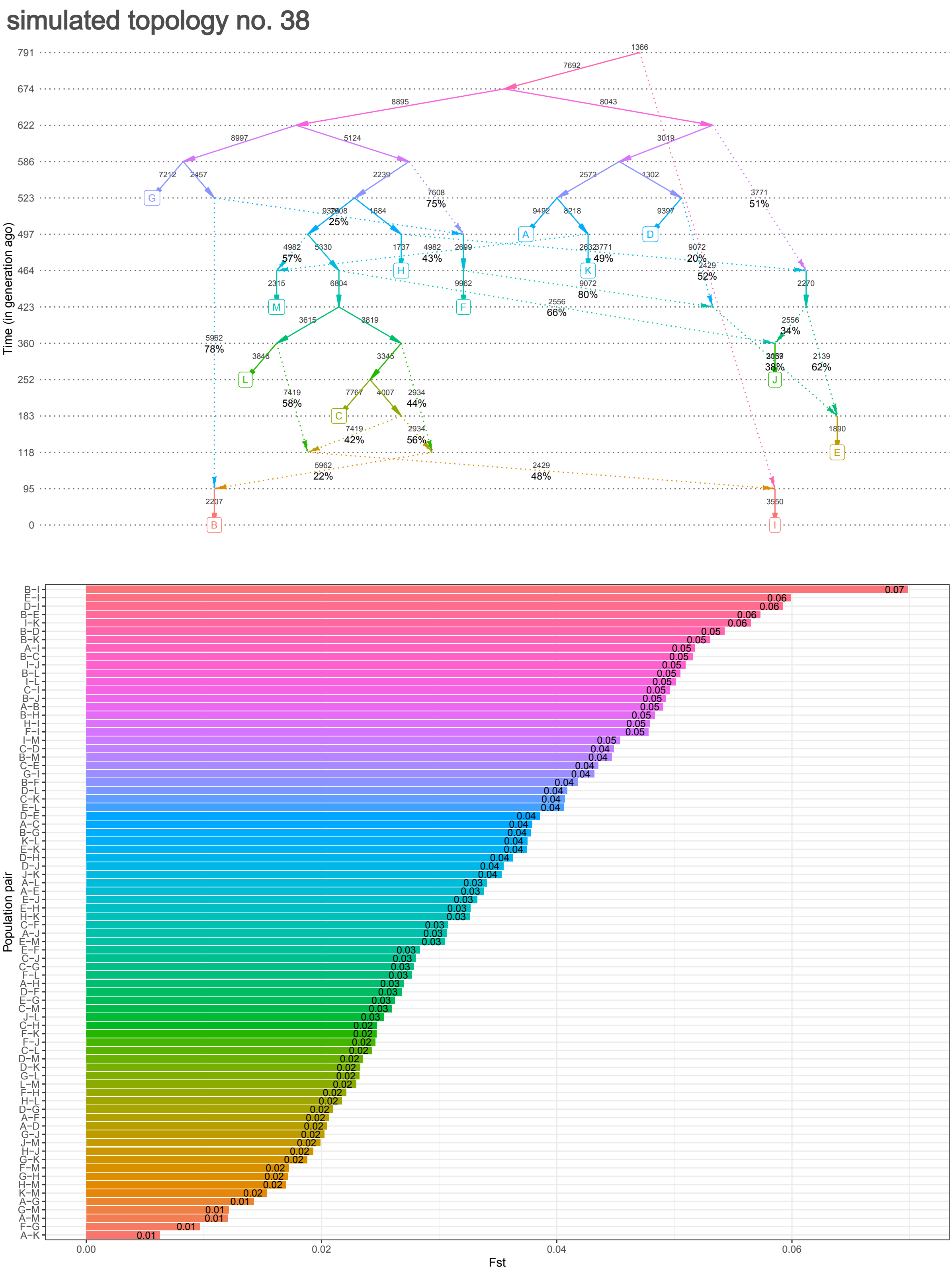
simulated topology no. 38
Time (in generation ago)
791
674
622
586
523
497
464
423
360
252
183
118
95
0
A
B
C
D
E
F
G
H
I
J
K
L
M
Population pair
Fst
0.00
0.02
0.04
0.06

**d**

#### simulated topology no. 38

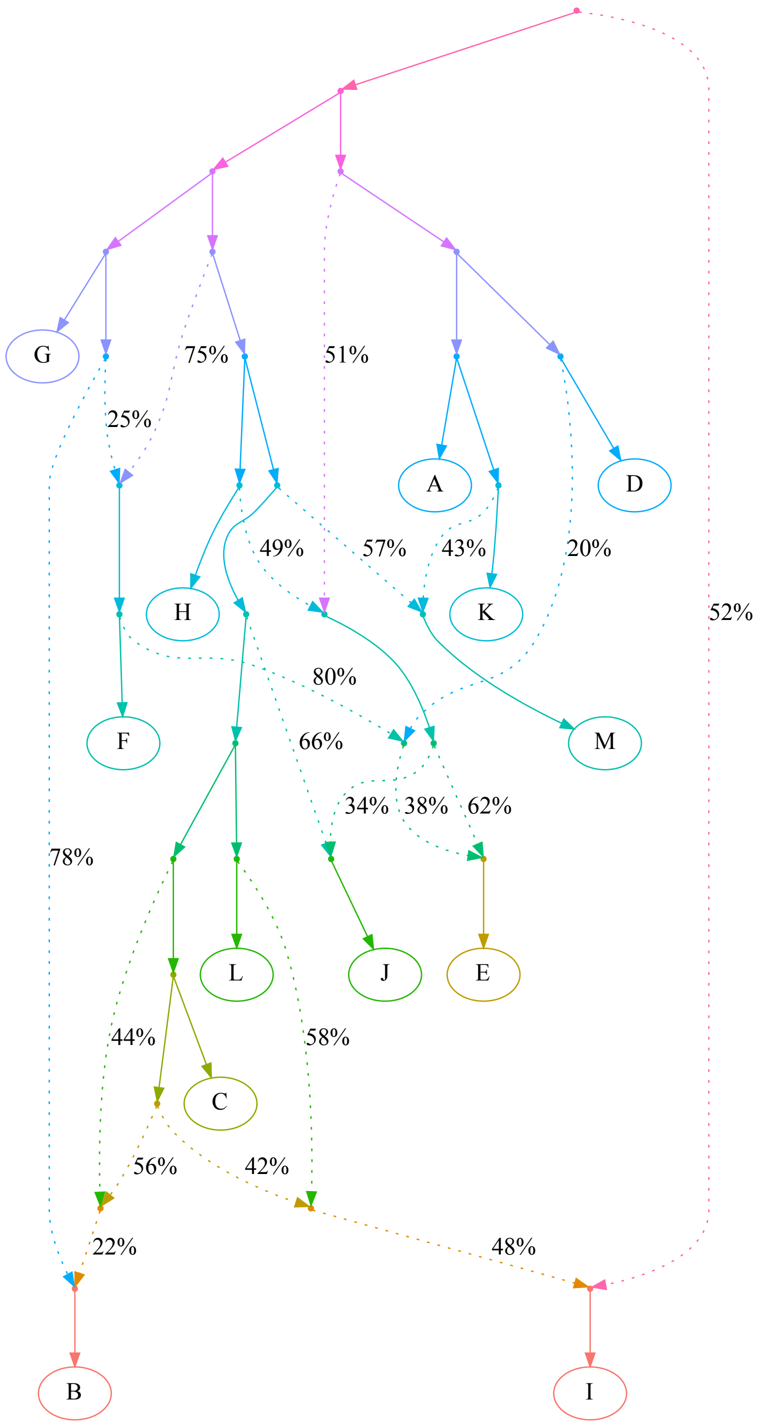

**e**

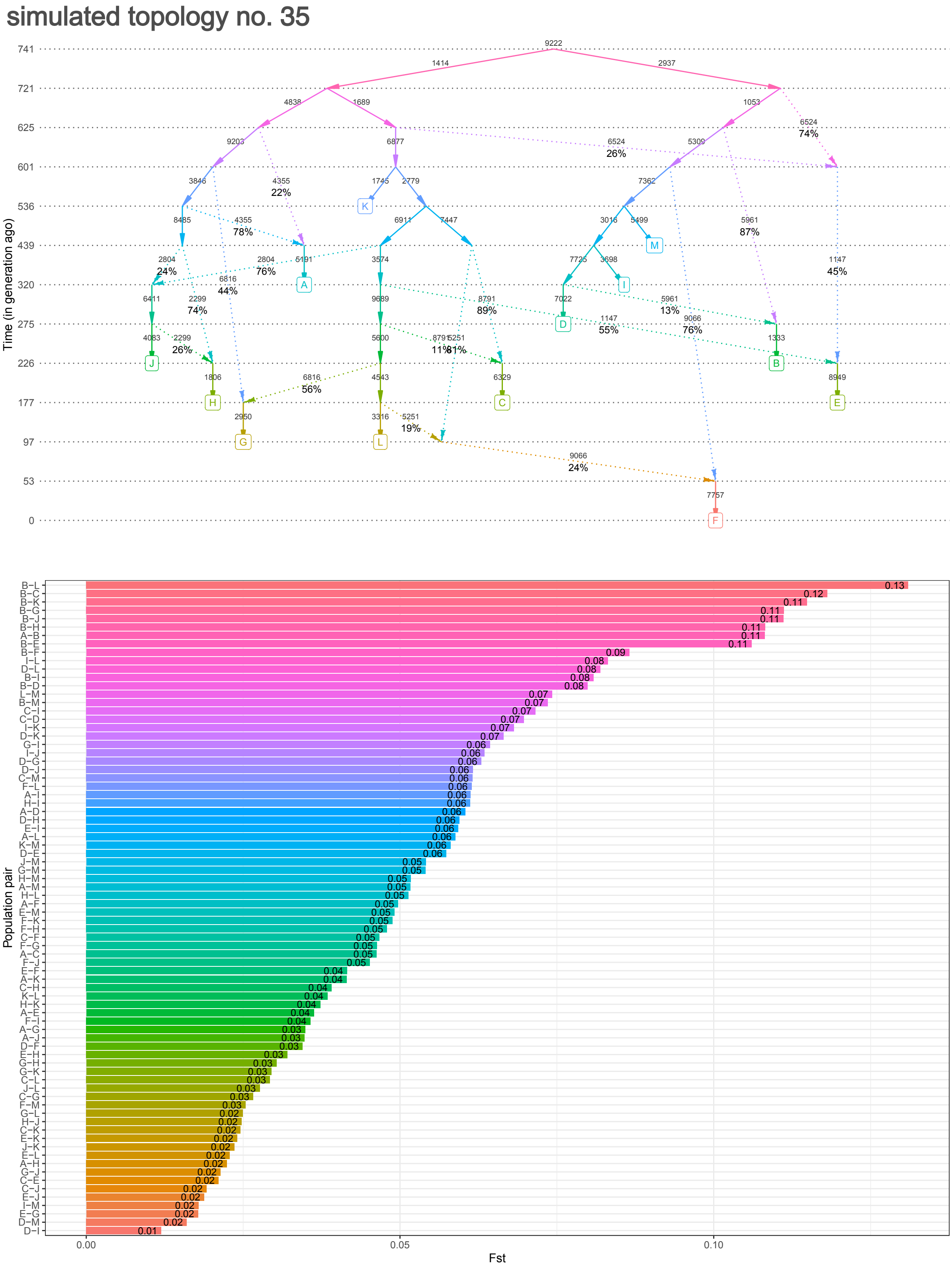
simulated topology no. 35
Time (in generation ago)
741
721
625
601
536
439
320
275
226
177
97
53
0
A
B
C
D
E
F
G
H
I
J
K
L
M
Population pair
Fst
0.00
0.05
0.10

**f**

#### simulated topology no. 35

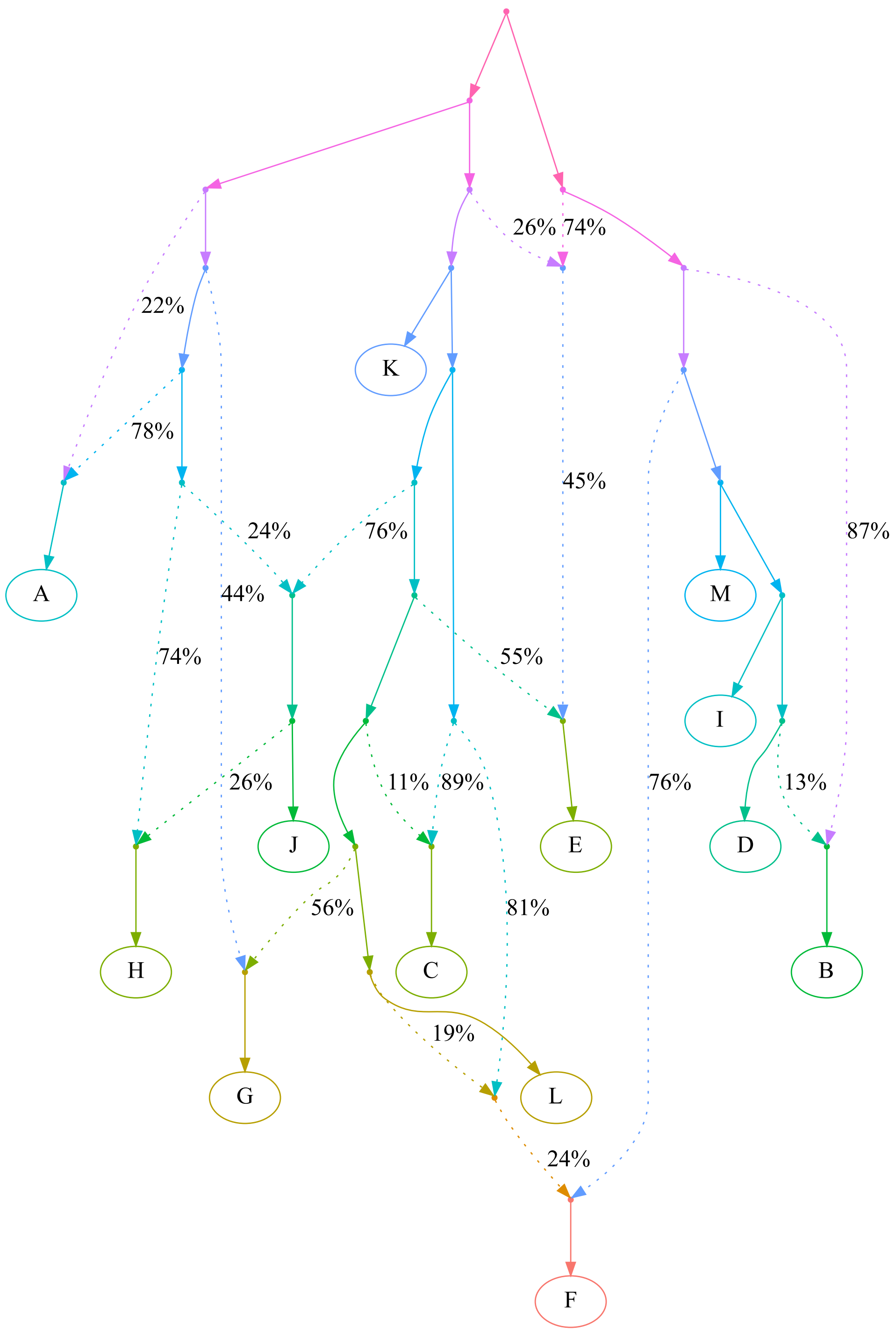

Simulated graph topologies most problematic for distal *qpAdm* protocols, i.e., those responsible for >20% of all FP 2-way *qpAdm* models in three or more sets of results. By a "set of results" we mean 40 genetic histories in the form of random admixture graph topologies of the same complexity simulated once and analyzed with a certain *qpAdm* protocol. The following topologies are shown: no. 28 (panels **a, b**; problematic under 8 of 80 "protocol/simulation setup/replicate" combinations, 4 of them including the rotating protocol), no. 38 (panels **c, d**; problematic under 8 of 80 "protocol/simulation setup/replicate" combinations, 7 of them including the rotating protocol), and no. 35 (panels **e, f**; problematic under 3 of 80 "protocol/simulation setup/replicate" combinations, all of them including the rotating protocol). In all cases, $F_{ST}$ values are shown for populations sampled at different times, simulation replicate no. 1, 300-Mbp-sized genomes, divergence times up to 800 generations, and diploid data. Time in generations is shown on the y-axis (for visual clarity, points on this axis are not spaced proportionately). Effective population sizes (in diploid individuals) are shown beside each edge. Sampled populations are labelled by letters in squares. Gene flows going from the past to the future were simulated as unsampled ghost populations. The same simulated histories are shown in a form more accessible to topological analysis, but without any additional information, in panels **b**, **d**, and **f**. The color gradients have no special meaning and are used for visual clarity only.

**Figure S13.**

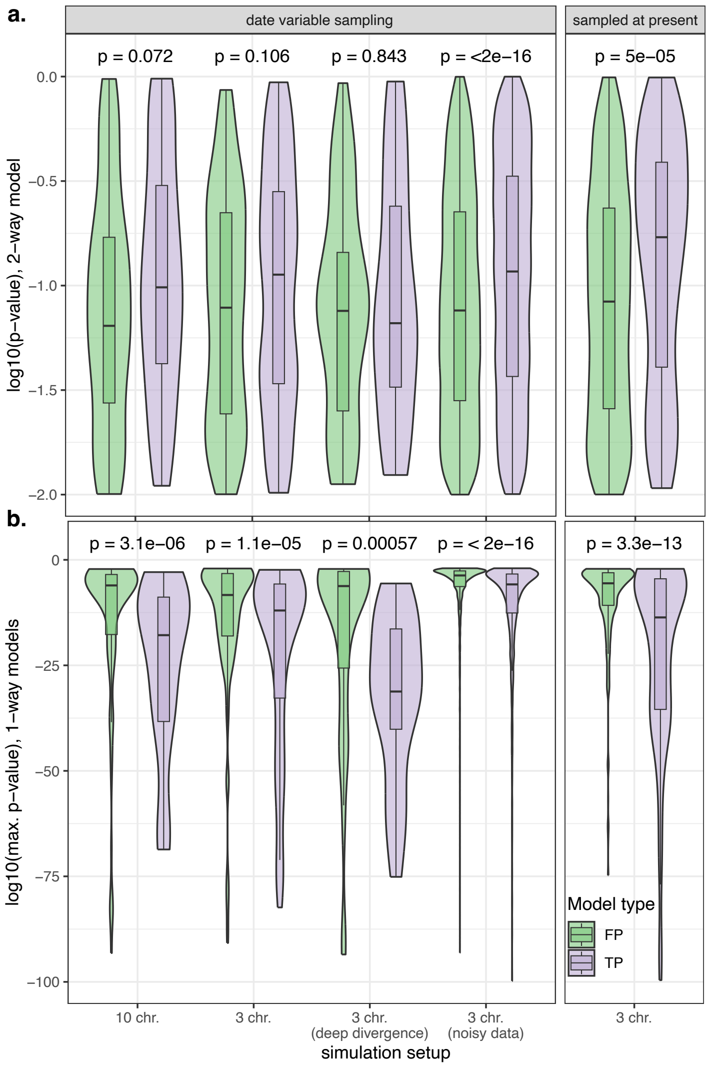


Comparing distributions of $log_{10}$(*p*-values) for true (TP) and false positive (FP) two-way *qpAdm* models (outcomes of the proximal rotating protocol). The distributions are summarized with boxplots and violin plots. In panel **a**, *p*-values for feasible two-way *qpAdm* models are shown. In panel **b**, the highest *p*-values in each group of three one-way models corresponding to a feasible two-way model are shown. Outliers having $log_{10}$(*p*-value) < -100 were removed (163 such models in total). The results are presented for five simulation setups labelled on the *x*-axis, for one simulation replicate per setup. Multiple subsampling replicates were considered in the case of low-quality data. The means were compared using the non-parametric Wilcoxon test (see *p*-values above the paired plots).

**Figure S14.**

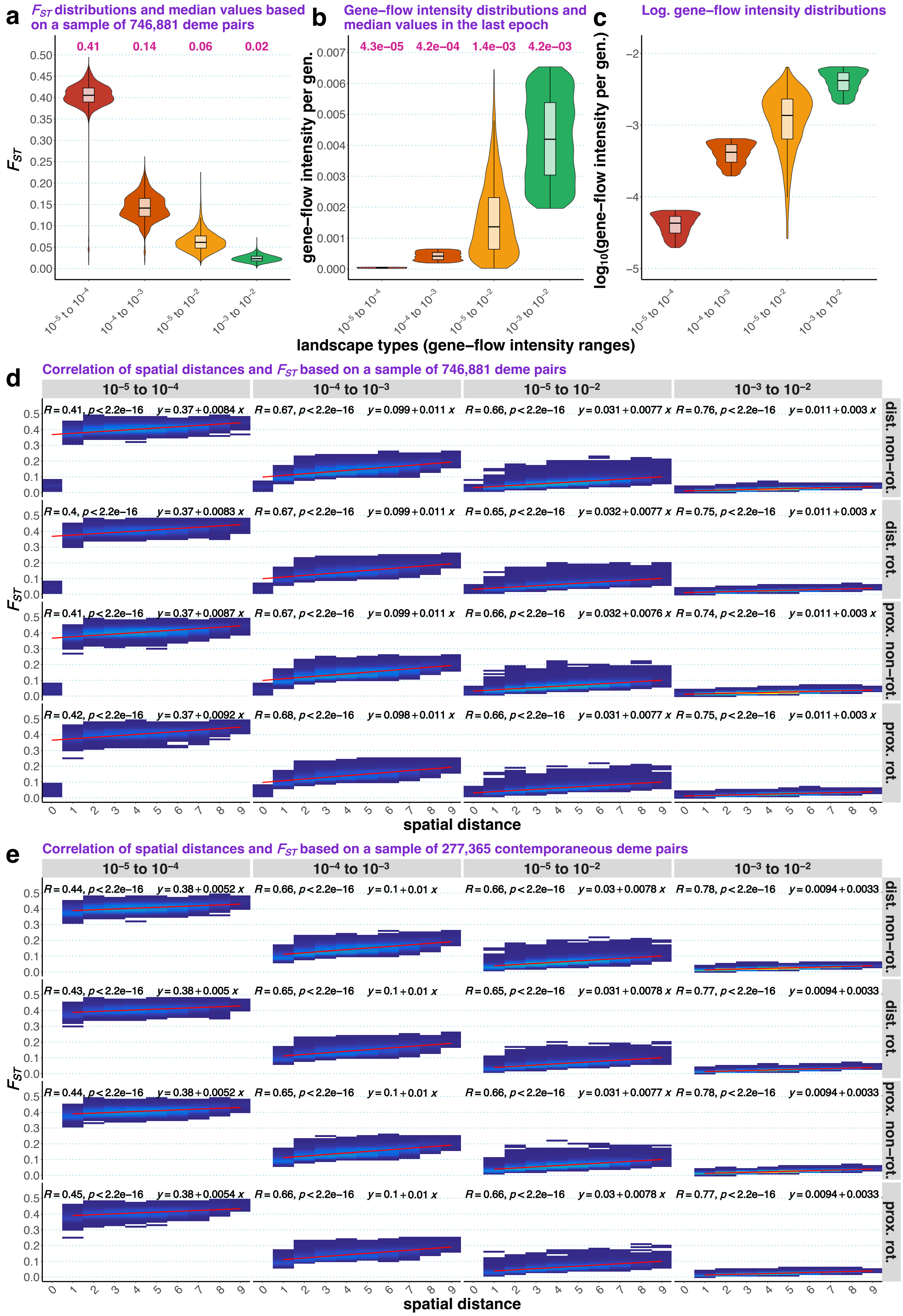
a
$F_{ST}$ distributions and median values based on a sample of 746,881 deme pairs
0.41 0.14 0.06 0.02
$F_{ST}$
b
Gene–flow intensity distributions and median values in the last epoch
4.3e–05 4.2e–04 1.4e–03 4.2e–03
gene–flow intensity per gen.
c
Log. gene–flow intensity distributions
$\log_{10}$(gene–flow intensity per gen.)
$10^{-5}$ to $10^{-4}$
$10^{-4}$ to $10^{-3}$
$10^{-5}$ to $10^{-2}$
$10^{-3}$ to $10^{-2}$
landscape types (gene–flow intensity ranges)
d
Correlation of spatial distances and $F_{ST}$ based on a sample of 746,881 deme pairs
$R=0.41$, $p<2.2e-16$ $y=0.37+0.0084x$
$R=0.67$, $p<2.2e-16$ $y=0.099+0.011x$
$R=0.66$, $p<2.2e-16$ $y=0.031+0.0077x$
$R=0.76$, $p<2.2e-16$ $y=0.011+0.003x$
$R=0.4$, $p<2.2e-16$ $y=0.37+0.0083x$
$R=0.67$, $p<2.2e-16$ $y=0.099+0.011x$
$R=0.65$, $p<2.2e-16$ $y=0.032+0.0077x$
$R=0.75$, $p<2.2e-16$ $y=0.011+0.003x$
$R=0.41$, $p<2.2e-16$ $y=0.37+0.0087x$
$R=0.67$, $p<2.2e-16$ $y=0.099+0.011x$
$R=0.66$, $p<2.2e-16$ $y=0.032+0.0076x$
$R=0.74$, $p<2.2e-16$ $y=0.011+0.003x$
$R=0.42$, $p<2.2e-16$ $y=0.37+0.0092x$
$R=0.68$, $p<2.2e-16$ $y=0.098+0.011x$
$R=0.66$, $p<2.2e-16$ $y=0.031+0.0077x$
$R=0.75$, $p<2.2e-16$ $y=0.011+0.003x$
dist. non–rot.
dist. rot.
prox. non–rot.
prox. rot.
spatial distance
e
Correlation of spatial distances and $F_{ST}$ based on a sample of 277,365 contemporaneous deme pairs
$R=0.44$, $p<2.2e-16$ $y=0.38+0.0052x$
$R=0.66$, $p<2.2e-16$ $y=0.1+0.01x$
$R=0.66$, $p<2.2e-16$ $y=0.03+0.0078x$
$R=0.78$, $p<2.2e-16$ $y=0.0094+0.0033$
$R=0.43$, $p<2.2e-16$ $y=0.38+0.005x$
$R=0.65$, $p<2.2e-16$ $y=0.1+0.01x$
$R=0.65$, $p<2.2e-16$ $y=0.031+0.0078x$
$R=0.77$, $p<2.2e-16$ $y=0.0094+0.0033$
$R=0.44$, $p<2.2e-16$ $y=0.38+0.0052x$
$R=0.65$, $p<2.2e-16$ $y=0.1+0.01x$
$R=0.66$, $p<2.2e-16$ $y=0.031+0.0077x$
$R=0.78$, $p<2.2e-16$ $y=0.0094+0.0033$
$R=0.45$, $p<2.2e-16$ $y=0.38+0.0054x$
$R=0.66$, $p<2.2e-16$ $y=0.1+0.01x$
$R=0.66$, $p<2.2e-16$ $y=0.03+0.0078x$
$R=0.77$, $p<2.2e-16$ $y=0.0094+0.0033$
dist. non–rot.
dist. rot.
prox. non–rot.
prox. rot.
spatial distance

(**a**) Distributions of $F_{ST}$ values in sets of 14 (in the case of rotating protocols) or 21 demes (in the case of non-rotating protocols) used for randomized *qpAdm* tests. The *qpAdm* protocols did not influence $F_{ST}$ distributions, that is why the results are stratified by SSL type only. Median $F_{ST}$ values are shown above the violin plots. (**b**) Distributions of per-generation gene-flow intensities in the last epoch (lasting for 769 generations, except for the "$10^{-3}$ to $10^{-2}$" landscape where it lasted 153 generations) for the same SSL types. Gene flows were simulated between the nearest neighbors only, and forward and reverse intensities were considered separately in these distributions. Median per-generation gene-flow intensity values are shown above the violin plots. (**c**) The same plot as in **b**, but the y-axis scale is logarithmic. (**d**) Density plots illustrating correlation of spatial distances and $F_{ST}$ in a sample of deme pairs derived from *qpAdm* experiments, stratified by SSL types and *qpAdm* protocols. The space is subdivided into rectangular bins, and density of deme pairs in this space is shown with a linear color scale. Pearson's correlation coefficients and linear equations are shown above each plot. (**e**) The same plot as in **d**, but considering contemporaneous deme pairs only.

**Figure S15.**

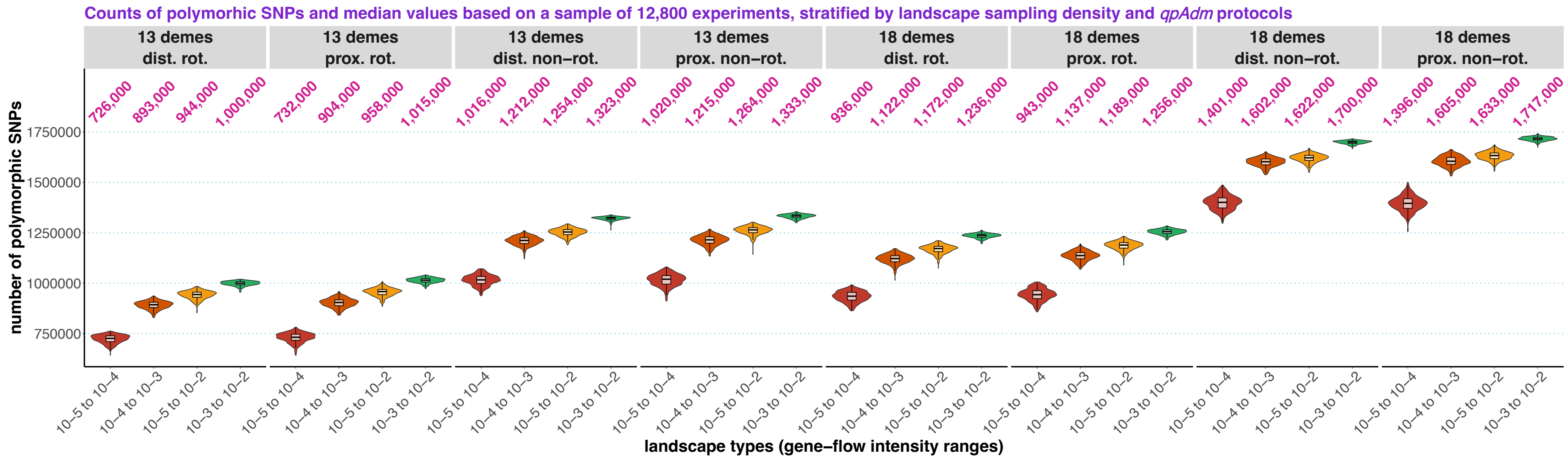


Counts of SNPs polymorphic in sets of 14 or 19 (in the case of rotating protocols), 21 or 31 demes (in the case of non-rotating protocols) used for randomized *qpAdm* tests. The results are stratified by SSL type, landscape sampling density (13 or 18 demes in rotating sets and 10 or 15 demes in proxy and "right" sets, respectively), and *qpAdm* protocol. Median SNP counts rounded to thousands are shown above the violin plots.

**Figure S16.**

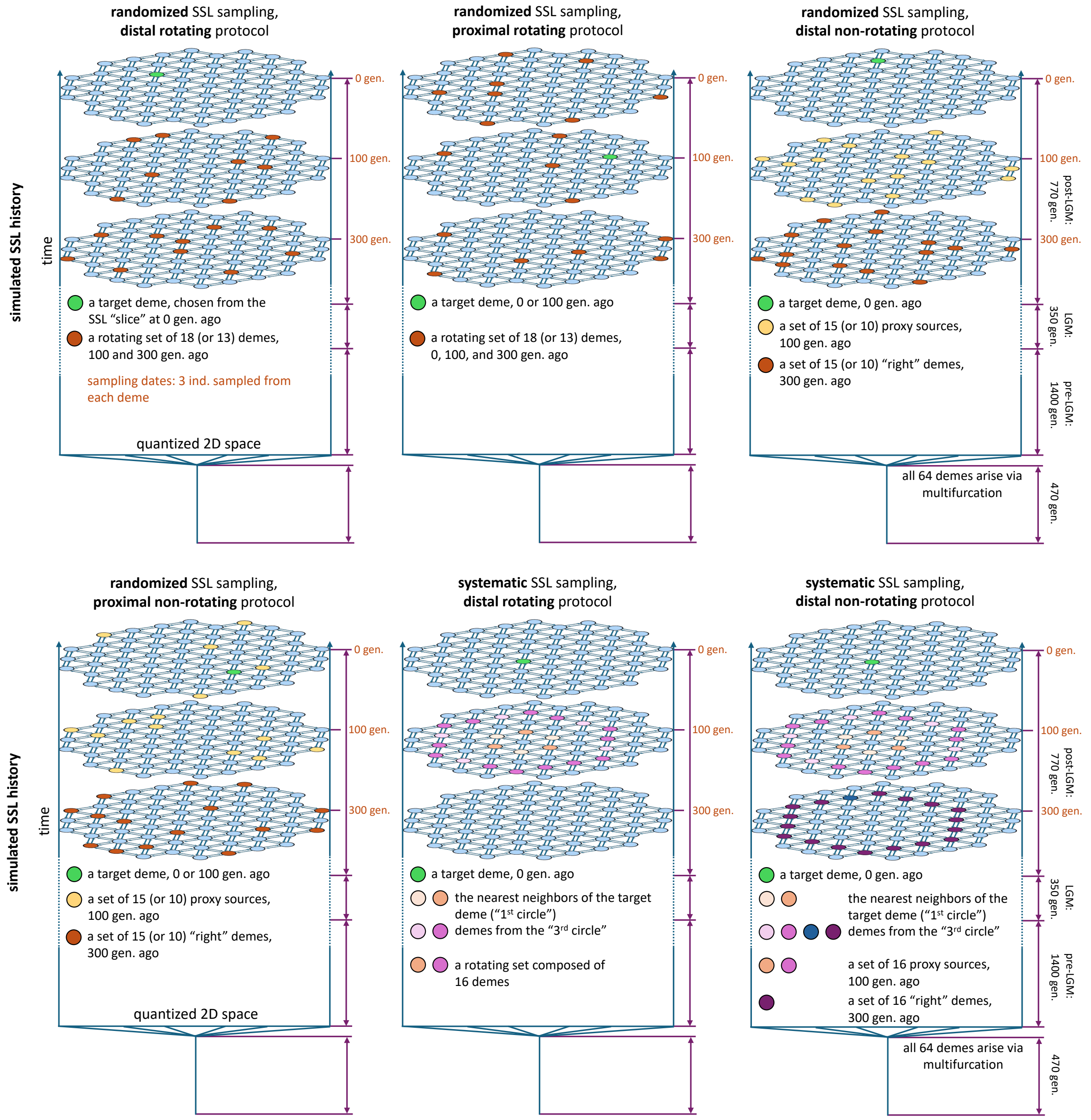


Diagrams summarizing *qpAdm* protocols tested in this study that are based on randomized or systematic SSL sampling (four and two protocols, respectively). Various categories of demes (targets, proxy sources, those belonging to rotating and "right" sets) are shown in various colors and labelled according to the legends in each panel.

**Figure S17.**

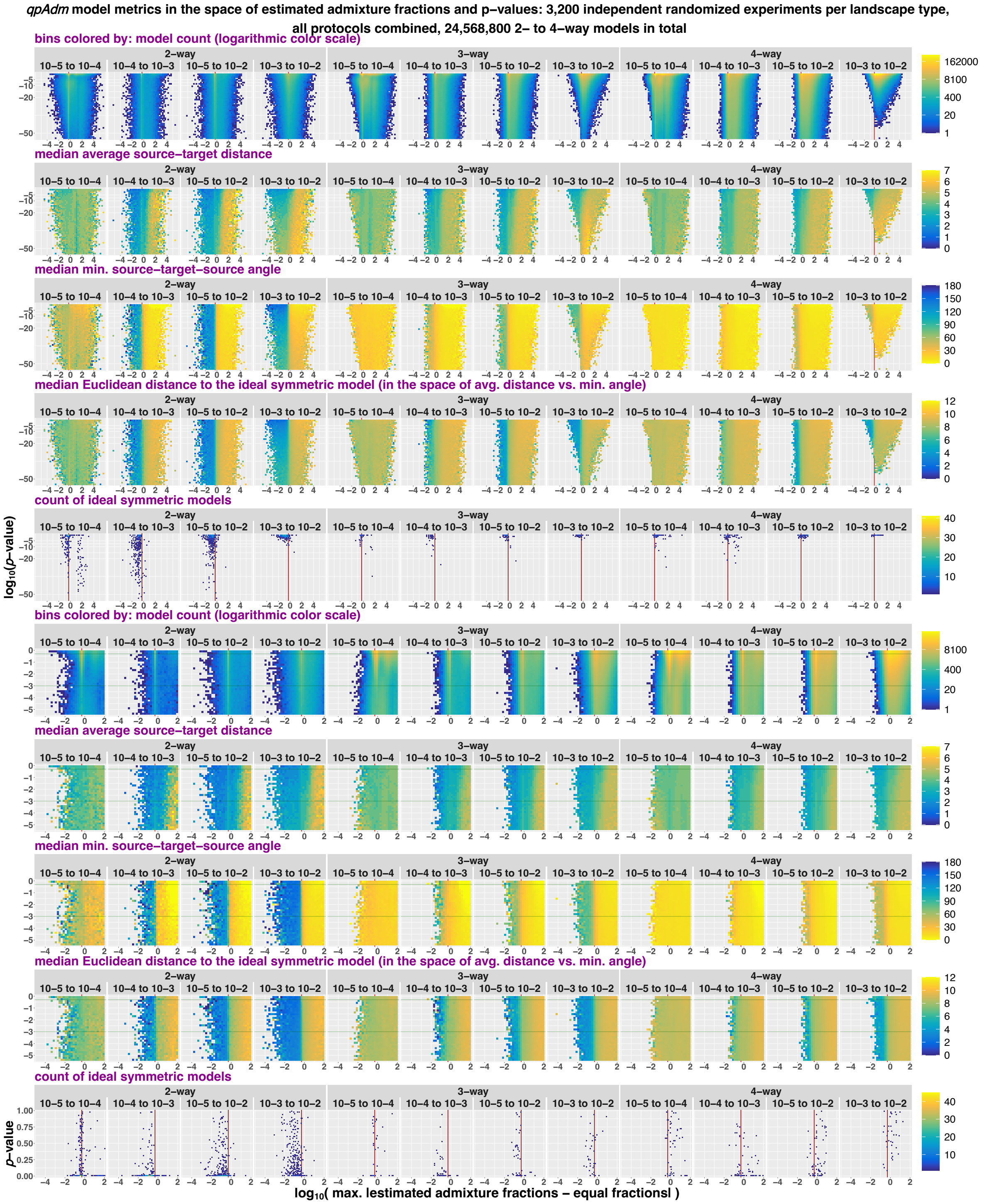


Randomized two to four-way *qpAdm* models in the space of max|*EAF–EF*| and *p*-values (both axes are logarithmic). Results for all *qpAdm* protocols are combined but stratified by model complexity and landscape type. Two sections of the space are shown: *p*-values from $10^{-55}$ to 1 (the upper five rows of plots) and from $3.16\times10^{-6}$ to 1 (the four rows below them). The space is divided into rectangular bins (50×50 for the larger section and 35×35 for the smaller section), and they are colored by density of individual *qpAdm* models populating this space (logarithmic color scale) or by median values of model optimality metrics in those bins: average ST distance, min. STS angle, and Euclidean distance to the ideal symmetric model.

Density of ideal symmetric models (in the case of four-way models, the most optimal non-ideal models available on our SSL) in this space is also shown (in 75×75 bins). To assess if distributions of ideal symmetric models over *p*-values are uniform, we use non-logarithmic *p*-value scale (the bottom row of plots). The red vertical lines (or the red tick marks) mark max|*EAF*–*EF*| = 0.5, and the green horizontal lines mark the lowest and highest *p*-value thresholds used in this study (0.001 and 0.5).

**Figure S18.**

**a**

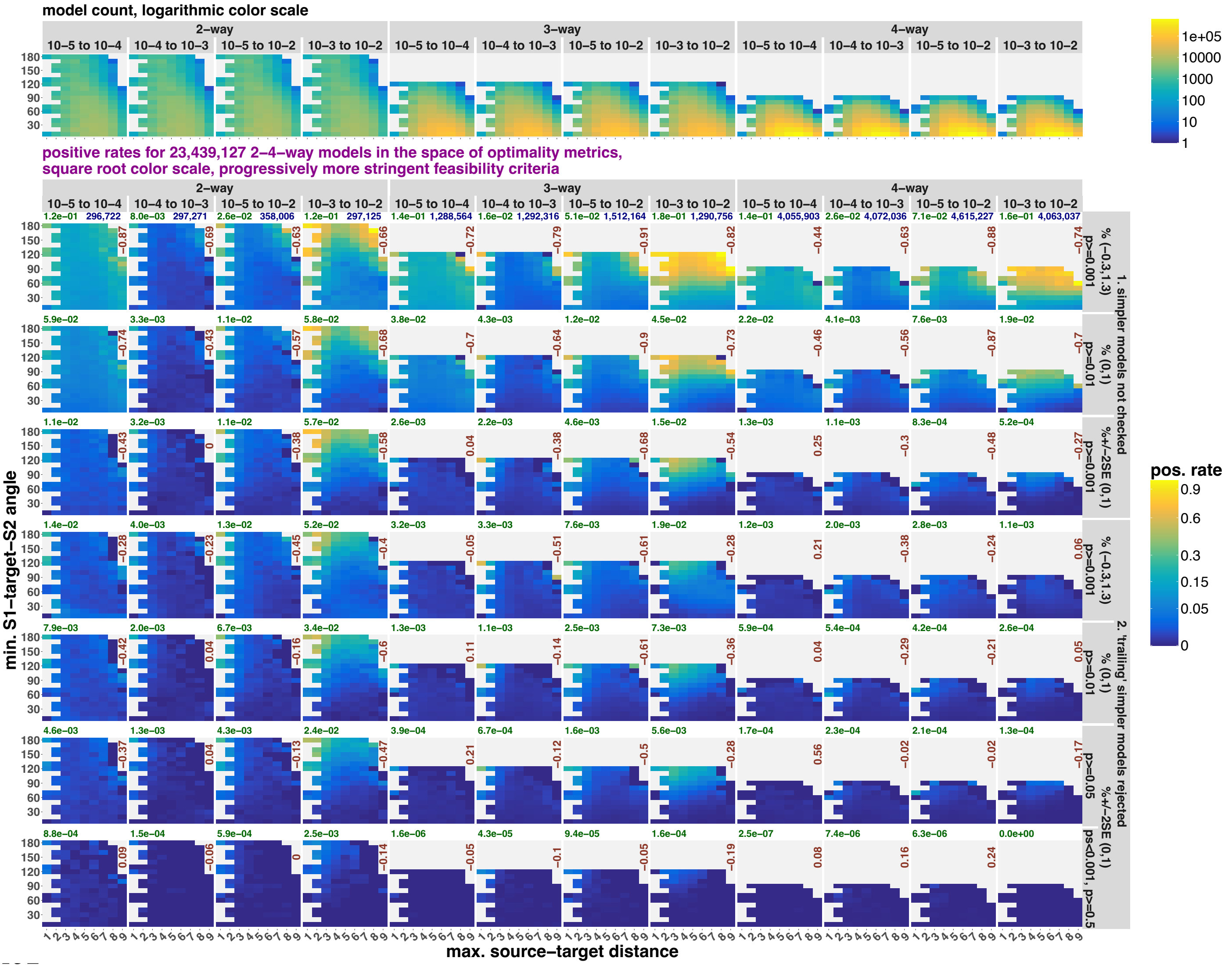

model count, logarithmic color scale
2–way
3–way
4–way
10–5 to 10–4
10–4 to 10–3
10–5 to 10–2
10–3 to 10–2
positive rates for 23,439,127 2–4–way models in the space of optimality metrics,
square root color scale, progressively more stringent feasibility criteria
min. S1–target–S2 angle
max. source–target distance
pos. rate
1. simpler models not checked
2. 'trailing' simpler models rejected

**b**

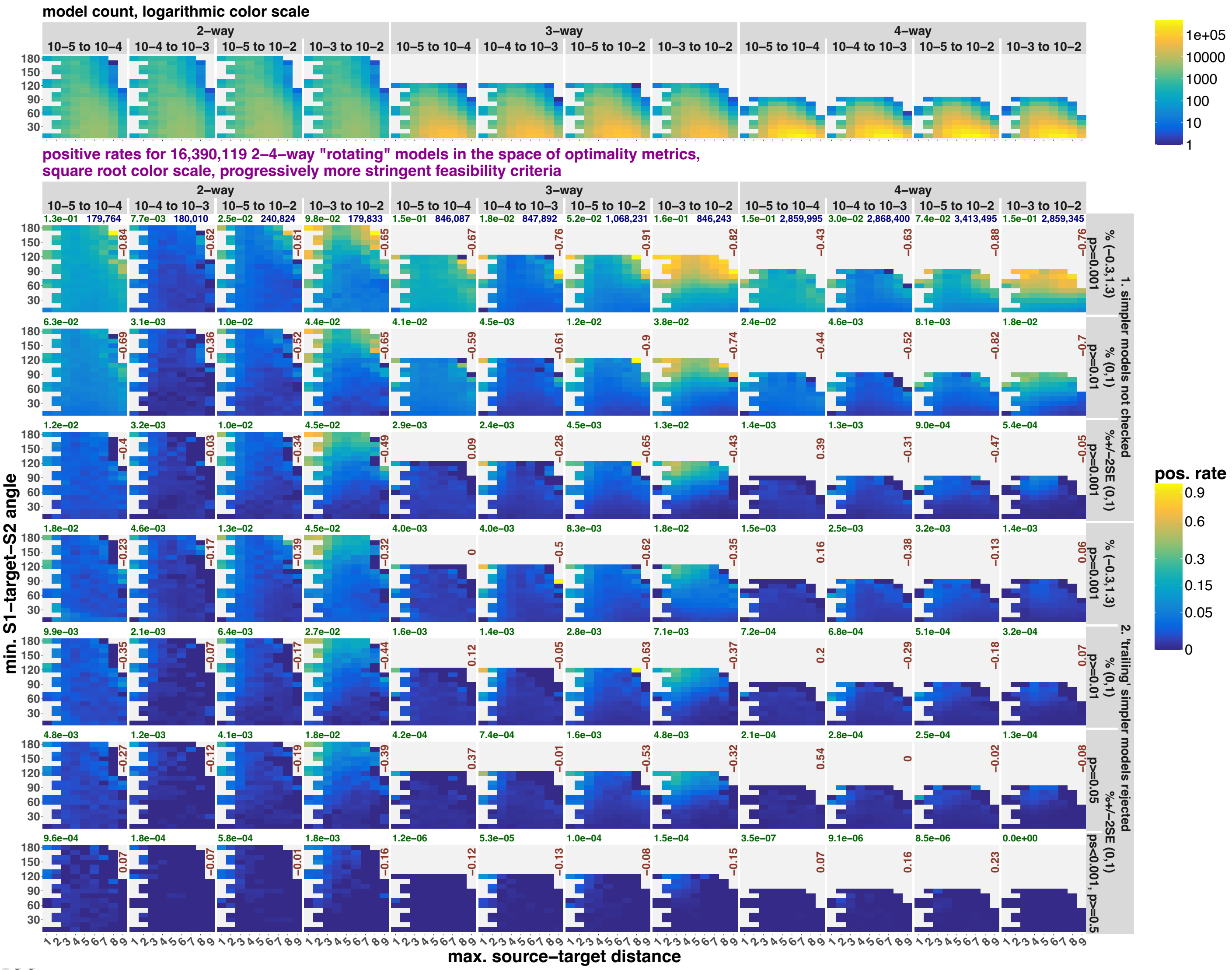

model count, logarithmic color scale
2–way
3–way
4–way
10–5 to 10–4
10–4 to 10–3
10–5 to 10–2
10–3 to 10–2
1e+05
10000
1000
100
10
1
positive rates for 16,390,119 2–4–way "rotating" models in the space of optimality metrics,
square root color scale, progressively more stringent feasibility criteria
pos. rate
0.9
0.6
0.3
0.15
0.05
0
min. S1–target–S2 angle
max. source–target distance
1. simpler models not checked
2. 'trailing' simpler models rejected
% (−0.3,1.3) p≥0.001
% (0,1) p≥0.01
%+/−2SE (0,1) p≥0.001
% (−0.3,1.3) p≥0.001
% (0,1) p≥0.01
p≥0.05
%+/−2SE (0,1) ps<0.001, p≥0.5

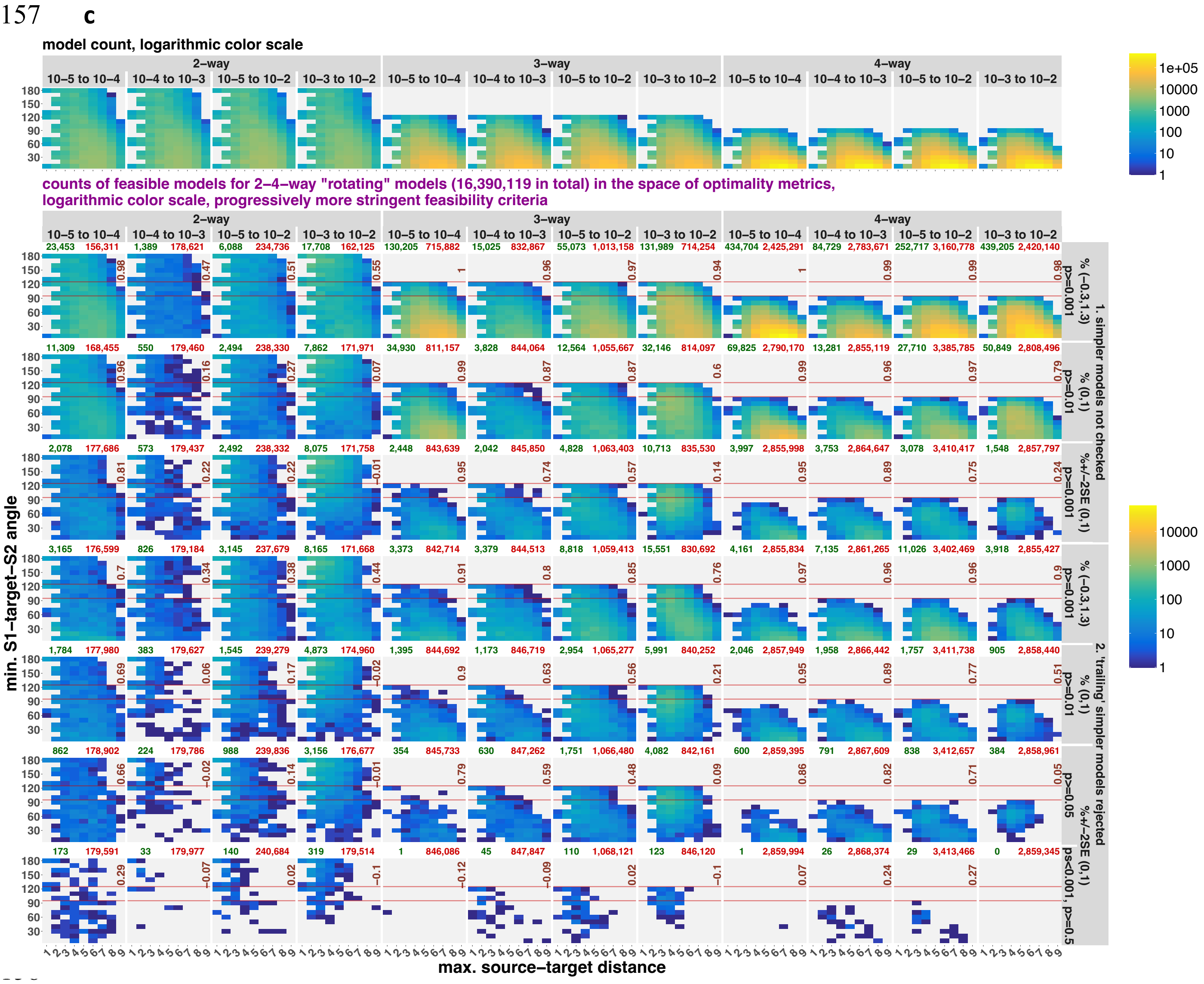
c
model count, logarithmic color scale
2–way
3–way
4–way
10–5 to 10–4
10–4 to 10–3
10–5 to 10–2
10–3 to 10–2
1e+05
10000
1000
100
10
1
counts of feasible models for 2–4–way "rotating" models (16,390,119 in total) in the space of optimality metrics,
logarithmic color scale, progressively more stringent feasibility criteria
23,453 156,311 1,389 178,621 6,088 234,736 17,708 162,125 130,205 715,882 15,025 832,867 55,073 1,013,158 131,989 714,254 434,704 2,425,291 84,729 2,783,671 252,717 3,160,778 439,205 2,420,140
11,309 168,455 550 179,460 2,494 238,330 7,862 171,971 34,930 811,157 3,828 844,064 12,564 1,055,667 32,146 814,097 69,825 2,790,170 13,281 2,855,119 27,710 3,385,785 50,849 2,808,496
2,078 177,686 573 179,437 2,492 238,332 8,075 171,758 2,448 843,639 2,042 845,850 4,828 1,063,403 10,713 835,530 3,997 2,855,998 3,753 2,864,647 3,078 3,410,417 1,548 2,857,797
3,165 176,599 826 179,184 3,145 237,679 8,165 171,668 3,373 842,714 3,379 844,513 8,818 1,059,413 15,551 830,692 4,161 2,855,834 7,135 2,861,265 11,026 3,402,469 3,918 2,855,427
1,784 177,980 383 179,627 1,545 239,279 4,873 174,960 1,395 844,692 1,173 846,719 2,954 1,065,277 5,991 840,252 2,046 2,857,949 1,958 2,866,442 1,757 3,411,738 905 2,858,440
862 178,902 224 179,786 988 239,836 3,156 176,677 354 845,733 630 847,262 1,751 1,066,480 4,082 842,161 600 2,859,395 791 2,867,609 838 3,412,657 384 2,858,961
173 179,591 33 179,977 140 240,684 319 179,514 1 846,086 45 847,847 110 1,068,121 123 846,120 1 2,859,994 26 2,868,374 29 3,413,466 0 2,859,345
min. S1–target–S2 angle
max. source–target distance
% (−0.3,1.3) p>=0.001
% (0,1) p>=0.01
%+/−2SE (0,1) p>=0.001
% (−0.3,1.3) p>=0.001
% (0,1) p>=0.01
%+/−2SE (0,1) p>=0.05
ps<0.001, p>=0.5
1. simpler models not checked
2. 'trailing' simpler models rejected

**d**

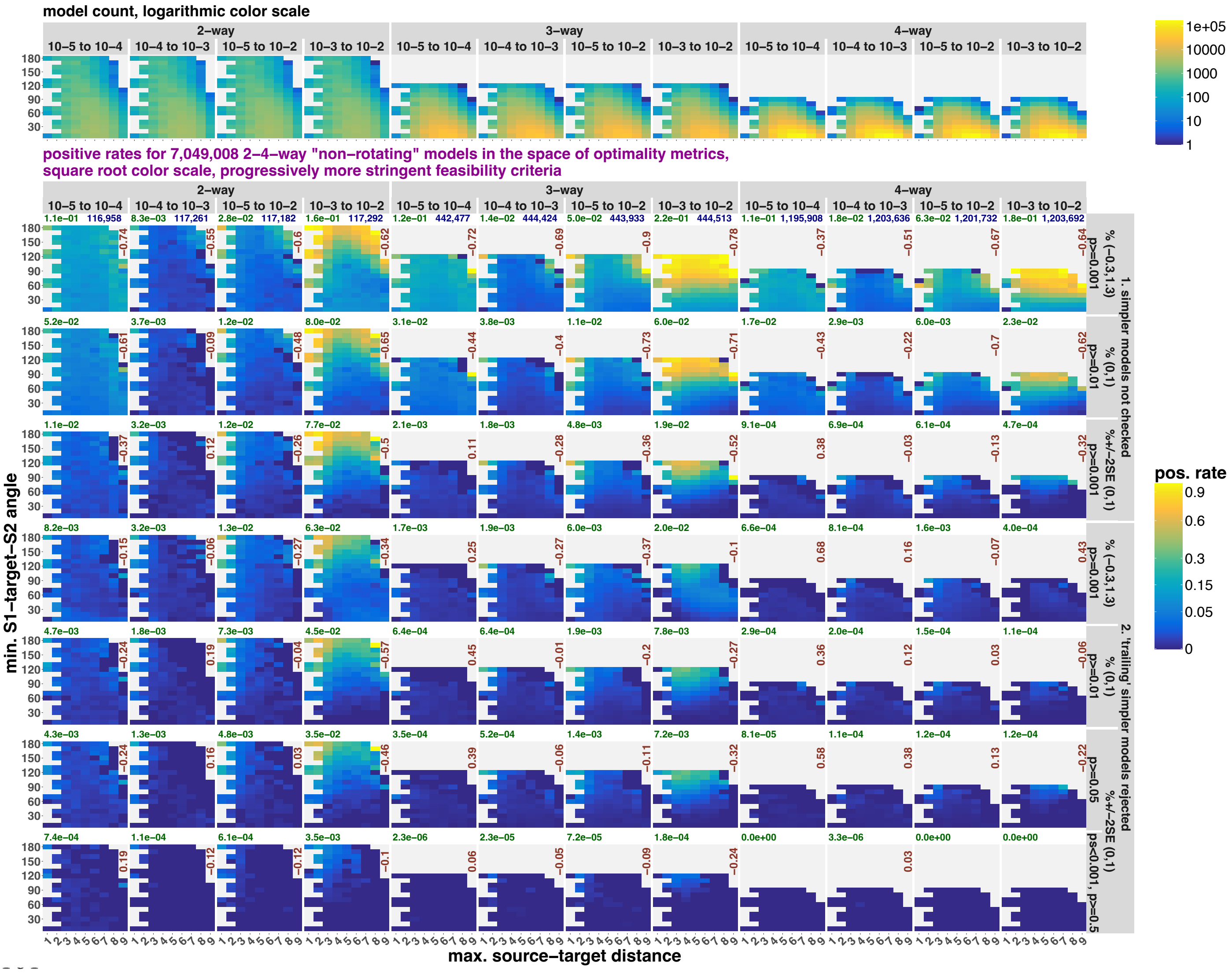

model count, logarithmic color scale
2–way
3–way
4–way
10–5 to 10–4
10–4 to 10–3
10–5 to 10–2
10–3 to 10–2
positive rates for 7,049,008 2–4–way "non–rotating" models in the space of optimality metrics, square root color scale, progressively more stringent feasibility criteria
min. S1–target–S2 angle
max. source–target distance
pos. rate
1. simpler models not checked
2. 'trailing' simpler models rejected

**e**

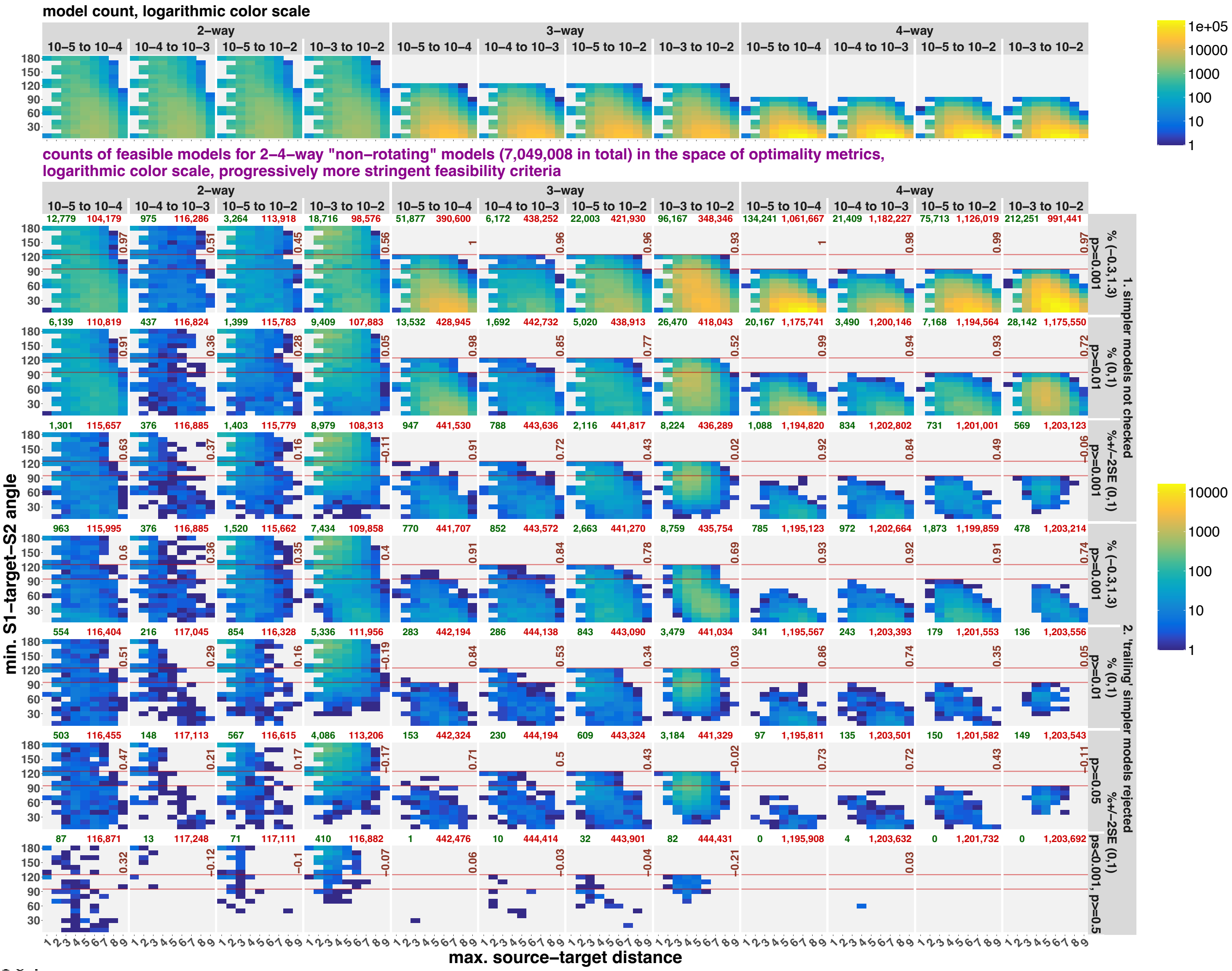


Randomized two to four-way *qpAdm* models in the space of model optimality metrics: max. ST distance and min. STS angle (models with the latter metric undefined were excluded from this analysis). Results are either combined for all *qpAdm* protocols (panel **a**) or shown separately for rotating (**b, c**) and non-rotating protocols (**d, e**). In all cases, the results are stratified by model complexity and landscape type. The space was divided into 9 bins on the x-axis (bin width = 1), and 18 bins on the y-axis (intervals [0°, 10°], (10°, 20°], and so forth). The upper row of plots shows density of all randomized models in those two-dimensional bins, and the other rows show either *qpAdm* positive rates (fractions of *qpAdm* models satisfying the selected feasibility criteria; panels **a, b, d**) or density of fitting models (**c, e**). The selected feasibility criteria are listed in the captions on the right (7 of 36 criteria tested in this study, from the least stringent on top to the most stringent at the bottom). Logarithmic color scales are used for showing density of individual *qpAdm* models populating the space, and square root color scales for *qpAdm* positive rate. Spearman's correlation coefficients for counts of all models vs. counts of fitting models or *qpAdm* positive rates in the two-dimensional bins are shown in each panel in brown (bins not populated by any models were not considered). The red horizontal lines mark the "ideal" min. STS angles for three- and four-way models (120° and 90°). The numbers above plots show either overall *qpAdm* positive rates (in green) in those analyses and total counts of models tested in blue (**a, b, d**), or total counts of fitting and non-fitting models in green and red, respectively (**c, e**).

**Figure S19.**

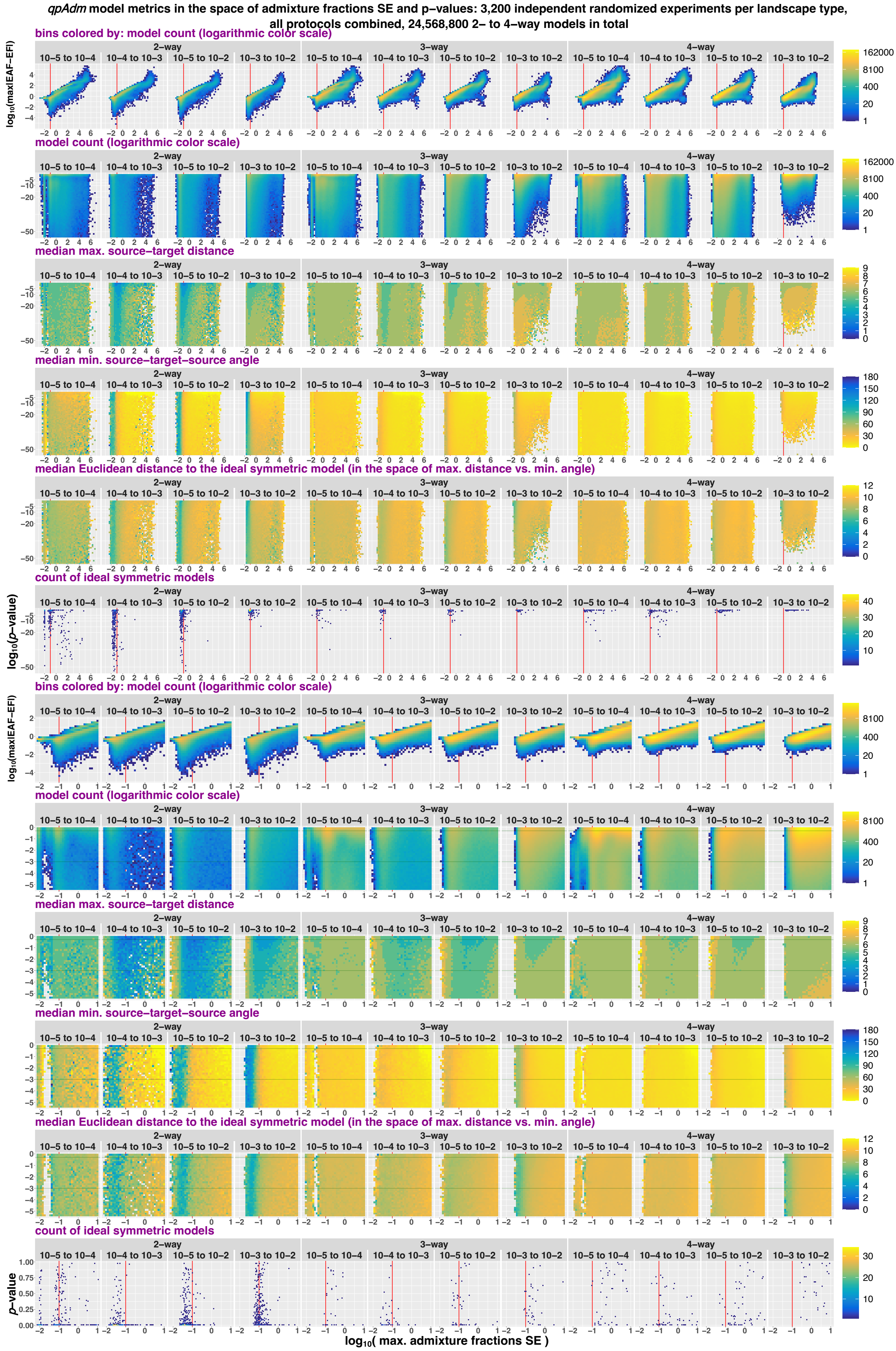


Randomized two to four-way *qpAdm* models in the space of max. SE of EAF and *p*-values (both axes are logarithmic). Results for all *qpAdm* protocols are combined but stratified by model complexity and landscape type. Two sections of the space are shown: *p*-values from $10^{-55}$ to 1 (the upper five rows of plots) and from $3.16\times10^{-6}$ to 1 (the four rows below them). The space is divided into rectangular bins (50×50 for the larger section and 35×35 for the smaller section), and they are colored by density of individual *qpAdm* models populating this space (logarithmic color scale) or by median values of model optimality metrics in those bins: max. ST distance, min. STS angle, and Euclidean distance to the ideal symmetric model. Density of ideal symmetric models (in the case of four-way models, the most optimal non-ideal models available on our SSL) in this space is also shown (in 75×75 bins). To assess if distributions of ideal symmetric models over *p*-values are uniform, we use non-logarithmic *p*-value scale (the bottom row of plots). The red vertical lines (or the red tick marks) mark max. SE = 0.1, and the green horizontal lines mark the lowest and highest *p*-value thresholds used in this study (0.001 and 0.5). We also visualize the relationship between max. SE and max|*EAF*–*EF*| (density of individual *qpAdm* models populating that space).

**Figure S20.**

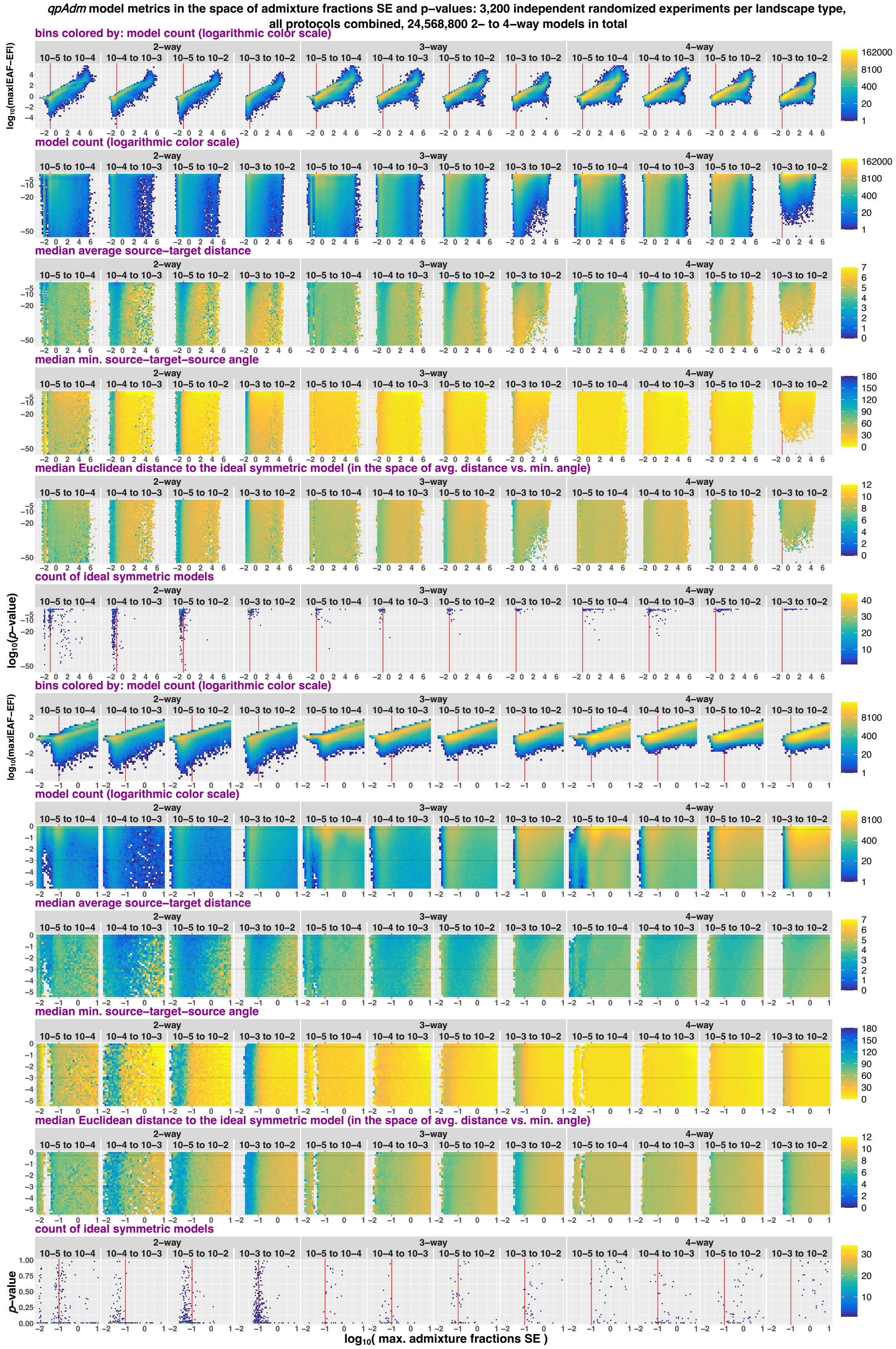


The same results as in **Fig. S19** but showing average ST distance instead of max. ST distance.

**Figure S21.**

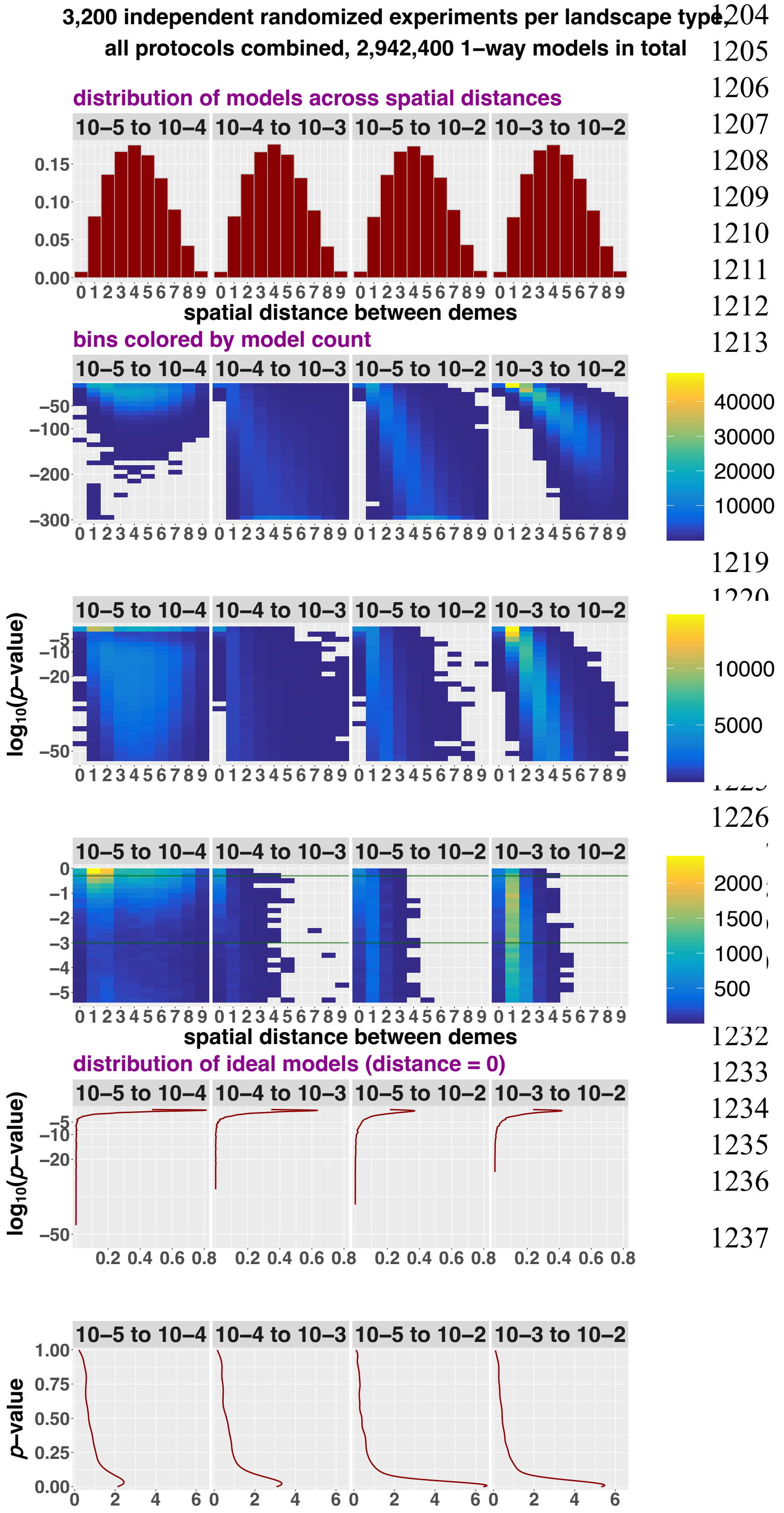


Randomized one-way *qpAdm* models in the space of ST distance and *p*-values (the *p*-value axis is logarithmic). Results for all *qpAdm* protocols are combined but stratified by landscape type. Three sections of the space are shown: *p*-values from $10^{-300}$ to 1, from $10^{-55}$ to 1, and from $3.16\times10^{-6}$ to 1. The space is divided into rectangular bins (10 intervals on the x-axis and 27 or 30 on the y-axis) colored by density of individual *qpAdm* models populating this space (logarithmic color scale). Distributions (probability density) of ideal models defined as those with ST distance = 0 are also shown on the logarithmic and non-logarithmic *p*-value axes. The green horizontal lines mark the lowest and highest *p*-value thresholds used in this study (0.001 and 0.5).

**Figure S22.**

**a**

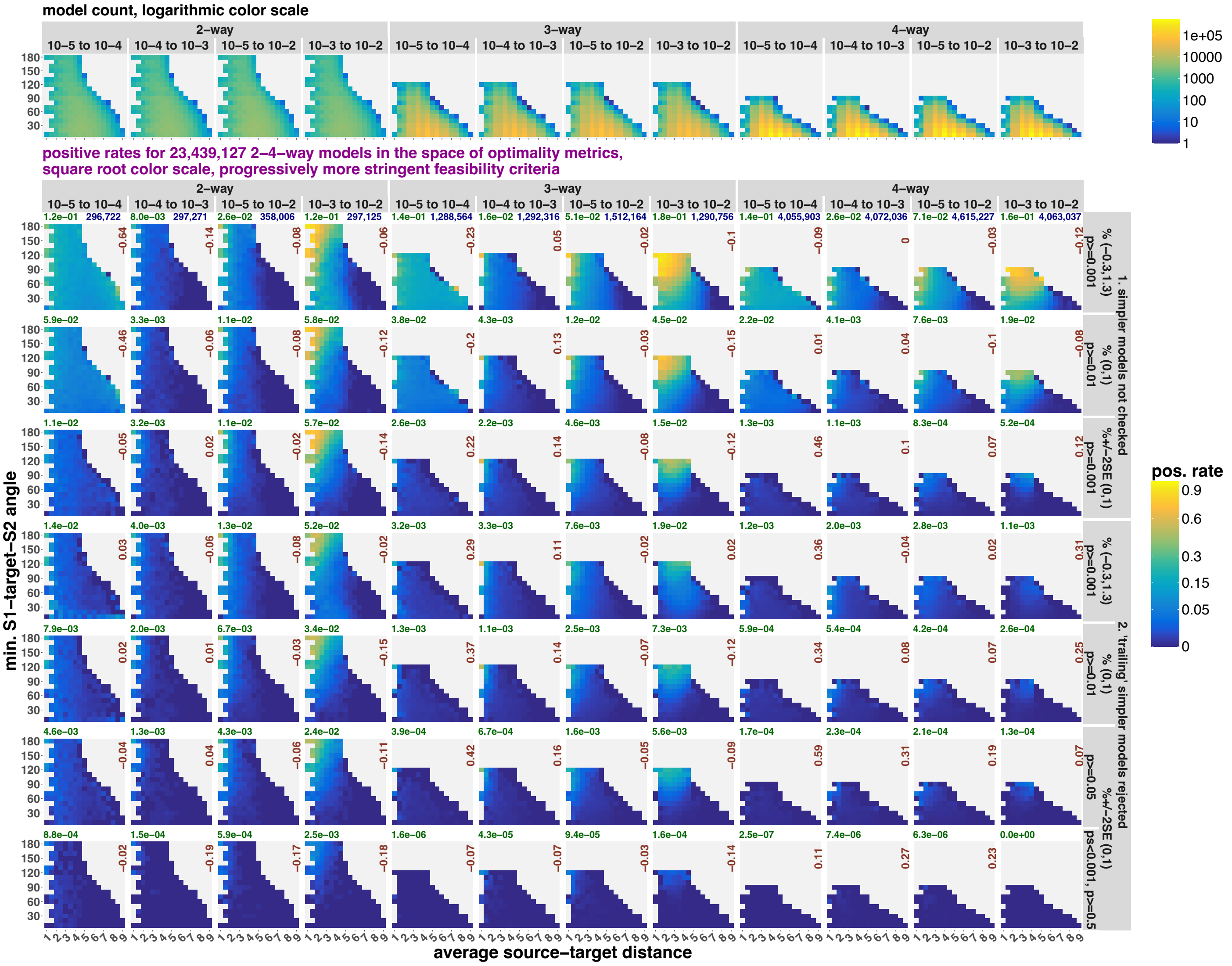

model count, logarithmic color scale
2–way
3–way
4–way
10–5 to 10–4
10–4 to 10–3
10–5 to 10–2
10–3 to 10–2
positive rates for 23,439,127 2–4–way models in the space of optimality metrics,
square root color scale, progressively more stringent feasibility criteria
min. S1–target–S2 angle
average source–target distance
pos. rate
1. simpler models not checked
2. 'trailing' simpler models rejected

**b**

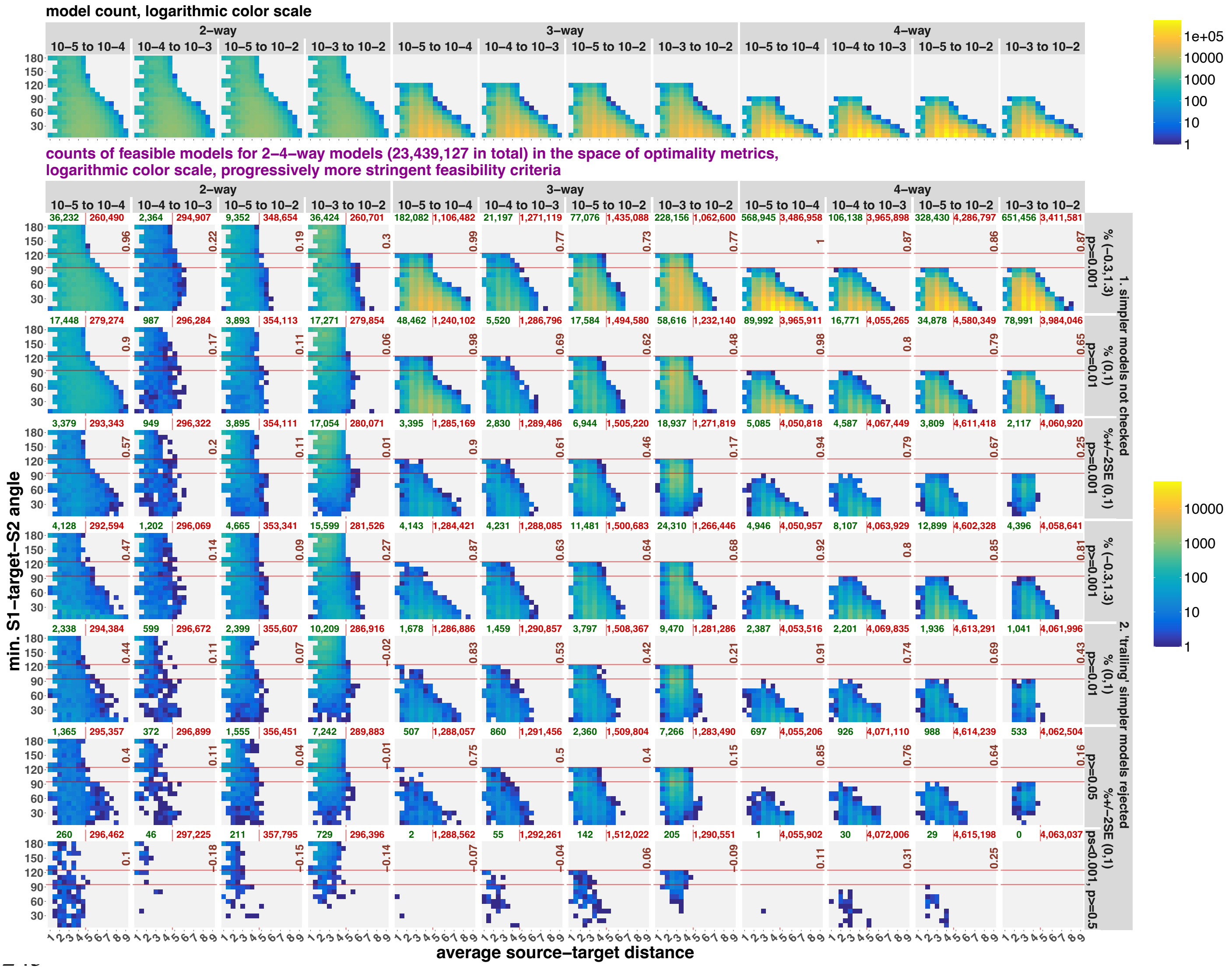

model count, logarithmic color scale
2–way
3–way
4–way
10–5 to 10–4
10–4 to 10–3
10–5 to 10–2
10–3 to 10–2
counts of feasible models for 2–4–way models (23,439,127 in total) in the space of optimality metrics,
logarithmic color scale, progressively more stringent feasibility criteria
min. S1–target–S2 angle
average source–target distance
1. simpler models not checked
2. 'trailing' simpler models rejected
% (−0.3,1.3) p>=0.001
% (0,1) p>=0.01
%+/−2SE (0,1) p>=0.001
% (−0.3,1.3) p>=0.001
% (0,1) p>=0.01
%+/−2SE (0,1) p>=0.05
ps<0.001, p>=0.5

**c**

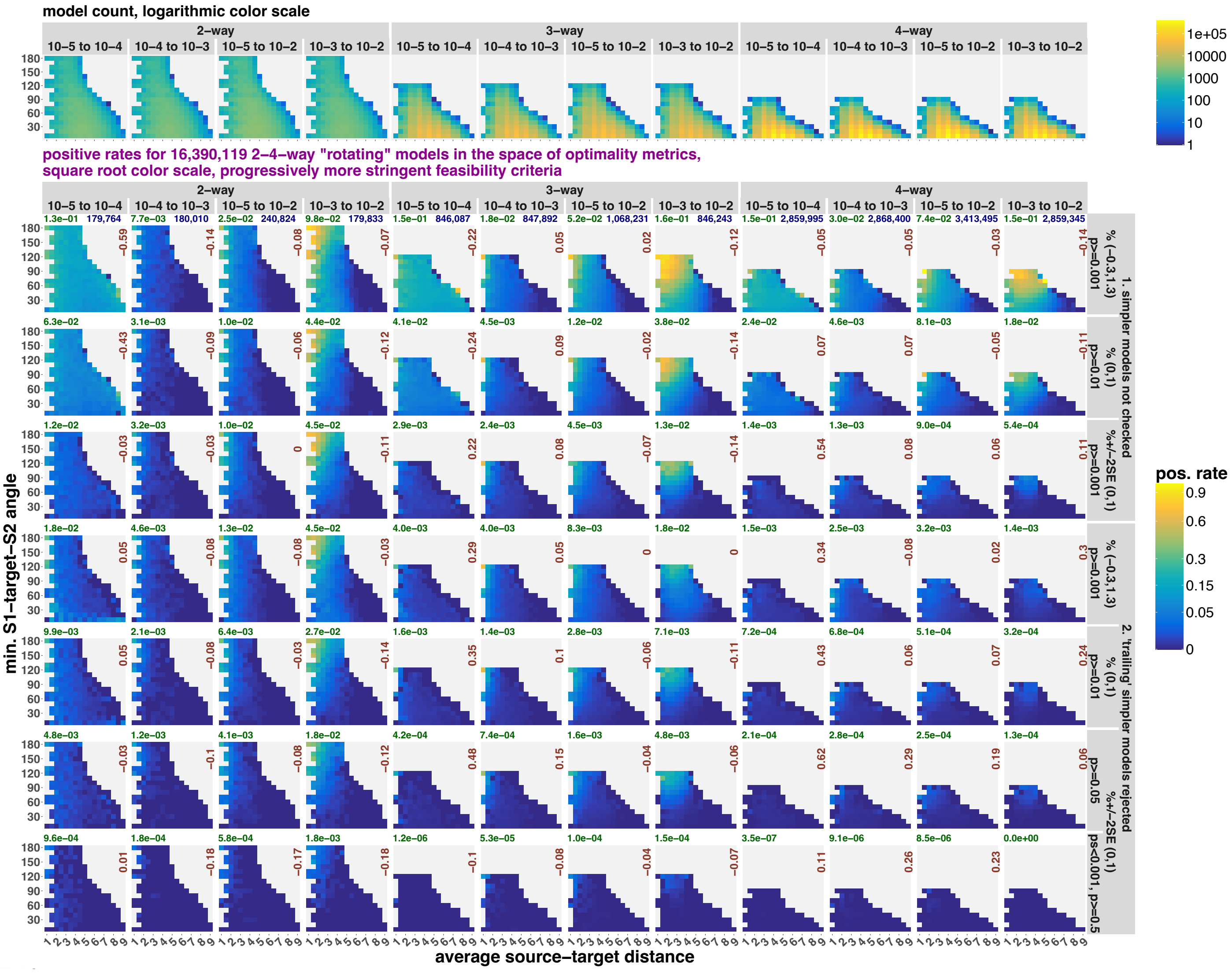

model count, logarithmic color scale
2–way
3–way
4–way
10–5 to 10–4
10–4 to 10–3
10–5 to 10–2
10–3 to 10–2
1e+05
10000
1000
100
10
1
positive rates for 16,390,119 2–4–way "rotating" models in the space of optimality metrics,
square root color scale, progressively more stringent feasibility criteria
pos. rate
0.9
0.6
0.3
0.15
0.05
0
min. S1–target–S2 angle
average source–target distance
1. simpler models not checked
2. 'trailing' simpler models rejected
% (−0.3,1.3) p>=0.001
% (0,1) p>=0.01
%+/−2SE (0,1) p>=0.001
% (0,1) p>=0.05
%+/−2SE (0,1) p>=0.001, p>=0.5
179,764
180,010
240,824
179,833
846,087
847,892
1,068,231
846,243
2,859,995
2,868,400
3,413,495
2,859,345

**d**

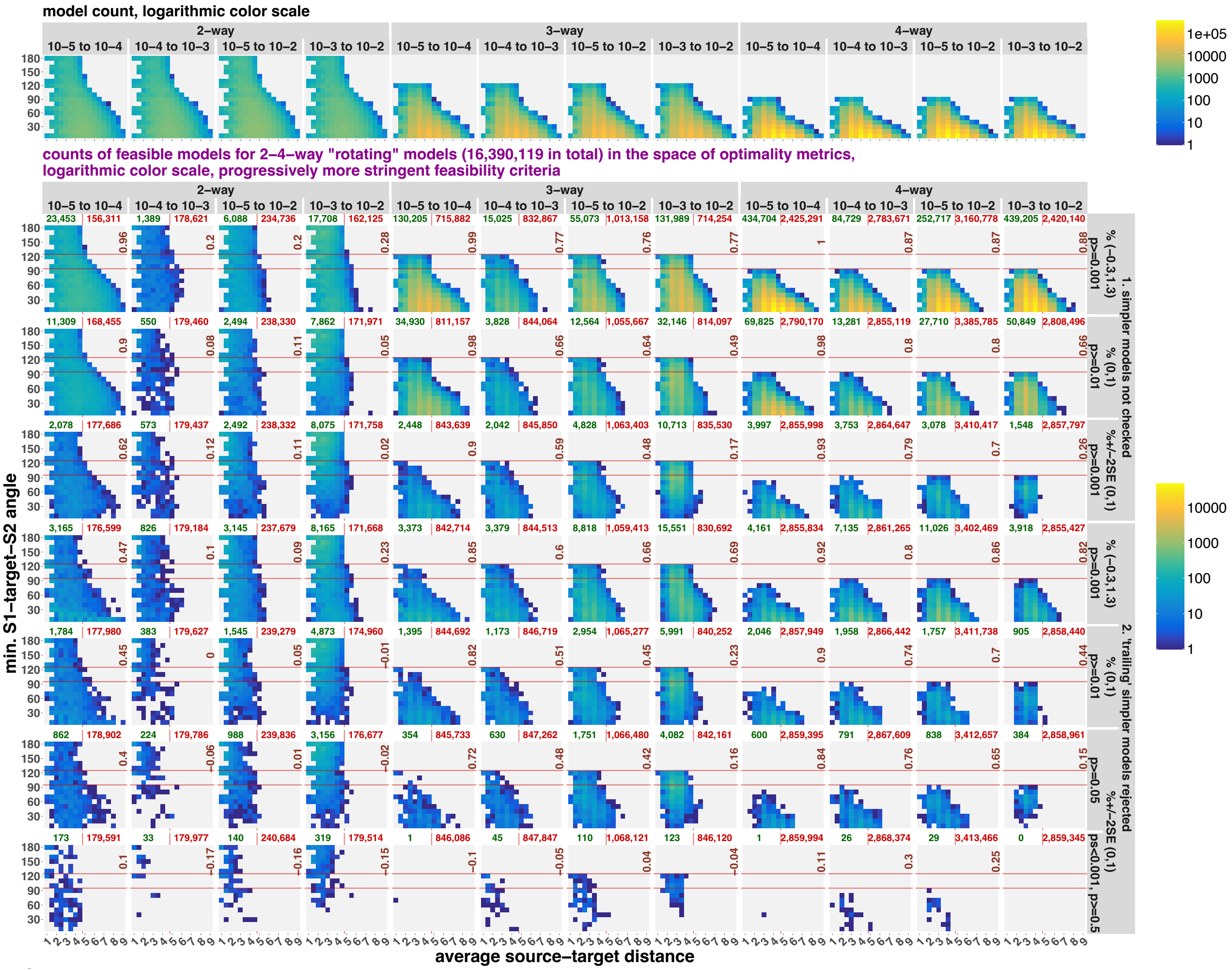

model count, logarithmic color scale
2–way
3–way
4–way
10–5 to 10–4
10–4 to 10–3
10–5 to 10–2
10–3 to 10–2
counts of feasible models for 2–4–way "rotating" models (16,390,119 in total) in the space of optimality metrics, logarithmic color scale, progressively more stringent feasibility criteria
min. S1–target–S2 angle
average source–target distance
1. simpler models not checked
2. 'trailing' simpler models rejected
% (−0.3,1.3) p>=0.001
% (0,1) p>=0.01
%+/−2SE (0,1)
p>=0.05
ps<0.001, p>0.5

**e**

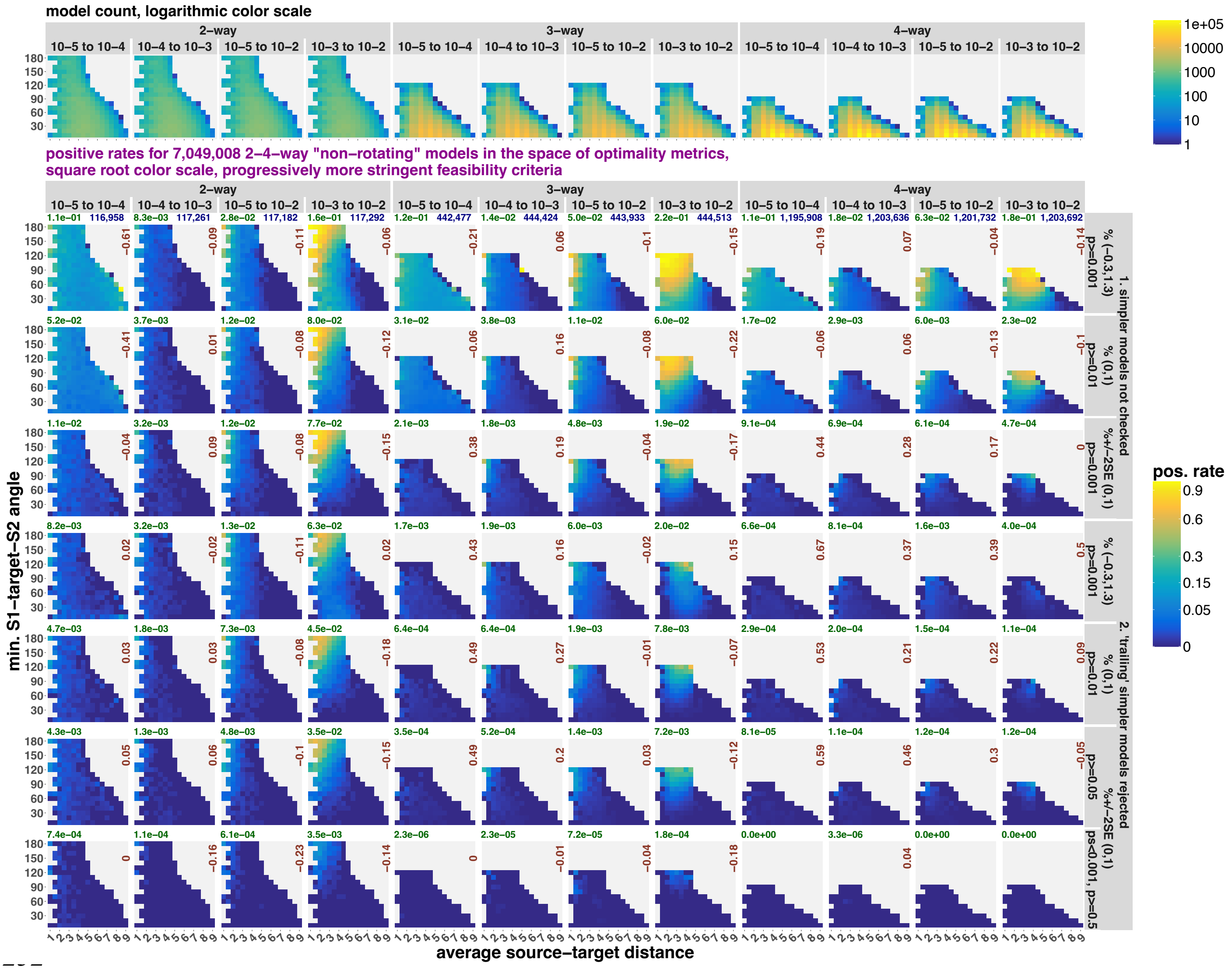

model count, logarithmic color scale
2–way
3–way
4–way
10–5 to 10–4
10–4 to 10–3
10–5 to 10–2
10–3 to 10–2
positive rates for 7,049,008 2–4–way "non–rotating" models in the space of optimality metrics,
square root color scale, progressively more stringent feasibility criteria
min. S1–target–S2 angle
average source–target distance
pos. rate
1. simpler models not checked
2. 'trailing' simpler models rejected
% (−0.3,1.3) p>=0.001
% (0,1) p>=0.01
%+/−2SE (0,1) p>=0.001
% (0,1) p>=0.05
%+/−2SE (0,1) ps<0.001, p>=0.5

**f**

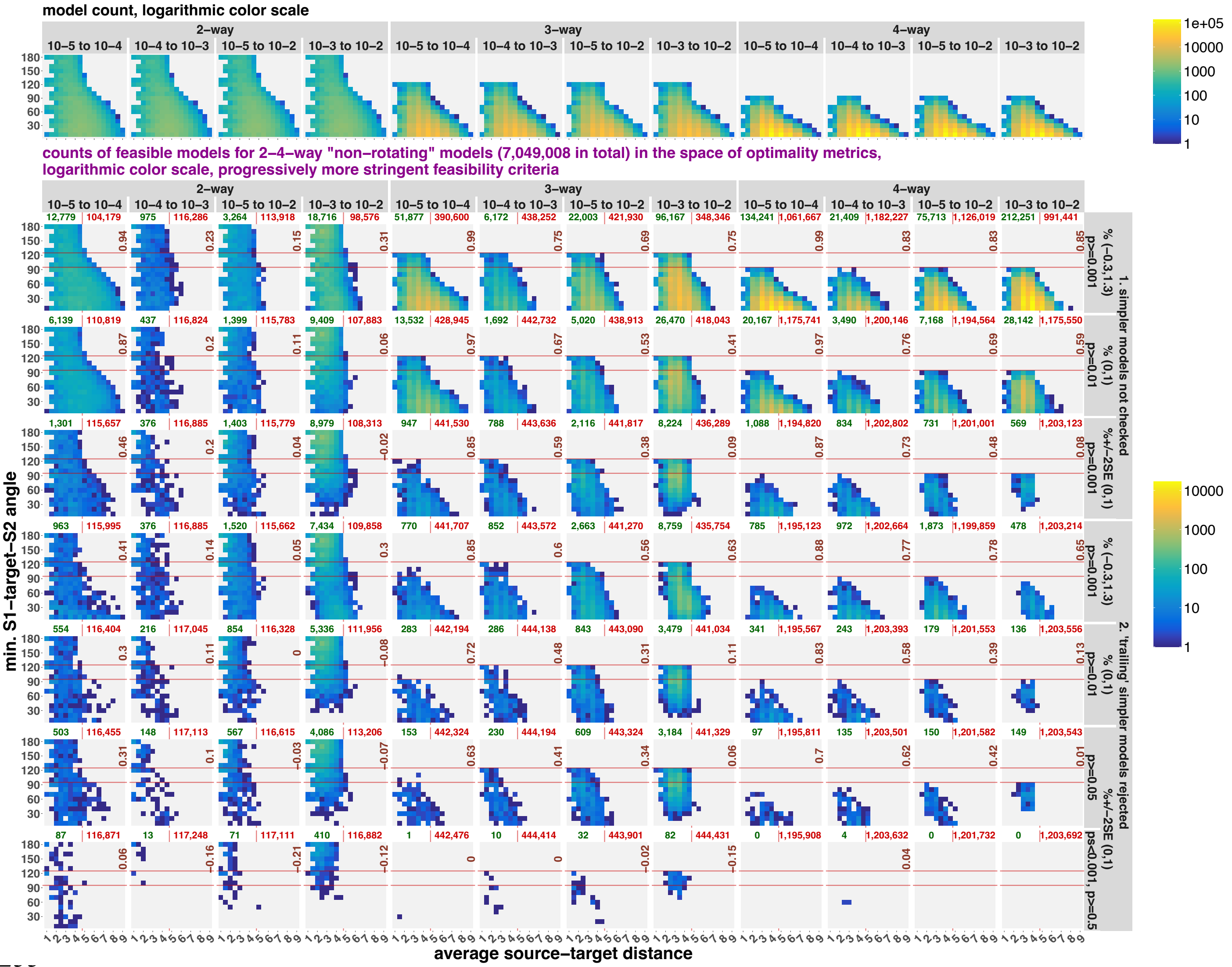


Randomized two to four-way *qpAdm* models in the space of model optimality metrics: average ST distance and min. STS angle (models with the latter metric undefined were excluded from this analysis). Results are either combined for all *qpAdm* protocols (panels **a, b**) or shown separately for rotating (**c, d**) and non-rotating protocols (**e, f**). In all cases, the results are stratified by model complexity and landscape type. The space was divided into 18 bins on both the x-axis (bin width = 0.5) and y-axis (intervals [0°, 10°], (10°, 20°], and so forth). The upper row of plots shows density of all randomized models in those two-dimensional bins, and the other rows show either *qpAdm* positive rates (fractions of *qpAdm* models satisfying the selected feasibility criteria; panels **a, c, e**) or density of fitting models (**b, d, f**). The selected feasibility criteria are listed in the captions on the right (7 of 36 criteria tested in this study, from the least stringent on top to the most stringent at the bottom). Logarithmic color scales are used for showing density of individual *qpAdm* models populating the space, and square root color scales for *qpAdm* positive rate. Spearman's correlation coefficients for counts of all models vs. counts of fitting models or *qpAdm* positive rates in the two-dimensional bins are shown in each panel in brown (bins not populated by any models were not considered). The red horizontal lines mark the "ideal" min. STS angles for three- and four-way models (120° and 90°), and the red vertical lines mark the average ST distance (4.5) that equals the radius of the landscape: a boundary separating "highly asymmetric" models to the right from more optimal models to the left. The numbers above the plots show either overall *qpAdm* positive rates (in green) and total counts of models tested in blue (**a, b, d**), or total counts of fitting and non-fitting models in green and red, respectively (**c, e**).

**Figure S23.**

**a**

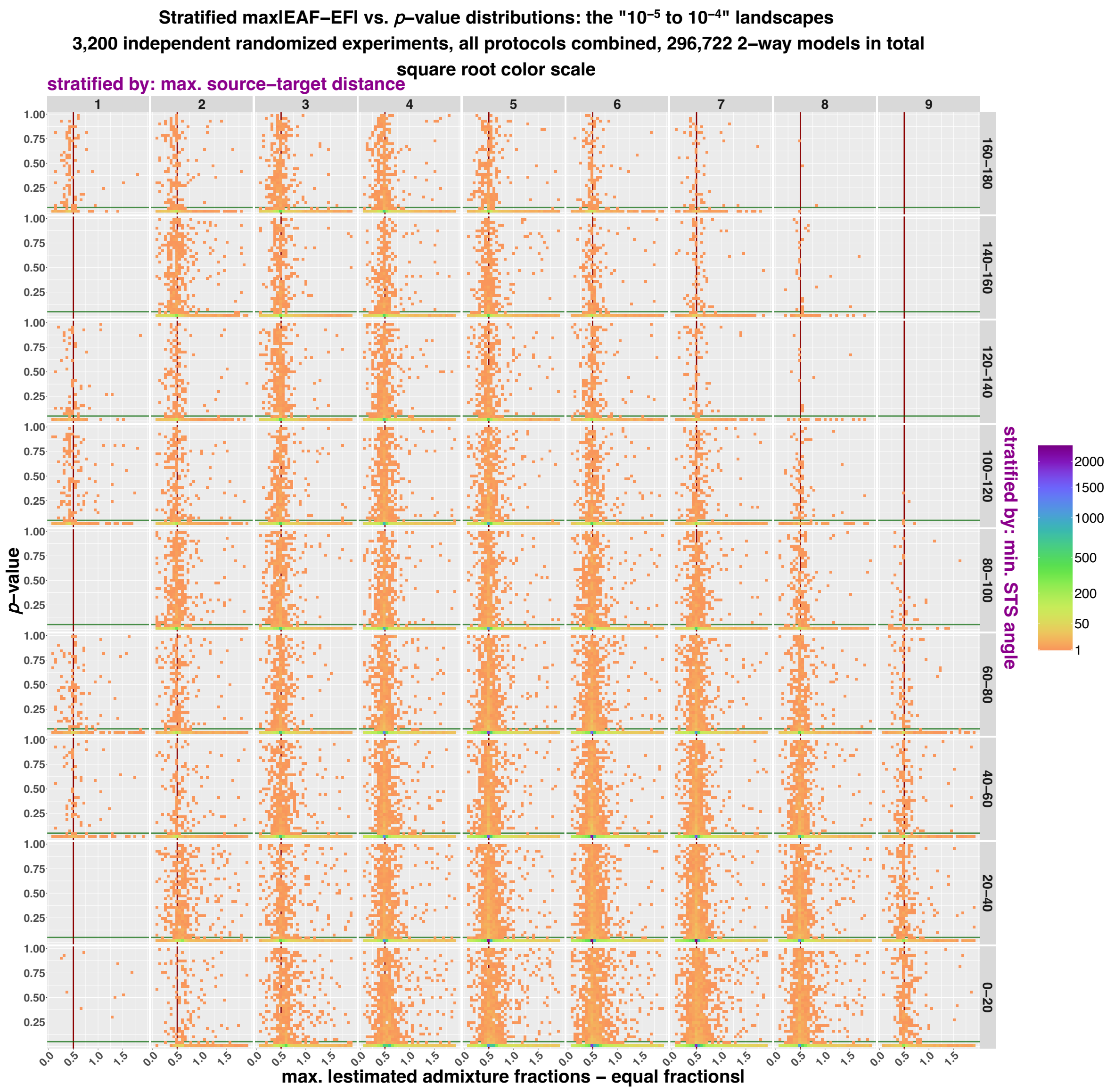

Stratified max|EAF−EF| vs. p−value distributions: the "10⁻⁵ to 10⁻⁴" landscapes
3,200 independent randomized experiments, all protocols combined, 296,722 2−way models in total
square root color scale
stratified by: max. source−target distance
1
2
3
4
5
6
7
8
9
160−180
140−160
120−140
100−120
80−100
60−80
40−60
20−40
0−20
stratified by: min. STS angle
p−value
1.00
0.75
0.50
0.25
0.0
0.5
1.0
1.5
max. |estimated admixture fractions − equal fractions|
2000
1500
1000
500
200
50
1

**b**

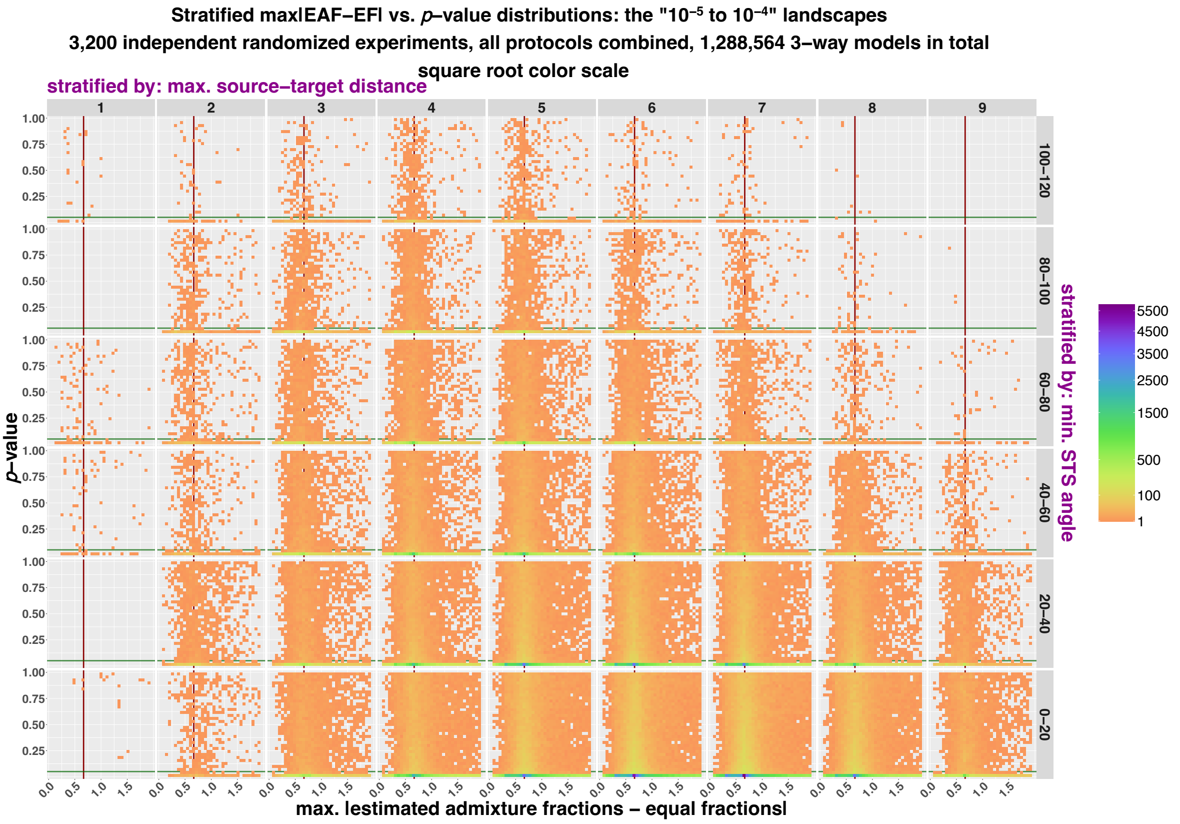
Stratified max|EAF−EF| vs. p–value distributions: the "10⁻⁵ to 10⁻⁴" landscapes
3,200 independent randomized experiments, all protocols combined, 1,288,564 3–way models in total
square root color scale
stratified by: max. source–target distance
1 2 3 4 5 6 7 8 9
100–120
80–100
60–80
40–60
20–40
0–20
stratified by: min. STS angle
p–value
max. |estimated admixture fractions – equal fractions|
5500
4500
3500
2500
1500
500
100
1

**c**

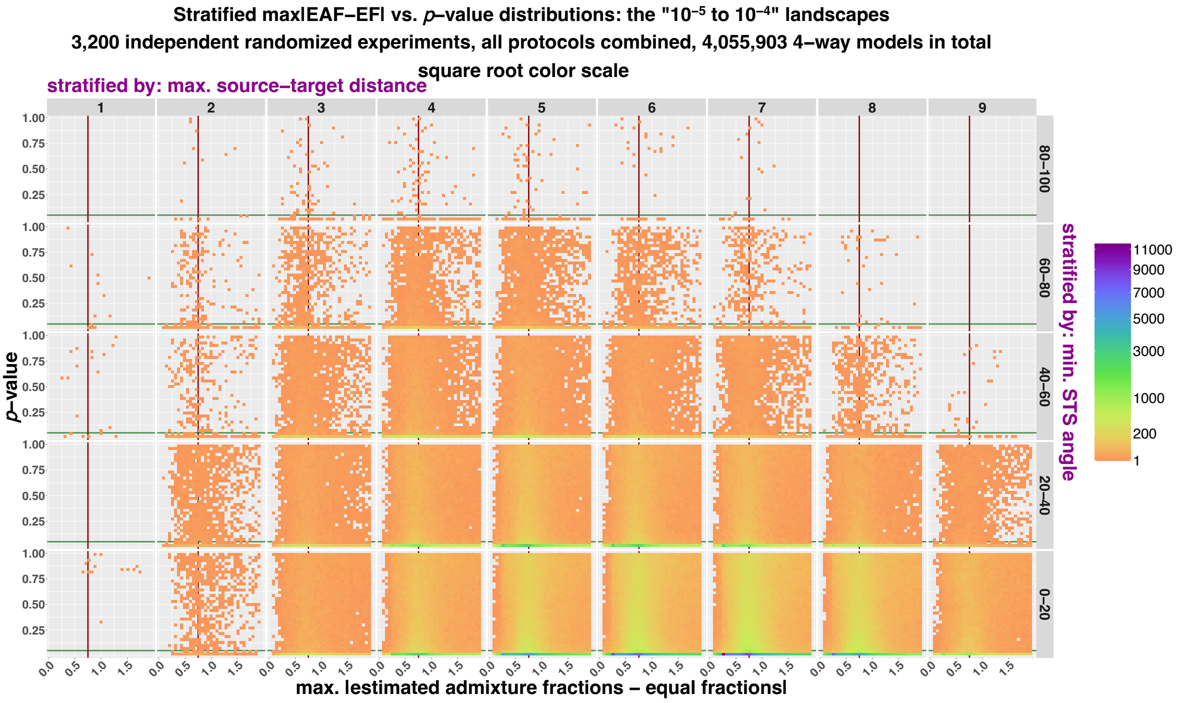
Stratified max|EAF−EF| vs. p–value distributions: the "10⁻⁵ to 10⁻⁴" landscapes
3,200 independent randomized experiments, all protocols combined, 4,055,903 4–way models in total
square root color scale
stratified by: max. source–target distance
1 2 3 4 5 6 7 8 9
80–100
60–80
40–60
20–40
0–20
stratified by: min. STS angle
p–value
max. |estimated admixture fractions – equal fractions|
11000
9000
7000
5000
3000
1000
200
1

**d**

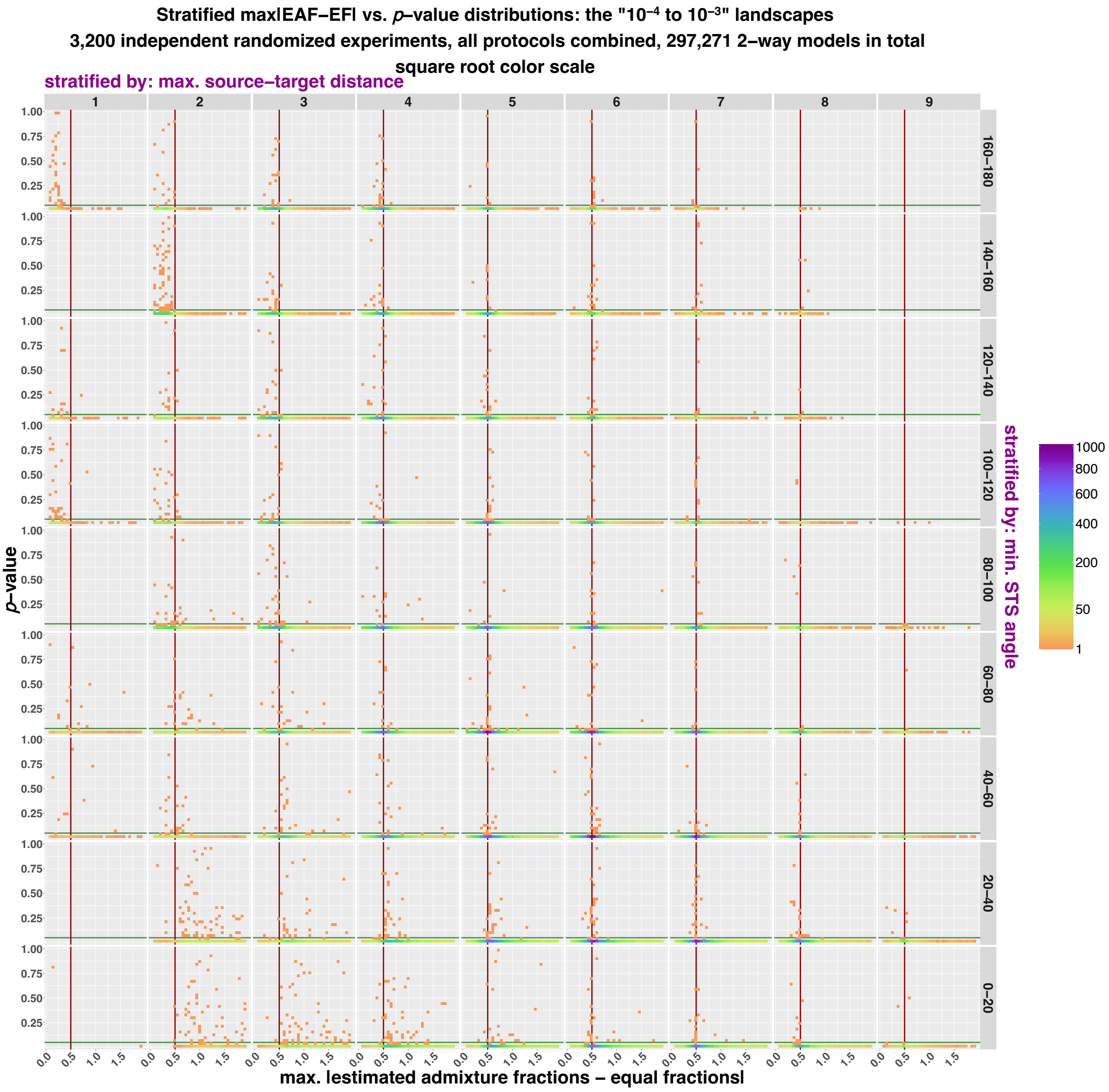
Stratified max|EAF−EF| vs. p−value distributions: the "10⁻⁴ to 10⁻³" landscapes
3,200 independent randomized experiments, all protocols combined, 297,271 2−way models in total
square root color scale
stratified by: max. source−target distance
1
2
3
4
5
6
7
8
9
stratified by: min. STS angle
160−180
140−160
120−140
100−120
80−100
60−80
40−60
20−40
0−20
p−value
1.00
0.75
0.50
0.25
0.0
0.5
1.0
1.5
max. |estimated admixture fractions − equal fractions|
1000
800
600
400
200
50
1

**e**

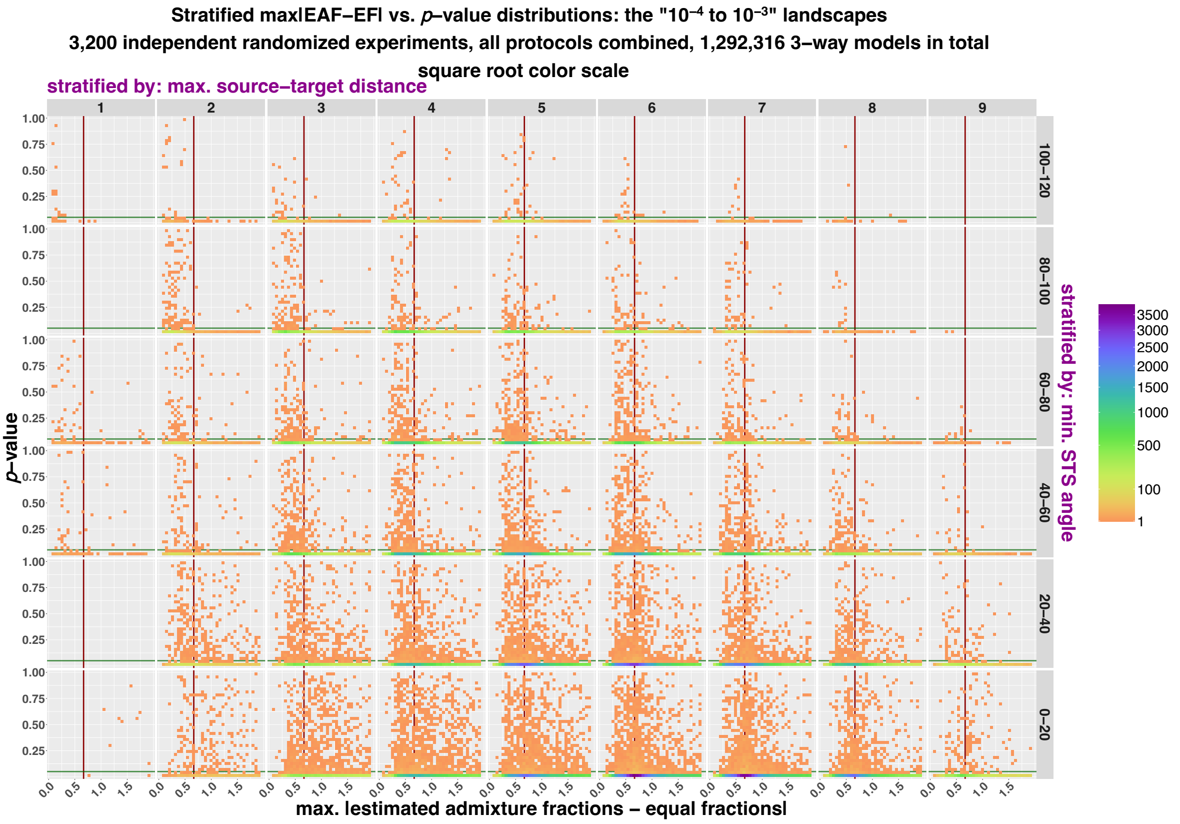
Stratified max|EAF−EF| vs. p–value distributions: the "10⁻⁴ to 10⁻³" landscapes
3,200 independent randomized experiments, all protocols combined, 1,292,316 3–way models in total
square root color scale
stratified by: max. source–target distance
1 2 3 4 5 6 7 8 9
100–120
80–100
60–80
40–60
20–40
0–20
stratified by: min. STS angle
p–value
max. |estimated admixture fractions – equal fractions|
3500 3000 2500 2000 1500 1000 500 100 1

**f**

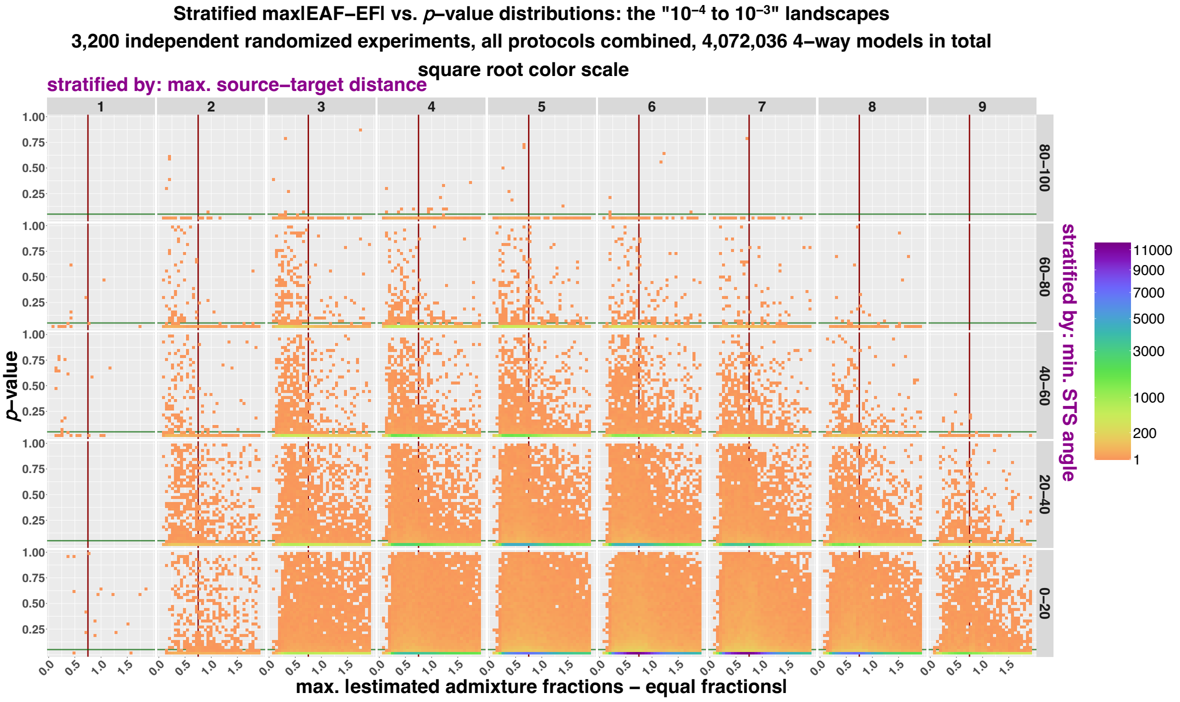
Stratified max|EAF−EF| vs. p–value distributions: the "10⁻⁴ to 10⁻³" landscapes
3,200 independent randomized experiments, all protocols combined, 4,072,036 4–way models in total
square root color scale
stratified by: max. source–target distance
1 2 3 4 5 6 7 8 9
80–100
60–80
40–60
20–40
0–20
stratified by: min. STS angle
p–value
max. |estimated admixture fractions – equal fractions|
11000 9000 7000 5000 3000 1000 200 1

**g**

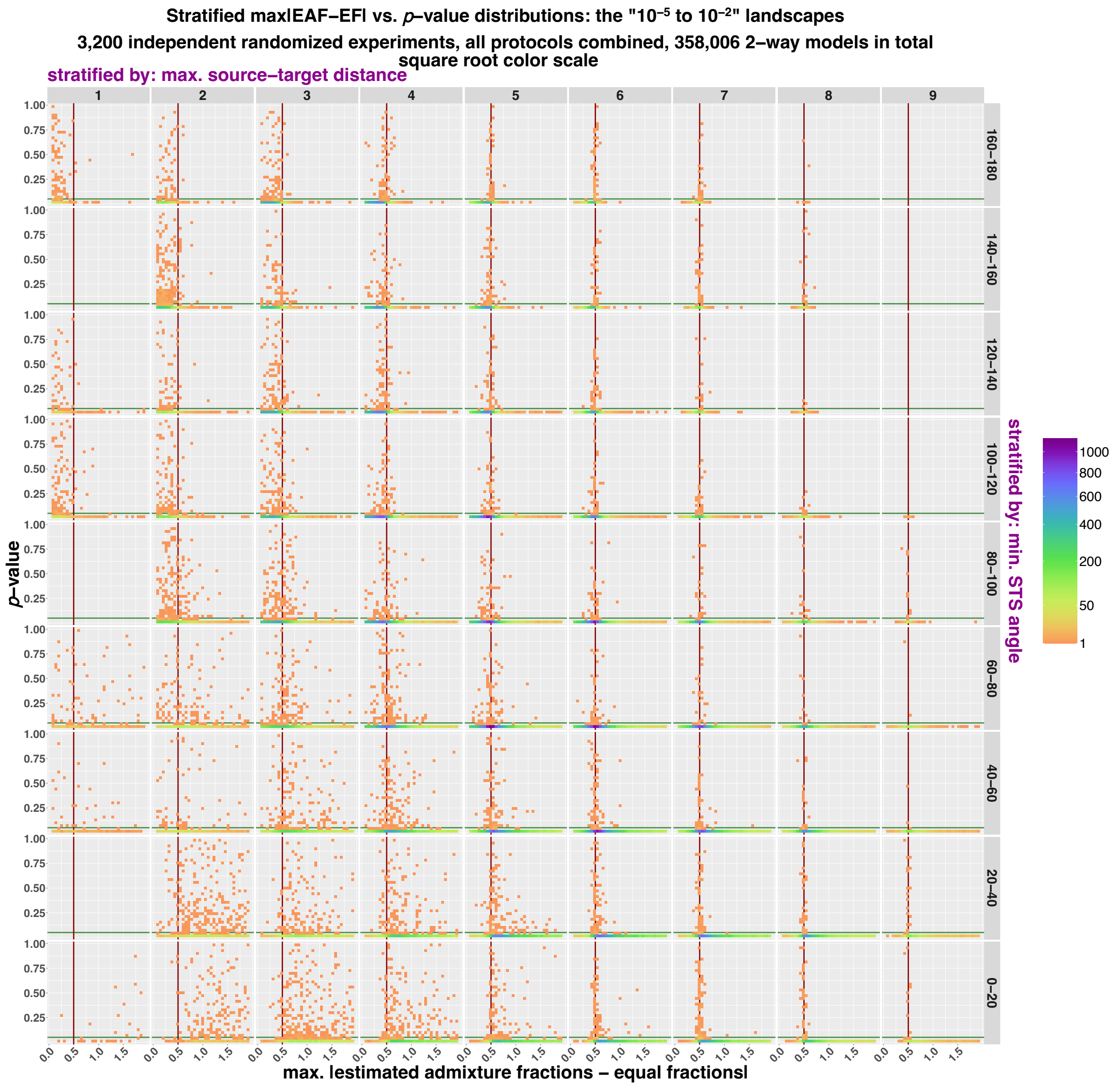
Stratified max|EAF−EF| vs. p–value distributions: the "10⁻⁵ to 10⁻²" landscapes
3,200 independent randomized experiments, all protocols combined, 358,006 2–way models in total
square root color scale
stratified by: max. source–target distance
1
2
3
4
5
6
7
8
9
160–180
140–160
120–140
100–120
80–100
60–80
40–60
20–40
0–20
stratified by: min. STS angle
p–value
1.00
0.75
0.50
0.25
0.0
0.5
1.0
1.5
max. |estimated admixture fractions – equal fractions|
1000
800
600
400
200
50
1

**h**

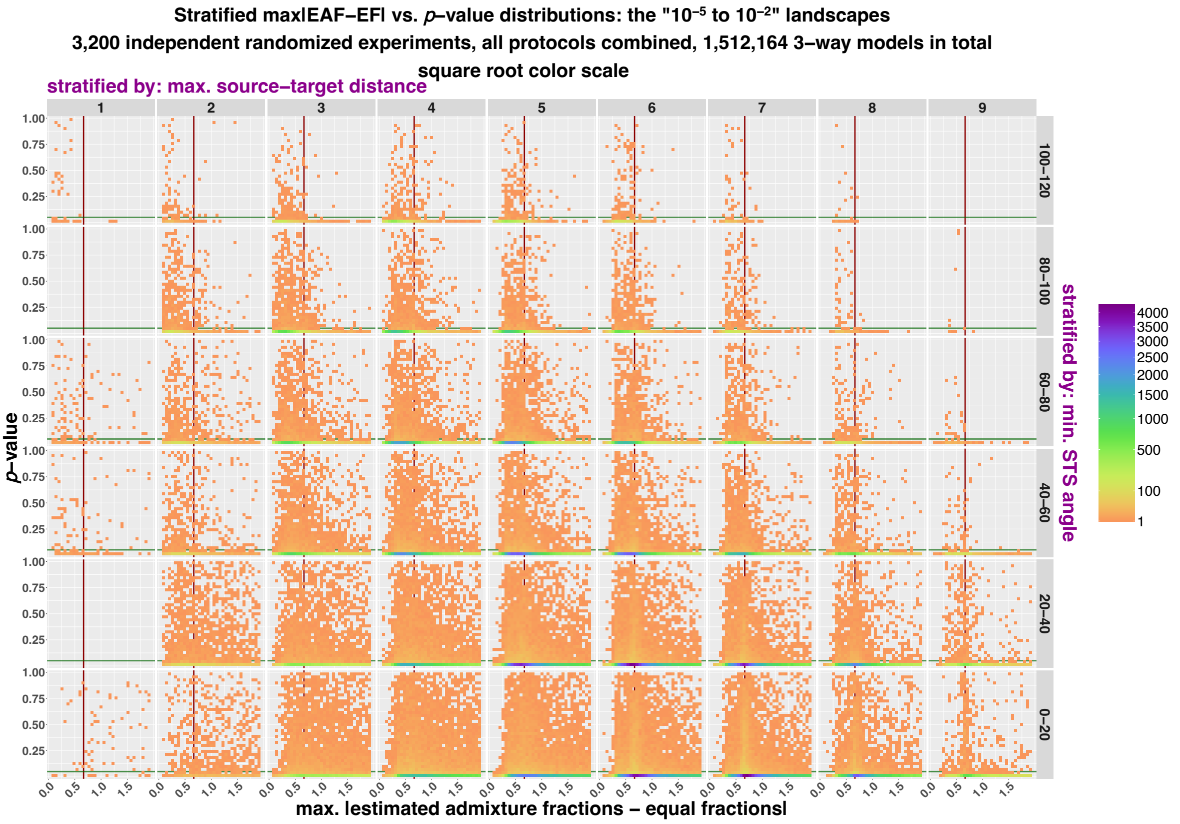
Stratified max|EAF−EF| vs. p–value distributions: the "10⁻⁵ to 10⁻²" landscapes
3,200 independent randomized experiments, all protocols combined, 1,512,164 3–way models in total
square root color scale
stratified by: max. source–target distance
1
2
3
4
5
6
7
8
9
100–120
80–100
60–80
40–60
20–40
0–20
stratified by: min. STS angle
p–value
1.00
0.75
0.50
0.25
0.0
0.5
1.0
1.5
max. |estimated admixture fractions – equal fractions|
4000
3500
3000
2500
2000
1500
1000
500
100
1

**i**

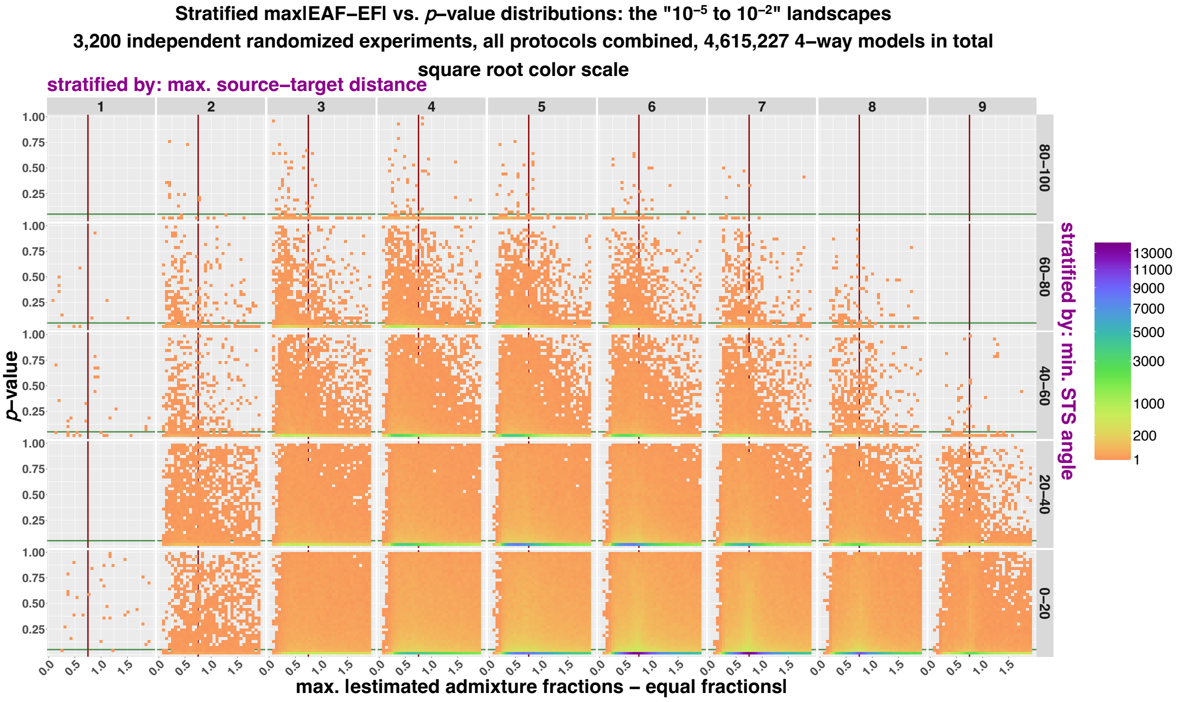
Stratified max|EAF−EF| vs. p–value distributions: the "10⁻⁵ to 10⁻²" landscapes
3,200 independent randomized experiments, all protocols combined, 4,615,227 4–way models in total
square root color scale
stratified by: max. source–target distance
1
2
3
4
5
6
7
8
9
80–100
60–80
40–60
20–40
0–20
stratified by: min. STS angle
p–value
1.00
0.75
0.50
0.25
0.0
0.5
1.0
1.5
max. |estimated admixture fractions – equal fractions|
13000
11000
9000
7000
5000
3000
1000
200
1

**j**

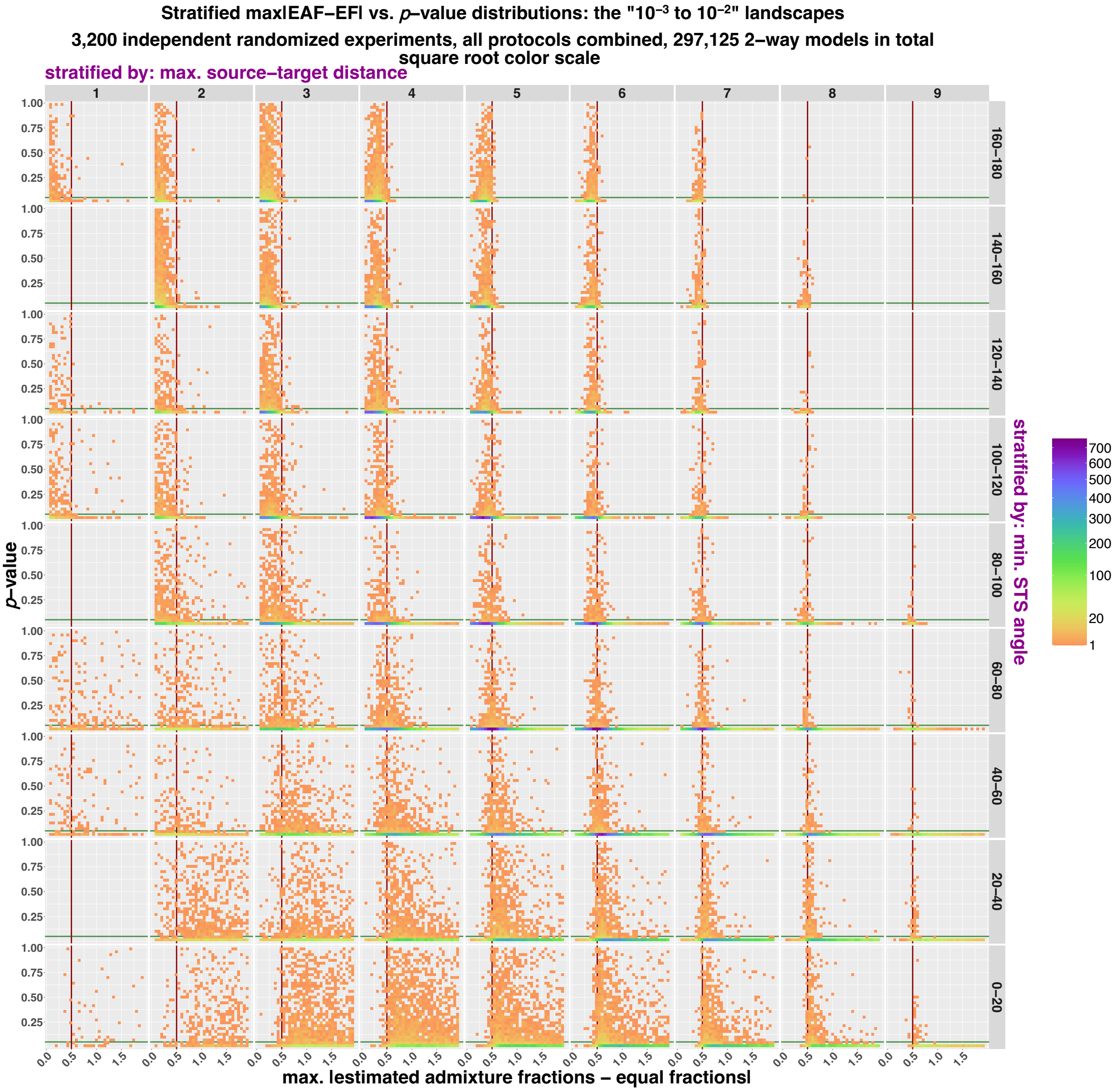
Stratified max|EAF–EF| vs. p–value distributions: the "10⁻³ to 10⁻²" landscapes
3,200 independent randomized experiments, all protocols combined, 297,125 2–way models in total
square root color scale
stratified by: max. source–target distance
1
2
3
4
5
6
7
8
9
160–180
140–160
120–140
100–120
80–100
60–80
40–60
20–40
0–20
stratified by: min. STS angle
p–value
1.00
0.75
0.50
0.25
0.0
0.5
1.0
1.5
max. |estimated admixture fractions – equal fractions|
700
600
500
400
300
200
100
20
1

**k**

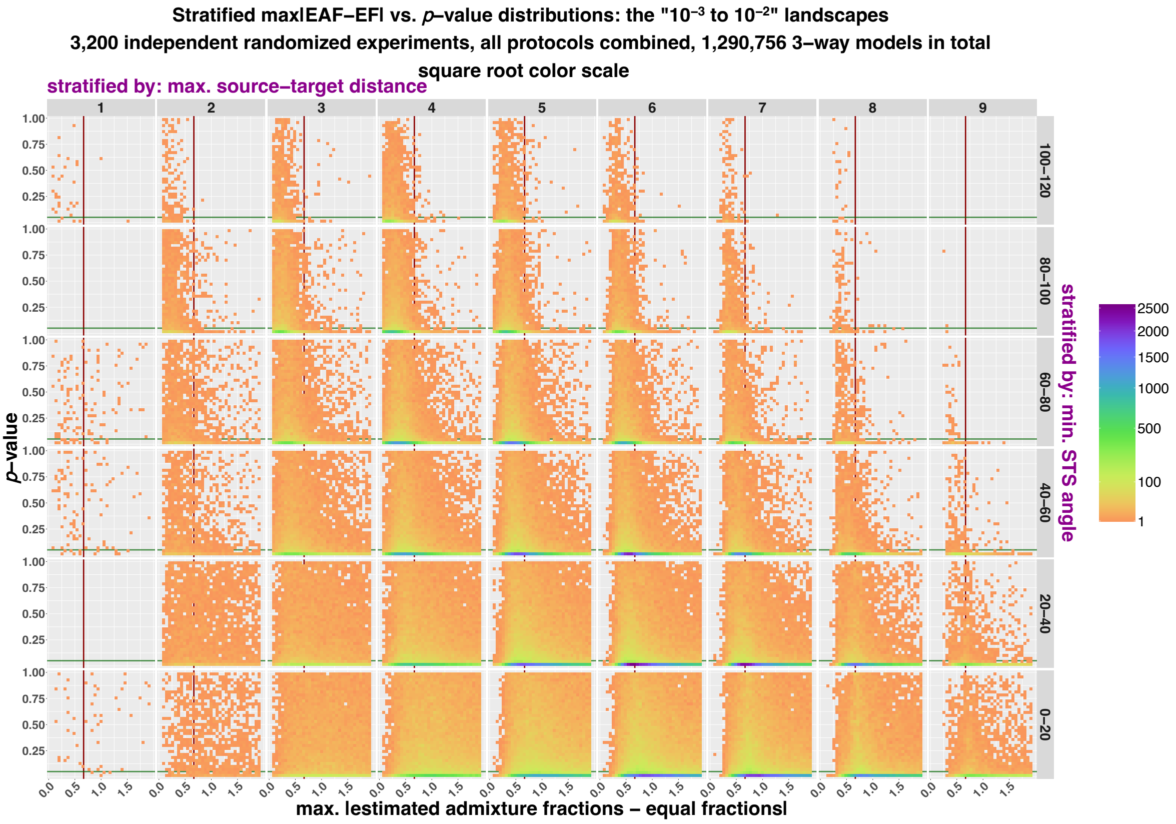


**l**

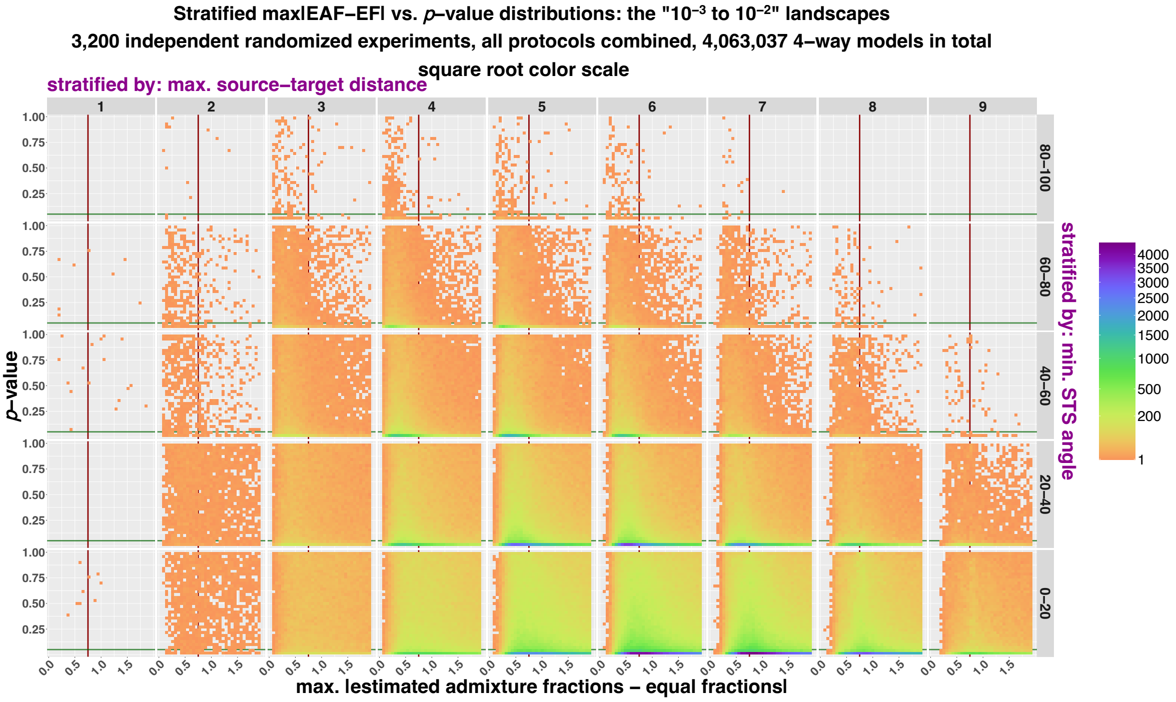


Distributions of randomized two (**a, d, g, j**), three- (**b, e, h, k**), and four-way (**c, f, i, l**) *qpAdm* models in the space of max|*EAF–EF*| and *p*-values (max|*EAF–EF*| <2; both axes are non-logarithmic). Results are shown for the "$10^{-5}$ to $10^{-4}$" (**a-c**), "$10^{-4}$ to $10^{-3}$" (**d-f**), "$10^{-5}$ to $10^{-2}$" (**g-i**), and "$10^{-3}$ to $10^{-2}$" (**j-l**) landscapes, for all the *qpAdm* protocols combined, and are stratified by binned model optimality metrics: max. ST distance (in relative units) and min. STS angle (in degrees). Models with the latter metric undefined were excluded from this analysis. The space is divided into 35×35 rectangular bins, and they are colored by density of individual *qpAdm* models populating this space (square-root color scale). The red vertical lines (or the red tick marks) mark max$|EAF–EF|$ = 1 – $EF$; for three-way and more complex models this boundary is similar but not identical to the boundary between EAF within (0, 1) and outside (0, 1). The green horizontal lines mark a $p$-value threshold = 0.05.

**Figure S24.**

**a**

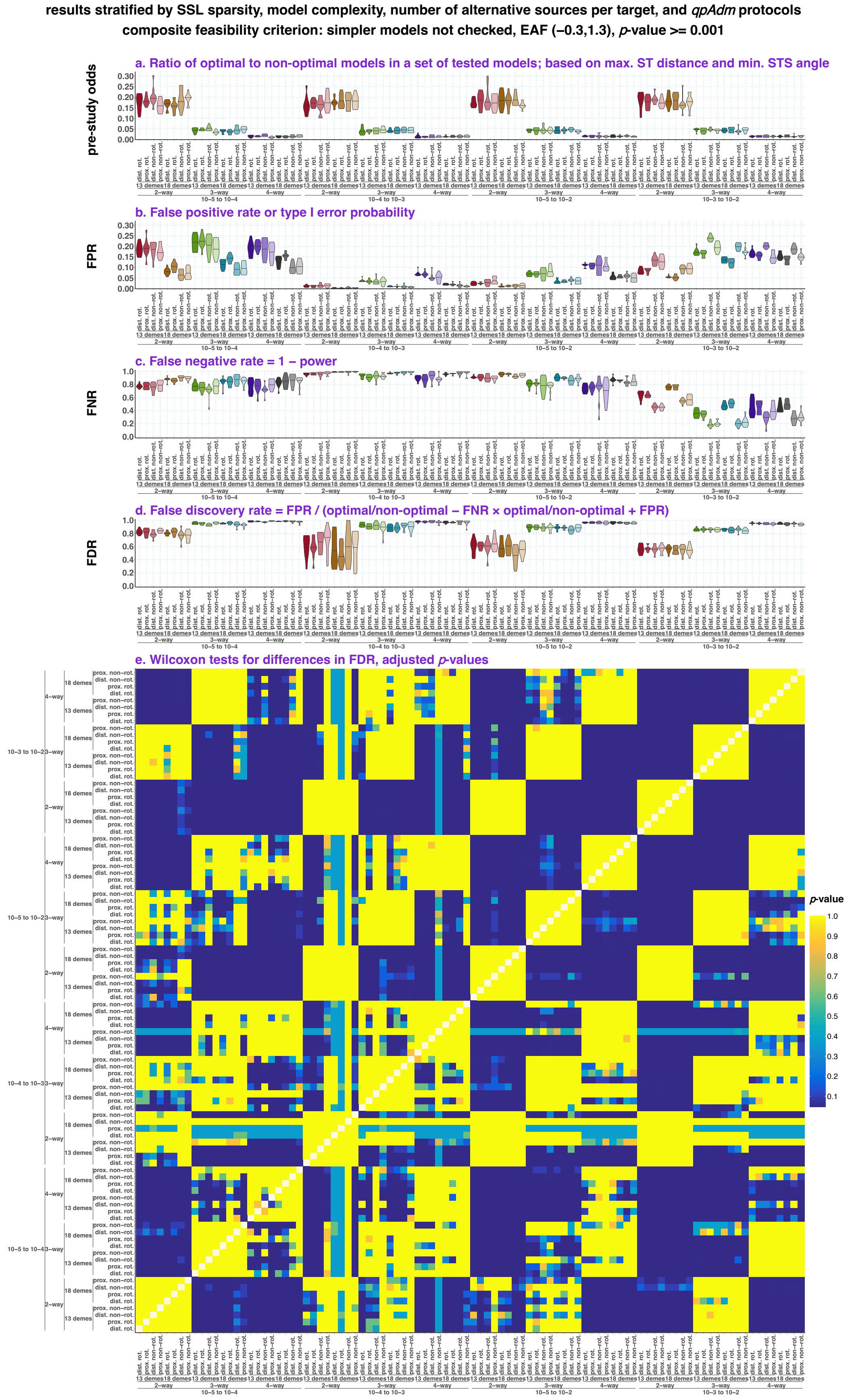

results stratified by SSL sparsity, model complexity, number of alternative sources per target, and qpAdm protocols
composite feasibility criterion: simpler models not checked, EAF (−0.3,1.3), p-value >= 0.001
a. Ratio of optimal to non-optimal models in a set of tested models; based on max. ST distance and min. STS angle
pre-study odds
b. False positive rate or type I error probability
FPR
c. False negative rate = 1 − power
FNR
d. False discovery rate = FPR / (optimal/non-optimal − FNR × optimal/non-optimal + FPR)
FDR
e. Wilcoxon tests for differences in FDR, adjusted p-values
p-value
dist. rot.
prox. rot.
dist. non-rot.
prox. non-rot.
13 demes
18 demes
2-way
3-way
4-way
10−5 to 10−4
10−4 to 10−3
10−5 to 10−2
10−3 to 10−2
10−5 to 10−4

**b**

**results stratified by SSL sparsity, model complexity, number of alternative sources per target, and *qpAdm* protocols**
**composite feasibility criterion: simpler models not checked, EAF (0,1), *p*-value >= 0.01**

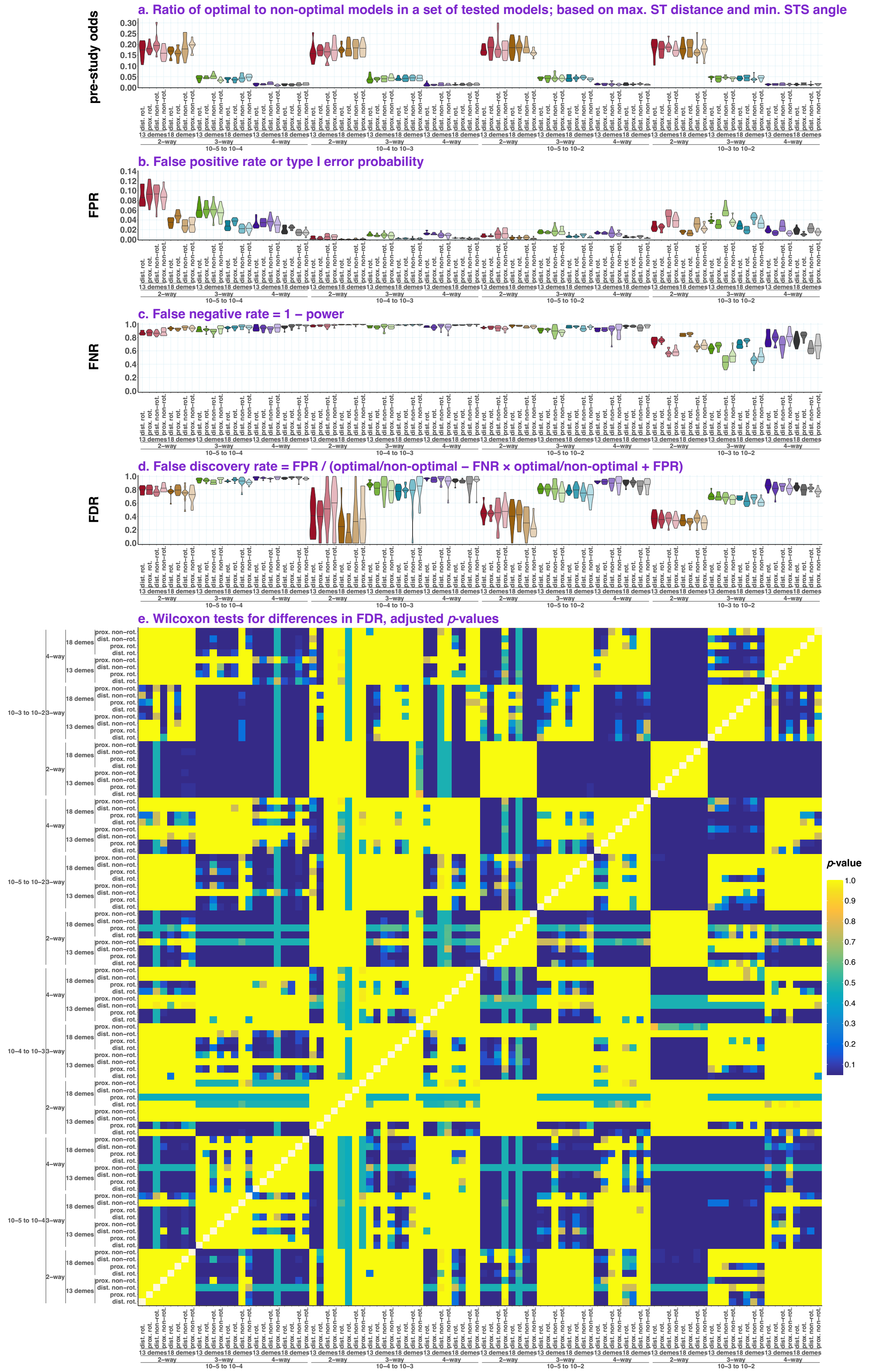

**c**

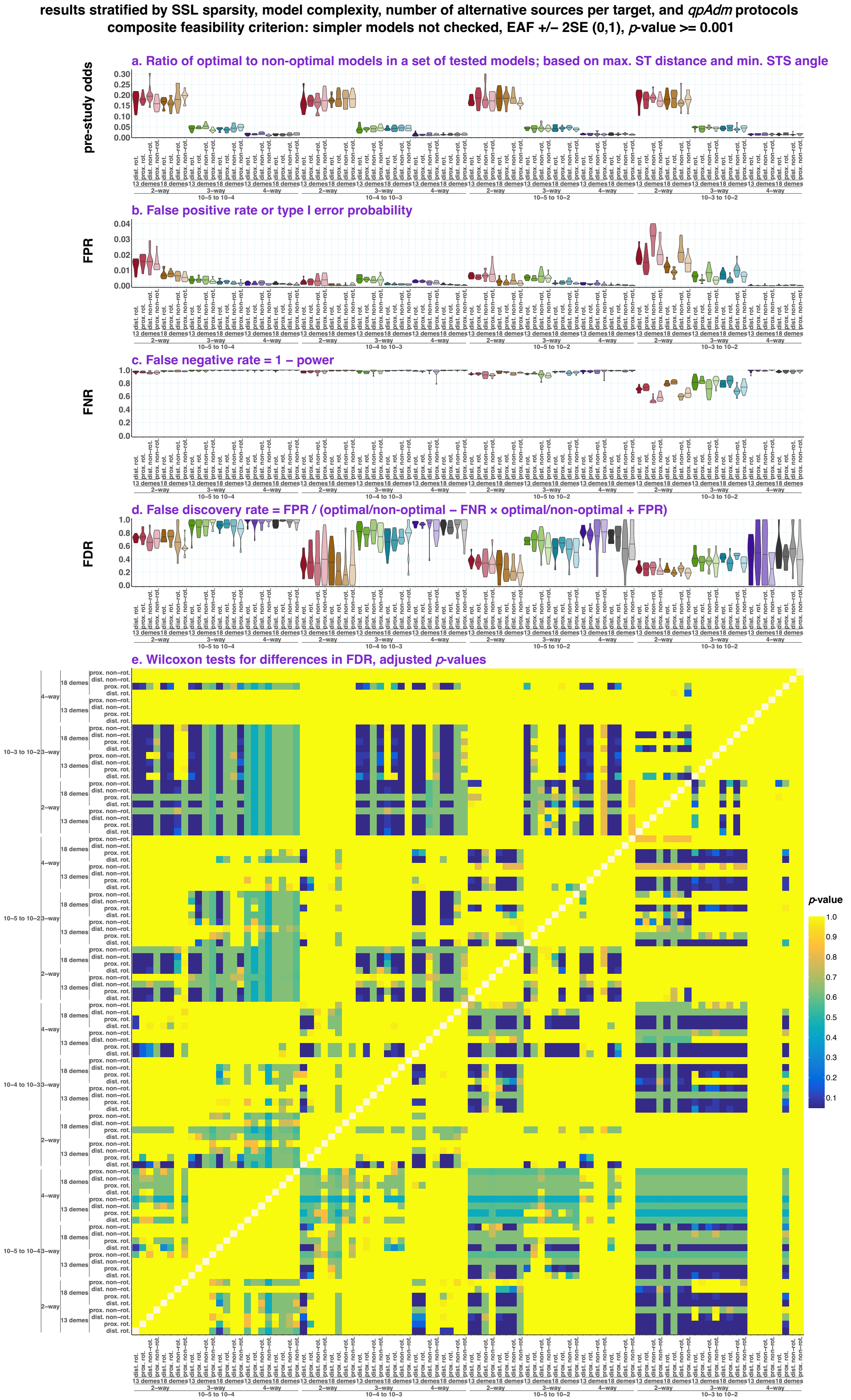
results stratified by SSL sparsity, model complexity, number of alternative sources per target, and qpAdm protocols
composite feasibility criterion: simpler models not checked, EAF +/– 2SE (0,1), p-value >= 0.001
a. Ratio of optimal to non-optimal models in a set of tested models; based on max. ST distance and min. STS angle
pre-study odds
b. False positive rate or type I error probability
FPR
c. False negative rate = 1 – power
FNR
d. False discovery rate = FPR / (optimal/non-optimal – FNR × optimal/non-optimal + FPR)
FDR
e. Wilcoxon tests for differences in FDR, adjusted p-values
p-value
10−5 to 10−4
10−4 to 10−3
10−5 to 10−2
10−3 to 10−2
2−way
3−way
4−way
13 demes
18 demes
dist. rot.
prox. rot.
dist. non−rot.
prox. non−rot.

**d**

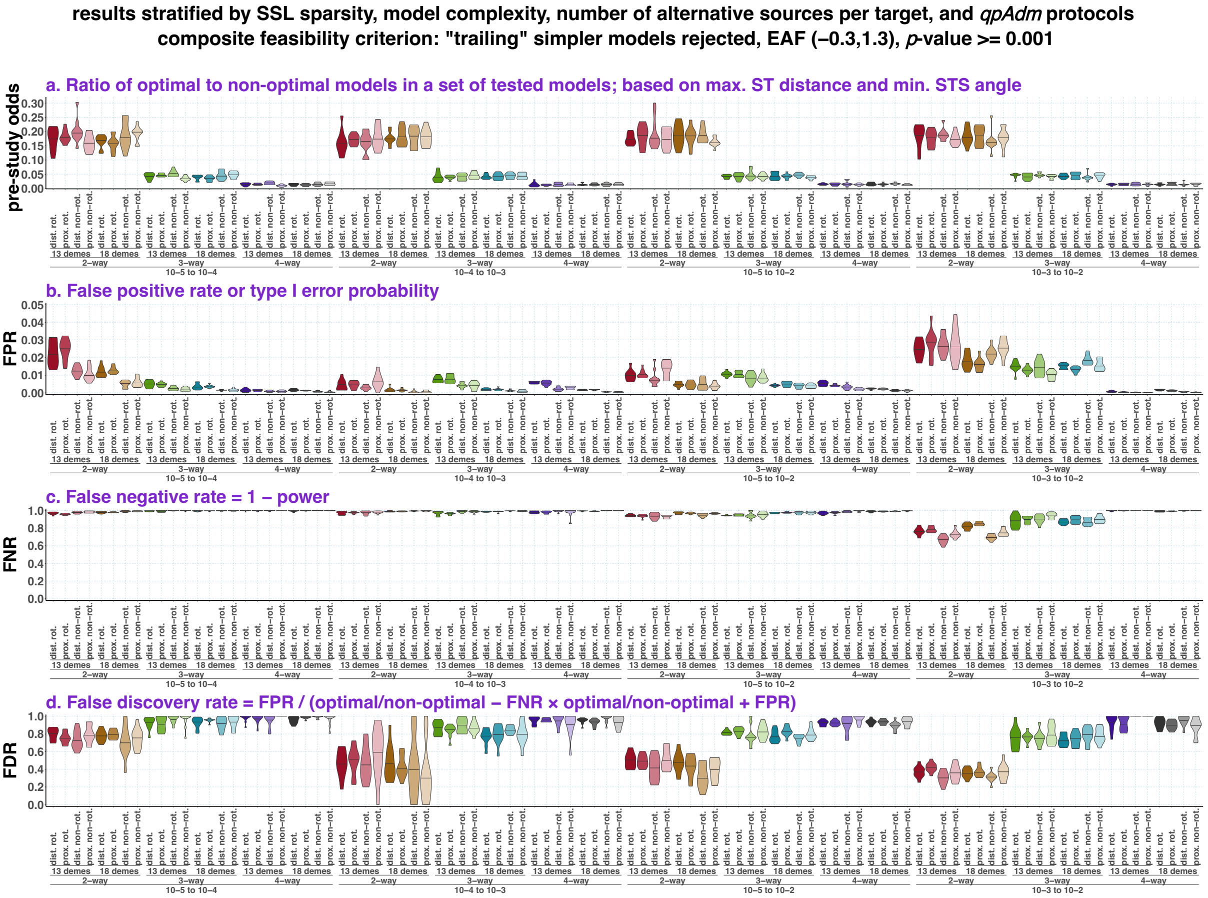

results stratified by SSL sparsity, model complexity, number of alternative sources per target, and *qpAdm* protocols
composite feasibility criterion: "trailing" simpler models rejected, EAF (−0.3,1.3), *p*-value >= 0.001
a. Ratio of optimal to non-optimal models in a set of tested models; based on max. ST distance and min. STS angle
pre-study odds
b. False positive rate or type I error probability
FPR
c. False negative rate = 1 – power
FNR
d. False discovery rate = FPR / (optimal/non-optimal – FNR × optimal/non-optimal + FPR)
FDR
dist. rot.
prox. rot.
dist. non-rot.
prox. non-rot.
13 demes
18 demes
2-way
3-way
4-way
10–5 to 10–4
10–4 to 10–3
10–5 to 10–2
10–3 to 10–2

**e**

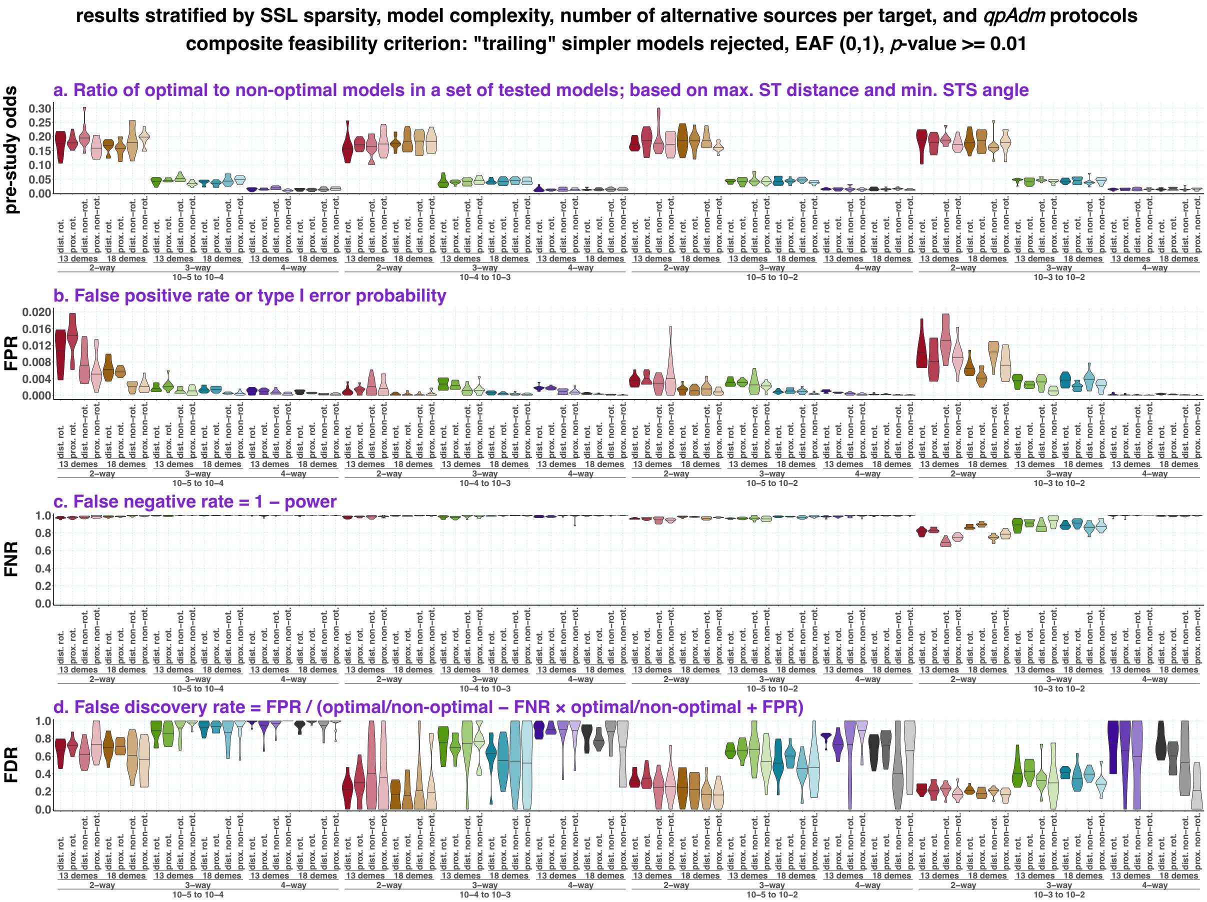
results stratified by SSL sparsity, model complexity, number of alternative sources per target, and qpAdm protocols
composite feasibility criterion: "trailing" simpler models rejected, EAF (0,1), p-value >= 0.01
a. Ratio of optimal to non-optimal models in a set of tested models; based on max. ST distance and min. STS angle
pre-study odds
b. False positive rate or type I error probability
FPR
c. False negative rate = 1 – power
FNR
d. False discovery rate = FPR / (optimal/non-optimal – FNR × optimal/non-optimal + FPR)
FDR
dist. rot.
prox. rot.
dist. non-rot.
prox. non-rot.
13 demes
18 demes
2–way
3–way
4–way
10–5 to 10–4
10–4 to 10–3
10–5 to 10–2
10–3 to 10–2

**f**

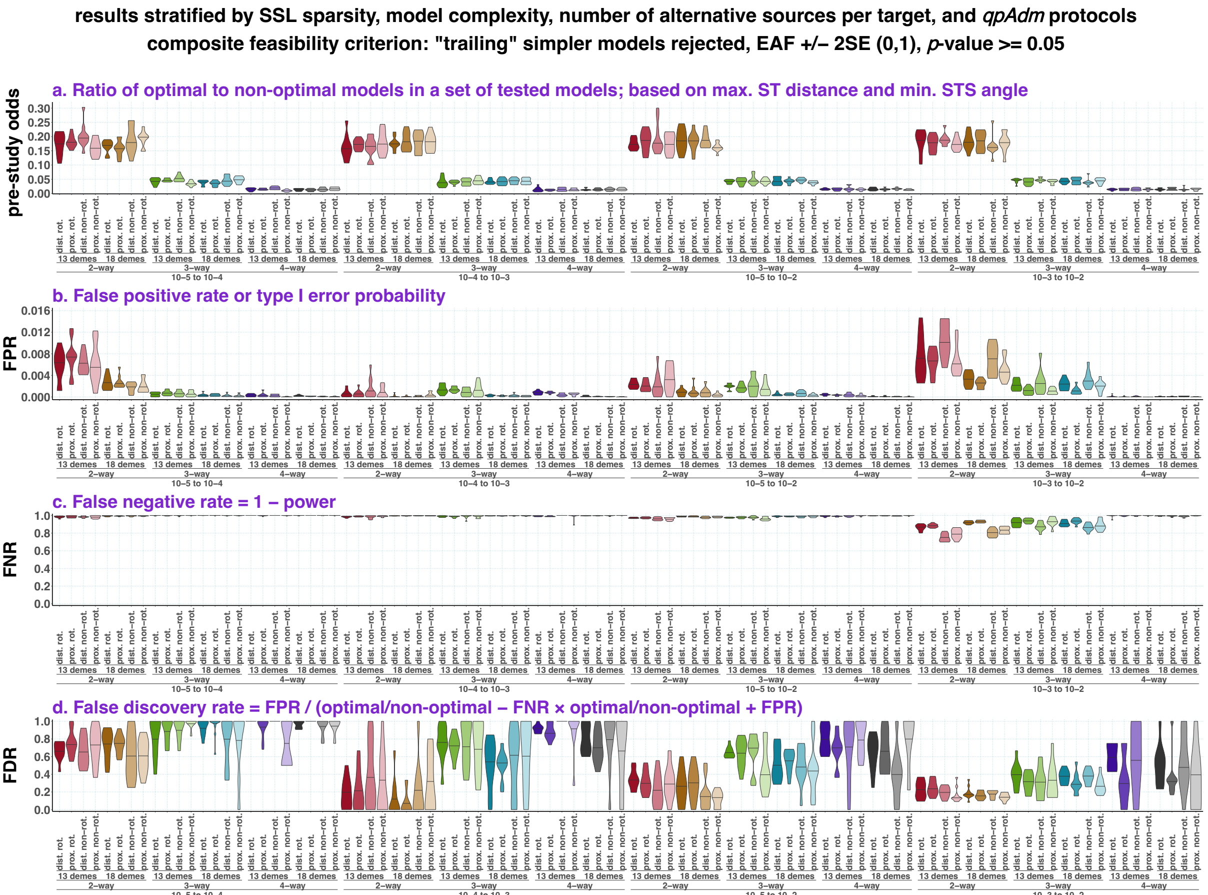


Pre-study odds, FPR, FNR, and FDR are shown for six composite model feasibility criteria (**a-f**). The violin plots with medians visualize distributions of these metrics across simulation replicates. The results are grouped by SSL sparsity, model complexity, the number of alternative sources per target (13 or 18 in the rotating protocols, corresponding to 10 and 15 in the non-rotating ones), and by *qpAdm* protocols. Results of pairwise comparisons of the FDR distributions are shown in the matrix form in panels **a-c** (*p*-values of non-paired Wilcoxon tests were adjusted for multiple tests using the Holm method). These statistical tests were not performed for some feasibility criteria due to the presence of missing FDR values.

**Figure S25.**

**a**

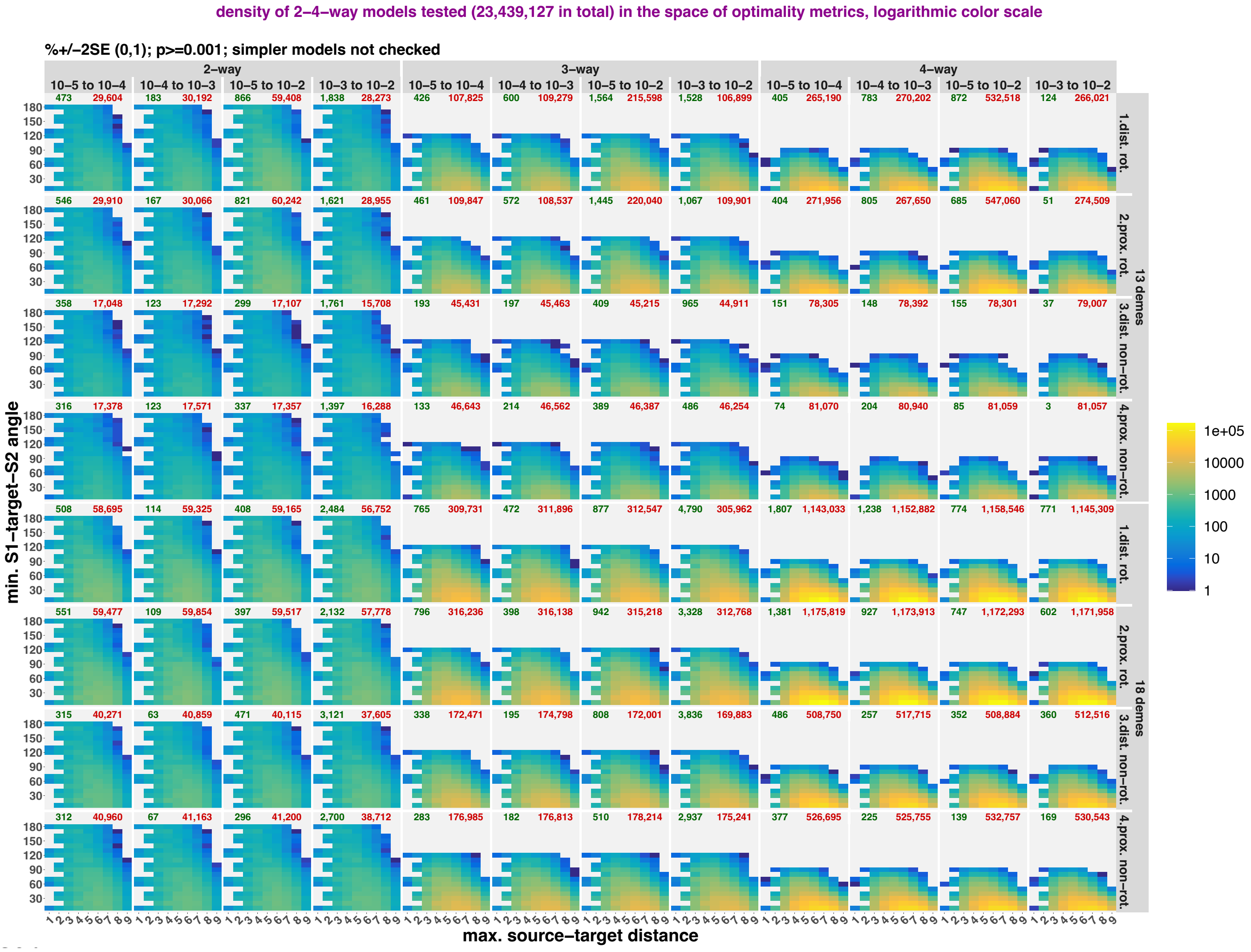

density of 2–4–way models tested (23,439,127 in total) in the space of optimality metrics, logarithmic color scale
%+/−2SE (0,1); p>=0.001; simpler models not checked
2–way
3–way
4–way
10–5 to 10–4
10–4 to 10–3
10–5 to 10–2
10–3 to 10–2
473 29,604
183 30,192
866 59,408
1,838 28,273
426 107,825
600 109,279
1,564 215,598
1,528 106,899
405 265,190
783 270,202
872 532,518
124 266,021
546 29,910
167 30,066
821 60,242
1,621 28,955
461 109,847
572 108,537
1,445 220,040
1,067 109,901
404 271,956
805 267,650
685 547,060
51 274,509
358 17,048
123 17,292
299 17,107
1,761 15,708
193 45,431
197 45,463
409 45,215
965 44,911
151 78,305
148 78,392
155 78,301
37 79,007
316 17,378
123 17,571
337 17,357
1,397 16,288
133 46,643
214 46,562
389 46,387
486 46,254
74 81,070
204 80,940
85 81,059
3 81,057
508 58,695
114 59,325
408 59,165
2,484 56,752
765 309,731
472 311,896
877 312,547
4,790 305,962
1,807 1,143,033
1,238 1,152,882
774 1,158,546
771 1,145,309
551 59,477
109 59,854
397 59,517
2,132 57,778
796 316,236
398 316,138
942 315,218
3,328 312,768
1,381 1,175,819
927 1,173,913
747 1,172,293
602 1,171,958
315 40,271
63 40,859
471 40,115
3,121 37,605
338 172,471
195 174,798
808 172,001
3,836 169,883
486 508,750
257 517,715
352 508,884
360 512,516
312 40,960
67 41,163
296 41,200
2,700 38,712
283 176,985
182 176,813
510 178,214
2,937 175,241
377 526,695
225 525,755
139 532,757
169 530,543
1.dist. rot.
2.prox. rot.
3.dist. non–rot.
4.prox. non–rot.
13 demes
18 demes
min. S1–target–S2 angle
180
150
120
90
60
30
max. source–target distance
1 2 3 4 5 6 7 8 9
1e+05
10000
1000
100
10
1

**b**

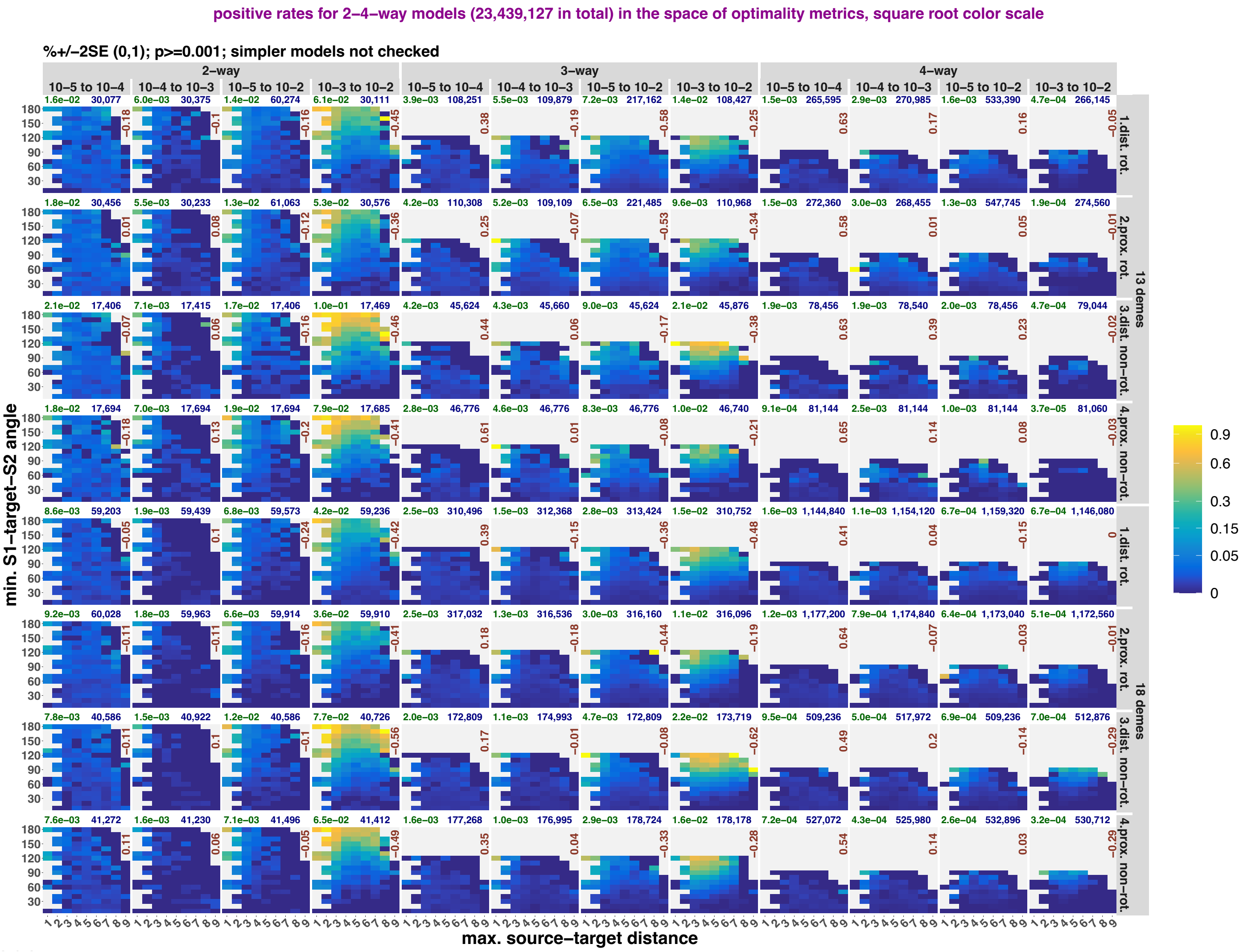

positive rates for 2–4–way models (23,439,127 in total) in the space of optimality metrics, square root color scale
%+/−2SE (0,1); p>=0.001; simpler models not checked
2–way
3–way
4–way
10–5 to 10–4
10–4 to 10–3
10–5 to 10–2
10–3 to 10–2
1.dist. rot.
2.prox. rot.
3.dist. non–rot.
4.prox. non–rot.
13 demes
18 demes
min. S1–target–S2 angle
max. source–target distance
0.9
0.6
0.3
0.15
0.05
0

**c**

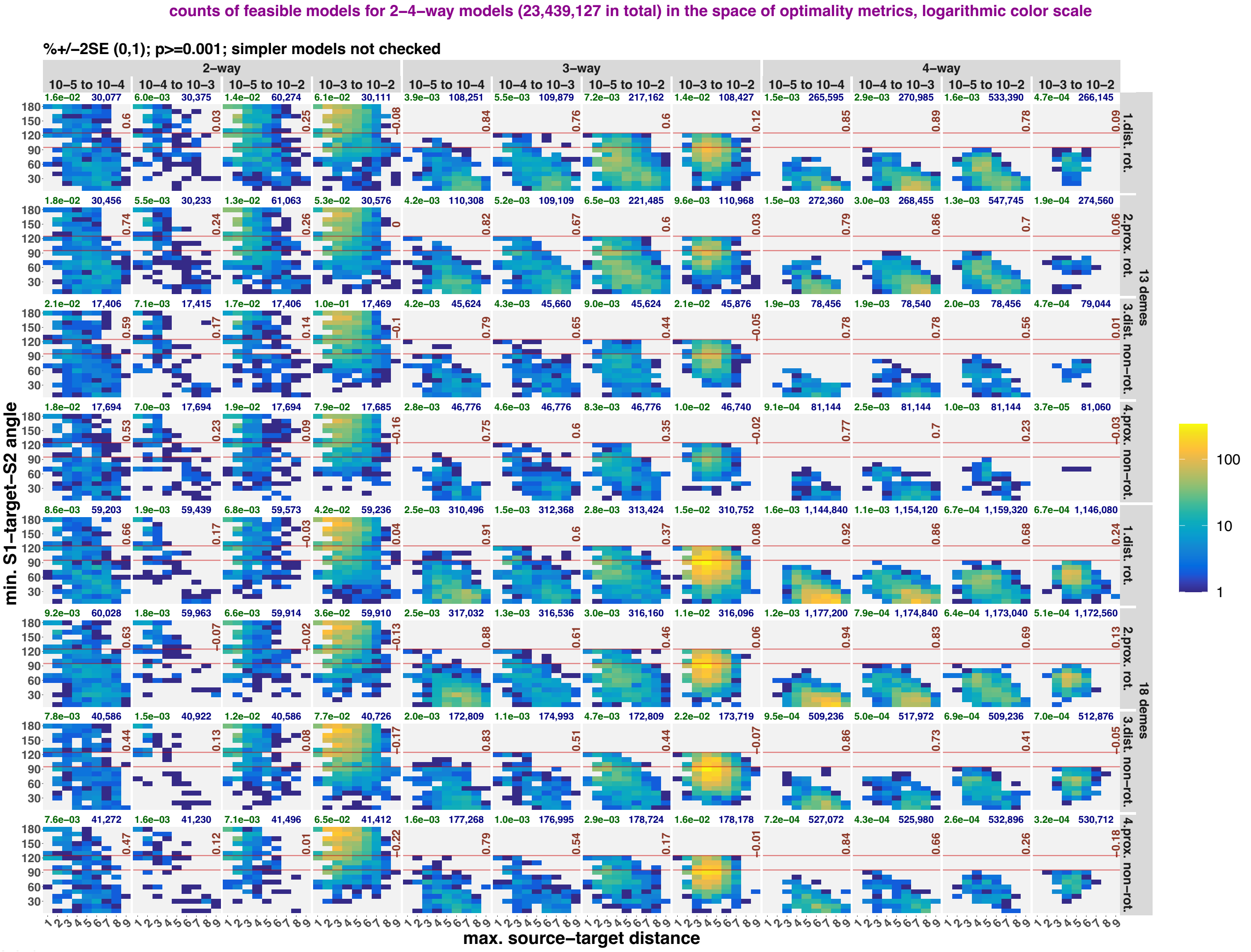


Models in the space of optimality metrics (max. ST distance and min. STS angle) following the approach from **Fig. S18**, but the results are shown for a single composite feasibility criterion deemed the best performing and stratified not only by model complexity and landscape type, but also by *qpAdm* protocols and landscape sampling density (13 or 18 demes in rotating sets and 10 or 15 demes in non-rotating “right” sets, respectively). Abbreviations: rot., rotating; non-rot., non-rotating; dist., distal; prox., proximal. The spaces are colored either by density of all randomized models in two-dimensional bins (panel **a**), or by *qpAdm* positive rates (fractions of *qpAdm* models satisfying the selected composite feasibility criterion; panel **b**), or by density of fitting models (**c**). Logarithmic color scales are used for showing density of individual *qpAdm* models populating the space, and square root color scales for *qpAdm* positive rate. Spearman’s correlation coefficients for counts of all models vs. counts of fitting models or *qpAdm* positive rates in the two-dimensional bins are shown in each panel in brown (bins not populated by any models were not considered). The red horizontal lines mark the “ideal” min. STS angles for three- and four-way models (120° and 90°). The numbers above the plots show either overall *qpAdm* positive rates (in green) and total counts of models tested in blue (**b, c**), or total counts of fitting and non-fitting models in green and red, respectively (**a**).

**Figure S26.**

**a**

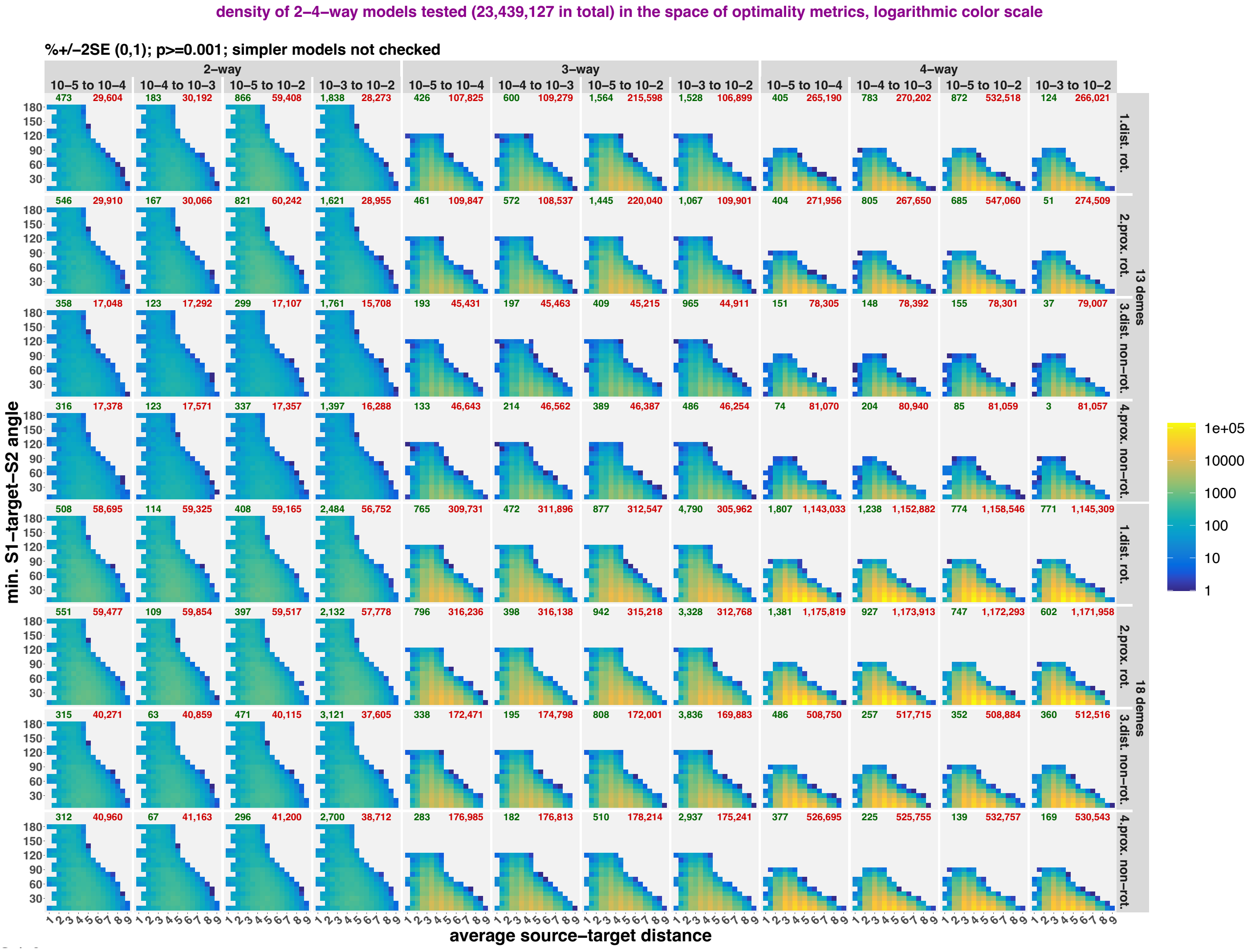

density of 2–4–way models tested (23,439,127 in total) in the space of optimality metrics, logarithmic color scale
%+/–2SE (0,1); p>=0.001; simpler models not checked
2–way
3–way
4–way
10–5 to 10–4
10–4 to 10–3
10–5 to 10–2
10–3 to 10–2
473 29,604
183 30,192
866 59,408
1,838 28,273
426 107,825
600 109,279
1,564 215,598
1,528 106,899
405 265,190
783 270,202
872 532,518
124 266,021
546 29,910
167 30,066
821 60,242
1,621 28,955
461 109,847
572 108,537
1,445 220,040
1,067 109,901
404 271,956
805 267,650
685 547,060
51 274,509
358 17,048
123 17,292
299 17,107
1,761 15,708
193 45,431
197 45,463
409 45,215
965 44,911
151 78,305
148 78,392
155 78,301
37 79,007
316 17,378
123 17,571
337 17,357
1,397 16,288
133 46,643
214 46,562
389 46,387
486 46,254
74 81,070
204 80,940
85 81,059
3 81,057
508 58,695
114 59,325
408 59,165
2,484 56,752
765 309,731
472 311,896
877 312,547
4,790 305,962
1,807 1,143,033
1,238 1,152,882
774 1,158,546
771 1,145,309
551 59,477
109 59,854
397 59,517
2,132 57,778
796 316,236
398 316,138
942 315,218
3,328 312,768
1,381 1,175,819
927 1,173,913
747 1,172,293
602 1,171,958
315 40,271
63 40,859
471 40,115
3,121 37,605
338 172,471
195 174,798
808 172,001
3,836 169,883
486 508,750
257 517,715
352 508,884
360 512,516
312 40,960
67 41,163
296 41,200
2,700 38,712
283 176,985
182 176,813
510 178,214
2,937 175,241
377 526,695
225 525,755
139 532,757
169 530,543
1.dist. rot.
2.prox. rot.
3.dist. non–rot.
4.prox. non–rot.
13 demes
18 demes
min. S1–target–S2 angle
180
150
120
90
60
30
average source–target distance
1 2 3 4 5 6 7 8 9
1e+05
10000
1000
100
10
1

1378 **b**

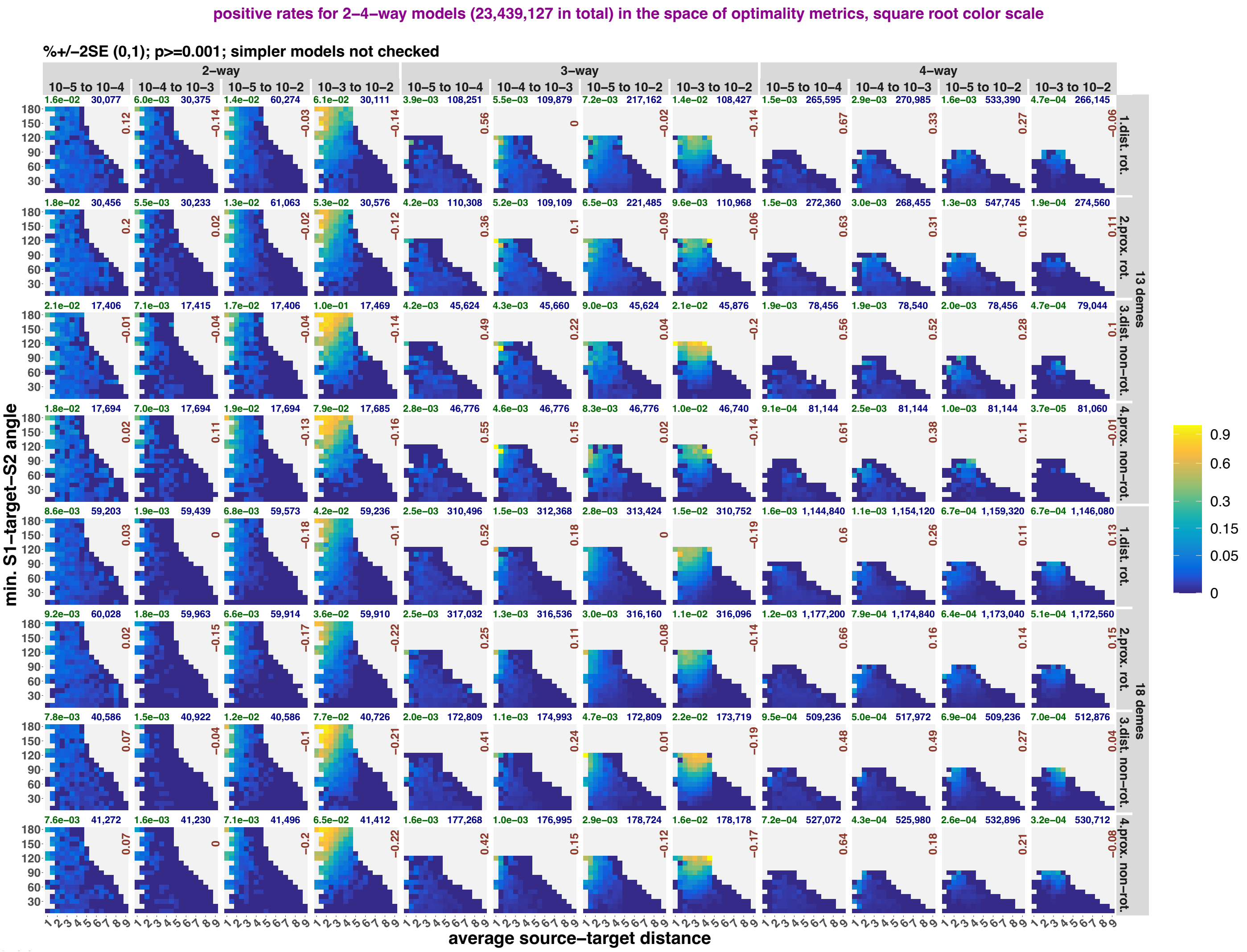
positive rates for 2–4–way models (23,439,127 in total) in the space of optimality metrics, square root color scale
%+/−2SE (0,1); p>=0.001; simpler models not checked
2–way
3–way
4–way
10–5 to 10–4
10–4 to 10–3
10–5 to 10–2
10–3 to 10–2
min. S1–target–S2 angle
average source–target distance
1.dist. rot.
2.prox. rot.
3.dist. non–rot.
4.prox. non–rot.
13 demes
18 demes
0.9
0.6
0.3
0.15
0.05
0

1380

**c**

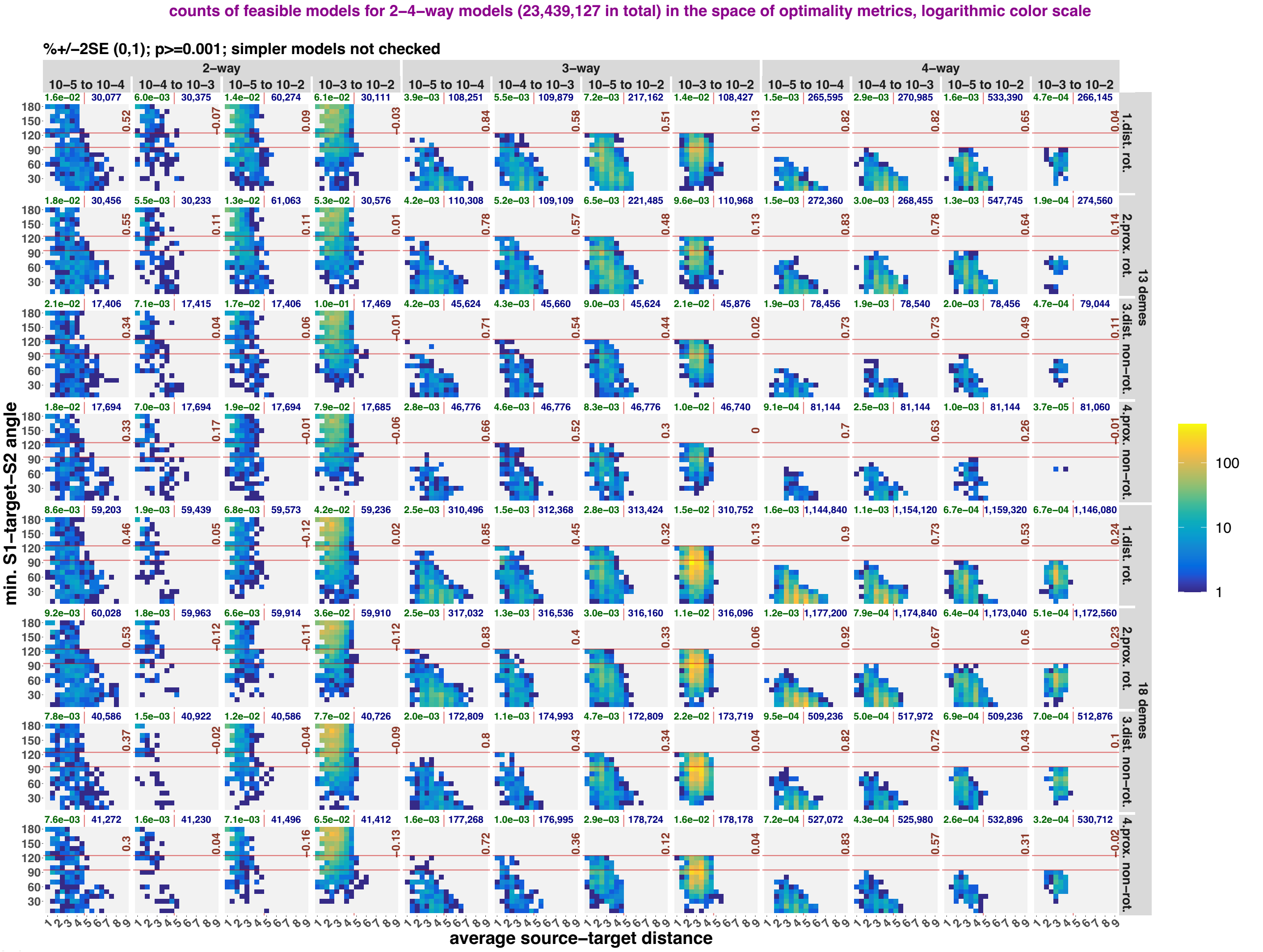


Models in the space of optimality metrics (average ST distance and min. STS angle) following the approach from **Fig. S19**, but the results are shown for a single composite feasibility criterion deemed the best performing and stratified not only by model complexity and landscape type, but also by *qpAdm* protocols and landscape sampling density (13 or 18 demes in rotating sets and 10 or 15 demes in non-rotating "right" sets, respectively). Abbreviations: rot., rotating; non-rot., non-rotating; dist., distal; prox., proximal. The spaces are colored either by density of all randomized models in two-dimensional bins (panel **a**), or by *qpAdm* positive rates (fractions of *qpAdm* models satisfying the selected composite feasibility criterion; panel **b**), or by density of fitting models (**c**). Logarithmic color scales are used for showing density of individual *qpAdm* models populating the space, and square root color scales for *qpAdm* positive rate. Spearman's correlation coefficients for counts of all models vs. counts of fitting models or *qpAdm* positive rates in the two-dimensional bins are shown in each panel in brown (bins not populated by any models were not considered). The red horizontal lines mark the "ideal" min. STS angles for three- and four-way models (120° and 90°), and the red vertical lines mark the average ST distance (4.5) that equals the radius of the landscape: a boundary separating "highly asymmetric" models to the right from more optimal models to the left. The numbers above the plots show either overall *qpAdm* positive rates (in green) and total counts of models tested in blue (**b, c**), or total counts of fitting and non-fitting models in green and red, respectively (**a**).

**Figure S27.**

**a**

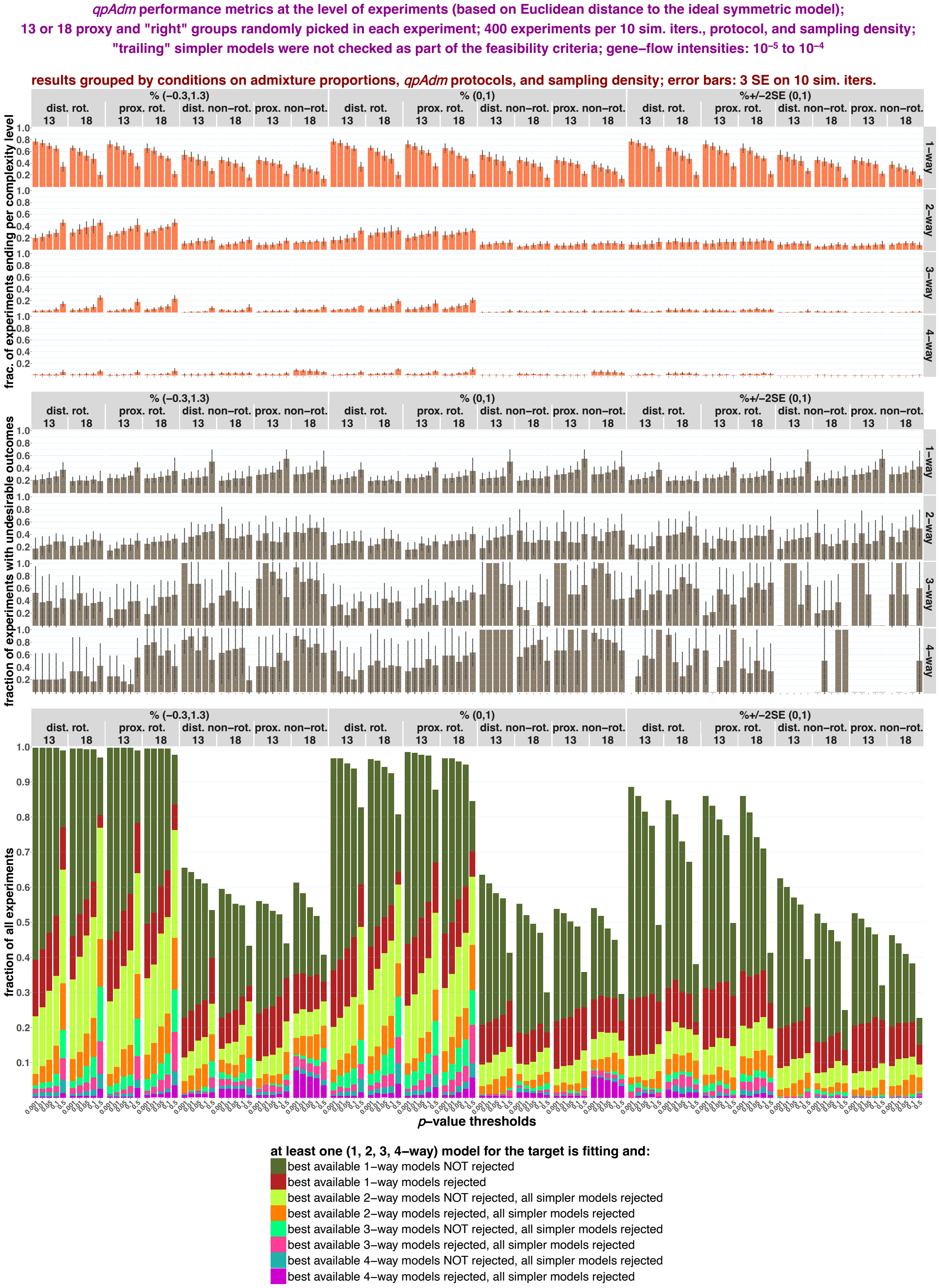
qpAdm performance metrics at the level of experiments (based on Euclidean distance to the ideal symmetric model);
13 or 18 proxy and "right" groups randomly picked in each experiment; 400 experiments per 10 sim. iters., protocol, and sampling density;
"trailing" simpler models were not checked as part of the feasibility criteria; gene–flow intensities: 10⁻⁵ to 10⁻⁴
results grouped by conditions on admixture proportions, qpAdm protocols, and sampling density; error bars: 3 SE on 10 sim. iters.
% (−0.3,1.3)
% (0,1)
%+/−2SE (0,1)
dist. rot.
prox. rot.
dist. non−rot.
prox. non−rot.
13
18
1–way
2–way
3–way
4–way
frac. of experiments ending per complexity level
fraction of experiments with undesirable outcomes
fraction of all experiments
p–value thresholds
at least one (1, 2, 3, 4–way) model for the target is fitting and:
best available 1–way models NOT rejected
best available 1–way models rejected
best available 2–way models NOT rejected, all simpler models rejected
best available 2–way models rejected, all simpler models rejected
best available 3–way models NOT rejected, all simpler models rejected
best available 3–way models rejected, all simpler models rejected
best available 4–way models NOT rejected, all simpler models rejected
best available 4–way models rejected, all simpler models rejected

**b**

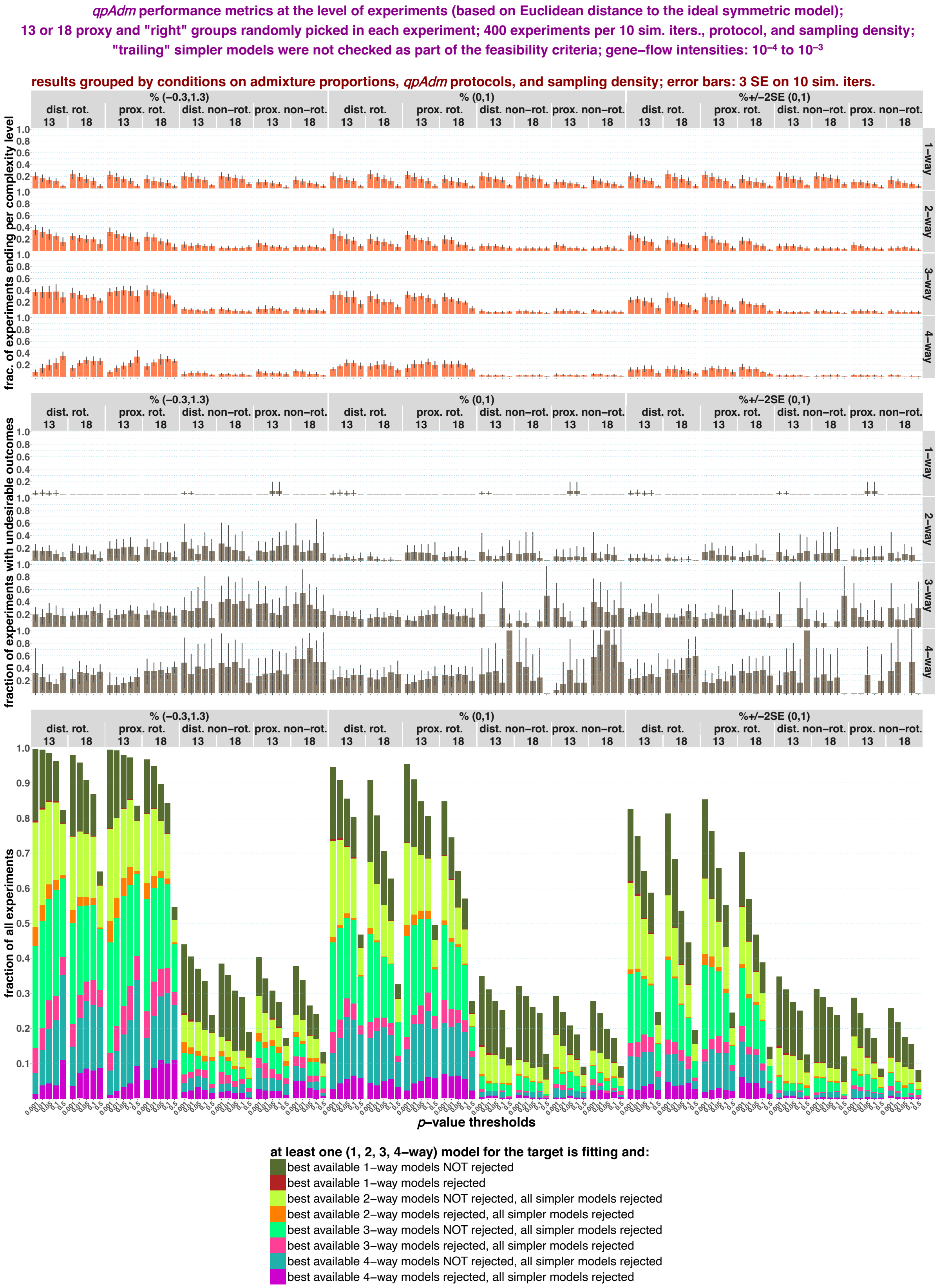

qpAdm performance metrics at the level of experiments (based on Euclidean distance to the ideal symmetric model);
13 or 18 proxy and "right" groups randomly picked in each experiment; 400 experiments per 10 sim. iters., protocol, and sampling density;
"trailing" simpler models were not checked as part of the feasibility criteria; gene–flow intensities: 10⁻⁴ to 10⁻³
results grouped by conditions on admixture proportions, qpAdm protocols, and sampling density; error bars: 3 SE on 10 sim. iters.
% (−0.3,1.3)
% (0,1)
%+/−2SE (0,1)
dist. rot.
prox. rot.
dist. non−rot.
prox. non−rot.
13
18
1−way
2−way
3−way
4−way
frac. of experiments ending per complexity level
fraction of experiments with undesirable outcomes
fraction of all experiments
p−value thresholds
0.001
0.01
0.05
0.1
0.5
at least one (1, 2, 3, 4−way) model for the target is fitting and:
best available 1−way models NOT rejected
best available 1−way models rejected
best available 2−way models NOT rejected, all simpler models rejected
best available 2−way models rejected, all simpler models rejected
best available 3−way models NOT rejected, all simpler models rejected
best available 3−way models rejected, all simpler models rejected
best available 4−way models NOT rejected, all simpler models rejected
best available 4−way models rejected, all simpler models rejected

c

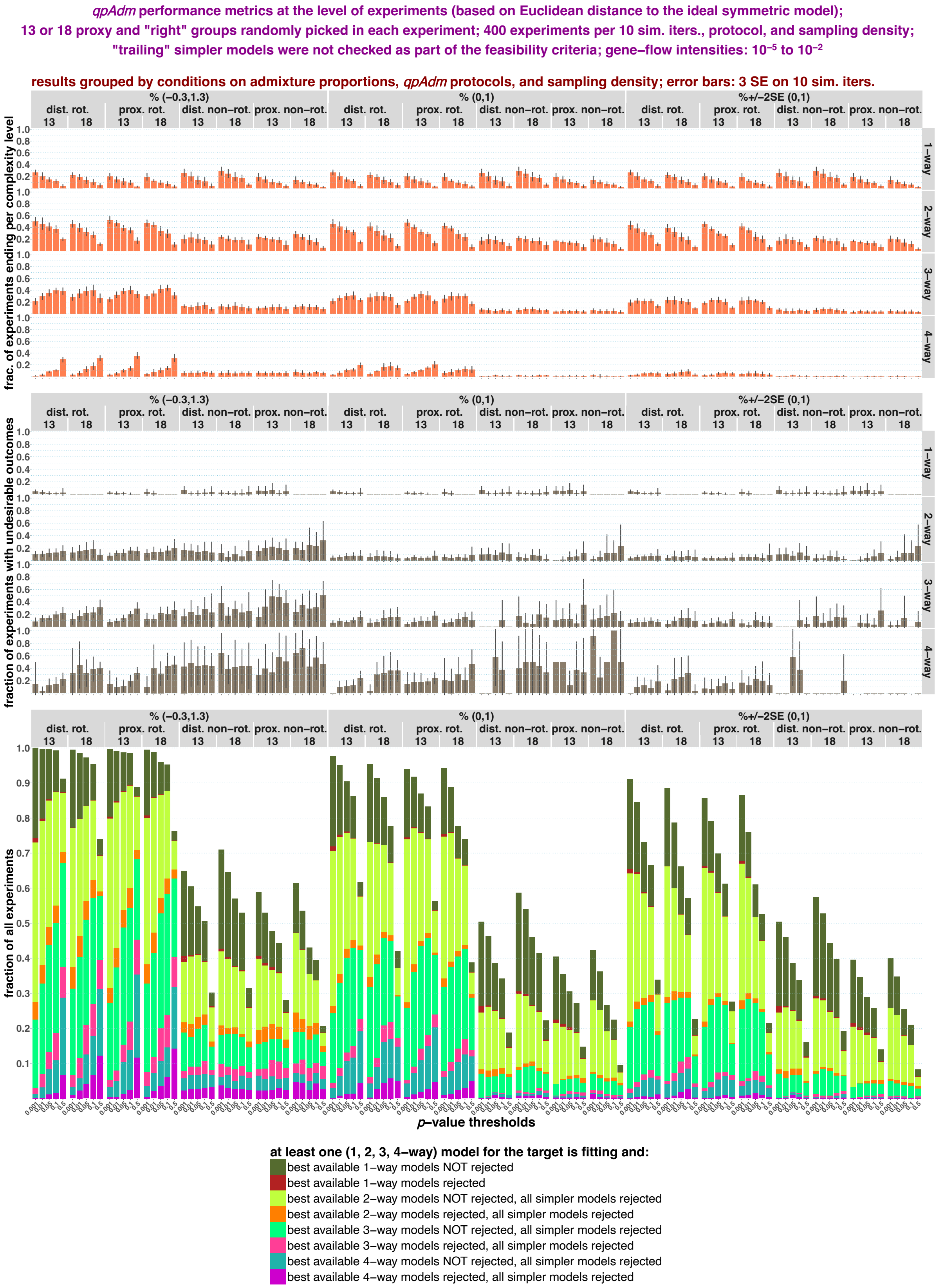

qpAdm performance metrics at the level of experiments (based on Euclidean distance to the ideal symmetric model);
13 or 18 proxy and "right" groups randomly picked in each experiment; 400 experiments per 10 sim. iters., protocol, and sampling density;
"trailing" simpler models were not checked as part of the feasibility criteria; gene−flow intensities: 10⁻⁵ to 10⁻²
results grouped by conditions on admixture proportions, qpAdm protocols, and sampling density; error bars: 3 SE on 10 sim. iters.
% (−0.3,1.3)
% (0,1)
%+/−2SE (0,1)
dist. rot.
prox. rot.
dist. non−rot.
prox. non−rot.
13
18
1−way
2−way
3−way
4−way
frac. of experiments ending per complexity level
fraction of experiments with undesirable outcomes
fraction of all experiments
p−value thresholds
at least one (1, 2, 3, 4−way) model for the target is fitting and:
best available 1−way models NOT rejected
best available 1−way models rejected
best available 2−way models NOT rejected, all simpler models rejected
best available 2−way models rejected, all simpler models rejected
best available 3−way models NOT rejected, all simpler models rejected
best available 3−way models rejected, all simpler models rejected
best available 4−way models NOT rejected, all simpler models rejected
best available 4−way models rejected, all simpler models rejected

**d**

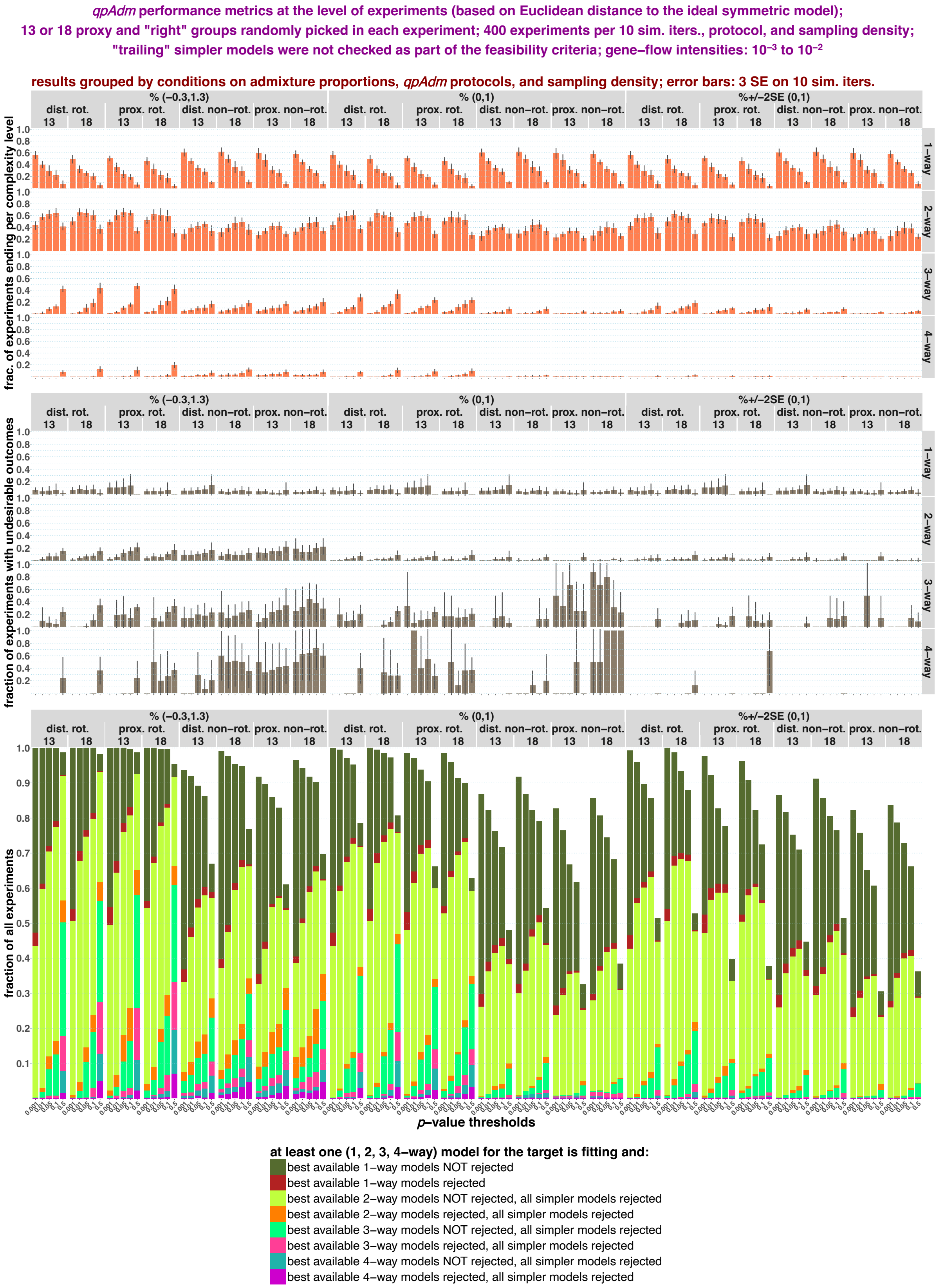
qpAdm performance metrics at the level of experiments (based on Euclidean distance to the ideal symmetric model);
13 or 18 proxy and "right" groups randomly picked in each experiment; 400 experiments per 10 sim. iters., protocol, and sampling density;
"trailing" simpler models were not checked as part of the feasibility criteria; gene−flow intensities: 10⁻³ to 10⁻²
results grouped by conditions on admixture proportions, qpAdm protocols, and sampling density; error bars: 3 SE on 10 sim. iters.
% (−0.3,1.3)
% (0,1)
%+/−2SE (0,1)
dist. rot.
prox. rot.
dist. non−rot.
prox. non−rot.
13
18
1−way
2−way
3−way
4−way
frac. of experiments ending per complexity level
fraction of experiments with undesirable outcomes
fraction of all experiments
p−value thresholds
at least one (1, 2, 3, 4−way) model for the target is fitting and:
best available 1−way models NOT rejected
best available 1−way models rejected
best available 2−way models NOT rejected, all simpler models rejected
best available 2−way models rejected, all simpler models rejected
best available 3−way models NOT rejected, all simpler models rejected
best available 3−way models rejected, all simpler models rejected
best available 4−way models NOT rejected, all simpler models rejected
best available 4−way models rejected, all simpler models rejected

**e**

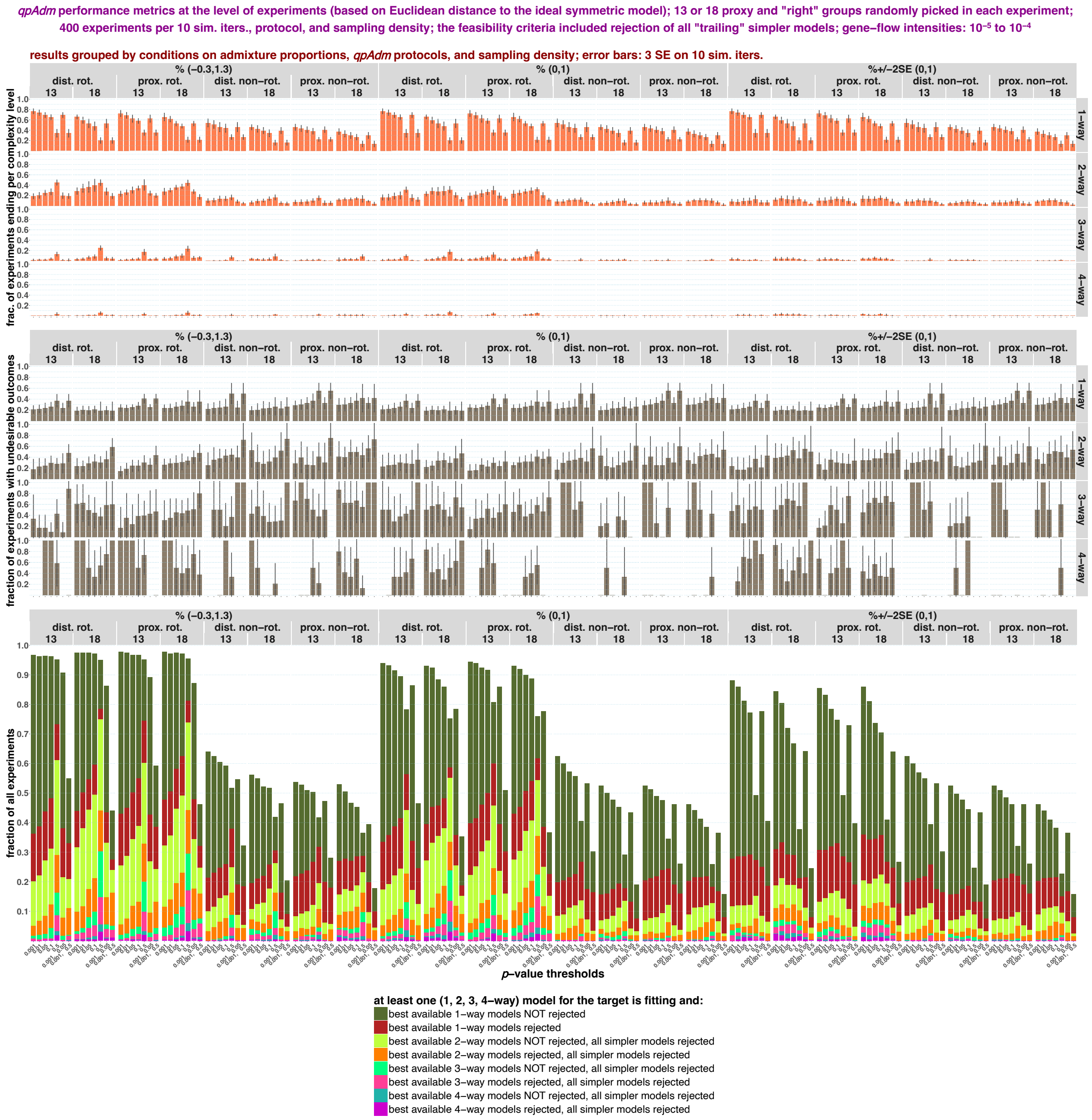
qpAdm performance metrics at the level of experiments (based on Euclidean distance to the ideal symmetric model); 13 or 18 proxy and "right" groups randomly picked in each experiment;
400 experiments per 10 sim. iters., protocol, and sampling density; the feasibility criteria included rejection of all "trailing" simpler models; gene–flow intensities: 10⁻⁵ to 10⁻⁴
results grouped by conditions on admixture proportions, qpAdm protocols, and sampling density; error bars: 3 SE on 10 sim. iters.
% (−0.3,1.3)
% (0,1)
%+/−2SE (0,1)
dist. rot.
prox. rot.
dist. non−rot.
prox. non−rot.
13
18
1−way
2−way
3−way
4−way
frac. of experiments ending per complexity level
fraction of experiments with undesirable outcomes
fraction of all experiments
p–value thresholds
at least one (1, 2, 3, 4–way) model for the target is fitting and:
best available 1–way models NOT rejected
best available 1–way models rejected
best available 2–way models NOT rejected, all simpler models rejected
best available 2–way models rejected, all simpler models rejected
best available 3–way models NOT rejected, all simpler models rejected
best available 3–way models rejected, all simpler models rejected
best available 4–way models NOT rejected, all simpler models rejected
best available 4–way models rejected, all simpler models rejected

**f**

*qpAdm* performance metrics at the level of experiments (based on Euclidean distance to the ideal symmetric model); 13 or 18 proxy and "right" groups randomly picked in each experiment; 400 experiments per 10 sim. iters., protocol, and sampling density; the feasibility criteria included rejection of all "trailing" simpler models; gene–flow intensities: $10^{-4}$ to $10^{-3}$

results grouped by conditions on admixture proportions, *qpAdm* protocols, and sampling density; error bars: 3 SE on 10 sim. iters.

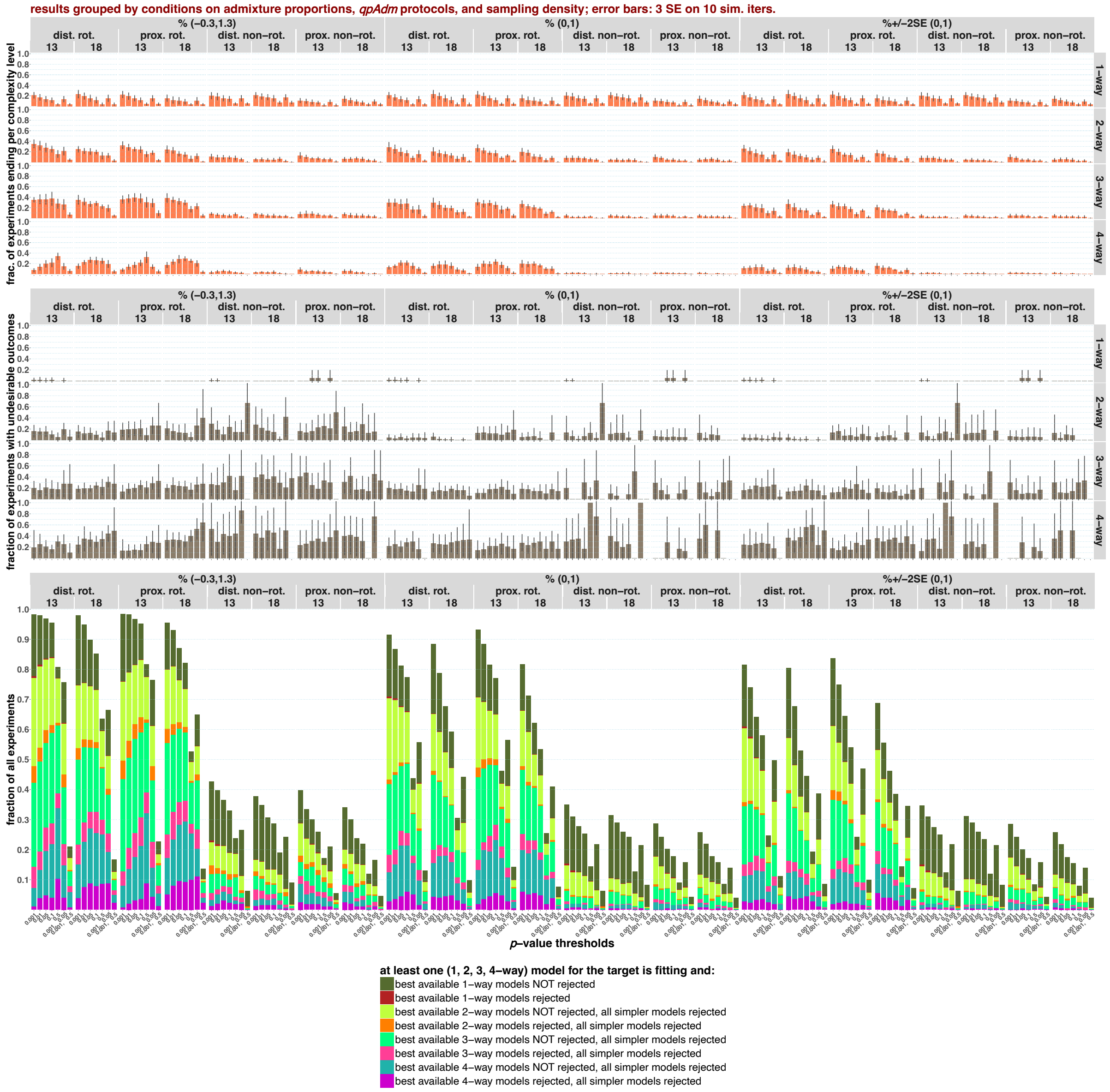

**g**

*qpAdm* performance metrics at the level of experiments (based on Euclidean distance to the ideal symmetric model); 13 or 18 proxy and "right" groups randomly picked in each experiment; 400 experiments per 10 sim. iters., protocol, and sampling density; the feasibility criteria included rejection of all "trailing" simpler models; gene–flow intensities: $10^{-5}$ to $10^{-2}$

results grouped by conditions on admixture proportions, *qpAdm* protocols, and sampling density; error bars: 3 SE on 10 sim. iters.

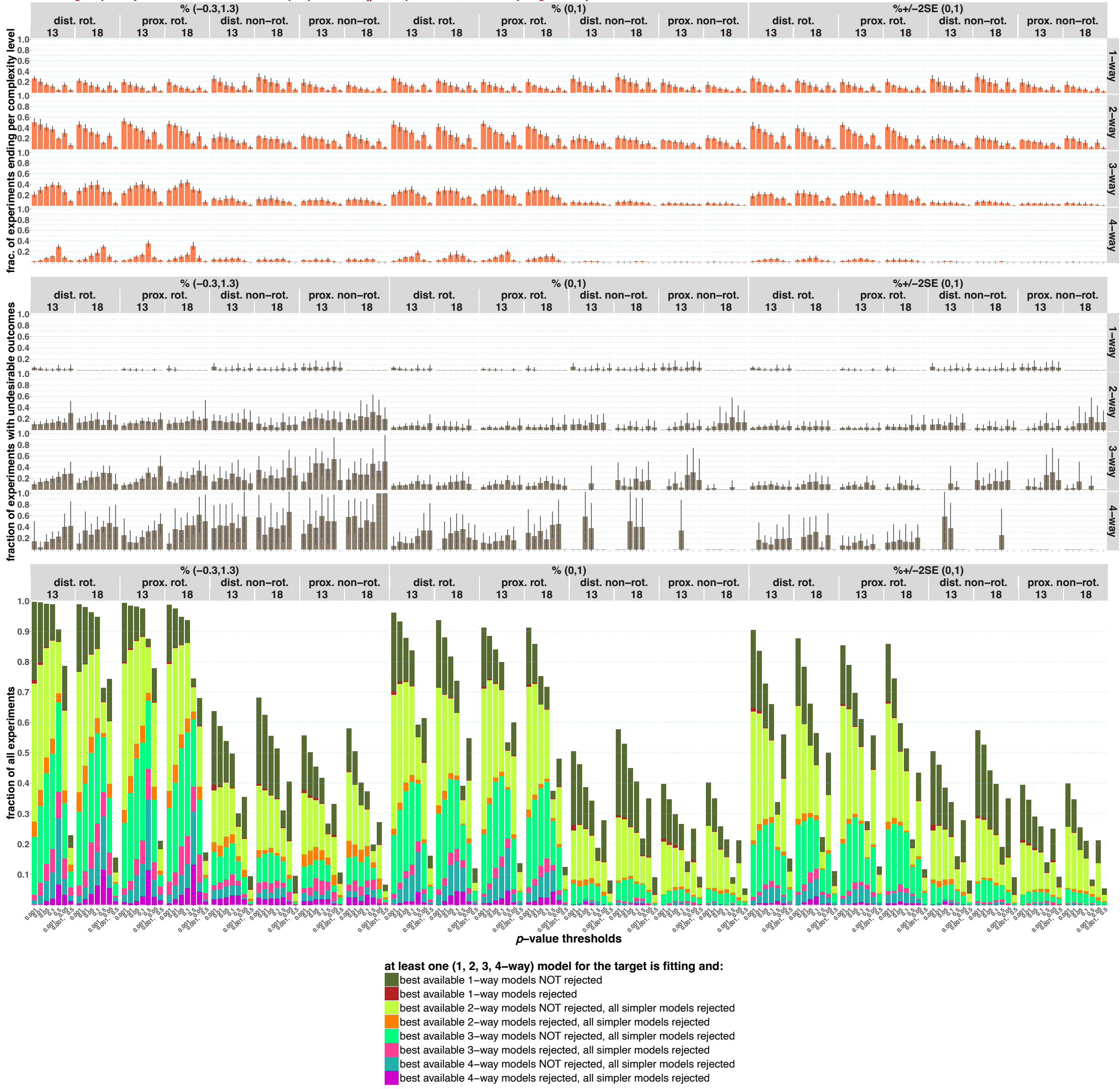

**h**

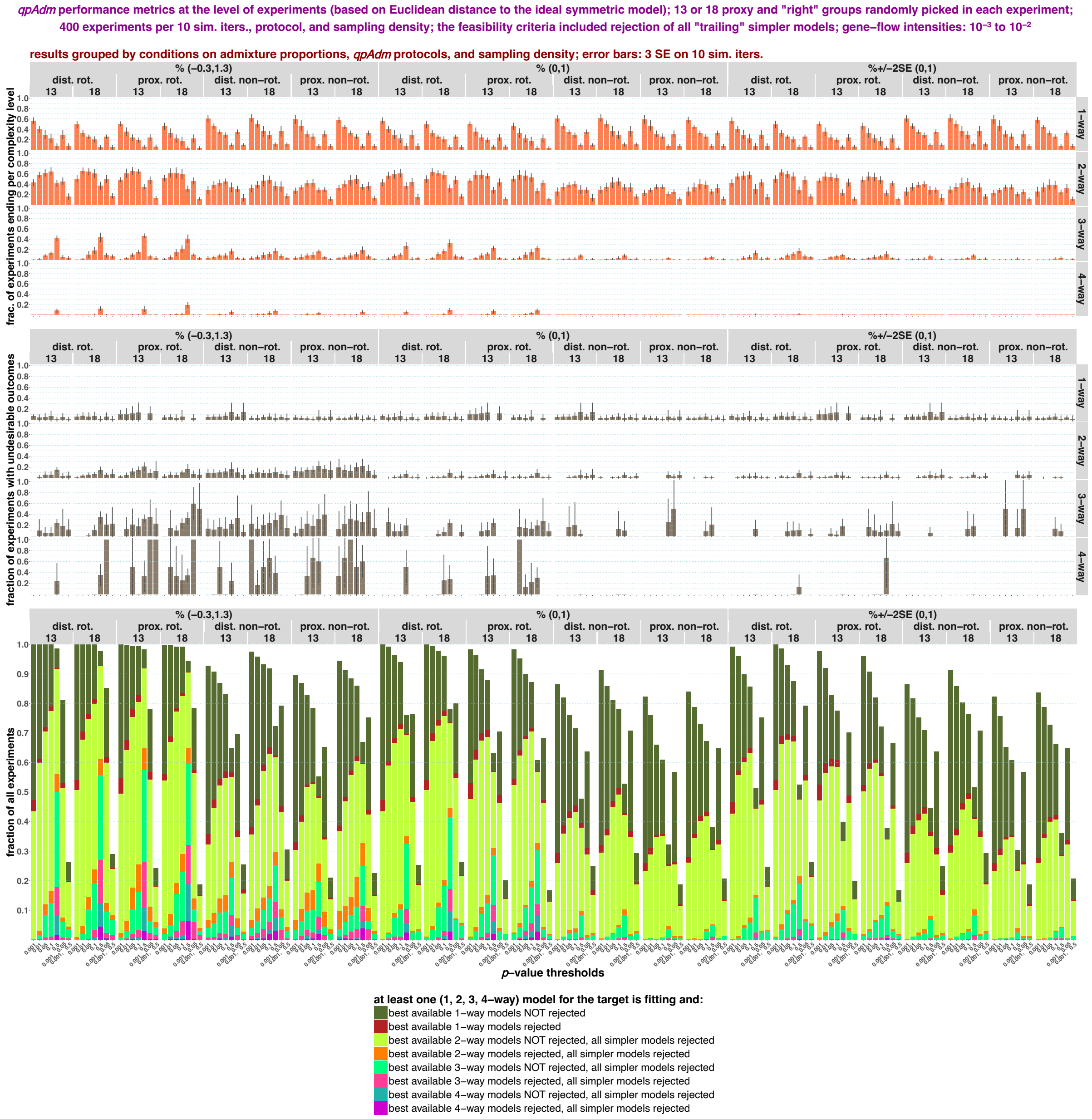


Randomized *qpAdm* results at the level of experiments visualized for all landscape types and all composite feasibility criteria. The results are stratified by *qpAdm* protocols and landscape sampling density (13 or 18 demes in rotating sets and 10 or 15 demes in non-rotating "right" sets, respectively), and also by model complexity level at which experiments generate positive results, by conditions on EAF, and by *p*-value thresholds (results for all the thresholds tested in this study are shown). We show as bar plots fractions of experiments producing positive results (at least one fitting model per target) at each complexity level and fractions of those experiments with positive outcomes classified as potentially misleading or "undesirable". See Methods for a definition based on Euclidean distances in the space of optimality metrics: max. ST distance and min. STS angle. In the case of "bad" experiments, the most optimal model (according to ST distances and angles) available in the chosen deme set was rejected, but a much less optimal model emerged as fitting. The error bars show 3 SE intervals calculated on 10 simulation replicates. Fractions of experiments falling into these different classes (producing positive results at complexity levels from one to four and classified as either misleading or not misleading according to rejection of most optimal models available) are visualized by the stacked bar plots. Error bars are not shown for visual clarity in the stacked bar plots. The results are shown separately for the “$10^{-5}$ to $10^{-4}$” (**a, e**), “$10^{-4}$ to $10^{-3}$” (**b, f**), “$10^{-5}$ to $10^{-2}$” (**c, g**), and “$10^{-3}$ to $10^{-2}$” (**d, h**) landscape types and for composite feasibility criteria either not considering “trailing” simpler models (**a-d**) or including *p*-value thresholds for rejection of these models (**e-h**).

**Figure S28.**

**a**

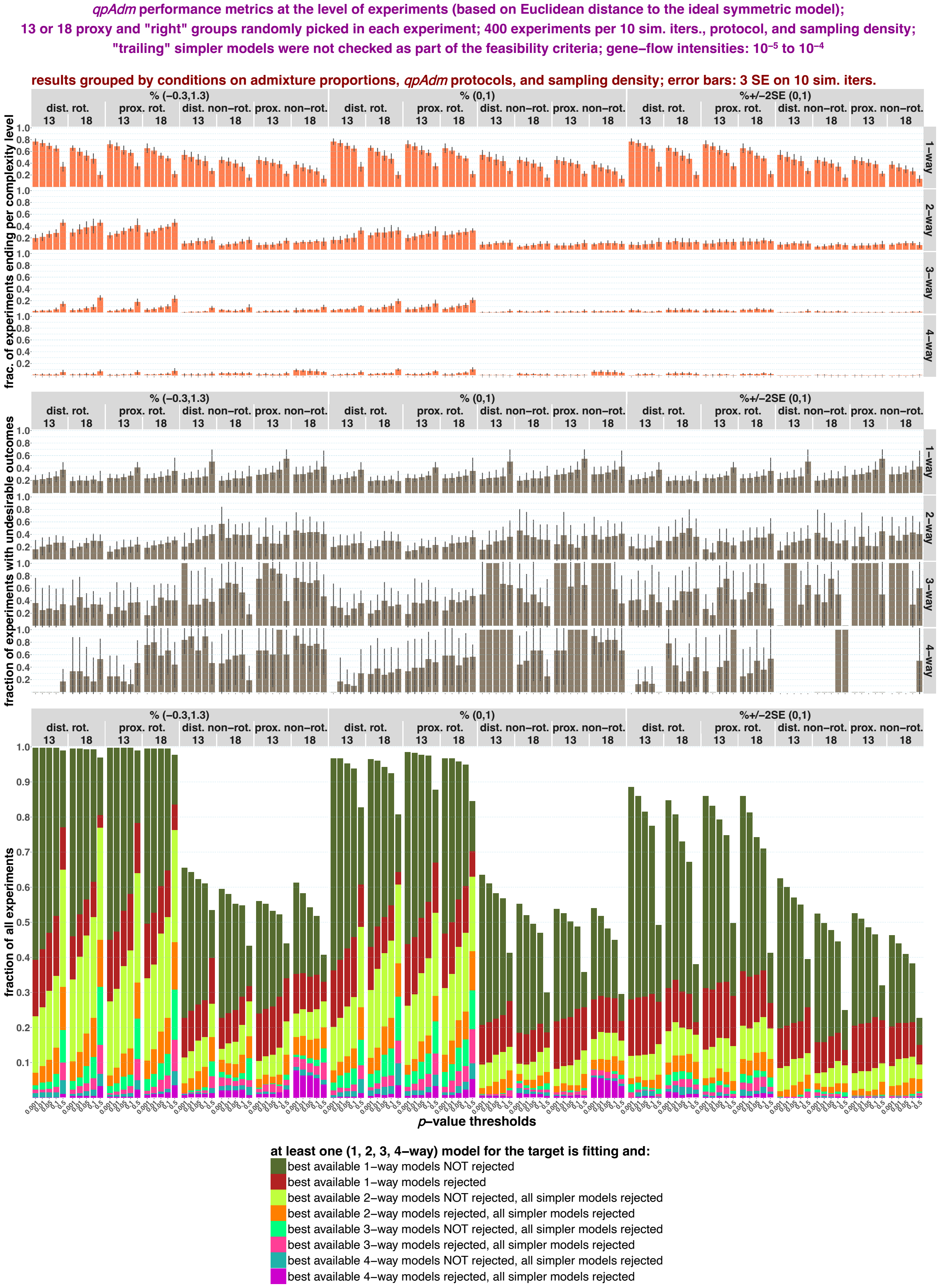

*qpAdm* performance metrics at the level of experiments (based on Euclidean distance to the ideal symmetric model);
13 or 18 proxy and "right" groups randomly picked in each experiment; 400 experiments per 10 sim. iters., protocol, and sampling density;
"trailing" simpler models were not checked as part of the feasibility criteria; gene–flow intensities: $10^{-5}$ to $10^{-4}$
results grouped by conditions on admixture proportions, *qpAdm* protocols, and sampling density; error bars: 3 SE on 10 sim. iters.
% (−0.3,1.3)
% (0,1)
%+/−2SE (0,1)
dist. rot.
prox. rot.
dist. non−rot.
prox. non−rot.
13
18
1–way
2–way
3–way
4–way
frac. of experiments ending per complexity level
fraction of experiments with undesirable outcomes
fraction of all experiments
*p*–value thresholds
at least one (1, 2, 3, 4–way) model for the target is fitting and:
best available 1–way models NOT rejected
best available 1–way models rejected
best available 2–way models NOT rejected, all simpler models rejected
best available 2–way models rejected, all simpler models rejected
best available 3–way models NOT rejected, all simpler models rejected
best available 3–way models rejected, all simpler models rejected
best available 4–way models NOT rejected, all simpler models rejected
best available 4–way models rejected, all simpler models rejected

**b**

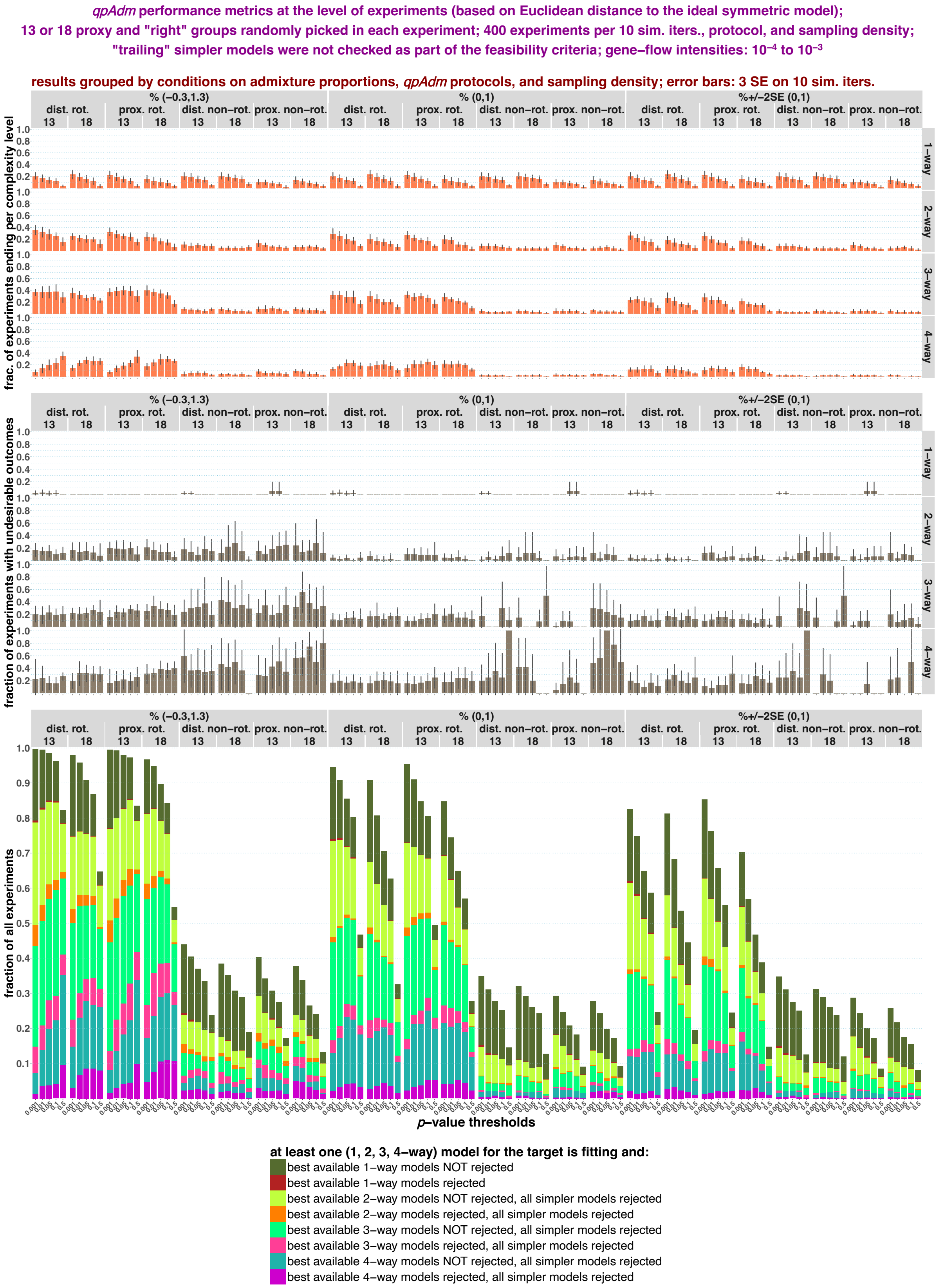
qpAdm performance metrics at the level of experiments (based on Euclidean distance to the ideal symmetric model);
13 or 18 proxy and "right" groups randomly picked in each experiment; 400 experiments per 10 sim. iters., protocol, and sampling density;
"trailing" simpler models were not checked as part of the feasibility criteria; gene–flow intensities: 10⁻⁴ to 10⁻³
results grouped by conditions on admixture proportions, qpAdm protocols, and sampling density; error bars: 3 SE on 10 sim. iters.
% (−0.3,1.3)
% (0,1)
%+/−2SE (0,1)
dist. rot.
prox. rot.
dist. non−rot.
prox. non−rot.
13
18
1−way
2−way
3−way
4−way
frac. of experiments ending per complexity level
fraction of experiments with undesirable outcomes
fraction of all experiments
p−value thresholds
at least one (1, 2, 3, 4−way) model for the target is fitting and:
best available 1−way models NOT rejected
best available 1−way models rejected
best available 2−way models NOT rejected, all simpler models rejected
best available 2−way models rejected, all simpler models rejected
best available 3−way models NOT rejected, all simpler models rejected
best available 3−way models rejected, all simpler models rejected
best available 4−way models NOT rejected, all simpler models rejected
best available 4−way models rejected, all simpler models rejected

**c**

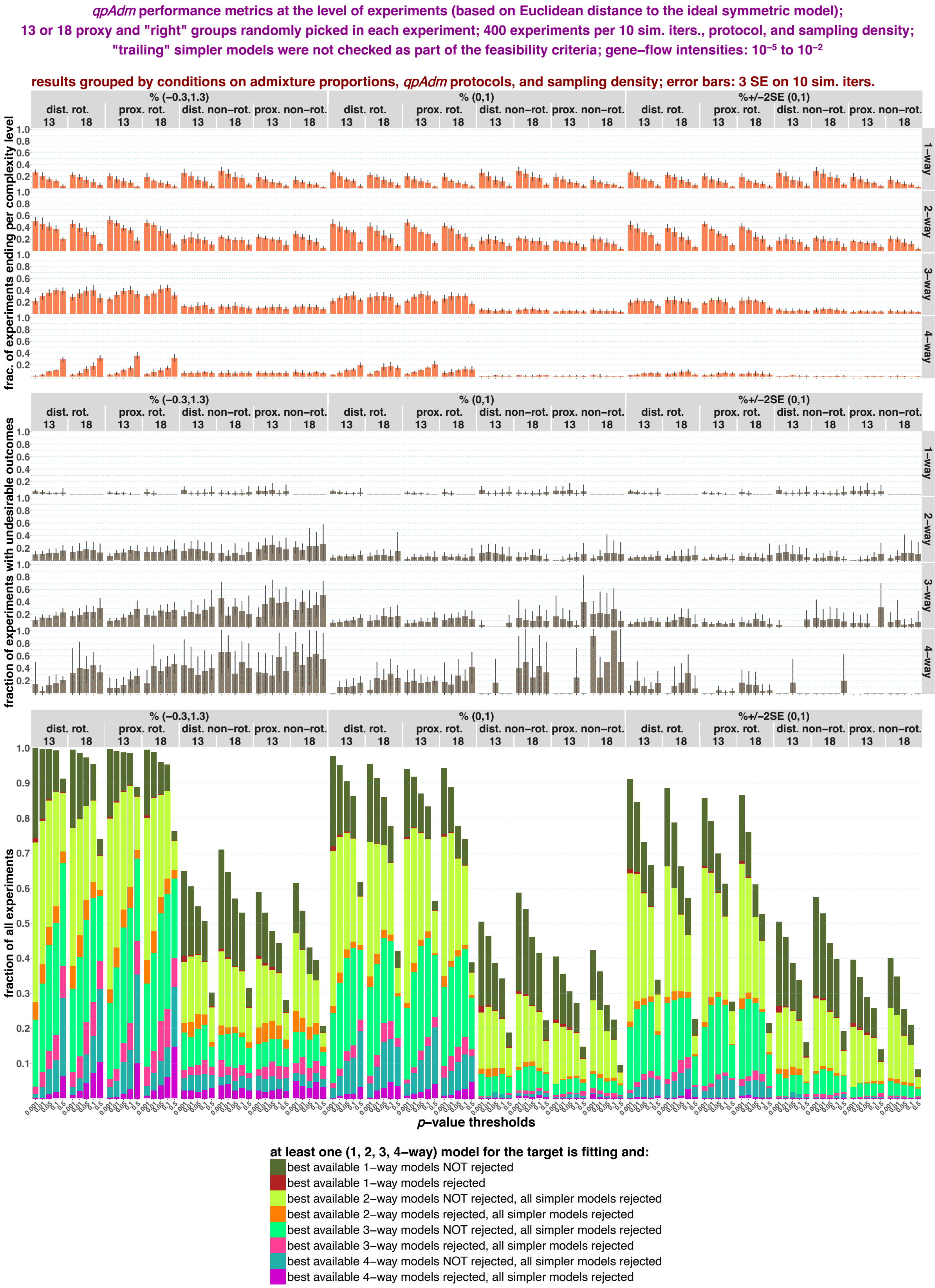
qpAdm performance metrics at the level of experiments (based on Euclidean distance to the ideal symmetric model);
13 or 18 proxy and "right" groups randomly picked in each experiment; 400 experiments per 10 sim. iters., protocol, and sampling density;
"trailing" simpler models were not checked as part of the feasibility criteria; gene–flow intensities: 10⁻⁵ to 10⁻²
results grouped by conditions on admixture proportions, qpAdm protocols, and sampling density; error bars: 3 SE on 10 sim. iters.
% (−0.3,1.3)
% (0,1)
%+/−2SE (0,1)
dist. rot.
prox. rot.
dist. non−rot.
prox. non−rot.
13
18
1−way
2−way
3−way
4−way
frac. of experiments ending per complexity level
fraction of experiments with undesirable outcomes
fraction of all experiments
p−value thresholds
at least one (1, 2, 3, 4−way) model for the target is fitting and:
best available 1−way models NOT rejected
best available 1−way models rejected
best available 2−way models NOT rejected, all simpler models rejected
best available 2−way models rejected, all simpler models rejected
best available 3−way models NOT rejected, all simpler models rejected
best available 3−way models rejected, all simpler models rejected
best available 4−way models NOT rejected, all simpler models rejected
best available 4−way models rejected, all simpler models rejected

**d**

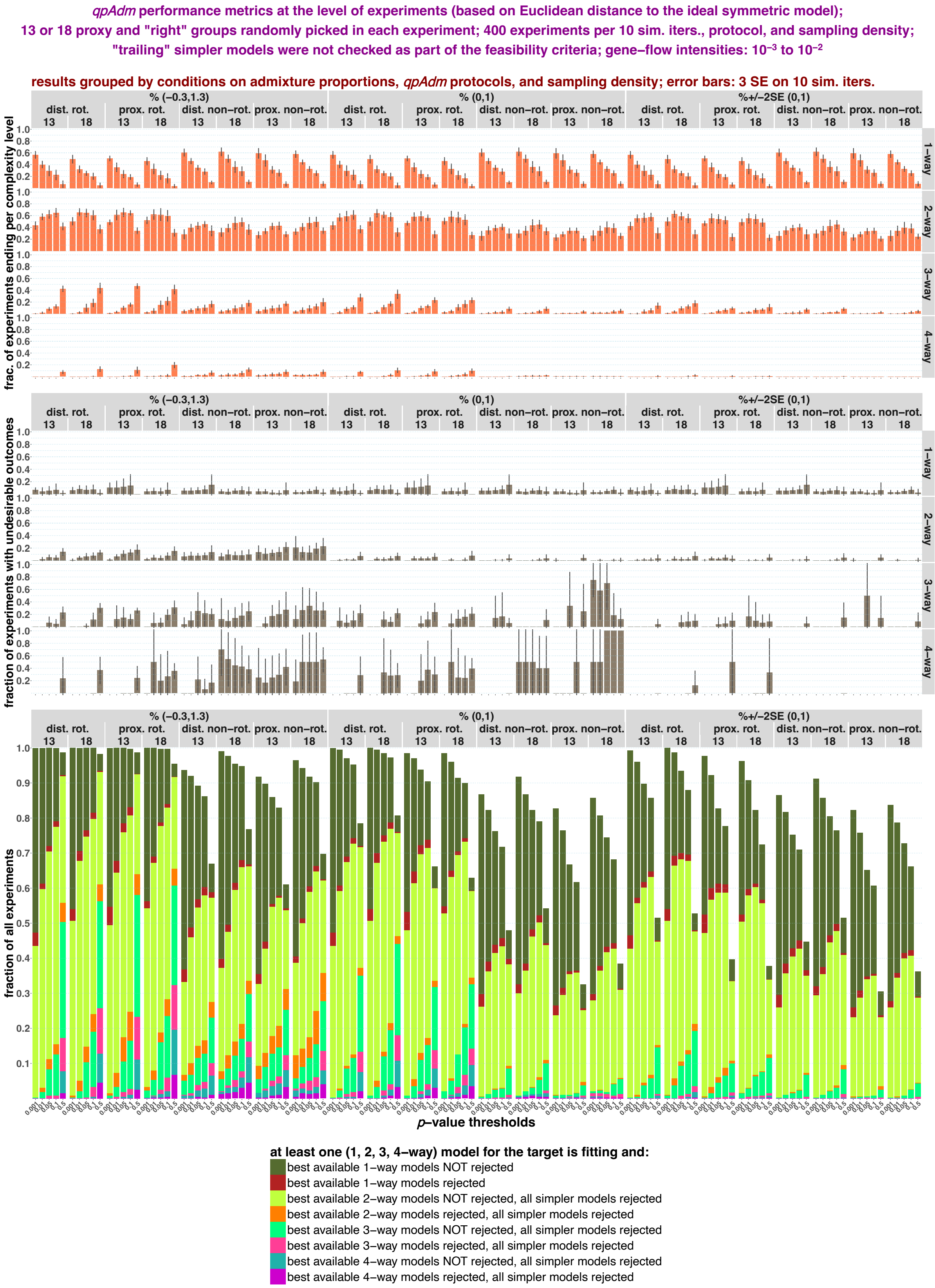

*qpAdm* performance metrics at the level of experiments (based on Euclidean distance to the ideal symmetric model);
13 or 18 proxy and "right" groups randomly picked in each experiment; 400 experiments per 10 sim. iters., protocol, and sampling density;
"trailing" simpler models were not checked as part of the feasibility criteria; gene–flow intensities: $10^{-3}$ to $10^{-2}$
results grouped by conditions on admixture proportions, *qpAdm* protocols, and sampling density; error bars: 3 SE on 10 sim. iters.
% (−0.3,1.3)
% (0,1)
%+/−2SE (0,1)
dist. rot.
prox. rot.
dist. non−rot.
prox. non−rot.
13
18
1−way
2−way
3−way
4−way
frac. of experiments ending per complexity level
fraction of experiments with undesirable outcomes
fraction of all experiments
p–value thresholds
0.001
0.01
0.05
0.1
0.5
at least one (1, 2, 3, 4−way) model for the target is fitting and:
best available 1−way models NOT rejected
best available 1−way models rejected
best available 2−way models NOT rejected, all simpler models rejected
best available 2−way models rejected, all simpler models rejected
best available 3−way models NOT rejected, all simpler models rejected
best available 3−way models rejected, all simpler models rejected
best available 4−way models NOT rejected, all simpler models rejected
best available 4−way models rejected, all simpler models rejected

**e**

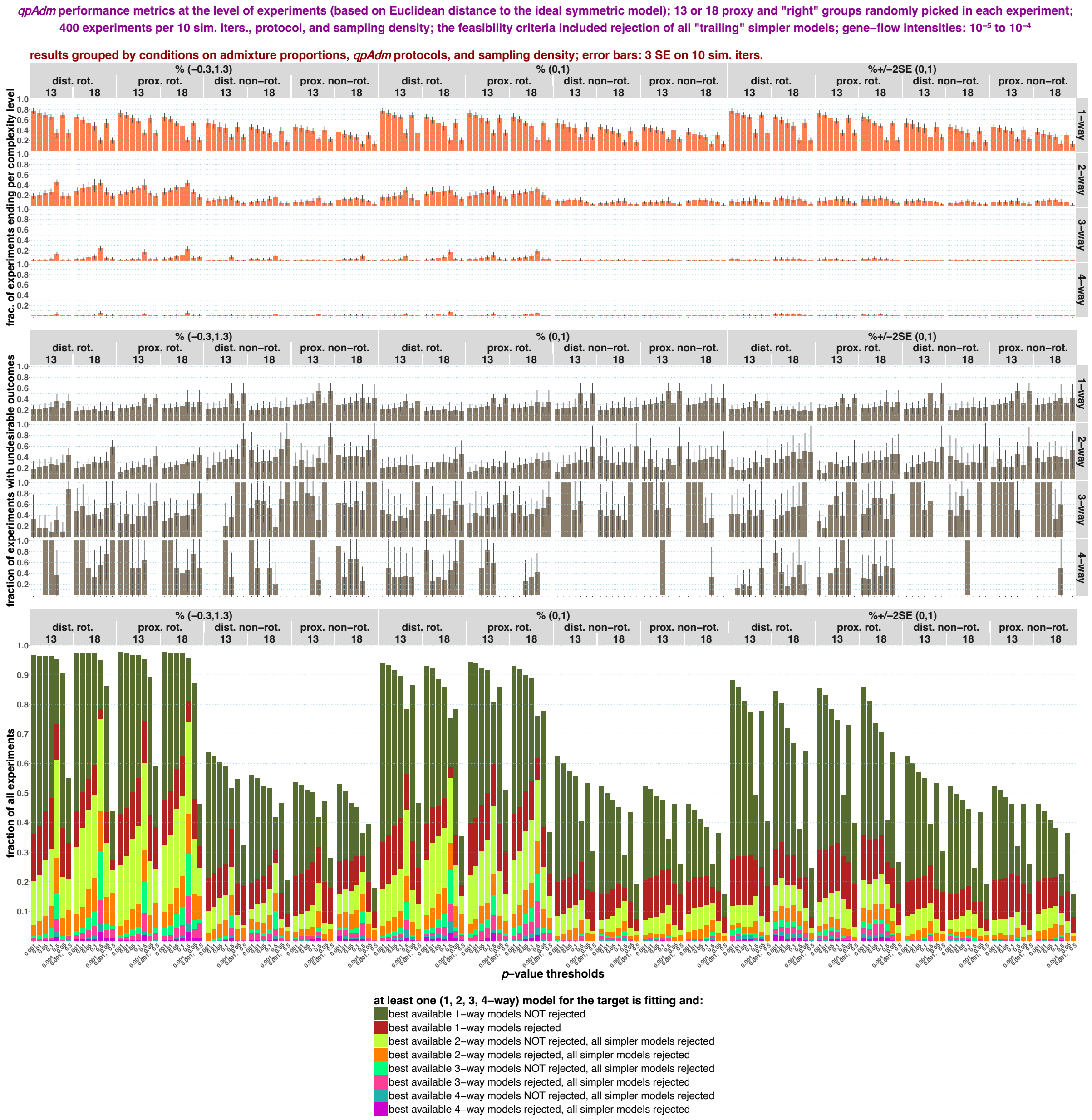
qpAdm performance metrics at the level of experiments (based on Euclidean distance to the ideal symmetric model); 13 or 18 proxy and "right" groups randomly picked in each experiment;
400 experiments per 10 sim. iters., protocol, and sampling density; the feasibility criteria included rejection of all "trailing" simpler models; gene–flow intensities: 10⁻⁵ to 10⁻⁴
results grouped by conditions on admixture proportions, qpAdm protocols, and sampling density; error bars: 3 SE on 10 sim. iters.
% (−0.3,1.3)
% (0,1)
%+/−2SE (0,1)
dist. rot.
prox. rot.
dist. non−rot.
prox. non−rot.
13
18
1−way
2−way
3−way
4−way
frac. of experiments ending per complexity level
fraction of experiments with undesirable outcomes
fraction of all experiments
p−value thresholds
at least one (1, 2, 3, 4−way) model for the target is fitting and:
best available 1−way models NOT rejected
best available 1−way models rejected
best available 2−way models NOT rejected, all simpler models rejected
best available 2−way models rejected, all simpler models rejected
best available 3−way models NOT rejected, all simpler models rejected
best available 3−way models rejected, all simpler models rejected
best available 4−way models NOT rejected, all simpler models rejected
best available 4−way models rejected, all simpler models rejected

1460 **f**

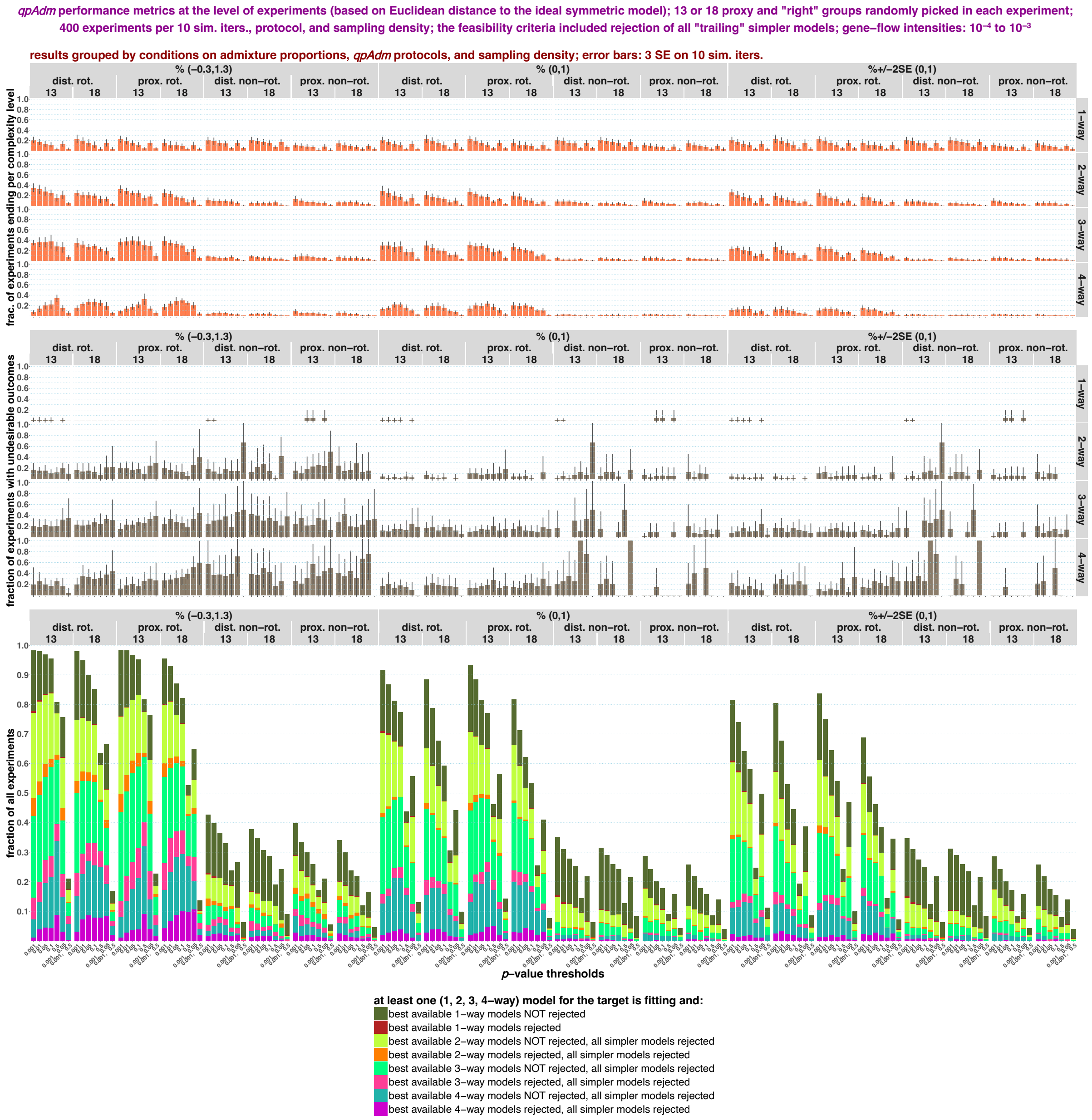


1461
1462

**g**

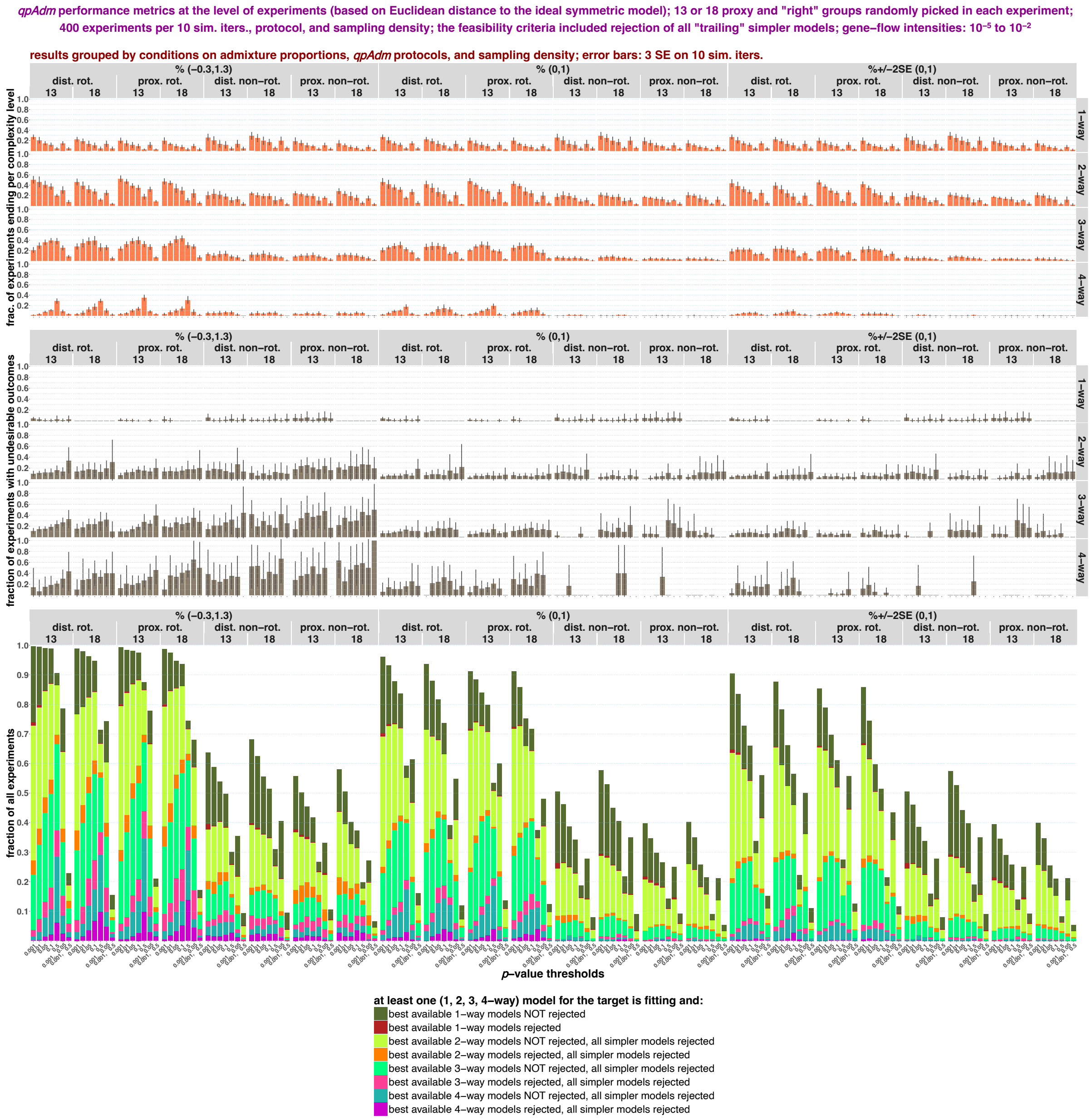
qpAdm performance metrics at the level of experiments (based on Euclidean distance to the ideal symmetric model); 13 or 18 proxy and "right" groups randomly picked in each experiment; 400 experiments per 10 sim. iters., protocol, and sampling density; the feasibility criteria included rejection of all "trailing" simpler models; gene–flow intensities: 10⁻⁵ to 10⁻²
results grouped by conditions on admixture proportions, qpAdm protocols, and sampling density; error bars: 3 SE on 10 sim. iters.
% (−0.3,1.3)
% (0,1)
%+/−2SE (0,1)
dist. rot.
prox. rot.
dist. non−rot.
prox. non−rot.
13
18
1−way
2−way
3−way
4−way
frac. of experiments ending per complexity level
fraction of experiments with undesirable outcomes
fraction of all experiments
p−value thresholds
at least one (1, 2, 3, 4−way) model for the target is fitting and:
best available 1−way models NOT rejected
best available 1−way models rejected
best available 2−way models NOT rejected, all simpler models rejected
best available 2−way models rejected, all simpler models rejected
best available 3−way models NOT rejected, all simpler models rejected
best available 3−way models rejected, all simpler models rejected
best available 4−way models NOT rejected, all simpler models rejected
best available 4−way models rejected, all simpler models rejected

**g**

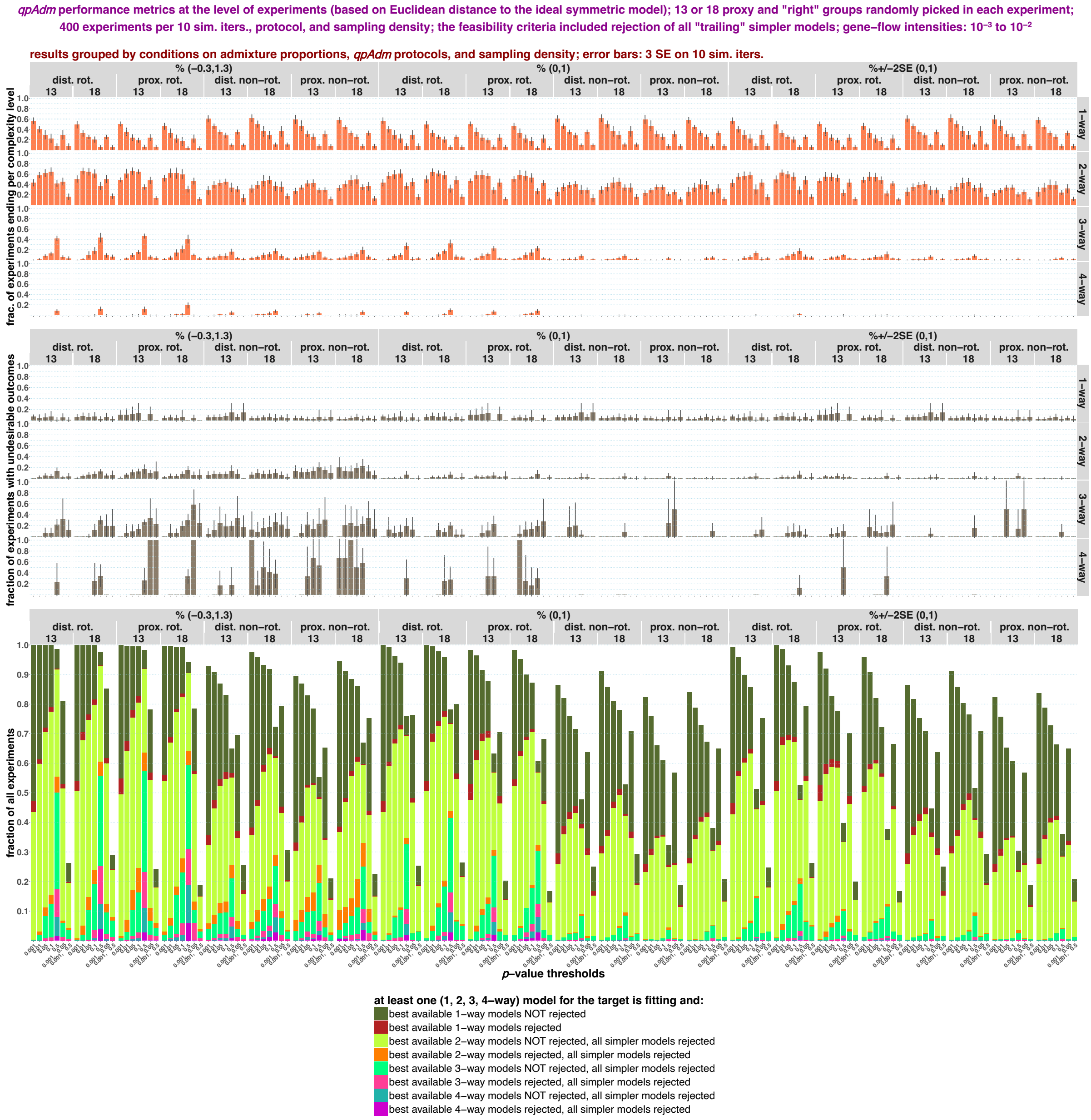


The same results as in **Fig. S27** but fractions of positive experiments with potentially misleading outcomes are based on the space of average ST distance vs. min. STS angle instead of max. ST distance vs. min. STS angle. The results are shown separately for the "$10^{-5}$ to $10^{-4}$" (**a, e**), "$10^{-4}$ to $10^{-3}$" (**b, f**), "$10^{-5}$ to $10^{-2}$" (**c, g**), and "$10^{-3}$ to $10^{-2}$" (**d, h**) landscape types and for composite feasibility criteria either not considering "trailing" simpler models (**a-d**) or including *p*-value thresholds for rejection of these models (**e-h**).

**Figure S29.**

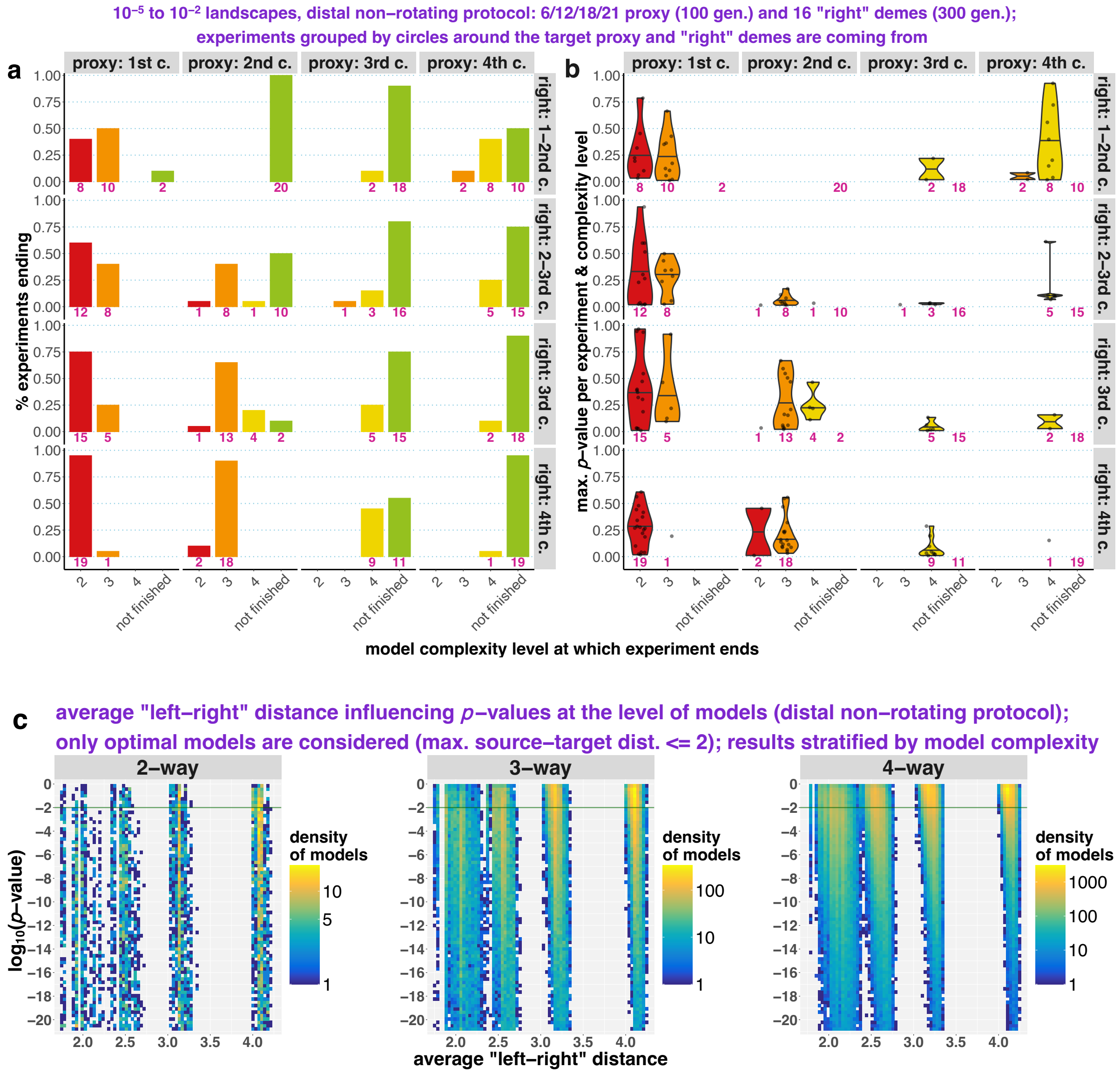

10⁻⁵ to 10⁻² landscapes, distal non–rotating protocol: 6/12/18/21 proxy (100 gen.) and 16 "right" demes (300 gen.);
experiments grouped by circles around the target proxy and "right" demes are coming from
a
b
proxy: 1st c.
proxy: 2nd c.
proxy: 3rd c.
proxy: 4th c.
right: 1–2nd c.
right: 2–3rd c.
right: 3rd c.
right: 4th c.
% experiments ending
max. p–value per experiment & complexity level
not finished
model complexity level at which experiment ends
c
average "left–right" distance influencing p–values at the level of models (distal non–rotating protocol);
only optimal models are considered (max. source–target dist. <= 2); results stratified by model complexity
2–way
3–way
4–way
log10(p–value)
density of models
average "left–right" distance

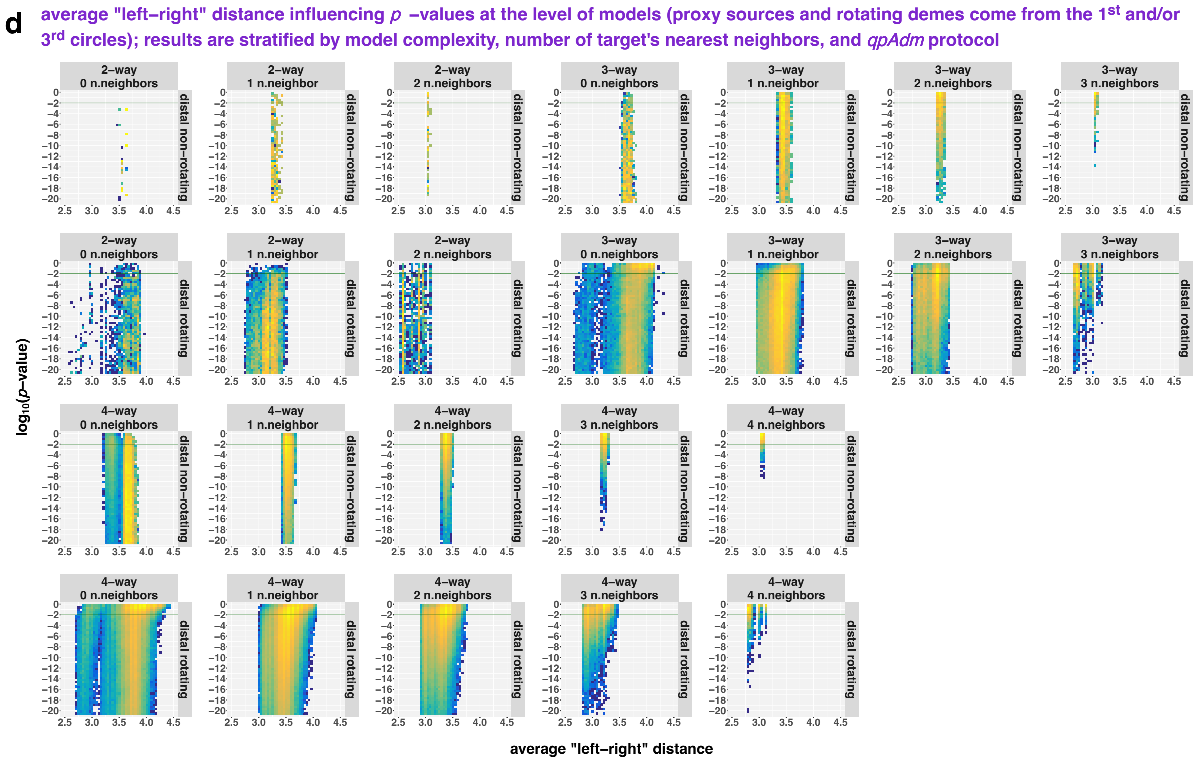


Illustrating effects of average “left-right” spatial distance on distributions of fitting *qpAdm* models by model complexity. Results are interpreted either at the level of experiments (**a, b**) or at the level of individual models (**c, d**). We used systematic SSL sampling and the distal non-rotating and rotating protocols relying on the following composite feasibility criterion: *p*-value threshold at 0.01, EAF between 0 and 1. In panel **a**, distributions of non-rotating experiments (**Fig. S16**) over model complexity levels at which they end are shown, and experiments are grouped by target-proxy distance (proxy sources came from the 1$^{st}$, or 2$^{nd}$, or 3$^{rd}$, or 4$^{th}$ circles of demes around two alternative targets in the center of SSL) and by target-“right” distance (16 “right” groups came either from the 1$^{st}$ and 2$^{nd}$, or 2$^{nd}$ and 3$^{rd}$, or 3$^{rd}$, or 4$^{th}$ circle around the targets). In panel **b**, each non-rotating experiment is represented by only one fitting model with the highest *p*-value. We show distributions of *p*-values for these models (the scale on the left) as violin plots stratified by three variables: target-“right” distance (on the right), target-proxy distance (on top), and model complexity level at which experiments end (the scale at the bottom). The pink numbers stand for the number of experiments in the respective categories in all simulation replicates combined. In panel **c**, results of the same systematic non-rotating protocol (**Fig. S16**) are presented at the level of two- to four-way models where sets of alternative proxy sources encompassed all demes at a distance of 1 or 2 from the two alternative targets. Models are placed in the coordinates of average “left-right” distance and *qpAdm* *p*-values (the latter axis is logarithmic and only the highest *p*-values $>10^{-21}$ are shown). Density of models in this space (in 75×75 bins) is visualized with the logarithmic color scales, and results are stratified by model complexity. For a given model, the shortest paths on the unweighted and undirected graph representation of SSL were calculated between all demes in the “left” set and all demes in the “right” set and averaged. In panel **d**, similar results are presented for the systematic non-rotating and rotating protocols (**Fig. S16**) with sets of alternative proxy sources and rotating sets composed of 16 demes distributed on the 1$^{st}$ and 3$^{rd}$ circles around 10 targets. Density of models in this space (in 50×50 bins) is visualized with logarithmic color scales (the scales in panels are independent). The results are stratified by model complexity, number of target's nearest neighbors in a model ("n.neighbors") and by *qpAdm* protocol.

**Figure S30.**

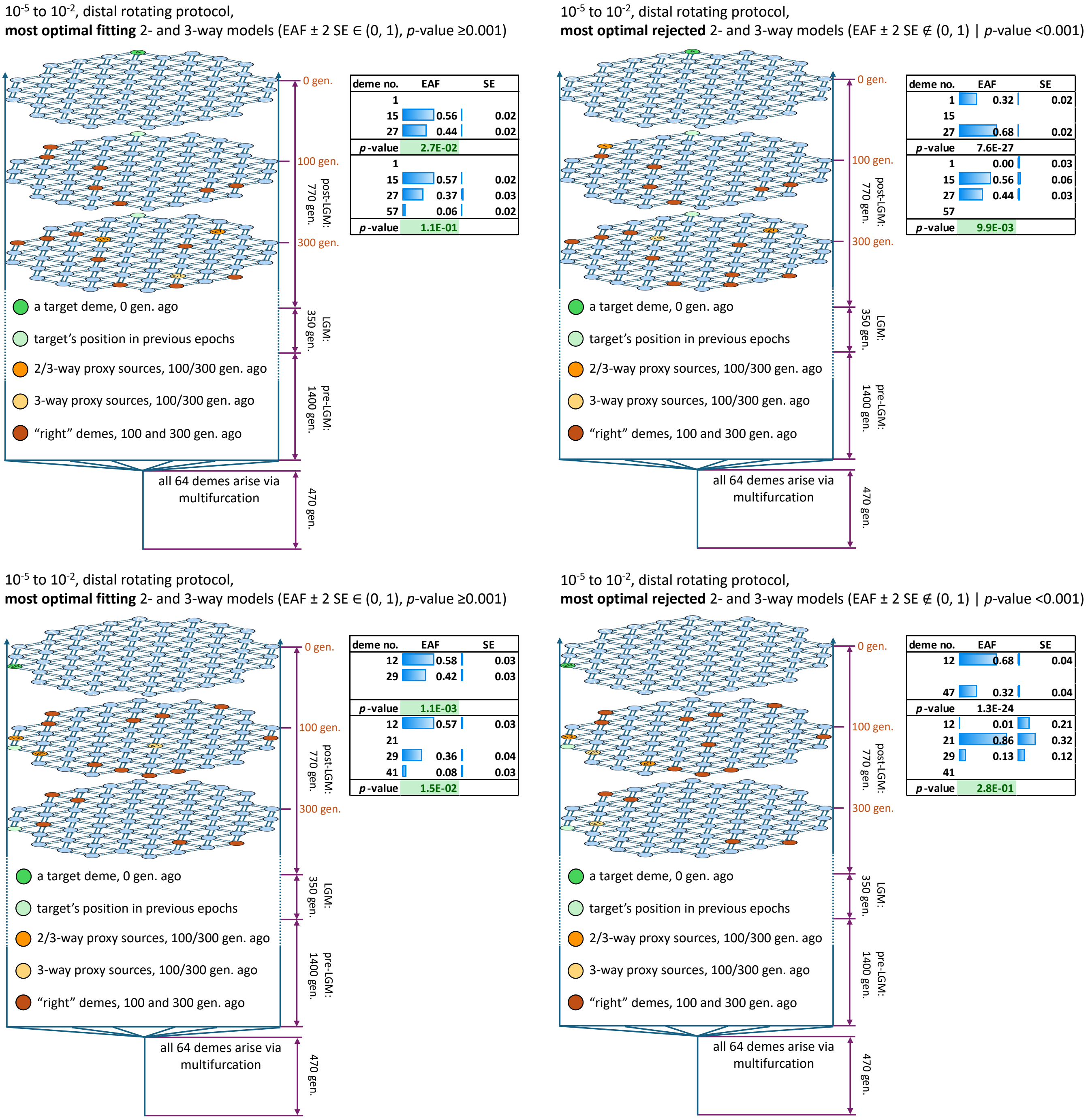


Diagrams summarizing potentially misleading outcomes of two *qpAdm* experiments when three-way models are considered instead of two-way models (two replicates of the "$10^{-5}$ to $10^{-2}$" landscape, distal rotating protocol, 18 demes in rotating sets). These experiments are used as examples of the large class of experiments with potentially misleading outcomes although, strictly speaking, the experiments end at two-way models (one such fitting model was found in each case) and should not have been evaluated at the stage of three-way models; however, the patterns shown here are common for all "bad" experiments. The experiments relied on a composite feasibility criterion that was deemed the best performing in this study (EAF ± 2 SE ∈ (0, 1), *p*-value ≥0.001, "trailing" models not checked). The diagrams on the left visualize deme arrangements in one most optimal two-way and one most optimal three-way fitting model in each experiment, and the diagrams on the right show deme arrangements in the corresponding most optimal non-fitting models. Model “optimality” was defined as Euclidean distance (in the space of optimality metrics) to the ideal symmetric model of the same complexity. Various categories of demes (targets, proxy sources in two- or three-way models, “right” sets) are color-coded and labelled according to the legends in each panel. The small tables beside each panel show EAF, corresponding SE, and *p*-values for the two- and three-way models visualized. *P*-values above the threshold are highlighted in green. Target and source demes are numbered both in the diagrams and tables.

**Figure S31.**

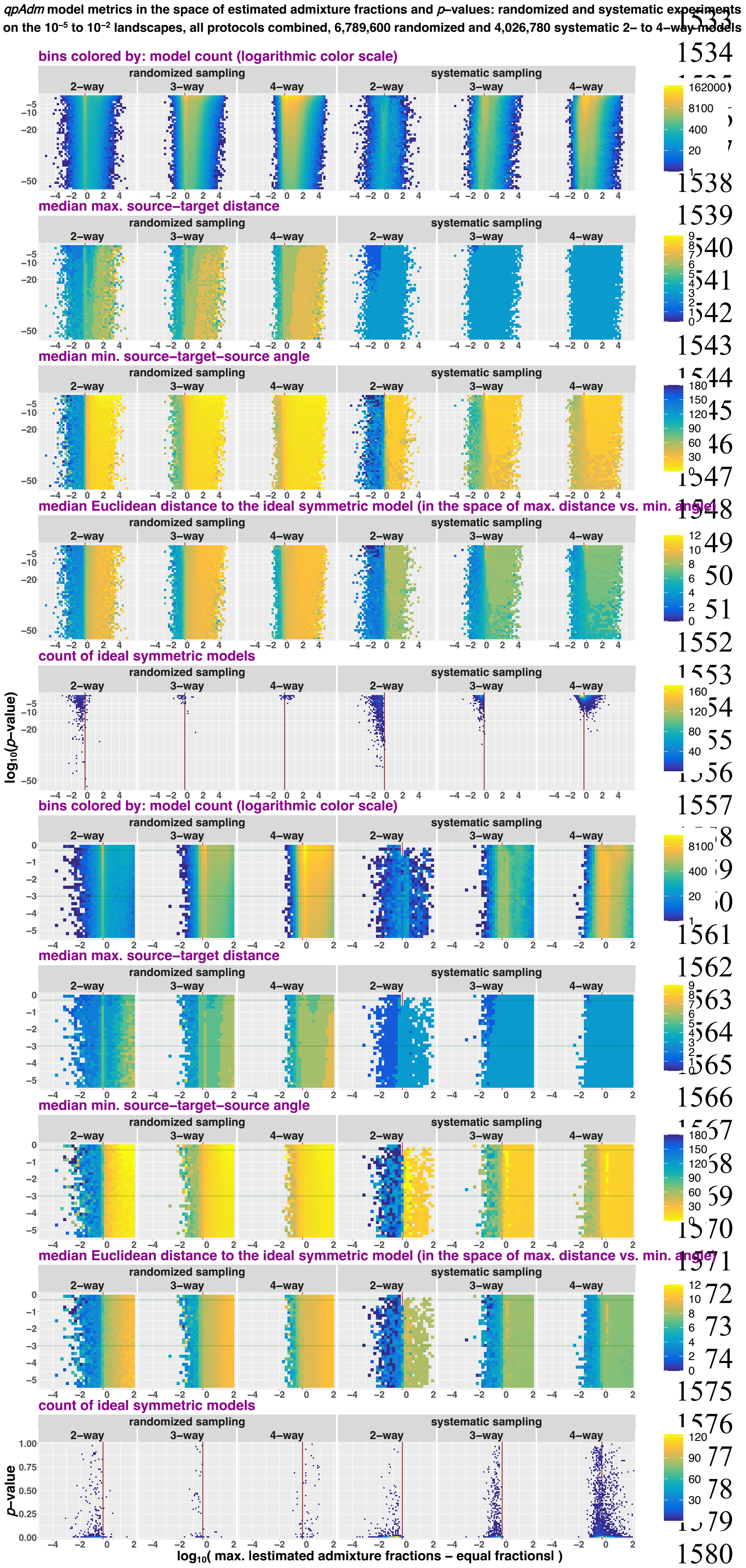


Comparing results of randomized and systematic deme sampling on the "$10^{-5}$ to $10^{-2}$" landscapes in the space of max|*EAF–EF*| and *p*-values (both axes are logarithmic). Results for all randomized or systematic *qpAdm* protocols are combined but stratified by model complexity. Two sections of the space are shown: *p*-values from $10^{-55}$ to 1 (the upper five rows of plots) and from $3.16\times10^{-6}$ to 1 (the four rows below them). The space is divided into rectangular bins (50×50 for the larger section and 35×35 for the smaller section), and they are colored by density of individual *qpAdm* models populating this space (logarithmic color scale) or by median values of model optimality metrics in those bins: max. ST distance, min. STS angle, and Euclidean distance to the ideal symmetric model. Density of ideal symmetric models (in the case of four-way models, the most optimal non-ideal models available on our SSL) in this space is also shown (in 75×75 bins). To assess if distributions of ideal symmetric models over *p*-values are uniform, we use non-logarithmic *p*-value scale (the bottom row of plots). The red vertical lines (or the red tick marks) mark max|*EAF–EF*| = 0.5, and the green horizontal lines mark the lowest and highest *p*-value thresholds used in this study (0.001 and 0.5).

#### **Figure S32.**

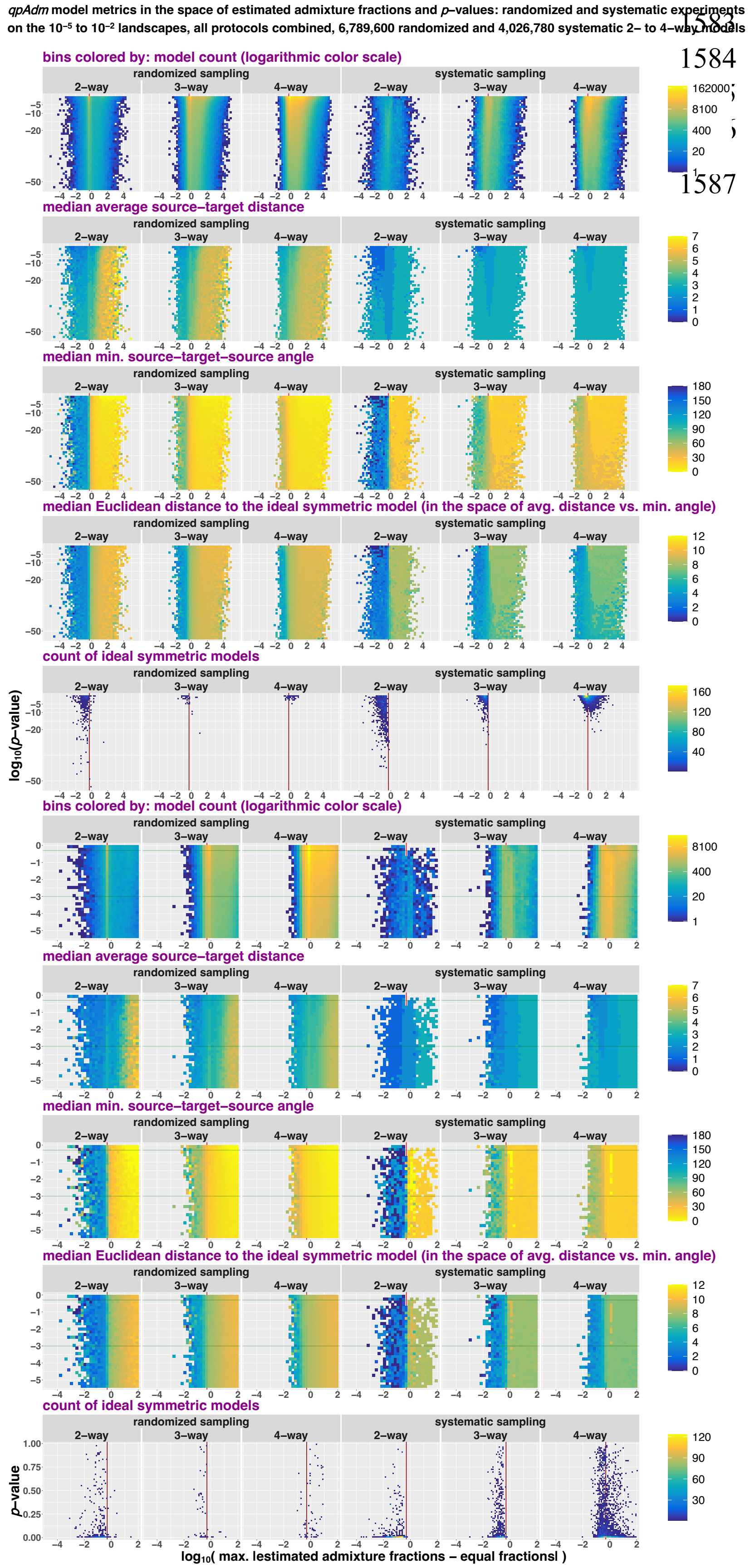


The same results as in **Fig. S31** but showing average ST distance instead of max. ST distance.

**Figure S33.**

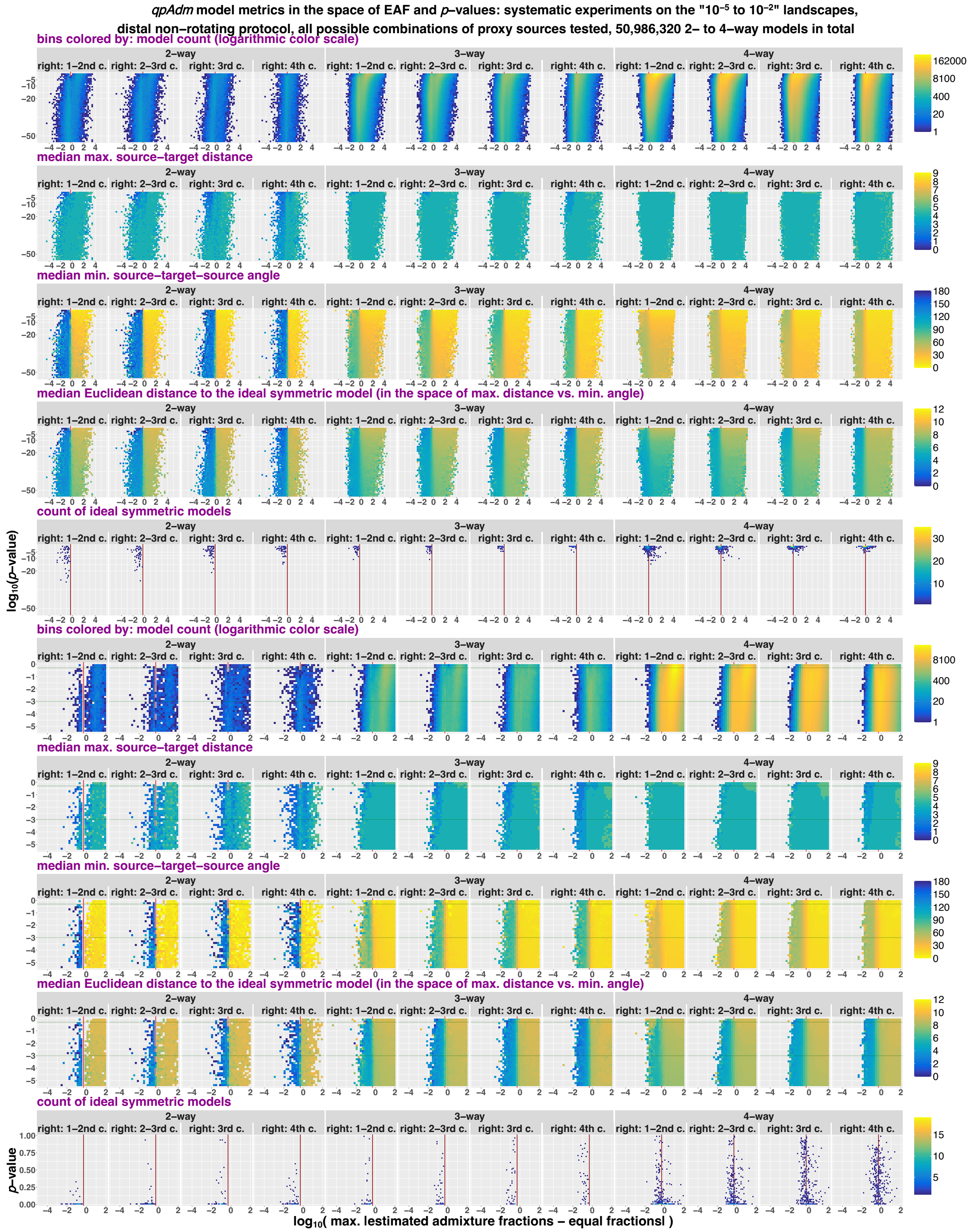


Systematic *qpAdm* results: the distal non-rotating protocol and all possible combinations of proxy sources tested for two targets (demes no. 32 and 33 in the center of the "$10^{-5}$ to $10^{-2}$" landscapes). Two to four-way *qpAdm* models are shown in the space of max|*EAF*–*EF*| and *p*-values (both axes are logarithmic). The results are stratified by model complexity and target-"right" distance: 16 "right" groups came either from the 1st and 2nd, or 2nd and 3rd, or 3rd, or 4th circle around the target. Two sections of the space are shown: *p*-values from $10^{-55}$ to 1 (the upper five rows of plots) and from $3.16\times10^{-6}$ to 1 (the four rows below them). The space is divided into rectangular bins (50×50 for the larger section and 35×35 for the smaller section), and they are colored by density of individual *qpAdm* models populating this space (logarithmic color scale) or by median values of model optimality metrics in those bins: max. ST distance, min. STS angle, and Euclidean distance to the ideal symmetric model. Density of ideal symmetric models (in the case of four-way models, the most optimal non-ideal models available on our SSL) in this space is also shown (in 75×75 bins). To assess if distributions of ideal symmetric models over *p*-values are uniform, we use non-logarithmic *p*-value scale (the bottom row of plots). The red vertical lines (or the red tick marks) mark max|*EAF–EF*| = 0.5, and the green horizontal lines mark the lowest and highest *p*-value thresholds used in this study (0.001 and 0.5).

1607 **Figure S34.**

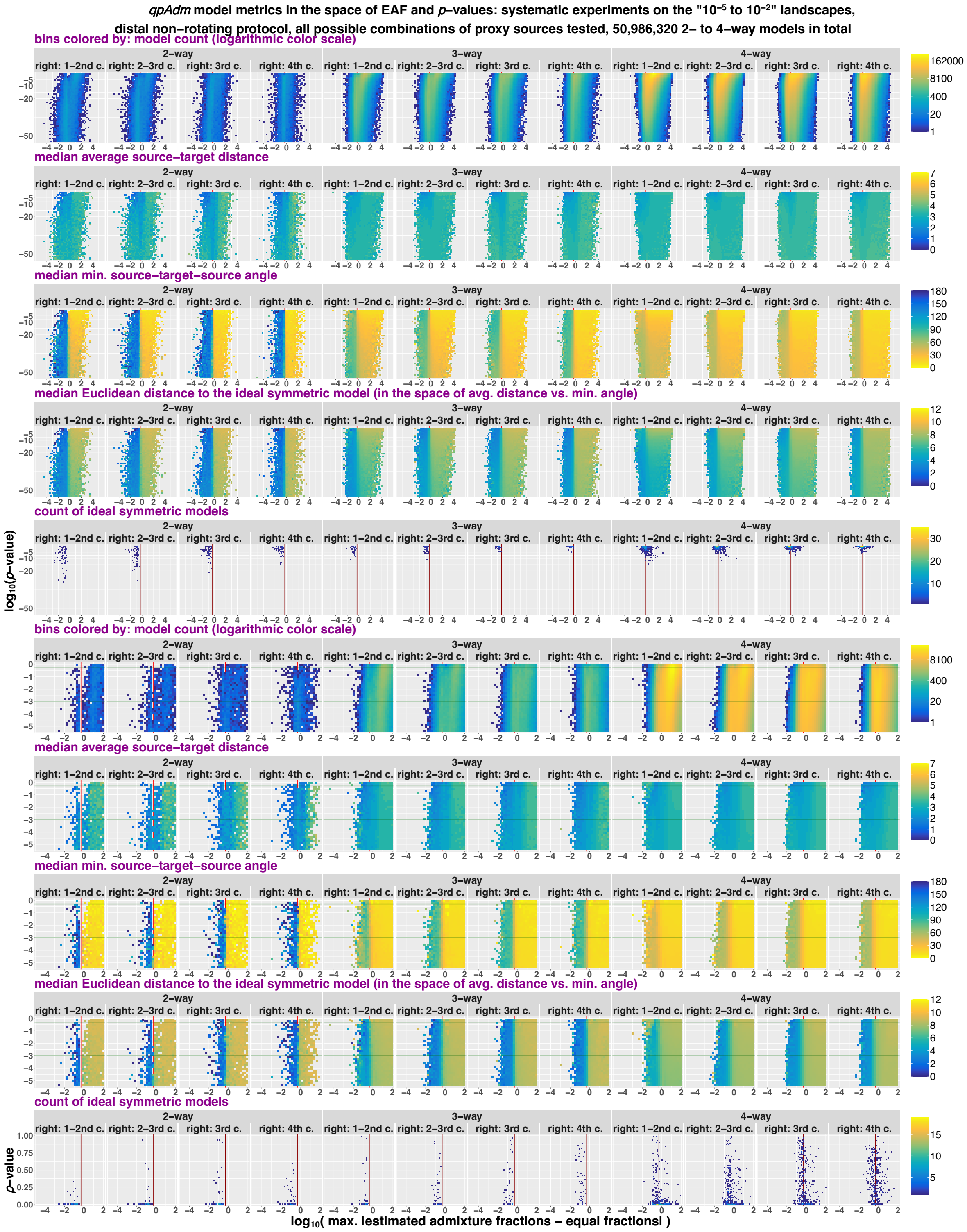


1608

1609 The same results as in **Fig. S33** but showing average ST distance instead of max. ST distance.

**Figure S35.**

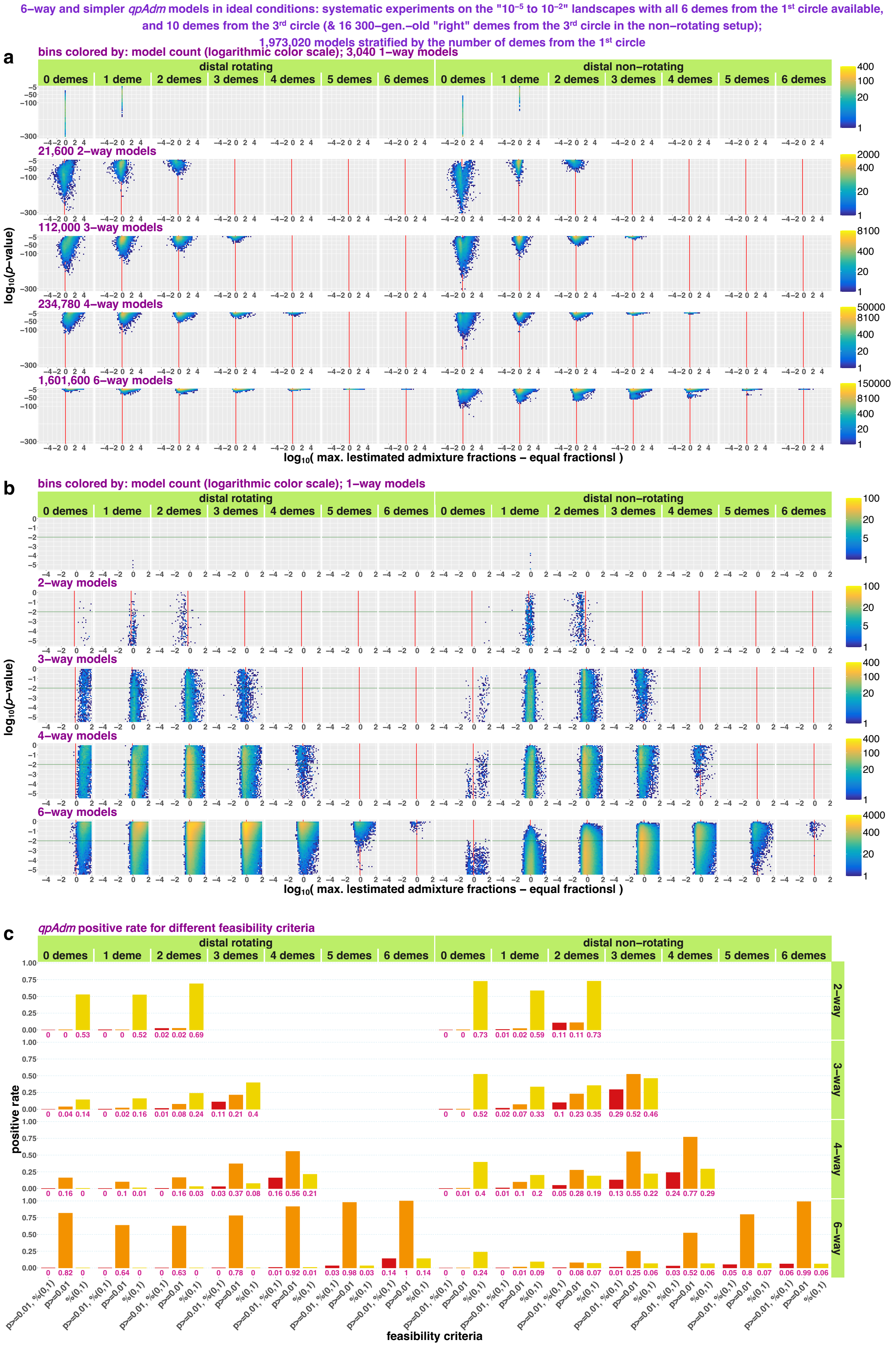

6–way and simpler qpAdm models in ideal conditions: systematic experiments on the "10⁻⁵ to 10⁻²" landscapes with all 6 demes from the 1st circle available,
and 10 demes from the 3rd circle (& 16 300–gen.–old "right" demes from the 3rd circle in the non–rotating setup);
1,973,020 models stratified by the number of demes from the 1st circle
a
bins colored by: model count (logarithmic color scale); 3,040 1–way models
distal rotating
distal non–rotating
0 demes
1 deme
2 demes
3 demes
4 demes
5 demes
6 demes
21,600 2–way models
112,000 3–way models
234,780 4–way models
1,601,600 6–way models
log10(p–value)
log10( max. |estimated admixture fractions – equal fractions| )
b
bins colored by: model count (logarithmic color scale); 1–way models
2–way models
3–way models
4–way models
6–way models
c
qpAdm positive rate for different feasibility criteria
positive rate
2–way
3–way
4–way
6–way
p>=0.01, %(0,1)
p>=0.01
%(0,1)
feasibility criteria

Systematic two to four- and six-way *qpAdm* models on the "$10^{-5}$ to $10^{-2}$" landscapes in the space of max|*EAF–EF*| and *p*-values (**a, b**). Results are stratified by model complexity, *qpAdm* protocol, and number of target's nearest neighbors in a model, which is the only optimality metric in this analysis. Two parts of the whole space are shown: *p*-values from $10^{-55}$ to 1 (**a**) and from $3.16\times10^{-6}$ to 1 (**b**). The space is divided into rectangular bins (50×50), and they are colored by density of individual *qpAdm* models populating this space (see the logarithmic color scales on the right). The red vertical lines (or the red tick marks) mark max|*EAF–EF*| = 1 – *EF*; for three-way and more complex models this boundary is similar but not identical to the boundary between EAF within (0, 1) and outside (0, 1). The green horizontal lines mark the *p*-value threshold used in all our systematic experiments (0.01). In panel **c**, we show as bar plots corresponding fractions of models that satisfy three kinds of feasibility criteria marked on the x-axis. The fractions are also indicated below the bars.

**Figure S36.**

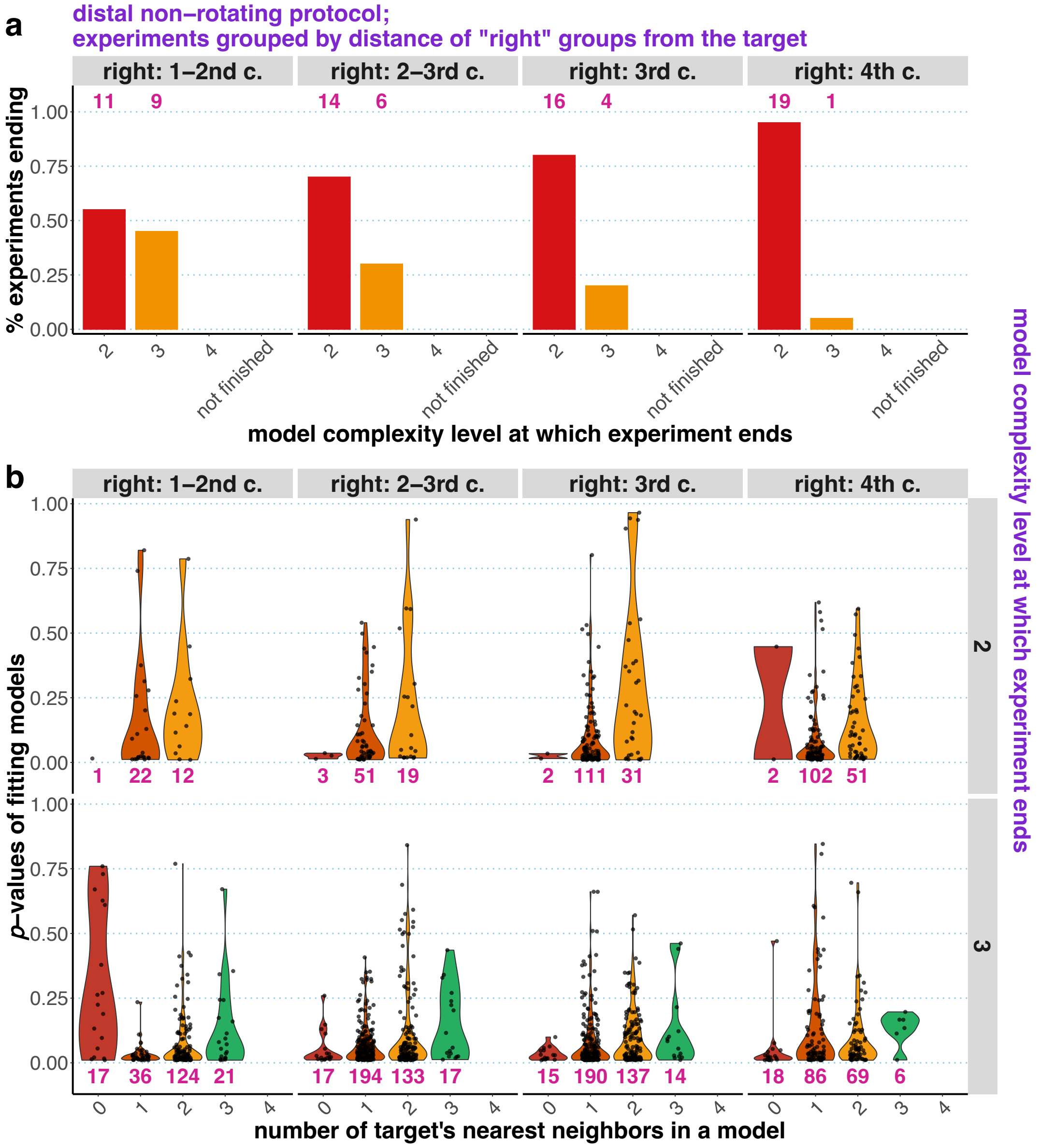


Systematic *qpAdm* results for the distal non-rotating protocol and all possible combinations of proxy sources interpreted at the level of experiments. The protocol relied on the following composite feasibility criterion: *p*-value threshold at 0.01, estimated admixture fractions between 0 and 1. In panel **a**, distributions of experiments over model complexity level at which they end are shown, and experiments are grouped by target-"right" distance: 16 "right" groups came either from the 1$^{st}$ and 2$^{nd}$, or 2$^{nd}$ and 3$^{rd}$, or 3$^{rd}$, or 4$^{th}$ circle around the target. In panel **b**, we show distributions of *p*-values for fitting models (the scale on the left) as violin plots stratified by three variables: model complexity level at which an experiment ends (on the right), target-"right" distance (on top), and number of target's nearest neighbors included in a model (the scale at the bottom). The pink numbers stand for the number of experiments (in panel **a**) or models (in panel **b**) in the respective categories in all simulation replicates combined.

**Figure S37.**

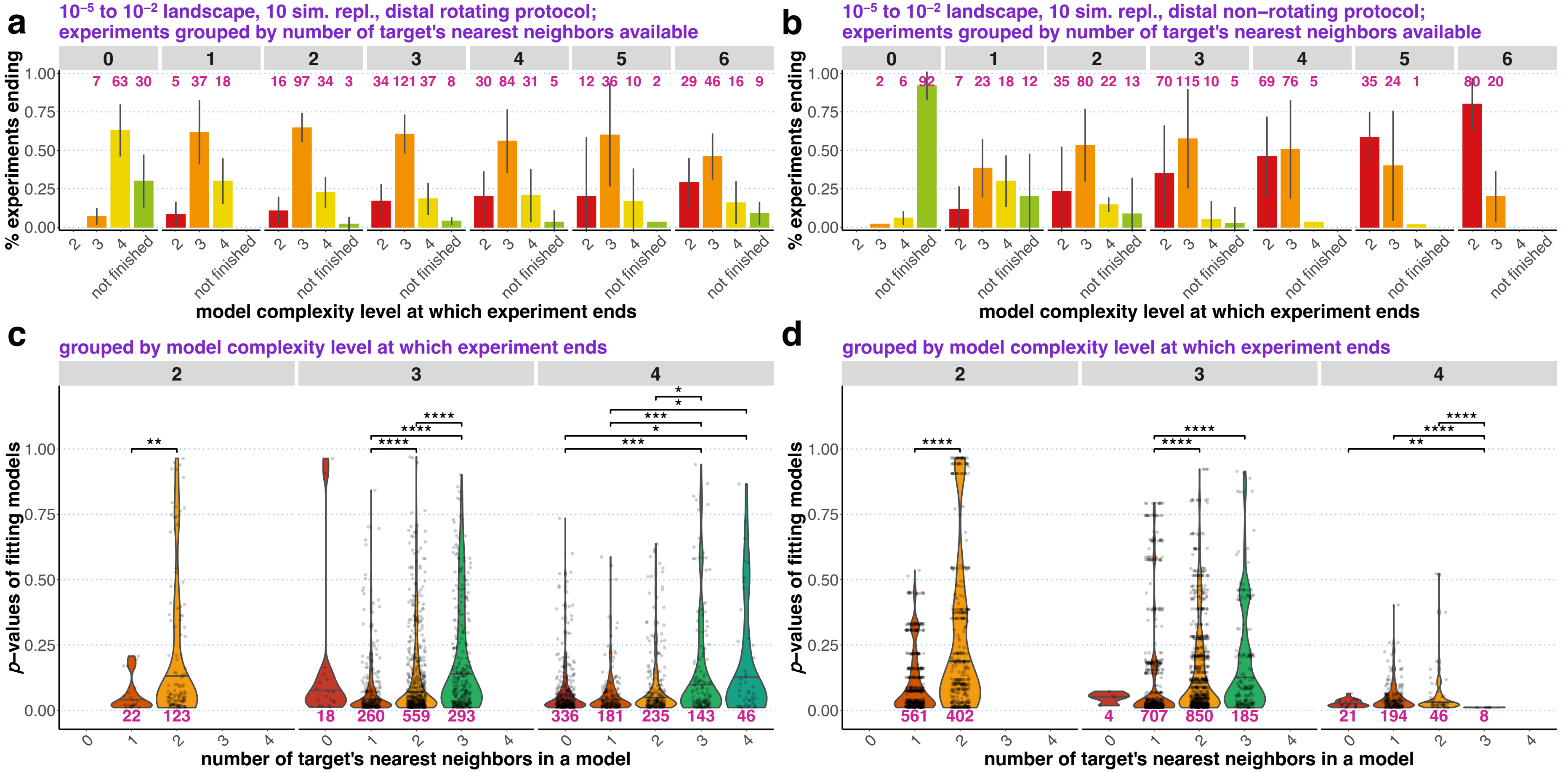


*qpAdm* performance in the case of systematic and symmetric landscape sampling, interpreted at the level of experiments. Results for the distal rotating (**a, c**) and distal non-rotating (**b, d**) protocols are shown. The protocols relied on the following composite feasibility criterion: *p*-value threshold at 0.01, estimated admixture fractions between 0 and 1. In panels **a** and **b**, distributions of experiments over model complexity level at which they end are shown, and experiments are grouped by the number of target's nearest neighbors (demes from the 1st circle) available. The other proxy sources and "right" groups were taken from the 3rd circle around the target. The error bars show standard deviation calculated on simulation replicates. In panels **c** and **d**, we show distributions of *p*-values for fitting models (the scale on the left) as violin plots with medians, stratified by two variables: model complexity level at which an experiment ends (the scale on top), and number of target's nearest neighbors included in a model (the scale at the bottom). For each model complexity level, all pairs of the distributions were compared using the two-sided non-paired Wilcoxon test, and only significant *p*-values (adjusted for multiple comparisons with the Holm method) are shown. The asterisks stand for the following significance levels: *, ≤0.05; **, ≤0.01; ***, ≤0.001; ****, ≤0.0001. The pink numbers stand for the number of experiments (in panels **a, b**) or models (in panels **c, d**) in the respective categories in all simulation replicates combined.

**Figure S38.**

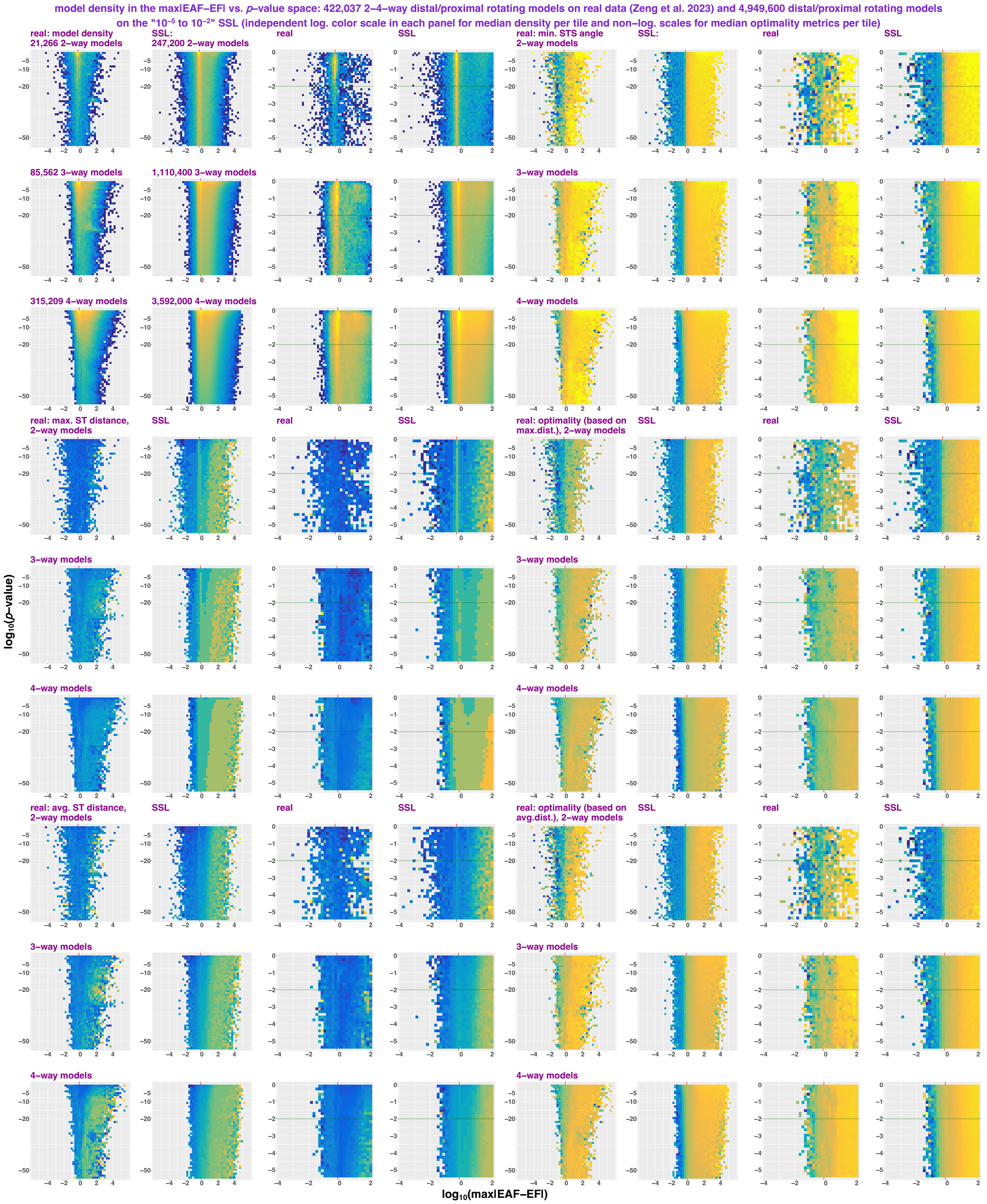


Comparing *qpAdm* models analyzed by the high-throughput protocol from Zeng *et al.* (2023) and results of the randomized *qpAdm* protocol on the "$10^{-5}$ to $10^{-2}$" landscapes. In both cases, two- to four-way models from distal and proximal rotating experiments are visualized in the space of max|*EAF–EF*| and *p*-values (both axes are logarithmic). Results are stratified by model complexity. Two sections of the space are shown: *p*-values from $10^{-55}$ to 1 (columns no. 1-2 and 5-6 from the left) and from $3.16\times10^{-6}$ to 1 (columns no. 3-4 and 7-8 from the left). The space is divided into rectangular bins (50×50 for the larger section and 35×35 for the smaller section), and they are colored by density of individual *qpAdm* models populating this space (logarithmic color scale) or by median values of model optimality metrics in those bins: max. ST distance, average ST distance, min. STS angle, and Euclidean distance to the ideal symmetric model ("optimality") in the space of max. or average ST distances and min. STS angles. The red vertical lines (or the red tick marks) mark max|*EAF*–*EF*| = 1 – *EF*; for three-way and more complex models this boundary is similar but not identical to the boundary between EAF within (0, 1) and outside (0, 1). The green horizontal lines mark the *p*-value threshold of 0.01. The distances and angles on the real data (great circle distances in km and angles between two bearings) are based on centroids of groups calculated using an *R* implementation of the *Mean Center* tool from *ArcGIS Pro*. We considered ST distances ≤50 km to be negligible, and for such models min. STS angles and hence distances to the ideal symmetric models were not defined, following the approach we applied to zero-length ST distances on the simulated data.

**Figure S39.**

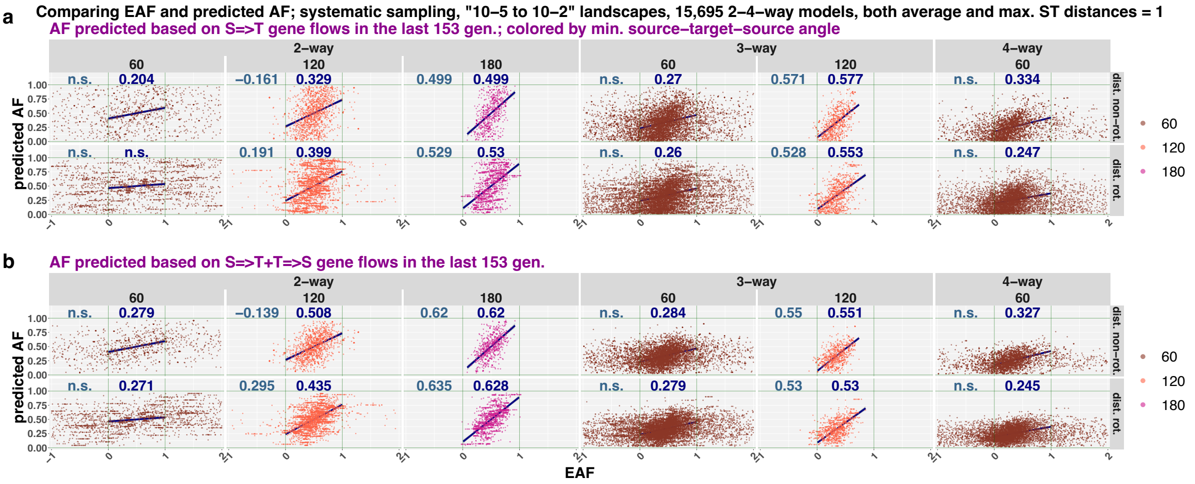


Scatterplots illustrating correlations between EAF and predicted AF on systematically sampled “$10^{-5}$ to $10^{-2}$” landscapes. Models composed of target’s nearest neighbors only were considered, and AF were predicted based on simulated source-to-target gene-flow intensities or on sums of source-to-target and target-to-source gene-flow intensities (see the text for details). Each *n*-way model in these scatterplots is represented by *n* points, one per source. The results are stratified by model complexity, by min. STS angle, and *qpAdm* protocol. Pearson’s correlation coefficients were calculated for EAF vs. predicted AF: for EAF between 0 and 1 (the numbers in navy blue above the plots) or across the whole range of EAF (the numbers in slate gray; however, only models with EAF between -1 and 2 are shown). Non-significant correlation coefficients are marked as “n.s.” Linear trend lines and 95% confidence intervals are also shown for EAF between 0 and 1.

**Figure S40.**

**a**

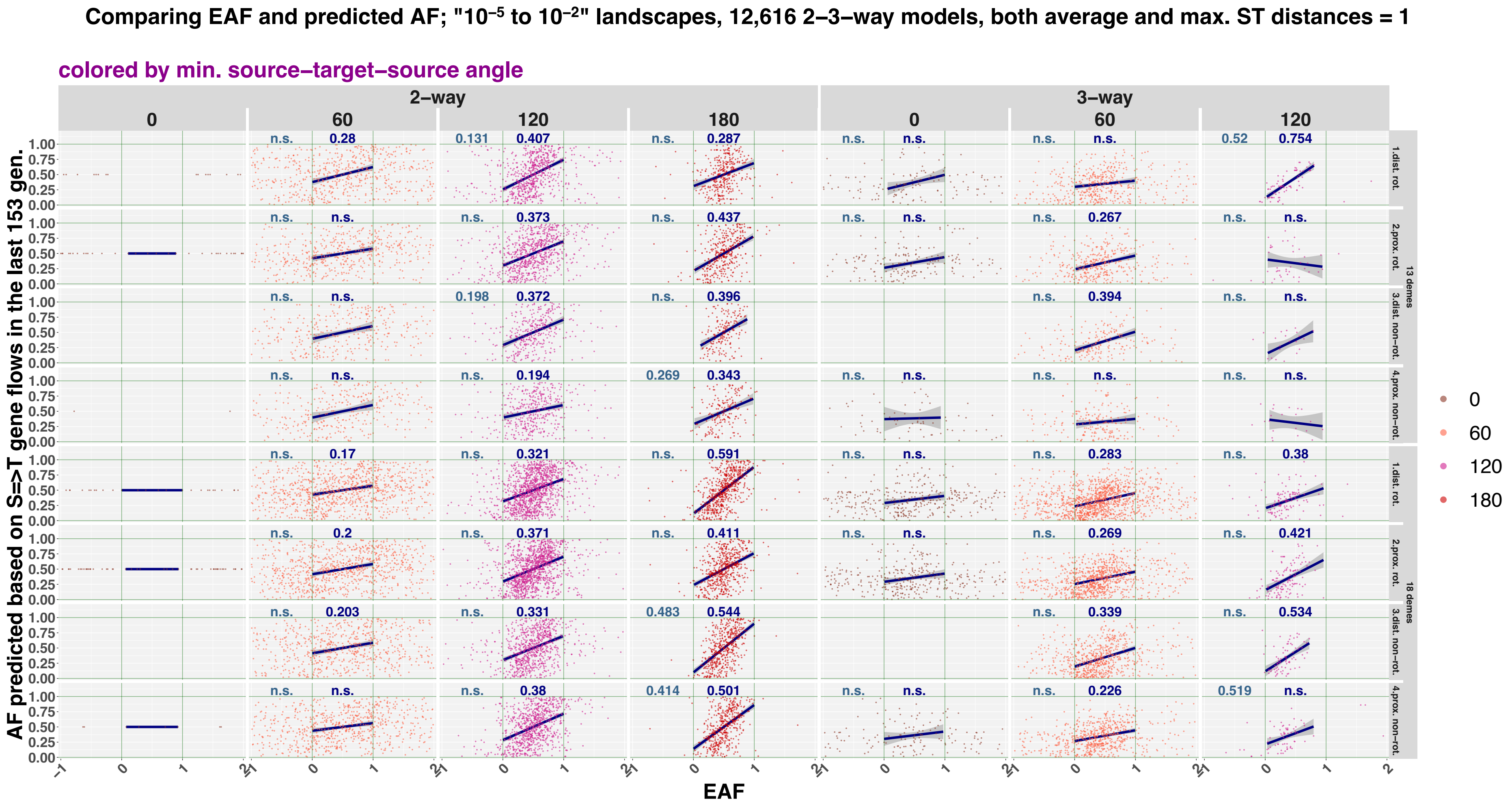

Comparing EAF and predicted AF; "10⁻⁵ to 10⁻²" landscapes, 12,616 2–3–way models, both average and max. ST distances = 1
colored by min. source–target–source angle
2–way
3–way
0
60
120
180
AF predicted based on S=>T gene flows in the last 153 gen.
EAF
1.dist. rot.
2.prox. rot.
3.dist. non–rot.
4.prox. non–rot.
13 demes
18 demes

**b**

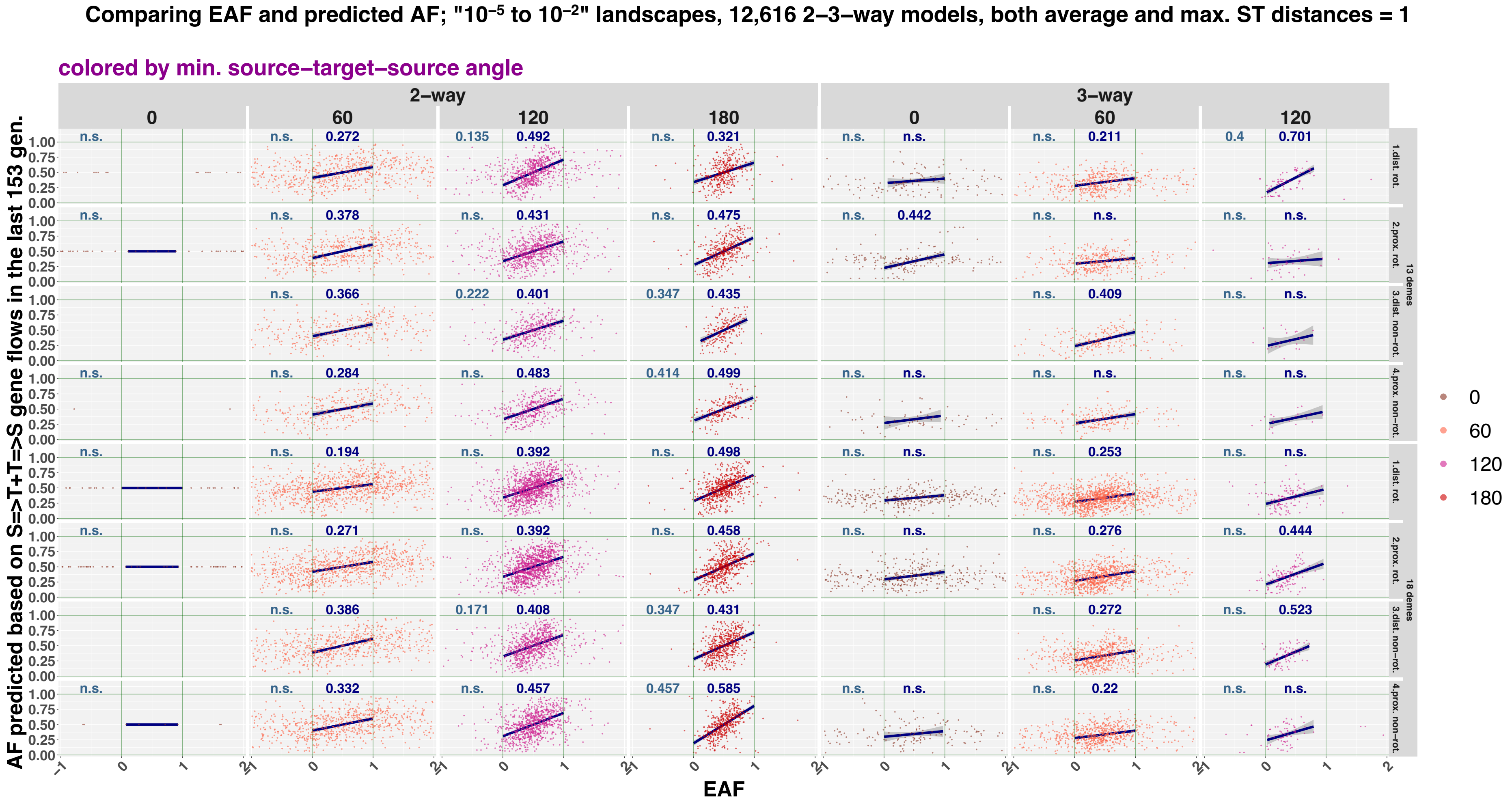


Scatterplots illustrating correlations between EAF and predicted AF on randomly sampled “$10^{-5}$ to $10^{-2}$” landscapes. Models composed of target’s nearest neighbors only were considered, and AF were predicted based on simulated source-to-target gene-flow intensities (**a**) or on sums of source-to-target and target-to-source gene-flow intensities (**b**), see the text for details. Each *n*-way model in these scatterplots is represented by *n* points, one per source. The results are stratified by model complexity, by min. STS angle, *qpAdm* protocol, and landscape sampling density (13 or 18 demes in rotating sets and 10 or 15 demes in non-rotating "right" sets, respectively). Pearson's correlation coefficients were calculated for EAF vs. predicted AF: for EAF between 0 and 1 (the numbers in navy blue above the plots) or across the whole range of EAF (the numbers in slate gray; however, only models with EAF between -1 and 2 are shown). Non-significant correlation coefficients are marked as "n.s." Linear trend lines and 95% confidence intervals are also shown for EAF between 0 and 1.

**Table S1.**

**a**

| | | no. admixture events in target's history | model class: consistently negative | model class: occasionally fitting and false | model class: occasionally fitting and true | model class: consistently negative | model class: occasionally fitting and false | model class: occasionally fitting and true |
|---|---|---|---|---|---|---|---|---|
| proximal | rotating | 0 | 10,468 | 1,990 | 0 | | | |
| | | 1 | 4,390 | 576 | 325 | | | |
| | | 2 | 1,766 | 277 | 229 | | | |
| | | 3 | 536 | 109 | 65 | | | |
| | | 4 | 155 | 10 | 26 | | | |
| | | 5 | 20 | 10 | 0 | | | |
| | | | 82.7% | 14.2% | 3.1% | | | |
| | non-rotating | 0 | 388 | 66 | 0 | | | |
| | | 1 | 645 | 138 | 56 | | | |
| | | 2 | 552 | 116 | 82 | | | |
| | | 3 | 173 | 51 | 40 | | | |
| | | 4 | 64 | 9 | 4 | | | |
| | | 5 | 4 | 8 | 0 | | | |
| | | | 76.2% | 16.2% | 7.6% | | | |
| distal | rotating | 0 | 1,659 | 73 | 0 | | | |
| | | 1 | 2,371 | 197 | 193 | | | |
| | | 2 | 3,655 | 300 | 505 | | | |
| | | 3 | 1,973 | 167 | 318 | | | |
| | | 4 | 1,294 | 69 | 294 | | | |
| | | 5 | 235 | 25 | 40 | | | |
| | | | 83.7% | 6.2% | 10.1% | | | |
| | non-rotating | 0 | 65 | 6 | 0 | | | |
| | | 1 | 129 | 48 | 34 | | | |
| | | 2 | 474 | 87 | 114 | | | |
| | | 3 | 306 | 59 | 76 | | | |
| | | 4 | 227 | 30 | 86 | | | |
| | | 5 | 41 | 6 | 16 | | | |
| | | | 68.8% | 13.1% | 18.1% | | | |
| | | | | | total: | 82.0% | 11.5% | 6.5% |

**b**

| temporal stratification | | distal | | proximal | | |
|---|---|---|---|---|---|---|
| | *qpAdm* protocol | non-rotating | rotating | non-rotating | rotating | |
| number of models: unique target/source/"right"/AGS topology sets^ | total | 1,242 | 11,187 | 1,826 | 17,335 | consistently negative |
| | | 236 | 831 | 388 | 2,972 | occasionally fitting and false |
| | | 326 | 1,350 | 182 | 645 | occasionally fitting and true |
| target sampling date, generations | median | 0 | 364 | 679 | 1331 | consistently negative* |
| | | 38 | 334 | 578 | 1436 | occasionally fitting and false* |
| | | 0 | 158 | 666 | 821 | occasionally fitting and true* |
| | average | 225 | 460 | 713 | 1277 | consistently negative* |
| | | 244 | 435 | 608 | 1412 | occasionally fitting and false* |
| | | 183 | 313 | 613 | 864 | occasionally fitting and true* |
| number of admixture events in target's history | median | 2 | 2 | 1 | 0 | consistently negative** |
| | | 2 | 2 | 1 | 0 | occasionally fitting and false** |
| | | 3 | 2 | 2 | 1 | occasionally fitting and true** |
| | average | 2.5 | 2.0 | 1.4 | 0.6 | consistently negative** |
| | | 2.3 | 2.0 | 1.5 | 0.5 | occasionally fitting and false** |
| | | 2.8 | 2.6 | 2.0 | 1.7 | occasionally fitting and true** |
| number of admixture events in target's history – average number of admixture events in history of proxy sources | median | 1 | 1 | -1 | -1 | consistently negative*** |
| | | 0.5 | 1 | -0.75 | -1 | occasionally fitting and false*** |
| | | 1.5 | 2 | -0.25 | 0 | occasionally fitting and true*** |
| | average | 0.9 | 1.2 | -0.8 | -0.9 | consistently negative*** |
| | | 0.6 | 1.1 | -0.7 | -0.8 | occasionally fitting and false*** |
| | | 1.3 | 1.8 | -0.1 | 0.2 | occasionally fitting and true*** |

^ "non-rotating" models (105 per topology and 4,200 in total) were tested by both protocols (with "right" sets being different) and thus were counted twice

* all but one *p*-value of the two-sided non-paired Wilcoxon test <0.044 (adjusted for multiple testing using the Holm method); model classes were compared

** all *p*-values of the two-sided non-paired Wilcoxon test <2.21e-8 (adjusted for multiple testing using the Holm method); model classes were compared

*** all *p*-values of the two-sided non-paired Wilcoxon test <8.82e-4 (adjusted for multiple testing using the Holm method); model classes were compared

**c**

| variable 1: | | number of admixture events in target's history – average number of admixture events in history of proxy sources | | | | | | | |
|---|---|---|---|---|---|---|---|---|---|
| | amount of data | 3,000-Mbp-sized genomes, 3,000 gen. divergence time | | | | | | | |
| | temporal stratification | distal | | | | proximal | | | |
| | *qpAdm* protocol | non-rotating | | rotating | | non-rotating | | rotating | |
| variable 2: | data quality | high quality | noisy | high quality | noisy | high quality | noisy | high quality | noisy |
| **max\|*EAF–EF*\|** | **p >= 1e-55** | 0.13 | 0.09 | 0.04 | -0.02 | 0.19 | 0.01 | -0.11 | -0.11 |
| | **p >= 1e-5** | 0.24 | 0.16 | 0.14 | 0.06 | 0.30 | 0.24 | 0.00 | -0.09 |
| | **max\|*EAF–EF*\| <= 0.5** | -0.08 | -0.10 | -0.10 | -0.12 | -0.16 | -0.13 | -0.05 | -0.02 |
| | **p >= 1e-55; max\|*EAF–EF*\| <= 0.5** | -0.46 | -0.21 | -0.45 | -0.36 | -0.06 | -0.11 | -0.24 | -0.05 |
| | **p >= 1e-5; max\|*EAF–EF*\| <= 0.5** | -0.49 | -0.43 | -0.22 | -0.39 | 0.07 | -0.10 | -0.17 | -0.18 |
| ***p*-value** | **p >= 1e-55** | -0.16 | -0.07 | 0.03 | 0.01 | -0.16 | -0.19 | -0.03 | 0.03 |
| | **p >= 1e-5** | -0.08 | -0.11 | 0.03 | 0.02 | -0.02 | -0.10 | -0.03 | -0.05 |
| | **max\|*EAF–EF*\| <= 0.5** | -0.11 | -0.10 | -0.14 | -0.10 | -0.19 | -0.18 | 0.00 | 0.03 |
| | **p >= 1e-55; max\|*EAF–EF*\| <= 0.5** | -0.32 | -0.15 | -0.41 | -0.40 | -0.07 | -0.10 | -0.21 | -0.01 |
| | **p >= 1e-5; max\|*EAF–EF*\| <= 0.5** | -0.18 | -0.20 | -0.10 | -0.27 | 0.12 | -0.08 | -0.07 | -0.16 |

*p*-value > 0.05/80 (Bonferroni correction)

Quantifying complexity of the AGS simulated histories as number of admixture events on all paths from a target population to the root of the simulation (or, in short, number of admixture events in target's history). (**a**) Distributions of unique models tested over complexities of their target's histories; the results are stratified by distal/proximal models, by *qpAdm* protocols (rotating/non-rotating), and by model classes (consistently negative or non-fitting in magenta, occasionally fitting and false in red, occasionally fitting and true in green). Blue bars in the right half of the table visualize fractions in the whole dataset, while magenta, red, and green bars show fractions within each category separately. Relative sizes of the three model classes in percentages are also shown. The median number of admixture events in target's history for the whole dataset (1) is highlighted in bold. (**b**) Comparing other metrics informative about admixture history in the three *qpAdm* model classes: consistently negative or non-fitting, occasionally fitting and false, and occasionally fitting and true models. Median and average values of target sampling dates, numbers of admixture events in target's history, and differences between number of admixture events in target's history and average number of admixture events in sources' histories ($AEH_T$ – *av.* $AEH_S$) are shown. The results are stratified by distal/proximal models and by *qpAdm* protocols (rotating/non-rotating). Footnotes show results of the two-sided non-paired Wilcoxon test comparing the metrics across the model classes. The size of those model classes is presented too. (**c**) Spearman's correlation coefficient (ρ) between the $AEH_T$ – *av.* $AEH_S$ metric and max|*EAF–EF*| or *p*-values in various sections of the max|*EAF–EF*| vs. *p*-value space (listed on the left). Results are shown for the 3,000-Mbp simulations and stratified by distal/proximal models, by *qpAdm* protocols (rotating/non-rotating), and by data quality levels. Non-significant ρ values (with Bonferroni correction) are underlined.

**Table S2.**

| simulation scheme | max. depth of the simulation, gen. | pseudohaploid noisy data | genome size, Mbp | feasible *qpAdm* models/ all 2-way models | two-way models supported by: | | | | |
|---|---|---|---|---|---|---|---|---|---|
| | | | | proximal rotating *qpAdm* | 1 | 1 | 1 | 1 | 1 |
| | | | | "admixture" *f3*- statistics | | 2 | | | |
| | | | | 3D PCA | | | 2 | | 2 |
| | | | | unsupervised *ADMIXTURE* | | | | 2 | 3 |
| date variable, no. 1 | 800 | no | 300 | 385/ 34,320 (1.1%) | 55.1% | 0.5% | 33.5% | 15.3% | 10.9% |
| | | | | | | | | | 22.6% |
| | | | | | | 54.5% | 21.6% | 39.7% | 4.4% |
| | | | | | | | | | 17.1% |
| | | | | | 44.9% | 2.1% | 21.0% | 15.6% | 10.6% |
| | | | | | | | | | 10.4% |
| | | | | | | 42.9% | 23.9% | 29.4% | 4.9% |
| | | | | | | | | | 19.0% |
| | | | | FDR | 55.1% | 20.0% | 61.4% | 49.6% | 50.6% |
| | | | | FOR | 0% | 95.4% | 53.2% | 65.3% | 76.3% |
| date variable (no. 2) | 800 | no | 1000 | 198/ 34,320 (0.6%) | 57.6% | 0.5% | 42.4% | 9.1% | 8.1% |
| | | | | | | | | | 34.3% |
| | | | | | | 57.1% | 15.2% | 48.5% | 1.0% |
| | | | | | | | | | 14.1% |
| | | | | | 42.4% | 3.0% | 17.7% | 13.1% | 10.1% |
| | | | | | | | | | 7.6% |
| | | | | | | 39.4% | 24.7% | 29.3% | 3.0% |
| | | | | | | | | | 21.7% |
| | | | | FDR | 57.6% | 14.3% | 70.6% | 40.9% | 44.4% |
| | | | | FOR | 0% | 92.9% | 58.3% | 69.0% | 76.2% |
| date variable, no. 4 | 3000 | no | 300 | 92/ 34,320 (0.3%) | 66.3% | 1.1% | 44.6% | 13.0% | 13.0% |
| | | | | | | | | | 31.5% |
| | | | | | | 65.2% | 21.7% | 53.3% | 0.0% |
| | | | | | | | | | 21.7% |
| | | | | | 33.7% | 2.2% | 13.0% | 15.2% | 8.7% |
| | | | | | | | | | 4.3% |
| | | | | | | 31.5% | 20.7% | 18.5% | 6.5% |
| | | | | | | | | | 14.1% |
| | | | | FDR | 66.3% | 33.3% | 77.4% | 46.2% | 60.0% |
| | | | | FOR | 0% | 93.5% | 61.3% | 54.8% | 74.2% |
| at "present", no. 5 | 800 | no | 300 | 569/ 34,320 (1.7%) | 64.0% | 0.4% | 31.3% | 10.4% | 7.9% |
| | | | | | | | | | 23.4% |
| | | | | | | 63.6% | 32.7% | 53.6% | 2.5% |
| | | | | | | | | | 30.2% |
| | | | | | 36.0% | 0.0% | 21.3% | 17.0% | 11.1% |
| | | | | | | | | | 10.2% |
| | | | | | | 36.0% | 14.8% | 19.0% | 6.0% |
| | | | | | | | | | 8.8% |
| | | | | FDR | 64.0% | N/A | 59.5% | 37.8% | 41.7% |
| | | | | FOR | 0% | 100.0% | 41.0% | 52.7% | 69.3% |

Assessing FDR of the proximal rotating *qpAdm* protocol and of more complex admixture inference pipelines based on it. For each simulation setup, the analysis relies on simulation replicate no. 1 only since PCA and *ADMIXTURE* results were not generated for multiple replicates (see Discussion). For constructing the pipelines, we used three additional methods: "admixture" $f_3$-statistics, 3D PCA with all individuals co-analyzed, and unsupervised *ADMIXTURE* with all individuals co-analyzed. In each pipeline, for declaring a positive result support of an admixture model by all methods was required, hence the order of methods is not important except for the first method, which was proximal rotating *qpAdm* in all cases. In each column, methods comprising a pipeline are color-coded and numbered by their order. All feasible two-way *qpAdm* models emerging as outcomes of the proximal rotating protocol were classified into FP and TP (highlighted in red and green in the leftmost column, respectively). The other columns are structured like bifurcating trees: FP *qpAdm* models supported by method no. 2; FP *qpAdm* models not supported by method no. 2; TP *qpAdm* models supported by method no. 2; TP *qpAdm* models not supported by method no. 2. The same principle is used for representing results of more complex pipelines. All model counts are normalized by the number of feasible *qpAdm* models (FP + TP), outcomes of the first method. Percentages of models supported/not supported by the last method in the pipeline are highlighted in green and red, respectively. FDR values are shown for these different pipelines. Fractions of TP *qpAdm* models that are pruned out by progressively more stringent support requirements are also shown (false omission rate or FOR).

**Table S3.**

| | | 0.01 | | | | 0.1 | | | | *p*-value theshold for 2-way models |
|---|---|---|---|---|---|---|---|---|---|---|
| **simulation setup** | **model supported by:** | **0.01** | **1.E-10** | **1.E-20** | **1.E-50** | **0.01** | **1.E-10** | **1.E-20** | **1.E-50** | ***p*-value theshold for 1-way models** |
| sampling dates variable, 1,000 Mbp | *qpAdm* | 57.6% | 46.2% | 45.5% | 44.1% | 54.9% | 44.1% | 44.3% | 38.0% | |
| sampling dates variable, 300 Mbp, noisy data | *qpAdm* | 53.3% | 44.1% | 41.1% | 38.7% | 48.0% | 37.4% | 33.6% | 32.6% | |
| sampling dates variable, 300 Mbp | *qpAdm* | 55.1% | 50.2% | 47.6% | 43.7% | 51.9% | 44.4% | 41.1% | 33.3% | |
| | *qpAdm* + PCA | 61.4% | 53.8% | 49.3% | 43.6% | 60.0% | 51.3% | 47.8% | 41.5% | |
| | *qpAdm* + PCA + *ADMIXTURE* | 50.6% | 43.7% | 37.7% | 35.4% | 50.0% | 42.1% | 36.4% | 32.1% | **false discovery rate (FDR)** |
| sampling dates variable, deep divergence, 300 Mbp | *qpAdm* | 66.3% | 58.7% | 58.6% | 58.7% | 68.6% | 59.3% | 60.0% | 59.1% | |
| | *qpAdm* + PCA | 77.4% | 72.7% | 72.1% | 71.1% | 73.1% | 68.2% | 66.7% | 63.2% | |
| | *qpAdm* + PCA + *ADMIXTURE* | 60.0% | 57.9% | 55.6% | 50.0% | 50.0% | 50.0% | 44.4% | 28.6% | |
| sampled at "present", 300 Mbp | *qpAdm* | 64.0% | 56.8% | 50.8% | 50.5% | 55.9% | 48.8% | 44.5% | 39.0% | |
| | *qpAdm* + PCA | 59.5% | 52.9% | 49.2% | 44.1% | 53.2% | 45.1% | 43.4% | 32.2% | |
| | *qpAdm* + PCA + *ADMIXTURE* | 41.7% | 36.9% | 35.4% | 37.3% | 36.1% | 30.0% | 29.8% | 26.7% | |
| sampling dates variable, 1,000 Mbp | *qpAdm* | 0.0% | 8.3% | 14.3% | 21.4% | 51.2% | 54.8% | 59.5% | 63.1% | |
| sampling dates variable, 300 Mbp, noisy data | *qpAdm* | 0.0% | 37.8% | 53.6% | 75.0% | 46.6% | 66.7% | 74.8% | 86.7% | |
| sampling dates variable, 300 Mbp | *qpAdm* | 0.0% | 15.0% | 24.3% | 35.8% | 48.0% | 57.2% | 61.8% | 66.5% | |
| | *qpAdm* + PCA | 53.2% | 57.2% | 60.1% | 67.1% | 74.6% | 77.5% | 79.2% | 82.1% | |
| | *qpAdm* + PCA + *ADMIXTURE* | 76.3% | 76.9% | 78.0% | 82.1% | 86.7% | 87.3% | 87.9% | 89.0% | **false omission rate (FOR)** |
| sampling dates variable, deep divergence, 300 Mbp | *qpAdm* | 0.0% | 0.0% | 6.5% | 16.1% | 64.5% | 64.5% | 67.7% | 71.0% | |
| | *qpAdm* + PCA | 61.3% | 61.3% | 61.3% | 64.5% | 77.4% | 77.4% | 77.4% | 77.4% | |
| | *qpAdm* + PCA + *ADMIXTURE* | 74.2% | 74.2% | 74.2% | 74.2% | 83.9% | 83.9% | 83.9% | 83.9% | |
| sampled at "present", 300 Mbp | *qpAdm* | 0.0% | 18.0% | 24.4% | 50.2% | 38.5% | 48.8% | 53.2% | 68.8% | |
| | *qpAdm* + PCA | 41.0% | 51.7% | 54.6% | 67.8% | 64.4% | 69.8% | 72.7% | 80.5% | |
| | *qpAdm* + PCA + *ADMIXTURE* | 69.3% | 74.1% | 75.1% | 84.4% | 81.0% | 82.9% | 83.9% | 89.3% | |

False discovery and false omission rates (FDR, FOR) of the proximal rotating *qpAdm* protocol for a range of model positivity criteria. Simulation setups and methods used to declare a positive result are shown on the left. Two *p*-value thresholds were tested for accepting two-way models (0.01 and 0.1), and four *p*-value thresholds were tested for rejecting one-way models (0.01, $10^{-10}$, $10^{-20}$, and $10^{-50}$). For each simulation setup, the analysis relies on simulation replicate no. 1 only since PCA and *ADMIXTURE* results were not generated for multiple replicates (see Discussion).

**Table S4.**

| simulation scheme | max. depth of the simulation, gen. | pseudohaploid noisy data | genome size, Mbp | two-way feasible models binned by: model type | protocol | bin size: median (min.-max.) | fraction of models that are distal: median | min.-max. | FDR for proximal & distal models: median | min.-max. | FDR for distal models: median | min.-max. | supported by 3D PCA: bin size: | fraction of models that are distal: | FDR for proximal & distal models: | FDR for distal models: | supported by unsupervised *ADMIXTURE*: bin size: | fraction of models that are distal: | FDR for proximal & distal models: | FDR for distal models: |
|---|---|---|---|---|---|---|---|---|---|---|---|---|---|---|---|---|---|---|---|---|
| date variable, no. 1 (noisy) | 800 | yes | 300 | FP | non-rotating | 75 (62-95) | 37.3% | 30.6%-42.9% | 37.4% | 29.5%-42.3% | 24.2% | 17.6%-28.1% | | | | | | | | |
| | | | | FP | rotating | 642 (579-733) | 20.9% | 17.8%-24.6% | 53.0% | 51.6%-54.9% | 25.1% | 22.4%-29.0% | | | | | | | | |
| | | | | TP | non-rotating | 146 (115-149) | 67.0% | 60.0%-76.4% | | | | | | | | | | | | |
| | | | | TP | rotating | 554 (492-613) | 68.6% | 66.7%-73.8% | | | | | | | | | | | | |
| date variable, no. 2 (noisy) | 800 | yes | 1000 | FP | non-rotating | 86 (76-102) | 36.2% | 28.2%-47.3% | 39.3% | 34.1%-41.2% | 25.1% | 18.6%-32.6% | | | | | | | | |
| | | | | FP | rotating | 417 (355-500) | 19.4% | 17.7%-23.5% | 54.6% | 51.7%-59.7% | 24.0% | 21.8%-27.9% | | | | | | | | |
| | | | | TP | non-rotating | 133 (126-164) | 66.8% | 62.7%-72.5% | | | | | | | | | | | | |
| | | | | TP | rotating | 340 (297-405) | 74.0% | 70.1%-81.3%% | | | | | | | | | | | | |
| date variable, no. 1 | 800 | no | 300 | FP | non-rotating | 76 (62-92) | 39.2% | 34.5%-48.0% | 42.6% | 35.8%-45.5% | 31.2% | 24.5%-34.4% | 20<br>72 | 30.0%<br>40.3% | 35.1% | 18.2% | 2<br>90 | 100.0%<br>36.7% | 4.5% | 6.1% |
| | | | | FP | rotating | 247 (194-268) | 16.4% | 10.5%-19.1% | 57.4% | 55.6%-62.6% | 22.1% | 14.4%-27.9% | 129<br>83 | 7.8%<br>36.1% | 61.4% | 14.3% | 59<br>153 | 3.4%<br>24.8% | 49.6% | 4.4% |
| | | | | TP | non-rotating | 105 (93-115) | 65.5% | 59.5%-71.2% | | | | | 37<br>72 | 73.0%<br>59.7% | | | 42<br>67 | 73.8%<br>58.2% | | |
| | | | | TP | rotating | 165 (149-202) | 80.8% | 78.1%-84.1% | | | | | 81<br>92 | 74.1%<br>89.1% | | | 60<br>113 | 71.7%<br>87.6% | | |
| date variable, no. 2 | 800 | no | 1000 | FP | non-rotating | 43 (32-62) | 40.4% | 26.8%-53.1% | 38.4% | 31.3%-41.5% | 26.5% | 16.4%-30.4% | 14<br>18 | 35.7%<br>66.7% | 40.0% | 23.8% | 1<br>31 | 0.0%<br>54.8% | 3.2% | 0.0% |
| | | | | FP | rotating | 127 (106-139) | 10.5% | 6.0%-16.7% | 62.0% | 57.6%-68.1% | 16.4% | 12.1%-26.7% | 84<br>30 | 3.6%<br>26.7% | 70.6% | 10.3% | 18<br>96 | 0.0%<br>11.5% | 40.9% | 0.0% |
| | | | | TP | non-rotating | 75 (65-95) | 66.0% | 58.9%-74.7% | | | | | 21<br>49 | 76.2%<br>73.5% | | | 30<br>40 | 80.0%<br>70.0% | | |
| | | | | TP | rotating | 75 (65-84) | 83.6% | 80.0%-88.1% | | | | | 35<br>49 | 74.3%<br>91.8% | | | 26<br>58 | 69.2%<br>91.4% | | |
| date variable, no. 4 | 3000 | no | 300 | FP | non-rotating | 36 | 38.9% | | 36.0% | | 24.1% | | 12<br>24 | 33.3%<br>41.7% | 34.3% | 20.0% | 5<br>31 | 20.0%<br>41.9% | 12.8% | 5.3% |
| | | | | FP | rotating | 61 | 16.4% | | 66.3% | | 25.6% | | 41<br>20 | 7.3%<br>35.0% | 77.4% | 23.1% | 12<br>49 | 8.3%<br>18.4% | 46.2% | 7.7% |
| | | | | TP | non-rotating | 64 | 68.8% | | | | | | 23<br>41 | 69.6%<br>68.3% | | | 34<br>30 | 52.9%<br>86.7% | | |
| | | | | TP | rotating | 31 | 93.5% | | | | | | 12<br>19 | 83.3%<br>100.0% | | | 14<br>17 | 85.7%<br>100.0% | | |
| at "present", no. 5 | 800 | no | 300 | FP | non-rotating | 99 | 40.4% | | 42.1% | | 30.3% | | 53<br>46 | 64.2%<br>13.0% | 36.1% | 34.0% | 18<br>81 | 72.2%<br>33.3% | 20.5% | 19.1% |
| | | | | FP | rotating | 364 | 17.6% | | 64.0% | | 29.8% | | 178<br>186 | 22.5%<br>12.9% | 59.5% | 29.9% | 59<br>305 | 22.0%<br>16.7% | 37.8% | 15.1% |
| | | | | TP | non-rotating | 136 | 67.6% | | | | | | 94<br>42 | 70.2%<br>61.9% | | | 70<br>66 | 78.6%<br>56.1% | | |
| | | | | TP | rotating | 205 | 73.7% | | | | | | 121<br>84 | 77.7%<br>67.9% | | | 97<br>108 | 75.3%<br>72.2% | | |

Assessing the effect of temporal stratification of targets and proxy sources on FDR of non-rotating and rotating *qpAdm* protocols and their combinations with PCA and *ADMIXTURE* analyses**.** Another kind of temporal stratification, stratification of the "right" and "left" sets, was a part of the non-rotating protocols, but not of the rotating ones. In the left half of the table, feasible two-way models are binned by model type (false or true positive) and *qpAdm* protocol (rotating or non-rotating), and for each bin its size (number of models) and fraction of models that are distal (magenta bars) are shown. FDR values are also visualized by cell background color. In the right half of the table these bins are further subdivided into models supported/not supported (highlighted in green and red, respectively) by another method (PCA or *ADMIXTURE*). In the case of simulation setups no. 1 – 3, 10 simulation replicates and 10 subsampling replicates derived from simulation replicate no. 1 were generated, and median, minimal, and maximal values across the replicates are shown for sizes of the bins, for fractions of models that are distal, and for FDR.

**Table S5.**

| two-way models supported by: | simulation scheme | max. depth of the simulation, gen. | pseudohaploid noisy data | genome size, Mbp | feasible *qpAdm* models/ all 2-way models | non-rotating *qpAdm* with temporal stratification of the R and L sets, but with no stratification of targets and proxy sources | | | | one or two alternative sources "on the left", all others "on the right" | | | | one or two alternative sources "on the left", all others "on the right", one at a time | | | |
|---|---|---|---|---|---|---|---|---|---|---|---|---|---|---|---|---|---|
| proximal non-rotating *qpAdm* | | | | | | 1 | 1 | 1 | 1 | 1 | 1 | 1 | 1 | 1 | 1 | 1 | 1 |
| *qpAdm* model competition | | | | | | | | | | 2 | 2 | 2 | 2 | 2 | 2 | 2 | 2 |
| "admixture" *f3*- statistics | | | | | | | 2 | | | | | | | | | | |
| 3D PCA | | | | | | | | 2 | | | 3 | | 3 | | 3 | | 3 |
| unsupervised *ADMIXTURE* | | | | | | | | | 2 | | | 3 | 4 | | | 3 | 4 |
| | date variable, no. 1 | 800 | no | 300 | 201/ 4,200 (4.8%) | 45.8% | 0.0% | 10.0% | 1.0% | 16.4% | 6.0%<br>10.4% | 0.0%<br>16.4% | 0.0%<br>6.0%<br>0.0%<br>10.4% | 15.9% | 5.0%<br>10.9% | 0.0%<br>15.9% | 0.0%<br>5.0%<br>0.0%<br>10.9% |
| | | | | | | | 45.8% | 35.8% | 44.8% | 29.4% | 4.0%<br>25.4% | 1.0%<br>28.4% | 0.0%<br>4.0%<br>1.0%<br>24.4% | 29.9% | 5.0%<br>24.9% | 1.0%<br>28.9% | 0.0%<br>5.0%<br>1.0%<br>23.9% |
| | | | | | | 54.2% | 2.5% | 18.4% | 20.9% | 19.4% | 7.5%<br>11.9% | 10.0%<br>9.5% | 4.5%<br>3.0%<br>5.5%<br>6.5% | 22.4% | 8.5%<br>13.9% | 10.4%<br>11.9% | 5.5%<br>3.0%<br>5.0%<br>9.0% |
| | | | | | | | 51.7% | 35.8% | 33.3% | 34.8% | 10.9%<br>23.9% | 10.9%<br>23.9% | 4.0%<br>7.0%<br>7.0%<br>16.9% | 31.8% | 10.0%<br>21.9% | 10.4%<br>21.4% | 3.0%<br>7.0%<br>7.5%<br>14.4% |
| | | | | | FDR | 45.8% | 0% | 35.1% | 4.5% | 45.8% | 44.4% | 0% | 0% | 41.6% | 37.0% | 0% | 0% |
| | | | | | FOR | 0% | 95.4% | 66.1% | 61.5% | 64.2% | 86.2% | 81.7% | 91.7% | 58.7% | 84.4% | 80.7% | 89.9% |
| | date variable, no. 4 | 3000 | no | 300 | 100/ 4,200 (2.4%) | 36.0% | 0.0% | 12.0% | 5.0% | 8.0% | 6.0%<br>2.0% | 2.0%<br>6.0% | 2.0%<br>4.0%<br>0.0%<br>2.0% | 8.0% | 6.0%<br>2.0% | 2.0%<br>6.0% | 2.0%<br>4.0%<br>0.0%<br>2.0% |
| | | | | | | | 36.0% | 24.0% | 31.0% | 28.0% | 6.0%<br>22.0% | 3.0%<br>25.0% | 3.0%<br>3.0%<br>0.0%<br>22.0% | 28.0% | 6.0%<br>22.0% | 3.0%<br>25.0% | 3.0%<br>3.0%<br>0.0%<br>22.0% |
| | | | | | | 64.0% | 4.0% | 23.0% | 34.0% | 20.0% | 5.0%<br>15.0% | 7.0%<br>13.0% | 4.0%<br>1.0%<br>3.0%<br>12.0% | 18.0% | 5.0%<br>13.0% | 7.0%<br>11.0% | 4.0%<br>1.0%<br>3.0%<br>10.0% |
| | | | | | | | 60.0% | 41.0% | 30.0% | 44.0% | 18.0%<br>26.0% | 27.0%<br>17.0% | 15.0%<br>3.0%<br>12.0%<br>14.0% | 46.0% | 18.0%<br>28.0% | 27.0%<br>19.0% | 15.0%<br>3.0%<br>12.0%<br>16.0% |
| | | | | | FDR | 36.0% | 0% | 34.3% | 12.8% | 28.6% | 54.5% | 22.2% | 33.3% | 30.8% | 54.5% | 22.2% | 33.3% |
| | | | | | FOR | 0% | 93.8% | 64.1% | 46.9% | 68.8% | 92.2% | 89.1% | 93.8% | 71.9% | 92.2% | 89.1% | 93.8% |
| | at "present", no. 5 | 800 | no | 300 | 235/ 4,200 (5.6%) | 42.1% | 0.0% | 22.6% | 7.7% | 13.6% | 9.4%<br>4.3% | 5.1%<br>8.5% | 3.8%<br>5.5%<br>1.3%<br>3.0% | 11.9% | 7.7%<br>4.3% | 4.3%<br>7.7% | 3.0%<br>4.7%<br>1.3%<br>3.0% |
| | | | | | | | 42.1% | 19.6% | 34.5% | 28.5% | 13.2%<br>15.3% | 2.6%<br>26.0% | 2.1%<br>11.1%<br>0.4%<br>14.9% | 30.2% | 14.9%<br>15.3% | 3.4%<br>26.8% | 3.0%<br>11.9%<br>0.4%<br>14.9% |
| | | | | | | 57.9% | 2.6% | 40.0% | 29.8% | 22.1% | 15.7%<br>6.4% | 11.9%<br>10.2% | 10.2%<br>5.5%<br>1.7%<br>4.7% | 23.8% | 17.0%<br>6.8% | 13.2%<br>10.6% | 11.1%<br>6.0%<br>2.1%<br>4.7% |
| | | | | | | | 55.3% | 17.9% | 28.1% | 35.7% | 24.3%<br>11.5% | 17.9%<br>17.9% | 14.5%<br>9.8%<br>3.4%<br>8.1% | 34.0% | 23.0%<br>11.1% | 16.6%<br>17.4% | 13.6%<br>9.4%<br>3.0%<br>8.1% |
| | | | | | FDR | 42.1% | 0% | 36.1% | 20.5% | 38.1% | 37.3% | 30.0% | 27.3% | 33.3% | 31.0% | 24.4% | 21.2% |
| | | | | | FOR | 0% | 95.6% | 30.9% | 48.5% | 61.8% | 72.8% | 79.4% | 82.4% | 58.8% | 70.6% | 77.2% | 80.9% |

Assessing FDR of the proximal non-rotating *qpAdm* protocol combined with model competition (Narasimhan *et al.* 2019). For each simulation setup, the analysis relies on simulation replicate no. 1 only. For constructing pipelines resembling those common in the archaeogenetic literature, we used five methods: proximal non-rotating *qpAdm*, *qpAdm* model competition (two alternative protocols), "admixture" $f_3$-statistics, 3D PCA with all individuals co-analyzed, and unsupervised *ADMIXTURE* with all individuals co-analyzed. For declaring a positive result, support of an admixture model by all methods in the pipeline was required, hence the order of methods is not important except for the first method, which was the proximal non-rotating *qpAdm* in all cases. In each column, methods comprising a pipeline are color-coded and numbered by their order. All feasible two-way *qpAdm* models emerging as outcomes of the proximal non-rotating protocol were classified into FP and TP (highlighted in red and green in the leftmost column, respectively). The other columns are structured like bifurcating trees: FP *qpAdm* models supported by method no. 2; FP *qpAdm* models not supported by method no. 2; TP *qpAdm* models supported by method no. 2; TP *qpAdm* models not supported by method no. 2. The same principle is used for representing results of more complex pipelines. All model counts are normalized by the number of feasible *qpAdm* models (FP + TP) that are outcomes of the first method. Percentages of models supported/not supported by the last method in the pipeline are highlighted in green and red, respectively. FDR values are shown for these different pipelines. Fractions of TP *qpAdm* models that are pruned out by progressively more stringent support requirements are also shown (false omission rate or FOR).

**Table S6.**

| gene-flow intensity limits | ~10-5 to ~10-4, per gen. | random sampling parameters | ~10-4 to ~10-3, per gen. | random sampling parameters | ~10-3 to ~10-2, per gen. | random sampling parameters | ~10-5 to ~10-2, per gen. | ~10-5 to ~10-2, per epoch | random sampling parameters |
|---|---|---|---|---|---|---|---|---|---|
| pre-LGM, min. | 2.1E-05 | uniform distr., | 2.1E-04 | uniform distr., | 2.1E-03 | uniform distr., | 2.1E-05 | 0.006 | mean=0, SD=0.3; |
| pre-LGM, max. | 7.1E-05 | once per era | 7.1E-04 | once per era | 3.6E-03 | 5x per era | 3.6E-03 | 1 | once per era |
| LGM, min. | 2.9E-06 | uniform distr., | 2.9E-06 | uniform distr., | 2.9E-06 | uniform distr., | 2.9E-06 | 0.0002 | mean=0, SD=0.3; |
| LGM, max. | 3.6E-05 | once per era | 3.6E-04 | once per era | 3.6E-03 | 5x per era | 3.6E-03 | 0.25 | once per era |
| post-LGM, min. | 2.3E-05 | uniform distr., | 2.3E-04 | uniform distr., | 2.3E-03 | uniform distr., | 2.3E-05 | 0.003 | mean=0, SD=0.3; |
| post-LGM, max. | 7.7E-05 | once per era | 7.7E-04 | once per era | 7.7E-03 | 5x per era | 7.7E-03 | 1 | once per era |

Gene-flow parameters of the simulated SSL histories. Maximal and minimal gene-flow intensities (percentage of individuals in each generation coming from a given neighboring deme) are shown for each of the three eras (pre-LGM, LGM, and post-LGM). Gene-flow intensities were sampled randomly within the specified intervals from either uniform or (in the case of the "$10^{-5}$ to $10^{-2}$" landscapes) from normal distributions centered at 0. For this landscape type, both per-generation and per-epoch gene-flow intensities are shown since the latter are comparable to the parameters of the normal distributions shown in the rightmost column. The random sampling was repeated either once per era or once per epoch (that is, five times per era at equal intervals), as shown in the table.

#### Table S7.

| | | variable 2: | max. \|estimated admixture fractions - equal fractions\| | | | | | *p*-value | | | | | max. SE | | | | | *p*-value | | |
|---|---|---|---|---|---|---|---|---|---|---|---|---|---|---|---|---|---|---|---|---|
| total number of randomized models: | | | 21,663,216 | 12,439,194 | 2,392,357 | 2,040,355 | 1,058,613 | 21,663,216 | 12,439,194 | 2,392,357 | 2,040,355 | 1,058,613 | 21,663,216 | 12,439,194 | 4,378,913 | 3,566,530 | 1,533,911 | 4,378,913 | 3,566,530 | 1,533,911 |
| total number of systematic models: | | | 3,576,667 | 1,158,133 | 1,267,603 | 1,064,846 | 240,961 | 3,576,667 | 1,158,133 | 1,267,603 | 1,064,846 | 240,961 | N/A | | | | | N/A | | |
| variable 1: | model complexity | gene-flow intensity per gen. | p >= 1e-55 | p >= 1e-5 | max. \|*EAF–EF*\| <= 0.5 | p >= 1e-55; max. \|*EAF–EF*\| <= 0.5 | p >= 1e-5; max. \|*EAF–EF*\| <= 0.5 | p >= 1e-55 | p >= 1e-5 | max. \|*EAF–EF*\| <= 0.5 | p >= 1e-55; max. \|*EAF–EF*\| <= 0.5 | p >= 1e-5; max. \|*EAF–EF*\| <= 0.5 | p >= 1e-55 | p >= 1e-5 | max. SE <= 0.1 | p >= 1e-55; max. SE <= 0.1 | p >= 1e-5; max. SE <= 0.1 | max. SE <= 0.1 | p >= 1e-55; max. SE <= 0.1 | p >= 1e-5; max. SE <= 0.1 |
| minimal source-target-source angle | 2-way | 10-5 to 10-4 | -0.13 | -0.24 | -0.02 | -0.01 | 0.00 | -0.03 | -0.02 | 0.04 | 0.04 | 0.02 | -0.10 | -0.22 | -0.01 | -0.01 | -0.01 | 0.06 | 0.08 | 0.02 |
| | | 10-4 to 10-3 | **-0.65** | **-0.80** | -0.14 | -0.19 | -0.25 | -0.10 | -0.33 | 0.10 | 0.03 | 0.07 | **-0.59** | **-0.74** | -0.12 | -0.16 | -0.15 | 0.07 | 0.02 | -0.01 |
| | | 10-5 to 10-2 | **-0.77** | **-0.78** | -0.20 | -0.23 | -0.24 | -0.07 | -0.10 | 0.24 | 0.01 | -0.04 | **-0.64** | **-0.68** | -0.29 | -0.28 | -0.17 | 0.06 | 0.02 | 0.00 |
| | | 10-5 to 10-2 syst. | **-0.81** | **-0.77** | -0.25 | -0.29 | -0.19 | -0.15 | 0.04 | 0.23 | 0.08 | 0.02 | | | | | | | | |
| | | 10-3 to 10-2 | **-0.79** | **-0.81** | -0.29 | -0.31 | -0.37 | -0.13 | -0.06 | 0.42 | 0.37 | 0.08 | **-0.66** | **-0.67** | -0.37 | -0.37 | -0.30 | 0.13 | 0.11 | 0.02 |
| | 3-way | 10-5 to 10-4 | -0.11 | -0.18 | -0.02 | -0.02 | -0.02 | -0.08 | -0.06 | 0.01 | 0.01 | 0.01 | -0.11 | -0.17 | 0.02 | 0.02 | 0.00 | 0.03 | 0.04 | 0.00 |
| | | 10-4 to 10-3 | -0.46 | **-0.61** | -0.14 | -0.16 | -0.17 | -0.15 | -0.30 | -0.03 | -0.01 | 0.05 | -0.45 | **-0.59** | -0.08 | -0.10 | -0.10 | -0.04 | -0.02 | -0.02 |
| | | 10-5 to 10-2 | **-0.51** | **-0.55** | -0.31 | -0.29 | -0.22 | -0.15 | -0.10 | -0.11 | -0.11 | -0.04 | -0.48 | **-0.53** | -0.24 | -0.24 | -0.18 | -0.08 | -0.05 | 0.00 |
| | | 10-5 to 10-2 syst. | **-0.65** | **-0.51** | -0.31 | -0.34 | -0.36 | -0.28 | -0.04 | 0.03 | 0.02 | -0.01 | | | | | | | | |
| | | 10-3 to 10-2 | **-0.52** | **-0.53** | -0.39 | -0.39 | -0.38 | -0.18 | -0.10 | -0.05 | -0.05 | -0.06 | -0.49 | -0.49 | -0.28 | -0.28 | -0.27 | 0.02 | 0.02 | 0.00 |
| | 4-way | 10-5 to 10-4 | -0.10 | -0.13 | -0.01 | -0.01 | -0.02 | -0.10 | -0.08 | 0.00 | 0.00 | 0.00 | -0.10 | -0.13 | 0.02 | 0.03 | 0.01 | 0.02 | 0.03 | 0.02 |
| | | 10-4 to 10-3 | -0.32 | -0.44 | -0.10 | -0.11 | -0.13 | -0.18 | -0.22 | -0.09 | -0.05 | 0.02 | -0.31 | -0.43 | -0.06 | -0.07 | -0.06 | -0.07 | -0.05 | -0.02 |
| | | 10-5 to 10-2 | -0.34 | -0.35 | -0.19 | -0.18 | -0.13 | -0.18 | -0.10 | -0.22 | -0.20 | -0.07 | -0.32 | -0.35 | -0.19 | -0.18 | -0.15 | -0.13 | -0.11 | -0.03 |
| | | 10-5 to 10-2 syst. | -0.39 | -0.23 | -0.30 | -0.31 | -0.30 | -0.35 | -0.08 | -0.13 | -0.11 | -0.11 | | | | | | | | |
| | | 10-3 to 10-2 | -0.31 | -0.31 | -0.28 | -0.28 | -0.27 | -0.13 | -0.11 | -0.15 | -0.15 | -0.13 | -0.27 | -0.28 | -0.18 | -0.18 | -0.17 | -0.08 | -0.08 | -0.06 |
| maximal source-target distance | 2-way | 10-5 to 10-4 | 0.02 | 0.03 | 0.01 | 0.01 | 0.00 | -0.06 | -0.01 | -0.05 | -0.05 | -0.02 | 0.03 | 0.05 | -0.03 | -0.03 | 0.07 | -0.06 | -0.07 | -0.01 |
| | | 10-4 to 10-3 | 0.15 | 0.12 | 0.19 | 0.31 | 0.40 | -0.11 | -0.03 | -0.25 | -0.11 | -0.02 | -0.02 | -0.21 | -0.13 | -0.26 | -0.45 | -0.25 | -0.09 | -0.02 |
| | | 10-5 to 10-2 | 0.25 | 0.18 | 0.33 | 0.48 | **0.59** | -0.18 | -0.07 | -0.34 | -0.11 | -0.02 | -0.07 | -0.24 | -0.18 | -0.40 | **-0.64** | -0.31 | -0.14 | -0.05 |
| | | 10-5 to 10-2 syst. | 0.19 | **0.67** | 0.10 | 0.28 | **0.77** | -0.18 | -0.10 | -0.30 | -0.31 | -0.12 | | | | | | | | |
| | | 10-3 to 10-2 | 0.29 | 0.28 | 0.38 | 0.41 | **0.55** | -0.29 | -0.12 | -0.37 | -0.30 | -0.09 | -0.01 | -0.12 | -0.17 | -0.20 | -0.40 | -0.28 | -0.24 | -0.07 |
| | 3-way | 10-5 to 10-4 | 0.01 | 0.01 | -0.06 | -0.06 | -0.01 | -0.02 | -0.01 | -0.01 | -0.01 | -0.01 | 0.02 | 0.02 | 0.01 | 0.02 | 0.06 | -0.05 | -0.06 | -0.04 |
| | | 10-4 to 10-3 | 0.11 | 0.08 | 0.05 | 0.07 | 0.05 | -0.07 | -0.06 | -0.14 | -0.06 | -0.10 | 0.05 | -0.02 | -0.11 | -0.16 | -0.20 | -0.11 | -0.03 | -0.05 |
| | | 10-5 to 10-2 | 0.19 | 0.14 | 0.23 | 0.22 | 0.16 | -0.15 | -0.06 | -0.16 | -0.10 | -0.06 | 0.07 | 0.01 | -0.14 | -0.18 | -0.28 | -0.13 | -0.07 | -0.01 |
| | | 10-5 to 10-2 syst. | 0.05 | 0.15 | 0.02 | 0.03 | 0.21 | -0.09 | -0.06 | -0.15 | -0.15 | -0.12 | | | | | | | | |
| | | 10-3 to 10-2 | 0.19 | 0.20 | 0.36 | 0.36 | 0.34 | -0.23 | -0.13 | -0.13 | -0.13 | -0.08 | 0.03 | 0.05 | -0.06 | -0.06 | -0.09 | -0.13 | -0.13 | -0.04 |
| | 4-way | 10-5 to 10-4 | 0.00 | 0.00 | -0.06 | -0.06 | -0.03 | -0.01 | 0.00 | 0.00 | -0.01 | -0.02 | 0.01 | 0.01 | 0.02 | 0.02 | 0.03 | -0.06 | -0.06 | -0.07 |
| | | 10-4 to 10-3 | 0.11 | 0.07 | 0.01 | 0.02 | 0.04 | -0.06 | -0.06 | -0.05 | -0.01 | -0.05 | 0.06 | 0.02 | -0.10 | -0.11 | -0.12 | -0.04 | 0.00 | -0.05 |
| | | 10-5 to 10-2 | 0.17 | 0.14 | 0.11 | 0.11 | 0.06 | -0.13 | -0.06 | -0.03 | -0.02 | -0.01 | 0.09 | 0.06 | -0.10 | -0.11 | -0.16 | -0.02 | -0.01 | 0.02 |
| | | 10-5 to 10-2 syst. | 0.01 | 0.02 | 0.02 | 0.02 | 0.08 | -0.03 | -0.01 | -0.06 | -0.06 | -0.04 | | | | | | | | |
| | | 10-3 to 10-2 | 0.18 | 0.18 | 0.22 | 0.22 | 0.21 | -0.15 | -0.12 | -0.07 | -0.07 | -0.05 | 0.07 | 0.07 | 0.00 | 0.00 | -0.01 | 0.02 | 0.02 | 0.03 |
| average source-target distance | 2-way | 10-5 to 10-4 | 0.11 | 0.19 | 0.03 | 0.02 | -0.03 | -0.12 | -0.05 | -0.15 | -0.16 | -0.06 | 0.11 | 0.23 | -0.07 | -0.05 | 0.16 | -0.19 | -0.22 | -0.06 |
| | | 10-4 to 10-3 | 0.39 | 0.40 | 0.11 | 0.15 | 0.20 | -0.22 | -0.15 | -0.46 | -0.27 | -0.18 | 0.30 | 0.19 | -0.02 | -0.10 | -0.30 | -0.49 | -0.28 | -0.17 |
| | | 10-5 to 10-2 | 0.47 | 0.44 | 0.10 | 0.25 | 0.39 | -0.28 | -0.15 | **-0.62** | -0.32 | -0.15 | 0.24 | 0.12 | -0.01 | -0.27 | **-0.53** | **-0.60** | -0.34 | -0.18 |
| | | 10-5 to 10-2 syst. | 0.22 | **0.80** | -0.32 | -0.19 | **0.76** | -0.37 | -0.08 | **-0.75** | **-0.58** | -0.13 | | | | | | | | |
| | | 10-3 to 10-2 | 0.46 | 0.50 | 0.10 | 0.12 | 0.30 | -0.45 | -0.18 | **-0.67** | **-0.60** | -0.22 | 0.26 | 0.18 | -0.07 | -0.12 | -0.38 | **-0.60** | **-0.55** | -0.20 |
| | 3-way | 10-5 to 10-4 | 0.12 | 0.17 | -0.08 | -0.08 | -0.03 | -0.08 | -0.05 | -0.12 | -0.12 | -0.08 | 0.13 | 0.19 | -0.02 | 0.01 | 0.21 | -0.25 | -0.26 | -0.14 |
| | | 10-4 to 10-3 | 0.40 | 0.44 | 0.03 | 0.07 | 0.09 | -0.16 | -0.13 | -0.36 | -0.20 | -0.19 | 0.34 | 0.34 | -0.03 | -0.09 | -0.19 | -0.36 | -0.22 | -0.16 |
| | | 10-5 to 10-2 | 0.46 | 0.47 | 0.15 | 0.16 | 0.14 | -0.26 | -0.13 | -0.49 | -0.37 | -0.17 | 0.32 | 0.32 | -0.12 | -0.18 | -0.31 | -0.41 | -0.30 | -0.12 |
| | | 10-5 to 10-2 syst. | 0.12 | **0.61** | -0.40 | -0.45 | 0.19 | -0.50 | 0.01 | **-0.73** | **-0.69** | -0.24 | | | | | | | | |
| | | 10-3 to 10-2 | 0.43 | 0.46 | 0.27 | 0.27 | 0.27 | -0.33 | -0.19 | -0.40 | -0.40 | -0.27 | 0.22 | 0.25 | -0.09 | -0.09 | -0.14 | -0.36 | -0.36 | -0.19 |
| | 4-way | 10-5 to 10-4 | 0.12 | 0.15 | -0.09 | -0.09 | -0.06 | -0.06 | -0.04 | -0.10 | -0.10 | -0.09 | 0.13 | 0.15 | 0.05 | 0.08 | 0.16 | -0.27 | -0.27 | -0.21 |
| | | 10-4 to 10-3 | 0.38 | 0.41 | 0.03 | 0.05 | 0.10 | -0.14 | -0.11 | -0.26 | -0.17 | -0.14 | 0.31 | 0.34 | -0.07 | -0.09 | -0.13 | -0.26 | -0.19 | -0.14 |
| | | 10-5 to 10-2 | 0.43 | 0.45 | 0.12 | 0.12 | 0.10 | -0.23 | -0.11 | -0.33 | -0.30 | -0.13 | 0.31 | 0.33 | -0.14 | -0.15 | -0.20 | -0.26 | -0.22 | -0.07 |
| | | 10-5 to 10-2 syst. | 0.19 | **0.54** | -0.44 | -0.45 | 0.01 | -0.47 | 0.01 | **-0.73** | **-0.72** | -0.36 | | | | | | | | |
| | | 10-3 to 10-2 | 0.40 | 0.41 | 0.22 | 0.22 | 0.22 | -0.22 | -0.18 | -0.29 | -0.29 | -0.25 | 0.23 | 0.24 | -0.07 | -0.07 | -0.07 | -0.14 | -0.14 | -0.10 |
| distance to the ideal symmetric model (in the space of max. distance vs. min. angle) | 2-way | 10-5 to 10-4 | 0.10 | 0.18 | 0.02 | 0.02 | 0.02 | 0.01 | 0.02 | -0.03 | -0.03 | -0.01 | 0.08 | 0.17 | 0.00 | 0.00 | 0.01 | -0.05 | -0.07 | -0.01 |
| | | 10-4 to 10-3 | **0.58** | **0.72** | 0.20 | 0.30 | 0.43 | 0.04 | 0.22 | -0.16 | -0.06 | -0.09 | 0.48 | **0.62** | 0.04 | 0.04 | -0.01 | -0.14 | -0.03 | -0.02 |
| | | 10-5 to 10-2 | **0.72** | **0.73** | 0.32 | 0.44 | **0.51** | -0.01 | 0.05 | -0.31 | -0.06 | -0.01 | **0.50** | **0.53** | 0.14 | 0.07 | -0.09 | -0.16 | -0.06 | -0.03 |
| | | 10-5 to 10-2 syst. | **0.82** | **0.78** | 0.26 | 0.33 | 0.33 | 0.14 | -0.04 | -0.25 | -0.12 | -0.04 | | | | | | | | |
| | | 10-3 to 10-2 | **0.72** | **0.76** | 0.42 | 0.46 | **0.60** | -0.02 | 0.00 | -0.46 | -0.39 | -0.11 | **0.51** | **0.52** | 0.19 | 0.18 | 0.07 | -0.19 | -0.16 | -0.04 |
| | 3-way | 10-5 to 10-4 | 0.07 | 0.11 | -0.02 | -0.02 | 0.00 | 0.05 | 0.04 | -0.01 | -0.01 | -0.01 | 0.08 | 0.11 | 0.00 | -0.01 | 0.01 | -0.01 | -0.02 | 0.00 |
| | | 10-4 to 10-3 | 0.39 | 0.50 | 0.11 | 0.14 | 0.15 | 0.07 | 0.15 | -0.04 | -0.01 | -0.07 | 0.36 | 0.45 | 0.01 | 0.01 | 0.00 | -0.02 | 0.02 | 0.01 |
| | | 10-5 to 10-2 | 0.47 | 0.49 | 0.34 | 0.32 | 0.24 | 0.02 | 0.03 | 0.03 | 0.06 | 0.03 | 0.39 | 0.43 | 0.12 | 0.11 | 0.02 | 0.00 | 0.01 | 0.00 |
| | | 10-5 to 10-2 syst. | **0.65** | **0.51** | 0.31 | 0.34 | 0.37 | 0.28 | 0.03 | -0.04 | -0.02 | 0.00 | | | | | | | | |
| | | 10-3 to 10-2 | 0.47 | 0.49 | 0.46 | 0.46 | 0.45 | 0.00 | 0.00 | 0.00 | 0.00 | 0.03 | 0.37 | 0.39 | 0.17 | 0.17 | 0.16 | -0.07 | -0.07 | -0.02 |
| | 4-way | 10-5 to 10-4 | 0.06 | 0.07 | -0.03 | -0.03 | -0.01 | 0.06 | 0.05 | 0.00 | 0.00 | -0.01 | 0.06 | 0.07 | 0.00 | 0.00 | 0.00 | 0.00 | -0.01 | -0.03 |
| | | 10-4 to 10-3 | 0.28 | 0.35 | 0.08 | 0.09 | 0.11 | 0.08 | 0.10 | 0.04 | 0.04 | -0.04 | 0.26 | 0.32 | 0.00 | -0.01 | -0.01 | 0.04 | 0.05 | 0.02 |
| | | 10-5 to 10-2 | 0.34 | 0.34 | 0.19 | 0.18 | 0.12 | 0.04 | 0.02 | 0.17 | 0.15 | 0.06 | 0.28 | 0.29 | 0.09 | 0.09 | 0.04 | 0.08 | 0.07 | 0.04 |
| | | 10-5 to 10-2 syst. | 0.39 | 0.23 | 0.30 | 0.31 | 0.31 | 0.35 | 0.08 | 0.13 | 0.11 | 0.11 | | | | | | | | |
| | | 10-3 to 10-2 | 0.32 | 0.33 | 0.31 | 0.31 | 0.31 | 0.00 | 0.00 | 0.10 | 0.10 | 0.09 | 0.23 | 0.24 | 0.12 | 0.12 | 0.12 | 0.06 | 0.06 | 0.05 |
| distance to the ideal symmetric model (in the space of avg. distance vs. min. angle) | 2-way | 10-5 to 10-4 | 0.13 | 0.24 | 0.03 | 0.03 | 0.03 | 0.01 | 0.01 | -0.06 | -0.06 | -0.02 | 0.11 | 0.22 | -0.01 | -0.01 | 0.02 | -0.07 | -0.10 | -0.02 |
| | | 10-4 to 10-3 | **0.67** | **0.81** | 0.17 | 0.24 | 0.33 | 0.05 | 0.26 | -0.19 | -0.06 | -0.09 | **0.60** | **0.74** | 0.09 | 0.13 | 0.10 | -0.17 | -0.04 | -0.01 |
| | | 10-5 to 10-2 | **0.79** | **0.81** | 0.22 | 0.30 | 0.36 | 0.00 | 0.07 | -0.37 | -0.08 | 0.01 | **0.65** | **0.68** | 0.26 | 0.22 | 0.08 | -0.19 | -0.07 | -0.01 |
| | | 10-5 to 10-2 syst. | **0.81** | **0.80** | 0.19 | 0.23 | 0.29 | 0.11 | -0.04 | -0.34 | -0.20 | -0.04 | | | | | | | | |
| | | 10-3 to 10-2 | **0.80** | **0.84** | 0.29 | 0.32 | 0.45 | -0.02 | 0.01 | **-0.54** | -0.48 | -0.14 | **0.65** | **0.67** | 0.30 | 0.29 | 0.20 | -0.25 | -0.21 | -0.05 |
| | 3-way | 10-5 to 10-4 | 0.11 | 0.18 | -0.01 | -0.01 | 0.01 | 0.06 | 0.05 | -0.04 | -0.04 | -0.02 | 0.11 | 0.17 | -0.03 | -0.03 | 0.01 | -0.05 | -0.06 | 0.00 |
| | | 10-4 to 10-3 | **0.53** | **0.66** | 0.12 | 0.15 | 0.18 | 0.09 | 0.22 | -0.06 | -0.03 | -0.06 | 0.50 | **0.63** | 0.06 | 0.07 | 0.06 | -0.04 | 0.00 | 0.01 |
| | | 10-5 to 10-2 | **0.59** | **0.64** | 0.31 | 0.30 | 0.23 | 0.05 | 0.05 | 0.01 | 0.05 | 0.03 | **0.53** | **0.59** | 0.19 | 0.19 | 0.12 | -0.01 | 0.00 | -0.01 |
| | | 10-5 to 10-2 syst. | **0.66** | **0.57** | 0.26 | 0.28 | 0.38 | 0.23 | 0.04 | -0.11 | -0.10 | -0.01 | | | | | | | | |
| | | 10-3 to 10-2 | **0.59** | **0.60** | 0.42 | 0.42 | 0.41 | 0.05 | 0.03 | -0.01 | -0.01 | 0.02 | 0.50 | **0.52** | 0.23 | 0.23 | 0.22 | -0.09 | -0.09 | -0.04 |
| | 4-way | 10-5 to 10-4 | 0.11 | 0.14 | -0.02 | -0.02 | 0.00 | 0.07 | 0.06 | -0.03 | -0.03 | -0.02 | 0.11 | 0.14 | -0.02 | -0.03 | 0.00 | -0.04 | -0.05 | -0.04 |
| | | 10-4 to 10-3 | 0.41 | **0.53** | 0.10 | 0.11 | 0.14 | 0.12 | 0.16 | 0.02 | 0.01 | -0.04 | 0.38 | **0.50** | 0.04 | 0.04 | 0.03 | 0.02 | 0.02 | 0.01 |
| | | 10-5 to 10-2 | 0.45 | 0.49 | 0.20 | 0.19 | 0.14 | 0.08 | 0.05 | 0.16 | 0.15 | 0.06 | 0.40 | 0.44 | 0.15 | 0.14 | 0.10 | 0.06 | 0.06 | 0.02 |
| | | 10-5 to 10-2 syst. | 0.42 | 0.30 | 0.26 | 0.26 | 0.31 | 0.30 | 0.08 | 0.05 | 0.03 | 0.07 | | | | | | | | |
| | | 10-3 to 10-2 | 0.42 | 0.43 | 0.30 | 0.30 | 0.30 | 0.04 | 0.03 | 0.10 | 0.10 | 0.09 | 0.33 | 0.34 | 0.15 | 0.15 | 0.14 | 0.04 | 0.04 | 0.03 |

*p*-value > 0.05/1230 (Bonferroni correction)

**|Spearman correlation coefficient| >= 0.5**

Spearman's correlation coefficient (ρ) between several model optimality metrics (listed on the left) and max|*EAF–EF*|, *p*-values, or max. SE in various sections of the respective two-dimensional spaces (in columns). Results are shown for both randomized and systematic *qpAdm* experiments (the latter labelled "syst.") stratified by model complexity and landscape type. Non-significant ρ values (with Bonferroni correction) are underlined, and absolute ρ values ≥0.5 are shown in bold.

#### Table S8.

| part 1 | | | | variable 2: | max. \|estimated admixture fractions - equal fractions\| | | | | | *p*-value | | | | | max. SE | | | *p*-value | | |
|---|---|---|---|---|---|---|---|---|---|---|---|---|---|---|---|---|---|---|---|---|
| total number of randomized models: | | | | | 21,663,216 | 12,439,194 | 2,392,357 | 2,040,355 | 1,058,613 | 21,663,216 | 12,439,194 | 2,392,357 | 2,040,355 | 1,058,613 | 4,378,913 | 3,566,530 | 1,533,911 | 4,378,913 | 3,566,530 | 1,533,911 |
| total number of systematic models (10-5 to 10-2): | | | | | 3,576,667 | 1,158,133 | 1,267,603 | 1,064,846 | 240,961 | 3,576,667 | 1,158,133 | 1,267,603 | 1,064,846 | 240,961 | N/A | | | N/A | | |
| variable 1: | model complexity | gene-flow intensity per gen. | *qpAdm* protocol | sampling density | p >= 1e-55 | p >= 1e-5 | max. \|*EAF–EF*\| <= 0.5 | p >= 1e-55; max. \|*EAF–EF*\| <= 0.5 | p >= 1e-5; max. \|*EAF–EF*\| <= 0.5 | p >= 1e-55 | p >= 1e-5 | max. \|*EAF–EF*\| <= 0.5 | p >= 1e-55; max. \|*EAF–EF*\| <= 0.5 | p >= 1e-5; max. \|*EAF–EF*\| <= 0.5 | max. SE <= 0.1 | p >= 1e-55; max. SE <= 0.1 | p >= 1e-5; max. SE <= 0.1 | max. SE <= 0.1 | p >= 1e-55; max. SE <= 0.1 | p >= 1e-5; max. SE <= 0.1 |
| minimal source-target-source angle | 2-way | 10-5 to 10-4 | dist. rot. | 13 demes | -0.10 | -0.19 | 0.03 | 0.03 | 0.01 | -0.05 | -0.01 | 0.03 | 0.02 | 0.01 | -0.02 | -0.03 | -0.01 | 0.04 | 0.04 | 0.01 |
| | | | | 18 demes | -0.15 | -0.30 | -0.03 | -0.02 | 0.00 | -0.04 | -0.03 | 0.05 | 0.05 | 0.05 | 0.02 | 0.02 | -0.04 | 0.08 | 0.09 | 0.03 |
| | | | prox. rot. | 13 demes | -0.12 | -0.20 | -0.03 | -0.02 | 0.01 | -0.01 | 0.02 | 0.05 | 0.06 | 0.06 | 0.04 | 0.04 | 0.02 | 0.07 | 0.10 | 0.05 |
| | | | | 18 demes | -0.14 | -0.30 | -0.02 | -0.02 | 0.01 | -0.04 | -0.02 | 0.06 | 0.04 | 0.01 | 0.01 | 0.01 | -0.02 | 0.09 | 0.09 | 0.04 |
| | | | dist. non-rot. | 13 demes | -0.11 | -0.16 | -0.03 | -0.03 | -0.03 | 0.00 | -0.03 | 0.03 | 0.03 | -0.01 | 0.01 | 0.01 | 0.04 | 0.09 | 0.12 | 0.01 |
| | | | | 18 demes | -0.13 | -0.27 | -0.03 | -0.03 | 0.03 | -0.02 | 0.02 | 0.04 | 0.04 | 0.07 | -0.04 | -0.04 | -0.03 | 0.04 | 0.08 | 0.04 |
| | | | prox. non-rot. | 13 demes | -0.13 | -0.19 | -0.03 | -0.02 | -0.02 | -0.06 | -0.04 | 0.01 | -0.02 | 0.01 | -0.02 | -0.02 | 0.05 | 0.03 | 0.03 | -0.01 |
| | | | | 18 demes | -0.14 | -0.33 | -0.03 | -0.03 | -0.06 | -0.04 | -0.07 | 0.05 | 0.07 | 0.05 | -0.06 | -0.06 | -0.02 | 0.03 | 0.08 | 0.03 |
| | | 10-4 to 10-3 | dist. rot. | 13 demes | **-0.61** | **-0.77** | -0.16 | -0.20 | -0.25 | -0.10 | -0.19 | 0.12 | 0.06 | 0.16 | -0.12 | -0.14 | -0.14 | 0.07 | 0.01 | 0.11 |
| | | | | 18 demes | **-0.70** | **-0.81** | -0.16 | -0.25 | -0.46 | -0.12 | **-0.51** | 0.15 | 0.02 | 0.12 | -0.10 | -0.18 | -0.14 | 0.10 | 0.05 | 0.00 |
| | | | prox. rot. | 13 demes | **-0.63** | **-0.76** | -0.08 | -0.11 | -0.17 | -0.11 | -0.30 | 0.11 | 0.07 | 0.16 | -0.14 | -0.18 | -0.17 | 0.07 | 0.04 | 0.00 |
| | | | | 18 demes | **-0.70** | **-0.83** | -0.16 | -0.24 | -0.24 | -0.14 | **-0.53** | 0.14 | 0.02 | 0.22 | -0.12 | -0.14 | -0.21 | 0.09 | 0.01 | 0.01 |
| | | | dist. non-rot. | 13 demes | **-0.56** | **-0.64** | -0.13 | -0.16 | -0.28 | -0.03 | 0.02 | 0.06 | 0.07 | -0.13 | -0.12 | -0.13 | -0.13 | 0.06 | 0.02 | -0.05 |
| | | | | 18 demes | **-0.64** | **-0.67** | -0.13 | -0.20 | -0.20 | -0.01 | 0.08 | 0.11 | 0.01 | 0.09 | -0.12 | -0.14 | -0.06 | 0.07 | 0.01 | 0.08 |
| | | | prox. non-rot. | 13 demes | **-0.58** | **-0.75** | -0.16 | -0.21 | -0.25 | -0.06 | -0.30 | 0.10 | 0.12 | 0.04 | -0.13 | -0.15 | -0.16 | 0.09 | 0.06 | -0.10 |
| | | | | 18 demes | **-0.70** | **-0.83** | -0.15 | -0.22 | -0.28 | -0.15 | **-0.53** | 0.11 | -0.03 | -0.03 | -0.11 | -0.13 | 0.05 | 0.08 | -0.01 | -0.09 |
| | | 10-5 to 10-2 | dist. rot. | 13 demes | **-0.75** | **-0.77** | -0.19 | -0.22 | -0.21 | -0.07 | -0.08 | 0.21 | 0.01 | -0.04 | -0.28 | -0.28 | -0.18 | 0.05 | 0.03 | -0.01 |
| | | | | 18 demes | **-0.78** | **-0.79** | -0.22 | -0.26 | -0.15 | -0.08 | -0.09 | 0.22 | 0.00 | -0.03 | -0.31 | -0.30 | -0.19 | 0.03 | 0.02 | 0.01 |
| | | | | systematic | **-0.82** | **-0.62** | -0.25 | -0.26 | -0.13 | -0.21 | -0.05 | 0.14 | 0.07 | 0.07 | | | | | | |
| | | | prox. rot. | 13 demes | **-0.75** | **-0.77** | -0.18 | -0.20 | -0.22 | -0.09 | -0.12 | 0.23 | 0.02 | -0.05 | -0.28 | -0.28 | -0.17 | 0.06 | 0.01 | -0.01 |
| | | | | 18 demes | **-0.78** | **-0.80** | -0.23 | -0.27 | -0.25 | -0.09 | -0.15 | 0.26 | -0.01 | -0.10 | -0.29 | -0.29 | -0.16 | 0.05 | 0.01 | -0.04 |
| | | | dist. non-rot. | 13 demes | **-0.75** | **-0.74** | -0.25 | -0.29 | -0.45 | -0.06 | -0.08 | 0.23 | 0.01 | -0.14 | -0.28 | -0.28 | -0.16 | 0.08 | 0.01 | -0.13 |
| | | | | 18 demes | **-0.79** | **-0.78** | -0.23 | -0.29 | -0.26 | -0.02 | 0.05 | 0.29 | 0.01 | 0.01 | -0.30 | -0.28 | -0.15 | 0.10 | 0.05 | 0.05 |
| | | | | systematic | **-0.81** | **-0.82** | -0.28 | -0.33 | -0.27 | -0.08 | 0.05 | 0.33 | 0.12 | -0.02 | | | | | | |
| | | | prox. non-rot. | 13 demes | **-0.76** | **-0.75** | -0.16 | -0.18 | -0.33 | -0.07 | -0.12 | 0.28 | 0.02 | -0.01 | -0.28 | -0.27 | -0.14 | 0.08 | 0.00 | -0.01 |
| | | | | 18 demes | **-0.80** | **-0.81** | -0.19 | -0.23 | -0.24 | -0.07 | -0.15 | 0.37 | 0.07 | 0.13 | -0.30 | -0.29 | -0.16 | 0.08 | 0.03 | 0.14 |
| | | 10-3 to 10-2 | dist. rot. | 13 demes | **-0.77** | **-0.80** | -0.28 | -0.29 | -0.33 | -0.11 | -0.06 | 0.41 | 0.38 | 0.07 | -0.34 | -0.34 | -0.26 | 0.16 | 0.15 | 0.04 |
| | | | | 18 demes | **-0.80** | **-0.81** | -0.32 | -0.35 | -0.42 | -0.14 | -0.06 | 0.40 | 0.31 | 0.03 | -0.39 | -0.39 | -0.28 | 0.12 | 0.09 | 0.01 |
| | | | prox. rot. | 13 demes | **-0.76** | **-0.79** | -0.22 | -0.23 | -0.25 | -0.15 | -0.07 | 0.39 | 0.36 | 0.04 | -0.32 | -0.32 | -0.25 | 0.11 | 0.10 | 0.01 |
| | | | | 18 demes | **-0.79** | **-0.81** | -0.24 | -0.27 | -0.25 | -0.13 | -0.06 | 0.47 | 0.39 | 0.05 | -0.38 | -0.39 | -0.32 | 0.13 | 0.10 | 0.01 |
| | | | dist. non-rot. | 13 demes | **-0.78** | **-0.81** | -0.35 | -0.36 | -0.44 | -0.11 | -0.04 | 0.45 | 0.44 | 0.16 | -0.37 | -0.37 | -0.33 | 0.18 | 0.17 | 0.03 |
| | | | | 18 demes | **-0.80** | **-0.82** | -0.35 | -0.37 | -0.48 | -0.13 | -0.03 | 0.44 | 0.40 | 0.09 | -0.40 | -0.40 | -0.33 | 0.14 | 0.12 | 0.02 |
| | | | prox. non-rot. | 13 demes | **-0.77** | **-0.81** | -0.25 | -0.25 | -0.36 | -0.15 | -0.10 | 0.47 | 0.46 | 0.19 | -0.35 | -0.35 | -0.34 | 0.15 | 0.14 | 0.05 |
| | | | | 18 demes | **-0.79** | **-0.82** | -0.29 | -0.31 | -0.40 | -0.18 | -0.10 | 0.49 | 0.45 | 0.14 | -0.40 | -0.40 | -0.36 | 0.12 | 0.10 | 0.03 |
| | 3-way | 10-5 to 10-4 | dist. rot. | 13 demes | -0.08 | -0.11 | 0.01 | 0.01 | 0.03 | -0.07 | -0.05 | 0.00 | 0.00 | 0.00 | 0.01 | 0.01 | -0.01 | 0.05 | 0.04 | 0.00 |
| | | | | 18 demes | -0.12 | -0.21 | -0.02 | -0.02 | -0.03 | -0.10 | -0.08 | 0.01 | 0.01 | 0.02 | 0.03 | 0.03 | 0.00 | 0.03 | 0.03 | 0.00 |
| | | | prox. rot. | 13 demes | -0.10 | -0.13 | -0.06 | -0.06 | -0.06 | -0.06 | -0.03 | 0.00 | 0.00 | 0.01 | -0.02 | -0.01 | -0.04 | -0.02 | -0.01 | 0.00 |
| | | | | 18 demes | -0.12 | -0.22 | -0.02 | -0.02 | -0.01 | -0.10 | -0.08 | -0.01 | -0.01 | 0.01 | 0.04 | 0.05 | 0.04 | 0.04 | 0.04 | 0.03 |
| | | | dist. non-rot. | 13 demes | -0.07 | -0.10 | -0.06 | -0.06 | -0.04 | -0.03 | -0.03 | 0.02 | 0.02 | 0.03 | 0.06 | 0.06 | 0.02 | 0.08 | 0.09 | 0.00 |
| | | | | 18 demes | -0.10 | -0.17 | -0.02 | -0.02 | 0.01 | -0.05 | -0.01 | 0.01 | 0.01 | 0.09 | 0.01 | 0.01 | -0.01 | 0.04 | 0.06 | 0.03 |
| | | | prox. non-rot. | 13 demes | -0.10 | -0.14 | -0.01 | -0.01 | -0.01 | -0.10 | -0.07 | -0.01 | -0.02 | -0.02 | 0.05 | 0.05 | 0.06 | 0.01 | 0.01 | -0.05 |
| | | | | 18 demes | -0.12 | -0.26 | -0.01 | -0.02 | -0.04 | -0.10 | -0.13 | 0.05 | 0.06 | 0.02 | -0.02 | -0.01 | -0.02 | 0.03 | 0.05 | 0.00 |
| | | 10-4 to 10-3 | dist. rot. | 13 demes | -0.40 | **-0.51** | -0.17 | -0.19 | -0.22 | -0.16 | -0.19 | -0.01 | -0.02 | 0.09 | -0.09 | -0.10 | -0.06 | -0.03 | -0.04 | 0.03 |
| | | | | 18 demes | -0.49 | **-0.65** | -0.15 | -0.15 | -0.19 | -0.17 | -0.42 | 0.02 | 0.00 | -0.01 | -0.08 | -0.10 | -0.11 | -0.02 | -0.01 | -0.04 |
| | | | prox. rot. | 13 demes | -0.41 | **-0.54** | -0.14 | -0.15 | -0.19 | -0.16 | -0.24 | 0.00 | 0.02 | 0.09 | -0.09 | -0.09 | -0.09 | -0.01 | 0.01 | -0.02 |
| | | | | 18 demes | **-0.51** | **-0.66** | -0.14 | -0.16 | -0.10 | -0.19 | -0.41 | -0.03 | -0.01 | 0.01 | -0.08 | -0.09 | -0.17 | -0.05 | -0.03 | 0.00 |
| | | | dist. non-rot. | 13 demes | -0.33 | -0.40 | -0.10 | -0.11 | -0.16 | -0.08 | -0.02 | -0.05 | 0.02 | 0.03 | -0.05 | -0.05 | -0.03 | -0.07 | -0.04 | -0.09 |
| | | | | 18 demes | -0.43 | -0.47 | -0.10 | -0.13 | -0.16 | -0.05 | 0.01 | -0.02 | 0.02 | 0.11 | -0.07 | -0.08 | -0.05 | -0.04 | -0.02 | -0.01 |
| | | | prox. non-rot. | 13 demes | -0.41 | **-0.55** | -0.14 | -0.16 | -0.14 | -0.12 | -0.25 | -0.01 | 0.04 | 0.10 | -0.09 | -0.10 | -0.08 | -0.01 | 0.02 | -0.04 |
| | | | | 18 demes | **-0.52** | **-0.68** | -0.15 | -0.21 | -0.19 | -0.19 | -0.44 | -0.02 | -0.02 | 0.10 | -0.07 | -0.11 | -0.02 | -0.03 | -0.05 | -0.02 |
| | | 10-5 to 10-2 | dist. rot. | 13 demes | -0.49 | **-0.53** | -0.29 | -0.28 | -0.22 | -0.15 | -0.09 | -0.10 | -0.11 | -0.03 | -0.21 | -0.22 | -0.16 | -0.05 | -0.04 | 0.00 |
| | | | | 18 demes | **-0.52** | **-0.56** | -0.32 | -0.30 | -0.24 | -0.17 | -0.11 | -0.12 | -0.12 | -0.06 | -0.25 | -0.26 | -0.20 | -0.11 | -0.07 | -0.01 |
| | | | | systematic | **-0.60** | -0.27 | -0.29 | -0.29 | -0.37 | -0.38 | -0.09 | -0.12 | -0.07 | -0.02 | | | | | | |
| | | | prox. rot. | 13 demes | -0.48 | **-0.52** | -0.25 | -0.24 | -0.18 | -0.17 | -0.11 | -0.12 | -0.12 | -0.06 | -0.20 | -0.20 | -0.15 | -0.09 | -0.06 | -0.01 |
| | | | | 18 demes | **-0.53** | **-0.57** | -0.32 | -0.29 | -0.18 | -0.17 | -0.14 | -0.12 | -0.11 | -0.08 | -0.25 | -0.26 | -0.18 | -0.10 | -0.05 | -0.02 |
| | | | dist. non-rot. | 13 demes | -0.49 | -0.48 | -0.31 | -0.31 | -0.20 | -0.12 | -0.06 | -0.08 | -0.10 | -0.06 | -0.26 | -0.27 | -0.24 | -0.07 | -0.07 | -0.09 |
| | | | | 18 demes | **-0.53** | **-0.55** | -0.32 | -0.30 | -0.27 | -0.09 | 0.01 | -0.05 | -0.07 | -0.02 | -0.25 | -0.26 | -0.23 | -0.06 | -0.03 | 0.01 |
| | | | | systematic | **-0.70** | **-0.63** | -0.34 | -0.37 | -0.35 | -0.19 | 0.01 | 0.15 | 0.07 | -0.02 | | | | | | |
| | | | prox. non-rot. | 13 demes | -0.50 | **-0.51** | -0.29 | -0.28 | -0.25 | -0.16 | -0.10 | -0.05 | -0.06 | 0.05 | -0.20 | -0.20 | -0.14 | -0.08 | -0.07 | -0.01 |
| | | | | 18 demes | **-0.54** | **-0.58** | -0.33 | -0.32 | -0.28 | -0.17 | -0.15 | -0.04 | -0.08 | -0.04 | -0.25 | -0.27 | -0.22 | -0.09 | -0.06 | 0.09 |
| | | 10-3 to 10-2 | dist. rot. | 13 demes | **-0.51** | **-0.52** | -0.37 | -0.37 | -0.36 | -0.15 | -0.08 | -0.05 | -0.05 | -0.05 | -0.22 | -0.22 | -0.21 | 0.03 | 0.03 | 0.02 |
| | | | | 18 demes | **-0.53** | **-0.53** | -0.40 | -0.40 | -0.39 | -0.21 | -0.10 | -0.07 | -0.07 | -0.09 | -0.28 | -0.28 | -0.26 | -0.01 | -0.01 | -0.02 |
| | | | prox. rot. | 13 demes | -0.49 | **-0.50** | -0.33 | -0.33 | -0.32 | -0.19 | -0.11 | -0.10 | -0.10 | -0.10 | -0.22 | -0.22 | -0.22 | -0.06 | -0.06 | -0.07 |
| | | | | 18 demes | **-0.52** | **-0.53** | -0.40 | -0.40 | -0.39 | -0.19 | -0.09 | -0.06 | -0.06 | -0.05 | -0.28 | -0.28 | -0.27 | 0.01 | 0.01 | -0.01 |
| | | | dist. non-rot. | 13 demes | **-0.50** | **-0.51** | -0.37 | -0.37 | -0.37 | -0.16 | -0.11 | -0.07 | -0.07 | -0.08 | -0.28 | -0.28 | -0.28 | 0.06 | 0.06 | 0.03 |
| | | | | 18 demes | **-0.52** | **-0.53** | -0.43 | -0.43 | -0.44 | -0.19 | -0.11 | -0.04 | -0.04 | -0.05 | -0.33 | -0.33 | -0.34 | 0.02 | 0.02 | 0.00 |
| | | | prox. non-rot. | 13 demes | **-0.51** | **-0.51** | -0.37 | -0.37 | -0.37 | -0.18 | -0.14 | -0.05 | -0.05 | -0.05 | -0.24 | -0.24 | -0.24 | 0.07 | 0.07 | 0.05 |
| | | | | 18 demes | **-0.54** | **-0.54** | -0.37 | -0.37 | -0.37 | -0.21 | -0.15 | -0.02 | -0.02 | -0.02 | -0.32 | -0.32 | -0.32 | 0.05 | 0.05 | 0.02 |
| | 4-way | 10-5 to 10-4 | dist. rot. | 13 demes | -0.05 | -0.06 | -0.01 | -0.01 | -0.01 | -0.07 | -0.06 | -0.03 | -0.03 | -0.04 | 0.10 | 0.10 | 0.08 | -0.05 | -0.05 | -0.12 |
| | | | | 18 demes | -0.11 | -0.14 | -0.02 | -0.02 | -0.04 | -0.10 | -0.10 | 0.00 | 0.00 | 0.00 | 0.05 | 0.05 | 0.02 | 0.03 | 0.03 | 0.04 |
| | | | prox. rot. | 13 demes | -0.08 | -0.09 | -0.04 | -0.04 | -0.05 | -0.06 | -0.04 | -0.01 | -0.01 | 0.00 | -0.01 | -0.01 | 0.04 | -0.05 | -0.05 | -0.03 |
| | | | | 18 demes | -0.12 | -0.16 | -0.01 | -0.01 | 0.00 | -0.12 | -0.10 | 0.01 | 0.01 | 0.01 | 0.02 | 0.03 | 0.01 | 0.01 | 0.03 | 0.03 |
| | | | dist. non-rot. | 13 demes | -0.05 | -0.06 | -0.06 | -0.06 | -0.05 | -0.03 | -0.02 | 0.04 | 0.04 | 0.04 | 0.04 | 0.04 | 0.04 | 0.04 | 0.04 | 0.05 |
| | | | | 18 demes | -0.07 | -0.09 | -0.01 | -0.01 | 0.02 | -0.04 | -0.02 | -0.02 | -0.02 | 0.07 | 0.01 | 0.01 | -0.01 | 0.03 | 0.04 | 0.05 |
| | | | prox. non-rot. | 13 demes | -0.10 | -0.10 | 0.00 | 0.00 | 0.00 | -0.10 | -0.08 | -0.01 | -0.01 | -0.01 | 0.08 | 0.07 | 0.10 | 0.01 | 0.01 | -0.01 |
| | | | | 18 demes | -0.11 | -0.20 | 0.00 | 0.00 | -0.03 | -0.12 | -0.14 | 0.03 | 0.04 | -0.02 | 0.01 | 0.03 | 0.00 | 0.03 | 0.05 | 0.00 |
| | | 10-4 to 10-3 | dist. rot. | 13 demes | -0.25 | -0.31 | -0.16 | -0.16 | -0.17 | -0.18 | -0.16 | -0.07 | -0.06 | 0.04 | -0.05 | -0.05 | -0.02 | -0.06 | -0.06 | -0.01 |
| | | | | 18 demes | -0.33 | -0.47 | -0.10 | -0.11 | -0.12 | -0.20 | -0.30 | -0.09 | -0.07 | -0.03 | -0.07 | -0.08 | -0.08 | -0.06 | -0.04 | -0.04 |
| | | | prox. rot. | 13 demes | -0.27 | -0.34 | -0.11 | -0.11 | -0.11 | -0.19 | -0.19 | -0.02 | -0.01 | 0.04 | -0.06 | -0.06 | -0.08 | -0.02 | -0.01 | -0.03 |
| | | | | 18 demes | -0.35 | -0.49 | -0.10 | -0.12 | -0.10 | -0.22 | -0.30 | -0.11 | -0.07 | 0.03 | -0.06 | -0.06 | -0.07 | -0.09 | -0.06 | 0.01 |
| | | | dist. non-rot. | 13 demes | -0.18 | -0.20 | -0.09 | -0.09 | -0.12 | -0.08 | -0.03 | -0.07 | -0.01 | 0.07 | -0.02 | -0.01 | -0.02 | -0.12 | -0.07 | -0.08 |
| | | | | 18 demes | -0.25 | -0.27 | -0.08 | -0.10 | -0.19 | -0.06 | -0.01 | -0.04 | 0.00 | 0.09 | -0.05 | -0.05 | -0.02 | -0.07 | -0.04 | -0.01 |
| | | | prox. non-rot. | 13 demes | -0.30 | -0.38 | -0.15 | -0.15 | -0.22 | -0.16 | -0.19 | -0.05 | -0.02 | 0.10 | -0.05 | -0.06 | -0.02 | -0.04 | -0.02 | 0.04 |
| | | | | 18 demes | -0.37 | **-0.54** | -0.11 | -0.13 | -0.12 | -0.21 | -0.35 | -0.07 | -0.01 | 0.00 | -0.06 | -0.07 | -0.05 | -0.08 | -0.06 | -0.03 |
| | | 10-5 to 10-2 | dist. rot. | 13 demes | -0.31 | -0.33 | -0.17 | -0.16 | -0.12 | -0.15 | -0.09 | -0.18 | -0.17 | -0.08 | -0.17 | -0.17 | -0.15 | -0.10 | -0.10 | -0.05 |
| | | | | 18 demes | -0.34 | -0.37 | -0.20 | -0.19 | -0.13 | -0.21 | -0.11 | -0.25 | -0.22 | -0.07 | -0.19 | -0.19 | -0.14 | -0.16 | -0.13 | -0.03 |
| | | | | systematic | -0.30 | -0.06 | -0.25 | -0.25 | -0.32 | -0.44 | -0.09 | -0.27 | -0.23 | -0.13 | | | | | | |
| | | | prox. rot. | 13 demes | -0.31 | -0.32 | -0.15 | -0.15 | -0.13 | -0.18 | -0.11 | -0.18 | -0.17 | -0.07 | -0.13 | -0.13 | -0.11 | -0.10 | -0.10 | -0.03 |
| | | | | 18 demes | -0.35 | -0.37 | -0.19 | -0.17 | -0.09 | -0.21 | -0.13 | -0.25 | -0.21 | -0.08 | -0.21 | -0.20 | -0.16 | -0.14 | -0.12 | -0.07 |
| | | | dist. non-rot. | 13 demes | -0.31 | -0.29 | -0.22 | -0.22 | -0.10 | -0.12 | -0.06 | -0.21 | -0.21 | -0.08 | -0.21 | -0.21 | -0.20 | -0.14 | -0.14 | -0.08 |
| | | | | 18 demes | -0.33 | -0.34 | -0.23 | -0.22 | -0.20 | -0.11 | -0.03 | -0.17 | -0.15 | -0.04 | -0.20 | -0.19 | -0.17 | -0.10 | -0.09 | -0.03 |
| | | | | systematic | -0.50 | -0.39 | -0.34 | -0.35 | -0.30 | -0.25 | -0.04 | -0.05 | -0.06 | -0.10 | | | | | | |
| | | | prox. non-rot. | 13 demes | -0.32 | -0.32 | -0.18 | -0.18 | -0.12 | -0.18 | -0.10 | -0.18 | -0.17 | 0.01 | -0.17 | -0.16 | -0.13 | -0.14 | -0.13 | -0.03 |
| | | | | 18 demes | -0.36 | -0.39 | -0.21 | -0.20 | -0.17 | -0.21 | -0.14 | -0.21 | -0.19 | -0.10 | -0.19 | -0.19 | -0.18 | -0.13 | -0.11 | 0.04 |
| | | 10-3 to 10-2 | dist. rot. | 13 demes | -0.30 | -0.30 | -0.25 | -0.25 | -0.25 | -0.09 | -0.09 | -0.12 | -0.12 | -0.12 | -0.15 | -0.15 | -0.14 | -0.13 | -0.13 | -0.14 |
| | | | | 18 demes | -0.31 | -0.31 | -0.28 | -0.28 | -0.27 | -0.15 | -0.11 | -0.19 | -0.19 | -0.15 | -0.20 | -0.20 | -0.20 | -0.12 | -0.12 | -0.09 |
| | | | prox. rot. | 13 demes | -0.27 | -0.28 | -0.23 | -0.23 | -0.23 | -0.13 | -0.11 | -0.14 | -0.14 | -0.13 | -0.14 | -0.14 | -0.14 | -0.09 | -0.09 | -0.10 |
| | | | | 18 demes | -0.31 | -0.31 | -0.28 | -0.28 | -0.27 | -0.14 | -0.10 | -0.15 | -0.15 | -0.12 | -0.18 | -0.18 | -0.17 | -0.09 | -0.09 | -0.08 |
| | | | dist. non-rot. | 13 demes | -0.29 | -0.29 | -0.28 | -0.28 | -0.28 | -0.11 | -0.11 | -0.17 | -0.17 | -0.17 | -0.10 | -0.10 | -0.10 | -0.08 | -0.08 | -0.10 |
| | | | | 18 demes | -0.32 | -0.32 | -0.30 | -0.30 | -0.30 | -0.15 | -0.13 | -0.15 | -0.15 | -0.15 | -0.20 | -0.20 | -0.20 | -0.04 | -0.04 | -0.05 |
| | | | prox. non-rot. | 13 demes | -0.29 | -0.29 | -0.22 | -0.22 | -0.22 | -0.13 | -0.13 | -0.13 | -0.13 | -0.13 | -0.04 | -0.04 | -0.04 | -0.12 | -0.12 | -0.12 |
| | | | | 18 demes | -0.33 | -0.33 | -0.30 | -0.30 | -0.31 | -0.17 | -0.16 | -0.15 | -0.15 | -0.15 | -0.15 | -0.15 | -0.15 | 0.00 | 0.00 | -0.01 |

*p*-value > 0.05/7980 (Bonferroni correction)

**|Spearman correlation coefficient| >= 0.5**

| part 2 | | | | variable 2: | max. \|estimated admixture fractions - equal fractions\| | | | | | *p*-value | | | | | max. SE | | | *p*-value | | |
|---|---|---|---|---|---|---|---|---|---|---|---|---|---|---|---|---|---|---|---|---|
| total number of randomized models: | | | | | 21,663,216 | 12,439,194 | 2,392,357 | 2,040,355 | 1,058,613 | 21,663,216 | 12,439,194 | 2,392,357 | 2,040,355 | 1,058,613 | 4,378,913 | 3,566,530 | 1,533,911 | 4,378,913 | 3,566,530 | 1,533,911 |
| total number of systematic models (10-5 to 10-2): | | | | | 3,576,667 | 1,158,133 | 1,267,603 | 1,064,846 | 240,961 | 3,576,667 | 1,158,133 | 1,267,603 | 1,064,846 | 240,961 | N/A | | | N/A | | |
| variable 1: | model complexity | gene-flow intensity per gen. | *qpAdm* protocol | sampling density | p >= 1e-55 | p >= 1e-5 | max. \|*EAF–EF*\| <= 0.5 | p >= 1e-55; max. \|*EAF–EF*\| <= 0.5 | p >= 1e-5; max. \|*EAF–EF*\| <= 0.5 | p >= 1e-55 | p >= 1e-5 | max. \|*EAF–EF*\| <= 0.5 | p >= 1e-55; max. \|*EAF–EF*\| <= 0.5 | p >= 1e-5; max. \|*EAF–EF*\| <= 0.5 | max. SE <= 0.1 | p >= 1e-55; max. SE <= 0.1 | p >= 1e-5; max. SE <= 0.1 | max. SE <= 0.1 | p >= 1e-55; max. SE <= 0.1 | p >= 1e-5; max. SE <= 0.1 |
| maximal source-target distance | 2-way | 10-5 to 10-4 | dist. rot. | 13 demes | 0.01 | 0.02 | -0.03 | -0.03 | -0.01 | -0.04 | -0.02 | -0.04 | -0.05 | -0.02 | 0.05 | 0.06 | 0.11 | -0.05 | -0.06 | -0.03 |
| | | | | 18 demes | 0.06 | 0.08 | 0.00 | -0.01 | 0.01 | -0.08 | -0.04 | -0.08 | -0.09 | -0.06 | -0.04 | -0.03 | 0.06 | -0.11 | -0.11 | -0.03 |
| | | | prox. rot. | 13 demes | 0.00 | 0.00 | 0.01 | 0.00 | 0.02 | -0.04 | -0.02 | -0.04 | -0.03 | -0.03 | -0.03 | -0.02 | 0.04 | -0.05 | -0.04 | -0.03 |
| | | | | 18 demes | -0.01 | -0.03 | 0.02 | 0.02 | 0.02 | -0.06 | -0.01 | -0.03 | -0.03 | -0.02 | -0.03 | -0.03 | 0.07 | -0.04 | -0.05 | -0.01 |
| | | | dist. non-rot. | 13 demes | 0.05 | 0.05 | 0.02 | 0.03 | 0.01 | -0.07 | -0.01 | -0.07 | -0.06 | -0.03 | -0.08 | -0.08 | 0.01 | -0.10 | -0.11 | -0.05 |
| | | | | 18 demes | 0.04 | 0.08 | 0.00 | 0.00 | -0.05 | -0.06 | -0.04 | -0.05 | -0.06 | -0.04 | -0.05 | -0.05 | 0.07 | -0.07 | -0.09 | -0.04 |
| | | | prox. non-rot. | 13 demes | 0.02 | 0.00 | 0.07 | 0.07 | 0.05 | -0.02 | 0.02 | 0.00 | 0.00 | 0.05 | -0.06 | -0.06 | -0.01 | -0.02 | -0.02 | 0.06 |
| | | | | 18 demes | 0.01 | 0.04 | 0.04 | 0.05 | 0.07 | -0.06 | 0.00 | -0.06 | -0.08 | -0.01 | -0.05 | -0.04 | 0.06 | -0.05 | -0.07 | 0.01 |
| | | 10-4 to 10-3 | dist. rot. | 13 demes | 0.14 | 0.14 | 0.17 | 0.26 | 0.38 | -0.14 | -0.06 | -0.24 | -0.18 | -0.06 | -0.12 | -0.20 | -0.40 | -0.23 | -0.14 | -0.02 |
| | | | | 18 demes | 0.18 | 0.14 | 0.24 | 0.41 | 0.46 | -0.10 | 0.00 | -0.33 | -0.13 | -0.03 | -0.16 | -0.38 | **-0.59** | -0.31 | -0.09 | -0.02 |
| | | | prox. rot. | 13 demes | 0.15 | 0.12 | 0.15 | 0.23 | 0.37 | -0.15 | -0.05 | -0.25 | -0.15 | -0.08 | -0.10 | -0.18 | -0.42 | -0.25 | -0.12 | -0.03 |
| | | | | 18 demes | 0.16 | 0.09 | 0.23 | 0.43 | **0.50** | -0.09 | 0.00 | -0.28 | -0.09 | 0.06 | -0.13 | -0.36 | **-0.53** | -0.28 | -0.06 | -0.01 |
| | | | dist. non-rot. | 13 demes | 0.15 | 0.15 | 0.16 | 0.23 | 0.43 | -0.15 | -0.03 | -0.22 | -0.13 | 0.02 | -0.10 | -0.21 | -0.39 | -0.25 | -0.14 | 0.04 |
| | | | | 18 demes | 0.16 | 0.16 | 0.14 | 0.31 | 0.39 | -0.08 | -0.02 | -0.26 | -0.06 | -0.03 | -0.12 | -0.30 | -0.48 | -0.25 | -0.05 | -0.02 |
| | | | prox. non-rot. | 13 demes | 0.10 | 0.06 | 0.18 | 0.25 | 0.33 | -0.15 | -0.11 | -0.21 | -0.13 | -0.12 | -0.12 | -0.20 | -0.34 | -0.23 | -0.13 | -0.10 |
| | | | | 18 demes | 0.15 | 0.14 | 0.21 | 0.40 | 0.45 | -0.09 | -0.03 | -0.25 | -0.05 | 0.02 | -0.15 | -0.36 | **-0.56** | -0.26 | -0.06 | 0.00 |
| | | 10-5 to 10-2 | dist. rot. | 13 demes | 0.25 | 0.18 | 0.30 | 0.43 | **0.55** | -0.20 | -0.08 | -0.33 | -0.12 | -0.04 | -0.18 | -0.37 | **-0.62** | -0.30 | -0.15 | -0.05 |
| | | | | 18 demes | 0.26 | 0.17 | 0.34 | **0.53** | **0.60** | -0.17 | -0.07 | -0.34 | -0.09 | -0.03 | -0.18 | -0.47 | **-0.72** | -0.32 | -0.14 | -0.06 |
| | | | | systematic | 0.18 | **0.67** | 0.11 | 0.28 | **0.70** | -0.17 | -0.08 | -0.30 | -0.32 | -0.23 | | | | | | |
| | | | prox. rot. | 13 demes | 0.23 | 0.19 | 0.29 | 0.44 | **0.58** | -0.20 | -0.05 | -0.35 | -0.16 | -0.01 | -0.18 | -0.35 | **-0.60** | -0.30 | -0.15 | -0.05 |
| | | | | 18 demes | 0.22 | 0.14 | 0.37 | **0.55** | **0.62** | -0.19 | -0.10 | -0.35 | -0.13 | -0.04 | -0.19 | -0.49 | **-0.72** | -0.32 | -0.16 | -0.06 |
| | | | dist. non-rot. | 13 demes | 0.28 | 0.21 | 0.31 | 0.42 | **0.54** | -0.21 | -0.05 | -0.32 | -0.11 | -0.01 | -0.18 | -0.35 | **-0.63** | -0.34 | -0.18 | -0.02 |
| | | | | 18 demes | 0.29 | 0.22 | 0.35 | **0.52** | **0.58** | -0.17 | -0.05 | -0.33 | -0.06 | 0.03 | -0.20 | -0.49 | **-0.75** | -0.35 | -0.14 | 0.00 |
| | | | | systematic | 0.19 | **0.65** | 0.08 | 0.29 | **0.78** | -0.21 | -0.10 | -0.31 | -0.31 | -0.12 | | | | | | |
| | | | prox. non-rot. | 13 demes | 0.25 | 0.14 | 0.32 | 0.44 | **0.64** | -0.22 | -0.12 | -0.35 | -0.14 | -0.04 | -0.19 | -0.35 | **-0.58** | -0.31 | -0.15 | -0.08 |
| | | | | 18 demes | 0.27 | 0.17 | 0.35 | **0.54** | **0.68** | -0.16 | -0.11 | -0.41 | -0.11 | -0.09 | -0.17 | -0.45 | **-0.68** | -0.34 | -0.11 | -0.11 |
| | | 10-3 to 10-2 | dist. rot. | 13 demes | 0.29 | 0.27 | 0.39 | 0.40 | **0.55** | -0.31 | -0.11 | -0.33 | -0.29 | -0.07 | -0.20 | -0.22 | -0.44 | -0.26 | -0.24 | -0.06 |
| | | | | 18 demes | 0.30 | 0.27 | 0.40 | 0.45 | **0.60** | -0.29 | -0.12 | -0.36 | -0.26 | -0.08 | -0.19 | -0.25 | **-0.51** | -0.32 | -0.25 | -0.08 |
| | | | prox. rot. | 13 demes | 0.27 | 0.27 | 0.34 | 0.36 | **0.52** | -0.29 | -0.12 | -0.38 | -0.34 | -0.11 | -0.17 | -0.19 | -0.37 | -0.25 | -0.22 | -0.06 |
| | | | | 18 demes | 0.29 | 0.24 | 0.38 | 0.44 | **0.59** | -0.28 | -0.09 | -0.40 | -0.29 | -0.05 | -0.17 | -0.22 | -0.47 | -0.27 | -0.21 | -0.04 |
| | | | dist. non-rot. | 13 demes | 0.29 | 0.31 | 0.37 | 0.38 | 0.49 | -0.34 | -0.15 | -0.38 | -0.36 | -0.11 | -0.17 | -0.18 | -0.32 | -0.29 | -0.28 | -0.09 |
| | | | | 18 demes | 0.31 | 0.33 | 0.40 | 0.43 | **0.58** | -0.31 | -0.15 | -0.38 | -0.32 | -0.11 | -0.19 | -0.22 | -0.45 | -0.32 | -0.28 | -0.10 |
| | | | prox. non-rot. | 13 demes | 0.29 | 0.30 | 0.35 | 0.36 | **0.52** | -0.31 | -0.14 | -0.39 | -0.37 | -0.13 | -0.17 | -0.17 | -0.29 | -0.25 | -0.24 | -0.07 |
| | | | | 18 demes | 0.30 | 0.29 | 0.39 | 0.41 | **0.52** | -0.30 | -0.14 | -0.41 | -0.35 | -0.10 | -0.16 | -0.18 | -0.36 | -0.29 | -0.26 | -0.10 |
| | 3-way | 10-5 to 10-4 | dist. rot. | 13 demes | -0.02 | -0.01 | -0.07 | -0.06 | -0.02 | -0.01 | -0.02 | -0.04 | -0.04 | -0.04 | 0.12 | 0.12 | 0.14 | -0.10 | -0.10 | -0.11 |
| | | | | 18 demes | 0.06 | 0.05 | -0.06 | -0.06 | -0.03 | -0.03 | -0.03 | -0.03 | -0.03 | -0.05 | 0.03 | 0.04 | 0.04 | -0.04 | -0.04 | -0.05 |
| | | | prox. rot. | 13 demes | -0.02 | -0.01 | -0.04 | -0.04 | 0.02 | -0.01 | -0.02 | 0.00 | -0.01 | -0.02 | 0.04 | 0.04 | 0.12 | -0.09 | -0.09 | -0.05 |
| | | | | 18 demes | -0.03 | -0.05 | -0.06 | -0.06 | -0.03 | -0.03 | -0.01 | 0.01 | 0.01 | -0.02 | 0.01 | 0.01 | 0.05 | -0.05 | -0.05 | -0.05 |
| | | | dist. non-rot. | 13 demes | 0.03 | 0.03 | -0.01 | -0.01 | 0.00 | -0.04 | -0.02 | 0.00 | 0.01 | -0.02 | -0.05 | -0.05 | 0.01 | -0.10 | -0.11 | -0.10 |
| | | | | 18 demes | 0.04 | 0.06 | -0.06 | -0.07 | -0.04 | -0.03 | -0.01 | 0.01 | 0.01 | -0.02 | -0.02 | -0.02 | 0.06 | -0.07 | -0.07 | -0.03 |
| | | | prox. non-rot. | 13 demes | 0.00 | -0.03 | 0.02 | 0.02 | 0.01 | 0.01 | 0.01 | 0.08 | 0.09 | 0.08 | -0.01 | -0.01 | 0.03 | -0.02 | -0.02 | -0.01 |
| | | | | 18 demes | 0.00 | 0.02 | -0.05 | -0.05 | 0.03 | -0.03 | 0.00 | -0.08 | -0.09 | 0.00 | -0.01 | 0.00 | 0.04 | -0.04 | -0.05 | 0.01 |
| | | 10-4 to 10-3 | dist. rot. | 13 demes | 0.10 | 0.09 | 0.07 | 0.07 | 0.05 | -0.10 | -0.07 | -0.15 | -0.10 | -0.15 | -0.08 | -0.10 | -0.18 | -0.12 | -0.07 | -0.05 |
| | | | | 18 demes | 0.14 | 0.09 | 0.07 | 0.09 | 0.11 | -0.07 | -0.05 | -0.20 | -0.06 | -0.12 | -0.14 | -0.21 | -0.29 | -0.16 | -0.04 | -0.07 |
| | | | prox. rot. | 13 demes | 0.11 | 0.08 | 0.04 | 0.05 | 0.05 | -0.10 | -0.07 | -0.13 | -0.07 | -0.12 | -0.08 | -0.10 | -0.17 | -0.11 | -0.05 | -0.05 |
| | | | | 18 demes | 0.12 | 0.08 | 0.04 | 0.07 | 0.04 | -0.07 | -0.04 | -0.17 | -0.12 | -0.06 | -0.13 | -0.20 | -0.23 | -0.12 | -0.02 | -0.02 |
| | | | dist. non-rot. | 13 demes | 0.11 | 0.08 | 0.02 | 0.02 | 0.02 | -0.09 | -0.03 | -0.15 | -0.12 | -0.09 | -0.10 | -0.14 | -0.17 | -0.10 | -0.05 | 0.02 |
| | | | | 18 demes | 0.10 | 0.07 | 0.02 | 0.06 | 0.09 | -0.04 | -0.06 | -0.14 | -0.04 | -0.16 | -0.10 | -0.17 | -0.21 | -0.11 | -0.02 | -0.07 |
| | | | prox. non-rot. | 13 demes | 0.09 | 0.04 | 0.04 | 0.06 | 0.02 | -0.11 | -0.09 | -0.10 | -0.08 | -0.14 | -0.09 | -0.12 | -0.18 | -0.09 | -0.06 | -0.08 |
| | | | | 18 demes | 0.10 | 0.09 | 0.07 | 0.10 | 0.06 | -0.06 | -0.07 | -0.15 | -0.06 | -0.13 | -0.12 | -0.20 | -0.27 | -0.13 | -0.01 | -0.05 |
| | | 10-5 to 10-2 | dist. rot. | 13 demes | 0.19 | 0.15 | 0.23 | 0.23 | 0.19 | -0.17 | -0.07 | -0.13 | -0.08 | -0.05 | -0.12 | -0.14 | -0.23 | -0.11 | -0.07 | 0.00 |
| | | | | 18 demes | 0.20 | 0.14 | 0.25 | 0.22 | 0.15 | -0.15 | -0.06 | -0.17 | -0.11 | -0.02 | -0.14 | -0.21 | -0.30 | -0.14 | -0.07 | -0.03 |
| | | | | systematic | 0.06 | 0.17 | 0.05 | 0.07 | 0.26 | -0.08 | -0.02 | -0.16 | -0.15 | -0.13 | | | | | | |
| | | | prox. rot. | 13 demes | 0.16 | 0.14 | 0.20 | 0.19 | 0.15 | -0.17 | -0.05 | -0.15 | -0.12 | -0.07 | -0.14 | -0.16 | -0.26 | -0.11 | -0.08 | 0.00 |
| | | | | 18 demes | 0.17 | 0.11 | 0.25 | 0.22 | 0.14 | -0.16 | -0.07 | -0.13 | -0.08 | -0.07 | -0.15 | -0.21 | -0.32 | -0.15 | -0.09 | -0.01 |
| | | | dist. non-rot. | 13 demes | 0.22 | 0.17 | 0.24 | 0.24 | 0.18 | -0.18 | -0.05 | -0.18 | -0.13 | -0.09 | -0.12 | -0.14 | -0.24 | -0.14 | -0.08 | 0.01 |
| | | | | 18 demes | 0.23 | 0.17 | 0.23 | 0.23 | 0.15 | -0.16 | -0.05 | -0.24 | -0.16 | -0.09 | -0.16 | -0.21 | -0.33 | -0.19 | -0.09 | 0.03 |
| | | | | systematic | 0.03 | 0.13 | 0.01 | 0.02 | 0.19 | -0.11 | -0.09 | -0.15 | -0.15 | -0.13 | | | | | | |
| | | | prox. non-rot. | 13 demes | 0.18 | 0.12 | 0.23 | 0.23 | 0.20 | -0.18 | -0.09 | -0.16 | -0.12 | -0.13 | -0.14 | -0.15 | -0.24 | -0.08 | -0.04 | 0.02 |
| | | | | 18 demes | 0.22 | 0.16 | 0.22 | 0.22 | 0.16 | -0.15 | -0.10 | -0.22 | -0.10 | -0.06 | -0.14 | -0.20 | -0.32 | -0.17 | -0.06 | -0.08 |
| | | 10-3 to 10-2 | dist. rot. | 13 demes | 0.19 | 0.19 | 0.34 | 0.34 | 0.32 | -0.21 | -0.11 | -0.10 | -0.10 | -0.04 | -0.06 | -0.06 | -0.08 | -0.09 | -0.09 | -0.02 |
| | | | | 18 demes | 0.20 | 0.20 | 0.36 | 0.36 | 0.34 | -0.25 | -0.12 | -0.16 | -0.16 | -0.08 | -0.09 | -0.09 | -0.14 | -0.15 | -0.15 | -0.03 |
| | | | prox. rot. | 13 demes | 0.17 | 0.19 | 0.33 | 0.33 | 0.33 | -0.19 | -0.11 | -0.11 | -0.11 | -0.05 | -0.08 | -0.08 | -0.10 | -0.08 | -0.08 | 0.00 |
| | | | | 18 demes | 0.18 | 0.19 | 0.38 | 0.38 | 0.36 | -0.24 | -0.11 | -0.13 | -0.13 | -0.06 | -0.05 | -0.05 | -0.09 | -0.13 | -0.13 | -0.01 |
| | | | dist. non-rot. | 13 demes | 0.21 | 0.21 | 0.34 | 0.34 | 0.33 | -0.24 | -0.18 | -0.15 | -0.15 | -0.12 | -0.06 | -0.06 | -0.07 | -0.17 | -0.17 | -0.10 |
| | | | | 18 demes | 0.20 | 0.21 | 0.35 | 0.35 | 0.34 | -0.27 | -0.17 | -0.18 | -0.18 | -0.13 | -0.07 | -0.07 | -0.09 | -0.18 | -0.18 | -0.08 |
| | | | prox. non-rot. | 13 demes | 0.19 | 0.20 | 0.35 | 0.35 | 0.35 | -0.23 | -0.17 | -0.13 | -0.13 | -0.11 | -0.05 | -0.05 | -0.06 | -0.15 | -0.15 | -0.09 |
| | | | | 18 demes | 0.19 | 0.20 | 0.36 | 0.36 | 0.35 | -0.28 | -0.19 | -0.16 | -0.16 | -0.13 | -0.02 | -0.02 | -0.05 | -0.21 | -0.21 | -0.10 |
| | 4-way | 10-5 to 10-4 | dist. rot. | 13 demes | -0.02 | -0.02 | -0.04 | -0.04 | -0.03 | 0.00 | -0.01 | -0.02 | -0.02 | -0.02 | 0.08 | 0.08 | 0.08 | -0.15 | -0.15 | -0.14 |
| | | | | 18 demes | 0.05 | 0.03 | -0.07 | -0.07 | -0.05 | -0.02 | -0.01 | -0.03 | -0.02 | -0.04 | 0.04 | 0.03 | 0.00 | -0.01 | -0.01 | -0.04 |
| | | | prox. rot. | 13 demes | -0.03 | -0.02 | 0.00 | 0.00 | 0.02 | -0.01 | -0.01 | -0.02 | -0.02 | -0.01 | 0.11 | 0.11 | 0.11 | -0.17 | -0.17 | -0.16 |
| | | | | 18 demes | -0.04 | -0.05 | -0.07 | -0.07 | -0.04 | -0.02 | -0.01 | 0.02 | 0.01 | -0.02 | 0.01 | 0.02 | 0.03 | -0.09 | -0.09 | -0.12 |
| | | | dist. non-rot. | 13 demes | 0.02 | 0.02 | 0.02 | 0.02 | 0.02 | -0.03 | -0.03 | 0.01 | 0.01 | -0.01 | -0.03 | -0.03 | -0.02 | -0.10 | -0.10 | -0.14 |
| | | | | 18 demes | 0.03 | 0.03 | -0.07 | -0.07 | -0.06 | -0.01 | -0.01 | 0.04 | 0.04 | -0.05 | 0.00 | 0.01 | 0.04 | -0.07 | -0.08 | -0.08 |
| | | | prox. non-rot. | 13 demes | -0.01 | -0.03 | 0.05 | 0.05 | 0.06 | 0.00 | 0.00 | 0.10 | 0.09 | 0.09 | 0.00 | -0.01 | 0.05 | -0.06 | -0.06 | -0.06 |
| | | | | 18 demes | -0.01 | 0.00 | -0.08 | -0.09 | -0.03 | -0.01 | 0.01 | -0.08 | -0.09 | -0.02 | 0.01 | 0.01 | 0.00 | -0.03 | -0.03 | 0.02 |
| | | 10-4 to 10-3 | dist. rot. | 13 demes | 0.08 | 0.08 | 0.01 | 0.00 | 0.04 | -0.08 | -0.06 | -0.04 | -0.03 | -0.10 | -0.07 | -0.07 | -0.11 | -0.04 | -0.02 | -0.01 |
| | | | | 18 demes | 0.13 | 0.09 | 0.03 | 0.04 | 0.06 | -0.07 | -0.08 | -0.08 | 0.00 | -0.07 | -0.12 | -0.14 | -0.17 | -0.07 | -0.03 | -0.09 |
| | | | prox. rot. | 13 demes | 0.09 | 0.08 | 0.01 | 0.01 | 0.03 | -0.08 | -0.07 | -0.07 | -0.05 | -0.10 | -0.08 | -0.08 | -0.08 | -0.04 | -0.02 | -0.04 |
| | | | | 18 demes | 0.11 | 0.07 | 0.00 | 0.01 | 0.03 | -0.08 | -0.06 | -0.08 | -0.07 | -0.04 | -0.10 | -0.12 | -0.13 | -0.03 | -0.01 | -0.02 |
| | | | dist. non-rot. | 13 demes | 0.10 | 0.07 | 0.03 | 0.03 | 0.04 | -0.07 | -0.02 | -0.06 | -0.05 | -0.05 | -0.10 | -0.11 | -0.10 | -0.01 | 0.00 | 0.02 |
| | | | | 18 demes | 0.09 | 0.05 | 0.00 | 0.02 | 0.02 | -0.04 | -0.06 | -0.06 | 0.01 | -0.09 | -0.09 | -0.12 | -0.11 | -0.03 | -0.01 | -0.09 |
| | | | prox. non-rot. | 13 demes | 0.09 | 0.04 | 0.06 | 0.06 | 0.10 | -0.09 | -0.08 | -0.04 | -0.03 | -0.14 | -0.08 | -0.09 | -0.12 | -0.01 | -0.01 | -0.06 |
| | | | | 18 demes | 0.10 | 0.06 | 0.02 | 0.03 | 0.03 | -0.06 | -0.10 | -0.04 | 0.00 | -0.10 | -0.10 | -0.13 | -0.15 | -0.04 | 0.01 | -0.10 |
| | | 10-5 to 10-2 | dist. rot. | 13 demes | 0.16 | 0.14 | 0.12 | 0.12 | 0.09 | -0.12 | -0.06 | -0.01 | -0.01 | -0.01 | -0.07 | -0.07 | -0.10 | 0.01 | 0.02 | 0.03 |
| | | | | 18 demes | 0.18 | 0.14 | 0.13 | 0.12 | 0.06 | -0.14 | -0.08 | -0.02 | -0.01 | 0.01 | -0.11 | -0.12 | -0.17 | -0.02 | -0.01 | -0.02 |
| | | | | systematic | 0.02 | 0.02 | 0.05 | 0.05 | 0.08 | -0.02 | -0.01 | -0.05 | -0.05 | -0.06 | | | | | | |
| | | | prox. rot. | 13 demes | 0.13 | 0.13 | 0.08 | 0.08 | 0.04 | -0.13 | -0.06 | -0.05 | -0.05 | -0.05 | -0.12 | -0.12 | -0.15 | 0.00 | 0.00 | 0.04 |
| | | | | 18 demes | 0.16 | 0.12 | 0.10 | 0.09 | 0.05 | -0.15 | -0.08 | -0.04 | -0.03 | -0.05 | -0.11 | -0.12 | -0.17 | -0.03 | -0.03 | 0.02 |
| | | | dist. non-rot. | 13 demes | 0.22 | 0.18 | 0.13 | 0.13 | 0.07 | -0.15 | -0.06 | -0.02 | -0.02 | -0.01 | -0.08 | -0.08 | -0.15 | 0.00 | 0.01 | 0.03 |
| | | | | 18 demes | 0.21 | 0.18 | 0.14 | 0.13 | 0.09 | -0.15 | -0.05 | -0.11 | -0.10 | -0.03 | -0.11 | -0.12 | -0.20 | -0.07 | -0.04 | 0.04 |
| | | | | systematic | 0.01 | 0.03 | 0.01 | 0.01 | 0.07 | -0.04 | -0.02 | -0.06 | -0.06 | -0.03 | | | | | | |
| | | | prox. non-rot. | 13 demes | 0.15 | 0.13 | 0.12 | 0.12 | 0.08 | -0.14 | -0.08 | -0.09 | -0.08 | -0.12 | -0.09 | -0.09 | -0.14 | 0.01 | 0.02 | 0.03 |
| | | | | 18 demes | 0.20 | 0.17 | 0.10 | 0.09 | 0.04 | -0.15 | -0.10 | -0.10 | -0.08 | -0.07 | -0.11 | -0.12 | -0.19 | -0.05 | -0.02 | -0.07 |
| | | 10-3 to 10-2 | dist. rot. | 13 demes | 0.17 | 0.17 | 0.22 | 0.22 | 0.22 | -0.11 | -0.10 | 0.00 | 0.00 | 0.01 | 0.04 | 0.04 | 0.03 | 0.09 | 0.09 | 0.10 |
| | | | | 18 demes | 0.18 | 0.18 | 0.21 | 0.21 | 0.20 | -0.16 | -0.12 | -0.08 | -0.08 | -0.07 | -0.02 | -0.02 | -0.02 | 0.01 | 0.01 | 0.02 |
| | | | prox. rot. | 13 demes | 0.17 | 0.17 | 0.20 | 0.20 | 0.20 | -0.09 | -0.07 | 0.01 | 0.01 | 0.02 | 0.05 | 0.05 | 0.05 | 0.18 | 0.18 | 0.17 |
| | | | | 18 demes | 0.18 | 0.17 | 0.24 | 0.24 | 0.24 | -0.16 | -0.12 | -0.07 | -0.07 | -0.05 | 0.01 | 0.01 | 0.00 | 0.00 | 0.00 | 0.00 |
| | | | dist. non-rot. | 13 demes | 0.19 | 0.19 | 0.24 | 0.24 | 0.24 | -0.15 | -0.14 | -0.12 | -0.12 | -0.11 | -0.05 | -0.05 | -0.05 | 0.03 | 0.03 | 0.05 |
| | | | | 18 demes | 0.19 | 0.19 | 0.19 | 0.19 | 0.19 | -0.17 | -0.15 | -0.12 | -0.12 | -0.11 | -0.01 | -0.01 | -0.02 | -0.02 | -0.02 | 0.00 |
| | | | prox. non-rot. | 13 demes | 0.18 | 0.18 | 0.19 | 0.19 | 0.19 | -0.15 | -0.14 | -0.15 | -0.15 | -0.15 | -0.22 | -0.22 | -0.22 | -0.08 | -0.08 | -0.08 |
| | | | | 18 demes | 0.20 | 0.20 | 0.26 | 0.26 | 0.26 | -0.19 | -0.18 | -0.14 | -0.14 | -0.13 | -0.03 | -0.03 | -0.03 | -0.06 | -0.06 | -0.05 |

*p*-value > 0.05/7980 (Bonferroni correction)

**|Spearman correlation coefficient| >= 0.5**

| part 3 | | | | variable 2: | max. \|estimated admixture fractions - equal fractions\| | | | | | *p*-value | | | | | max. SE | | | *p*-value | | |
|---|---|---|---|---|---|---|---|---|---|---|---|---|---|---|---|---|---|---|---|---|
| total number of randomized models: | | | | | 21,663,216 | 12,439,194 | 2,392,357 | 2,040,355 | 1,058,613 | 21,663,216 | 12,439,194 | 2,392,357 | 2,040,355 | 1,058,613 | 4,378,913 | 3,566,530 | 1,533,911 | 4,378,913 | 3,566,530 | 1,533,911 |
| total number of systematic models (10-5 to 10-2): | | | | | 3,576,667 | 1,158,133 | 1,267,603 | 1,064,846 | 240,961 | 3,576,667 | 1,158,133 | 1,267,603 | 1,064,846 | 240,961 | N/A | | | N/A | | |
| variable 1: | model complexity | gene-flow intensity per gen. | *qpAdm* protocol | sampling density | p >= 1e-55 | p >= 1e-5 | max. \|*EAF–EF*\| <= 0.5 | p >= 1e-55; max. \|*EAF–EF*\| <= 0.5 | p >= 1e-5; max. \|*EAF–EF*\| <= 0.5 | p >= 1e-55 | p >= 1e-5 | max. \|*EAF–EF*\| <= 0.5 | p >= 1e-55; max. \|*EAF–EF*\| <= 0.5 | p >= 1e-5; max. \|*EAF–EF*\| <= 0.5 | max. SE <= 0.1 | p >= 1e-55; max. SE <= 0.1 | p >= 1e-5; max. SE <= 0.1 | max. SE <= 0.1 | p >= 1e-55; max. SE <= 0.1 | p >= 1e-5; max. SE <= 0.1 |
| average source-target distance | 2-way | 10-5 to 10-4 | dist. rot. | 13 demes | 0.09 | 0.16 | -0.02 | -0.02 | -0.01 | -0.09 | -0.06 | -0.13 | -0.13 | -0.07 | 0.01 | 0.02 | 0.18 | -0.18 | -0.19 | -0.07 |
| | | | | 18 demes | 0.15 | 0.26 | 0.01 | -0.01 | -0.05 | -0.15 | -0.09 | -0.22 | -0.23 | -0.12 | -0.08 | -0.06 | 0.20 | -0.29 | -0.31 | -0.11 |
| | | | prox. rot. | 13 demes | 0.08 | 0.14 | 0.01 | 0.00 | 0.02 | -0.08 | -0.03 | -0.12 | -0.12 | -0.04 | -0.05 | -0.04 | 0.12 | -0.15 | -0.15 | -0.04 |
| | | | | 18 demes | 0.08 | 0.14 | 0.04 | 0.02 | -0.02 | -0.12 | -0.05 | -0.14 | -0.14 | -0.07 | -0.06 | -0.05 | 0.16 | -0.18 | -0.20 | -0.08 |
| | | | dist. non-rot. | 13 demes | 0.13 | 0.17 | 0.04 | 0.04 | -0.01 | -0.13 | -0.03 | -0.15 | -0.15 | -0.05 | -0.12 | -0.12 | 0.05 | -0.22 | -0.25 | -0.07 |
| | | | | 18 demes | 0.14 | 0.25 | 0.03 | 0.02 | -0.16 | -0.14 | -0.11 | -0.18 | -0.20 | -0.13 | -0.07 | -0.07 | 0.16 | -0.22 | -0.26 | -0.12 |
| | | | prox. non-rot. | 13 demes | 0.10 | 0.14 | 0.07 | 0.07 | 0.05 | -0.06 | -0.01 | -0.10 | -0.10 | 0.00 | -0.07 | -0.07 | 0.06 | -0.11 | -0.13 | 0.02 |
| | | | | 18 demes | 0.10 | 0.21 | 0.08 | 0.08 | 0.03 | -0.12 | -0.01 | -0.15 | -0.19 | -0.05 | -0.06 | -0.05 | 0.15 | -0.15 | -0.20 | -0.05 |
| | | 10-4 to 10-3 | dist. rot. | 13 demes | 0.35 | 0.42 | 0.10 | 0.15 | 0.15 | -0.24 | -0.18 | -0.44 | -0.32 | -0.22 | -0.02 | -0.06 | -0.23 | -0.45 | -0.32 | -0.20 |
| | | | | 18 demes | 0.44 | 0.42 | 0.13 | 0.24 | 0.28 | -0.22 | -0.08 | **-0.54** | -0.30 | -0.17 | -0.03 | -0.22 | -0.49 | **-0.56** | -0.30 | -0.14 |
| | | | prox. rot. | 13 demes | 0.38 | 0.42 | 0.08 | 0.10 | 0.18 | -0.25 | -0.19 | -0.46 | -0.31 | -0.28 | 0.00 | -0.03 | -0.25 | -0.48 | -0.33 | -0.22 |
| | | | | 18 demes | 0.42 | 0.43 | 0.14 | 0.30 | 0.34 | -0.19 | -0.05 | **-0.53** | -0.23 | -0.12 | -0.02 | -0.21 | -0.42 | **-0.55** | -0.25 | -0.17 |
| | | | dist. non-rot. | 13 demes | 0.36 | 0.34 | 0.07 | 0.06 | 0.21 | -0.30 | -0.26 | -0.45 | -0.33 | -0.13 | -0.01 | -0.06 | -0.21 | -0.49 | -0.34 | -0.15 |
| | | | | 18 demes | 0.40 | 0.29 | 0.06 | 0.14 | 0.26 | -0.25 | -0.20 | -0.49 | -0.26 | -0.13 | -0.01 | -0.14 | -0.36 | **-0.51** | -0.24 | -0.14 |
| | | | prox. non-rot. | 13 demes | 0.32 | 0.34 | 0.13 | 0.15 | 0.15 | -0.22 | -0.15 | -0.41 | -0.26 | -0.25 | -0.05 | -0.09 | -0.18 | -0.44 | -0.27 | -0.24 |
| | | | | 18 demes | 0.41 | 0.46 | 0.13 | 0.25 | 0.31 | -0.18 | -0.08 | -0.48 | -0.22 | -0.10 | -0.04 | -0.21 | -0.46 | **-0.50** | -0.24 | -0.12 |
| | | 10-5 to 10-2 | dist. rot. | 13 demes | 0.46 | 0.44 | 0.07 | 0.20 | 0.36 | -0.31 | -0.16 | **-0.61** | -0.36 | -0.16 | -0.01 | -0.22 | **-0.51** | **-0.59** | -0.38 | -0.18 |
| | | | | 18 demes | 0.49 | 0.44 | 0.11 | 0.31 | 0.44 | -0.27 | -0.15 | **-0.65** | -0.31 | -0.15 | 0.01 | -0.35 | **-0.62** | **-0.62** | -0.33 | -0.16 |
| | | | | systematic | 0.30 | **0.86** | -0.27 | -0.12 | **0.69** | -0.30 | -0.02 | **-0.74** | **-0.54** | -0.23 | | | | | | |
| | | | prox. rot. | 13 demes | 0.44 | 0.46 | 0.09 | 0.24 | 0.38 | -0.28 | -0.11 | **-0.61** | -0.35 | -0.14 | -0.03 | -0.23 | **-0.51** | **-0.57** | -0.34 | -0.16 |
| | | | | 18 demes | 0.46 | 0.42 | 0.14 | 0.36 | 0.46 | -0.27 | -0.15 | **-0.64** | -0.31 | -0.16 | -0.02 | -0.37 | **-0.62** | **-0.62** | -0.34 | -0.17 |
| | | | dist. non-rot. | 13 demes | 0.46 | 0.42 | 0.08 | 0.16 | 0.32 | -0.35 | -0.19 | **-0.61** | -0.37 | -0.19 | -0.01 | -0.23 | **-0.52** | **-0.61** | -0.42 | -0.19 |
| | | | | 18 demes | **0.50** | 0.41 | 0.09 | 0.28 | 0.41 | -0.32 | -0.22 | **-0.67** | -0.32 | -0.12 | 0.00 | -0.35 | **-0.66** | **-0.66** | -0.38 | -0.16 |
| | | | | systematic | 0.10 | **0.71** | -0.38 | -0.28 | **0.77** | -0.47 | -0.10 | **-0.75** | **-0.63** | -0.12 | | | | | | |
| | | | prox. non-rot. | 13 demes | 0.44 | 0.37 | 0.09 | 0.20 | 0.48 | -0.33 | -0.18 | **-0.63** | -0.36 | -0.16 | -0.03 | -0.23 | -0.47 | **-0.60** | -0.36 | -0.20 |
| | | | | 18 demes | **0.51** | 0.47 | 0.13 | 0.35 | **0.54** | -0.25 | -0.12 | **-0.68** | -0.30 | -0.21 | 0.02 | -0.35 | **-0.60** | **-0.64** | -0.30 | -0.23 |
| | | 10-3 to 10-2 | dist. rot. | 13 demes | 0.45 | 0.48 | 0.11 | 0.12 | 0.31 | -0.48 | -0.18 | **-0.65** | **-0.61** | -0.22 | -0.09 | -0.13 | -0.43 | **-0.61** | **-0.58** | -0.21 |
| | | | | 18 demes | 0.47 | 0.49 | 0.09 | 0.14 | 0.37 | -0.45 | -0.17 | **-0.69** | **-0.59** | -0.20 | -0.06 | -0.16 | -0.49 | **-0.65** | **-0.56** | -0.20 |
| | | | prox. rot. | 13 demes | 0.44 | 0.49 | 0.09 | 0.10 | 0.31 | -0.45 | -0.15 | **-0.65** | **-0.61** | -0.20 | -0.09 | -0.13 | -0.36 | **-0.57** | **-0.54** | -0.16 |
| | | | | 18 demes | 0.46 | 0.48 | 0.11 | 0.16 | 0.37 | -0.42 | -0.14 | **-0.68** | **-0.57** | -0.16 | -0.07 | -0.15 | -0.45 | **-0.61** | **-0.51** | -0.14 |
| | | | dist. non-rot. | 13 demes | 0.42 | 0.48 | 0.09 | 0.09 | 0.19 | **-0.52** | -0.24 | **-0.67** | **-0.66** | -0.28 | -0.10 | -0.12 | -0.35 | **-0.61** | **-0.60** | -0.25 |
| | | | | 18 demes | 0.47 | **0.53** | 0.09 | 0.11 | 0.31 | -0.49 | -0.25 | **-0.72** | **-0.67** | -0.30 | -0.07 | -0.12 | -0.43 | **-0.66** | **-0.62** | -0.27 |
| | | | prox. non-rot. | 13 demes | 0.45 | **0.52** | 0.08 | 0.09 | 0.28 | -0.48 | -0.21 | **-0.68** | **-0.66** | -0.29 | -0.10 | -0.11 | -0.31 | **-0.59** | **-0.58** | -0.23 |
| | | | | 18 demes | 0.45 | **0.50** | 0.12 | 0.14 | 0.26 | -0.46 | -0.19 | **-0.69** | **-0.64** | -0.25 | -0.08 | -0.12 | -0.37 | **-0.62** | **-0.58** | -0.24 |
| | 3-way | 10-5 to 10-4 | dist. rot. | 13 demes | 0.09 | 0.13 | -0.09 | -0.09 | -0.02 | -0.05 | -0.05 | -0.14 | -0.14 | -0.09 | 0.18 | 0.18 | 0.26 | -0.27 | -0.26 | -0.18 |
| | | | | 18 demes | 0.16 | 0.24 | -0.09 | -0.09 | -0.04 | -0.10 | -0.08 | -0.18 | -0.18 | -0.15 | 0.04 | 0.07 | 0.21 | -0.28 | -0.28 | -0.15 |
| | | | prox. rot. | 13 demes | 0.07 | 0.11 | -0.07 | -0.07 | -0.01 | -0.04 | -0.04 | -0.11 | -0.12 | -0.07 | 0.08 | 0.08 | 0.24 | -0.21 | -0.22 | -0.11 |
| | | | | 18 demes | 0.08 | 0.13 | -0.08 | -0.08 | -0.03 | -0.09 | -0.04 | -0.10 | -0.10 | -0.07 | -0.03 | 0.00 | 0.19 | -0.23 | -0.23 | -0.13 |
| | | | dist. non-rot. | 13 demes | 0.13 | 0.16 | -0.04 | -0.04 | -0.03 | -0.10 | -0.05 | -0.11 | -0.11 | -0.08 | -0.07 | -0.06 | 0.11 | -0.29 | -0.30 | -0.17 |
| | | | | 18 demes | 0.16 | 0.25 | -0.07 | -0.08 | -0.09 | -0.11 | -0.09 | -0.08 | -0.08 | -0.10 | -0.07 | -0.05 | 0.17 | -0.29 | -0.30 | -0.15 |
| | | | prox. non-rot. | 13 demes | 0.10 | 0.12 | -0.04 | -0.04 | -0.01 | -0.04 | -0.03 | -0.06 | -0.06 | -0.03 | -0.03 | -0.02 | 0.11 | -0.15 | -0.15 | -0.07 |
| | | | | 18 demes | 0.10 | 0.19 | -0.07 | -0.07 | -0.01 | -0.08 | -0.03 | -0.14 | -0.17 | -0.05 | -0.04 | -0.02 | 0.18 | -0.21 | -0.23 | -0.10 |
| | | 10-4 to 10-3 | dist. rot. | 13 demes | 0.33 | 0.42 | 0.02 | 0.05 | 0.08 | -0.19 | -0.16 | -0.35 | -0.28 | -0.30 | -0.01 | -0.05 | -0.12 | -0.35 | -0.28 | -0.21 |
| | | | | 18 demes | 0.44 | 0.46 | 0.05 | 0.11 | 0.17 | -0.17 | -0.10 | -0.44 | -0.24 | -0.19 | -0.05 | -0.16 | -0.33 | -0.42 | -0.26 | -0.16 |
| | | | prox. rot. | 13 demes | 0.36 | 0.42 | 0.00 | 0.03 | 0.08 | -0.21 | -0.17 | -0.36 | -0.26 | -0.23 | -0.01 | -0.03 | -0.13 | -0.38 | -0.29 | -0.22 |
| | | | | 18 demes | 0.43 | 0.46 | 0.04 | 0.11 | 0.10 | -0.16 | -0.05 | -0.44 | -0.25 | -0.10 | -0.05 | -0.16 | -0.29 | -0.38 | -0.21 | -0.14 |
| | | | dist. non-rot. | 13 demes | 0.37 | 0.40 | -0.01 | 0.01 | 0.00 | -0.24 | -0.21 | -0.37 | -0.32 | -0.23 | -0.03 | -0.07 | -0.14 | -0.37 | -0.30 | -0.10 |
| | | | | 18 demes | 0.41 | 0.35 | 0.02 | 0.10 | 0.16 | -0.19 | -0.24 | -0.38 | -0.16 | -0.28 | -0.01 | -0.11 | -0.25 | -0.38 | -0.21 | -0.16 |
| | | | prox. non-rot. | 13 demes | 0.33 | 0.37 | 0.05 | 0.09 | 0.02 | -0.19 | -0.11 | -0.32 | -0.26 | -0.26 | -0.04 | -0.07 | -0.15 | -0.31 | -0.25 | -0.19 |
| | | | | 18 demes | 0.41 | 0.48 | 0.05 | 0.12 | 0.09 | -0.15 | -0.09 | -0.39 | -0.19 | -0.24 | -0.04 | -0.15 | -0.30 | -0.38 | -0.20 | -0.13 |
| | | 10-5 to 10-2 | dist. rot. | 13 demes | 0.44 | 0.47 | 0.15 | 0.16 | 0.17 | -0.28 | -0.14 | -0.45 | -0.37 | -0.20 | -0.11 | -0.14 | -0.27 | -0.39 | -0.32 | -0.14 |
| | | | | 18 demes | 0.47 | 0.49 | 0.16 | 0.18 | 0.12 | -0.25 | -0.13 | **-0.54** | -0.40 | -0.13 | -0.12 | -0.22 | -0.36 | -0.44 | -0.30 | -0.13 |
| | | | | systematic | 0.28 | **0.75** | -0.34 | -0.36 | 0.22 | -0.35 | 0.19 | **-0.72** | **-0.64** | -0.24 | | | | | | |
| | | | prox. rot. | 13 demes | 0.41 | 0.46 | 0.13 | 0.14 | 0.11 | -0.26 | -0.11 | -0.44 | -0.38 | -0.18 | -0.13 | -0.16 | -0.29 | -0.35 | -0.29 | -0.12 |
| | | | | 18 demes | 0.46 | 0.47 | 0.16 | 0.19 | 0.13 | -0.26 | -0.13 | **-0.51** | -0.38 | -0.18 | -0.14 | -0.23 | -0.38 | -0.44 | -0.32 | -0.13 |
| | | | dist. non-rot. | 13 demes | 0.46 | 0.48 | 0.13 | 0.14 | 0.13 | -0.32 | -0.16 | **-0.50** | -0.43 | -0.26 | -0.12 | -0.15 | -0.29 | -0.43 | -0.35 | -0.13 |
| | | | | 18 demes | **0.51** | **0.50** | 0.14 | 0.16 | 0.17 | -0.31 | -0.19 | **-0.58** | -0.45 | -0.21 | -0.14 | -0.22 | -0.42 | -0.49 | -0.36 | -0.10 |
| | | | | systematic | -0.02 | 0.42 | -0.43 | -0.49 | 0.20 | **-0.64** | -0.16 | **-0.75** | **-0.74** | -0.27 | | | | | | |
| | | | prox. non-rot. | 13 demes | 0.43 | 0.42 | 0.14 | 0.14 | 0.17 | -0.31 | -0.17 | **-0.51** | -0.44 | -0.29 | -0.14 | -0.17 | -0.29 | -0.36 | -0.29 | -0.11 |
| | | | | 18 demes | 0.50 | **0.50** | 0.13 | 0.17 | 0.20 | -0.24 | -0.11 | **-0.57** | -0.41 | -0.20 | -0.12 | -0.21 | -0.36 | -0.44 | -0.27 | -0.19 |
| | | 10-3 to 10-2 | dist. rot. | 13 demes | 0.42 | 0.45 | 0.26 | 0.26 | 0.26 | -0.31 | -0.18 | -0.36 | -0.36 | -0.22 | -0.11 | -0.11 | -0.15 | -0.33 | -0.33 | -0.19 |
| | | | | 18 demes | 0.44 | 0.48 | 0.29 | 0.28 | 0.29 | -0.36 | -0.18 | -0.45 | -0.45 | -0.26 | -0.11 | -0.11 | -0.21 | -0.40 | -0.40 | -0.18 |
| | | | prox. rot. | 13 demes | 0.41 | 0.44 | 0.27 | 0.27 | 0.27 | -0.28 | -0.17 | -0.36 | -0.36 | -0.23 | -0.10 | -0.10 | -0.13 | -0.28 | -0.28 | -0.14 |
| | | | | 18 demes | 0.42 | 0.45 | 0.25 | 0.25 | 0.27 | -0.34 | -0.17 | -0.43 | -0.43 | -0.26 | -0.09 | -0.09 | -0.16 | -0.34 | -0.34 | -0.13 |
| | | | dist. non-rot. | 13 demes | 0.43 | 0.45 | 0.26 | 0.26 | 0.24 | -0.35 | -0.26 | -0.36 | -0.36 | -0.29 | -0.11 | -0.11 | -0.13 | -0.36 | -0.36 | -0.25 |
| | | | | 18 demes | 0.46 | 0.49 | 0.29 | 0.29 | 0.27 | -0.40 | -0.27 | -0.50 | -0.50 | -0.41 | -0.09 | -0.09 | -0.15 | -0.46 | -0.46 | -0.30 |
| | | | prox. non-rot. | 13 demes | 0.44 | 0.45 | 0.25 | 0.25 | 0.25 | -0.30 | -0.23 | -0.35 | -0.35 | -0.31 | -0.09 | -0.09 | -0.11 | -0.31 | -0.31 | -0.23 |
| | | | | 18 demes | 0.42 | 0.45 | 0.25 | 0.25 | 0.24 | -0.37 | -0.26 | -0.43 | -0.43 | -0.37 | -0.06 | -0.06 | -0.11 | -0.44 | -0.44 | -0.29 |
| | 4-way | 10-5 to 10-4 | dist. rot. | 13 demes | 0.10 | 0.11 | -0.05 | -0.05 | -0.01 | -0.03 | -0.04 | -0.08 | -0.08 | -0.07 | 0.15 | 0.15 | 0.16 | -0.30 | -0.30 | -0.25 |
| | | | | 18 demes | 0.17 | 0.20 | -0.10 | -0.11 | -0.08 | -0.08 | -0.06 | -0.18 | -0.18 | -0.17 | 0.10 | 0.12 | 0.12 | -0.22 | -0.21 | -0.18 |
| | | | prox. rot. | 13 demes | 0.07 | 0.09 | -0.04 | -0.04 | -0.01 | -0.03 | -0.03 | -0.11 | -0.11 | -0.08 | 0.20 | 0.20 | 0.22 | -0.30 | -0.30 | -0.24 |
| | | | | 18 demes | 0.09 | 0.11 | -0.09 | -0.09 | -0.05 | -0.07 | -0.04 | -0.08 | -0.09 | -0.07 | 0.05 | 0.08 | 0.17 | -0.29 | -0.28 | -0.25 |
| | | | dist. non-rot. | 13 demes | 0.13 | 0.15 | -0.02 | -0.02 | -0.02 | -0.09 | -0.06 | -0.08 | -0.08 | -0.04 | 0.01 | 0.01 | 0.08 | -0.33 | -0.32 | -0.28 |
| | | | | 18 demes | 0.16 | 0.22 | -0.09 | -0.10 | -0.13 | -0.09 | -0.07 | -0.05 | -0.04 | -0.13 | 0.00 | 0.03 | 0.15 | -0.31 | -0.31 | -0.23 |
| | | | prox. non-rot. | 13 demes | 0.10 | 0.10 | -0.03 | -0.03 | -0.01 | -0.04 | -0.03 | -0.05 | -0.05 | -0.04 | 0.04 | 0.05 | 0.17 | -0.21 | -0.22 | -0.19 |
| | | | | 18 demes | 0.10 | 0.16 | -0.12 | -0.12 | -0.08 | -0.05 | -0.02 | -0.12 | -0.13 | -0.06 | 0.00 | 0.04 | 0.17 | -0.24 | -0.24 | -0.13 |
| | | 10-4 to 10-3 | dist. rot. | 13 demes | 0.30 | 0.36 | 0.01 | 0.01 | 0.06 | -0.16 | -0.12 | -0.27 | -0.26 | -0.26 | -0.05 | -0.06 | -0.08 | -0.26 | -0.24 | -0.16 |
| | | | | 18 demes | 0.40 | 0.45 | 0.05 | 0.07 | 0.14 | -0.16 | -0.12 | -0.32 | -0.22 | -0.18 | -0.09 | -0.13 | -0.22 | -0.32 | -0.25 | -0.17 |
| | | | prox. rot. | 13 demes | 0.32 | 0.38 | 0.01 | 0.02 | 0.08 | -0.18 | -0.15 | -0.31 | -0.28 | -0.22 | -0.05 | -0.05 | -0.05 | -0.29 | -0.27 | -0.18 |
| | | | | 18 demes | 0.39 | 0.43 | 0.04 | 0.06 | 0.10 | -0.15 | -0.08 | -0.32 | -0.23 | -0.11 | -0.09 | -0.13 | -0.19 | -0.25 | -0.18 | -0.11 |
| | | | dist. non-rot. | 13 demes | 0.35 | 0.40 | 0.01 | 0.02 | 0.06 | -0.21 | -0.17 | -0.33 | -0.31 | -0.21 | -0.06 | -0.07 | -0.08 | -0.28 | -0.25 | -0.08 |
| | | | | 18 demes | 0.39 | 0.39 | 0.02 | 0.08 | 0.12 | -0.17 | -0.22 | -0.29 | -0.14 | -0.22 | -0.04 | -0.09 | -0.16 | -0.27 | -0.19 | -0.16 |
| | | | prox. non-rot. | 13 demes | 0.32 | 0.35 | 0.07 | 0.08 | 0.13 | -0.17 | -0.11 | -0.27 | -0.25 | -0.27 | -0.07 | -0.08 | -0.12 | -0.23 | -0.22 | -0.15 |
| | | | | 18 demes | 0.38 | 0.44 | 0.03 | 0.07 | 0.11 | -0.14 | -0.12 | -0.28 | -0.17 | -0.21 | -0.07 | -0.11 | -0.20 | -0.27 | -0.18 | -0.16 |
| | | 10-5 to 10-2 | dist. rot. | 13 demes | 0.40 | 0.43 | 0.11 | 0.11 | 0.10 | -0.23 | -0.13 | -0.28 | -0.28 | -0.17 | -0.12 | -0.12 | -0.16 | -0.21 | -0.20 | -0.10 |
| | | | | 18 demes | 0.44 | 0.46 | 0.13 | 0.14 | 0.11 | -0.24 | -0.13 | -0.36 | -0.31 | -0.10 | -0.16 | -0.17 | -0.22 | -0.28 | -0.25 | -0.11 |
| | | | | systematic | 0.34 | **0.63** | -0.40 | -0.40 | -0.05 | -0.29 | 0.15 | **-0.68** | **-0.65** | -0.31 | | | | | | |
| | | | prox. rot. | 13 demes | 0.37 | 0.40 | 0.10 | 0.10 | 0.07 | -0.21 | -0.11 | -0.29 | -0.29 | -0.17 | -0.14 | -0.14 | -0.18 | -0.19 | -0.18 | -0.07 |
| | | | | 18 demes | 0.42 | 0.45 | 0.10 | 0.11 | 0.10 | -0.25 | -0.13 | -0.37 | -0.33 | -0.16 | -0.16 | -0.17 | -0.25 | -0.28 | -0.25 | -0.09 |
| | | | dist. non-rot. | 13 demes | 0.45 | 0.47 | 0.11 | 0.11 | 0.12 | -0.28 | -0.15 | -0.33 | -0.32 | -0.15 | -0.14 | -0.15 | -0.20 | -0.24 | -0.22 | -0.08 |
| | | | | 18 demes | 0.49 | **0.51** | 0.14 | 0.15 | 0.16 | -0.30 | -0.15 | -0.46 | -0.42 | -0.19 | -0.16 | -0.18 | -0.28 | -0.34 | -0.30 | -0.06 |
| | | | | systematic | 0.06 | 0.35 | -0.44 | -0.45 | 0.06 | **-0.67** | -0.24 | **-0.77** | **-0.77** | -0.39 | | | | | | |
| | | | prox. non-rot. | 13 demes | 0.40 | 0.42 | 0.11 | 0.11 | 0.13 | -0.27 | -0.18 | -0.42 | -0.41 | -0.31 | -0.13 | -0.13 | -0.18 | -0.20 | -0.19 | -0.09 |
| | | | | 18 demes | 0.46 | 0.49 | 0.11 | 0.12 | 0.10 | -0.24 | -0.13 | -0.44 | -0.40 | -0.22 | -0.15 | -0.16 | -0.22 | -0.29 | -0.25 | -0.17 |
| | | 10-3 to 10-2 | dist. rot. | 13 demes | 0.38 | 0.38 | 0.21 | 0.21 | 0.21 | -0.18 | -0.16 | -0.20 | -0.20 | -0.18 | -0.07 | -0.07 | -0.08 | -0.10 | -0.10 | -0.09 |
| | | | | 18 demes | 0.42 | 0.42 | 0.21 | 0.21 | 0.21 | -0.25 | -0.19 | -0.32 | -0.32 | -0.25 | -0.09 | -0.09 | -0.10 | -0.18 | -0.18 | -0.12 |
| | | | prox. rot. | 13 demes | 0.38 | 0.38 | 0.21 | 0.21 | 0.21 | -0.15 | -0.13 | -0.15 | -0.15 | -0.14 | 0.03 | 0.03 | 0.04 | 0.00 | 0.00 | 0.05 |
| | | | | 18 demes | 0.39 | 0.40 | 0.23 | 0.23 | 0.23 | -0.23 | -0.18 | -0.33 | -0.33 | -0.27 | -0.06 | -0.06 | -0.07 | -0.16 | -0.16 | -0.12 |
| | | | dist. non-rot. | 13 demes | 0.39 | 0.39 | 0.22 | 0.22 | 0.22 | -0.21 | -0.21 | -0.25 | -0.25 | -0.25 | -0.16 | -0.16 | -0.16 | -0.11 | -0.11 | -0.09 |
| | | | | 18 demes | 0.43 | 0.44 | 0.21 | 0.21 | 0.21 | -0.27 | -0.24 | -0.37 | -0.37 | -0.36 | -0.08 | -0.08 | -0.08 | -0.20 | -0.20 | -0.18 |
| | | | prox. non-rot. | 13 demes | 0.39 | 0.39 | 0.21 | 0.21 | 0.21 | -0.19 | -0.18 | -0.29 | -0.29 | -0.29 | -0.11 | -0.11 | -0.11 | -0.12 | -0.12 | -0.12 |
| | | | | 18 demes | 0.40 | 0.41 | 0.21 | 0.21 | 0.21 | -0.25 | -0.24 | -0.37 | -0.37 | -0.37 | -0.07 | -0.07 | -0.07 | -0.22 | -0.22 | -0.20 |

*p*-value > 0.05/7980 (Bonferroni correction)

**|Spearman correlation coefficient| >= 0.5**

| **part 4** | | | | **variable 2:** | **max. \|estimated admixture fractions - equal fractions\|** | | | | | ***p*-value** | | | | | **max. SE** | | | ***p*-value** | | |
|---|---|---|---|---|---|---|---|---|---|---|---|---|---|---|---|---|---|---|---|---|
| **total number of randomized models:** | | | | | 21,663,216 | 12,439,194 | 2,392,357 | 2,040,355 | 1,058,613 | 21,663,216 | 12,439,194 | 2,392,357 | 2,040,355 | 1,058,613 | 4,378,913 | 3,566,530 | 1,533,911 | 4,378,913 | 3,566,530 | 1,533,911 |
| **total number of systematic models (10-5 to 10-2):** | | | | | 3,576,667 | 1,158,133 | 1,267,603 | 1,064,846 | 240,961 | 3,576,667 | 1,158,133 | 1,267,603 | 1,064,846 | 240,961 | N/A | | | N/A | | |
| **variable 1:** | **model complexity** | **gene-flow intensity per gen.** | ***qpAdm* protocol** | **sampling density** | p >= 1e-55 | p >= 1e-5 | max. \|*EAF–EF*\| <= 0.5 | p >= 1e-55; max. \|*EAF–EF*\| <= 0.5 | p >= 1e-5; max. \|*EAF–EF*\| <= 0.5 | p >= 1e-55 | p >= 1e-5 | max. \|*EAF–EF*\| <= 0.5 | p >= 1e-55; max. \|*EAF–EF*\| <= 0.5 | p >= 1e-5; max. \|*EAF–EF*\| <= 0.5 | max. SE <= 0.1 | p >= 1e-55; max. SE <= 0.1 | p >= 1e-5; max. SE <= 0.1 | max. SE <= 0.1 | p >= 1e-55; max. SE <= 0.1 | p >= 1e-5; max. SE <= 0.1 |
| **distance to the ideal symmetric model (in the space of max. distance vs. min. angle)** | **2-way** | **10-5 to 10-4** | dist. rot. | 13 demes | 0.07 | 0.13 | -0.03 | -0.03 | 0.01 | 0.03 | 0.00 | -0.02 | -0.01 | -0.01 | 0.05 | 0.05 | 0.03 | -0.01 | -0.02 | 0.00 |
| | | | | 18 demes | 0.13 | 0.26 | 0.02 | 0.01 | 0.02 | 0.01 | 0.02 | -0.05 | -0.06 | -0.05 | -0.02 | -0.02 | 0.04 | -0.08 | -0.10 | -0.02 |
| | | | prox. rot. | 13 demes | 0.08 | 0.14 | 0.03 | 0.03 | 0.01 | 0.00 | -0.02 | -0.04 | -0.04 | -0.05 | -0.04 | -0.04 | -0.03 | -0.05 | -0.07 | -0.05 |
| | | | | 18 demes | 0.09 | 0.19 | 0.02 | 0.02 | 0.01 | 0.02 | 0.02 | -0.03 | -0.02 | 0.00 | -0.02 | -0.02 | 0.02 | -0.06 | -0.06 | -0.02 |
| | | | dist. non-rot. | 13 demes | 0.09 | 0.13 | 0.04 | 0.04 | 0.03 | -0.02 | 0.02 | -0.03 | -0.02 | 0.01 | -0.04 | -0.04 | -0.04 | -0.09 | -0.11 | -0.01 |
| | | | | 18 demes | 0.11 | 0.22 | 0.03 | 0.03 | -0.01 | 0.01 | -0.01 | -0.02 | -0.03 | -0.03 | 0.01 | 0.01 | 0.03 | -0.04 | -0.07 | -0.02 |
| | | | prox. non-rot. | 13 demes | 0.10 | 0.12 | 0.06 | 0.06 | 0.06 | 0.05 | 0.05 | 0.01 | 0.03 | 0.03 | -0.02 | -0.02 | -0.06 | -0.01 | -0.01 | 0.05 |
| | | | | 18 demes | 0.11 | 0.25 | 0.04 | 0.04 | 0.09 | 0.01 | 0.07 | -0.05 | -0.07 | -0.03 | 0.02 | 0.03 | 0.03 | -0.03 | -0.07 | 0.00 |
| | | **10-4 to 10-3** | dist. rot. | 13 demes | **0.53** | **0.69** | 0.20 | 0.28 | 0.42 | 0.03 | 0.09 | -0.17 | -0.10 | -0.15 | 0.03 | 0.04 | 0.01 | -0.13 | -0.05 | -0.12 |
| | | | | 18 demes | **0.65** | **0.72** | 0.24 | 0.40 | **0.56** | 0.05 | 0.37 | -0.23 | -0.06 | -0.13 | 0.01 | 0.01 | -0.03 | -0.19 | -0.06 | -0.02 |
| | | | prox. rot. | 13 demes | **0.56** | **0.69** | 0.14 | 0.21 | 0.32 | 0.03 | 0.19 | -0.17 | -0.09 | -0.18 | 0.06 | 0.08 | 0.04 | -0.14 | -0.06 | -0.03 |
| | | | | 18 demes | **0.65** | **0.78** | 0.22 | 0.39 | 0.48 | 0.07 | 0.40 | -0.21 | -0.05 | -0.29 | 0.04 | 0.00 | 0.04 | -0.17 | -0.02 | -0.11 |
| | | | dist. non-rot. | 13 demes | 0.50 | **0.60** | 0.17 | 0.24 | 0.46 | -0.03 | -0.03 | -0.12 | -0.09 | 0.10 | 0.05 | 0.03 | -0.03 | -0.13 | -0.05 | 0.05 |
| | | | | 18 demes | **0.58** | **0.66** | 0.17 | 0.33 | 0.46 | -0.01 | -0.08 | -0.17 | -0.03 | -0.10 | 0.04 | 0.01 | -0.15 | -0.14 | -0.01 | -0.05 |
| | | | prox. non-rot. | 13 demes | 0.50 | **0.64** | 0.19 | 0.27 | 0.38 | 0.00 | 0.19 | -0.14 | -0.13 | -0.10 | 0.04 | 0.04 | 0.00 | -0.15 | -0.07 | 0.05 |
| | | | | 18 demes | **0.62** | **0.72** | 0.21 | 0.38 | **0.51** | 0.09 | 0.34 | -0.17 | 0.00 | 0.01 | 0.02 | -0.02 | -0.26 | -0.15 | 0.01 | 0.04 |
| | | **10-5 to 10-2** | dist. rot. | 13 demes | **0.69** | **0.72** | 0.30 | 0.40 | 0.46 | -0.02 | 0.03 | -0.29 | -0.05 | 0.00 | 0.14 | 0.08 | -0.06 | -0.15 | -0.07 | -0.02 |
| | | | | 18 demes | **0.74** | **0.75** | 0.35 | 0.50 | 0.50 | 0.00 | 0.05 | -0.30 | -0.04 | -0.02 | 0.16 | 0.06 | -0.11 | -0.13 | -0.05 | -0.02 |
| | | | | systematic | **0.82** | **0.62** | 0.26 | 0.30 | 0.27 | 0.19 | 0.05 | -0.17 | -0.11 | -0.12 | | | | | | |
| | | | prox. rot. | 13 demes | **0.69** | **0.72** | 0.28 | 0.38 | 0.45 | 0.00 | 0.07 | -0.30 | -0.08 | 0.00 | 0.13 | 0.09 | -0.08 | -0.15 | -0.05 | -0.01 |
| | | | | 18 demes | **0.73** | **0.74** | 0.36 | **0.50** | **0.55** | 0.00 | 0.08 | -0.33 | -0.04 | 0.02 | 0.14 | 0.05 | -0.11 | -0.15 | -0.05 | 0.00 |
| | | | dist. non-rot. | 13 demes | **0.70** | **0.69** | 0.35 | 0.44 | **0.64** | -0.03 | 0.06 | -0.30 | -0.05 | 0.09 | 0.14 | 0.09 | -0.09 | -0.19 | -0.07 | 0.10 |
| | | | | 18 demes | **0.75** | **0.78** | 0.36 | **0.53** | **0.57** | -0.05 | -0.08 | -0.35 | -0.05 | -0.07 | 0.14 | 0.05 | -0.16 | -0.20 | -0.08 | -0.09 |
| | | | | systematic | **0.82** | **0.84** | 0.29 | 0.37 | 0.40 | 0.06 | -0.05 | -0.35 | -0.16 | 0.00 | | | | | | |
| | | | prox. non-rot. | 13 demes | **0.70** | **0.68** | 0.28 | 0.37 | **0.57** | -0.02 | 0.05 | -0.34 | -0.08 | -0.02 | 0.13 | 0.07 | -0.11 | -0.17 | -0.04 | -0.02 |
| | | | | 18 demes | **0.75** | **0.75** | 0.32 | 0.48 | **0.54** | -0.01 | 0.06 | -0.44 | -0.10 | -0.17 | 0.15 | 0.06 | -0.12 | -0.18 | -0.05 | -0.16 |
| | | **10-3 to 10-2** | dist. rot. | 13 demes | **0.72** | **0.76** | 0.42 | 0.44 | **0.57** | -0.04 | 0.00 | -0.44 | -0.40 | -0.10 | 0.16 | 0.15 | 0.02 | -0.21 | -0.19 | -0.06 |
| | | | | 18 demes | **0.74** | **0.77** | 0.45 | **0.52** | **0.66** | -0.02 | 0.00 | -0.45 | -0.34 | -0.08 | 0.20 | 0.18 | 0.03 | -0.20 | -0.15 | -0.05 |
| | | | prox. rot. | 13 demes | **0.69** | **0.73** | 0.35 | 0.38 | **0.50** | -0.01 | 0.01 | -0.43 | -0.39 | -0.07 | 0.15 | 0.15 | 0.03 | -0.17 | -0.15 | -0.03 |
| | | | | 18 demes | **0.73** | **0.76** | 0.39 | 0.45 | **0.58** | -0.02 | 0.01 | **-0.50** | -0.40 | -0.05 | 0.20 | 0.19 | 0.05 | -0.19 | -0.14 | -0.01 |
| | | | dist. non-rot. | 13 demes | **0.72** | **0.77** | 0.45 | 0.46 | **0.60** | -0.06 | -0.02 | -0.49 | -0.47 | -0.17 | 0.19 | 0.18 | 0.10 | -0.23 | -0.22 | -0.05 |
| | | | | 18 demes | **0.73** | **0.78** | 0.47 | **0.51** | **0.69** | -0.04 | -0.03 | -0.49 | -0.43 | -0.14 | 0.20 | 0.19 | 0.07 | -0.21 | -0.19 | -0.06 |
| | | | prox. non-rot. | 13 demes | **0.70** | **0.76** | 0.37 | 0.38 | **0.59** | -0.02 | 0.02 | -0.50 | -0.48 | -0.19 | 0.16 | 0.16 | 0.11 | -0.18 | -0.18 | -0.05 |
| | | | | 18 demes | **0.73** | **0.76** | 0.43 | 0.46 | **0.62** | 0.00 | 0.01 | **-0.52** | -0.47 | -0.15 | 0.20 | 0.20 | 0.12 | -0.18 | -0.16 | -0.05 |
| | **3-way** | **10-5 to 10-4** | dist. rot. | 13 demes | 0.03 | 0.05 | -0.04 | -0.04 | -0.03 | 0.06 | 0.03 | -0.02 | -0.02 | -0.01 | 0.04 | 0.04 | 0.03 | -0.02 | -0.02 | -0.01 |
| | | | | 18 demes | 0.11 | 0.16 | -0.03 | -0.03 | 0.00 | 0.05 | 0.04 | -0.02 | -0.02 | -0.04 | 0.00 | 0.00 | 0.02 | -0.02 | -0.02 | 0.00 |
| | | | prox. rot. | 13 demes | 0.05 | 0.07 | 0.03 | 0.03 | 0.05 | 0.04 | 0.01 | 0.01 | 0.01 | -0.01 | 0.01 | 0.01 | 0.05 | 0.00 | 0.00 | 0.00 |
| | | | | 18 demes | 0.06 | 0.10 | -0.03 | -0.03 | -0.01 | 0.06 | 0.05 | 0.02 | 0.02 | -0.02 | -0.03 | -0.03 | -0.03 | -0.02 | -0.02 | -0.02 |
| | | | dist. non-rot. | 13 demes | 0.06 | 0.07 | 0.04 | 0.04 | 0.04 | 0.00 | 0.01 | -0.01 | -0.01 | -0.03 | -0.07 | -0.07 | -0.02 | -0.09 | -0.10 | -0.03 |
| | | | | 18 demes | 0.08 | 0.12 | -0.02 | -0.02 | -0.01 | 0.03 | 0.02 | 0.01 | 0.01 | -0.06 | -0.02 | -0.02 | 0.00 | -0.02 | -0.03 | 0.00 |
| | | | prox. non-rot. | 13 demes | 0.06 | 0.07 | 0.02 | 0.02 | 0.02 | 0.08 | 0.06 | 0.05 | 0.06 | 0.06 | -0.05 | -0.05 | -0.05 | 0.01 | 0.01 | 0.07 |
| | | | | 18 demes | 0.07 | 0.17 | -0.02 | -0.02 | 0.04 | 0.06 | 0.09 | -0.07 | -0.09 | -0.01 | 0.01 | 0.01 | 0.02 | -0.02 | -0.04 | 0.03 |
| | | **10-4 to 10-3** | dist. rot. | 13 demes | 0.33 | 0.42 | 0.14 | 0.17 | 0.19 | 0.05 | 0.07 | -0.07 | -0.02 | -0.13 | 0.03 | 0.03 | -0.03 | -0.03 | 0.00 | -0.05 |
| | | | | 18 demes | 0.43 | **0.55** | 0.13 | 0.15 | 0.21 | 0.08 | 0.24 | -0.10 | -0.01 | -0.02 | -0.01 | -0.01 | -0.02 | -0.06 | 0.00 | 0.02 |
| | | | prox. rot. | 13 demes | 0.35 | 0.45 | 0.11 | 0.12 | 0.15 | 0.05 | 0.11 | -0.05 | -0.04 | -0.12 | 0.04 | 0.03 | 0.02 | -0.04 | -0.02 | 0.00 |
| | | | | 18 demes | 0.45 | **0.57** | 0.12 | 0.15 | 0.09 | 0.08 | 0.23 | -0.05 | -0.03 | -0.01 | 0.01 | -0.01 | 0.06 | -0.02 | 0.01 | -0.04 |
| | | | dist. non-rot. | 13 demes | 0.30 | 0.35 | 0.08 | 0.09 | 0.13 | 0.01 | -0.01 | -0.03 | -0.06 | -0.06 | -0.01 | -0.03 | -0.06 | 0.01 | 0.02 | 0.09 |
| | | | | 18 demes | 0.36 | 0.40 | 0.07 | 0.12 | 0.16 | 0.01 | -0.03 | -0.05 | -0.03 | -0.14 | 0.00 | -0.02 | -0.08 | -0.01 | 0.03 | 0.01 |
| | | | prox. non-rot. | 13 demes | 0.33 | 0.41 | 0.11 | 0.14 | 0.12 | 0.03 | 0.12 | -0.03 | -0.06 | -0.12 | 0.03 | 0.02 | -0.02 | -0.03 | -0.02 | 0.04 |
| | | | | 18 demes | 0.43 | **0.53** | 0.14 | 0.19 | 0.18 | 0.10 | 0.21 | -0.04 | 0.01 | -0.10 | 0.00 | -0.01 | -0.13 | -0.03 | 0.06 | 0.02 |
| | | **10-5 to 10-2** | dist. rot. | 13 demes | 0.45 | 0.48 | 0.32 | 0.31 | 0.26 | 0.01 | 0.02 | 0.03 | 0.07 | 0.02 | 0.11 | 0.11 | 0.03 | -0.02 | 0.00 | 0.00 |
| | | | | 18 demes | 0.48 | **0.51** | 0.36 | 0.33 | 0.25 | 0.04 | 0.05 | 0.04 | 0.07 | 0.06 | 0.13 | 0.11 | 0.02 | 0.02 | 0.03 | 0.03 |
| | | | | systematic | **0.60** | 0.27 | 0.29 | 0.30 | 0.39 | 0.38 | 0.09 | 0.11 | 0.06 | 0.01 | | | | | | |
| | | | prox. rot. | 13 demes | 0.43 | 0.47 | 0.28 | 0.27 | 0.20 | 0.03 | 0.04 | 0.04 | 0.06 | 0.03 | 0.08 | 0.07 | -0.01 | 0.01 | 0.02 | 0.01 |
| | | | | 18 demes | 0.48 | **0.51** | 0.36 | 0.32 | 0.20 | 0.03 | 0.05 | 0.06 | 0.07 | 0.05 | 0.12 | 0.11 | 0.00 | 0.01 | 0.00 | 0.01 |
| | | | dist. non-rot. | 13 demes | 0.46 | 0.45 | 0.35 | 0.35 | 0.25 | -0.02 | 0.03 | 0.00 | 0.04 | 0.03 | 0.15 | 0.15 | 0.10 | -0.02 | 0.02 | 0.10 |
| | | | | 18 demes | 0.50 | **0.53** | 0.35 | 0.34 | 0.28 | -0.03 | -0.04 | -0.05 | 0.01 | 0.00 | 0.13 | 0.12 | 0.04 | -0.05 | -0.02 | 0.00 |
| | | | | systematic | **0.70** | **0.63** | 0.34 | 0.37 | 0.36 | 0.19 | -0.01 | -0.15 | -0.07 | 0.01 | | | | | | |
| | | | prox. non-rot. | 13 demes | 0.45 | 0.45 | 0.32 | 0.32 | 0.28 | 0.01 | 0.01 | -0.02 | 0.02 | -0.08 | 0.08 | 0.08 | 0.01 | 0.02 | 0.04 | 0.02 |
| | | | | 18 demes | **0.50** | **0.53** | 0.35 | 0.35 | 0.29 | 0.03 | 0.03 | -0.05 | 0.04 | 0.02 | 0.13 | 0.12 | 0.03 | -0.02 | 0.02 | -0.10 |
| | | **10-3 to 10-2** | dist. rot. | 13 demes | 0.46 | 0.48 | 0.43 | 0.43 | 0.42 | -0.01 | -0.01 | 0.01 | 0.01 | 0.04 | 0.13 | 0.13 | 0.12 | -0.07 | -0.07 | -0.03 |
| | | | | 18 demes | 0.48 | 0.50 | 0.47 | 0.47 | 0.45 | 0.01 | 0.01 | 0.00 | 0.00 | 0.05 | 0.17 | 0.17 | 0.14 | -0.06 | -0.06 | 0.00 |
| | | | prox. rot. | 13 demes | 0.44 | 0.46 | 0.40 | 0.40 | 0.39 | 0.03 | 0.02 | 0.04 | 0.04 | 0.07 | 0.12 | 0.12 | 0.11 | 0.00 | 0.00 | 0.05 |
| | | | | 18 demes | 0.46 | 0.49 | 0.48 | 0.48 | 0.46 | -0.01 | 0.00 | 0.01 | 0.01 | 0.04 | 0.17 | 0.17 | 0.16 | -0.07 | -0.07 | 0.01 |
| | | | dist. non-rot. | 13 demes | 0.46 | 0.48 | 0.44 | 0.44 | 0.44 | -0.01 | -0.01 | 0.01 | 0.01 | 0.02 | 0.17 | 0.17 | 0.17 | -0.11 | -0.11 | -0.06 |
| | | | | 18 demes | 0.47 | 0.49 | 0.49 | 0.49 | 0.49 | -0.02 | -0.01 | -0.03 | -0.03 | 0.00 | 0.20 | 0.20 | 0.21 | -0.10 | -0.10 | -0.04 |
| | | | prox. non-rot. | 13 demes | 0.46 | 0.47 | 0.44 | 0.44 | 0.44 | 0.00 | 0.00 | 0.00 | 0.00 | 0.01 | 0.14 | 0.14 | 0.14 | -0.11 | -0.11 | -0.07 |
| | | | | 18 demes | 0.48 | 0.49 | 0.46 | 0.46 | 0.45 | -0.01 | 0.00 | -0.04 | -0.04 | -0.02 | 0.20 | 0.20 | 0.20 | -0.13 | -0.13 | -0.06 |
| | **4-way** | **10-5 to 10-4** | dist. rot. | 13 demes | 0.01 | 0.01 | -0.02 | -0.02 | -0.02 | 0.05 | 0.04 | 0.00 | 0.00 | 0.01 | -0.08 | -0.08 | -0.09 | 0.01 | 0.01 | 0.05 |
| | | | | 18 demes | 0.09 | 0.11 | -0.04 | -0.04 | -0.01 | 0.05 | 0.05 | -0.03 | -0.02 | -0.04 | 0.02 | 0.01 | 0.01 | 0.00 | 0.00 | -0.05 |
| | | | prox. rot. | 13 demes | 0.02 | 0.03 | 0.03 | 0.03 | 0.04 | 0.03 | 0.02 | 0.01 | 0.01 | 0.00 | 0.04 | 0.04 | -0.01 | -0.02 | -0.02 | -0.04 |
| | | | | 18 demes | 0.04 | 0.05 | -0.04 | -0.04 | -0.03 | 0.07 | 0.06 | 0.01 | 0.01 | -0.02 | -0.02 | -0.02 | -0.03 | 0.00 | -0.01 | -0.04 |
| | | | dist. non-rot. | 13 demes | 0.04 | 0.04 | 0.05 | 0.05 | 0.04 | 0.00 | 0.00 | -0.03 | -0.03 | -0.04 | -0.06 | -0.06 | -0.04 | -0.07 | -0.07 | -0.09 |
| | | | | 18 demes | 0.06 | 0.07 | -0.03 | -0.03 | -0.03 | 0.03 | 0.02 | 0.04 | 0.05 | -0.06 | -0.01 | -0.01 | 0.00 | 0.01 | -0.01 | -0.03 |
| | | | prox. non-rot. | 13 demes | 0.05 | 0.04 | 0.02 | 0.02 | 0.03 | 0.07 | 0.05 | 0.06 | 0.06 | 0.05 | -0.06 | -0.06 | -0.04 | 0.00 | 0.00 | 0.01 |
| | | | | 18 demes | 0.06 | 0.12 | -0.05 | -0.05 | 0.00 | 0.08 | 0.09 | -0.07 | -0.07 | 0.00 | 0.00 | -0.01 | 0.01 | -0.03 | -0.04 | -0.01 |
| | | **10-4 to 10-3** | dist. rot. | 13 demes | 0.21 | 0.25 | 0.11 | 0.11 | 0.14 | 0.06 | 0.06 | 0.02 | 0.02 | -0.09 | 0.00 | 0.00 | -0.04 | 0.02 | 0.03 | 0.00 |
| | | | | 18 demes | 0.30 | 0.39 | 0.08 | 0.09 | 0.12 | 0.08 | 0.14 | 0.03 | 0.05 | 0.00 | 0.00 | -0.01 | -0.01 | 0.01 | 0.03 | 0.03 |
| | | | prox. rot. | 13 demes | 0.24 | 0.28 | 0.07 | 0.08 | 0.08 | 0.07 | 0.07 | -0.02 | -0.02 | -0.08 | 0.01 | 0.01 | 0.05 | 0.00 | 0.01 | 0.02 |
| | | | | 18 demes | 0.32 | 0.41 | 0.08 | 0.09 | 0.08 | 0.09 | 0.14 | 0.05 | 0.02 | -0.03 | 0.00 | -0.01 | -0.01 | 0.04 | 0.04 | -0.03 |
| | | | dist. non-rot. | 13 demes | 0.19 | 0.20 | 0.08 | 0.08 | 0.11 | 0.02 | 0.00 | 0.01 | -0.02 | -0.08 | -0.03 | -0.05 | -0.04 | 0.09 | 0.07 | 0.09 |
| | | | | 18 demes | 0.23 | 0.24 | 0.05 | 0.08 | 0.15 | 0.02 | -0.02 | 0.00 | 0.01 | -0.11 | -0.01 | -0.03 | -0.07 | 0.04 | 0.05 | 0.00 |
| | | | prox. non-rot. | 13 demes | 0.25 | 0.27 | 0.13 | 0.14 | 0.22 | 0.05 | 0.07 | 0.02 | 0.00 | -0.14 | 0.00 | -0.01 | -0.04 | 0.05 | 0.03 | -0.02 |
| | | | | 18 demes | 0.31 | 0.39 | 0.09 | 0.10 | 0.10 | 0.09 | 0.12 | 0.04 | 0.02 | -0.04 | -0.01 | -0.01 | -0.07 | 0.05 | 0.08 | 0.02 |
| | | **10-5 to 10-2** | dist. rot. | 13 demes | 0.31 | 0.32 | 0.18 | 0.17 | 0.13 | 0.02 | 0.02 | 0.14 | 0.13 | 0.07 | 0.10 | 0.10 | 0.06 | 0.07 | 0.08 | 0.05 |
| | | | | 18 demes | 0.34 | 0.35 | 0.21 | 0.20 | 0.12 | 0.05 | 0.03 | 0.19 | 0.18 | 0.07 | 0.09 | 0.08 | 0.02 | 0.11 | 0.10 | 0.05 |
| | | | | systematic | 0.30 | 0.06 | 0.25 | 0.25 | 0.32 | 0.44 | 0.09 | 0.26 | 0.23 | 0.13 | | | | | | |
| | | | prox. rot. | 13 demes | 0.29 | 0.30 | 0.14 | 0.14 | 0.11 | 0.03 | 0.03 | 0.12 | 0.12 | 0.04 | 0.04 | 0.03 | 0.00 | 0.07 | 0.07 | 0.04 |
| | | | | 18 demes | 0.34 | 0.35 | 0.19 | 0.16 | 0.09 | 0.05 | 0.03 | 0.18 | 0.16 | 0.05 | 0.10 | 0.09 | 0.04 | 0.08 | 0.07 | 0.06 |
| | | | dist. non-rot. | 13 demes | 0.34 | 0.32 | 0.22 | 0.22 | 0.10 | -0.01 | 0.01 | 0.15 | 0.16 | 0.07 | 0.13 | 0.12 | 0.10 | 0.10 | 0.10 | 0.10 |
| | | | | 18 demes | 0.35 | 0.37 | 0.24 | 0.23 | 0.19 | -0.02 | -0.01 | 0.09 | 0.08 | 0.03 | 0.10 | 0.09 | 0.04 | 0.02 | 0.03 | 0.04 |
| | | | | systematic | 0.50 | 0.39 | 0.34 | 0.35 | 0.30 | 0.25 | 0.04 | 0.05 | 0.05 | 0.10 | | | | | | |
| | | | prox. non-rot. | 13 demes | 0.32 | 0.31 | 0.19 | 0.18 | 0.12 | 0.03 | 0.01 | 0.11 | 0.11 | -0.06 | 0.07 | 0.07 | 0.02 | 0.09 | 0.09 | 0.03 |
| | | | | 18 demes | 0.37 | 0.39 | 0.21 | 0.19 | 0.15 | 0.03 | 0.02 | 0.13 | 0.12 | 0.06 | 0.09 | 0.08 | 0.04 | 0.06 | 0.06 | -0.06 |
| | | **10-3 to 10-2** | dist. rot. | 13 demes | 0.31 | 0.32 | 0.29 | 0.29 | 0.29 | -0.01 | -0.01 | 0.10 | 0.10 | 0.10 | 0.13 | 0.13 | 0.12 | 0.13 | 0.13 | 0.15 |
| | | | | 18 demes | 0.33 | 0.33 | 0.31 | 0.31 | 0.30 | 0.01 | 0.00 | 0.12 | 0.12 | 0.10 | 0.13 | 0.13 | 0.13 | 0.08 | 0.08 | 0.07 |
| | | | prox. rot. | 13 demes | 0.29 | 0.29 | 0.26 | 0.26 | 0.27 | 0.03 | 0.03 | 0.11 | 0.11 | 0.11 | 0.13 | 0.13 | 0.12 | 0.15 | 0.15 | 0.15 |
| | | | | 18 demes | 0.32 | 0.32 | 0.32 | 0.32 | 0.31 | -0.01 | -0.01 | 0.10 | 0.10 | 0.08 | 0.13 | 0.13 | 0.12 | 0.06 | 0.06 | 0.05 |
| | | | dist. non-rot. | 13 demes | 0.31 | 0.31 | 0.33 | 0.33 | 0.33 | -0.01 | -0.01 | 0.09 | 0.09 | 0.09 | 0.04 | 0.04 | 0.04 | 0.06 | 0.06 | 0.08 |
| | | | | 18 demes | 0.33 | 0.33 | 0.32 | 0.32 | 0.32 | -0.01 | -0.01 | 0.07 | 0.07 | 0.07 | 0.13 | 0.13 | 0.13 | 0.01 | 0.01 | 0.02 |
| | | | prox. non-rot. | 13 demes | 0.31 | 0.31 | 0.25 | 0.25 | 0.25 | 0.00 | 0.00 | 0.05 | 0.05 | 0.05 | -0.09 | -0.09 | -0.09 | 0.04 | 0.04 | 0.04 |
| | | | | 18 demes | 0.34 | 0.35 | 0.35 | 0.35 | 0.35 | 0.00 | 0.00 | 0.07 | 0.07 | 0.07 | 0.08 | 0.08 | 0.08 | -0.04 | -0.04 | -0.03 |

*p*-value > 0.05/7980 (Bonferroni correction)

**|Spearman correlation coefficient| >= 0.5**

| part 5 | | | | variable 2: | max. \|estimated admixture fractions - equal fractions\| | | | | | *p*-value | | | | | max. SE | | | *p*-value | | |
|---|---|---|---|---|---|---|---|---|---|---|---|---|---|---|---|---|---|---|---|---|
| total number of randomized models: | | | | | 21,663,216 | 12,439,194 | 2,392,357 | 2,040,355 | 1,058,613 | 21,663,216 | 12,439,194 | 2,392,357 | 2,040,355 | 1,058,613 | 4,378,913 | 3,566,530 | 1,533,911 | 4,378,913 | 3,566,530 | 1,533,911 |
| total number of systematic models (10-5 to 10-2): | | | | | 3,576,667 | 1,158,133 | 1,267,603 | 1,064,846 | 240,961 | 3,576,667 | 1,158,133 | 1,267,603 | 1,064,846 | 240,961 | N/A | | | N/A | | |
| variable 1: | model complexity | gene-flow intensity per gen. | *qpAdm* protocol | sampling density | p >= 1e-55 | p >= 1e-5 | max. \|*EAF–EF*\| <= 0.5 | p >= 1e-55; max. \|*EAF–EF*\| <= 0.5 | p >= 1e-5; max. \|*EAF –EF*\| <= 0.5 | p >= 1e-55 | p >= 1e-5 | max. \|*EAF–EF*\| <= 0.5 | p >= 1e-55; max. \|*EAF–EF*\| <= 0.5 | p >= 1e-5; max. \|*EAF –EF*\| <= 0.5 | max. SE <= 0.1 | p >= 1e-55; max. SE <= 0.1 | p >= 1e-5; max. SE <= 0.1 | max. SE <= 0.1 | p >= 1e-55; max. SE <= 0.1 | p >= 1e-5; max. SE <= 0.1 |
| distance to the ideal symmetric model (in the space of avg. distance vs. min. angle) | 2-way | 10-5 to 10-4 | dist. rot. | 13 demes | 0.10 | 0.19 | -0.02 | -0.02 | 0.02 | 0.03 | 0.00 | -0.04 | -0.03 | -0.02 | 0.03 | 0.03 | 0.02 | -0.03 | -0.04 | -0.01 |
| | | | | 18 demes | 0.16 | 0.32 | 0.04 | 0.02 | 0.02 | 0.01 | 0.02 | -0.08 | -0.09 | -0.06 | -0.03 | -0.03 | 0.05 | -0.11 | -0.13 | -0.04 |
| | | | prox. rot. | 13 demes | 0.11 | 0.19 | 0.04 | 0.03 | 0.02 | 0.00 | -0.02 | -0.06 | -0.07 | -0.05 | -0.05 | -0.05 | -0.01 | -0.08 | -0.10 | -0.04 |
| | | | | 18 demes | 0.13 | 0.27 | 0.04 | 0.03 | 0.01 | 0.01 | 0.02 | -0.07 | -0.05 | -0.01 | -0.02 | -0.03 | 0.03 | -0.10 | -0.10 | -0.03 |
| | | | dist. non-rot. | 13 demes | 0.12 | 0.17 | 0.05 | 0.05 | 0.05 | -0.02 | 0.02 | -0.05 | -0.04 | 0.01 | -0.05 | -0.04 | -0.04 | -0.11 | -0.14 | 0.00 |
| | | | | 18 demes | 0.14 | 0.28 | 0.05 | 0.05 | -0.01 | 0.01 | -0.02 | -0.05 | -0.05 | -0.06 | 0.01 | 0.01 | 0.03 | -0.06 | -0.10 | -0.04 |
| | | | prox. non-rot. | 13 demes | 0.13 | 0.18 | 0.05 | 0.05 | 0.06 | 0.05 | 0.04 | -0.02 | 0.01 | 0.01 | -0.01 | -0.01 | -0.05 | -0.04 | -0.03 | 0.03 |
| | | | | 18 demes | 0.14 | 0.31 | 0.05 | 0.05 | 0.09 | 0.01 | 0.08 | -0.07 | -0.09 | -0.03 | 0.03 | 0.04 | 0.03 | -0.05 | -0.09 | -0.01 |
| | | 10-4 to 10-3 | dist. rot. | 13 demes | **0.62** | **0.79** | 0.17 | 0.24 | 0.33 | 0.04 | 0.13 | -0.21 | -0.10 | -0.17 | 0.09 | 0.11 | 0.10 | -0.17 | -0.05 | -0.12 |
| | | | | 18 demes | **0.73** | **0.81** | 0.19 | 0.31 | **0.51** | 0.07 | 0.43 | -0.25 | -0.04 | -0.12 | 0.07 | 0.13 | 0.08 | -0.22 | -0.06 | -0.01 |
| | | | prox. rot. | 13 demes | **0.65** | **0.79** | 0.11 | 0.15 | 0.24 | 0.05 | 0.23 | -0.20 | -0.10 | -0.18 | 0.11 | 0.16 | 0.13 | -0.17 | -0.07 | -0.02 |
| | | | | 18 demes | **0.73** | **0.84** | 0.19 | 0.30 | 0.34 | 0.09 | 0.45 | -0.25 | -0.04 | -0.26 | 0.10 | 0.10 | 0.15 | -0.21 | -0.02 | -0.06 |
| | | | dist. non-rot. | 13 demes | **0.58** | **0.67** | 0.16 | 0.19 | 0.35 | -0.02 | -0.05 | -0.15 | -0.10 | 0.12 | 0.09 | 0.10 | 0.08 | -0.16 | -0.06 | 0.05 |
| | | | | 18 demes | **0.67** | **0.72** | 0.15 | 0.26 | 0.30 | -0.02 | -0.10 | -0.21 | -0.03 | -0.12 | 0.09 | 0.10 | -0.01 | -0.18 | -0.02 | -0.09 |
| | | | prox. non-rot. | 13 demes | **0.59** | **0.76** | 0.18 | 0.23 | 0.31 | 0.01 | 0.24 | -0.18 | -0.15 | -0.07 | 0.09 | 0.12 | 0.12 | -0.19 | -0.09 | 0.08 |
| | | | | 18 demes | **0.72** | **0.83** | 0.18 | 0.29 | 0.38 | 0.11 | 0.40 | -0.21 | 0.00 | 0.01 | 0.08 | 0.08 | -0.13 | -0.19 | 0.00 | 0.06 |
| | | 10-5 to 10-2 | dist. rot. | 13 demes | **0.78** | **0.80** | 0.20 | 0.28 | 0.32 | 0.00 | 0.05 | -0.34 | -0.08 | 0.02 | 0.25 | 0.22 | 0.09 | -0.18 | -0.08 | -0.01 |
| | | | | 18 demes | **0.81** | **0.83** | 0.24 | 0.35 | 0.29 | 0.02 | 0.06 | -0.36 | -0.06 | 0.01 | 0.28 | 0.22 | 0.08 | -0.16 | -0.06 | -0.02 |
| | | | | systematic | **0.82** | **0.68** | 0.19 | 0.21 | 0.23 | 0.17 | 0.06 | -0.27 | -0.17 | -0.12 | | | | | | |
| | | | prox. rot. | 13 demes | **0.77** | **0.80** | 0.19 | 0.26 | 0.31 | 0.02 | 0.09 | -0.35 | -0.09 | 0.02 | 0.24 | 0.22 | 0.09 | -0.19 | -0.06 | -0.01 |
| | | | | 18 demes | **0.81** | **0.83** | 0.25 | 0.36 | 0.38 | 0.02 | 0.11 | -0.39 | -0.06 | 0.07 | 0.26 | 0.21 | 0.07 | -0.19 | -0.05 | 0.03 |
| | | | dist. non-rot. | 13 demes | **0.78** | **0.77** | 0.26 | 0.34 | **0.54** | -0.02 | 0.06 | -0.35 | -0.08 | 0.10 | 0.25 | 0.22 | 0.07 | -0.21 | -0.08 | 0.11 |
| | | | | 18 demes | **0.82** | **0.82** | 0.25 | 0.38 | 0.40 | -0.04 | -0.07 | -0.42 | -0.08 | -0.04 | 0.26 | 0.20 | 0.03 | -0.24 | -0.09 | -0.07 |
| | | | | systematic | **0.80** | **0.85** | 0.21 | 0.25 | 0.37 | 0.02 | -0.05 | -0.44 | -0.25 | 0.00 | | | | | | |
| | | | prox. non-rot. | 13 demes | **0.78** | **0.77** | 0.17 | 0.23 | 0.43 | -0.01 | 0.08 | -0.40 | -0.11 | -0.01 | 0.24 | 0.21 | 0.06 | -0.22 | -0.06 | -0.01 |
| | | | | 18 demes | **0.83** | **0.85** | 0.20 | 0.32 | 0.37 | 0.00 | 0.11 | **-0.50** | -0.14 | -0.16 | 0.27 | 0.22 | 0.06 | -0.22 | -0.06 | -0.15 |
| | | 10-3 to 10-2 | dist. rot. | 13 demes | **0.79** | **0.83** | 0.29 | 0.31 | 0.42 | -0.04 | 0.01 | **-0.53** | -0.49 | -0.12 | 0.28 | 0.27 | 0.15 | -0.28 | -0.25 | -0.07 |
| | | | | 18 demes | **0.82** | **0.84** | 0.32 | 0.37 | **0.52** | 0.00 | 0.02 | **-0.53** | -0.42 | -0.08 | 0.33 | 0.31 | 0.17 | -0.25 | -0.19 | -0.04 |
| | | | prox. rot. | 13 demes | **0.78** | **0.82** | 0.23 | 0.24 | 0.34 | -0.01 | 0.03 | **-0.51** | -0.47 | -0.09 | 0.26 | 0.25 | 0.15 | -0.22 | -0.20 | -0.03 |
| | | | | 18 demes | **0.81** | **0.84** | 0.25 | 0.29 | 0.39 | -0.01 | 0.02 | **-0.59** | -0.49 | -0.09 | 0.32 | 0.31 | 0.20 | -0.25 | -0.19 | -0.03 |
| | | | dist. non-rot. | 13 demes | **0.79** | **0.84** | 0.34 | 0.35 | 0.47 | -0.06 | -0.02 | **-0.57** | **-0.56** | -0.21 | 0.29 | 0.29 | 0.22 | -0.29 | -0.28 | -0.08 |
| | | | | 18 demes | **0.81** | **0.85** | 0.33 | 0.37 | **0.55** | -0.04 | -0.03 | **-0.58** | **-0.52** | -0.17 | 0.33 | 0.32 | 0.22 | -0.27 | -0.24 | -0.07 |
| | | | prox. non-rot. | 13 demes | **0.78** | **0.84** | 0.24 | 0.25 | 0.43 | -0.02 | 0.03 | **-0.59** | **-0.57** | -0.25 | 0.28 | 0.27 | 0.23 | -0.26 | -0.25 | -0.08 |
| | | | | 18 demes | **0.81** | **0.85** | 0.29 | 0.32 | 0.47 | 0.01 | 0.02 | **-0.61** | **-0.56** | -0.21 | 0.32 | 0.32 | 0.25 | -0.24 | -0.21 | -0.07 |
| | 3-way | 10-5 to 10-4 | dist. rot. | 13 demes | 0.07 | 0.11 | -0.03 | -0.03 | -0.03 | 0.06 | 0.03 | -0.04 | -0.04 | -0.02 | 0.01 | 0.02 | 0.02 | -0.05 | -0.04 | 0.00 |
| | | | | 18 demes | 0.14 | 0.24 | -0.02 | -0.02 | 0.01 | 0.06 | 0.06 | -0.07 | -0.07 | -0.06 | -0.03 | -0.03 | 0.01 | -0.06 | -0.06 | -0.01 |
| | | | prox. rot. | 13 demes | 0.09 | 0.13 | 0.03 | 0.03 | 0.05 | 0.05 | 0.01 | -0.02 | -0.02 | -0.02 | 0.01 | 0.01 | 0.05 | -0.01 | -0.02 | -0.01 |
| | | | | 18 demes | 0.12 | 0.20 | -0.01 | -0.01 | 0.00 | 0.06 | 0.06 | -0.01 | -0.02 | -0.03 | -0.06 | -0.06 | -0.03 | -0.06 | -0.06 | -0.02 |
| | | | dist. non-rot. | 13 demes | 0.09 | 0.12 | 0.04 | 0.04 | 0.03 | 0.00 | 0.02 | -0.04 | -0.03 | -0.03 | -0.08 | -0.08 | -0.02 | -0.12 | -0.13 | -0.01 |
| | | | | 18 demes | 0.12 | 0.19 | 0.00 | 0.00 | -0.02 | 0.03 | 0.01 | -0.02 | -0.02 | -0.08 | -0.03 | -0.03 | 0.01 | -0.06 | -0.08 | -0.02 |
| | | | prox. non-rot. | 13 demes | 0.11 | 0.13 | 0.00 | 0.00 | 0.01 | 0.08 | 0.06 | 0.00 | 0.01 | 0.02 | -0.07 | -0.06 | -0.06 | -0.01 | -0.01 | 0.05 |
| | | | | 18 demes | 0.12 | 0.25 | -0.01 | -0.01 | 0.04 | 0.07 | 0.11 | -0.09 | -0.11 | -0.02 | 0.00 | 0.00 | 0.02 | -0.05 | -0.08 | 0.02 |
| | | 10-4 to 10-3 | dist. rot. | 13 demes | 0.45 | **0.57** | 0.15 | 0.18 | 0.23 | 0.09 | 0.12 | -0.09 | -0.04 | -0.13 | 0.08 | 0.08 | 0.03 | -0.05 | -0.01 | -0.05 |
| | | | | 18 demes | **0.56** | **0.71** | 0.14 | 0.16 | 0.22 | 0.11 | 0.32 | -0.12 | -0.03 | -0.01 | 0.05 | 0.06 | 0.07 | -0.07 | -0.02 | 0.02 |
| | | | prox. rot. | 13 demes | 0.47 | **0.60** | 0.12 | 0.14 | 0.19 | 0.09 | 0.17 | -0.08 | -0.07 | -0.12 | 0.07 | 0.08 | 0.06 | -0.06 | -0.06 | 0.00 |
| | | | | 18 demes | **0.58** | **0.72** | 0.14 | 0.17 | 0.11 | 0.12 | 0.32 | -0.06 | -0.03 | -0.01 | 0.06 | 0.06 | 0.12 | -0.04 | 0.00 | -0.03 |
| | | | dist. non-rot. | 13 demes | 0.40 | 0.50 | 0.09 | 0.10 | 0.16 | 0.02 | -0.02 | -0.04 | -0.08 | -0.05 | 0.03 | 0.01 | 0.00 | -0.01 | -0.01 | 0.08 |
| | | | | 18 demes | **0.50** | **0.57** | 0.08 | 0.14 | 0.18 | 0.01 | -0.04 | -0.08 | -0.05 | -0.14 | 0.06 | 0.04 | -0.01 | -0.05 | 0.00 | 0.00 |
| | | | prox. non-rot. | 13 demes | 0.46 | **0.59** | 0.13 | 0.15 | 0.14 | 0.05 | 0.18 | -0.06 | -0.09 | -0.13 | 0.07 | 0.08 | 0.04 | -0.06 | -0.06 | 0.03 |
| | | | | 18 demes | **0.58** | **0.72** | 0.14 | 0.21 | 0.20 | 0.13 | 0.32 | -0.06 | -0.01 | -0.11 | 0.05 | 0.06 | -0.04 | -0.05 | 0.03 | 0.02 |
| | | 10-5 to 10-2 | dist. rot. | 13 demes | **0.56** | **0.62** | 0.29 | 0.29 | 0.24 | 0.05 | 0.04 | 0.01 | 0.05 | 0.01 | 0.18 | 0.18 | 0.11 | -0.03 | -0.02 | -0.01 |
| | | | | 18 demes | **0.61** | **0.67** | 0.34 | 0.32 | 0.25 | 0.08 | 0.07 | 0.03 | 0.06 | 0.05 | 0.21 | 0.21 | 0.13 | 0.01 | 0.02 | 0.01 |
| | | | | systematic | **0.63** | 0.36 | 0.25 | 0.25 | 0.39 | 0.34 | 0.13 | 0.03 | -0.01 | 0.00 | | | | | | |
| | | | prox. rot. | 13 demes | **0.55** | **0.61** | 0.26 | 0.25 | 0.19 | 0.07 | 0.07 | 0.04 | 0.05 | 0.03 | 0.16 | 0.15 | 0.09 | 0.00 | 0.00 | -0.01 |
| | | | | 18 demes | **0.61** | **0.67** | 0.34 | 0.30 | 0.19 | 0.07 | 0.08 | 0.03 | 0.05 | 0.06 | 0.20 | 0.20 | 0.11 | 0.00 | -0.01 | 0.00 |
| | | | dist. non-rot. | 13 demes | **0.56** | **0.58** | 0.32 | 0.32 | 0.23 | 0.01 | 0.02 | -0.01 | 0.03 | 0.03 | 0.21 | 0.22 | 0.18 | -0.03 | 0.00 | 0.08 |
| | | | | 18 demes | **0.62** | **0.66** | 0.33 | 0.32 | 0.29 | 0.00 | -0.03 | -0.06 | 0.01 | 0.00 | 0.21 | 0.21 | 0.15 | -0.05 | -0.03 | -0.02 |
| | | | | systematic | **0.70** | **0.65** | 0.29 | 0.31 | 0.37 | 0.13 | -0.02 | -0.23 | -0.16 | -0.01 | | | | | | |
| | | | prox. non-rot. | 13 demes | **0.57** | **0.59** | 0.29 | 0.29 | 0.26 | 0.04 | 0.03 | -0.05 | -0.01 | -0.08 | 0.15 | 0.15 | 0.08 | -0.01 | 0.01 | 0.00 |
| | | | | 18 demes | **0.63** | **0.68** | 0.33 | 0.33 | 0.30 | 0.06 | 0.08 | -0.07 | 0.02 | 0.02 | 0.21 | 0.21 | 0.14 | -0.02 | 0.01 | -0.11 |
| | | 10-3 to 10-2 | dist. rot. | 13 demes | **0.58** | **0.60** | 0.40 | 0.40 | 0.39 | 0.03 | 0.02 | -0.02 | -0.02 | 0.02 | 0.17 | 0.17 | 0.16 | -0.10 | -0.10 | -0.05 |
| | | | | 18 demes | **0.60** | **0.62** | 0.43 | 0.43 | 0.42 | 0.06 | 0.04 | 0.00 | 0.00 | 0.05 | 0.24 | 0.24 | 0.21 | -0.07 | -0.07 | -0.01 |
| | | | prox. rot. | 13 demes | **0.56** | **0.58** | 0.35 | 0.35 | 0.35 | 0.07 | 0.04 | 0.03 | 0.03 | 0.06 | 0.17 | 0.17 | 0.17 | -0.01 | -0.01 | 0.03 |
| | | | | 18 demes | **0.59** | **0.61** | 0.43 | 0.43 | 0.42 | 0.04 | 0.02 | -0.02 | -0.02 | 0.02 | 0.23 | 0.23 | 0.22 | -0.08 | -0.08 | -0.01 |
| | | | dist. non-rot. | 13 demes | **0.58** | **0.59** | 0.40 | 0.40 | 0.40 | 0.02 | 0.01 | 0.01 | 0.01 | 0.03 | 0.22 | 0.22 | 0.23 | -0.13 | -0.13 | -0.07 |
| | | | | 18 demes | **0.59** | **0.61** | 0.46 | 0.46 | 0.47 | 0.03 | 0.01 | -0.05 | -0.05 | -0.01 | 0.28 | 0.28 | 0.28 | -0.12 | -0.12 | -0.05 |
| | | | prox. non-rot. | 13 demes | **0.58** | **0.59** | 0.40 | 0.40 | 0.39 | 0.05 | 0.04 | -0.01 | -0.01 | 0.01 | 0.19 | 0.19 | 0.19 | -0.13 | -0.13 | -0.09 |
| | | | | 18 demes | **0.60** | **0.61** | 0.40 | 0.40 | 0.40 | 0.06 | 0.05 | -0.06 | -0.06 | -0.03 | 0.27 | 0.27 | 0.26 | -0.14 | -0.14 | -0.07 |
| | 4-way | 10-5 to 10-4 | dist. rot. | 13 demes | 0.06 | 0.07 | -0.01 | -0.01 | 0.00 | 0.06 | 0.05 | -0.01 | -0.01 | 0.01 | -0.08 | -0.08 | -0.07 | 0.01 | 0.01 | 0.07 |
| | | | | 18 demes | 0.13 | 0.17 | -0.03 | -0.03 | 0.01 | 0.07 | 0.07 | -0.07 | -0.07 | -0.06 | -0.03 | -0.03 | 0.00 | -0.05 | -0.05 | -0.06 |
| | | | prox. rot. | 13 demes | 0.08 | 0.08 | 0.03 | 0.03 | 0.04 | 0.04 | 0.03 | -0.01 | -0.01 | -0.01 | 0.06 | 0.06 | 0.00 | 0.01 | 0.01 | -0.02 |
| | | | | 18 demes | 0.11 | 0.15 | -0.02 | -0.02 | -0.02 | 0.08 | 0.08 | -0.02 | -0.03 | -0.03 | -0.04 | -0.04 | -0.02 | -0.04 | -0.05 | -0.04 |
| | | | dist. non-rot. | 13 demes | 0.07 | 0.08 | 0.05 | 0.05 | 0.04 | 0.01 | 0.01 | -0.05 | -0.05 | -0.03 | -0.06 | -0.06 | -0.05 | -0.09 | -0.09 | -0.09 |
| | | | | 18 demes | 0.10 | 0.13 | -0.02 | -0.01 | -0.04 | 0.03 | 0.02 | 0.01 | 0.02 | -0.08 | -0.03 | -0.03 | 0.00 | -0.05 | -0.06 | -0.06 |
| | | | prox. non-rot. | 13 demes | 0.10 | 0.10 | -0.01 | -0.01 | 0.00 | 0.08 | 0.06 | 0.01 | 0.01 | 0.01 | -0.08 | -0.07 | -0.07 | -0.01 | -0.01 | 0.00 |
| | | | | 18 demes | 0.11 | 0.20 | -0.04 | -0.04 | 0.01 | 0.09 | 0.12 | -0.07 | -0.08 | 0.00 | -0.02 | -0.03 | 0.00 | -0.06 | -0.08 | -0.01 |
| | | 10-4 to 10-3 | dist. rot. | 13 demes | 0.32 | 0.40 | 0.14 | 0.14 | 0.17 | 0.10 | 0.10 | -0.02 | -0.02 | -0.10 | 0.03 | 0.03 | 0.00 | -0.01 | 0.00 | -0.02 |
| | | | | 18 demes | 0.43 | **0.58** | 0.10 | 0.11 | 0.14 | 0.13 | 0.22 | 0.00 | 0.02 | 0.00 | 0.05 | 0.04 | 0.05 | -0.01 | 0.00 | 0.02 |
| | | | prox. rot. | 13 demes | 0.34 | 0.43 | 0.10 | 0.10 | 0.12 | 0.11 | 0.13 | -0.06 | -0.07 | -0.08 | 0.04 | 0.04 | 0.08 | -0.04 | -0.04 | 0.01 |
| | | | | 18 demes | 0.45 | **0.60** | 0.10 | 0.12 | 0.11 | 0.14 | 0.22 | 0.04 | 0.02 | -0.04 | 0.04 | 0.03 | 0.03 | 0.03 | 0.03 | -0.03 |
| | | | dist. non-rot. | 13 demes | 0.29 | 0.35 | 0.08 | 0.08 | 0.13 | 0.03 | 0.00 | -0.03 | -0.07 | -0.11 | -0.01 | -0.02 | -0.01 | 0.05 | 0.03 | 0.07 |
| | | | | 18 demes | 0.36 | 0.43 | 0.07 | 0.11 | 0.20 | 0.02 | -0.02 | -0.04 | -0.04 | -0.13 | 0.03 | 0.01 | -0.03 | 0.00 | 0.02 | -0.01 |
| | | | prox. non-rot. | 13 demes | 0.37 | 0.46 | 0.15 | 0.15 | 0.23 | 0.08 | 0.12 | -0.02 | -0.04 | -0.15 | 0.03 | 0.03 | -0.01 | -0.01 | -0.02 | -0.05 |
| | | | | 18 demes | 0.45 | **0.61** | 0.10 | 0.12 | 0.14 | 0.14 | 0.23 | 0.00 | -0.03 | -0.04 | 0.03 | 0.04 | -0.01 | 0.02 | 0.04 | 0.01 |
| | | 10-5 to 10-2 | dist. rot. | 13 demes | 0.42 | 0.45 | 0.18 | 0.17 | 0.14 | 0.06 | 0.05 | 0.12 | 0.11 | 0.05 | 0.14 | 0.14 | 0.11 | 0.05 | 0.05 | 0.03 |
| | | | | 18 demes | 0.46 | **0.51** | 0.22 | 0.21 | 0.14 | 0.11 | 0.06 | 0.19 | 0.17 | 0.06 | 0.15 | 0.14 | 0.09 | 0.09 | 0.08 | 0.02 |
| | | | | systematic | 0.34 | 0.16 | 0.20 | 0.20 | 0.31 | 0.41 | 0.12 | 0.19 | 0.15 | 0.10 | | | | | | |
| | | | prox. rot. | 13 demes | 0.41 | 0.44 | 0.16 | 0.16 | 0.14 | 0.08 | 0.06 | 0.12 | 0.11 | 0.04 | 0.09 | 0.08 | 0.06 | 0.05 | 0.05 | 0.01 |
| | | | | 18 demes | 0.46 | **0.51** | 0.20 | 0.18 | 0.10 | 0.10 | 0.06 | 0.17 | 0.15 | 0.06 | 0.16 | 0.15 | 0.11 | 0.07 | 0.06 | 0.04 |
| | | | dist. non-rot. | 13 demes | 0.43 | 0.43 | 0.23 | 0.22 | 0.11 | 0.02 | 0.02 | 0.14 | 0.15 | 0.06 | 0.16 | 0.16 | 0.15 | 0.08 | 0.09 | 0.08 |
| | | | | 18 demes | 0.46 | **0.50** | 0.24 | 0.24 | 0.22 | 0.01 | 0.00 | 0.09 | 0.08 | 0.02 | 0.15 | 0.15 | 0.11 | 0.02 | 0.02 | 0.02 |
| | | | | systematic | **0.51** | 0.43 | 0.29 | 0.30 | 0.30 | 0.18 | 0.01 | -0.03 | -0.03 | 0.06 | | | | | | |
| | | | prox. non-rot. | 13 demes | 0.44 | 0.46 | 0.19 | 0.18 | 0.13 | 0.06 | 0.02 | 0.11 | 0.10 | -0.06 | 0.12 | 0.12 | 0.08 | 0.08 | 0.08 | 0.01 |
| | | | | 18 demes | 0.49 | **0.55** | 0.22 | 0.21 | 0.17 | 0.09 | 0.07 | 0.13 | 0.13 | 0.07 | 0.15 | 0.15 | 0.12 | 0.05 | 0.05 | -0.07 |
| | | 10-3 to 10-2 | dist. rot. | 13 demes | 0.41 | 0.41 | 0.28 | 0.28 | 0.27 | 0.02 | 0.02 | 0.08 | 0.08 | 0.08 | 0.12 | 0.12 | 0.12 | 0.09 | 0.09 | 0.10 |
| | | | | 18 demes | 0.43 | 0.43 | 0.30 | 0.30 | 0.29 | 0.05 | 0.04 | 0.14 | 0.14 | 0.11 | 0.17 | 0.17 | 0.17 | 0.08 | 0.08 | 0.06 |
| | | | prox. rot. | 13 demes | 0.38 | 0.38 | 0.25 | 0.25 | 0.26 | 0.05 | 0.05 | 0.11 | 0.11 | 0.10 | 0.13 | 0.13 | 0.13 | 0.08 | 0.08 | 0.10 |
| | | | | 18 demes | 0.42 | 0.43 | 0.30 | 0.30 | 0.29 | 0.04 | 0.03 | 0.10 | 0.10 | 0.08 | 0.15 | 0.15 | 0.14 | 0.05 | 0.05 | 0.05 |
| | | | dist. non-rot. | 13 demes | 0.40 | 0.40 | 0.31 | 0.31 | 0.31 | 0.02 | 0.02 | 0.12 | 0.12 | 0.12 | 0.05 | 0.05 | 0.05 | 0.05 | 0.05 | 0.07 |
| | | | | 18 demes | 0.44 | 0.44 | 0.33 | 0.33 | 0.33 | 0.03 | 0.03 | 0.09 | 0.09 | 0.09 | 0.16 | 0.16 | 0.16 | -0.01 | -0.01 | 0.00 |
| | | | prox. non-rot. | 13 demes | 0.40 | 0.40 | 0.25 | 0.25 | 0.25 | 0.04 | 0.04 | 0.09 | 0.09 | 0.09 | 0.01 | 0.01 | 0.01 | 0.08 | 0.08 | 0.08 |
| | | | | 18 demes | 0.44 | 0.44 | 0.33 | 0.33 | 0.33 | 0.06 | 0.05 | 0.09 | 0.09 | 0.09 | 0.11 | 0.11 | 0.11 | -0.05 | -0.05 | -0.04 |

*p*-value > 0.05/7980 (Bonferroni correction)

**|Spearman correlation coefficient| >= 0.5**

Spearman's correlation coefficient (ρ) between several model optimality metrics (one metric in each part of the table) and max|*EAF–EF*|, *p*-values, or max. SE in various sections of the respective two-dimensional spaces (in columns). Results are shown for both randomized and systematic *qpAdm* experiments stratified by model complexity, landscape type, *qpAdm* protocol, and landscape sampling density (13 or 18 demes in rotating sets and 10 or 15 demes in non-rotating "right" sets, respectively). Non-significant ρ values (with Bonferroni correction) are underlined, and absolute ρ values ≥0.5 are shown in bold.

**Table S9.**

| gene-flow intensity per generation | model complexity | landscape sampling | percentage of ideal models rejected according to: p < 0.01 | p < 0.05 | *max* $\lvert EAF–EF \rvert$ > 1–*EF* | n |
|---|---|---|---|---|---|---|
| 10-5 to 10-4 | 2-way | randomized | 60.8% | 64.2% | 36.1% | 296 |
| | 3-way | randomized | 42.9% | 45.2% | 40.5% | 42 |
| | 4-way | randomized | 21.7% | 28.3% | 78.3% | 60 |
| 10-4 to 10-3 | 2-way | randomized | 80.8% | 85.3% | 7.2% | 334 |
| | 3-way | randomized | 47.7% | 59.1% | 9.1% | 44 |
| | 4-way | randomized | 44.3% | 55.7% | 50.0% | 70 |
| 10-5 to 10-2 | 2-way | randomized | 55.4% | 68.6% | 5.3% | 395 |
| | | **systematic** | 84.4% | 88.7% | 0.3% | 1,212 |
| | 3-way | randomized | 29.8% | 44.7% | 12.8% | 47 |
| | | **systematic** | 40.4% | 50.1% | 0.0% | 680 |
| | 4-way | randomized | 14.1% | 29.7% | 62.5% | 64 |
| | | **systematic** | 31.0% | 46.4% | 37.1% | 2,835 |
| 10-3 to 10-2 | 2-way | randomized | 17.1% | 30.1% | 6.6% | 286 |
| | 3-way | randomized | 8.6% | 14.3% | 11.4% | 35 |
| | 4-way | randomized | 0.0% | 0.0% | 72.2% | 36 |

Percentage of ideal symmetric models rejected according to various conditions on *p*-values and EAF listed in the table header. Results are shown for both randomized and systematic *qpAdm* experiments stratified by landscape type and model complexity. Total number of ideal symmetric models in each category (both rejected and non-rejected) is shown in the rightmost column.

**Table S10.**

| source-target distance | gene-flow intensity per generation: 10-5 to 10-4 | | 10-4 to 10-3 | | 10-5 to 10-2 | | 10-3 to 10-2 | |
|---|---|---|---|---|---|---|---|---|
| | **p >=0.01** | **p >=0.05** | **p >=0.01** | **p >=0.05** | **p >=0.01** | **p >=0.05** | **p >=0.01** | **p >=0.05** |
| | **55,510** | **47,730** | **5,320** | **4,212** | **8,404** | **5,840** | **21,378** | **13,726** |
| 0 | 8.6% | 8.7% | 81.0% | 84.9% | 50.2% | 55.7% | 16.4% | 19.9% |
| 1 | 18.9% | 19.1% | 14.8% | 12.2% | 42.1% | 38.1% | 68.7% | 68.2% |
| 2 | 17.5% | 17.6% | 3.1% | 2.1% | 7.0% | 5.9% | 13.7% | 11.1% |
| 3 | 9.8% | 9.9% | 0.6% | 0.4% | 0.5% | 0.3% | 1.1% | 0.8% |
| 4 | 10.8% | 10.7% | 0.4% | 0.4% | 0.1% | 0.0% | 0.1% | 0.1% |
| 5 | 10.7% | 10.5% | 0.1% | 0.0% | 0.0% | 0.0% | 0.0% | 0.0% |
| 6 | 9.4% | 9.3% | 0.0% | 0.0% | 0.0% | 0.0% | 0.0% | 0.0% |
| 7 | 7.9% | 7.8% | 0.0% | 0.0% | 0.0% | 0.0% | 0.0% | 0.0% |
| 8 | 5.0% | 5.1% | 0.0% | 0.0% | 0.0% | 0.0% | 0.0% | 0.0% |
| 9 | 1.4% | 1.4% | 0.0% | 0.0% | 0.0% | 0.0% | 0.0% | 0.0% |

Distributions of fitting one-way randomized *qpAdm* models over source-target distances. Results are stratified by landscape type. Total counts of fitting one-way models are shown in the table header.

### Table S11.

| optimality metrics space: | treatment of simpler models | conditions on EAF | *p*-value thresholds | overall *qpAdm* positive rate | | | | | | | | | | | | Spearman correlation coefficient: total count of models vs. count of fitting models in 2D bins | | | | | | | | | | | | Spearman correlation coefficient: total count of models vs. *qpAdm* positive rate in 2D bins | | | | | | | | | | | |
|---|---|---|---|---|---|---|---|---|---|---|---|---|---|---|---|---|---|---|---|---|---|---|---|---|---|---|---|---|---|---|---|---|---|---|---|---|---|---|---|
| | | | | 2-way | | | | 3-way | | | | 4-way | | | | 2-way | | | | 3-way | | | | 4-way | | | | 2-way | | | | 3-way | | | | 4-way | | | |
| | | | | 10-5 - 10-4 | 10-4 - 10-3 | 10-5 - 10-2 | 10-3 - 10-2 | 10-5 - 10-4 | 10-4 - 10-3 | 10-5 - 10-2 | 10-3 - 10-2 | 10-5 - 10-4 | 10-4 - 10-3 | 10-5 - 10-2 | 10-3 - 10-2 | 10-5 - 10-4 | 10-4 - 10-3 | 10-5 - 10-2 | 10-3 - 10-2 | 10-5 - 10-4 | 10-4 - 10-3 | 10-5 - 10-2 | 10-3 - 10-2 | 10-5 - 10-4 | 10-4 - 10-3 | 10-5 - 10-2 | 10-3 - 10-2 | 10-5 - 10-4 | 10-4 - 10-3 | 10-5 - 10-2 | 10-3 - 10-2 | 10-5 - 10-4 | 10-4 - 10-3 | 10-5 - 10-2 | 10-3 - 10-2 | 10-5 - 10-4 | 10-4 - 10-3 | 10-5 - 10-2 | 10-3 - 10-2 |
| max. source-target distance - min. STS angle | simpler models not checked | % (-0.3,1.3) | 0.001 | 1.2E-01 | 8.0E-03 | 2.6E-02 | 1.2E-01 | 1.4E-01 | 1.6E-02 | 5.1E-02 | 1.8E-01 | 1.4E-01 | 2.6E-02 | 7.1E-02 | 1.6E-01 | 0.98 | 0.53 | 0.50 | 0.56 | 1.00 | 0.96 | 0.97 | 0.94 | 1.00 | 0.99 | 0.99 | 0.98 | -0.87 | -0.69 | -0.63 | -0.66 | -0.72 | -0.79 | -0.91 | -0.82 | -0.44 | -0.63 | -0.88 | -0.74 |
| | | | 0.01 | 1.1E-01 | 5.5E-03 | 1.8E-02 | 9.0E-02 | 1.3E-01 | 1.2E-02 | 3.7E-02 | 1.4E-01 | 1.3E-01 | 1.9E-02 | 5.4E-02 | 1.4E-01 | 0.99 | 0.47 | 0.47 | 0.54 | 1.00 | 0.96 | 0.97 | 0.94 | 1.00 | 0.99 | 0.99 | 0.98 | -0.86 | -0.63 | -0.64 | -0.66 | -0.71 | -0.71 | -0.90 | -0.83 | -0.45 | -0.63 | -0.89 | -0.77 |
| | | | 0.05 | 9.9E-02 | 3.6E-03 | 1.2E-02 | 6.1E-02 | 1.2E-01 | 7.8E-03 | 2.6E-02 | 1.1E-01 | 1.1E-01 | 1.3E-02 | 3.9E-02 | 1.2E-01 | 0.98 | 0.42 | 0.45 | 0.53 | 1.00 | 0.95 | 0.97 | 0.94 | 1.00 | 0.98 | 0.99 | 0.98 | -0.87 | -0.48 | -0.58 | -0.61 | -0.70 | -0.70 | -0.91 | -0.84 | -0.45 | -0.62 | -0.90 | -0.82 |
| | | | 0.1 | 8.8E-02 | 2.7E-03 | 9.3E-03 | 4.8E-02 | 1.0E-01 | 6.0E-03 | 2.0E-02 | 9.1E-02 | 1.0E-01 | 1.0E-02 | 3.1E-02 | 1.0E-01 | 0.98 | 0.39 | 0.44 | 0.53 | 1.00 | 0.95 | 0.96 | 0.94 | 1.00 | 0.98 | 0.99 | 0.98 | -0.87 | -0.38 | -0.54 | -0.60 | -0.71 | -0.71 | -0.91 | -0.83 | -0.41 | -0.53 | -0.89 | -0.83 |
| | | | 0.5 | 3.9E-02 | 7.8E-04 | 2.9E-03 | 1.4E-02 | 4.9E-02 | 1.9E-03 | 6.1E-03 | 3.1E-02 | 4.9E-02 | 3.1E-03 | 9.3E-03 | 4.3E-02 | 0.97 | 0.27 | 0.43 | 0.48 | 0.99 | 0.91 | 0.96 | 0.94 | 0.99 | 0.97 | 0.99 | 0.98 | -0.81 | -0.18 | -0.42 | -0.50 | -0.68 | -0.24 | -0.85 | -0.87 | -0.52 | -0.46 | -0.80 | -0.88 |
| | | % (0,1) | 0.001 | 6.4E-02 | 4.8E-03 | 1.6E-02 | 8.0E-02 | 4.2E-02 | 6.1E-03 | 1.6E-02 | 5.5E-02 | 2.6E-02 | 5.9E-03 | 9.7E-03 | 2.2E-02 | 0.95 | 0.39 | 0.29 | 0.09 | 0.99 | 0.89 | 0.85 | 0.57 | 0.99 | 0.97 | 0.96 | 0.77 | -0.74 | -0.47 | -0.57 | -0.67 | -0.68 | -0.69 | -0.90 | -0.71 | -0.46 | -0.57 | -0.87 | -0.69 |
| | | | 0.01 | 5.9E-02 | 3.3E-03 | 1.1E-02 | 5.8E-02 | 3.8E-02 | 4.3E-03 | 1.2E-02 | 4.5E-02 | 2.2E-02 | 4.1E-03 | 7.6E-03 | 1.9E-02 | 0.95 | 0.34 | 0.29 | 0.06 | 0.99 | 0.88 | 0.85 | 0.57 | 0.99 | 0.96 | 0.96 | 0.77 | -0.74 | -0.43 | -0.57 | -0.68 | -0.70 | -0.64 | -0.90 | -0.73 | -0.46 | -0.56 | -0.87 | -0.70 |
| | | | 0.05 | 5.1E-02 | 2.2E-03 | 7.2E-03 | 4.0E-02 | 3.3E-02 | 2.8E-03 | 8.2E-03 | 3.5E-02 | 1.9E-02 | 2.8E-03 | 5.6E-03 | 1.6E-02 | 0.95 | 0.34 | 0.29 | 0.07 | 0.99 | 0.86 | 0.85 | 0.58 | 0.99 | 0.96 | 0.97 | 0.78 | -0.76 | -0.34 | -0.52 | -0.63 | -0.70 | -0.62 | -0.84 | -0.75 | -0.48 | -0.53 | -0.83 | -0.72 |
| | | | 0.1 | 4.6E-02 | 1.6E-03 | 5.6E-03 | 3.1E-02 | 3.0E-02 | 2.2E-03 | 6.5E-03 | 2.9E-02 | 1.7E-02 | 2.1E-03 | 4.5E-03 | 1.4E-02 | 0.95 | 0.29 | 0.29 | 0.08 | 0.99 | 0.85 | 0.84 | 0.59 | 0.99 | 0.96 | 0.97 | 0.79 | -0.76 | -0.20 | -0.52 | -0.63 | -0.68 | -0.55 | -0.84 | -0.75 | -0.48 | -0.52 | -0.84 | -0.72 |
| | | | 0.5 | 2.1E-02 | 4.3E-04 | 1.7E-03 | 8.9E-03 | 1.4E-02 | 6.8E-04 | 1.9E-03 | 1.0E-02 | 7.8E-03 | 6.2E-04 | 1.4E-03 | 5.8E-03 | 0.93 | 0.15 | 0.27 | 0.10 | 0.99 | 0.78 | 0.84 | 0.58 | 0.99 | 0.95 | 0.97 | 0.82 | -0.69 | -0.01 | -0.26 | -0.35 | -0.57 | -0.22 | -0.76 | -0.77 | -0.51 | -0.38 | -0.63 | -0.74 |
| | | %+/-2SE (0,1) | 0.001 | 1.1E-02 | 3.2E-03 | 1.1E-02 | 5.7E-02 | 2.6E-03 | 2.2E-03 | 4.6E-03 | 1.5E-02 | 1.3E-03 | 1.1E-03 | 8.3E-04 | 5.2E-04 | 0.77 | 0.34 | 0.23 | -0.06 | 0.95 | 0.75 | 0.53 | 0.12 | 0.95 | 0.90 | 0.72 | 0.18 | -0.43 | 0.00 | -0.38 | -0.58 | 0.04 | -0.38 | -0.68 | -0.54 | 0.25 | -0.30 | -0.48 | -0.27 |
| | | | 0.01 | 9.4E-03 | 2.1E-03 | 7.5E-03 | 4.1E-02 | 1.5E-03 | 1.4E-03 | 3.2E-03 | 1.2E-02 | 6.1E-04 | 6.6E-04 | 5.7E-04 | 4.5E-04 | 0.74 | 0.29 | 0.26 | -0.08 | 0.93 | 0.70 | 0.52 | 0.11 | 0.94 | 0.88 | 0.73 | 0.16 | -0.45 | -0.01 | -0.28 | -0.55 | 0.04 | -0.27 | -0.67 | -0.55 | 0.19 | -0.30 | -0.35 | -0.29 |
| | | | 0.05 | 7.9E-03 | 1.4E-03 | 4.8E-03 | 2.8E-02 | 8.7E-04 | 8.8E-04 | 2.1E-03 | 8.7E-03 | 2.7E-04 | 3.7E-04 | 3.9E-04 | 3.7E-04 | 0.74 | 0.22 | 0.25 | -0.07 | 0.90 | 0.67 | 0.51 | 0.11 | 0.92 | 0.87 | 0.72 | 0.14 | -0.45 | -0.02 | -0.23 | -0.50 | 0.08 | -0.22 | -0.61 | -0.56 | 0.52 | -0.11 | -0.27 | -0.28 |
| | | | 0.1 | 7.0E-03 | 1.1E-03 | 3.7E-03 | 2.2E-02 | 6.9E-04 | 6.5E-04 | 1.7E-03 | 7.0E-03 | 1.7E-04 | 2.7E-04 | 3.1E-04 | 3.1E-04 | 0.74 | 0.17 | 0.24 | -0.07 | 0.89 | 0.64 | 0.49 | 0.11 | 0.91 | 0.85 | 0.72 | 0.15 | -0.41 | 0.02 | -0.17 | -0.46 | 0.11 | -0.14 | -0.48 | -0.57 | 0.66 | -0.12 | -0.28 | -0.23 |
| | | | 0.5 | 2.8E-03 | 2.9E-04 | 1.0E-03 | 6.4E-03 | 2.2E-04 | 1.7E-04 | 4.4E-04 | 2.2E-03 | 2.5E-05 | 6.1E-05 | 7.8E-05 | 9.4E-05 | 0.65 | 0.04 | 0.12 | -0.12 | 0.87 | 0.51 | 0.47 | 0.09 | 0.82 | 0.78 | 0.71 | 0.15 | -0.31 | -0.01 | -0.07 | -0.35 | 0.45 | 0.06 | -0.36 | -0.50 | 0.65 | 0.24 | 0.07 | -0.04 |
| | "trailing" simpler models rejected | % (-0.3,1.3) | 0.001 | 1.4E-02 | 4.0E-03 | 1.3E-02 | 5.2E-02 | 3.2E-03 | 3.3E-03 | 7.6E-03 | 1.9E-02 | 1.2E-03 | 2.0E-03 | 2.8E-03 | 1.1E-03 | 0.69 | 0.39 | 0.37 | 0.43 | 0.92 | 0.82 | 0.84 | 0.75 | 0.97 | 0.96 | 0.96 | 0.91 | -0.28 | -0.23 | -0.45 | -0.40 | -0.05 | -0.51 | -0.61 | -0.28 | 0.21 | -0.38 | -0.24 | 0.06 |
| | | | 0.01 | 1.4E-02 | 3.0E-03 | 9.8E-03 | 4.4E-02 | 2.7E-03 | 2.5E-03 | 6.6E-03 | 2.0E-02 | 8.6E-04 | 1.7E-03 | 2.7E-03 | 1.7E-03 | 0.70 | 0.37 | 0.40 | 0.41 | 0.89 | 0.80 | 0.83 | 0.78 | 0.96 | 0.94 | 0.95 | 0.92 | -0.32 | -0.19 | -0.37 | -0.52 | 0.09 | -0.39 | -0.65 | -0.39 | -0.01 | -0.36 | -0.41 | 0.01 |
| | | | 0.05 | 1.5E-02 | 2.2E-03 | 7.0E-03 | 3.4E-02 | 2.9E-03 | 2.0E-03 | 5.5E-03 | 2.0E-02 | 7.4E-04 | 1.5E-03 | 2.7E-03 | 2.5E-03 | 0.75 | 0.31 | 0.37 | 0.42 | 0.89 | 0.78 | 0.85 | 0.79 | 0.94 | 0.94 | 0.96 | 0.92 | -0.39 | -0.21 | -0.38 | -0.43 | 0.00 | -0.11 | -0.68 | -0.54 | 0.34 | -0.27 | -0.56 | -0.15 |
| | | | 0.1 | 1.5E-02 | 1.6E-03 | 5.4E-03 | 2.8E-02 | 3.1E-03 | 1.7E-03 | 4.7E-03 | 1.8E-02 | 7.6E-04 | 1.3E-03 | 2.6E-03 | 2.9E-03 | 0.78 | 0.28 | 0.35 | 0.42 | 0.89 | 0.78 | 0.84 | 0.80 | 0.93 | 0.93 | 0.96 | 0.92 | -0.34 | -0.03 | -0.28 | -0.40 | 0.02 | -0.29 | -0.60 | -0.45 | 0.34 | -0.21 | -0.49 | -0.22 |
| | | | 0.5 | 1.0E-02 | 5.4E-04 | 1.8E-03 | 9.5E-03 | 3.8E-03 | 6.8E-04 | 1.8E-03 | 8.9E-03 | 1.1E-03 | 6.2E-04 | 1.5E-03 | 2.9E-03 | 0.82 | 0.23 | 0.32 | 0.37 | 0.95 | 0.76 | 0.84 | 0.85 | 0.96 | 0.93 | 0.96 | 0.96 | -0.37 | -0.02 | -0.21 | -0.36 | -0.20 | 0.02 | -0.46 | -0.67 | 0.10 | -0.27 | -0.41 | -0.52 |
| | | | 0.001, 0.05 | 8.6E-03 | 1.4E-03 | 4.5E-03 | 1.7E-02 | 9.0E-04 | 7.4E-04 | 1.5E-03 | 3.2E-03 | 6.9E-05 | 2.8E-04 | 3.2E-04 | 2.2E-05 | 0.59 | 0.15 | 0.31 | 0.35 | 0.62 | 0.57 | 0.66 | 0.59 | 0.71 | 0.83 | 0.90 | 0.73 | -0.19 | -0.01 | -0.04 | -0.21 | 0.24 | 0.07 | -0.42 | -0.11 | 0.52 | -0.06 | 0.09 | 0.64 |
| | | | 0.001, 0.5 | 3.0E-03 | 2.2E-04 | 7.7E-04 | 2.6E-03 | 2.3E-04 | 1.2E-04 | 1.8E-04 | 2.2E-04 | 1.6E-05 | 3.4E-05 | 2.4E-05 | 2.5E-07 | 0.50 | 0.01 | 0.15 | 0.11 | 0.52 | 0.19 | 0.44 | 0.36 | 0.43 | 0.55 | 0.65 | 0.08 | 0.05 | -0.01 | 0.07 | 0.00 | 0.29 | 0.02 | 0.09 | 0.19 | 0.37 | 0.22 | 0.46 | 0.08 |
| | | % (0,1) | 0.001 | 8.4E-03 | 2.7E-03 | 9.0E-03 | 4.2E-02 | 1.9E-03 | 1.5E-03 | 2.9E-03 | 6.5E-03 | 9.5E-04 | 6.9E-04 | 4.1E-04 | 1.5E-04 | 0.70 | 0.32 | 0.18 | -0.06 | 0.94 | 0.69 | 0.49 | 0.16 | 0.96 | 0.91 | 0.72 | 0.47 | -0.35 | 0.10 | -0.26 | -0.48 | 0.08 | -0.28 | -0.51 | -0.25 | 0.35 | -0.28 | -0.30 | 0.11 |
| | | | 0.01 | 7.9E-03 | 2.0E-03 | 6.7E-03 | 3.4E-02 | 1.3E-03 | 1.1E-03 | 2.5E-03 | 7.3E-03 | 5.9E-04 | 5.4E-04 | 4.2E-04 | 2.6E-04 | 0.69 | 0.24 | 0.21 | -0.10 | 0.91 | 0.65 | 0.52 | 0.18 | 0.95 | 0.89 | 0.75 | 0.50 | -0.42 | 0.04 | -0.16 | -0.60 | 0.11 | -0.14 | -0.61 | -0.36 | 0.04 | -0.29 | -0.21 | 0.05 |
| | | | 0.05 | 8.2E-03 | 1.4E-03 | 4.7E-03 | 2.6E-02 | 1.0E-03 | 8.8E-04 | 2.1E-03 | 7.2E-03 | 3.5E-04 | 4.0E-04 | 4.3E-04 | 4.0E-04 | 0.72 | 0.20 | 0.22 | -0.08 | 0.89 | 0.63 | 0.56 | 0.19 | 0.95 | 0.87 | 0.79 | 0.42 | -0.50 | 0.01 | -0.20 | -0.51 | 0.06 | -0.17 | -0.57 | -0.50 | 0.31 | -0.15 | -0.42 | -0.20 |
| | | | 0.1 | 8.2E-03 | 1.0E-03 | 3.7E-03 | 2.1E-02 | 1.0E-03 | 7.5E-04 | 1.8E-03 | 6.6E-03 | 2.8E-04 | 3.3E-04 | 4.3E-04 | 4.6E-04 | 0.75 | 0.17 | 0.19 | -0.05 | 0.89 | 0.63 | 0.53 | 0.20 | 0.94 | 0.86 | 0.83 | 0.44 | -0.45 | 0.06 | -0.18 | -0.49 | 0.03 | -0.23 | -0.43 | -0.44 | 0.43 | -0.15 | -0.33 | -0.18 |
| | | | 0.5 | 5.7E-03 | 3.3E-04 | 1.2E-03 | 6.9E-03 | 1.2E-03 | 2.9E-04 | 6.7E-04 | 3.2E-03 | 2.3E-04 | 1.5E-04 | 2.5E-04 | 4.2E-04 | 0.73 | 0.06 | 0.17 | -0.06 | 0.92 | 0.58 | 0.59 | 0.22 | 0.89 | 0.89 | 0.89 | 0.54 | -0.41 | -0.02 | -0.08 | -0.34 | -0.07 | 0.00 | -0.44 | -0.67 | 0.14 | -0.21 | -0.22 | -0.28 |
| | | | 0.001, 0.05 | 4.8E-03 | 9.8E-04 | 3.3E-03 | 1.5E-02 | 2.2E-04 | 3.6E-04 | 7.7E-04 | 1.6E-03 | 2.2E-05 | 9.0E-05 | 7.4E-05 | 1.7E-06 | 0.59 | 0.05 | 0.15 | -0.11 | 0.62 | 0.47 | 0.38 | -0.03 | 0.70 | 0.74 | 0.67 | 0.10 | -0.29 | 0.00 | 0.03 | -0.34 | 0.31 | 0.01 | -0.24 | -0.22 | 0.49 | 0.00 | 0.26 | 0.10 |
| | | | 0.001, 0.5 | 1.6E-03 | 1.7E-04 | 6.1E-04 | 2.5E-03 | 5.5E-05 | 5.6E-05 | 1.0E-04 | 1.6E-04 | 3.0E-06 | 1.1E-05 | 7.2E-06 | 0.0E+00 | 0.45 | -0.06 | 0.01 | -0.09 | 0.39 | 0.01 | 0.08 | -0.13 | 0.33 | 0.28 | 0.30 | NA | -0.03 | -0.07 | -0.02 | -0.14 | 0.20 | -0.05 | -0.04 | -0.18 | 0.32 | 0.16 | 0.26 | NA |
| | | %+/-2SE (0,1) | 0.001 | 5.4E-03 | 2.5E-03 | 8.6E-03 | 4.1E-02 | 1.3E-03 | 1.3E-03 | 2.5E-03 | 5.7E-03 | 5.9E-04 | 5.1E-04 | 2.8E-04 | 5.4E-05 | 0.69 | 0.30 | 0.16 | -0.05 | 0.94 | 0.67 | 0.46 | 0.04 | 0.95 | 0.86 | 0.64 | 0.10 | -0.32 | 0.09 | -0.26 | -0.45 | 0.14 | -0.26 | -0.48 | -0.21 | 0.59 | -0.17 | -0.20 | 0.03 |
| | | | 0.01 | 4.7E-03 | 1.9E-03 | 6.4E-03 | 3.3E-02 | 7.0E-04 | 8.9E-04 | 2.0E-03 | 6.1E-03 | 3.4E-04 | 3.5E-04 | 2.4E-04 | 9.2E-05 | 0.64 | 0.23 | 0.20 | -0.11 | 0.89 | 0.64 | 0.45 | 0.05 | 0.93 | 0.84 | 0.65 | 0.13 | -0.35 | 0.03 | -0.15 | -0.52 | 0.20 | -0.14 | -0.54 | -0.31 | 0.25 | -0.19 | -0.07 | -0.04 |
| | | | 0.05 | 4.6E-03 | 1.3E-03 | 4.3E-03 | 2.4E-02 | 3.9E-04 | 6.7E-04 | 1.6E-03 | 5.6E-03 | 1.7E-04 | 2.3E-04 | 2.1E-04 | 1.3E-04 | 0.65 | 0.19 | 0.22 | -0.08 | 0.83 | 0.61 | 0.48 | 0.08 | 0.90 | 0.82 | 0.69 | 0.02 | -0.37 | 0.04 | -0.13 | -0.47 | 0.21 | -0.12 | -0.50 | -0.28 | 0.56 | -0.02 | -0.02 | -0.17 |
| | | | 0.1 | 4.6E-03 | 9.6E-04 | 3.4E-03 | 2.0E-02 | 3.5E-04 | 5.1E-04 | 1.3E-03 | 5.0E-03 | 1.1E-04 | 1.8E-04 | 2.0E-04 | 1.4E-04 | 0.68 | 0.15 | 0.19 | -0.06 | 0.81 | 0.58 | 0.45 | 0.07 | 0.88 | 0.83 | 0.71 | 0.04 | -0.35 | 0.05 | -0.14 | -0.44 | 0.24 | -0.07 | -0.37 | -0.33 | 0.69 | 0.05 | -0.04 | -0.14 |
| | | | 0.5 | 2.5E-03 | 2.8E-04 | 1.0E-03 | 6.3E-03 | 1.6E-04 | 1.7E-04 | 4.1E-04 | 2.0E-03 | 1.7E-05 | 5.6E-05 | 7.0E-05 | 7.8E-05 | 0.60 | 0.05 | 0.11 | -0.14 | 0.81 | 0.53 | 0.46 | 0.05 | 0.76 | 0.77 | 0.72 | 0.15 | -0.27 | 0.01 | -0.08 | -0.36 | 0.53 | 0.06 | -0.35 | -0.51 | 0.62 | 0.23 | 0.07 | -0.04 |
| | | | 0.001, 0.05 | 2.7E-03 | 9.1E-04 | 3.2E-03 | 1.5E-02 | 6.3E-05 | 3.1E-04 | 6.8E-04 | 1.6E-03 | 1.2E-05 | 6.5E-05 | 5.5E-05 | 4.9E-07 | 0.55 | 0.03 | 0.16 | -0.09 | 0.54 | 0.46 | 0.36 | -0.06 | 0.66 | 0.70 | 0.58 | 0.08 | -0.23 | -0.02 | 0.03 | -0.31 | 0.38 | 0.03 | -0.25 | -0.24 | 0.61 | 0.17 | 0.21 | 0.08 |
| | | | 0.001, 0.5 | 8.8E-04 | 1.5E-04 | 5.9E-04 | 2.5E-03 | 1.6E-06 | 4.3E-05 | 9.4E-05 | 1.6E-04 | 2.5E-07 | 7.4E-06 | 6.3E-06 | 0.0E+00 | 0.37 | -0.05 | 0.03 | -0.09 | -0.05 | -0.04 | 0.06 | -0.14 | 0.08 | 0.24 | 0.27 | NA | 0.09 | -0.06 | 0.00 | -0.14 | -0.05 | -0.10 | -0.05 | -0.19 | 0.08 | 0.16 | 0.24 | NA |
| average source-target distance - min. STS angle | simpler models not checked | % (-0.3,1.3) | 0.001 | 1.2E-01 | 8.0E-03 | 2.6E-02 | 1.2E-01 | 1.4E-01 | 1.6E-02 | 5.1E-02 | 1.8E-01 | 1.4E-01 | 2.6E-02 | 7.1E-02 | 1.6E-01 | 0.96 | 0.22 | 0.19 | 0.30 | 0.99 | 0.77 | 0.73 | 0.77 | 1.00 | 0.87 | 0.86 | 0.87 | -0.64 | -0.14 | -0.08 | -0.06 | -0.23 | 0.05 | -0.02 | -0.10 | -0.09 | 0.00 | -0.03 | -0.12 |
| | | | 0.01 | 1.1E-01 | 5.5E-03 | 1.8E-02 | 9.0E-02 | 1.3E-01 | 1.2E-02 | 3.7E-02 | 1.4E-01 | 1.3E-01 | 1.9E-02 | 5.4E-02 | 1.4E-01 | 0.96 | 0.18 | 0.18 | 0.28 | 1.00 | 0.75 | 0.73 | 0.76 | 1.00 | 0.86 | 0.86 | 0.87 | -0.62 | -0.13 | -0.07 | -0.06 | -0.24 | 0.06 | 0.03 | -0.09 | -0.07 | 0.01 | -0.02 | -0.12 |
| | | | 0.05 | 9.9E-02 | 3.6E-03 | 1.2E-02 | 6.1E-02 | 1.2E-01 | 7.8E-03 | 2.6E-02 | 1.1E-01 | 1.1E-01 | 1.3E-02 | 3.9E-02 | 1.2E-01 | 0.96 | 0.15 | 0.13 | 0.26 | 1.00 | 0.74 | 0.72 | 0.75 | 1.00 | 0.85 | 0.85 | 0.87 | -0.63 | -0.11 | -0.10 | -0.05 | -0.27 | 0.08 | 0.03 | -0.09 | -0.10 | 0.07 | 0.01 | -0.12 |
| | | | 0.1 | 8.8E-02 | 2.7E-03 | 9.3E-03 | 4.8E-02 | 1.0E-01 | 6.0E-03 | 2.0E-02 | 9.1E-02 | 1.0E-01 | 1.0E-02 | 3.1E-02 | 1.0E-01 | 0.96 | 0.11 | 0.12 | 0.25 | 1.00 | 0.72 | 0.71 | 0.75 | 1.00 | 0.83 | 0.85 | 0.87 | -0.65 | -0.11 | -0.10 | -0.06 | -0.29 | 0.08 | 0.04 | -0.08 | -0.12 | 0.06 | 0.01 | -0.14 |
| | | | 0.5 | 3.9E-02 | 7.8E-04 | 2.9E-03 | 1.4E-02 | 4.9E-02 | 1.9E-03 | 6.1E-03 | 3.1E-02 | 4.9E-02 | 3.1E-03 | 9.3E-03 | 4.3E-02 | 0.93 | 0.08 | 0.14 | 0.19 | 0.99 | 0.70 | 0.69 | 0.72 | 0.99 | 0.81 | 0.83 | 0.85 | -0.55 | -0.07 | -0.08 | -0.07 | -0.14 | 0.23 | 0.05 | -0.05 | -0.21 | 0.10 | 0.00 | -0.14 |
| | | % (0,1) | 0.001 | 6.4E-02 | 4.8E-03 | 1.6E-02 | 8.0E-02 | 4.2E-02 | 6.1E-03 | 1.6E-02 | 5.5E-02 | 2.6E-02 | 5.9E-03 | 9.7E-03 | 2.2E-02 | 0.90 | 0.20 | 0.13 | 0.08 | 0.98 | 0.72 | 0.62 | 0.49 | 0.98 | 0.82 | 0.79 | 0.65 | -0.49 | -0.06 | -0.06 | -0.12 | -0.17 | 0.12 | -0.09 | -0.15 | -0.03 | 0.05 | -0.12 | -0.12 |
| | | | 0.01 | 5.9E-02 | 3.3E-03 | 1.1E-02 | 5.8E-02 | 3.8E-02 | 4.3E-03 | 1.2E-02 | 4.5E-02 | 2.2E-02 | 4.1E-03 | 7.6E-03 | 1.9E-02 | 0.90 | 0.17 | 0.11 | 0.06 | 0.98 | 0.69 | 0.62 | 0.48 | 0.98 | 0.80 | 0.79 | 0.65 | -0.46 | -0.06 | -0.08 | -0.12 | -0.20 | 0.13 | -0.03 | -0.15 | 0.01 | 0.04 | -0.10 | -0.08 |
| | | | 0.05 | 5.1E-02 | 2.2E-03 | 7.2E-03 | 4.0E-02 | 3.3E-02 | 2.8E-03 | 8.2E-03 | 3.5E-02 | 1.9E-02 | 2.8E-03 | 5.6E-03 | 1.6E-02 | 0.90 | 0.17 | 0.09 | 0.05 | 0.98 | 0.66 | 0.60 | 0.48 | 0.98 | 0.80 | 0.78 | 0.65 | -0.47 | -0.02 | -0.10 | -0.11 | -0.24 | 0.12 | -0.02 | -0.13 | 0.01 | 0.12 | -0.06 | -0.09 |
| | | | 0.1 | 4.6E-02 | 1.6E-03 | 5.6E-03 | 3.1E-02 | 3.0E-02 | 2.2E-03 | 6.5E-03 | 2.9E-02 | 1.7E-02 | 2.1E-03 | 4.5E-03 | 1.4E-02 | 0.90 | 0.11 | 0.09 | 0.05 | 0.98 | 0.65 | 0.59 | 0.48 | 0.98 | 0.80 | 0.76 | 0.65 | -0.49 | -0.03 | -0.10 | -0.12 | -0.22 | 0.12 | -0.01 | -0.13 | 0.03 | 0.13 | -0.08 | -0.09 |
| | | | 0.5 | 2.1E-02 | 4.3E-04 | 1.7E-03 | 8.9E-03 | 1.4E-02 | 6.8E-04 | 1.9E-03 | 1.0E-02 | 7.8E-03 | 6.2E-04 | 1.4E-03 | 5.8E-03 | 0.85 | 0.01 | 0.09 | 0.02 | 0.98 | 0.58 | 0.59 | 0.43 | 0.98 | 0.75 | 0.76 | 0.63 | -0.49 | -0.06 | -0.03 | -0.12 | -0.11 | 0.19 | 0.05 | -0.14 | -0.03 | 0.15 | 0.00 | -0.08 |
| | | %+/-2SE (0,1) | 0.001 | 1.1E-02 | 3.2E-03 | 1.1E-02 | 5.7E-02 | 2.6E-03 | 2.2E-03 | 4.6E-03 | 1.5E-02 | 1.3E-03 | 1.1E-03 | 8.3E-04 | 5.2E-04 | 0.57 | 0.20 | 0.11 | 0.01 | 0.90 | 0.61 | 0.46 | 0.17 | 0.94 | 0.79 | 0.67 | 0.25 | -0.05 | 0.02 | -0.02 | -0.14 | 0.22 | 0.14 | -0.08 | -0.12 | 0.46 | 0.10 | 0.07 | 0.12 |
| | | | 0.01 | 9.4E-03 | 2.1E-03 | 7.5E-03 | 4.1E-02 | 1.5E-03 | 1.4E-03 | 3.2E-03 | 1.2E-02 | 6.1E-04 | 6.6E-04 | 5.7E-04 | 4.5E-04 | 0.53 | 0.16 | 0.10 | -0.01 | 0.85 | 0.56 | 0.43 | 0.16 | 0.90 | 0.77 | 0.67 | 0.25 | -0.07 | 0.04 | -0.03 | -0.13 | 0.21 | 0.15 | -0.07 | -0.13 | 0.40 | 0.17 | 0.06 | 0.12 |
| | | | 0.05 | 7.9E-03 | 1.4E-03 | 4.8E-03 | 2.8E-02 | 8.7E-04 | 8.8E-04 | 2.1E-03 | 8.7E-03 | 2.7E-04 | 3.7E-04 | 3.9E-04 | 3.7E-04 | 0.53 | 0.13 | 0.08 | -0.01 | 0.83 | 0.52 | 0.42 | 0.16 | 0.86 | 0.76 | 0.65 | 0.24 | -0.10 | 0.05 | -0.05 | -0.11 | 0.28 | 0.16 | -0.06 | -0.15 | 0.52 | 0.25 | 0.08 | 0.12 |
| | | | 0.1 | 7.0E-03 | 1.1E-03 | 3.7E-03 | 2.2E-02 | 6.9E-04 | 6.5E-04 | 1.7E-03 | 7.0E-03 | 1.7E-04 | 2.7E-04 | 3.1E-04 | 3.1E-04 | 0.52 | 0.05 | 0.07 | -0.01 | 0.81 | 0.49 | 0.40 | 0.14 | 0.85 | 0.74 | 0.64 | 0.24 | -0.10 | -0.01 | -0.04 | -0.11 | 0.27 | 0.14 | 0.01 | -0.15 | 0.59 | 0.26 | 0.10 | 0.12 |
| | | | 0.5 | 2.8E-03 | 2.9E-04 | 1.0E-03 | 6.4E-03 | 2.2E-04 | 1.7E-04 | 4.4E-04 | 2.2E-03 | 2.5E-05 | 6.1E-05 | 7.8E-05 | 9.4E-05 | 0.45 | -0.06 | -0.01 | -0.10 | 0.74 | 0.38 | 0.36 | 0.07 | 0.70 | 0.64 | 0.57 | 0.18 | -0.09 | -0.08 | -0.07 | -0.17 | 0.52 | 0.17 | 0.02 | -0.21 | 0.61 | 0.44 | 0.22 | 0.07 |
| | "trailing" simpler models rejected | % (-0.3,1.3) | 0.001 | 1.4E-02 | 4.0E-03 | 1.3E-02 | 5.2E-02 | 3.2E-03 | 3.3E-03 | 7.6E-03 | 1.9E-02 | 1.2E-03 | 2.0E-03 | 2.8E-03 | 1.1E-03 | 0.47 | 0.14 | 0.09 | 0.27 | 0.87 | 0.63 | 0.64 | 0.68 | 0.92 | 0.80 | 0.85 | 0.81 | 0.03 | -0.06 | -0.08 | -0.02 | 0.29 | 0.11 | -0.02 | 0.02 | 0.36 | -0.04 | 0.02 | 0.31 |
| | | | 0.01 | 1.4E-02 | 3.0E-03 | 9.8E-03 | 4.4E-02 | 2.7E-03 | 2.5E-03 | 6.6E-03 | 2.0E-02 | 8.6E-04 | 1.7E-03 | 2.7E-03 | 1.7E-03 | 0.49 | 0.13 | 0.09 | 0.24 | 0.83 | 0.61 | 0.61 | 0.68 | 0.91 | 0.77 | 0.85 | 0.83 | 0.04 | -0.04 | -0.07 | -0.03 | 0.35 | 0.12 | -0.01 | -0.05 | 0.31 | -0.01 | -0.03 | 0.35 |
| | | | 0.05 | 1.5E-02 | 2.2E-03 | 7.0E-03 | 3.4E-02 | 2.9E-03 | 2.0E-03 | 5.5E-03 | 2.0E-02 | 7.4E-04 | 1.5E-03 | 2.7E-03 | 2.5E-03 | 0.57 | 0.10 | 0.04 | 0.23 | 0.84 | 0.60 | 0.62 | 0.67 | 0.89 | 0.78 | 0.84 | 0.81 | -0.07 | -0.07 | -0.12 | -0.03 | 0.27 | 0.17 | 0.01 | -0.10 | 0.48 | 0.09 | 0.02 | 0.24 |
| | | | 0.1 | 1.5E-02 | 1.6E-03 | 5.4E-03 | 2.8E-02 | 3.1E-03 | 1.7E-03 | 4.7E-03 | 1.8E-02 | 7.6E-04 | 1.3E-03 | 2.6E-03 | 2.9E-03 | 0.59 | 0.07 | 0.03 | 0.22 | 0.85 | 0.58 | 0.60 | 0.67 | 0.86 | 0.78 | 0.83 | 0.80 | -0.10 | -0.05 | -0.10 | -0.03 | 0.27 | 0.17 | 0.04 | -0.08 | 0.47 | 0.14 | 0.03 | 0.08 |
| | | | 0.5 | 1.0E-02 | 5.4E-04 | 1.8E-03 | 9.5E-03 | 3.8E-03 | 6.8E-04 | 1.8E-03 | 8.9E-03 | 1.1E-03 | 6.2E-04 | 1.5E-03 | 2.9E-03 | 0.69 | 0.04 | 0.05 | 0.15 | 0.91 | 0.55 | 0.56 | 0.67 | 0.91 | 0.76 | 0.77 | 0.82 | -0.22 | -0.06 | -0.07 | -0.05 | 0.29 | 0.20 | 0.03 | -0.08 | 0.33 | 0.24 | 0.00 | -0.02 |
| | | | 0.001, 0.05 | 8.6E-03 | 1.4E-03 | 4.5E-03 | 1.7E-02 | 9.0E-04 | 7.4E-04 | 1.5E-03 | 3.2E-03 | 6.9E-05 | 2.8E-04 | 3.2E-04 | 2.2E-05 | 0.32 | -0.02 | 0.00 | 0.20 | 0.56 | 0.41 | 0.50 | 0.56 | 0.55 | 0.69 | 0.83 | 0.57 | 0.08 | -0.09 | -0.10 | -0.01 | 0.37 | 0.15 | 0.02 | 0.08 | 0.42 | 0.10 | 0.38 | 0.52 |
| | | | 0.001, 0.5 | 3.0E-03 | 2.2E-04 | 7.7E-04 | 2.6E-03 | 2.3E-04 | 1.2E-04 | 1.8E-04 | 2.2E-04 | 1.6E-05 | 3.4E-05 | 2.4E-05 | 2.5E-07 | 0.24 | -0.14 | -0.09 | 0.00 | 0.41 | 0.20 | 0.30 | 0.34 | 0.33 | 0.52 | 0.57 | 0.16 | 0.05 | -0.16 | -0.13 | -0.07 | 0.29 | 0.10 | 0.06 | 0.19 | 0.28 | 0.31 | 0.48 | 0.16 |
| | | % (0,1) | 0.001 | 8.4E-03 | 2.7E-03 | 9.0E-03 | 4.2E-02 | 1.9E-03 | 1.5E-03 | 2.9E-03 | 6.5E-03 | 9.5E-04 | 6.9E-04 | 4.1E-04 | 1.5E-04 | 0.44 | 0.14 | 0.06 | 0.00 | 0.89 | 0.56 | 0.43 | 0.21 | 0.92 | 0.77 | 0.66 | 0.36 | 0.00 | 0.00 | -0.05 | -0.15 | 0.37 | 0.10 | -0.07 | -0.05 | 0.42 | 0.04 | 0.16 | 0.15 |
| | | | 0.01 | 7.9E-03 | 2.0E-03 | 6.7E-03 | 3.4E-02 | 1.3E-03 | 1.1E-03 | 2.5E-03 | 7.3E-03 | 5.9E-04 | 5.4E-04 | 4.2E-04 | 2.6E-04 | 0.44 | 0.11 | 0.07 | -0.02 | 0.83 | 0.53 | 0.42 | 0.21 | 0.91 | 0.74 | 0.69 | 0.43 | 0.02 | 0.01 | -0.03 | -0.15 | 0.37 | 0.14 | -0.07 | -0.12 | 0.34 | 0.08 | 0.07 | 0.25 |
| | | | 0.05 | 8.2E-03 | 1.4E-03 | 4.7E-03 | 2.6E-02 | 1.0E-03 | 8.8E-04 | 2.1E-03 | 7.2E-03 | 3.5E-04 | 4.0E-04 | 4.3E-04 | 4.0E-04 | 0.49 | 0.11 | 0.04 | -0.02 | 0.80 | 0.50 | 0.44 | 0.23 | 0.88 | 0.74 | 0.67 | 0.42 | -0.12 | 0.03 | -0.07 | -0.14 | 0.35 | 0.14 | -0.04 | -0.12 | 0.43 | 0.22 | 0.03 | 0.15 |
| | | | 0.1 | 8.2E-03 | 1.0E-03 | 3.7E-03 | 2.1E-02 | 1.0E-03 | 7.5E-04 | 1.8E-03 | 6.6E-03 | 2.8E-04 | 3.3E-04 | 4.3E-04 | 4.6E-04 | 0.51 | 0.04 | 0.02 | -0.01 | 0.79 | 0.51 | 0.41 | 0.22 | 0.86 | 0.73 | 0.70 | 0.39 | -0.07 | -0.01 | -0.07 | -0.13 | 0.28 | 0.15 | 0.00 | -0.13 | 0.57 | 0.14 | 0.04 | 0.09 |
| | | | 0.5 | 5.7E-03 | 3.3E-04 | 1.2E-03 | 6.9E-03 | 1.2E-03 | 2.9E-04 | 6.7E-04 | 3.2E-03 | 2.3E-04 | 1.5E-04 | 2.5E-04 | 4.2E-04 | 0.57 | -0.03 | 0.01 | -0.05 | 0.86 | 0.46 | 0.41 | 0.20 | 0.82 | 0.67 | 0.60 | 0.44 | -0.23 | -0.06 | -0.05 | -0.14 | 0.26 | 0.17 | 0.02 | -0.19 | 0.42 | 0.32 | 0.03 | 0.07 |
| | | | 0.001, 0.05 | 4.8E-03 | 9.8E-04 | 3.3E-03 | 1.5E-02 | 2.2E-04 | 3.6E-04 | 7.7E-04 | 1.6E-03 | 2.2E-05 | 9.0E-05 | 7.4E-05 | 1.7E-06 | 0.29 | -0.06 | -0.04 | -0.09 | 0.52 | 0.30 | 0.33 | 0.02 | 0.57 | 0.65 | 0.60 | 0.09 | 0.03 | -0.09 | -0.09 | -0.16 | 0.38 | 0.08 | -0.01 | -0.11 | 0.47 | 0.30 | 0.36 | 0.09 |
| | | | 0.001, 0.5 | 1.6E-03 | 1.7E-04 | 6.1E-04 | 2.5E-03 | 5.5E-05 | 5.6E-05 | 1.0E-04 | 1.6E-04 | 3.0E-06 | 1.1E-05 | 7.2E-06 | 0.0E+00 | 0.20 | -0.17 | -0.16 | -0.14 | 0.21 | 0.00 | 0.10 | -0.07 | 0.19 | 0.34 | 0.29 | NA | 0.02 | -0.17 | -0.18 | -0.18 | 0.12 | -0.03 | 0.00 | -0.12 | 0.19 | 0.27 | 0.26 | NA |
| | | %+/-2SE (0,1) | 0.001 | 5.4E-03 | 2.5E-03 | 8.6E-03 | 4.1E-02 | 1.3E-03 | 1.3E-03 | 2.5E-03 | 5.7E-03 | 5.9E-04 | 5.1E-04 | 2.8E-04 | 5.4E-05 | 0.44 | 0.14 | 0.05 | 0.01 | 0.88 | 0.56 | 0.41 | 0.11 | 0.92 | 0.79 | 0.65 | 0.19 | 0.06 | 0.01 | -0.04 | -0.13 | 0.38 | 0.13 | -0.06 | -0.06 | 0.66 | 0.14 | 0.27 | 0.14 |
| | | | 0.01 | 4.7E-03 | 1.9E-03 | 6.4E-03 | 3.3E-02 | 7.0E-04 | 8.9E-04 | 2.0E-03 | 6.1E-03 | 3.4E-04 | 3.5E-04 | 2.4E-04 | 9.2E-05 | 0.40 | 0.12 | 0.08 | -0.03 | 0.81 | 0.53 | 0.40 | 0.12 | 0.90 | 0.76 | 0.63 | 0.20 | 0.07 | 0.02 | -0.02 | -0.13 | 0.37 | 0.16 | -0.05 | -0.09 | 0.54 | 0.15 | 0.17 | 0.13 |
| | | | 0.05 | 4.6E-03 | 1.3E-03 | 4.3E-03 | 2.4E-02 | 3.9E-04 | 6.7E-04 | 1.6E-03 | 5.6E-03 | 1.7E-04 | 2.3E-04 | 2.1E-04 | 1.3E-04 | 0.40 | 0.11 | 0.04 | -0.01 | 0.75 | 0.50 | 0.40 | 0.15 | 0.85 | 0.76 | 0.64 | 0.16 | -0.04 | 0.04 | -0.06 | -0.11 | 0.42 | 0.16 | -0.05 | -0.09 | 0.59 | 0.31 | 0.19 | 0.07 |
| | | | 0.1 | 4.6E-03 | 9.6E-04 | 3.4E-03 | 2.0E-02 | 3.5E-04 | 5.1E-04 | 1.3E-03 | 5.0E-03 | 1.1E-04 | 1.8E-04 | 2.0E-04 | 1.4E-04 | 0.42 | 0.04 | 0.01 | -0.01 | 0.73 | 0.47 | 0.37 | 0.14 | 0.82 | 0.75 | 0.65 | 0.12 | -0.06 | -0.01 | -0.08 | -0.11 | 0.35 | 0.14 | 0.00 | -0.11 | 0.64 | 0.31 | 0.19 | 0.02 |
| | | | 0.5 | 2.5E-03 | 2.8E-04 | 1.0E-03 | 6.3E-03 | 1.6E-04 | 1.7E-04 | 4.1E-04 | 2.0E-03 | 1.7E-05 | 5.6E-05 | 7.0E-05 | 7.8E-05 | 0.40 | -0.05 | -0.02 | -0.11 | 0.68 | 0.39 | 0.35 | 0.05 | 0.65 | 0.58 | 0.57 | 0.18 | -0.10 | -0.07 | -0.08 | -0.18 | 0.51 | 0.17 | 0.02 | -0.20 | 0.56 | 0.39 | 0.25 | 0.08 |
| | | | 0.001, 0.05 | 2.7E-03 | 9.1E-04 | 3.2E-03 | 1.5E-02 | 6.3E-05 | 3.1E-04 | 6.8E-04 | 1.6E-03 | 1.2E-05 | 6.5E-05 | 5.5E-05 | 4.9E-07 | 0.24 | -0.06 | -0.03 | -0.07 | 0.42 | 0.31 | 0.31 | 0.01 | 0.63 | 0.66 | 0.55 | 0.09 | 0.03 | -0.09 | -0.09 | -0.14 | 0.35 | 0.09 | -0.02 | -0.11 | 0.61 | 0.35 | 0.36 | 0.09 |
| | | | 0.001, 0.5 | 8.8E-04 | 1.5E-04 | 5.9E-04 | 2.5E-03 | 1.6E-06 | 4.3E-05 | 9.4E-05 | 1.6E-04 | 2.5E-07 | 7.4E-06 | 6.3E-06 | 0.0E+00 | 0.10 | -0.18 | -0.15 | -0.14 | -0.07 | -0.04 | 0.06 | -0.09 | 0.11 | 0.31 | 0.25 | NA | -0.02 | -0.19 | -0.17 | -0.18 | -0.07 | -0.07 | -0.03 | -0.14 | 0.11 | 0.27 | 0.23 | NA |

*p*-value > 0.05/1728 (Bonferroni correction)

**|Spearman correlation coefficient| >= 0.5**

Overall *qpAdm* positive rates and Spearman's correlation coefficients (ρ) in the spaces of model optimality metrics. Correlation was assessed between total model count and fitting model count or *qpAdm* positive rate. Results are stratified by model complexity and landscape type (in columns), and by model feasibility criteria (in rows). The composite feasibility criteria visualized in **Fig. 6** and other figures are highlighted in green. Optimal combinations of model complexity and landscape sparsity are also highlighted in green in the table header. Non-significant ρ values (with Bonferroni correction) are underlined, and absolute ρ values ≥0.5 are shown in bold.

**Table S12.**

| variable 1: | model complexity | "right" demes | max. \|estimated admixture fractions - equal fractions\| | | | | | p-value | | | | |
|---|---|---|---|---|---|---|---|---|---|---|---|---|
| total number of systematic models: | | | 49,309,831 | 20,452,113 | 11,729,213 | 10,978,092 | 2,331,106 | 49,309,831 | 20,452,113 | 11,729,213 | 10,978,092 | 2,331,106 |
| | | | p >= 1e-55 | p >= 1e-5 | max. \|EAF–EF\| <= 0.5 | p >= 1e-55; max. \|EAF–EF\| <= 0.5 | p >= 1e-5; max. \|EAF–EF\| <= 0.5 | p >= 1e-55 | p >= 1e-5 | max. \|EAF–EF\| <= 0.5 | p >= 1e-55; max. \|EAF–EF\| <= 0.5 | p >= 1e-5; max. \|EAF–EF\| <= 0.5 |
| minimal source-target-source angle | 2-way | 1st-2nd circles | **-0.81** | -0.36 | -0.24 | -0.23 | 0.03 | -0.40 | -0.30 | 0.15 | 0.05 | 0.13 |
| | | 2nd-3rd circles | **-0.81** | **-0.53** | -0.27 | -0.22 | -0.10 | -0.30 | -0.23 | 0.30 | 0.12 | 0.25 |
| | | 3rd circle | **-0.82** | **-0.71** | -0.26 | -0.27 | -0.27 | -0.32 | -0.18 | 0.34 | 0.05 | 0.17 |
| | | 4th circle | **-0.84** | **-0.82** | -0.28 | -0.32 | -0.29 | -0.17 | -0.12 | 0.26 | 0.10 | 0.12 |
| | 3-way | 1st-2nd circles | **-0.70** | -0.35 | -0.21 | -0.22 | -0.28 | **-0.62** | -0.31 | -0.15 | -0.15 | -0.04 |
| | | 2nd-3rd circles | **-0.74** | -0.46 | -0.34 | -0.35 | -0.38 | -0.47 | -0.23 | 0.03 | 0.02 | -0.03 |
| | | 3rd circle | **-0.76** | **-0.58** | -0.38 | -0.40 | -0.40 | -0.39 | -0.18 | 0.19 | 0.13 | -0.04 |
| | | 4th circle | **-0.73** | **-0.63** | -0.37 | -0.40 | -0.36 | -0.28 | -0.10 | 0.11 | 0.02 | 0.00 |
| | 4-way | 1st-2nd circles | -0.43 | -0.23 | -0.17 | -0.17 | -0.20 | **-0.59** | -0.30 | -0.35 | -0.34 | -0.14 |
| | | 2nd-3rd circles | **-0.52** | -0.32 | -0.32 | -0.32 | -0.26 | **-0.50** | -0.23 | -0.21 | -0.21 | -0.08 |
| | | 3rd circle | **-0.58** | -0.45 | -0.38 | -0.38 | -0.36 | -0.44 | -0.18 | -0.05 | -0.05 | -0.07 |
| | | 4th circle | **-0.54** | -0.44 | -0.37 | -0.38 | -0.32 | -0.32 | -0.10 | -0.11 | -0.11 | -0.07 |
| maximal source-target distance | 2-way | 1st-2nd circles | 0.13 | 0.31 | 0.03 | 0.05 | 0.31 | -0.02 | 0.15 | -0.11 | -0.15 | -0.29 |
| | | 2nd-3rd circles | 0.11 | 0.49 | 0.02 | 0.01 | 0.42 | -0.08 | 0.12 | -0.15 | -0.18 | -0.17 |
| | | 3rd circle | 0.17 | **0.52** | -0.06 | 0.02 | **0.56** | -0.09 | 0.05 | -0.28 | -0.23 | -0.13 |
| | | 4th circle | 0.27 | 0.42 | 0.09 | 0.40 | **0.74** | -0.14 | -0.12 | -0.44 | -0.22 | -0.20 |
| | 3-way | 1st-2nd circles | 0.14 | 0.21 | 0.03 | 0.04 | 0.04 | 0.05 | 0.12 | -0.12 | -0.12 | -0.14 |
| | | 2nd-3rd circles | 0.11 | 0.31 | 0.01 | 0.01 | 0.15 | -0.01 | 0.11 | -0.13 | -0.14 | -0.14 |
| | | 3rd circle | 0.10 | 0.31 | -0.07 | -0.07 | 0.14 | -0.08 | 0.02 | -0.20 | -0.19 | -0.10 |
| | | 4th circle | 0.12 | 0.22 | 0.00 | 0.09 | 0.26 | -0.15 | -0.10 | -0.33 | -0.24 | -0.12 |
| | 4-way | 1st-2nd circles | 0.15 | 0.17 | 0.03 | 0.03 | 0.03 | 0.10 | 0.11 | -0.08 | -0.08 | -0.07 |
| | | 2nd-3rd circles | 0.12 | 0.22 | 0.01 | 0.01 | 0.09 | 0.04 | 0.10 | -0.12 | -0.12 | -0.10 |
| | | 3rd circle | 0.11 | 0.21 | -0.05 | -0.05 | 0.04 | -0.04 | 0.02 | -0.15 | -0.14 | -0.08 |
| | | 4th circle | 0.09 | 0.16 | -0.01 | 0.01 | 0.11 | -0.13 | -0.08 | -0.23 | -0.21 | -0.11 |
| average source-target distance | 2-way | 1st-2nd circles | 0.18 | 0.46 | 0.01 | 0.01 | 0.35 | -0.08 | 0.23 | -0.29 | -0.30 | -0.31 |
| | | 2nd-3rd circles | 0.13 | **0.70** | -0.07 | -0.15 | 0.40 | -0.18 | 0.19 | -0.38 | -0.39 | -0.18 |
| | | 3rd circle | 0.20 | **0.77** | -0.25 | -0.30 | **0.53** | -0.23 | 0.11 | **-0.57** | **-0.50** | -0.15 |
| | | 4th circle | 0.32 | **0.58** | -0.17 | 0.08 | **0.71** | -0.31 | -0.09 | **-0.77** | **-0.50** | -0.22 |
| | 3-way | 1st-2nd circles | 0.25 | 0.46 | 0.01 | 0.02 | 0.04 | -0.01 | 0.23 | -0.40 | -0.39 | -0.27 |
| | | 2nd-3rd circles | 0.18 | **0.62** | -0.10 | -0.11 | 0.20 | -0.15 | 0.19 | -0.47 | -0.46 | -0.24 |
| | | 3rd circle | 0.15 | **0.63** | -0.28 | -0.33 | 0.12 | -0.28 | 0.04 | **-0.58** | **-0.54** | -0.23 |
| | | 4th circle | 0.23 | **0.50** | -0.26 | -0.18 | 0.24 | -0.39 | -0.13 | **-0.75** | **-0.63** | -0.29 |
| | 4-way | 1st-2nd circles | 0.36 | 0.45 | -0.01 | -0.01 | 0.02 | 0.10 | 0.21 | -0.43 | -0.42 | -0.24 |
| | | 2nd-3rd circles | 0.28 | **0.54** | -0.11 | -0.10 | 0.17 | -0.08 | 0.16 | **-0.55** | **-0.55** | -0.28 |
| | | 3rd circle | 0.20 | **0.51** | -0.28 | -0.28 | 0.00 | -0.28 | -0.03 | **-0.58** | **-0.57** | -0.31 |
| | | 4th circle | 0.20 | 0.42 | -0.26 | -0.23 | 0.12 | -0.45 | -0.19 | **-0.74** | **-0.69** | -0.35 |
| distance to the ideal symmetric model (in the space of max. distance vs. min. angle) | 2-way | 1st-2nd circles | **0.81** | 0.40 | 0.23 | 0.23 | 0.09 | 0.38 | 0.33 | -0.17 | -0.12 | -0.25 |
| | | 2nd-3rd circles | **0.80** | **0.61** | 0.26 | 0.21 | 0.27 | 0.27 | 0.25 | -0.33 | -0.18 | -0.29 |
| | | 3rd circle | **0.82** | **0.79** | 0.21 | 0.24 | 0.47 | 0.27 | 0.18 | -0.42 | -0.16 | -0.21 |
| | | 4th circle | **0.86** | **0.85** | 0.29 | 0.47 | **0.59** | 0.13 | 0.08 | -0.40 | -0.19 | -0.17 |
| | 3-way | 1st-2nd circles | **0.71** | 0.38 | 0.22 | 0.22 | 0.29 | **0.62** | 0.33 | 0.11 | 0.11 | 0.01 |
| | | 2nd-3rd circles | **0.74** | **0.50** | 0.33 | 0.34 | 0.40 | 0.46 | 0.25 | -0.06 | -0.05 | -0.01 |
| | | 3rd circle | **0.76** | **0.62** | 0.35 | 0.36 | 0.43 | 0.37 | 0.18 | -0.23 | -0.18 | 0.00 |
| | | 4th circle | **0.72** | **0.65** | 0.36 | 0.41 | 0.42 | 0.25 | 0.08 | -0.18 | -0.09 | -0.03 |
| | 4-way | 1st-2nd circles | 0.44 | 0.25 | 0.17 | 0.17 | 0.20 | **0.59** | 0.32 | 0.32 | 0.31 | 0.12 |
| | | 2nd-3rd circles | **0.53** | 0.35 | 0.32 | 0.31 | 0.27 | 0.50 | 0.25 | 0.18 | 0.18 | 0.06 |
| | | 3rd circle | **0.58** | 0.47 | 0.36 | 0.36 | 0.36 | 0.42 | 0.18 | 0.02 | 0.02 | 0.06 |
| | | 4th circle | **0.54** | 0.46 | 0.36 | 0.37 | 0.33 | 0.29 | 0.08 | 0.06 | 0.07 | 0.05 |
| distance to the ideal symmetric model (in the space of avg. distance vs. min. angle) | 2-way | 1st-2nd circles | **0.82** | 0.44 | 0.24 | 0.23 | 0.03 | 0.37 | 0.36 | -0.23 | -0.16 | -0.20 |
| | | 2nd-3rd circles | **0.80** | **0.65** | 0.24 | 0.17 | 0.16 | 0.25 | 0.28 | -0.40 | -0.25 | -0.30 |
| | | 3rd circle | **0.81** | **0.83** | 0.17 | 0.15 | 0.34 | 0.25 | 0.20 | -0.49 | -0.23 | -0.19 |
| | | 4th circle | **0.85** | **0.86** | 0.21 | 0.32 | 0.41 | 0.12 | 0.10 | -0.48 | -0.24 | -0.16 |
| | 3-way | 1st-2nd circles | **0.72** | 0.41 | 0.22 | 0.22 | 0.28 | **0.61** | 0.35 | 0.07 | 0.07 | 0.01 |
| | | 2nd-3rd circles | **0.75** | **0.53** | 0.32 | 0.33 | 0.40 | 0.45 | 0.26 | -0.11 | -0.10 | 0.00 |
| | | 3rd circle | **0.77** | **0.64** | 0.32 | 0.33 | 0.41 | 0.36 | 0.18 | -0.28 | -0.23 | 0.00 |
| | | 4th circle | **0.74** | **0.67** | 0.32 | 0.37 | 0.39 | 0.24 | 0.09 | -0.24 | -0.12 | -0.03 |
| | 4-way | 1st-2nd circles | 0.46 | 0.28 | 0.17 | 0.17 | 0.20 | **0.60** | 0.33 | 0.29 | 0.29 | 0.11 |
| | | 2nd-3rd circles | **0.55** | 0.38 | 0.31 | 0.31 | 0.27 | 0.49 | 0.25 | 0.14 | 0.14 | 0.06 |
| | | 3rd circle | **0.60** | 0.49 | 0.34 | 0.34 | 0.36 | 0.40 | 0.17 | -0.02 | -0.02 | 0.05 |
| | | 4th circle | **0.56** | 0.48 | 0.34 | 0.35 | 0.32 | 0.27 | 0.07 | 0.02 | 0.03 | 0.04 |

*p*-value > 0.05/600 (Bonferroni correction)
**|Spearman correlation coefficient| >= 0.5**

Spearman's correlation coefficient (ρ) between several model optimality metrics (listed on the left) and max|*EAF–EF*| or *p*-values in various sections of the respective two-dimensional space (in columns). Results are shown for systematic *qpAdm* experiments (the distal non-rotating protocol, all possible proxy sources tested for two targets in the center of the "$10^{-5}$ to $10^{-2}$" landscapes) stratified by model complexity and target-"right" distance: 16 "right" groups came either from the 1st and 2nd, or 2nd and 3rd, or 3rd, or 4th circle around the target. Non-significant ρ values (with Bonferroni correction) are underlined, and absolute ρ values ≥0.5 are shown in bold.

**Table S13.**

| | | | | | | | | | | | | | | | | | | | |
|---|---|---|---|---|---|---|---|---|---|---|---|---|---|---|---|---|---|---|---|
| true models: max. ST distance = 1; false models: max. ST distance > 1 | | | | | | | | | | | | | | | | | | | |
| distal non-rotating protocol; all demes at 100 gen. before "present" used as proxy sources; "right" demes at 300 gen. before "present" | | | | | | | | | | | | | | | | | | | |
| | | composite feasibility criterion: | | | *p*-value >=0.01, EAF within (0, 1) | | | *p*-value >=0.1, EAF within (0, 1) | | | *p*-value >=0.01 | | | *p*-value >=0.1 | | | EAF within (0, 1) | | |
| "right"-target distance | model complexity | number of models tested | number of true models | pre-study odds (R) | FPR (alpha) =1-TNR =FP/N | FNR (beta) =1-TPR =1-power =FN/P | FDR =FPR/(R-FNR*R+FPR) =FP/PP | FPR (alpha) =1-TNR =FP/N | FNR (beta) =1-TPR =1-power =FN/P | FDR =FPR/(R-FNR*R+FPR) =FP/PP | FPR (alpha) =1-TNR =FP/N | FNR (beta) =1-TPR =1-power =FN/P | FDR =FPR/(R-FNR*R+FPR) =FP/PP | FPR (alpha) =1-TNR =FP/N | FNR (beta) =1-TPR =1-power =FN/P | FDR =FPR/(R-FNR*R+FPR) =FP/PP | FPR (alpha) =1-TNR =FP/N | FNR (beta) =1-TPR =1-power =FN/P | FDR =FPR/(R-FNR*R+FPR) =FP/PP |
| right: 1–2nd c. | 2-way | 39,060 | 300 | 0.77% | 0.1% | 96.0% | 65.7% | 0.0% | 97.3% | 52.9% | 2.2% | 96.0% | 98.6% | 1.1% | 97.3% | 98.1% | 56.4% | 28.3% | 99.0% |
| right: 2–3rd c. | | 39,060 | 300 | 0.77% | 0.1% | 93.7% | 74.0% | 0.0% | 97.0% | 67.9% | 1.4% | 93.7% | 96.7% | 0.7% | 97.0% | 96.6% | 62.2% | 29.7% | 99.1% |
| right: 3rd c. | | 39,060 | 300 | 0.77% | 0.3% | 89.7% | 78.5% | 0.1% | 93.7% | 64.2% | 1.6% | 89.7% | 95.3% | 0.7% | 93.7% | 93.2% | 64.9% | 26.3% | 99.1% |
| right: 4th c. | | 39,060 | 300 | 0.77% | 0.3% | 83.0% | 67.1% | 0.1% | 91.3% | 49.0% | 1.2% | 82.7% | 89.9% | 0.4% | 91.3% | 85.0% | 64.1% | 27.0% | 99.1% |
| right: 1–2nd c. | 3-way | 794,220 | 400 | 0.050% | 0.2% | 83.3% | 96.6% | 0.1% | 91.3% | 95.5% | 11.8% | 70.8% | 99.9% | 6.3% | 87.3% | 99.9% | 20.5% | 49.5% | 99.9% |
| right: 2–3rd c. | | 794,220 | 400 | 0.050% | 0.5% | 75.0% | 97.5% | 0.2% | 86.8% | 96.7% | 7.6% | 59.5% | 99.7% | 3.7% | 78.3% | 99.7% | 30.0% | 52.8% | 99.9% |
| right: 3rd c. | | 794,220 | 400 | 0.050% | 1.0% | 73.0% | 98.7% | 0.4% | 80.5% | 97.7% | 7.8% | 49.8% | 99.7% | 3.7% | 68.8% | 99.6% | 36.6% | 52.3% | 99.9% |
| right: 4th c. | | 794,220 | 400 | 0.050% | 1.1% | 63.3% | 98.4% | 0.4% | 74.5% | 97.2% | 6.3% | 28.8% | 99.4% | 2.6% | 53.3% | 99.1% | 36.4% | 54.0% | 99.9% |
| right: 1–2nd c. | 4-way | 11,913,300 | 300 | 0.003% | 0.2% | 78.7% | 99.8% | 0.1% | 86.0% | 99.6% | 30.6% | 44.3% | 100.0% | 17.4% | 70.7% | 100.0% | 3.6% | 68.0% | 100.0% |
| right: 2–3rd c. | | 11,913,300 | 300 | 0.003% | 0.5% | 71.7% | 99.9% | 0.2% | 83.3% | 99.8% | 19.3% | 25.7% | 100.0% | 9.8% | 51.3% | 100.0% | 11.2% | 65.7% | 100.0% |
| right: 3rd c. | | 11,913,300 | 300 | 0.003% | 1.4% | 70.3% | 99.9% | 0.6% | 77.3% | 99.9% | 18.2% | 14.7% | 100.0% | 8.5% | 35.0% | 100.0% | 18.7% | 69.0% | 100.0% |
| right: 4th c. | | 11,913,300 | 300 | 0.003% | 1.4% | 71.0% | 99.9% | 0.6% | 73.0% | 99.9% | 14.0% | 6.0% | 100.0% | 5.9% | 17.3% | 100.0% | 19.1% | 70.0% | 100.0% |
| distal non-rotating protocol; proxy sources in the 1st and 3rd circles around the target, at 100 gen. before "present"; "right" demes at 300 gen. BP | | | | | | | | | | | | | | | | | | | |
| right: 3rd c. | 2-way | 10,800 | 1,350 | 14.3% | 0.4% | 89.5% | 22.0% | 0.1% | 95.0% | 13.0% | 1.2% | 89.2% | 44.5% | 0.4% | 95.0% | 35.8% | 64.6% | 27.3% | 86.1% |
| | 3-way | 56,000 | 2,000 | 3.7% | 3.6% | 70.7% | 76.8% | 1.6% | 81.5% | 69.4% | 9.9% | 47.7% | 83.7% | 4.2% | 70.5% | 79.4% | 38.2% | 54.0% | 95.7% |
| | 4-way | 83,720 | 690 | 0.83% | 3.7% | 75.9% | 94.9% | 1.4% | 86.2% | 92.5% | 20.3% | 23.0% | 97.0% | 8.2% | 56.7% | 95.8% | 22.1% | 70.7% | 98.9% |
| | 6-way | 800,800 | 100 | 0.012% | 0.8% | 94.0% | 99.9% | 0.2% | 97.0% | 99.8% | 15.8% | 1.0% | 99.9% | 4.1% | 23.0% | 99.8% | 7.6% | 94.0% | 100.0% |
| distal rotating protocol; proxy sources/outgroups in the 1st and 3rd circles around the target, at 100 gen. before "present" | | | | | | | | | | | | | | | | | | | |
| | 2-way | 10,800 | 1,350 | 14.3% | 0.0% | 97.7% | 6.1% | 0.0% | 99.1% | 0.0% | 0.2% | 97.6% | 36.5% | 0.1% | 99.1% | 29.4% | 52.5% | 31.1% | 84.2% |
| | 3-way | 56,000 | 2,000 | 3.7% | 0.4% | 89.0% | 48.1% | 0.1% | 94.7% | 39.9% | 4.0% | 78.7% | 83.4% | 1.4% | 90.5% | 79.9% | 17.8% | 60.3% | 92.4% |
| | 4-way | 151,060 | 1,245 | 0.83% | 0.5% | 84.0% | 77.3% | 0.2% | 91.2% | 70.9% | 16.1% | 44.5% | 97.2% | 7.1% | 69.8% | 96.6% | 2.5% | 78.6% | 93.4% |
| | 6-way | 800,800 | 100 | 0.012% | 0.1% | 86.0% | 98.8% | 0.1% | 86.0% | 98.4% | 70.7% | 0.0% | 100.0% | 47.2% | 4.0% | 100.0% | 0.2% | 86.0% | 98.9% |
| Pearson's *r* for pre-study odds vs. FDR (across all the protocols and model complexities above): | | | | | | | -0.91 | | | -0.85 | | | -0.99 | | | -0.99 | | | -0.95 |

Confusion matrix for three high-throughput *qpAdm* protocols based on systematic landscape sampling. Criteria for true and false models are shown in the table header; details of the *qpAdm* protocols are also listed in the sub-headers; five alternative model feasibility criteria are listed in the column captions. The following metrics are shown: false positive rate (FPR or $\alpha$) = 1 – true negative rate (TNR), false negative rate (FNR or $\beta$) = 1 – true positive rate (TPR) = 1 – power, and false discovery rate (FDR). Results for all simulation replicates were considered together.

**Table S14.**

| | | real data: Zeng et al. 2023 (distal and proximal rotating) | | | | | | | | | | SSL: "10-5 to 10-2" (distal and proximal rotating) | | | | | | | | | |
|---|---|---|---|---|---|---|---|---|---|---|---|---|---|---|---|---|---|---|---|---|---|
| variable 2: | | max. \|estimated admixture fractions - equal fractions\| | | | | | *p*-value | | | | | max. \|estimated admixture fractions - equal fractions\| | | | | | *p*-value | | | | |
| total number of models: | | 393,379 | 202,156 | 51,162 | 46,262 | 25,475 | 393,379 | 202,156 | 51,162 | 46,262 | 25,475 | 4,454,794 | 2,030,393 | 479,095 | 401,509 | 165,261 | 4,454,794 | 2,030,393 | 479,095 | 401,509 | 165,261 |
| variable 1: | model complexity | p >= 1e-55 | p >= 1e-5 | max. \|*EAF–EF*\| <= 0.5 | p >= 1e-55; max. \|*EAF–EF*\| <= 0.5 | p >= 1e-5; max. \|*EAF –EF*\| <= 0.5 | p >= 1e-55 | p >= 1e-5 | max. \|*EAF–EF*\| <= 0.5 | p >= 1e-55; max. \|*EAF–EF*\| <= 0.5 | p >= 1e-5; max. \|*EAF –EF*\| <= 0.5 | p >= 1e-55 | p >= 1e-5 | max. \|*EAF–EF*\| <= 0.5 | p >= 1e-55; max. \|*EAF–EF*\| <= 0.5 | p >= 1e-5; max. \|*EAF –EF*\| <= 0.5 | p >= 1e-55 | p >= 1e-5 | max. \|*EAF–EF*\| <= 0.5 | p >= 1e-55; max. \|*EAF–EF*\| <= 0.5 | p >= 1e-5; max. \|*EAF –EF*\| <= 0.5 |
| min. source-target-source angle | 2-way | -0.44 | -0.32 | -0.03 | -0.06 | -0.05 | 0.11 | 0.06 | -0.07 | -0.01 | -0.10 | **-0.76** | **-0.78** | -0.20 | -0.23 | -0.21 | -0.08 | -0.11 | 0.22 | 0.00 | -0.05 |
| | 3-way | -0.38 | -0.44 | -0.05 | -0.07 | -0.07 | 0.02 | -0.05 | 0.06 | 0.03 | -0.04 | **-0.51** | **-0.55** | -0.30 | -0.28 | -0.20 | -0.16 | -0.10 | -0.13 | -0.12 | -0.06 |
| | 4-way | -0.39 | -0.45 | -0.14 | -0.15 | -0.15 | -0.06 | -0.10 | 0.02 | 0.01 | 0.00 | -0.33 | -0.35 | -0.18 | -0.17 | -0.12 | -0.19 | -0.10 | -0.23 | -0.21 | -0.08 |
| max. source-target distance | 2-way | 0.04 | -0.11 | -0.05 | -0.05 | -0.01 | -0.21 | -0.14 | -0.15 | -0.14 | -0.16 | 0.24 | 0.18 | 0.32 | 0.47 | **0.58** | -0.18 | -0.07 | -0.33 | -0.12 | -0.03 |
| | 3-way | 0.05 | -0.11 | 0.07 | 0.08 | 0.01 | -0.22 | -0.05 | -0.11 | -0.09 | -0.10 | 0.18 | 0.14 | 0.24 | 0.22 | 0.16 | -0.15 | -0.06 | -0.13 | -0.08 | -0.05 |
| | 4-way | 0.10 | 0.02 | 0.04 | 0.04 | 0.02 | -0.27 | -0.11 | -0.11 | -0.10 | -0.07 | 0.16 | 0.13 | 0.11 | 0.10 | 0.06 | -0.13 | -0.06 | -0.01 | -0.01 | -0.01 |
| average source-target distance | 2-way | 0.10 | -0.03 | -0.12 | -0.14 | -0.11 | -0.29 | -0.18 | -0.25 | -0.24 | -0.17 | 0.46 | 0.44 | 0.10 | 0.26 | 0.39 | -0.28 | -0.14 | **-0.61** | -0.33 | -0.15 |
| | 3-way | 0.14 | 0.03 | 0.05 | 0.05 | -0.04 | -0.26 | -0.08 | -0.21 | -0.18 | -0.11 | 0.45 | 0.47 | 0.15 | 0.16 | 0.13 | -0.25 | -0.12 | -0.46 | -0.35 | -0.16 |
| | 4-way | 0.20 | 0.14 | 0.03 | 0.03 | 0.00 | -0.31 | -0.14 | -0.18 | -0.17 | -0.09 | 0.42 | 0.43 | 0.11 | 0.12 | 0.10 | -0.22 | -0.11 | -0.31 | -0.27 | -0.12 |
| distance to the ideal symmetric model (in the space of max. distance vs. min. angle) | 2-way | 0.43 | 0.30 | 0.02 | 0.06 | 0.05 | -0.12 | -0.07 | 0.04 | -0.01 | 0.08 | **0.71** | **0.73** | 0.32 | 0.43 | 0.48 | -0.01 | 0.06 | -0.29 | -0.05 | 0.00 |
| | 3-way | 0.34 | 0.38 | 0.08 | 0.10 | 0.09 | -0.10 | 0.02 | -0.08 | -0.04 | 0.03 | 0.46 | 0.49 | 0.33 | 0.31 | 0.23 | 0.03 | 0.04 | 0.06 | 0.08 | 0.04 |
| | 4-way | 0.35 | 0.38 | 0.15 | 0.15 | 0.15 | -0.10 | 0.01 | -0.05 | -0.03 | -0.01 | 0.33 | 0.33 | 0.18 | 0.17 | 0.11 | 0.04 | 0.03 | 0.18 | 0.17 | 0.06 |
| distance to the ideal symmetric model (in the space of average distance vs. min. angle) | 2-way | 0.45 | 0.31 | 0.01 | 0.05 | 0.04 | -0.13 | -0.07 | 0.04 | -0.01 | 0.09 | **0.79** | **0.81** | 0.22 | 0.30 | 0.32 | 0.01 | 0.08 | -0.35 | -0.07 | 0.03 |
| | 3-way | 0.38 | 0.42 | 0.06 | 0.08 | 0.08 | -0.09 | 0.02 | -0.09 | -0.05 | 0.03 | **0.58** | **0.64** | 0.31 | 0.29 | 0.22 | 0.06 | 0.06 | 0.04 | 0.06 | 0.04 |
| | 4-way | 0.40 | 0.44 | 0.15 | 0.16 | 0.16 | -0.08 | 0.02 | -0.04 | -0.03 | -0.01 | 0.44 | 0.48 | 0.19 | 0.18 | 0.13 | 0.09 | 0.05 | 0.18 | 0.16 | 0.06 |

*p*-value > 0.05/300 (Bonferroni correction)

**|Spearman correlation coefficient| >= 0.5**

Spearman's correlation coefficient (ρ) between several model optimality metrics (listed on the left) and max|*EAF–EF*| or *p*-values in various sections of the respective two-dimensional space (in columns). Here we compare *qpAdm* models analyzed by the high-throughput protocol from Zeng *et al.* (2023) and results of the randomized *qpAdm* protocol on the "$10^{-5}$ to $10^{-2}$" landscapes. In both cases, two- to four-way models from distal and proximal rotating experiments were considered, and the results are stratified by model complexity. Non-significant ρ values (with Bonferroni correction) are underlined, and absolute ρ values ≥0.5 are shown in bold. The distances and angles on the real data (great circle distances in km and angles between two bearings) are based on centroids of groups calculated using an *R* implementation of the *Mean Center* tool from *ArcGIS Pro*. We considered ST distances ≤50 km to be negligible, and for such models min. STS angles and hence distances to the ideal symmetric models were not defined, following the approach we applied to zero-length ST distances on the simulated data.

**Table S15.**

| metric | Pearson's *r* | Spearman's *rho* | |
|---|---|---|---|
| *p* -value | 0.99 | 0.72 | * |
| log10( *p* -value ) | 0.73 | 0.72 | |
| log10( \|EAF1\| ) | 0.91 | 0.94 | |
| log10( SE ) | 0.79 | 0.86 | |

* prioritizes high *p* -values

Correlation of *p*-values and EAF in results generated with *ADMIXTOOLS 2* v. 2.0.6 and classic *ADMIXTOOLS* v. 7.0 on the same data (low-quality pseudohaploid data for 3,000-Mbp-sized genomes, 10 AGS topologies, one simulation and one data subsampling replicate per topology). The results are based on 8,580 unique two-way *qpAdm* models. EAF1, estimated admixture fraction for proxy source no. 1 in a two-way model.
